# Method for inferring migration routes from phylogenetic trees

**DOI:** 10.1101/2020.02.06.937508

**Authors:** Juanjo Bermúdez

## Abstract

Inferring migration routes from a phylogenetic tree is something that has been done since we have phylogenetic trees. The usual method is just human inference, also known as intuition. Here I present a method for automating this task. Applied to phylogenetic trees with thousands of branches it makes possible to see for the first time inferred routes that were previously too complex to estimate.

## I. Introduction

One can make a rough estimation of a migration route just taking a look at a phylogenetic tree of reduced size. Knowing the location of a set of related leaves it’s usually common sense to find the most likely location of the immediate ancestor:

- Shorter routes are more likely than larger ones.
- Locations at an equidistant distance from all leaves are more likely than locations with very asymmetric distances.
- Locations not implying crossing natural obstacles are also more likely.
- Locations with more density of near leaves are preferred over other ones.
- If we know the location of a common ancestor, locations in the path between the leaves and the location of the common ancestor are more likely than locations far apart.
- Simpler routes are preferred over complex ones implying many changes of direction.

Sometimes, time-specific knowledge is also employed, like knowing when the sea level opened a path between islands or continents.

For example, a tree like the one in figure 1 could be read as: “There was a population somewhere. One of its descendants actually lives in Kenya. That population had a split and some of its members remained in Kenya (NA19454) and other ones moved to the south of Africa. The ones who moved split into two groups but are now mixed between local populations in southern Africa.”

**Fig. 1.**
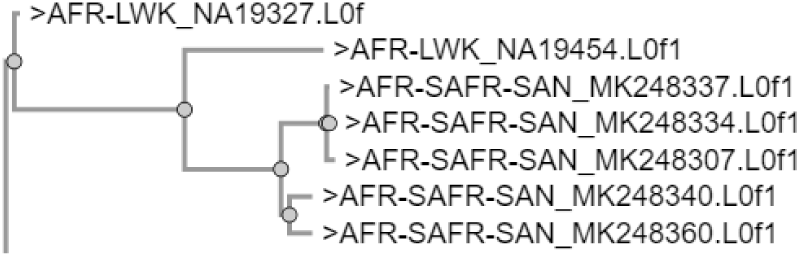
Phylogenetic tree of modern humans based on mitochondrial DNA. AFR-LWK are individuals actually living in Kenya and SAFR-SAN are members of the San tribes. All individuals are classified in the L0f haplogroup, with some of them more specifically included in L0f1.

And translated into a map that would look something like figure 2.

**Fig. 2.**
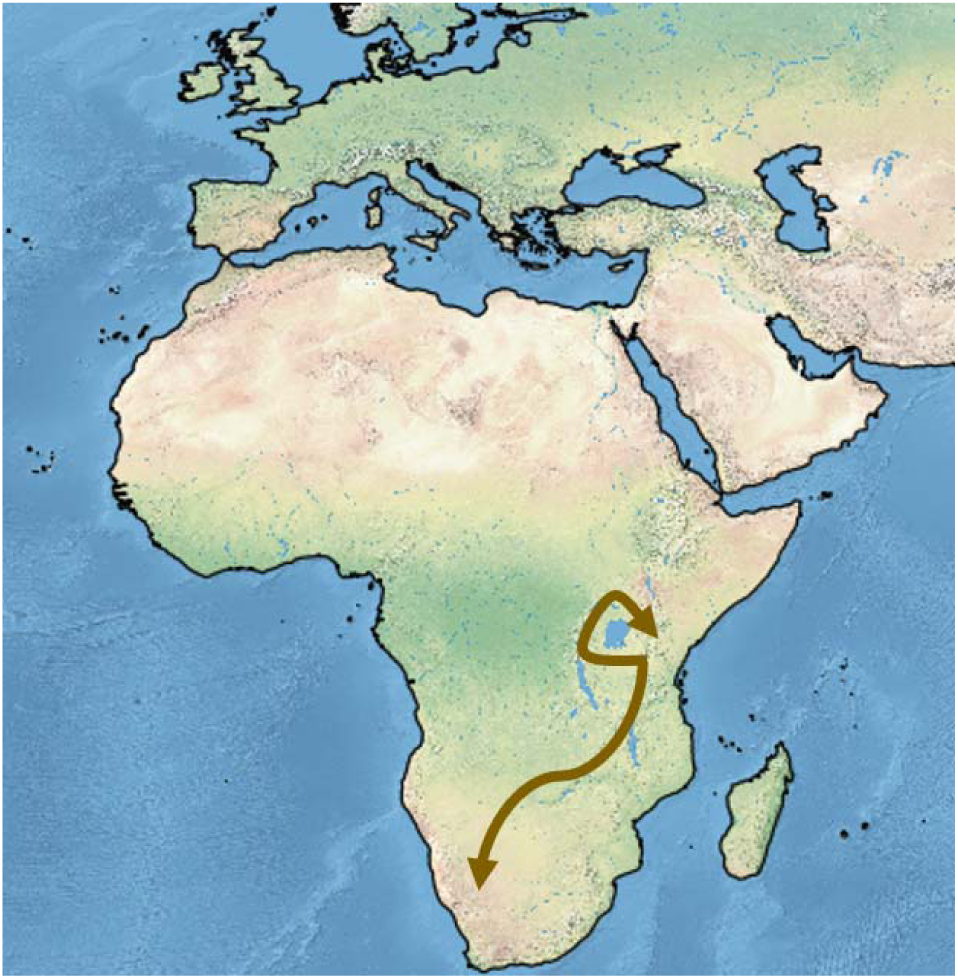
Inferred routes for populations related to figure 1.

This methodology can be automated. Some of the inferences will be less accurate than the ones made by humans but, in contrast, a bigger number of inferences can be made and we can dig deeper into large phylogenetic trees by inferring thousands of routes.

## II. Method

### 1. Network of available routes

First place, a network of available routes was created. This is an arbitrary step and many alternatives are possible.

**Fig. 3.**
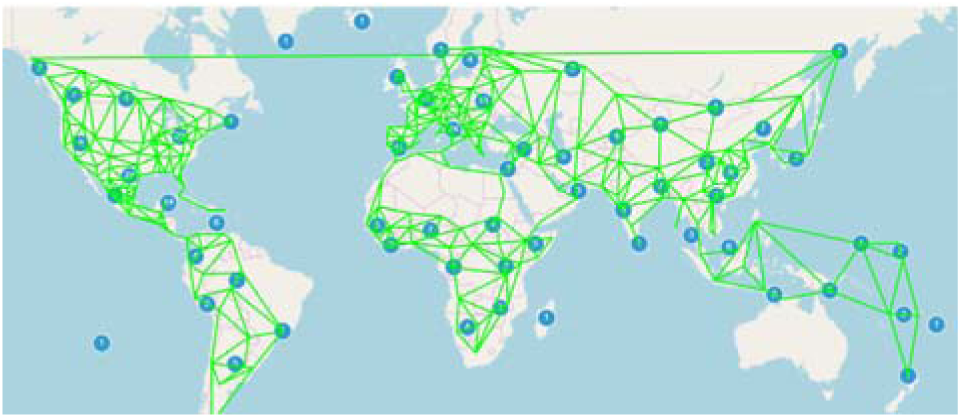
Network of available routes.

**Fig. 4.**
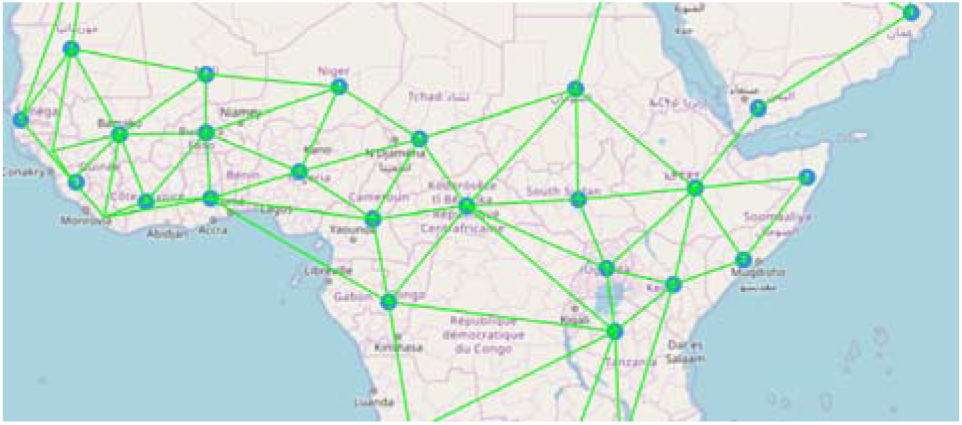
Network of available routes zoomed into an African region.

This network indicates the routes that can be taken from each marker on the map. Only marker-to-marker routes are allowed. We will generalize every location to its nearer marker.

These markers and the allowed routes were encoded in a Javascript script and an OpenLayers map [3] was used for visualization.

### 2. Inferring locations for internal nodes of the tree

Different algorithms for calculating the position of the internal nodes are possible. I made tests with the simplest one and that seems to be enough for our purposes. An implementation is available in the additional material.

First place, a marker id has to be passed as a parameter to the algorithm so that it becomes the location of the root node of the tree. This is an arbitrary parameter that we will use to test hypotheses.

Once we have assigned a location to the root node, we will traverse the tree in preorder. For every node, we will calculate the route from all its descendant leaf nodes to the location of the parent node. We will use the Dijkstra algorithm for that purpose. We will make then the gross assumption that the sought location will likely be around the nearest intersection of all these routes to the location of the leaves.

**Fig. 5.**
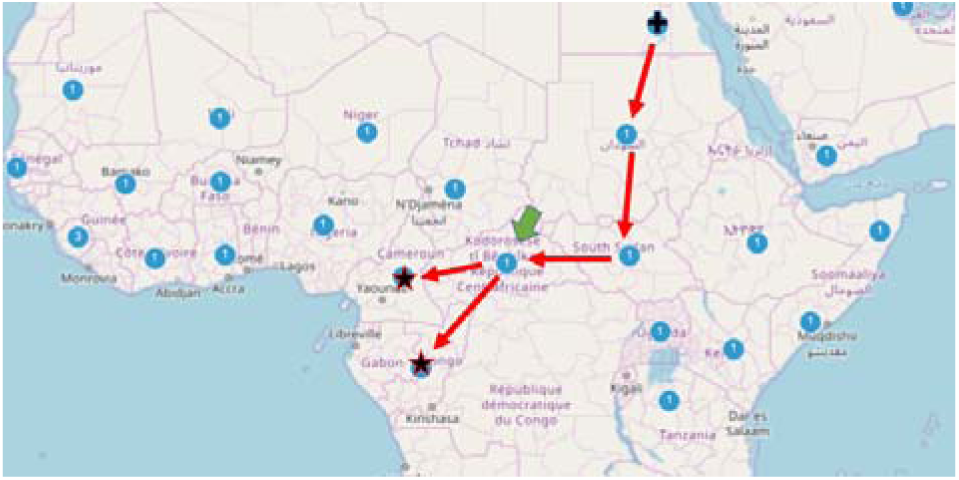
Route from the cross to the stars. The marker pointed by the green arrow is the selected one for the intermediate node.

**Fig. 6.**
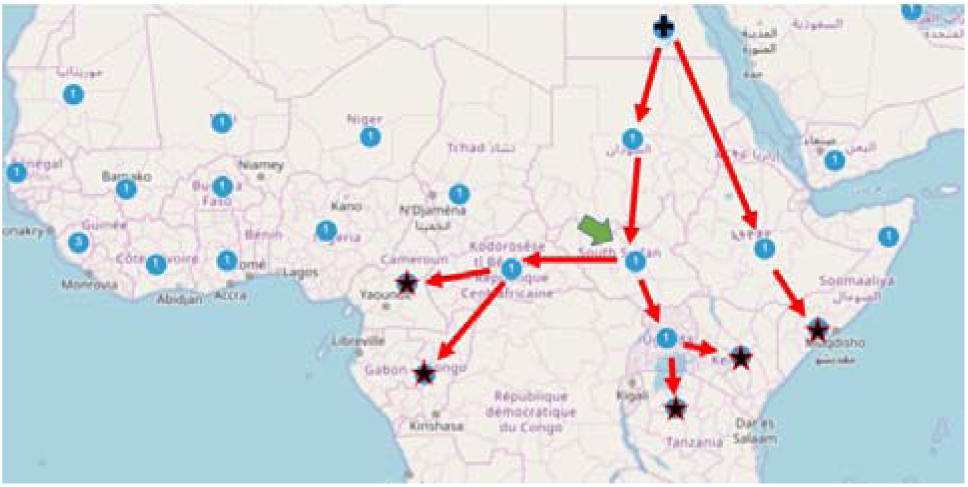
Route from the cross to the stars. The marker pointed by the green arrow is the selected one for the intermediate node.

An additional hack consists in going one or more steps back when the number of branches in an alternative route is higher than a given threshold. Another hack consists in using the mutations distance between the root and the leaves for weighting every route. This way, the nearest leaves (these that evolved earlier from the node) will have more influence on the location of the new node.

**Fig. 7.**
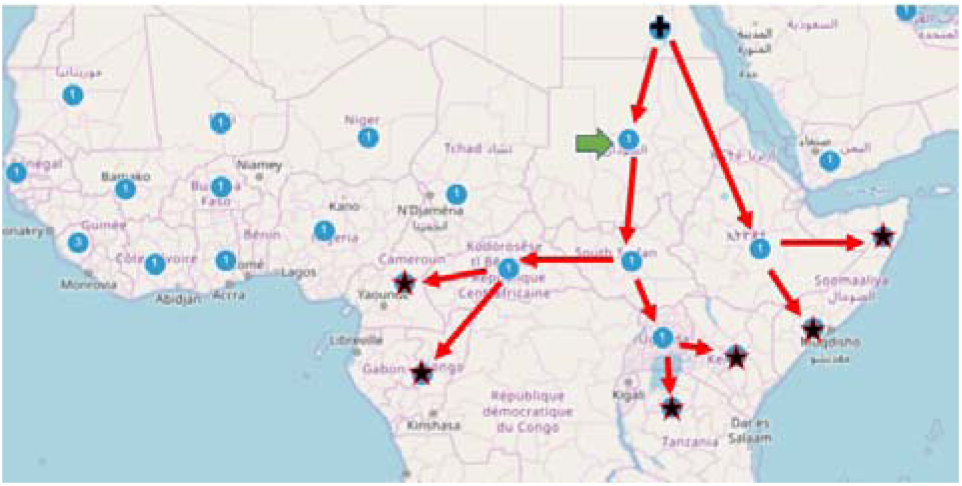
Route from the cross to the stars. The marker pointed by the green arrow is the selected one for the intermediate node.

### 3. Tracing routes

Once we have calculated the position for every node in the tree we can draw route-trees in different ways. I will call sometimes “route-trees” to these routes as they are a tree of routes.

#### 3.1. Routes from the root down to a depth limit

We can set as parameter a depth limit. The tool will only display routes for nodes whose distance in mutations to the root node is under that limit of base pairs (bp).

**Fig. 8.**
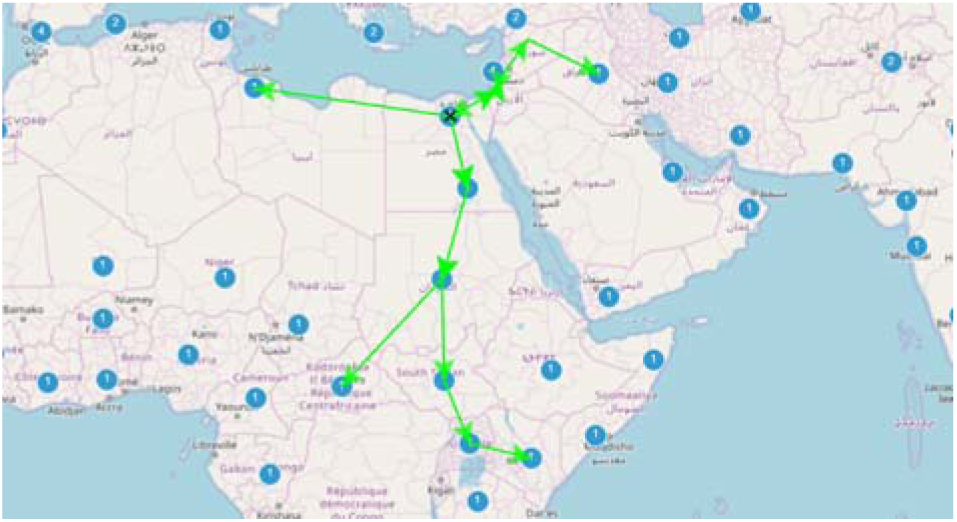
Routes of expansion of modern humans from a proposed location for the Mitochondrial Eve with depth limit 7.

**Fig. 9.**
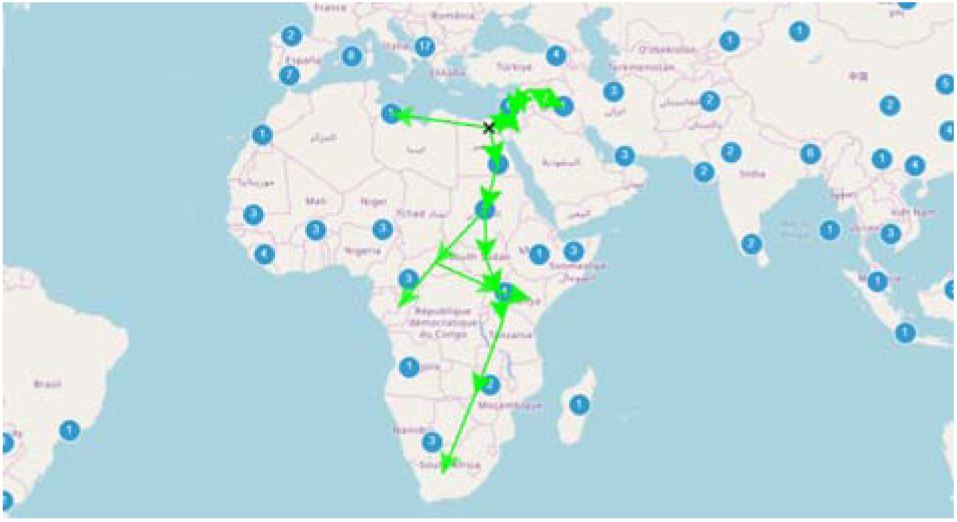
Routes of expansion of modern humans from a proposed location for the Mitochondrial Eve with depth limit 70.

#### 3.2. Routes from an arbitrary node in the phylogenetic tree down to a given depth

In the same way that we can set a depth limit, we can select a node in the phylogenetic tree to become the root of our route-tree. The depth limit will be calculated in relation to the new root.

**Fig. 10.**
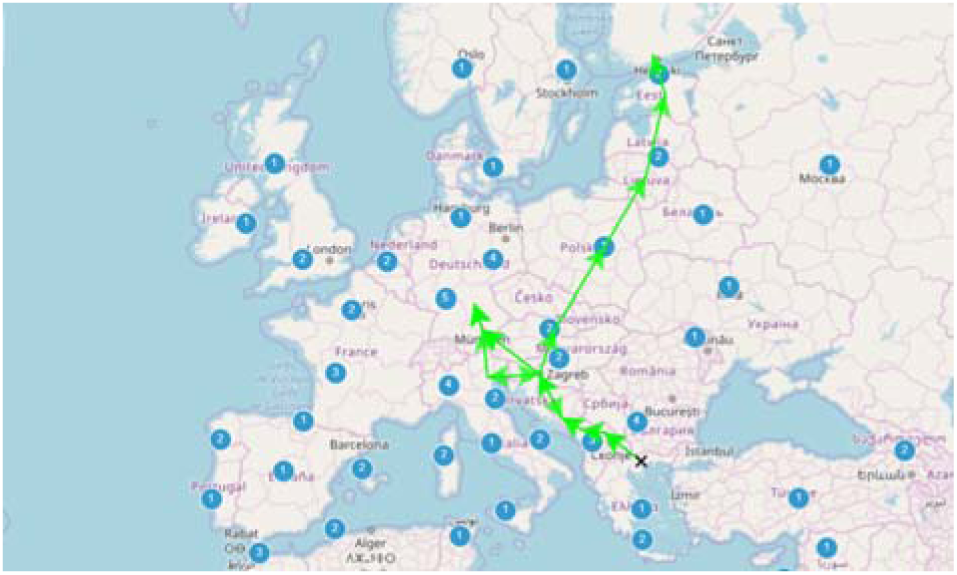
Route of migration of modern humans with depth limit 15 departing from a selected node in the phylogenetic tree.

**Fig. 11.**
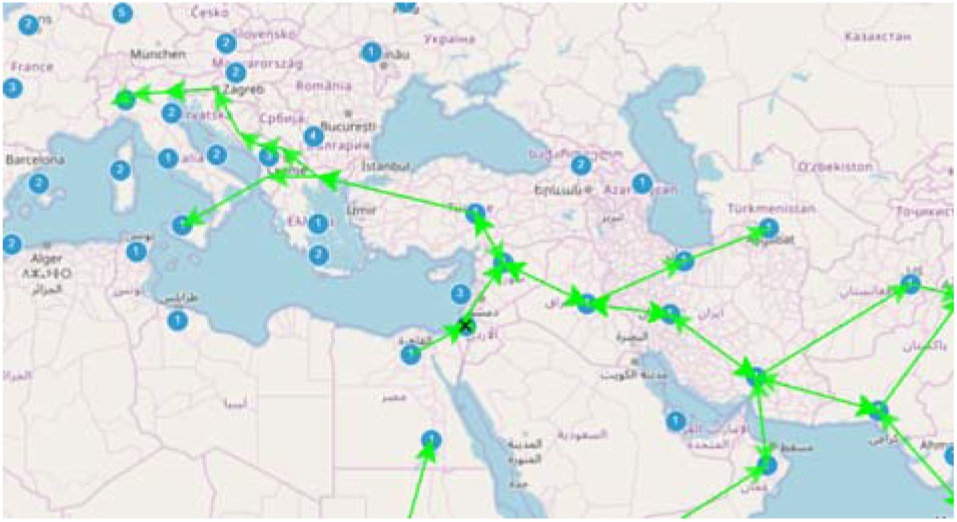
Routes of early migration of modern humans with depth limit 150 and initial depth 140.

#### 3.3. All routes to leaves from a node in the phylogenetic tree

**Fig. 12.**
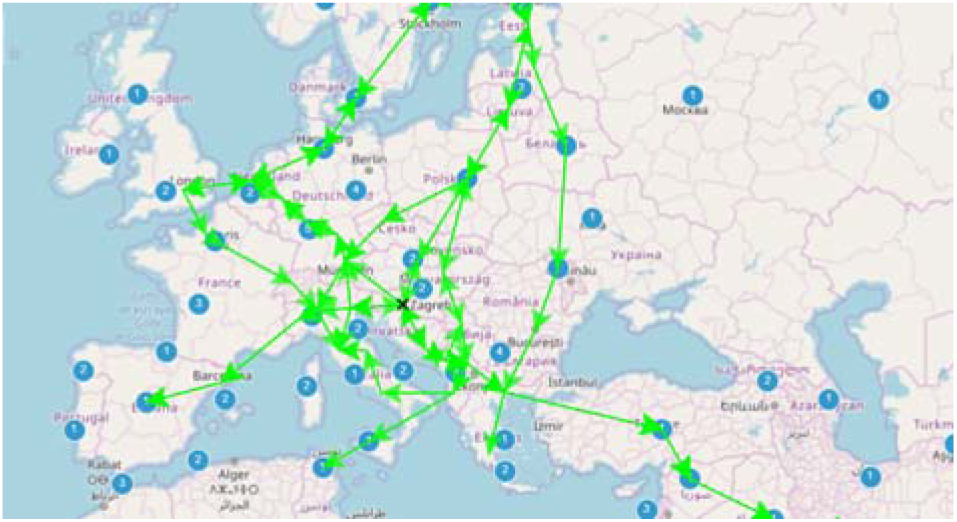
Routes to all leaves for a node deep in the mitochondrial tree of modern humans, corresponding to an expansion of a population from the Balcans to all over Europe and parts of Asia.

#### 3.4. All routes to a given destination in the map

**Fig. 13.**
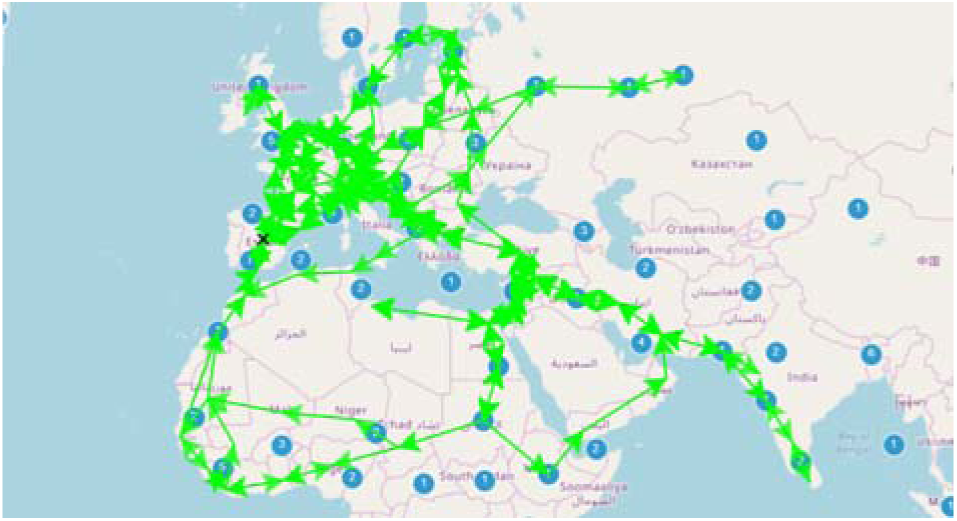
All routes having the Iberian population as destination in the tree of expansion of modern humans.

**Fig. 14.**
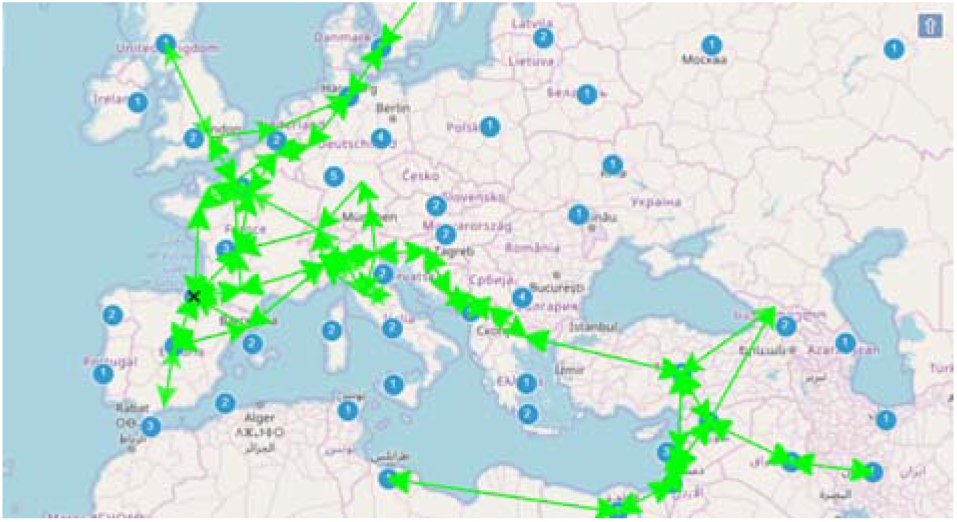
All routes having the Basque population as destination in the tree of expansion of modern humans.

**Fig. 15.**
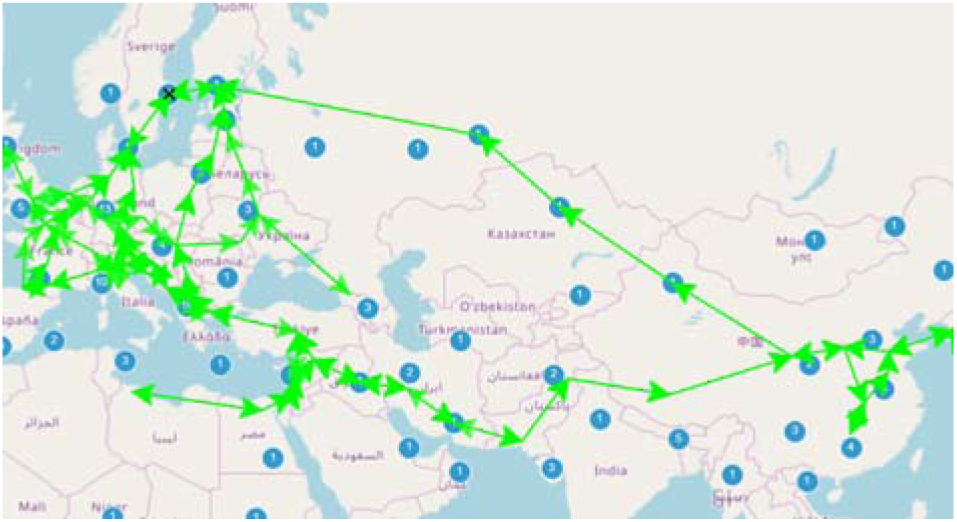
All routes having the Swedish population as destination in the tree of expansion of modern humans.

**Fig. 16.**
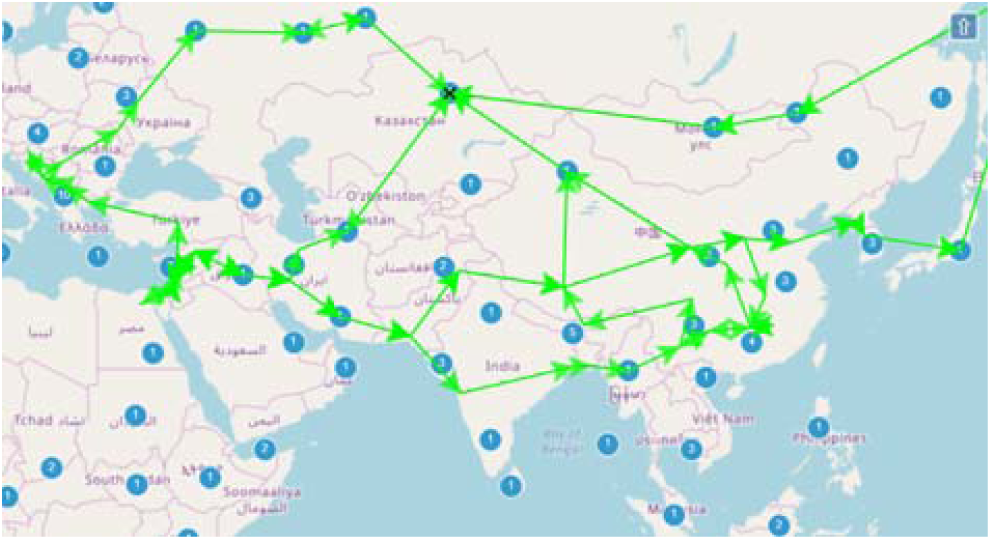
All routes having the population of Kazakhstan as destination in the tree of expansion of modern humans.

**Fig. 17.**
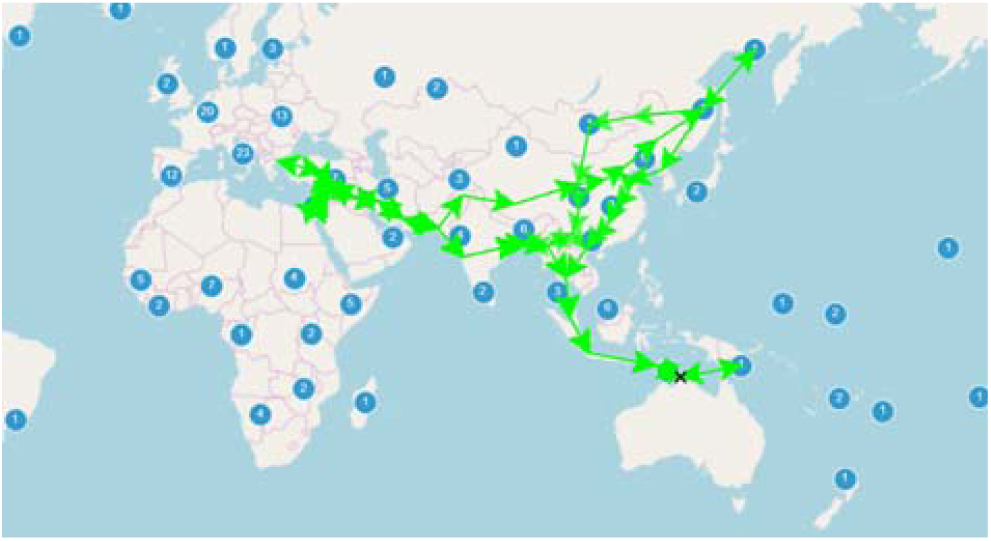
All routes having the Australasian population as a leaf in the tree of expansion of modern humans.

### 4. Uses

These functionalities can be used for different purposes. I will list next some of them.

#### 4.1. Making an animation of population expansions

**Fig. 18.**
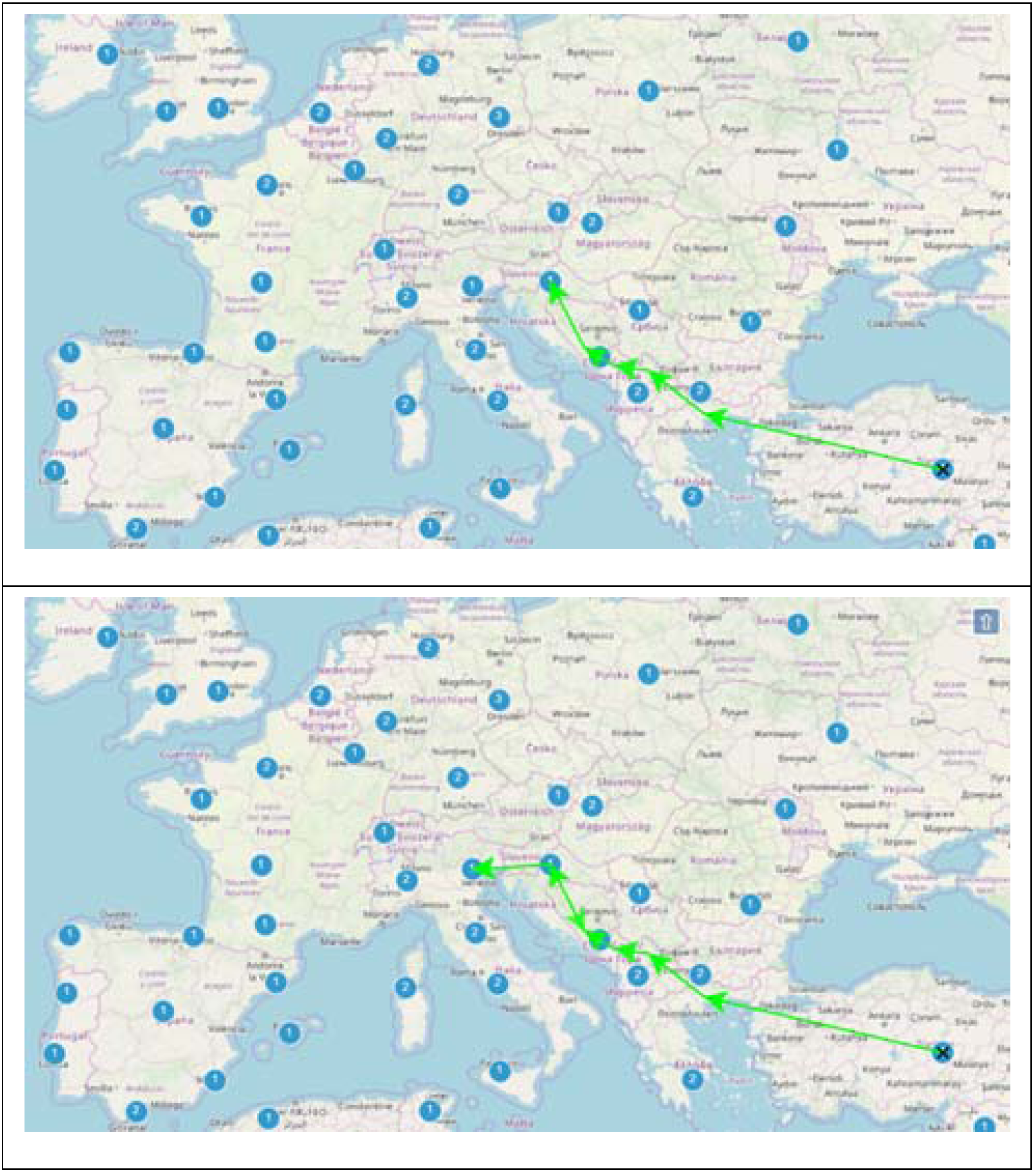
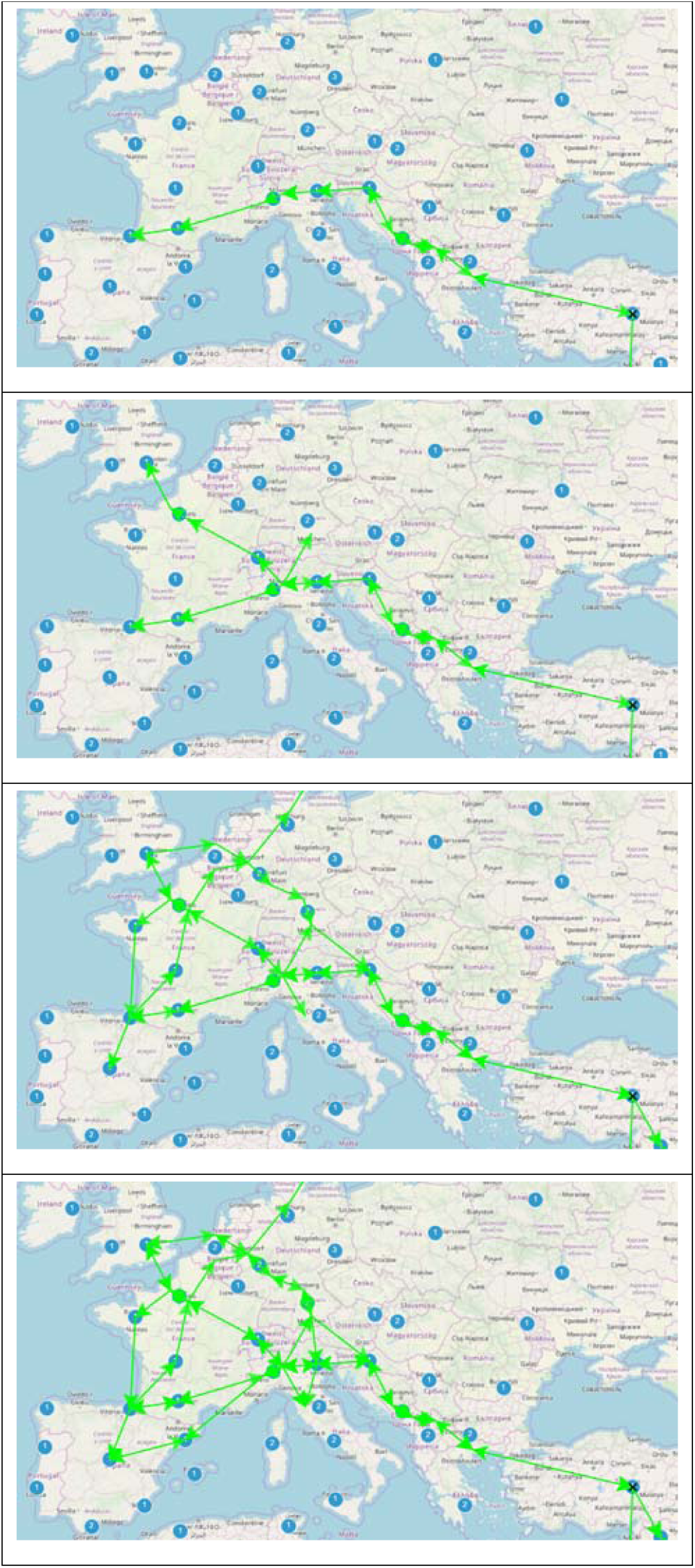
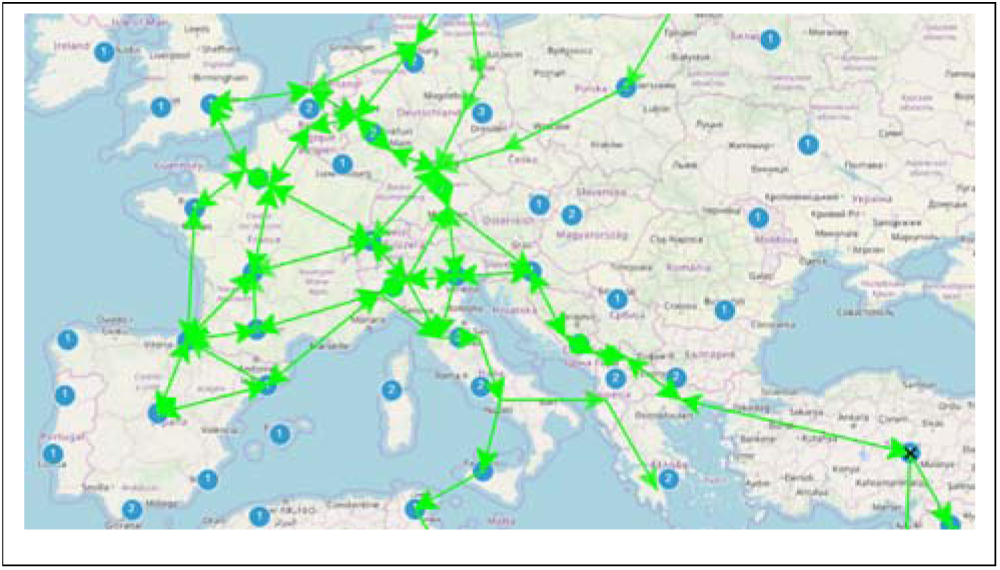
The sequence of expansions of some first modern humans arriving in Europe (depths 5, 15, 25, 28, 31 and unlimited).

From this route-tree of expansion of modern humans into Europe, some considerations are important. These considerations show some limitations of this method. But these are limitations that are not important if your main goal is just getting an overall picture of the migrations.

- The initial migration to the Basque country was more likely to all western Europe, as that would likely be the areas they populated by then. If it points only to that region that is because this is where they are actually located.
- The routes to Britain could mean routes to the ancestors of ancient Britons or routes to the ancestors of ancient Angles and Saxons. The ancestors of Angles and Saxons were not in Britain by then but (maybe) in the north of Germany. Therefore the arrow pointing to England maybe should be really pointing to the north of Germany.
- Data from only a few European countries were included in the phylogenetic tree. There is no data for French people, for example. The routes would likely be a little different with more data.

If these problems were important in your case, some restrictions could be easily incorporated into the algorithm, like placing people from England in north Germany until a given depth of the tree.

Another important consideration is that expansions over an area are usually not in a single branch of the phylogenetic tree. Many ancestor lines, for example, arrived in Europe. Some likely pertained to the same groups and followed the same routes. Other ones pertained to different waves or split later into different routes.

From these animations, we can conclude, for example, that many Europeans are descendants from the first hunter-gatherers arriving into Europe, who were closely related to the actual Basque people.

#### 4.2. Finding the likely location of the ancestors of a population

We can select a node in the phylogenetic tree from which all nodes of a selected population descend and see this way the likely location of the ancestorso of that population.

**Fig. 19.**
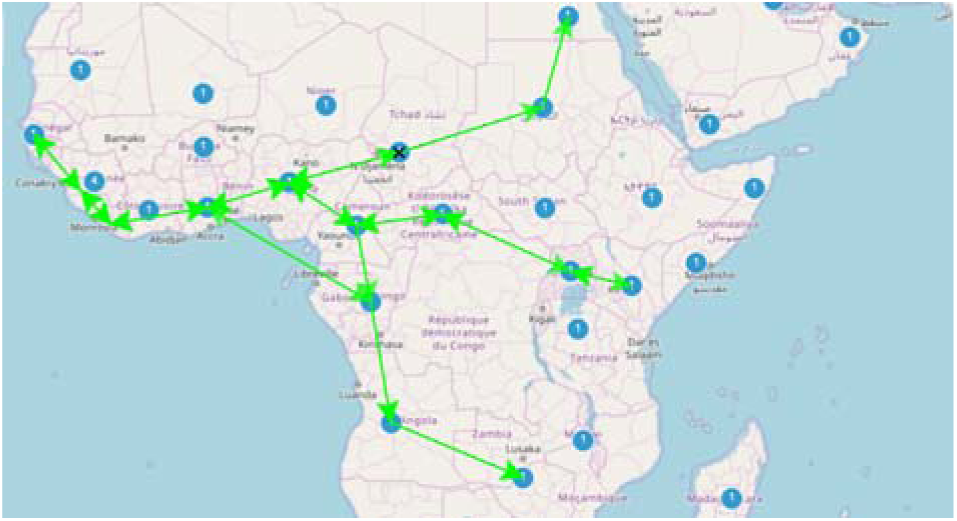
Likely origin of the haplogroup L2b and its expansion routes.

**Fig. 20.**
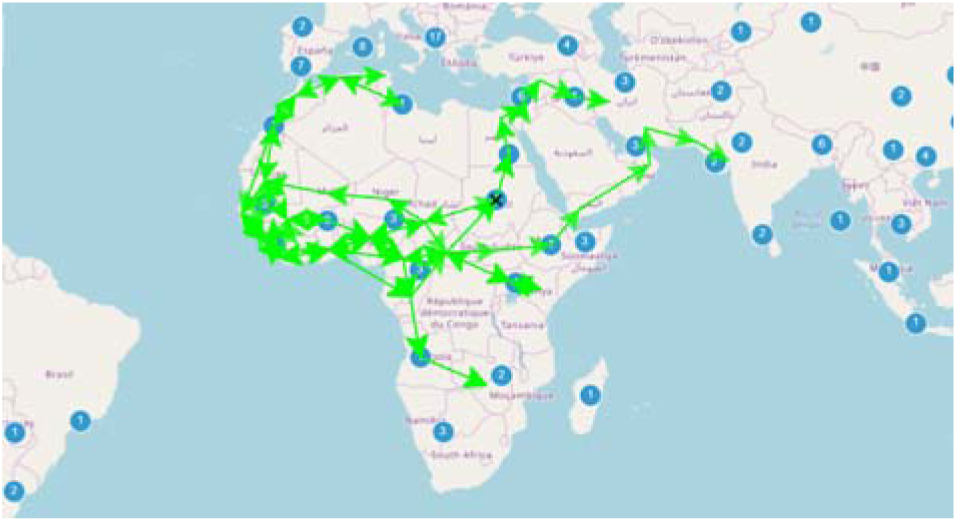
Likely origin of the haplogroup L2a and its expansion routes.

#### 4.3. Checking how some individuals became part of a population

**Fig. 21.**
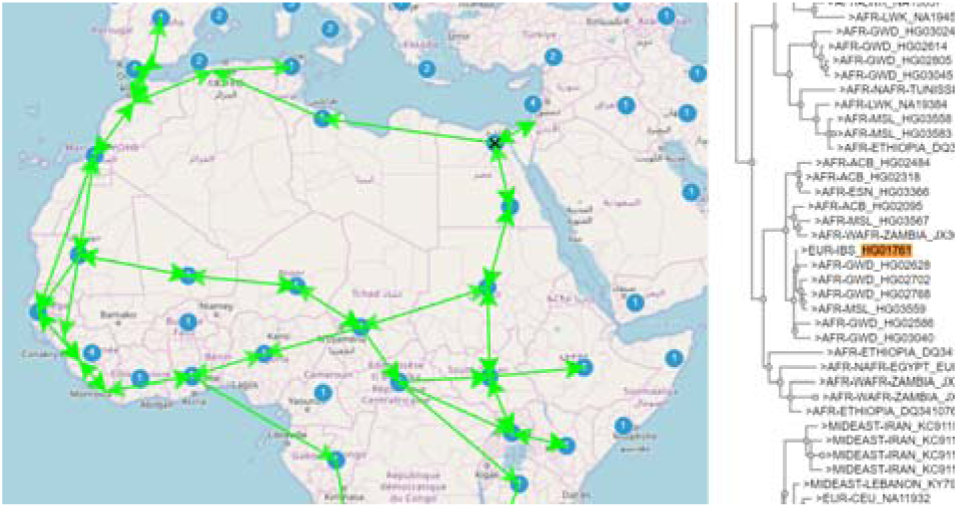
Routes associated with the branch of the phylogenetic tree on the right. An individual pertaining to an Iberian population is marked.

We can find in the phylogenetic tree an Iberian individual pertaining to the haplogroup L3. We would like to know how that person likely arrived there. There are two options: he arrived through the Pyrenees on an early OOA wave or he arrived through Gibraltar later. Drawing the inferred route-tree for an ancestor node in the phylogenetic tree we see that the most reasonable answer is through Gibraltar.

## III. CONCLUSIONS

Despite its limitations, this method lets us see by brute force what we could not see using more robust methods for inferring routes. Imprecisions interpreting some data are compensated by the abundance of data.

More research should be done on more complex algorithms following this idea.

## Supporting information

Some route-trees

## IV. Acknowledgments

Some data for this study were obtained from [1][2].

Some free software tools used were: [3][4]

The author of this research adheres to the Agile Science Manifesto v 1.0 [5] and this document was written in concordance with its arguments.

