## Supplementary material for "Method for inferring migration routes from phylogenetic trees": Some route-trees: bootstrap.min.js

```
/*!
  * Bootstrap v4.3.1 (https://getbootstrap.com/)
  * Copyright 2011-2019 The Bootstrap Authors (https://github.com/twbs/bootstrap/graphs/contributors)
  * Licensed under MIT (https://github.com/twbs/bootstrap/blob/master/LICENSE)
  */
!function(t,e){"object"==typeof exports&&"undefined"!=typeof module?e(exports,require("jquery"),require("popper.js")):"function"==typeof define&&define.amd?define(["exports","jquery","popper.js"],e):e((t=t||self).bootstrap={},t.jQuery,t.Popper)}(this,function(t,g,u){"use strict";function i(t,e){for(var n=0;n<e.length;n++){var i=e[n];i.enumerable=i.enumerable||!1,i.configurable=!0,"value"in i&&(i.writable=!0),Object.defineProperty(t,i.key,i)}}function s(t,e,n){return e&&i(t.prototype,e),n&&i(t,n),t}function l(o){for(var t=1;t<arguments.length;t++){var r=null!=arguments[t]?arguments[t]:{},e=Object.keys(r);"function"==typeof Object.getOwnPropertySymbols&&(e=e.concat(Object.getOwnPropertySymbols(r).filter(function(t){return Object.getOwnPropertyDescriptor(r,t).enumerable}))),e.forEach(function(t){var e,n,i;e=o,i=r[n=t],n in e?Object.defineProperty(e,n,{value:i,enumerable:!0,configurable:!0,writable:!0}):e[n]=i})}return o}g=g&&g.hasOwnProperty("default")?g.default:g,u=u&&u.hasOwnProperty("default")?u.default:u;var e="transitionend";function n(t){var e=this,n=!1;return g(this).one(_.TRANSITION_END,function(){n=!0}),setTimeout(function(){n||_.triggerTransitionEnd(e)},t),this}var _={TRANSITION_END:"bsTransitionEnd",getUID:function(t){for(;t+=~~(1e6*Math.random()),document.getElementById(t););return t},getSelectorFromElement:function(t){var e=t.getAttribute("data-target");if(!e||"#"===e){var n=t.getAttribute("href");e=n&&"#"!==n?n.trim():""}try{return document.querySelector(e)?e:null}catch(t){return null}},getTransitionDurationFromElement:function(t){if(!t)return 0;var e=g(t).css("transition-duration"),n=g(t).css("transition-delay"),i=parseFloat(e),o=parseFloat(n);return i||o?(e=e.split(",")[0],n=n.split(",")[0],1e3*(parseFloat(e)+parseFloat(n))):0},reflow:function(t){return t.offsetHeight},triggerTransitionEnd:function(t){g(t).trigger(e)},supportsTransitionEnd:function(){return Boolean(e)},isElement:function(t){return(t[0]||t).nodeType},typeCheckConfig:function(t,e,n){for(var i in n)if(Object.prototype.hasOwnProperty.call(n,i)){var o=n[i],r=e[i],s=r&&_.isElement(r)?"element":(a=r,{}.toString.call(a).match(/\s([a-z]+)/i)[1].toLowerCase());if(!new RegExp(o).test(s))throw new Error(t.toUpperCase()+': Option "'+i+'" provided type "'+s+'" but expected type "'+o+'".')}var a},findShadowRoot:function(t){if(!document.documentElement.attachShadow)return null;if("function"!=typeof t.getRootNode)return t instanceof ShadowRoot?t:t.parentNode?_.findShadowRoot(t.parentNode):null;var e=t.getRootNode();return e instanceof ShadowRoot?e:null}};g.fn.emulateTransitionEnd=n,g.event.special[_.TRANSITION_END]={bindType:e,delegateType:e,handle:function(t){if(g(t.target).is(this))return t.handleObj.handler.apply(this,arguments)}};var o="alert",r="bs.alert",a="."+r,c=g.fn[o],h={CLOSE:"close"+a,CLOSED:"closed"+a,CLICK_DATA_API:"click"+a+".data-api"},f="alert",d="fade",m="show",p=function(){function i(t){this._element=t}var t=i.prototype;return t.close=function(t){var e=this._element;t&&(e=this._getRootElement(t)),this._triggerCloseEvent(e).isDefaultPrevented()||this._removeElement(e)},t.dispose=function(){g.removeData(this._element,r),this._element=null},t._getRootElement=function(t){var e=_.getSelectorFromElement(t),n=!1;return e&&(n=document.querySelector(e)),n||(n=g(t).closest("."+f)[0]),n},t._triggerCloseEvent=function(t){var e=g.Event(h.CLOSE);return g(t).trigger(e),e},t._removeElement=function(e){var n=this;if(g(e).removeClass(m),g(e).hasClass(d)){var t=_.getTransitionDurationFromElement(e);g(e).one(_.TRANSITION_END,function(t){return n._destroyElement(e,t)}).emulateTransitionEnd(t)}else this._destroyElement(e)},t._destroyElement=function(t){g(t).detach().trigger(h.CLOSED).remove()},i._jQueryInterface=function(n){return this.each(function(){var t=g(this),e=t.data(r);e||(e=new i(this),t.data(r,e)),"close"===n&&e[n](this)})},i._handleDismiss=function(e){return function(t){t&&t.preventDefault(),e.close(this)}},s(i,null,[{key:"VERSION",get:function(){return"4.3.1"}}]),i}();g(document).on(h.CLICK_DATA_API,'[data-dismiss="alert"]',p._handleDismiss(new p)),g.fn[o]=p._jQueryInterface,g.fn[o].Constructor=p,g.fn[o].noConflict=function(){return g.fn[o]=c,p._jQueryInterface};var v="button",y="bs.button",E="."+y,C=".data-api",T=g.fn[v],S="active",b="btn",I="focus",D='[data-toggle^="button"]',w='[data-toggle="buttons"]',A='input:not([type="hidden"])',N=".active",O=".btn",k={CLICK_DATA_API:"click"+E+C,FOCUS_BLUR_DATA_API:"focus"+E+C+" blur"+E+C},P=function(){function n(t){this._element=t}var t=n.prototype;return t.toggle=function(){var t=!0,e=!0,n=g(this._element).closest(w)[0];if(n){var i=this._element.querySelector(A);if(i){if("radio"===i.type)if(i.checked&&this._element.classList.contains(S))t=!1;else{var o=n.querySelector(N);o&&g(o).removeClass(S)}if(t){if(i.hasAttribute("disabled")||n.hasAttribute("disabled")||i.classList.contains("disabled")||n.classList.contains("disabled"))return;i.checked=!this._element.classList.contains(S),g(i).trigger("change")}i.focus(),e=!1}}e&&this._element.setAttribute("aria-pressed",!this._element.classList.contains(S)),t&&g(this._element).toggleClass(S)},t.dispose=function(){g.removeData(this._element,y),this._element=null},n._jQueryInterface=function(e){return this.each(function(){var t=g(this).data(y);t||(t=new n(this),g(this).data(y,t)),"toggle"===e&&t[e]()})},s(n,null,[{key:"VERSION",get:function(){return"4.3.1"}}]),n}();g(document).on(k.CLICK_DATA_API,D,function(t){t.preventDefault();var e=t.target;g(e).hasClass(b)||(e=g(e).closest(O)),P._jQueryInterface.call(g(e),"toggle")}).on(k.FOCUS_BLUR_DATA_API,D,function(t){var e=g(t.target).closest(O)[0];g(e).toggleClass(I,/^focus(in)?$/.test(t.type))}),g.fn[v]=P._jQueryInterface,g.fn[v].Constructor=P,g.fn[v].noConflict=function(){return g.fn[v]=T,P._jQueryInterface};var L="carousel",j="bs.carousel",H="."+j,R=".data-api",x=g.fn[L],F={interval:5e3,keyboard:!0,slide:!1,pause:"hover",wrap:!0,touch:!0},U={interval:"(number|boolean)",keyboard:"boolean",slide:"(boolean|string)",pause:"(string|boolean)",wrap:"boolean",touch:"boolean"},W="next",q="prev",M="left",K="right",Q={SLIDE:"slide"+H,SLID:"slid"+H,KEYDOWN:"keydown"+H,MOUSEENTER:"mouseenter"+H,MOUSELEAVE:"mouseleave"+H,TOUCHSTART:"touchstart"+H,TOUCHMOVE:"touchmove"+H,TOUCHEND:"touchend"+H,POINTERDOWN:"pointerdown"+H,POINTERUP:"pointerup"+H,DRAG_START:"dragstart"+H,LOAD_DATA_API:"load"+H+R,CLICK_DATA_API:"click"+H+R},B="carousel",V="active",Y="slide",z="carousel-item-right",X="carousel-item-left",$="carousel-item-next",G="carousel-item-prev",J="pointer-event",Z=".active",tt=".active.carousel-item",et=".carousel-item",nt=".carousel-item img",it=".carousel-item-next, .carousel-item-prev",ot=".carousel-indicators",rt="[data-slide], [data-slide-to]",st='[data-ride="carousel"]',at={TOUCH:"touch",PEN:"pen"},lt=function(){function r(t,e){this._items=null,this._interval=null,this._activeElement=null,this._isPaused=!1,this._isSliding=!1,this.touchTimeout=null,this.touchStartX=0,this.touchDeltaX=0,this._config=this._getConfig(e),this._element=t,this._indicatorsElement=this._element.querySelector(ot),this._touchSupported="ontouchstart"in document.documentElement||0<navigator.maxTouchPoints,this._pointerEvent=Boolean(window.PointerEvent||window.MSPointerEvent),this._addEventListeners()}var t=r.prototype;return t.next=function(){this._isSliding||this._slide(W)},t.nextWhenVisible=function(){!document.hidden&&g(this._element).is(":visible")&&"hidden"!==g(this._element).css("visibility")&&this.next()},t.prev=function(){this._isSliding||this._slide(q)},t.pause=function(t){t||(this._isPaused=!0),this._element.querySelector(it)&&(_.triggerTransitionEnd(this._element),this.cycle(!0)),clearInterval(this._interval),this._interval=null},t.cycle=function(t){t||(this._isPaused=!1),this._interval&&(clearInterval(this._interval),this._interval=null),this._config.interval&&!this._isPaused&&(this._interval=setInterval((document.visibilityState?this.nextWhenVisible:this.next).bind(this),this._config.interval))},t.to=function(t){var e=this;this._activeElement=this._element.querySelector(tt);var n=this._getItemIndex(this._activeElement);if(!(t>this._items.length-1||t<0))if(this._isSliding)g(this._element).one(Q.SLID,function(){return e.to(t)});else{if(n===t)return this.pause(),void this.cycle();var i=n<t?W:q;this._slide(i,this._items[t])}},t.dispose=function(){g(this._element).off(H),g.removeData(this._element,j),this._items=null,this._config=null,this._element=null,this._interval=null,this._isPaused=null,this._isSliding=null,this._activeElement=null,this._indicatorsElement=null},t._getConfig=function(t){return t=l({},F,t),_.typeCheckConfig(L,t,U),t},t._handleSwipe=function(){var t=Math.abs(this.touchDeltaX);if(!(t<=40)){var e=t/this.touchDeltaX;0<e&&this.prev(),e<0&&this.next()}},t._addEventListeners=function(){var e=this;this._config.keyboard&&g(this._element).on(Q.KEYDOWN,function(t){return e._keydown(t)}),"hover"===this._config.pause&&g(this._element).on(Q.MOUSEENTER,function(t){return e.pause(t)}).on(Q.MOUSELEAVE,function(t){return e.cycle(t)}),this._config.touch&&this._addTouchEventListeners()},t._addTouchEventListeners=function(){var n=this;if(this._touchSupported){var e=function(t){n._pointerEvent&&at[t.originalEvent.pointerType.toUpperCase()]?n.touchStartX=t.originalEvent.clientX:n._pointerEvent||(n.touchStartX=t.originalEvent.touches[0].clientX)},i=function(t){n._pointerEvent&&at[t.originalEvent.pointerType.toUpperCase()]&&(n.touchDeltaX=t.originalEvent.clientX-n.touchStartX),n._handleSwipe(),"hover"===n._config.pause&&(n.pause(),n.touchTimeout&&clearTimeout(n.touchTimeout),n.touchTimeout=setTimeout(function(t){return n.cycle(t)},500+n._config.interval))};g(this._element.querySelectorAll(nt)).on(Q.DRAG_START,function(t){return t.preventDefault()}),this._pointerEvent?(g(this._element).on(Q.POINTERDOWN,function(t){return e(t)}),g(this._element).on(Q.POINTERUP,function(t){return i(t)}),this._element.classList.add(J)):(g(this._element).on(Q.TOUCHSTART,function(t){return e(t)}),g(this._element).on(Q.TOUCHMOVE,function(t){var e;(e=t).originalEvent.touches&&1<e.originalEvent.touches.length?n.touchDeltaX=0:n.touchDeltaX=e.originalEvent.touches[0].clientX-n.touchStartX}),g(this._element).on(Q.TOUCHEND,function(t){return i(t)}))}},t._keydown=function(t){if(!/input|textarea/i.test(t.target.tagName))switch(t.which){case 37:t.preventDefault(),this.prev();break;case 39:t.preventDefault(),this.next()}},t._getItemIndex=function(t){return this._items=t&&t.parentNode?[].slice.call(t.parentNode.querySelectorAll(et)):[],this._items.indexOf(t)},t._getItemByDirection=function(t,e){var n=t===W,i=t===q,o=this._getItemIndex(e),r=this._items.length-1;if((i&&0===o||n&&o===r)&&!this._config.wrap)return e;var s=(o+(t===q?-1:1))%this._items.length;return-1===s?this._items[this._items.length-1]:this._items[s]},t._triggerSlideEvent=function(t,e){var n=this._getItemIndex(t),i=this._getItemIndex(this._element.querySelector(tt)),o=g.Event(Q.SLIDE,{relatedTarget:t,direction:e,from:i,to:n});return g(this._element).trigger(o),o},t._setActiveIndicatorElement=function(t){if(this._indicatorsElement){var e=[].slice.call(this._indicatorsElement.querySelectorAll(Z));g(e).removeClass(V);var n=this._indicatorsElement.children[this._getItemIndex(t)];n&&g(n).addClass(V)}},t._slide=function(t,e){var n,i,o,r=this,s=this._element.querySelector(tt),a=this._getItemIndex(s),l=e||s&&this._getItemByDirection(t,s),c=this._getItemIndex(l),h=Boolean(this._interval);if(o=t===W?(n=X,i=$,M):(n=z,i=G,K),l&&g(l).hasClass(V))this._isSliding=!1;else if(!this._triggerSlideEvent(l,o).isDefaultPrevented()&&s&&l){this._isSliding=!0,h&&this.pause(),this._setActiveIndicatorElement(l);var u=g.Event(Q.SLID,{relatedTarget:l,direction:o,from:a,to:c});if(g(this._element).hasClass(Y)){g(l).addClass(i),_.reflow(l),g(s).addClass(n),g(l).addClass(n);var f=parseInt(l.getAttribute("data-interval"),10);this._config.interval=f?(this._config.defaultInterval=this._config.defaultInterval||this._config.interval,f):this._config.defaultInterval||this._config.interval;var d=_.getTransitionDurationFromElement(s);g(s).one(_.TRANSITION_END,function(){g(l).removeClass(n+" "+i).addClass(V),g(s).removeClass(V+" "+i+" "+n),r._isSliding=!1,setTimeout(function(){return g(r._element).trigger(u)},0)}).emulateTransitionEnd(d)}else g(s).removeClass(V),g(l).addClass(V),this._isSliding=!1,g(this._element).trigger(u);h&&this.cycle()}},r._jQueryInterface=function(i){return this.each(function(){var t=g(this).data(j),e=l({},F,g(this).data());"object"==typeof i&&(e=l({},e,i));var n="string"==typeof i?i:e.slide;if(t||(t=new r(this,e),g(this).data(j,t)),"number"==typeof i)t.to(i);else if("string"==typeof n){if("undefined"==typeof t[n])throw new TypeError('No method named "'+n+'"');t[n]()}else e.interval&&e.ride&&(t.pause(),t.cycle())})},r._dataApiClickHandler=function(t){var e=_.getSelectorFromElement(this);if(e){var n=g(e)[0];if(n&&g(n).hasClass(B)){var i=l({},g(n).data(),g(this).data()),o=this.getAttribute("data-slide-to");o&&(i.interval=!1),r._jQueryInterface.call(g(n),i),o&&g(n).data(j).to(o),t.preventDefault()}}},s(r,null,[{key:"VERSION",get:function(){return"4.3.1"}},{key:"Default",get:function(){return F}}]),r}();g(document).on(Q.CLICK_DATA_API,rt,lt._dataApiClickHandler),g(window).on(Q.LOAD_DATA_API,function(){for(var t=[].slice.call(document.querySelectorAll(st)),e=0,n=t.length;e<n;e++){var i=g(t[e]);lt._jQueryInterface.call(i,i.data())}}),g.fn[L]=lt._jQueryInterface,g.fn[L].Constructor=lt,g.fn[L].noConflict=function(){return g.fn[L]=x,lt._jQueryInterface};var ct="collapse",ht="bs.collapse",ut="."+ht,ft=g.fn[ct],dt={toggle:!0,parent:""},gt={toggle:"boolean",parent:"(string|element)"},_t={SHOW:"show"+ut,SHOWN:"shown"+ut,HIDE:"hide"+ut,HIDDEN:"hidden"+ut,CLICK_DATA_API:"click"+ut+".data-api"},mt="show",pt="collapse",vt="collapsing",yt="collapsed",Et="width",Ct="height",Tt=".show, .collapsing",St='[data-toggle="collapse"]',bt=function(){function a(e,t){this._isTransitioning=!1,this._element=e,this._config=this._getConfig(t),this._triggerArray=[].slice.call(document.querySelectorAll('[data-toggle="collapse"][href="#'+e.id+'"],[data-toggle="collapse"][data-target="#'+e.id+'"]'));for(var n=[].slice.call(document.querySelectorAll(St)),i=0,o=n.length;i<o;i++){var r=n[i],s=_.getSelectorFromElement(r),a=[].slice.call(document.querySelectorAll(s)).filter(function(t){return t===e});null!==s&&0<a.length&&(this._selector=s,this._triggerArray.push(r))}this._parent=this._config.parent?this._getParent():null,this._config.parent||this._addAriaAndCollapsedClass(this._element,this._triggerArray),this._config.toggle&&this.toggle()}var t=a.prototype;return t.toggle=function(){g(this._element).hasClass(mt)?this.hide():this.show()},t.show=function(){var t,e,n=this;if(!this._isTransitioning&&!g(this._element).hasClass(mt)&&(this._parent&&0===(t=[].slice.call(this._parent.querySelectorAll(Tt)).filter(function(t){return"string"==typeof n._config.parent?t.getAttribute("data-parent")===n._config.parent:t.classList.contains(pt)})).length&&(t=null),!(t&&(e=g(t).not(this._selector).data(ht))&&e._isTransitioning))){var i=g.Event(_t.SHOW);if(g(this._element).trigger(i),!i.isDefaultPrevented()){t&&(a._jQueryInterface.call(g(t).not(this._selector),"hide"),e||g(t).data(ht,null));var o=this._getDimension();g(this._element).removeClass(pt).addClass(vt),this._element.style[o]=0,this._triggerArray.length&&g(this._triggerArray).removeClass(yt).attr("aria-expanded",!0),this.setTransitioning(!0);var r="scroll"+(o[0].toUpperCase()+o.slice(1)),s=_.getTransitionDurationFromElement(this._element);g(this._element).one(_.TRANSITION_END,function(){g(n._element).removeClass(vt).addClass(pt).addClass(mt),n._element.style[o]="",n.setTransitioning(!1),g(n._element).trigger(_t.SHOWN)}).emulateTransitionEnd(s),this._element.style[o]=this._element[r]+"px"}}},t.hide=function(){var t=this;if(!this._isTransitioning&&g(this._element).hasClass(mt)){var e=g.Event(_t.HIDE);if(g(this._element).trigger(e),!e.isDefaultPrevented()){var n=this._getDimension();this._element.style[n]=this._element.getBoundingClientRect()[n]+"px",_.reflow(this._element),g(this._element).addClass(vt).removeClass(pt).removeClass(mt);var i=this._triggerArray.length;if(0<i)for(var o=0;o<i;o++){var r=this._triggerArray[o],s=_.getSelectorFromElement(r);if(null!==s)g([].slice.call(document.querySelectorAll(s))).hasClass(mt)||g(r).addClass(yt).attr("aria-expanded",!1)}this.setTransitioning(!0);this._element.style[n]="";var a=_.getTransitionDurationFromElement(this._element);g(this._element).one(_.TRANSITION_END,function(){t.setTransitioning(!1),g(t._element).removeClass(vt).addClass(pt).trigger(_t.HIDDEN)}).emulateTransitionEnd(a)}}},t.setTransitioning=function(t){this._isTransitioning=t},t.dispose=function(){g.removeData(this._element,ht),this._config=null,this._parent=null,this._element=null,this._triggerArray=null,this._isTransitioning=null},t._getConfig=function(t){return(t=l({},dt,t)).toggle=Boolean(t.toggle),_.typeCheckConfig(ct,t,gt),t},t._getDimension=function(){return g(this._element).hasClass(Et)?Et:Ct},t._getParent=function(){var t,n=this;_.isElement(this._config.parent)?(t=this._config.parent,"undefined"!=typeof this._config.parent.jquery&&(t=this._config.parent[0])):t=document.querySelector(this._config.parent);var e='[data-toggle="collapse"][data-parent="'+this._config.parent+'"]',i=[].slice.call(t.querySelectorAll(e));return g(i).each(function(t,e){n._addAriaAndCollapsedClass(a._getTargetFromElement(e),[e])}),t},t._addAriaAndCollapsedClass=function(t,e){var n=g(t).hasClass(mt);e.length&&g(e).toggleClass(yt,!n).attr("aria-expanded",n)},a._getTargetFromElement=function(t){var e=_.getSelectorFromElement(t);return e?document.querySelector(e):null},a._jQueryInterface=function(i){return this.each(function(){var t=g(this),e=t.data(ht),n=l({},dt,t.data(),"object"==typeof i&&i?i:{});if(!e&&n.toggle&&/show|hide/.test(i)&&(n.toggle=!1),e||(e=new a(this,n),t.data(ht,e)),"string"==typeof i){if("undefined"==typeof e[i])throw new TypeError('No method named "'+i+'"');e[i]()}})},s(a,null,[{key:"VERSION",get:function(){return"4.3.1"}},{key:"Default",get:function(){return dt}}]),a}();g(document).on(_t.CLICK_DATA_API,St,function(t){"A"===t.currentTarget.tagName&&t.preventDefault();var n=g(this),e=_.getSelectorFromElement(this),i=[].slice.call(document.querySelectorAll(e));g(i).each(function(){var t=g(this),e=t.data(ht)?"toggle":n.data();bt._jQueryInterface.call(t,e)})}),g.fn[ct]=bt._jQueryInterface,g.fn[ct].Constructor=bt,g.fn[ct].noConflict=function(){return g.fn[ct]=ft,bt._jQueryInterface};var It="dropdown",Dt="bs.dropdown",wt="."+Dt,At=".data-api",Nt=g.fn[It],Ot=new RegExp("38|40|27"),kt={HIDE:"hide"+wt,HIDDEN:"hidden"+wt,SHOW:"show"+wt,SHOWN:"shown"+wt,CLICK:"click"+wt,CLICK_DATA_API:"click"+wt+At,KEYDOWN_DATA_API:"keydown"+wt+At,KEYUP_DATA_API:"keyup"+wt+At},Pt="disabled",Lt="show",jt="dropup",Ht="dropright",Rt="dropleft",xt="dropdown-menu-right",Ft="position-static",Ut='[data-toggle="dropdown"]',Wt=".dropdown form",qt=".dropdown-menu",Mt=".navbar-nav",Kt=".dropdown-menu .dropdown-item:not(.disabled):not(:disabled)",Qt="top-start",Bt="top-end",Vt="bottom-start",Yt="bottom-end",zt="right-start",Xt="left-start",$t={offset:0,flip:!0,boundary:"scrollParent",reference:"toggle",display:"dynamic"},Gt={offset:"(number|string|function)",flip:"boolean",boundary:"(string|element)",reference:"(string|element)",display:"string"},Jt=function(){function c(t,e){this._element=t,this._popper=null,this._config=this._getConfig(e),this._menu=this._getMenuElement(),this._inNavbar=this._detectNavbar(),this._addEventListeners()}var t=c.prototype;return t.toggle=function(){if(!this._element.disabled&&!g(this._element).hasClass(Pt)){var t=c._getParentFromElement(this._element),e=g(this._menu).hasClass(Lt);if(c._clearMenus(),!e){var n={relatedTarget:this._element},i=g.Event(kt.SHOW,n);if(g(t).trigger(i),!i.isDefaultPrevented()){if(!this._inNavbar){if("undefined"==typeof u)throw new TypeError("Bootstrap's dropdowns require Popper.js (https://popper.js.org/)");var o=this._element;"parent"===this._config.reference?o=t:_.isElement(this._config.reference)&&(o=this._config.reference,"undefined"!=typeof this._config.reference.jquery&&(o=this._config.reference[0])),"scrollParent"!==this._config.boundary&&g(t).addClass(Ft),this._popper=new u(o,this._menu,this._getPopperConfig())}"ontouchstart"in document.documentElement&&0===g(t).closest(Mt).length&&g(document.body).children().on("mouseover",null,g.noop),this._element.focus(),this._element.setAttribute("aria-expanded",!0),g(this._menu).toggleClass(Lt),g(t).toggleClass(Lt).trigger(g.Event(kt.SHOWN,n))}}}},t.show=function(){if(!(this._element.disabled||g(this._element).hasClass(Pt)||g(this._menu).hasClass(Lt))){var t={relatedTarget:this._element},e=g.Event(kt.SHOW,t),n=c._getParentFromElement(this._element);g(n).trigger(e),e.isDefaultPrevented()||(g(this._menu).toggleClass(Lt),g(n).toggleClass(Lt).trigger(g.Event(kt.SHOWN,t)))}},t.hide=function(){if(!this._element.disabled&&!g(this._element).hasClass(Pt)&&g(this._menu).hasClass(Lt)){var t={relatedTarget:this._element},e=g.Event(kt.HIDE,t),n=c._getParentFromElement(this._element);g(n).trigger(e),e.isDefaultPrevented()||(g(this._menu).toggleClass(Lt),g(n).toggleClass(Lt).trigger(g.Event(kt.HIDDEN,t)))}},t.dispose=function(){g.removeData(this._element,Dt),g(this._element).off(wt),this._element=null,(this._menu=null)!==this._popper&&(this._popper.destroy(),this._popper=null)},t.update=function(){this._inNavbar=this._detectNavbar(),null!==this._popper&&this._popper.scheduleUpdate()},t._addEventListeners=function(){var e=this;g(this._element).on(kt.CLICK,function(t){t.preventDefault(),t.stopPropagation(),e.toggle()})},t._getConfig=function(t){return t=l({},this.constructor.Default,g(this._element).data(),t),_.typeCheckConfig(It,t,this.constructor.DefaultType),t},t._getMenuElement=function(){if(!this._menu){var t=c._getParentFromElement(this._element);t&&(this._menu=t.querySelector(qt))}return this._menu},t._getPlacement=function(){var t=g(this._element.parentNode),e=Vt;return t.hasClass(jt)?(e=Qt,g(this._menu).hasClass(xt)&&(e=Bt)):t.hasClass(Ht)?e=zt:t.hasClass(Rt)?e=Xt:g(this._menu).hasClass(xt)&&(e=Yt),e},t._detectNavbar=function(){return 0<g(this._element).closest(".navbar").length},t._getOffset=function(){var e=this,t={};return"function"==typeof this._config.offset?t.fn=function(t){return t.offsets=l({},t.offsets,e._config.offset(t.offsets,e._element)||{}),t}:t.offset=this._config.offset,t},t._getPopperConfig=function(){var t={placement:this._getPlacement(),modifiers:{offset:this._getOffset(),flip:{enabled:this._config.flip},preventOverflow:{boundariesElement:this._config.boundary}}};return"static"===this._config.display&&(t.modifiers.applyStyle={enabled:!1}),t},c._jQueryInterface=function(e){return this.each(function(){var t=g(this).data(Dt);if(t||(t=new c(this,"object"==typeof e?e:null),g(this).data(Dt,t)),"string"==typeof e){if("undefined"==typeof t[e])throw new TypeError('No method named "'+e+'"');t[e]()}})},c._clearMenus=function(t){if(!t||3!==t.which&&("keyup"!==t.type||9===t.which))for(var e=[].slice.call(document.querySelectorAll(Ut)),n=0,i=e.length;n<i;n++){var o=c._getParentFromElement(e[n]),r=g(e[n]).data(Dt),s={relatedTarget:e[n]};if(t&&"click"===t.type&&(s.clickEvent=t),r){var a=r._menu;if(g(o).hasClass(Lt)&&!(t&&("click"===t.type&&/input|textarea/i.test(t.target.tagName)||"keyup"===t.type&&9===t.which)&&g.contains(o,t.target))){var l=g.Event(kt.HIDE,s);g(o).trigger(l),l.isDefaultPrevented()||("ontouchstart"in document.documentElement&&g(document.body).children().off("mouseover",null,g.noop),e[n].setAttribute("aria-expanded","false"),g(a).removeClass(Lt),g(o).removeClass(Lt).trigger(g.Event(kt.HIDDEN,s)))}}}},c._getParentFromElement=function(t){var e,n=_.getSelectorFromElement(t);return n&&(e=document.querySelector(n)),e||t.parentNode},c._dataApiKeydownHandler=function(t){if((/input|textarea/i.test(t.target.tagName)?!(32===t.which||27!==t.which&&(40!==t.which&&38!==t.which||g(t.target).closest(qt).length)):Ot.test(t.which))&&(t.preventDefault(),t.stopPropagation(),!this.disabled&&!g(this).hasClass(Pt))){var e=c._getParentFromElement(this),n=g(e).hasClass(Lt);if(n&&(!n||27!==t.which&&32!==t.which)){var i=[].slice.call(e.querySelectorAll(Kt));if(0!==i.length){var o=i.indexOf(t.target);38===t.which&&0<o&&o--,40===t.which&&o<i.length-1&&o++,o<0&&(o=0),i[o].focus()}}else{if(27===t.which){var r=e.querySelector(Ut);g(r).trigger("focus")}g(this).trigger("click")}}},s(c,null,[{key:"VERSION",get:function(){return"4.3.1"}},{key:"Default",get:function(){return $t}},{key:"DefaultType",get:function(){return Gt}}]),c}();g(document).on(kt.KEYDOWN_DATA_API,Ut,Jt._dataApiKeydownHandler).on(kt.KEYDOWN_DATA_API,qt,Jt._dataApiKeydownHandler).on(kt.CLICK_DATA_API+" "+kt.KEYUP_DATA_API,Jt._clearMenus).on(kt.CLICK_DATA_API,Ut,function(t){t.preventDefault(),t.stopPropagation(),Jt._jQueryInterface.call(g(this),"toggle")}).on(kt.CLICK_DATA_API,Wt,function(t){t.stopPropagation()}),g.fn[It]=Jt._jQueryInterface,g.fn[It].Constructor=Jt,g.fn[It].noConflict=function(){return g.fn[It]=Nt,Jt._jQueryInterface};var Zt="modal",te="bs.modal",ee="."+te,ne=g.fn[Zt],ie={backdrop:!0,keyboard:!0,focus:!0,show:!0},oe={backdrop:"(boolean|string)",keyboard:"boolean",focus:"boolean",show:"boolean"},re={HIDE:"hide"+ee,HIDDEN:"hidden"+ee,SHOW:"show"+ee,SHOWN:"shown"+ee,FOCUSIN:"focusin"+ee,RESIZE:"resize"+ee,CLICK_DISMISS:"click.dismiss"+ee,KEYDOWN_DISMISS:"keydown.dismiss"+ee,MOUSEUP_DISMISS:"mouseup.dismiss"+ee,MOUSEDOWN_DISMISS:"mousedown.dismiss"+ee,CLICK_DATA_API:"click"+ee+".data-api"},se="modal-dialog-scrollable",ae="modal-scrollbar-measure",le="modal-backdrop",ce="modal-open",he="fade",ue="show",fe=".modal-dialog",de=".modal-body",ge='[data-toggle="modal"]',_e='[data-dismiss="modal"]',me=".fixed-top, .fixed-bottom, .is-fixed, .sticky-top",pe=".sticky-top",ve=function(){function o(t,e){this._config=this._getConfig(e),this._element=t,this._dialog=t.querySelector(fe),this._backdrop=null,this._isShown=!1,this._isBodyOverflowing=!1,this._ignoreBackdropClick=!1,this._isTransitioning=!1,this._scrollbarWidth=0}var t=o.prototype;return t.toggle=function(t){return this._isShown?this.hide():this.show(t)},t.show=function(t){var e=this;if(!this._isShown&&!this._isTransitioning){g(this._element).hasClass(he)&&(this._isTransitioning=!0);var n=g.Event(re.SHOW,{relatedTarget:t});g(this._element).trigger(n),this._isShown||n.isDefaultPrevented()||(this._isShown=!0,this._checkScrollbar(),this._setScrollbar(),this._adjustDialog(),this._setEscapeEvent(),this._setResizeEvent(),g(this._element).on(re.CLICK_DISMISS,_e,function(t){return e.hide(t)}),g(this._dialog).on(re.MOUSEDOWN_DISMISS,function(){g(e._element).one(re.MOUSEUP_DISMISS,function(t){g(t.target).is(e._element)&&(e._ignoreBackdropClick=!0)})}),this._showBackdrop(function(){return e._showElement(t)}))}},t.hide=function(t){var e=this;if(t&&t.preventDefault(),this._isShown&&!this._isTransitioning){var n=g.Event(re.HIDE);if(g(this._element).trigger(n),this._isShown&&!n.isDefaultPrevented()){this._isShown=!1;var i=g(this._element).hasClass(he);if(i&&(this._isTransitioning=!0),this._setEscapeEvent(),this._setResizeEvent(),g(document).off(re.FOCUSIN),g(this._element).removeClass(ue),g(this._element).off(re.CLICK_DISMISS),g(this._dialog).off(re.MOUSEDOWN_DISMISS),i){var o=_.getTransitionDurationFromElement(this._element);g(this._element).one(_.TRANSITION_END,function(t){return e._hideModal(t)}).emulateTransitionEnd(o)}else this._hideModal()}}},t.dispose=function(){[window,this._element,this._dialog].forEach(function(t){return g(t).off(ee)}),g(document).off(re.FOCUSIN),g.removeData(this._element,te),this._config=null,this._element=null,this._dialog=null,this._backdrop=null,this._isShown=null,this._isBodyOverflowing=null,this._ignoreBackdropClick=null,this._isTransitioning=null,this._scrollbarWidth=null},t.handleUpdate=function(){this._adjustDialog()},t._getConfig=function(t){return t=l({},ie,t),_.typeCheckConfig(Zt,t,oe),t},t._showElement=function(t){var e=this,n=g(this._element).hasClass(he);this._element.parentNode&&this._element.parentNode.nodeType===Node.ELEMENT_NODE||document.body.appendChild(this._element),this._element.style.display="block",this._element.removeAttribute("aria-hidden"),this._element.setAttribute("aria-modal",!0),g(this._dialog).hasClass(se)?this._dialog.querySelector(de).scrollTop=0:this._element.scrollTop=0,n&&_.reflow(this._element),g(this._element).addClass(ue),this._config.focus&&this._enforceFocus();var i=g.Event(re.SHOWN,{relatedTarget:t}),o=function(){e._config.focus&&e._element.focus(),e._isTransitioning=!1,g(e._element).trigger(i)};if(n){var r=_.getTransitionDurationFromElement(this._dialog);g(this._dialog).one(_.TRANSITION_END,o).emulateTransitionEnd(r)}else o()},t._enforceFocus=function(){var e=this;g(document).off(re.FOCUSIN).on(re.FOCUSIN,function(t){document!==t.target&&e._element!==t.target&&0===g(e._element).has(t.target).length&&e._element.focus()})},t._setEscapeEvent=function(){var e=this;this._isShown&&this._config.keyboard?g(this._element).on(re.KEYDOWN_DISMISS,function(t){27===t.which&&(t.preventDefault(),e.hide())}):this._isShown||g(this._element).off(re.KEYDOWN_DISMISS)},t._setResizeEvent=function(){var e=this;this._isShown?g(window).on(re.RESIZE,function(t){return e.handleUpdate(t)}):g(window).off(re.RESIZE)},t._hideModal=function(){var t=this;this._element.style.display="none",this._element.setAttribute("aria-hidden",!0),this._element.removeAttribute("aria-modal"),this._isTransitioning=!1,this._showBackdrop(function(){g(document.body).removeClass(ce),t._resetAdjustments(),t._resetScrollbar(),g(t._element).trigger(re.HIDDEN)})},t._removeBackdrop=function(){this._backdrop&&(g(this._backdrop).remove(),this._backdrop=null)},t._showBackdrop=function(t){var e=this,n=g(this._element).hasClass(he)?he:"";if(this._isShown&&this._config.backdrop){if(this._backdrop=document.createElement("div"),this._backdrop.className=le,n&&this._backdrop.classList.add(n),g(this._backdrop).appendTo(document.body),g(this._element).on(re.CLICK_DISMISS,function(t){e._ignoreBackdropClick?e._ignoreBackdropClick=!1:t.target===t.currentTarget&&("static"===e._config.backdrop?e._element.focus():e.hide())}),n&&_.reflow(this._backdrop),g(this._backdrop).addClass(ue),!t)return;if(!n)return void t();var i=_.getTransitionDurationFromElement(this._backdrop);g(this._backdrop).one(_.TRANSITION_END,t).emulateTransitionEnd(i)}else if(!this._isShown&&this._backdrop){g(this._backdrop).removeClass(ue);var o=function(){e._removeBackdrop(),t&&t()};if(g(this._element).hasClass(he)){var r=_.getTransitionDurationFromElement(this._backdrop);g(this._backdrop).one(_.TRANSITION_END,o).emulateTransitionEnd(r)}else o()}else t&&t()},t._adjustDialog=function(){var t=this._element.scrollHeight>document.documentElement.clientHeight;!this._isBodyOverflowing&&t&&(this._element.style.paddingLeft=this._scrollbarWidth+"px"),this._isBodyOverflowing&&!t&&(this._element.style.paddingRight=this._scrollbarWidth+"px")},t._resetAdjustments=function(){this._element.style.paddingLeft="",this._element.style.paddingRight=""},t._checkScrollbar=function(){var t=document.body.getBoundingClientRect();this._isBodyOverflowing=t.left+t.right<window.innerWidth,this._scrollbarWidth=this._getScrollbarWidth()},t._setScrollbar=function(){var o=this;if(this._isBodyOverflowing){var t=[].slice.call(document.querySelectorAll(me)),e=[].slice.call(document.querySelectorAll(pe));g(t).each(function(t,e){var n=e.style.paddingRight,i=g(e).css("padding-right");g(e).data("padding-right",n).css("padding-right",parseFloat(i)+o._scrollbarWidth+"px")}),g(e).each(function(t,e){var n=e.style.marginRight,i=g(e).css("margin-right");g(e).data("margin-right",n).css("margin-right",parseFloat(i)-o._scrollbarWidth+"px")});var n=document.body.style.paddingRight,i=g(document.body).css("padding-right");g(document.body).data("padding-right",n).css("padding-right",parseFloat(i)+this._scrollbarWidth+"px")}g(document.body).addClass(ce)},t._resetScrollbar=function(){var t=[].slice.call(document.querySelectorAll(me));g(t).each(function(t,e){var n=g(e).data("padding-right");g(e).removeData("padding-right"),e.style.paddingRight=n||""});var e=[].slice.call(document.querySelectorAll(""+pe));g(e).each(function(t,e){var n=g(e).data("margin-right");"undefined"!=typeof n&&g(e).css("margin-right",n).removeData("margin-right")});var n=g(document.body).data("padding-right");g(document.body).removeData("padding-right"),document.body.style.paddingRight=n||""},t._getScrollbarWidth=function(){var t=document.createElement("div");t.className=ae,document.body.appendChild(t);var e=t.getBoundingClientRect().width-t.clientWidth;return document.body.removeChild(t),e},o._jQueryInterface=function(n,i){return this.each(function(){var t=g(this).data(te),e=l({},ie,g(this).data(),"object"==typeof n&&n?n:{});if(t||(t=new o(this,e),g(this).data(te,t)),"string"==typeof n){if("undefined"==typeof t[n])throw new TypeError('No method named "'+n+'"');t[n](i)}else e.show&&t.show(i)})},s(o,null,[{key:"VERSION",get:function(){return"4.3.1"}},{key:"Default",get:function(){return ie}}]),o}();g(document).on(re.CLICK_DATA_API,ge,function(t){var e,n=this,i=_.getSelectorFromElement(this);i&&(e=document.querySelector(i));var o=g(e).data(te)?"toggle":l({},g(e).data(),g(this).data());"A"!==this.tagName&&"AREA"!==this.tagName||t.preventDefault();var r=g(e).one(re.SHOW,function(t){t.isDefaultPrevented()||r.one(re.HIDDEN,function(){g(n).is(":visible")&&n.focus()})});ve._jQueryInterface.call(g(e),o,this)}),g.fn[Zt]=ve._jQueryInterface,g.fn[Zt].Constructor=ve,g.fn[Zt].noConflict=function(){return g.fn[Zt]=ne,ve._jQueryInterface};var ye=["background","cite","href","itemtype","longdesc","poster","src","xlink:href"],Ee={"*":["class","dir","id","lang","role",/^aria-[\w-]*$/i],a:["target","href","title","rel"],area:[],b:[],br:[],col:[],code:[],div:[],em:[],hr:[],h1:[],h2:[],h3:[],h4:[],h5:[],h6:[],i:[],img:["src","alt","title","width","height"],li:[],ol:[],p:[],pre:[],s:[],small:[],span:[],sub:[],sup:[],strong:[],u:[],ul:[]},Ce=/^(?:(?:https?|mailto|ftp|tel|file):|[^&:/?#]*(?:[/?#]|$))/gi,Te=/^data:(?:image\/(?:bmp|gif|jpeg|jpg|png|tiff|webp)|video\/(?:mpeg|mp4|ogg|webm)|audio\/(?:mp3|oga|ogg|opus));base64,[a-z0-9+/]+=*$/i;function Se(t,s,e){if(0===t.length)return t;if(e&&"function"==typeof e)return e(t);for(var n=(new window.DOMParser).parseFromString(t,"text/html"),a=Object.keys(s),l=[].slice.call(n.body.querySelectorAll("*")),i=function(t,e){var n=l[t],i=n.nodeName.toLowerCase();if(-1===a.indexOf(n.nodeName.toLowerCase()))return n.parentNode.removeChild(n),"continue";var o=[].slice.call(n.attributes),r=[].concat(s["*"]||[],s[i]||[]);o.forEach(function(t){(function(t,e){var n=t.nodeName.toLowerCase();if(-1!==e.indexOf(n))return-1===ye.indexOf(n)||Boolean(t.nodeValue.match(Ce)||t.nodeValue.match(Te));for(var i=e.filter(function(t){return t instanceof RegExp}),o=0,r=i.length;o<r;o++)if(n.match(i[o]))return!0;return!1})(t,r)||n.removeAttribute(t.nodeName)})},o=0,r=l.length;o<r;o++)i(o);return n.body.innerHTML}var be="tooltip",Ie="bs.tooltip",De="."+Ie,we=g.fn[be],Ae="bs-tooltip",Ne=new RegExp("(^|\\s)"+Ae+"\\S+","g"),Oe=["sanitize","whiteList","sanitizeFn"],ke={animation:"boolean",template:"string",title:"(string|element|function)",trigger:"string",delay:"(number|object)",html:"boolean",selector:"(string|boolean)",placement:"(string|function)",offset:"(number|string|function)",container:"(string|element|boolean)",fallbackPlacement:"(string|array)",boundary:"(string|element)",sanitize:"boolean",sanitizeFn:"(null|function)",whiteList:"object"},Pe={AUTO:"auto",TOP:"top",RIGHT:"right",BOTTOM:"bottom",LEFT:"left"},Le={animation:!0,template:'<div class="tooltip" role="tooltip"><div class="arrow"></div><div class="tooltip-inner"></div></div>',trigger:"hover focus",title:"",delay:0,html:!1,selector:!1,placement:"top",offset:0,container:!1,fallbackPlacement:"flip",boundary:"scrollParent",sanitize:!0,sanitizeFn:null,whiteList:Ee},je="show",He="out",Re={HIDE:"hide"+De,HIDDEN:"hidden"+De,SHOW:"show"+De,SHOWN:"shown"+De,INSERTED:"inserted"+De,CLICK:"click"+De,FOCUSIN:"focusin"+De,FOCUSOUT:"focusout"+De,MOUSEENTER:"mouseenter"+De,MOUSELEAVE:"mouseleave"+De},xe="fade",Fe="show",Ue=".tooltip-inner",We=".arrow",qe="hover",Me="focus",Ke="click",Qe="manual",Be=function(){function i(t,e){if("undefined"==typeof u)throw new TypeError("Bootstrap's tooltips require Popper.js (https://popper.js.org/)");this._isEnabled=!0,this._timeout=0,this._hoverState="",this._activeTrigger={},this._popper=null,this.element=t,this.config=this._getConfig(e),this.tip=null,this._setListeners()}var t=i.prototype;return t.enable=function(){this._isEnabled=!0},t.disable=function(){this._isEnabled=!1},t.toggleEnabled=function(){this._isEnabled=!this._isEnabled},t.toggle=function(t){if(this._isEnabled)if(t){var e=this.constructor.DATA_KEY,n=g(t.currentTarget).data(e);n||(n=new this.constructor(t.currentTarget,this._getDelegateConfig()),g(t.currentTarget).data(e,n)),n._activeTrigger.click=!n._activeTrigger.click,n._isWithActiveTrigger()?n._enter(null,n):n._leave(null,n)}else{if(g(this.getTipElement()).hasClass(Fe))return void this._leave(null,this);this._enter(null,this)}},t.dispose=function(){clearTimeout(this._timeout),g.removeData(this.element,this.constructor.DATA_KEY),g(this.element).off(this.constructor.EVENT_KEY),g(this.element).closest(".modal").off("hide.bs.modal"),this.tip&&g(this.tip).remove(),this._isEnabled=null,this._timeout=null,this._hoverState=null,(this._activeTrigger=null)!==this._popper&&this._popper.destroy(),this._popper=null,this.element=null,this.config=null,this.tip=null},t.show=function(){var e=this;if("none"===g(this.element).css("display"))throw new Error("Please use show on visible elements");var t=g.Event(this.constructor.Event.SHOW);if(this.isWithContent()&&this._isEnabled){g(this.element).trigger(t);var n=_.findShadowRoot(this.element),i=g.contains(null!==n?n:this.element.ownerDocument.documentElement,this.element);if(t.isDefaultPrevented()||!i)return;var o=this.getTipElement(),r=_.getUID(this.constructor.NAME);o.setAttribute("id",r),this.element.setAttribute("aria-describedby",r),this.setContent(),this.config.animation&&g(o).addClass(xe);var s="function"==typeof this.config.placement?this.config.placement.call(this,o,this.element):this.config.placement,a=this._getAttachment(s);this.addAttachmentClass(a);var l=this._getContainer();g(o).data(this.constructor.DATA_KEY,this),g.contains(this.element.ownerDocument.documentElement,this.tip)||g(o).appendTo(l),g(this.element).trigger(this.constructor.Event.INSERTED),this._popper=new u(this.element,o,{placement:a,modifiers:{offset:this._getOffset(),flip:{behavior:this.config.fallbackPlacement},arrow:{element:We},preventOverflow:{boundariesElement:this.config.boundary}},onCreate:function(t){t.originalPlacement!==t.placement&&e._handlePopperPlacementChange(t)},onUpdate:function(t){return e._handlePopperPlacementChange(t)}}),g(o).addClass(Fe),"ontouchstart"in document.documentElement&&g(document.body).children().on("mouseover",null,g.noop);var c=function(){e.config.animation&&e._fixTransition();var t=e._hoverState;e._hoverState=null,g(e.element).trigger(e.constructor.Event.SHOWN),t===He&&e._leave(null,e)};if(g(this.tip).hasClass(xe)){var h=_.getTransitionDurationFromElement(this.tip);g(this.tip).one(_.TRANSITION_END,c).emulateTransitionEnd(h)}else c()}},t.hide=function(t){var e=this,n=this.getTipElement(),i=g.Event(this.constructor.Event.HIDE),o=function(){e._hoverState!==je&&n.parentNode&&n.parentNode.removeChild(n),e._cleanTipClass(),e.element.removeAttribute("aria-describedby"),g(e.element).trigger(e.constructor.Event.HIDDEN),null!==e._popper&&e._popper.destroy(),t&&t()};if(g(this.element).trigger(i),!i.isDefaultPrevented()){if(g(n).removeClass(Fe),"ontouchstart"in document.documentElement&&g(document.body).children().off("mouseover",null,g.noop),this._activeTrigger[Ke]=!1,this._activeTrigger[Me]=!1,this._activeTrigger[qe]=!1,g(this.tip).hasClass(xe)){var r=_.getTransitionDurationFromElement(n);g(n).one(_.TRANSITION_END,o).emulateTransitionEnd(r)}else o();this._hoverState=""}},t.update=function(){null!==this._popper&&this._popper.scheduleUpdate()},t.isWithContent=function(){return Boolean(this.getTitle())},t.addAttachmentClass=function(t){g(this.getTipElement()).addClass(Ae+"-"+t)},t.getTipElement=function(){return this.tip=this.tip||g(this.config.template)[0],this.tip},t.setContent=function(){var t=this.getTipElement();this.setElementContent(g(t.querySelectorAll(Ue)),this.getTitle()),g(t).removeClass(xe+" "+Fe)},t.setElementContent=function(t,e){"object"!=typeof e||!e.nodeType&&!e.jquery?this.config.html?(this.config.sanitize&&(e=Se(e,this.config.whiteList,this.config.sanitizeFn)),t.html(e)):t.text(e):this.config.html?g(e).parent().is(t)||t.empty().append(e):t.text(g(e).text())},t.getTitle=function(){var t=this.element.getAttribute("data-original-title");return t||(t="function"==typeof this.config.title?this.config.title.call(this.element):this.config.title),t},t._getOffset=function(){var e=this,t={};return"function"==typeof this.config.offset?t.fn=function(t){return t.offsets=l({},t.offsets,e.config.offset(t.offsets,e.element)||{}),t}:t.offset=this.config.offset,t},t._getContainer=function(){return!1===this.config.container?document.body:_.isElement(this.config.container)?g(this.config.container):g(document).find(this.config.container)},t._getAttachment=function(t){return Pe[t.toUpperCase()]},t._setListeners=function(){var i=this;this.config.trigger.split(" ").forEach(function(t){if("click"===t)g(i.element).on(i.constructor.Event.CLICK,i.config.selector,function(t){return i.toggle(t)});else if(t!==Qe){var e=t===qe?i.constructor.Event.MOUSEENTER:i.constructor.Event.FOCUSIN,n=t===qe?i.constructor.Event.MOUSELEAVE:i.constructor.Event.FOCUSOUT;g(i.element).on(e,i.config.selector,function(t){return i._enter(t)}).on(n,i.config.selector,function(t){return i._leave(t)})}}),g(this.element).closest(".modal").on("hide.bs.modal",function(){i.element&&i.hide()}),this.config.selector?this.config=l({},this.config,{trigger:"manual",selector:""}):this._fixTitle()},t._fixTitle=function(){var t=typeof this.element.getAttribute("data-original-title");(this.element.getAttribute("title")||"string"!==t)&&(this.element.setAttribute("data-original-title",this.element.getAttribute("title")||""),this.element.setAttribute("title",""))},t._enter=function(t,e){var n=this.constructor.DATA_KEY;(e=e||g(t.currentTarget).data(n))||(e=new this.constructor(t.currentTarget,this._getDelegateConfig()),g(t.currentTarget).data(n,e)),t&&(e._activeTrigger["focusin"===t.type?Me:qe]=!0),g(e.getTipElement()).hasClass(Fe)||e._hoverState===je?e._hoverState=je:(clearTimeout(e._timeout),e._hoverState=je,e.config.delay&&e.config.delay.show?e._timeout=setTimeout(function(){e._hoverState===je&&e.show()},e.config.delay.show):e.show())},t._leave=function(t,e){var n=this.constructor.DATA_KEY;(e=e||g(t.currentTarget).data(n))||(e=new this.constructor(t.currentTarget,this._getDelegateConfig()),g(t.currentTarget).data(n,e)),t&&(e._activeTrigger["focusout"===t.type?Me:qe]=!1),e._isWithActiveTrigger()||(clearTimeout(e._timeout),e._hoverState=He,e.config.delay&&e.config.delay.hide?e._timeout=setTimeout(function(){e._hoverState===He&&e.hide()},e.config.delay.hide):e.hide())},t._isWithActiveTrigger=function(){for(var t in this._activeTrigger)if(this._activeTrigger[t])return!0;return!1},t._getConfig=function(t){var e=g(this.element).data();return Object.keys(e).forEach(function(t){-1!==Oe.indexOf(t)&&delete e[t]}),"number"==typeof(t=l({},this.constructor.Default,e,"object"==typeof t&&t?t:{})).delay&&(t.delay={show:t.delay,hide:t.delay}),"number"==typeof t.title&&(t.title=t.title.toString()),"number"==typeof t.content&&(t.content=t.content.toString()),_.typeCheckConfig(be,t,this.constructor.DefaultType),t.sanitize&&(t.template=Se(t.template,t.whiteList,t.sanitizeFn)),t},t._getDelegateConfig=function(){var t={};if(this.config)for(var e in this.config)this.constructor.Default[e]!==this.config[e]&&(t[e]=this.config[e]);return t},t._cleanTipClass=function(){var t=g(this.getTipElement()),e=t.attr("class").match(Ne);null!==e&&e.length&&t.removeClass(e.join(""))},t._handlePopperPlacementChange=function(t){var e=t.instance;this.tip=e.popper,this._cleanTipClass(),this.addAttachmentClass(this._getAttachment(t.placement))},t._fixTransition=function(){var t=this.getTipElement(),e=this.config.animation;null===t.getAttribute("x-placement")&&(g(t).removeClass(xe),this.config.animation=!1,this.hide(),this.show(),this.config.animation=e)},i._jQueryInterface=function(n){return this.each(function(){var t=g(this).data(Ie),e="object"==typeof n&&n;if((t||!/dispose|hide/.test(n))&&(t||(t=new i(this,e),g(this).data(Ie,t)),"string"==typeof n)){if("undefined"==typeof t[n])throw new TypeError('No method named "'+n+'"');t[n]()}})},s(i,null,[{key:"VERSION",get:function(){return"4.3.1"}},{key:"Default",get:function(){return Le}},{key:"NAME",get:function(){return be}},{key:"DATA_KEY",get:function(){return Ie}},{key:"Event",get:function(){return Re}},{key:"EVENT_KEY",get:function(){return De}},{key:"DefaultType",get:function(){return ke}}]),i}();g.fn[be]=Be._jQueryInterface,g.fn[be].Constructor=Be,g.fn[be].noConflict=function(){return g.fn[be]=we,Be._jQueryInterface};var Ve="popover",Ye="bs.popover",ze="."+Ye,Xe=g.fn[Ve],$e="bs-popover",Ge=new RegExp("(^|\\s)"+$e+"\\S+","g"),Je=l({},Be.Default,{placement:"right",trigger:"click",content:"",template:'<div class="popover" role="tooltip"><div class="arrow"></div><h3 class="popover-header"></h3><div class="popover-body"></div></div>'}),Ze=l({},Be.DefaultType,{content:"(string|element|function)"}),tn="fade",en="show",nn=".popover-header",on=".popover-body",rn={HIDE:"hide"+ze,HIDDEN:"hidden"+ze,SHOW:"show"+ze,SHOWN:"shown"+ze,INSERTED:"inserted"+ze,CLICK:"click"+ze,FOCUSIN:"focusin"+ze,FOCUSOUT:"focusout"+ze,MOUSEENTER:"mouseenter"+ze,MOUSELEAVE:"mouseleave"+ze},sn=function(t){var e,n;function i(){return t.apply(this,arguments)||this}n=t,(e=i).prototype=Object.create(n.prototype),(e.prototype.constructor=e).__proto__=n;var o=i.prototype;return o.isWithContent=function(){return this.getTitle()||this._getContent()},o.addAttachmentClass=function(t){g(this.getTipElement()).addClass($e+"-"+t)},o.getTipElement=function(){return this.tip=this.tip||g(this.config.template)[0],this.tip},o.setContent=function(){var t=g(this.getTipElement());this.setElementContent(t.find(nn),this.getTitle());var e=this._getContent();"function"==typeof e&&(e=e.call(this.element)),this.setElementContent(t.find(on),e),t.removeClass(tn+" "+en)},o._getContent=function(){return this.element.getAttribute("data-content")||this.config.content},o._cleanTipClass=function(){var t=g(this.getTipElement()),e=t.attr("class").match(Ge);null!==e&&0<e.length&&t.removeClass(e.join(""))},i._jQueryInterface=function(n){return this.each(function(){var t=g(this).data(Ye),e="object"==typeof n?n:null;if((t||!/dispose|hide/.test(n))&&(t||(t=new i(this,e),g(this).data(Ye,t)),"string"==typeof n)){if("undefined"==typeof t[n])throw new TypeError('No method named "'+n+'"');t[n]()}})},s(i,null,[{key:"VERSION",get:function(){return"4.3.1"}},{key:"Default",get:function(){return Je}},{key:"NAME",get:function(){return Ve}},{key:"DATA_KEY",get:function(){return Ye}},{key:"Event",get:function(){return rn}},{key:"EVENT_KEY",get:function(){return ze}},{key:"DefaultType",get:function(){return Ze}}]),i}(Be);g.fn[Ve]=sn._jQueryInterface,g.fn[Ve].Constructor=sn,g.fn[Ve].noConflict=function(){return g.fn[Ve]=Xe,sn._jQueryInterface};var an="scrollspy",ln="bs.scrollspy",cn="."+ln,hn=g.fn[an],un={offset:10,method:"auto",target:""},fn={offset:"number",method:"string",target:"(string|element)"},dn={ACTIVATE:"activate"+cn,SCROLL:"scroll"+cn,LOAD_DATA_API:"load"+cn+".data-api"},gn="dropdown-item",_n="active",mn='[data-spy="scroll"]',pn=".nav, .list-group",vn=".nav-link",yn=".nav-item",En=".list-group-item",Cn=".dropdown",Tn=".dropdown-item",Sn=".dropdown-toggle",bn="offset",In="position",Dn=function(){function n(t,e){var n=this;this._element=t,this._scrollElement="BODY"===t.tagName?window:t,this._config=this._getConfig(e),this._selector=this._config.target+" "+vn+","+this._config.target+" "+En+","+this._config.target+" "+Tn,this._offsets=[],this._targets=[],this._activeTarget=null,this._scrollHeight=0,g(this._scrollElement).on(dn.SCROLL,function(t){return n._process(t)}),this.refresh(),this._process()}var t=n.prototype;return t.refresh=function(){var e=this,t=this._scrollElement===this._scrollElement.window?bn:In,o="auto"===this._config.method?t:this._config.method,r=o===In?this._getScrollTop():0;this._offsets=[],this._targets=[],this._scrollHeight=this._getScrollHeight(),[].slice.call(document.querySelectorAll(this._selector)).map(function(t){var e,n=_.getSelectorFromElement(t);if(n&&(e=document.querySelector(n)),e){var i=e.getBoundingClientRect();if(i.width||i.height)return[g(e)[o]().top+r,n]}return null}).filter(function(t){return t}).sort(function(t,e){return t[0]-e[0]}).forEach(function(t){e._offsets.push(t[0]),e._targets.push(t[1])})},t.dispose=function(){g.removeData(this._element,ln),g(this._scrollElement).off(cn),this._element=null,this._scrollElement=null,this._config=null,this._selector=null,this._offsets=null,this._targets=null,this._activeTarget=null,this._scrollHeight=null},t._getConfig=function(t){if("string"!=typeof(t=l({},un,"object"==typeof t&&t?t:{})).target){var e=g(t.target).attr("id");e||(e=_.getUID(an),g(t.target).attr("id",e)),t.target="#"+e}return _.typeCheckConfig(an,t,fn),t},t._getScrollTop=function(){return this._scrollElement===window?this._scrollElement.pageYOffset:this._scrollElement.scrollTop},t._getScrollHeight=function(){return this._scrollElement.scrollHeight||Math.max(document.body.scrollHeight,document.documentElement.scrollHeight)},t._getOffsetHeight=function(){return this._scrollElement===window?window.innerHeight:this._scrollElement.getBoundingClientRect().height},t._process=function(){var t=this._getScrollTop()+this._config.offset,e=this._getScrollHeight(),n=this._config.offset+e-this._getOffsetHeight();if(this._scrollHeight!==e&&this.refresh(),n<=t){var i=this._targets[this._targets.length-1];this._activeTarget!==i&&this._activate(i)}else{if(this._activeTarget&&t<this._offsets[0]&&0<this._offsets[0])return this._activeTarget=null,void this._clear();for(var o=this._offsets.length;o--;){this._activeTarget!==this._targets[o]&&t>=this._offsets[o]&&("undefined"==typeof this._offsets[o+1]||t<this._offsets[o+1])&&this._activate(this._targets[o])}}},t._activate=function(e){this._activeTarget=e,this._clear();var t=this._selector.split(",").map(function(t){return t+'[data-target="'+e+'"],'+t+'[href="'+e+'"]'}),n=g([].slice.call(document.querySelectorAll(t.join(","))));n.hasClass(gn)?(n.closest(Cn).find(Sn).addClass(_n),n.addClass(_n)):(n.addClass(_n),n.parents(pn).prev(vn+", "+En).addClass(_n),n.parents(pn).prev(yn).children(vn).addClass(_n)),g(this._scrollElement).trigger(dn.ACTIVATE,{relatedTarget:e})},t._clear=function(){[].slice.call(document.querySelectorAll(this._selector)).filter(function(t){return t.classList.contains(_n)}).forEach(function(t){return t.classList.remove(_n)})},n._jQueryInterface=function(e){return this.each(function(){var t=g(this).data(ln);if(t||(t=new n(this,"object"==typeof e&&e),g(this).data(ln,t)),"string"==typeof e){if("undefined"==typeof t[e])throw new TypeError('No method named "'+e+'"');t[e]()}})},s(n,null,[{key:"VERSION",get:function(){return"4.3.1"}},{key:"Default",get:function(){return un}}]),n}();g(window).on(dn.LOAD_DATA_API,function(){for(var t=[].slice.call(document.querySelectorAll(mn)),e=t.length;e--;){var n=g(t[e]);Dn._jQueryInterface.call(n,n.data())}}),g.fn[an]=Dn._jQueryInterface,g.fn[an].Constructor=Dn,g.fn[an].noConflict=function(){return g.fn[an]=hn,Dn._jQueryInterface};var wn="bs.tab",An="."+wn,Nn=g.fn.tab,On={HIDE:"hide"+An,HIDDEN:"hidden"+An,SHOW:"show"+An,SHOWN:"shown"+An,CLICK_DATA_API:"click"+An+".data-api"},kn="dropdown-menu",Pn="active",Ln="disabled",jn="fade",Hn="show",Rn=".dropdown",xn=".nav, .list-group",Fn=".active",Un="> li > .active",Wn='[data-toggle="tab"], [data-toggle="pill"], [data-toggle="list"]',qn=".dropdown-toggle",Mn="> .dropdown-menu .active",Kn=function(){function i(t){this._element=t}var t=i.prototype;return t.show=function(){var n=this;if(!(this._element.parentNode&&this._element.parentNode.nodeType===Node.ELEMENT_NODE&&g(this._element).hasClass(Pn)||g(this._element).hasClass(Ln))){var t,i,e=g(this._element).closest(xn)[0],o=_.getSelectorFromElement(this._element);if(e){var r="UL"===e.nodeName||"OL"===e.nodeName?Un:Fn;i=(i=g.makeArray(g(e).find(r)))[i.length-1]}var s=g.Event(On.HIDE,{relatedTarget:this._element}),a=g.Event(On.SHOW,{relatedTarget:i});if(i&&g(i).trigger(s),g(this._element).trigger(a),!a.isDefaultPrevented()&&!s.isDefaultPrevented()){o&&(t=document.querySelector(o)),this._activate(this._element,e);var l=function(){var t=g.Event(On.HIDDEN,{relatedTarget:n._element}),e=g.Event(On.SHOWN,{relatedTarget:i});g(i).trigger(t),g(n._element).trigger(e)};t?this._activate(t,t.parentNode,l):l()}}},t.dispose=function(){g.removeData(this._element,wn),this._element=null},t._activate=function(t,e,n){var i=this,o=(!e||"UL"!==e.nodeName&&"OL"!==e.nodeName?g(e).children(Fn):g(e).find(Un))[0],r=n&&o&&g(o).hasClass(jn),s=function(){return i._transitionComplete(t,o,n)};if(o&&r){var a=_.getTransitionDurationFromElement(o);g(o).removeClass(Hn).one(_.TRANSITION_END,s).emulateTransitionEnd(a)}else s()},t._transitionComplete=function(t,e,n){if(e){g(e).removeClass(Pn);var i=g(e.parentNode).find(Mn)[0];i&&g(i).removeClass(Pn),"tab"===e.getAttribute("role")&&e.setAttribute("aria-selected",!1)}if(g(t).addClass(Pn),"tab"===t.getAttribute("role")&&t.setAttribute("aria-selected",!0),_.reflow(t),t.classList.contains(jn)&&t.classList.add(Hn),t.parentNode&&g(t.parentNode).hasClass(kn)){var o=g(t).closest(Rn)[0];if(o){var r=[].slice.call(o.querySelectorAll(qn));g(r).addClass(Pn)}t.setAttribute("aria-expanded",!0)}n&&n()},i._jQueryInterface=function(n){return this.each(function(){var t=g(this),e=t.data(wn);if(e||(e=new i(this),t.data(wn,e)),"string"==typeof n){if("undefined"==typeof e[n])throw new TypeError('No method named "'+n+'"');e[n]()}})},s(i,null,[{key:"VERSION",get:function(){return"4.3.1"}}]),i}();g(document).on(On.CLICK_DATA_API,Wn,function(t){t.preventDefault(),Kn._jQueryInterface.call(g(this),"show")}),g.fn.tab=Kn._jQueryInterface,g.fn.tab.Constructor=Kn,g.fn.tab.noConflict=function(){return g.fn.tab=Nn,Kn._jQueryInterface};var Qn="toast",Bn="bs.toast",Vn="."+Bn,Yn=g.fn[Qn],zn={CLICK_DISMISS:"click.dismiss"+Vn,HIDE:"hide"+Vn,HIDDEN:"hidden"+Vn,SHOW:"show"+Vn,SHOWN:"shown"+Vn},Xn="fade",$n="hide",Gn="show",Jn="showing",Zn={animation:"boolean",autohide:"boolean",delay:"number"},ti={animation:!0,autohide:!0,delay:500},ei='[data-dismiss="toast"]',ni=function(){function i(t,e){this._element=t,this._config=this._getConfig(e),this._timeout=null,this._setListeners()}var t=i.prototype;return t.show=function(){var t=this;g(this._element).trigger(zn.SHOW),this._config.animation&&this._element.classList.add(Xn);var e=function(){t._element.classList.remove(Jn),t._element.classList.add(Gn),g(t._element).trigger(zn.SHOWN),t._config.autohide&&t.hide()};if(this._element.classList.remove($n),this._element.classList.add(Jn),this._config.animation){var n=_.getTransitionDurationFromElement(this._element);g(this._element).one(_.TRANSITION_END,e).emulateTransitionEnd(n)}else e()},t.hide=function(t){var e=this;this._element.classList.contains(Gn)&&(g(this._element).trigger(zn.HIDE),t?this._close():this._timeout=setTimeout(function(){e._close()},this._config.delay))},t.dispose=function(){clearTimeout(this._timeout),this._timeout=null,this._element.classList.contains(Gn)&&this._element.classList.remove(Gn),g(this._element).off(zn.CLICK_DISMISS),g.removeData(this._element,Bn),this._element=null,this._config=null},t._getConfig=function(t){return t=l({},ti,g(this._element).data(),"object"==typeof t&&t?t:{}),_.typeCheckConfig(Qn,t,this.constructor.DefaultType),t},t._setListeners=function(){var t=this;g(this._element).on(zn.CLICK_DISMISS,ei,function(){return t.hide(!0)})},t._close=function(){var t=this,e=function(){t._element.classList.add($n),g(t._element).trigger(zn.HIDDEN)};if(this._element.classList.remove(Gn),this._config.animation){var n=_.getTransitionDurationFromElement(this._element);g(this._element).one(_.TRANSITION_END,e).emulateTransitionEnd(n)}else e()},i._jQueryInterface=function(n){return this.each(function(){var t=g(this),e=t.data(Bn);if(e||(e=new i(this,"object"==typeof n&&n),t.data(Bn,e)),"string"==typeof n){if("undefined"==typeof e[n])throw new TypeError('No method named "'+n+'"');e[n](this)}})},s(i,null,[{key:"VERSION",get:function(){return"4.3.1"}},{key:"DefaultType",get:function(){return Zn}},{key:"Default",get:function(){return ti}}]),i}();g.fn[Qn]=ni._jQueryInterface,g.fn[Qn].Constructor=ni,g.fn[Qn].noConflict=function(){return g.fn[Qn]=Yn,ni._jQueryInterface},function(){if("undefined"==typeof g)throw new TypeError("Bootstrap's JavaScript requires jQuery. jQuery must be included before Bootstrap's JavaScript.");var t=g.fn.jquery.split(" ")[0].split(".");if(t[0]<2&&t[1]<9||1===t[0]&&9===t[1]&&t[2]<1||4<=t[0])throw new Error("Bootstrap's JavaScript requires at least jQuery v1.9.1 but less than v4.0.0")}(),t.Util=_,t.Alert=p,t.Button=P,t.Carousel=lt,t.Collapse=bt,t.Dropdown=Jt,t.Modal=ve,t.Popover=sn,t.Scrollspy=Dn,t.Tab=Kn,t.Toast=ni,t.Tooltip=Be,Object.defineProperty(t,"__esModule",{value:!0})});
//# sourceMappingURL=bootstrap.min.js.map
```
