## Supplementary material for "Method for inferring migration routes from phylogenetic trees": Some route-trees: d3.v3.min.js

```
!function(){function n(n){return n&&(n.ownerDocument||n.document||n).documentElement}function t(n){return n&&(n.ownerDocument&&n.ownerDocument.defaultView||n.document&&n||n.defaultView)}function e(n,t){return t>n?-1:n>t?1:n>=t?0:NaN}function r(n){return null===n?NaN:+n}function i(n){return!isNaN(n)}function u(n){return{left:function(t,e,r,i){for(arguments.length<3&&(r=0),arguments.length<4&&(i=t.length);i>r;){var u=r+i>>>1;n(t[u],e)<0?r=u+1:i=u}return r},right:function(t,e,r,i){for(arguments.length<3&&(r=0),arguments.length<4&&(i=t.length);i>r;){var u=r+i>>>1;n(t[u],e)>0?i=u:r=u+1}return r}}}function o(n){return n.length}function a(n){for(var t=1;n*t%1;)t*=10;return t}function l(n,t){for(var e in t)Object.defineProperty(n.prototype,e,{value:t[e],enumerable:!1})}function c(){this._=Object.create(null)}function f(n){return(n+="")===bo||n[0]===_o?_o+n:n}function s(n){return(n+="")[0]===_o?n.slice(1):n}function h(n){return f(n)in this._}function p(n){return(n=f(n))in this._&&delete this._[n]}function g(){var n=[];for(var t in this._)n.push(s(t));return n}function v(){var n=0;for(var t in this._)++n;return n}function d(){for(var n in this._)return!1;return!0}function y(){this._=Object.create(null)}function m(n){return n}function M(n,t,e){return function(){var r=e.apply(t,arguments);return r===t?n:r}}function x(n,t){if(t in n)return t;t=t.charAt(0).toUpperCase()+t.slice(1);for(var e=0,r=wo.length;r>e;++e){var i=wo[e]+t;if(i in n)return i}}function b(){}function _(){}function w(n){function t(){for(var t,r=e,i=-1,u=r.length;++i<u;)(t=r[i].on)&&t.apply(this,arguments);return n}var e=[],r=new c;return t.on=function(t,i){var u,o=r.get(t);return arguments.length<2?o&&o.on:(o&&(o.on=null,e=e.slice(0,u=e.indexOf(o)).concat(e.slice(u+1)),r.remove(t)),i&&e.push(r.set(t,{on:i})),n)},t}function S(){ao.event.preventDefault()}function k(){for(var n,t=ao.event;n=t.sourceEvent;)t=n;return t}function N(n){for(var t=new _,e=0,r=arguments.length;++e<r;)t[arguments[e]]=w(t);return t.of=function(e,r){return function(i){try{var u=i.sourceEvent=ao.event;i.target=n,ao.event=i,t[i.type].apply(e,r)}finally{ao.event=u}}},t}function E(n){return ko(n,Co),n}function A(n){return"function"==typeof n?n:function(){return No(n,this)}}function C(n){return"function"==typeof n?n:function(){return Eo(n,this)}}function z(n,t){function e(){this.removeAttribute(n)}function r(){this.removeAttributeNS(n.space,n.local)}function i(){this.setAttribute(n,t)}function u(){this.setAttributeNS(n.space,n.local,t)}function o(){var e=t.apply(this,arguments);null==e?this.removeAttribute(n):this.setAttribute(n,e)}function a(){var e=t.apply(this,arguments);null==e?this.removeAttributeNS(n.space,n.local):this.setAttributeNS(n.space,n.local,e)}return n=ao.ns.qualify(n),null==t?n.local?r:e:"function"==typeof t?n.local?a:o:n.local?u:i}function L(n){return n.trim().replace(/\s+/g," ")}function q(n){return new RegExp("(?:^|\\s+)"+ao.requote(n)+"(?:\\s+|$)","g")}function T(n){return(n+"").trim().split(/^|\s+/)}function R(n,t){function e(){for(var e=-1;++e<i;)n[e](this,t)}function r(){for(var e=-1,r=t.apply(this,arguments);++e<i;)n[e](this,r)}n=T(n).map(D);var i=n.length;return"function"==typeof t?r:e}function D(n){var t=q(n);return function(e,r){if(i=e.classList)return r?i.add(n):i.remove(n);var i=e.getAttribute("class")||"";r?(t.lastIndex=0,t.test(i)||e.setAttribute("class",L(i+" "+n))):e.setAttribute("class",L(i.replace(t," ")))}}function P(n,t,e){function r(){this.style.removeProperty(n)}function i(){this.style.setProperty(n,t,e)}function u(){var r=t.apply(this,arguments);null==r?this.style.removeProperty(n):this.style.setProperty(n,r,e)}return null==t?r:"function"==typeof t?u:i}function U(n,t){function e(){delete this[n]}function r(){this[n]=t}function i(){var e=t.apply(this,arguments);null==e?delete this[n]:this[n]=e}return null==t?e:"function"==typeof t?i:r}function j(n){function t(){var t=this.ownerDocument,e=this.namespaceURI;return e===zo&&t.documentElement.namespaceURI===zo?t.createElement(n):t.createElementNS(e,n)}function e(){return this.ownerDocument.createElementNS(n.space,n.local)}return"function"==typeof n?n:(n=ao.ns.qualify(n)).local?e:t}function F(){var n=this.parentNode;n&&n.removeChild(this)}function H(n){return{__data__:n}}function O(n){return function(){return Ao(this,n)}}function I(n){return arguments.length||(n=e),function(t,e){return t&&e?n(t.__data__,e.__data__):!t-!e}}function Y(n,t){for(var e=0,r=n.length;r>e;e++)for(var i,u=n[e],o=0,a=u.length;a>o;o++)(i=u[o])&&t(i,o,e);return n}function Z(n){return ko(n,qo),n}function V(n){var t,e;return function(r,i,u){var o,a=n[u].update,l=a.length;for(u!=e&&(e=u,t=0),i>=t&&(t=i+1);!(o=a[t])&&++t<l;);return o}}function X(n,t,e){function r(){var t=this[o];t&&(this.removeEventListener(n,t,t.$),delete this[o])}function i(){var i=l(t,co(arguments));r.call(this),this.addEventListener(n,this[o]=i,i.$=e),i._=t}function u(){var t,e=new RegExp("^__on([^.]+)"+ao.requote(n)+"$");for(var r in this)if(t=r.match(e)){var i=this[r];this.removeEventListener(t[1],i,i.$),delete this[r]}}var o="__on"+n,a=n.indexOf("."),l=$;a>0&&(n=n.slice(0,a));var c=To.get(n);return c&&(n=c,l=B),a?t?i:r:t?b:u}function $(n,t){return function(e){var r=ao.event;ao.event=e,t[0]=this.__data__;try{n.apply(this,t)}finally{ao.event=r}}}function B(n,t){var e=$(n,t);return function(n){var t=this,r=n.relatedTarget;r&&(r===t||8&r.compareDocumentPosition(t))||e.call(t,n)}}function W(e){var r=".dragsuppress-"+ ++Do,i="click"+r,u=ao.select(t(e)).on("touchmove"+r,S).on("dragstart"+r,S).on("selectstart"+r,S);if(null==Ro&&(Ro="onselectstart"in e?!1:x(e.style,"userSelect")),Ro){var o=n(e).style,a=o[Ro];o[Ro]="none"}return function(n){if(u.on(r,null),Ro&&(o[Ro]=a),n){var t=function(){u.on(i,null)};u.on(i,function(){S(),t()},!0),setTimeout(t,0)}}}function J(n,e){e.changedTouches&&(e=e.changedTouches[0]);var r=n.ownerSVGElement||n;if(r.createSVGPoint){var i=r.createSVGPoint();if(0>Po){var u=t(n);if(u.scrollX||u.scrollY){r=ao.select("body").append("svg").style({position:"absolute",top:0,left:0,margin:0,padding:0,border:"none"},"important");var o=r[0][0].getScreenCTM();Po=!(o.f||o.e),r.remove()}}return Po?(i.x=e.pageX,i.y=e.pageY):(i.x=e.clientX,i.y=e.clientY),i=i.matrixTransform(n.getScreenCTM().inverse()),[i.x,i.y]}var a=n.getBoundingClientRect();return[e.clientX-a.left-n.clientLeft,e.clientY-a.top-n.clientTop]}function G(){return ao.event.changedTouches[0].identifier}function K(n){return n>0?1:0>n?-1:0}function Q(n,t,e){return(t[0]-n[0])*(e[1]-n[1])-(t[1]-n[1])*(e[0]-n[0])}function nn(n){return n>1?0:-1>n?Fo:Math.acos(n)}function tn(n){return n>1?Io:-1>n?-Io:Math.asin(n)}function en(n){return((n=Math.exp(n))-1/n)/2}function rn(n){return((n=Math.exp(n))+1/n)/2}function un(n){return((n=Math.exp(2*n))-1)/(n+1)}function on(n){return(n=Math.sin(n/2))*n}function an(){}function ln(n,t,e){return this instanceof ln?(this.h=+n,this.s=+t,void(this.l=+e)):arguments.length<2?n instanceof ln?new ln(n.h,n.s,n.l):_n(""+n,wn,ln):new ln(n,t,e)}function cn(n,t,e){function r(n){return n>360?n-=360:0>n&&(n+=360),60>n?u+(o-u)*n/60:180>n?o:240>n?u+(o-u)*(240-n)/60:u}function i(n){return Math.round(255*r(n))}var u,o;return n=isNaN(n)?0:(n%=360)<0?n+360:n,t=isNaN(t)?0:0>t?0:t>1?1:t,e=0>e?0:e>1?1:e,o=.5>=e?e*(1+t):e+t-e*t,u=2*e-o,new mn(i(n+120),i(n),i(n-120))}function fn(n,t,e){return this instanceof fn?(this.h=+n,this.c=+t,void(this.l=+e)):arguments.length<2?n instanceof fn?new fn(n.h,n.c,n.l):n instanceof hn?gn(n.l,n.a,n.b):gn((n=Sn((n=ao.rgb(n)).r,n.g,n.b)).l,n.a,n.b):new fn(n,t,e)}function sn(n,t,e){return isNaN(n)&&(n=0),isNaN(t)&&(t=0),new hn(e,Math.cos(n*=Yo)*t,Math.sin(n)*t)}function hn(n,t,e){return this instanceof hn?(this.l=+n,this.a=+t,void(this.b=+e)):arguments.length<2?n instanceof hn?new hn(n.l,n.a,n.b):n instanceof fn?sn(n.h,n.c,n.l):Sn((n=mn(n)).r,n.g,n.b):new hn(n,t,e)}function pn(n,t,e){var r=(n+16)/116,i=r+t/500,u=r-e/200;return i=vn(i)*na,r=vn(r)*ta,u=vn(u)*ea,new mn(yn(3.2404542*i-1.5371385*r-.4985314*u),yn(-.969266*i+1.8760108*r+.041556*u),yn(.0556434*i-.2040259*r+1.0572252*u))}function gn(n,t,e){return n>0?new fn(Math.atan2(e,t)*Zo,Math.sqrt(t*t+e*e),n):new fn(NaN,NaN,n)}function vn(n){return n>.206893034?n*n*n:(n-4/29)/7.787037}function dn(n){return n>.008856?Math.pow(n,1/3):7.787037*n+4/29}function yn(n){return Math.round(255*(.00304>=n?12.92*n:1.055*Math.pow(n,1/2.4)-.055))}function mn(n,t,e){return this instanceof mn?(this.r=~~n,this.g=~~t,void(this.b=~~e)):arguments.length<2?n instanceof mn?new mn(n.r,n.g,n.b):_n(""+n,mn,cn):new mn(n,t,e)}function Mn(n){return new mn(n>>16,n>>8&255,255&n)}function xn(n){return Mn(n)+""}function bn(n){return 16>n?"0"+Math.max(0,n).toString(16):Math.min(255,n).toString(16)}function _n(n,t,e){var r,i,u,o=0,a=0,l=0;if(r=/([a-z]+)\((.*)\)/.exec(n=n.toLowerCase()))switch(i=r[2].split(","),r[1]){case"hsl":return e(parseFloat(i[0]),parseFloat(i[1])/100,parseFloat(i[2])/100);case"rgb":return t(Nn(i[0]),Nn(i[1]),Nn(i[2]))}return(u=ua.get(n))?t(u.r,u.g,u.b):(null==n||"#"!==n.charAt(0)||isNaN(u=parseInt(n.slice(1),16))||(4===n.length?(o=(3840&u)>>4,o=o>>4|o,a=240&u,a=a>>4|a,l=15&u,l=l<<4|l):7===n.length&&(o=(16711680&u)>>16,a=(65280&u)>>8,l=255&u)),t(o,a,l))}function wn(n,t,e){var r,i,u=Math.min(n/=255,t/=255,e/=255),o=Math.max(n,t,e),a=o-u,l=(o+u)/2;return a?(i=.5>l?a/(o+u):a/(2-o-u),r=n==o?(t-e)/a+(e>t?6:0):t==o?(e-n)/a+2:(n-t)/a+4,r*=60):(r=NaN,i=l>0&&1>l?0:r),new ln(r,i,l)}function Sn(n,t,e){n=kn(n),t=kn(t),e=kn(e);var r=dn((.4124564*n+.3575761*t+.1804375*e)/na),i=dn((.2126729*n+.7151522*t+.072175*e)/ta),u=dn((.0193339*n+.119192*t+.9503041*e)/ea);return hn(116*i-16,500*(r-i),200*(i-u))}function kn(n){return(n/=255)<=.04045?n/12.92:Math.pow((n+.055)/1.055,2.4)}function Nn(n){var t=parseFloat(n);return"%"===n.charAt(n.length-1)?Math.round(2.55*t):t}function En(n){return"function"==typeof n?n:function(){return n}}function An(n){return function(t,e,r){return 2===arguments.length&&"function"==typeof e&&(r=e,e=null),Cn(t,e,n,r)}}function Cn(n,t,e,r){function i(){var n,t=l.status;if(!t&&Ln(l)||t>=200&&300>t||304===t){try{n=e.call(u,l)}catch(r){return void o.error.call(u,r)}o.load.call(u,n)}else o.error.call(u,l)}var u={},o=ao.dispatch("beforesend","progress","load","error"),a={},l=new XMLHttpRequest,c=null;return!this.XDomainRequest||"withCredentials"in l||!/^(http(s)?:)?\/\//.test(n)||(l=new XDomainRequest),"onload"in l?l.onload=l.onerror=i:l.onreadystatechange=function(){l.readyState>3&&i()},l.onprogress=function(n){var t=ao.event;ao.event=n;try{o.progress.call(u,l)}finally{ao.event=t}},u.header=function(n,t){return n=(n+"").toLowerCase(),arguments.length<2?a[n]:(null==t?delete a[n]:a[n]=t+"",u)},u.mimeType=function(n){return arguments.length?(t=null==n?null:n+"",u):t},u.responseType=function(n){return arguments.length?(c=n,u):c},u.response=function(n){return e=n,u},["get","post"].forEach(function(n){u[n]=function(){return u.send.apply(u,[n].concat(co(arguments)))}}),u.send=function(e,r,i){if(2===arguments.length&&"function"==typeof r&&(i=r,r=null),l.open(e,n,!0),null==t||"accept"in a||(a.accept=t+",*/*"),l.setRequestHeader)for(var f in a)l.setRequestHeader(f,a[f]);return null!=t&&l.overrideMimeType&&l.overrideMimeType(t),null!=c&&(l.responseType=c),null!=i&&u.on("error",i).on("load",function(n){i(null,n)}),o.beforesend.call(u,l),l.send(null==r?null:r),u},u.abort=function(){return l.abort(),u},ao.rebind(u,o,"on"),null==r?u:u.get(zn(r))}function zn(n){return 1===n.length?function(t,e){n(null==t?e:null)}:n}function Ln(n){var t=n.responseType;return t&&"text"!==t?n.response:n.responseText}function qn(n,t,e){var r=arguments.length;2>r&&(t=0),3>r&&(e=Date.now());var i=e+t,u={c:n,t:i,n:null};return aa?aa.n=u:oa=u,aa=u,la||(ca=clearTimeout(ca),la=1,fa(Tn)),u}function Tn(){var n=Rn(),t=Dn()-n;t>24?(isFinite(t)&&(clearTimeout(ca),ca=setTimeout(Tn,t)),la=0):(la=1,fa(Tn))}function Rn(){for(var n=Date.now(),t=oa;t;)n>=t.t&&t.c(n-t.t)&&(t.c=null),t=t.n;return n}function Dn(){for(var n,t=oa,e=1/0;t;)t.c?(t.t<e&&(e=t.t),t=(n=t).n):t=n?n.n=t.n:oa=t.n;return aa=n,e}function Pn(n,t){return t-(n?Math.ceil(Math.log(n)/Math.LN10):1)}function Un(n,t){var e=Math.pow(10,3*xo(8-t));return{scale:t>8?function(n){return n/e}:function(n){return n*e},symbol:n}}function jn(n){var t=n.decimal,e=n.thousands,r=n.grouping,i=n.currency,u=r&&e?function(n,t){for(var i=n.length,u=[],o=0,a=r[0],l=0;i>0&&a>0&&(l+a+1>t&&(a=Math.max(1,t-l)),u.push(n.substring(i-=a,i+a)),!((l+=a+1)>t));)a=r[o=(o+1)%r.length];return u.reverse().join(e)}:m;return function(n){var e=ha.exec(n),r=e[1]||" ",o=e[2]||">",a=e[3]||"-",l=e[4]||"",c=e[5],f=+e[6],s=e[7],h=e[8],p=e[9],g=1,v="",d="",y=!1,m=!0;switch(h&&(h=+h.substring(1)),(c||"0"===r&&"="===o)&&(c=r="0",o="="),p){case"n":s=!0,p="g";break;case"%":g=100,d="%",p="f";break;case"p":g=100,d="%",p="r";break;case"b":case"o":case"x":case"X":"#"===l&&(v="0"+p.toLowerCase());case"c":m=!1;case"d":y=!0,h=0;break;case"s":g=-1,p="r"}"$"===l&&(v=i[0],d=i[1]),"r"!=p||h||(p="g"),null!=h&&("g"==p?h=Math.max(1,Math.min(21,h)):"e"!=p&&"f"!=p||(h=Math.max(0,Math.min(20,h)))),p=pa.get(p)||Fn;var M=c&&s;return function(n){var e=d;if(y&&n%1)return"";var i=0>n||0===n&&0>1/n?(n=-n,"-"):"-"===a?"":a;if(0>g){var l=ao.formatPrefix(n,h);n=l.scale(n),e=l.symbol+d}else n*=g;n=p(n,h);var x,b,_=n.lastIndexOf(".");if(0>_){var w=m?n.lastIndexOf("e"):-1;0>w?(x=n,b=""):(x=n.substring(0,w),b=n.substring(w))}else x=n.substring(0,_),b=t+n.substring(_+1);!c&&s&&(x=u(x,1/0));var S=v.length+x.length+b.length+(M?0:i.length),k=f>S?new Array(S=f-S+1).join(r):"";return M&&(x=u(k+x,k.length?f-b.length:1/0)),i+=v,n=x+b,("<"===o?i+n+k:">"===o?k+i+n:"^"===o?k.substring(0,S>>=1)+i+n+k.substring(S):i+(M?n:k+n))+e}}}function Fn(n){return n+""}function Hn(){this._=new Date(arguments.length>1?Date.UTC.apply(this,arguments):arguments[0])}function On(n,t,e){function r(t){var e=n(t),r=u(e,1);return r-t>t-e?e:r}function i(e){return t(e=n(new va(e-1)),1),e}function u(n,e){return t(n=new va(+n),e),n}function o(n,r,u){var o=i(n),a=[];if(u>1)for(;r>o;)e(o)%u||a.push(new Date(+o)),t(o,1);else for(;r>o;)a.push(new Date(+o)),t(o,1);return a}function a(n,t,e){try{va=Hn;var r=new Hn;return r._=n,o(r,t,e)}finally{va=Date}}n.floor=n,n.round=r,n.ceil=i,n.offset=u,n.range=o;var l=n.utc=In(n);return l.floor=l,l.round=In(r),l.ceil=In(i),l.offset=In(u),l.range=a,n}function In(n){return function(t,e){try{va=Hn;var r=new Hn;return r._=t,n(r,e)._}finally{va=Date}}}function Yn(n){function t(n){function t(t){for(var e,i,u,o=[],a=-1,l=0;++a<r;)37===n.charCodeAt(a)&&(o.push(n.slice(l,a)),null!=(i=ya[e=n.charAt(++a)])&&(e=n.charAt(++a)),(u=A[e])&&(e=u(t,null==i?"e"===e?" ":"0":i)),o.push(e),l=a+1);return o.push(n.slice(l,a)),o.join("")}var r=n.length;return t.parse=function(t){var r={y:1900,m:0,d:1,H:0,M:0,S:0,L:0,Z:null},i=e(r,n,t,0);if(i!=t.length)return null;"p"in r&&(r.H=r.H%12+12*r.p);var u=null!=r.Z&&va!==Hn,o=new(u?Hn:va);return"j"in r?o.setFullYear(r.y,0,r.j):"W"in r||"U"in r?("w"in r||(r.w="W"in r?1:0),o.setFullYear(r.y,0,1),o.setFullYear(r.y,0,"W"in r?(r.w+6)%7+7*r.W-(o.getDay()+5)%7:r.w+7*r.U-(o.getDay()+6)%7)):o.setFullYear(r.y,r.m,r.d),o.setHours(r.H+(r.Z/100|0),r.M+r.Z%100,r.S,r.L),u?o._:o},t.toString=function(){return n},t}function e(n,t,e,r){for(var i,u,o,a=0,l=t.length,c=e.length;l>a;){if(r>=c)return-1;if(i=t.charCodeAt(a++),37===i){if(o=t.charAt(a++),u=C[o in ya?t.charAt(a++):o],!u||(r=u(n,e,r))<0)return-1}else if(i!=e.charCodeAt(r++))return-1}return r}function r(n,t,e){_.lastIndex=0;var r=_.exec(t.slice(e));return r?(n.w=w.get(r[0].toLowerCase()),e+r[0].length):-1}function i(n,t,e){x.lastIndex=0;var r=x.exec(t.slice(e));return r?(n.w=b.get(r[0].toLowerCase()),e+r[0].length):-1}function u(n,t,e){N.lastIndex=0;var r=N.exec(t.slice(e));return r?(n.m=E.get(r[0].toLowerCase()),e+r[0].length):-1}function o(n,t,e){S.lastIndex=0;var r=S.exec(t.slice(e));return r?(n.m=k.get(r[0].toLowerCase()),e+r[0].length):-1}function a(n,t,r){return e(n,A.c.toString(),t,r)}function l(n,t,r){return e(n,A.x.toString(),t,r)}function c(n,t,r){return e(n,A.X.toString(),t,r)}function f(n,t,e){var r=M.get(t.slice(e,e+=2).toLowerCase());return null==r?-1:(n.p=r,e)}var s=n.dateTime,h=n.date,p=n.time,g=n.periods,v=n.days,d=n.shortDays,y=n.months,m=n.shortMonths;t.utc=function(n){function e(n){try{va=Hn;var t=new va;return t._=n,r(t)}finally{va=Date}}var r=t(n);return e.parse=function(n){try{va=Hn;var t=r.parse(n);return t&&t._}finally{va=Date}},e.toString=r.toString,e},t.multi=t.utc.multi=ct;var M=ao.map(),x=Vn(v),b=Xn(v),_=Vn(d),w=Xn(d),S=Vn(y),k=Xn(y),N=Vn(m),E=Xn(m);g.forEach(function(n,t){M.set(n.toLowerCase(),t)});var A={a:function(n){return d[n.getDay()]},A:function(n){return v[n.getDay()]},b:function(n){return m[n.getMonth()]},B:function(n){return y[n.getMonth()]},c:t(s),d:function(n,t){return Zn(n.getDate(),t,2)},e:function(n,t){return Zn(n.getDate(),t,2)},H:function(n,t){return Zn(n.getHours(),t,2)},I:function(n,t){return Zn(n.getHours()%12||12,t,2)},j:function(n,t){return Zn(1+ga.dayOfYear(n),t,3)},L:function(n,t){return Zn(n.getMilliseconds(),t,3)},m:function(n,t){return Zn(n.getMonth()+1,t,2)},M:function(n,t){return Zn(n.getMinutes(),t,2)},p:function(n){return g[+(n.getHours()>=12)]},S:function(n,t){return Zn(n.getSeconds(),t,2)},U:function(n,t){return Zn(ga.sundayOfYear(n),t,2)},w:function(n){return n.getDay()},W:function(n,t){return Zn(ga.mondayOfYear(n),t,2)},x:t(h),X:t(p),y:function(n,t){return Zn(n.getFullYear()%100,t,2)},Y:function(n,t){return Zn(n.getFullYear()%1e4,t,4)},Z:at,"%":function(){return"%"}},C={a:r,A:i,b:u,B:o,c:a,d:tt,e:tt,H:rt,I:rt,j:et,L:ot,m:nt,M:it,p:f,S:ut,U:Bn,w:$n,W:Wn,x:l,X:c,y:Gn,Y:Jn,Z:Kn,"%":lt};return t}function Zn(n,t,e){var r=0>n?"-":"",i=(r?-n:n)+"",u=i.length;return r+(e>u?new Array(e-u+1).join(t)+i:i)}function Vn(n){return new RegExp("^(?:"+n.map(ao.requote).join("|")+")","i")}function Xn(n){for(var t=new c,e=-1,r=n.length;++e<r;)t.set(n[e].toLowerCase(),e);return t}function $n(n,t,e){ma.lastIndex=0;var r=ma.exec(t.slice(e,e+1));return r?(n.w=+r[0],e+r[0].length):-1}function Bn(n,t,e){ma.lastIndex=0;var r=ma.exec(t.slice(e));return r?(n.U=+r[0],e+r[0].length):-1}function Wn(n,t,e){ma.lastIndex=0;var r=ma.exec(t.slice(e));return r?(n.W=+r[0],e+r[0].length):-1}function Jn(n,t,e){ma.lastIndex=0;var r=ma.exec(t.slice(e,e+4));return r?(n.y=+r[0],e+r[0].length):-1}function Gn(n,t,e){ma.lastIndex=0;var r=ma.exec(t.slice(e,e+2));return r?(n.y=Qn(+r[0]),e+r[0].length):-1}function Kn(n,t,e){return/^[+-]\d{4}$/.test(t=t.slice(e,e+5))?(n.Z=-t,e+5):-1}function Qn(n){return n+(n>68?1900:2e3)}function nt(n,t,e){ma.lastIndex=0;var r=ma.exec(t.slice(e,e+2));return r?(n.m=r[0]-1,e+r[0].length):-1}function tt(n,t,e){ma.lastIndex=0;var r=ma.exec(t.slice(e,e+2));return r?(n.d=+r[0],e+r[0].length):-1}function et(n,t,e){ma.lastIndex=0;var r=ma.exec(t.slice(e,e+3));return r?(n.j=+r[0],e+r[0].length):-1}function rt(n,t,e){ma.lastIndex=0;var r=ma.exec(t.slice(e,e+2));return r?(n.H=+r[0],e+r[0].length):-1}function it(n,t,e){ma.lastIndex=0;var r=ma.exec(t.slice(e,e+2));return r?(n.M=+r[0],e+r[0].length):-1}function ut(n,t,e){ma.lastIndex=0;var r=ma.exec(t.slice(e,e+2));return r?(n.S=+r[0],e+r[0].length):-1}function ot(n,t,e){ma.lastIndex=0;var r=ma.exec(t.slice(e,e+3));return r?(n.L=+r[0],e+r[0].length):-1}function at(n){var t=n.getTimezoneOffset(),e=t>0?"-":"+",r=xo(t)/60|0,i=xo(t)%60;return e+Zn(r,"0",2)+Zn(i,"0",2)}function lt(n,t,e){Ma.lastIndex=0;var r=Ma.exec(t.slice(e,e+1));return r?e+r[0].length:-1}function ct(n){for(var t=n.length,e=-1;++e<t;)n[e][0]=this(n[e][0]);return function(t){for(var e=0,r=n[e];!r[1](t);)r=n[++e];return r[0](t)}}function ft(){}function st(n,t,e){var r=e.s=n+t,i=r-n,u=r-i;e.t=n-u+(t-i)}function ht(n,t){n&&wa.hasOwnProperty(n.type)&&wa[n.type](n,t)}function pt(n,t,e){var r,i=-1,u=n.length-e;for(t.lineStart();++i<u;)r=n[i],t.point(r[0],r[1],r[2]);t.lineEnd()}function gt(n,t){var e=-1,r=n.length;for(t.polygonStart();++e<r;)pt(n[e],t,1);t.polygonEnd()}function vt(){function n(n,t){n*=Yo,t=t*Yo/2+Fo/4;var e=n-r,o=e>=0?1:-1,a=o*e,l=Math.cos(t),c=Math.sin(t),f=u*c,s=i*l+f*Math.cos(a),h=f*o*Math.sin(a);ka.add(Math.atan2(h,s)),r=n,i=l,u=c}var t,e,r,i,u;Na.point=function(o,a){Na.point=n,r=(t=o)*Yo,i=Math.cos(a=(e=a)*Yo/2+Fo/4),u=Math.sin(a)},Na.lineEnd=function(){n(t,e)}}function dt(n){var t=n[0],e=n[1],r=Math.cos(e);return[r*Math.cos(t),r*Math.sin(t),Math.sin(e)]}function yt(n,t){return n[0]*t[0]+n[1]*t[1]+n[2]*t[2]}function mt(n,t){return[n[1]*t[2]-n[2]*t[1],n[2]*t[0]-n[0]*t[2],n[0]*t[1]-n[1]*t[0]]}function Mt(n,t){n[0]+=t[0],n[1]+=t[1],n[2]+=t[2]}function xt(n,t){return[n[0]*t,n[1]*t,n[2]*t]}function bt(n){var t=Math.sqrt(n[0]*n[0]+n[1]*n[1]+n[2]*n[2]);n[0]/=t,n[1]/=t,n[2]/=t}function _t(n){return[Math.atan2(n[1],n[0]),tn(n[2])]}function wt(n,t){return xo(n[0]-t[0])<Uo&&xo(n[1]-t[1])<Uo}function St(n,t){n*=Yo;var e=Math.cos(t*=Yo);kt(e*Math.cos(n),e*Math.sin(n),Math.sin(t))}function kt(n,t,e){++Ea,Ca+=(n-Ca)/Ea,za+=(t-za)/Ea,La+=(e-La)/Ea}function Nt(){function n(n,i){n*=Yo;var u=Math.cos(i*=Yo),o=u*Math.cos(n),a=u*Math.sin(n),l=Math.sin(i),c=Math.atan2(Math.sqrt((c=e*l-r*a)*c+(c=r*o-t*l)*c+(c=t*a-e*o)*c),t*o+e*a+r*l);Aa+=c,qa+=c*(t+(t=o)),Ta+=c*(e+(e=a)),Ra+=c*(r+(r=l)),kt(t,e,r)}var t,e,r;ja.point=function(i,u){i*=Yo;var o=Math.cos(u*=Yo);t=o*Math.cos(i),e=o*Math.sin(i),r=Math.sin(u),ja.point=n,kt(t,e,r)}}function Et(){ja.point=St}function At(){function n(n,t){n*=Yo;var e=Math.cos(t*=Yo),o=e*Math.cos(n),a=e*Math.sin(n),l=Math.sin(t),c=i*l-u*a,f=u*o-r*l,s=r*a-i*o,h=Math.sqrt(c*c+f*f+s*s),p=r*o+i*a+u*l,g=h&&-nn(p)/h,v=Math.atan2(h,p);Da+=g*c,Pa+=g*f,Ua+=g*s,Aa+=v,qa+=v*(r+(r=o)),Ta+=v*(i+(i=a)),Ra+=v*(u+(u=l)),kt(r,i,u)}var t,e,r,i,u;ja.point=function(o,a){t=o,e=a,ja.point=n,o*=Yo;var l=Math.cos(a*=Yo);r=l*Math.cos(o),i=l*Math.sin(o),u=Math.sin(a),kt(r,i,u)},ja.lineEnd=function(){n(t,e),ja.lineEnd=Et,ja.point=St}}function Ct(n,t){function e(e,r){return e=n(e,r),t(e[0],e[1])}return n.invert&&t.invert&&(e.invert=function(e,r){return e=t.invert(e,r),e&&n.invert(e[0],e[1])}),e}function zt(){return!0}function Lt(n,t,e,r,i){var u=[],o=[];if(n.forEach(function(n){if(!((t=n.length-1)<=0)){var t,e=n[0],r=n[t];if(wt(e,r)){i.lineStart();for(var a=0;t>a;++a)i.point((e=n[a])[0],e[1]);return void i.lineEnd()}var l=new Tt(e,n,null,!0),c=new Tt(e,null,l,!1);l.o=c,u.push(l),o.push(c),l=new Tt(r,n,null,!1),c=new Tt(r,null,l,!0),l.o=c,u.push(l),o.push(c)}}),o.sort(t),qt(u),qt(o),u.length){for(var a=0,l=e,c=o.length;c>a;++a)o[a].e=l=!l;for(var f,s,h=u[0];;){for(var p=h,g=!0;p.v;)if((p=p.n)===h)return;f=p.z,i.lineStart();do{if(p.v=p.o.v=!0,p.e){if(g)for(var a=0,c=f.length;c>a;++a)i.point((s=f[a])[0],s[1]);else r(p.x,p.n.x,1,i);p=p.n}else{if(g){f=p.p.z;for(var a=f.length-1;a>=0;--a)i.point((s=f[a])[0],s[1])}else r(p.x,p.p.x,-1,i);p=p.p}p=p.o,f=p.z,g=!g}while(!p.v);i.lineEnd()}}}function qt(n){if(t=n.length){for(var t,e,r=0,i=n[0];++r<t;)i.n=e=n[r],e.p=i,i=e;i.n=e=n[0],e.p=i}}function Tt(n,t,e,r){this.x=n,this.z=t,this.o=e,this.e=r,this.v=!1,this.n=this.p=null}function Rt(n,t,e,r){return function(i,u){function o(t,e){var r=i(t,e);n(t=r[0],e=r[1])&&u.point(t,e)}function a(n,t){var e=i(n,t);d.point(e[0],e[1])}function l(){m.point=a,d.lineStart()}function c(){m.point=o,d.lineEnd()}function f(n,t){v.push([n,t]);var e=i(n,t);x.point(e[0],e[1])}function s(){x.lineStart(),v=[]}function h(){f(v[0][0],v[0][1]),x.lineEnd();var n,t=x.clean(),e=M.buffer(),r=e.length;if(v.pop(),g.push(v),v=null,r)if(1&t){n=e[0];var i,r=n.length-1,o=-1;if(r>0){for(b||(u.polygonStart(),b=!0),u.lineStart();++o<r;)u.point((i=n[o])[0],i[1]);u.lineEnd()}}else r>1&&2&t&&e.push(e.pop().concat(e.shift())),p.push(e.filter(Dt))}var p,g,v,d=t(u),y=i.invert(r[0],r[1]),m={point:o,lineStart:l,lineEnd:c,polygonStart:function(){m.point=f,m.lineStart=s,m.lineEnd=h,p=[],g=[]},polygonEnd:function(){m.point=o,m.lineStart=l,m.lineEnd=c,p=ao.merge(p);var n=Ot(y,g);p.length?(b||(u.polygonStart(),b=!0),Lt(p,Ut,n,e,u)):n&&(b||(u.polygonStart(),b=!0),u.lineStart(),e(null,null,1,u),u.lineEnd()),b&&(u.polygonEnd(),b=!1),p=g=null},sphere:function(){u.polygonStart(),u.lineStart(),e(null,null,1,u),u.lineEnd(),u.polygonEnd()}},M=Pt(),x=t(M),b=!1;return m}}function Dt(n){return n.length>1}function Pt(){var n,t=[];return{lineStart:function(){t.push(n=[])},point:function(t,e){n.push([t,e])},lineEnd:b,buffer:function(){var e=t;return t=[],n=null,e},rejoin:function(){t.length>1&&t.push(t.pop().concat(t.shift()))}}}function Ut(n,t){return((n=n.x)[0]<0?n[1]-Io-Uo:Io-n[1])-((t=t.x)[0]<0?t[1]-Io-Uo:Io-t[1])}function jt(n){var t,e=NaN,r=NaN,i=NaN;return{lineStart:function(){n.lineStart(),t=1},point:function(u,o){var a=u>0?Fo:-Fo,l=xo(u-e);xo(l-Fo)<Uo?(n.point(e,r=(r+o)/2>0?Io:-Io),n.point(i,r),n.lineEnd(),n.lineStart(),n.point(a,r),n.point(u,r),t=0):i!==a&&l>=Fo&&(xo(e-i)<Uo&&(e-=i*Uo),xo(u-a)<Uo&&(u-=a*Uo),r=Ft(e,r,u,o),n.point(i,r),n.lineEnd(),n.lineStart(),n.point(a,r),t=0),n.point(e=u,r=o),i=a},lineEnd:function(){n.lineEnd(),e=r=NaN},clean:function(){return 2-t}}}function Ft(n,t,e,r){var i,u,o=Math.sin(n-e);return xo(o)>Uo?Math.atan((Math.sin(t)*(u=Math.cos(r))*Math.sin(e)-Math.sin(r)*(i=Math.cos(t))*Math.sin(n))/(i*u*o)):(t+r)/2}function Ht(n,t,e,r){var i;if(null==n)i=e*Io,r.point(-Fo,i),r.point(0,i),r.point(Fo,i),r.point(Fo,0),r.point(Fo,-i),r.point(0,-i),r.point(-Fo,-i),r.point(-Fo,0),r.point(-Fo,i);else if(xo(n[0]-t[0])>Uo){var u=n[0]<t[0]?Fo:-Fo;i=e*u/2,r.point(-u,i),r.point(0,i),r.point(u,i)}else r.point(t[0],t[1])}function Ot(n,t){var e=n[0],r=n[1],i=[Math.sin(e),-Math.cos(e),0],u=0,o=0;ka.reset();for(var a=0,l=t.length;l>a;++a){var c=t[a],f=c.length;if(f)for(var s=c[0],h=s[0],p=s[1]/2+Fo/4,g=Math.sin(p),v=Math.cos(p),d=1;;){d===f&&(d=0),n=c[d];var y=n[0],m=n[1]/2+Fo/4,M=Math.sin(m),x=Math.cos(m),b=y-h,_=b>=0?1:-1,w=_*b,S=w>Fo,k=g*M;if(ka.add(Math.atan2(k*_*Math.sin(w),v*x+k*Math.cos(w))),u+=S?b+_*Ho:b,S^h>=e^y>=e){var N=mt(dt(s),dt(n));bt(N);var E=mt(i,N);bt(E);var A=(S^b>=0?-1:1)*tn(E[2]);(r>A||r===A&&(N[0]||N[1]))&&(o+=S^b>=0?1:-1)}if(!d++)break;h=y,g=M,v=x,s=n}}return(-Uo>u||Uo>u&&-Uo>ka)^1&o}function It(n){function t(n,t){return Math.cos(n)*Math.cos(t)>u}function e(n){var e,u,l,c,f;return{lineStart:function(){c=l=!1,f=1},point:function(s,h){var p,g=[s,h],v=t(s,h),d=o?v?0:i(s,h):v?i(s+(0>s?Fo:-Fo),h):0;if(!e&&(c=l=v)&&n.lineStart(),v!==l&&(p=r(e,g),(wt(e,p)||wt(g,p))&&(g[0]+=Uo,g[1]+=Uo,v=t(g[0],g[1]))),v!==l)f=0,v?(n.lineStart(),p=r(g,e),n.point(p[0],p[1])):(p=r(e,g),n.point(p[0],p[1]),n.lineEnd()),e=p;else if(a&&e&&o^v){var y;d&u||!(y=r(g,e,!0))||(f=0,o?(n.lineStart(),n.point(y[0][0],y[0][1]),n.point(y[1][0],y[1][1]),n.lineEnd()):(n.point(y[1][0],y[1][1]),n.lineEnd(),n.lineStart(),n.point(y[0][0],y[0][1])))}!v||e&&wt(e,g)||n.point(g[0],g[1]),e=g,l=v,u=d},lineEnd:function(){l&&n.lineEnd(),e=null},clean:function(){return f|(c&&l)<<1}}}function r(n,t,e){var r=dt(n),i=dt(t),o=[1,0,0],a=mt(r,i),l=yt(a,a),c=a[0],f=l-c*c;if(!f)return!e&&n;var s=u*l/f,h=-u*c/f,p=mt(o,a),g=xt(o,s),v=xt(a,h);Mt(g,v);var d=p,y=yt(g,d),m=yt(d,d),M=y*y-m*(yt(g,g)-1);if(!(0>M)){var x=Math.sqrt(M),b=xt(d,(-y-x)/m);if(Mt(b,g),b=_t(b),!e)return b;var _,w=n[0],S=t[0],k=n[1],N=t[1];w>S&&(_=w,w=S,S=_);var E=S-w,A=xo(E-Fo)<Uo,C=A||Uo>E;if(!A&&k>N&&(_=k,k=N,N=_),C?A?k+N>0^b[1]<(xo(b[0]-w)<Uo?k:N):k<=b[1]&&b[1]<=N:E>Fo^(w<=b[0]&&b[0]<=S)){var z=xt(d,(-y+x)/m);return Mt(z,g),[b,_t(z)]}}}function i(t,e){var r=o?n:Fo-n,i=0;return-r>t?i|=1:t>r&&(i|=2),-r>e?i|=4:e>r&&(i|=8),i}var u=Math.cos(n),o=u>0,a=xo(u)>Uo,l=ve(n,6*Yo);return Rt(t,e,l,o?[0,-n]:[-Fo,n-Fo])}function Yt(n,t,e,r){return function(i){var u,o=i.a,a=i.b,l=o.x,c=o.y,f=a.x,s=a.y,h=0,p=1,g=f-l,v=s-c;if(u=n-l,g||!(u>0)){if(u/=g,0>g){if(h>u)return;p>u&&(p=u)}else if(g>0){if(u>p)return;u>h&&(h=u)}if(u=e-l,g||!(0>u)){if(u/=g,0>g){if(u>p)return;u>h&&(h=u)}else if(g>0){if(h>u)return;p>u&&(p=u)}if(u=t-c,v||!(u>0)){if(u/=v,0>v){if(h>u)return;p>u&&(p=u)}else if(v>0){if(u>p)return;u>h&&(h=u)}if(u=r-c,v||!(0>u)){if(u/=v,0>v){if(u>p)return;u>h&&(h=u)}else if(v>0){if(h>u)return;p>u&&(p=u)}return h>0&&(i.a={x:l+h*g,y:c+h*v}),1>p&&(i.b={x:l+p*g,y:c+p*v}),i}}}}}}function Zt(n,t,e,r){function i(r,i){return xo(r[0]-n)<Uo?i>0?0:3:xo(r[0]-e)<Uo?i>0?2:1:xo(r[1]-t)<Uo?i>0?1:0:i>0?3:2}function u(n,t){return o(n.x,t.x)}function o(n,t){var e=i(n,1),r=i(t,1);return e!==r?e-r:0===e?t[1]-n[1]:1===e?n[0]-t[0]:2===e?n[1]-t[1]:t[0]-n[0]}return function(a){function l(n){for(var t=0,e=d.length,r=n[1],i=0;e>i;++i)for(var u,o=1,a=d[i],l=a.length,c=a[0];l>o;++o)u=a[o],c[1]<=r?u[1]>r&&Q(c,u,n)>0&&++t:u[1]<=r&&Q(c,u,n)<0&&--t,c=u;return 0!==t}function c(u,a,l,c){var f=0,s=0;if(null==u||(f=i(u,l))!==(s=i(a,l))||o(u,a)<0^l>0){do c.point(0===f||3===f?n:e,f>1?r:t);while((f=(f+l+4)%4)!==s)}else c.point(a[0],a[1])}function f(i,u){return i>=n&&e>=i&&u>=t&&r>=u}function s(n,t){f(n,t)&&a.point(n,t)}function h(){C.point=g,d&&d.push(y=[]),S=!0,w=!1,b=_=NaN}function p(){v&&(g(m,M),x&&w&&E.rejoin(),v.push(E.buffer())),C.point=s,w&&a.lineEnd()}function g(n,t){n=Math.max(-Ha,Math.min(Ha,n)),t=Math.max(-Ha,Math.min(Ha,t));var e=f(n,t);if(d&&y.push([n,t]),S)m=n,M=t,x=e,S=!1,e&&(a.lineStart(),a.point(n,t));else if(e&&w)a.point(n,t);else{var r={a:{x:b,y:_},b:{x:n,y:t}};A(r)?(w||(a.lineStart(),a.point(r.a.x,r.a.y)),a.point(r.b.x,r.b.y),e||a.lineEnd(),k=!1):e&&(a.lineStart(),a.point(n,t),k=!1)}b=n,_=t,w=e}var v,d,y,m,M,x,b,_,w,S,k,N=a,E=Pt(),A=Yt(n,t,e,r),C={point:s,lineStart:h,lineEnd:p,polygonStart:function(){a=E,v=[],d=[],k=!0},polygonEnd:function(){a=N,v=ao.merge(v);var t=l([n,r]),e=k&&t,i=v.length;(e||i)&&(a.polygonStart(),e&&(a.lineStart(),c(null,null,1,a),a.lineEnd()),i&&Lt(v,u,t,c,a),a.polygonEnd()),v=d=y=null}};return C}}function Vt(n){var t=0,e=Fo/3,r=ae(n),i=r(t,e);return i.parallels=function(n){return arguments.length?r(t=n[0]*Fo/180,e=n[1]*Fo/180):[t/Fo*180,e/Fo*180]},i}function Xt(n,t){function e(n,t){var e=Math.sqrt(u-2*i*Math.sin(t))/i;return[e*Math.sin(n*=i),o-e*Math.cos(n)]}var r=Math.sin(n),i=(r+Math.sin(t))/2,u=1+r*(2*i-r),o=Math.sqrt(u)/i;return e.invert=function(n,t){var e=o-t;return[Math.atan2(n,e)/i,tn((u-(n*n+e*e)*i*i)/(2*i))]},e}function $t(){function n(n,t){Ia+=i*n-r*t,r=n,i=t}var t,e,r,i;$a.point=function(u,o){$a.point=n,t=r=u,e=i=o},$a.lineEnd=function(){n(t,e)}}function Bt(n,t){Ya>n&&(Ya=n),n>Va&&(Va=n),Za>t&&(Za=t),t>Xa&&(Xa=t)}function Wt(){function n(n,t){o.push("M",n,",",t,u)}function t(n,t){o.push("M",n,",",t),a.point=e}function e(n,t){o.push("L",n,",",t)}function r(){a.point=n}function i(){o.push("Z")}var u=Jt(4.5),o=[],a={point:n,lineStart:function(){a.point=t},lineEnd:r,polygonStart:function(){a.lineEnd=i},polygonEnd:function(){a.lineEnd=r,a.point=n},pointRadius:function(n){return u=Jt(n),a},result:function(){if(o.length){var n=o.join("");return o=[],n}}};return a}function Jt(n){return"m0,"+n+"a"+n+","+n+" 0 1,1 0,"+-2*n+"a"+n+","+n+" 0 1,1 0,"+2*n+"z"}function Gt(n,t){Ca+=n,za+=t,++La}function Kt(){function n(n,r){var i=n-t,u=r-e,o=Math.sqrt(i*i+u*u);qa+=o*(t+n)/2,Ta+=o*(e+r)/2,Ra+=o,Gt(t=n,e=r)}var t,e;Wa.point=function(r,i){Wa.point=n,Gt(t=r,e=i)}}function Qt(){Wa.point=Gt}function ne(){function n(n,t){var e=n-r,u=t-i,o=Math.sqrt(e*e+u*u);qa+=o*(r+n)/2,Ta+=o*(i+t)/2,Ra+=o,o=i*n-r*t,Da+=o*(r+n),Pa+=o*(i+t),Ua+=3*o,Gt(r=n,i=t)}var t,e,r,i;Wa.point=function(u,o){Wa.point=n,Gt(t=r=u,e=i=o)},Wa.lineEnd=function(){n(t,e)}}function te(n){function t(t,e){n.moveTo(t+o,e),n.arc(t,e,o,0,Ho)}function e(t,e){n.moveTo(t,e),a.point=r}function r(t,e){n.lineTo(t,e)}function i(){a.point=t}function u(){n.closePath()}var o=4.5,a={point:t,lineStart:function(){a.point=e},lineEnd:i,polygonStart:function(){a.lineEnd=u},polygonEnd:function(){a.lineEnd=i,a.point=t},pointRadius:function(n){return o=n,a},result:b};return a}function ee(n){function t(n){return(a?r:e)(n)}function e(t){return ue(t,function(e,r){e=n(e,r),t.point(e[0],e[1])})}function r(t){function e(e,r){e=n(e,r),t.point(e[0],e[1])}function r(){M=NaN,S.point=u,t.lineStart()}function u(e,r){var u=dt([e,r]),o=n(e,r);i(M,x,m,b,_,w,M=o[0],x=o[1],m=e,b=u[0],_=u[1],w=u[2],a,t),t.point(M,x)}function o(){S.point=e,t.lineEnd()}function l(){
r(),S.point=c,S.lineEnd=f}function c(n,t){u(s=n,h=t),p=M,g=x,v=b,d=_,y=w,S.point=u}function f(){i(M,x,m,b,_,w,p,g,s,v,d,y,a,t),S.lineEnd=o,o()}var s,h,p,g,v,d,y,m,M,x,b,_,w,S={point:e,lineStart:r,lineEnd:o,polygonStart:function(){t.polygonStart(),S.lineStart=l},polygonEnd:function(){t.polygonEnd(),S.lineStart=r}};return S}function i(t,e,r,a,l,c,f,s,h,p,g,v,d,y){var m=f-t,M=s-e,x=m*m+M*M;if(x>4*u&&d--){var b=a+p,_=l+g,w=c+v,S=Math.sqrt(b*b+_*_+w*w),k=Math.asin(w/=S),N=xo(xo(w)-1)<Uo||xo(r-h)<Uo?(r+h)/2:Math.atan2(_,b),E=n(N,k),A=E[0],C=E[1],z=A-t,L=C-e,q=M*z-m*L;(q*q/x>u||xo((m*z+M*L)/x-.5)>.3||o>a*p+l*g+c*v)&&(i(t,e,r,a,l,c,A,C,N,b/=S,_/=S,w,d,y),y.point(A,C),i(A,C,N,b,_,w,f,s,h,p,g,v,d,y))}}var u=.5,o=Math.cos(30*Yo),a=16;return t.precision=function(n){return arguments.length?(a=(u=n*n)>0&&16,t):Math.sqrt(u)},t}function re(n){var t=ee(function(t,e){return n([t*Zo,e*Zo])});return function(n){return le(t(n))}}function ie(n){this.stream=n}function ue(n,t){return{point:t,sphere:function(){n.sphere()},lineStart:function(){n.lineStart()},lineEnd:function(){n.lineEnd()},polygonStart:function(){n.polygonStart()},polygonEnd:function(){n.polygonEnd()}}}function oe(n){return ae(function(){return n})()}function ae(n){function t(n){return n=a(n[0]*Yo,n[1]*Yo),[n[0]*h+l,c-n[1]*h]}function e(n){return n=a.invert((n[0]-l)/h,(c-n[1])/h),n&&[n[0]*Zo,n[1]*Zo]}function r(){a=Ct(o=se(y,M,x),u);var n=u(v,d);return l=p-n[0]*h,c=g+n[1]*h,i()}function i(){return f&&(f.valid=!1,f=null),t}var u,o,a,l,c,f,s=ee(function(n,t){return n=u(n,t),[n[0]*h+l,c-n[1]*h]}),h=150,p=480,g=250,v=0,d=0,y=0,M=0,x=0,b=Fa,_=m,w=null,S=null;return t.stream=function(n){return f&&(f.valid=!1),f=le(b(o,s(_(n)))),f.valid=!0,f},t.clipAngle=function(n){return arguments.length?(b=null==n?(w=n,Fa):It((w=+n)*Yo),i()):w},t.clipExtent=function(n){return arguments.length?(S=n,_=n?Zt(n[0][0],n[0][1],n[1][0],n[1][1]):m,i()):S},t.scale=function(n){return arguments.length?(h=+n,r()):h},t.translate=function(n){return arguments.length?(p=+n[0],g=+n[1],r()):[p,g]},t.center=function(n){return arguments.length?(v=n[0]%360*Yo,d=n[1]%360*Yo,r()):[v*Zo,d*Zo]},t.rotate=function(n){return arguments.length?(y=n[0]%360*Yo,M=n[1]%360*Yo,x=n.length>2?n[2]%360*Yo:0,r()):[y*Zo,M*Zo,x*Zo]},ao.rebind(t,s,"precision"),function(){return u=n.apply(this,arguments),t.invert=u.invert&&e,r()}}function le(n){return ue(n,function(t,e){n.point(t*Yo,e*Yo)})}function ce(n,t){return[n,t]}function fe(n,t){return[n>Fo?n-Ho:-Fo>n?n+Ho:n,t]}function se(n,t,e){return n?t||e?Ct(pe(n),ge(t,e)):pe(n):t||e?ge(t,e):fe}function he(n){return function(t,e){return t+=n,[t>Fo?t-Ho:-Fo>t?t+Ho:t,e]}}function pe(n){var t=he(n);return t.invert=he(-n),t}function ge(n,t){function e(n,t){var e=Math.cos(t),a=Math.cos(n)*e,l=Math.sin(n)*e,c=Math.sin(t),f=c*r+a*i;return[Math.atan2(l*u-f*o,a*r-c*i),tn(f*u+l*o)]}var r=Math.cos(n),i=Math.sin(n),u=Math.cos(t),o=Math.sin(t);return e.invert=function(n,t){var e=Math.cos(t),a=Math.cos(n)*e,l=Math.sin(n)*e,c=Math.sin(t),f=c*u-l*o;return[Math.atan2(l*u+c*o,a*r+f*i),tn(f*r-a*i)]},e}function ve(n,t){var e=Math.cos(n),r=Math.sin(n);return function(i,u,o,a){var l=o*t;null!=i?(i=de(e,i),u=de(e,u),(o>0?u>i:i>u)&&(i+=o*Ho)):(i=n+o*Ho,u=n-.5*l);for(var c,f=i;o>0?f>u:u>f;f-=l)a.point((c=_t([e,-r*Math.cos(f),-r*Math.sin(f)]))[0],c[1])}}function de(n,t){var e=dt(t);e[0]-=n,bt(e);var r=nn(-e[1]);return((-e[2]<0?-r:r)+2*Math.PI-Uo)%(2*Math.PI)}function ye(n,t,e){var r=ao.range(n,t-Uo,e).concat(t);return function(n){return r.map(function(t){return[n,t]})}}function me(n,t,e){var r=ao.range(n,t-Uo,e).concat(t);return function(n){return r.map(function(t){return[t,n]})}}function Me(n){return n.source}function xe(n){return n.target}function be(n,t,e,r){var i=Math.cos(t),u=Math.sin(t),o=Math.cos(r),a=Math.sin(r),l=i*Math.cos(n),c=i*Math.sin(n),f=o*Math.cos(e),s=o*Math.sin(e),h=2*Math.asin(Math.sqrt(on(r-t)+i*o*on(e-n))),p=1/Math.sin(h),g=h?function(n){var t=Math.sin(n*=h)*p,e=Math.sin(h-n)*p,r=e*l+t*f,i=e*c+t*s,o=e*u+t*a;return[Math.atan2(i,r)*Zo,Math.atan2(o,Math.sqrt(r*r+i*i))*Zo]}:function(){return[n*Zo,t*Zo]};return g.distance=h,g}function _e(){function n(n,i){var u=Math.sin(i*=Yo),o=Math.cos(i),a=xo((n*=Yo)-t),l=Math.cos(a);Ja+=Math.atan2(Math.sqrt((a=o*Math.sin(a))*a+(a=r*u-e*o*l)*a),e*u+r*o*l),t=n,e=u,r=o}var t,e,r;Ga.point=function(i,u){t=i*Yo,e=Math.sin(u*=Yo),r=Math.cos(u),Ga.point=n},Ga.lineEnd=function(){Ga.point=Ga.lineEnd=b}}function we(n,t){function e(t,e){var r=Math.cos(t),i=Math.cos(e),u=n(r*i);return[u*i*Math.sin(t),u*Math.sin(e)]}return e.invert=function(n,e){var r=Math.sqrt(n*n+e*e),i=t(r),u=Math.sin(i),o=Math.cos(i);return[Math.atan2(n*u,r*o),Math.asin(r&&e*u/r)]},e}function Se(n,t){function e(n,t){o>0?-Io+Uo>t&&(t=-Io+Uo):t>Io-Uo&&(t=Io-Uo);var e=o/Math.pow(i(t),u);return[e*Math.sin(u*n),o-e*Math.cos(u*n)]}var r=Math.cos(n),i=function(n){return Math.tan(Fo/4+n/2)},u=n===t?Math.sin(n):Math.log(r/Math.cos(t))/Math.log(i(t)/i(n)),o=r*Math.pow(i(n),u)/u;return u?(e.invert=function(n,t){var e=o-t,r=K(u)*Math.sqrt(n*n+e*e);return[Math.atan2(n,e)/u,2*Math.atan(Math.pow(o/r,1/u))-Io]},e):Ne}function ke(n,t){function e(n,t){var e=u-t;return[e*Math.sin(i*n),u-e*Math.cos(i*n)]}var r=Math.cos(n),i=n===t?Math.sin(n):(r-Math.cos(t))/(t-n),u=r/i+n;return xo(i)<Uo?ce:(e.invert=function(n,t){var e=u-t;return[Math.atan2(n,e)/i,u-K(i)*Math.sqrt(n*n+e*e)]},e)}function Ne(n,t){return[n,Math.log(Math.tan(Fo/4+t/2))]}function Ee(n){var t,e=oe(n),r=e.scale,i=e.translate,u=e.clipExtent;return e.scale=function(){var n=r.apply(e,arguments);return n===e?t?e.clipExtent(null):e:n},e.translate=function(){var n=i.apply(e,arguments);return n===e?t?e.clipExtent(null):e:n},e.clipExtent=function(n){var o=u.apply(e,arguments);if(o===e){if(t=null==n){var a=Fo*r(),l=i();u([[l[0]-a,l[1]-a],[l[0]+a,l[1]+a]])}}else t&&(o=null);return o},e.clipExtent(null)}function Ae(n,t){return[Math.log(Math.tan(Fo/4+t/2)),-n]}function Ce(n){return n[0]}function ze(n){return n[1]}function Le(n){for(var t=n.length,e=[0,1],r=2,i=2;t>i;i++){for(;r>1&&Q(n[e[r-2]],n[e[r-1]],n[i])<=0;)--r;e[r++]=i}return e.slice(0,r)}function qe(n,t){return n[0]-t[0]||n[1]-t[1]}function Te(n,t,e){return(e[0]-t[0])*(n[1]-t[1])<(e[1]-t[1])*(n[0]-t[0])}function Re(n,t,e,r){var i=n[0],u=e[0],o=t[0]-i,a=r[0]-u,l=n[1],c=e[1],f=t[1]-l,s=r[1]-c,h=(a*(l-c)-s*(i-u))/(s*o-a*f);return[i+h*o,l+h*f]}function De(n){var t=n[0],e=n[n.length-1];return!(t[0]-e[0]||t[1]-e[1])}function Pe(){rr(this),this.edge=this.site=this.circle=null}function Ue(n){var t=cl.pop()||new Pe;return t.site=n,t}function je(n){Be(n),ol.remove(n),cl.push(n),rr(n)}function Fe(n){var t=n.circle,e=t.x,r=t.cy,i={x:e,y:r},u=n.P,o=n.N,a=[n];je(n);for(var l=u;l.circle&&xo(e-l.circle.x)<Uo&&xo(r-l.circle.cy)<Uo;)u=l.P,a.unshift(l),je(l),l=u;a.unshift(l),Be(l);for(var c=o;c.circle&&xo(e-c.circle.x)<Uo&&xo(r-c.circle.cy)<Uo;)o=c.N,a.push(c),je(c),c=o;a.push(c),Be(c);var f,s=a.length;for(f=1;s>f;++f)c=a[f],l=a[f-1],nr(c.edge,l.site,c.site,i);l=a[0],c=a[s-1],c.edge=Ke(l.site,c.site,null,i),$e(l),$e(c)}function He(n){for(var t,e,r,i,u=n.x,o=n.y,a=ol._;a;)if(r=Oe(a,o)-u,r>Uo)a=a.L;else{if(i=u-Ie(a,o),!(i>Uo)){r>-Uo?(t=a.P,e=a):i>-Uo?(t=a,e=a.N):t=e=a;break}if(!a.R){t=a;break}a=a.R}var l=Ue(n);if(ol.insert(t,l),t||e){if(t===e)return Be(t),e=Ue(t.site),ol.insert(l,e),l.edge=e.edge=Ke(t.site,l.site),$e(t),void $e(e);if(!e)return void(l.edge=Ke(t.site,l.site));Be(t),Be(e);var c=t.site,f=c.x,s=c.y,h=n.x-f,p=n.y-s,g=e.site,v=g.x-f,d=g.y-s,y=2*(h*d-p*v),m=h*h+p*p,M=v*v+d*d,x={x:(d*m-p*M)/y+f,y:(h*M-v*m)/y+s};nr(e.edge,c,g,x),l.edge=Ke(c,n,null,x),e.edge=Ke(n,g,null,x),$e(t),$e(e)}}function Oe(n,t){var e=n.site,r=e.x,i=e.y,u=i-t;if(!u)return r;var o=n.P;if(!o)return-(1/0);e=o.site;var a=e.x,l=e.y,c=l-t;if(!c)return a;var f=a-r,s=1/u-1/c,h=f/c;return s?(-h+Math.sqrt(h*h-2*s*(f*f/(-2*c)-l+c/2+i-u/2)))/s+r:(r+a)/2}function Ie(n,t){var e=n.N;if(e)return Oe(e,t);var r=n.site;return r.y===t?r.x:1/0}function Ye(n){this.site=n,this.edges=[]}function Ze(n){for(var t,e,r,i,u,o,a,l,c,f,s=n[0][0],h=n[1][0],p=n[0][1],g=n[1][1],v=ul,d=v.length;d--;)if(u=v[d],u&&u.prepare())for(a=u.edges,l=a.length,o=0;l>o;)f=a[o].end(),r=f.x,i=f.y,c=a[++o%l].start(),t=c.x,e=c.y,(xo(r-t)>Uo||xo(i-e)>Uo)&&(a.splice(o,0,new tr(Qe(u.site,f,xo(r-s)<Uo&&g-i>Uo?{x:s,y:xo(t-s)<Uo?e:g}:xo(i-g)<Uo&&h-r>Uo?{x:xo(e-g)<Uo?t:h,y:g}:xo(r-h)<Uo&&i-p>Uo?{x:h,y:xo(t-h)<Uo?e:p}:xo(i-p)<Uo&&r-s>Uo?{x:xo(e-p)<Uo?t:s,y:p}:null),u.site,null)),++l)}function Ve(n,t){return t.angle-n.angle}function Xe(){rr(this),this.x=this.y=this.arc=this.site=this.cy=null}function $e(n){var t=n.P,e=n.N;if(t&&e){var r=t.site,i=n.site,u=e.site;if(r!==u){var o=i.x,a=i.y,l=r.x-o,c=r.y-a,f=u.x-o,s=u.y-a,h=2*(l*s-c*f);if(!(h>=-jo)){var p=l*l+c*c,g=f*f+s*s,v=(s*p-c*g)/h,d=(l*g-f*p)/h,s=d+a,y=fl.pop()||new Xe;y.arc=n,y.site=i,y.x=v+o,y.y=s+Math.sqrt(v*v+d*d),y.cy=s,n.circle=y;for(var m=null,M=ll._;M;)if(y.y<M.y||y.y===M.y&&y.x<=M.x){if(!M.L){m=M.P;break}M=M.L}else{if(!M.R){m=M;break}M=M.R}ll.insert(m,y),m||(al=y)}}}}function Be(n){var t=n.circle;t&&(t.P||(al=t.N),ll.remove(t),fl.push(t),rr(t),n.circle=null)}function We(n){for(var t,e=il,r=Yt(n[0][0],n[0][1],n[1][0],n[1][1]),i=e.length;i--;)t=e[i],(!Je(t,n)||!r(t)||xo(t.a.x-t.b.x)<Uo&&xo(t.a.y-t.b.y)<Uo)&&(t.a=t.b=null,e.splice(i,1))}function Je(n,t){var e=n.b;if(e)return!0;var r,i,u=n.a,o=t[0][0],a=t[1][0],l=t[0][1],c=t[1][1],f=n.l,s=n.r,h=f.x,p=f.y,g=s.x,v=s.y,d=(h+g)/2,y=(p+v)/2;if(v===p){if(o>d||d>=a)return;if(h>g){if(u){if(u.y>=c)return}else u={x:d,y:l};e={x:d,y:c}}else{if(u){if(u.y<l)return}else u={x:d,y:c};e={x:d,y:l}}}else if(r=(h-g)/(v-p),i=y-r*d,-1>r||r>1)if(h>g){if(u){if(u.y>=c)return}else u={x:(l-i)/r,y:l};e={x:(c-i)/r,y:c}}else{if(u){if(u.y<l)return}else u={x:(c-i)/r,y:c};e={x:(l-i)/r,y:l}}else if(v>p){if(u){if(u.x>=a)return}else u={x:o,y:r*o+i};e={x:a,y:r*a+i}}else{if(u){if(u.x<o)return}else u={x:a,y:r*a+i};e={x:o,y:r*o+i}}return n.a=u,n.b=e,!0}function Ge(n,t){this.l=n,this.r=t,this.a=this.b=null}function Ke(n,t,e,r){var i=new Ge(n,t);return il.push(i),e&&nr(i,n,t,e),r&&nr(i,t,n,r),ul[n.i].edges.push(new tr(i,n,t)),ul[t.i].edges.push(new tr(i,t,n)),i}function Qe(n,t,e){var r=new Ge(n,null);return r.a=t,r.b=e,il.push(r),r}function nr(n,t,e,r){n.a||n.b?n.l===e?n.b=r:n.a=r:(n.a=r,n.l=t,n.r=e)}function tr(n,t,e){var r=n.a,i=n.b;this.edge=n,this.site=t,this.angle=e?Math.atan2(e.y-t.y,e.x-t.x):n.l===t?Math.atan2(i.x-r.x,r.y-i.y):Math.atan2(r.x-i.x,i.y-r.y)}function er(){this._=null}function rr(n){n.U=n.C=n.L=n.R=n.P=n.N=null}function ir(n,t){var e=t,r=t.R,i=e.U;i?i.L===e?i.L=r:i.R=r:n._=r,r.U=i,e.U=r,e.R=r.L,e.R&&(e.R.U=e),r.L=e}function ur(n,t){var e=t,r=t.L,i=e.U;i?i.L===e?i.L=r:i.R=r:n._=r,r.U=i,e.U=r,e.L=r.R,e.L&&(e.L.U=e),r.R=e}function or(n){for(;n.L;)n=n.L;return n}function ar(n,t){var e,r,i,u=n.sort(lr).pop();for(il=[],ul=new Array(n.length),ol=new er,ll=new er;;)if(i=al,u&&(!i||u.y<i.y||u.y===i.y&&u.x<i.x))u.x===e&&u.y===r||(ul[u.i]=new Ye(u),He(u),e=u.x,r=u.y),u=n.pop();else{if(!i)break;Fe(i.arc)}t&&(We(t),Ze(t));var o={cells:ul,edges:il};return ol=ll=il=ul=null,o}function lr(n,t){return t.y-n.y||t.x-n.x}function cr(n,t,e){return(n.x-e.x)*(t.y-n.y)-(n.x-t.x)*(e.y-n.y)}function fr(n){return n.x}function sr(n){return n.y}function hr(){return{leaf:!0,nodes:[],point:null,x:null,y:null}}function pr(n,t,e,r,i,u){if(!n(t,e,r,i,u)){var o=.5*(e+i),a=.5*(r+u),l=t.nodes;l[0]&&pr(n,l[0],e,r,o,a),l[1]&&pr(n,l[1],o,r,i,a),l[2]&&pr(n,l[2],e,a,o,u),l[3]&&pr(n,l[3],o,a,i,u)}}function gr(n,t,e,r,i,u,o){var a,l=1/0;return function c(n,f,s,h,p){if(!(f>u||s>o||r>h||i>p)){if(g=n.point){var g,v=t-n.x,d=e-n.y,y=v*v+d*d;if(l>y){var m=Math.sqrt(l=y);r=t-m,i=e-m,u=t+m,o=e+m,a=g}}for(var M=n.nodes,x=.5*(f+h),b=.5*(s+p),_=t>=x,w=e>=b,S=w<<1|_,k=S+4;k>S;++S)if(n=M[3&S])switch(3&S){case 0:c(n,f,s,x,b);break;case 1:c(n,x,s,h,b);break;case 2:c(n,f,b,x,p);break;case 3:c(n,x,b,h,p)}}}(n,r,i,u,o),a}function vr(n,t){n=ao.rgb(n),t=ao.rgb(t);var e=n.r,r=n.g,i=n.b,u=t.r-e,o=t.g-r,a=t.b-i;return function(n){return"#"+bn(Math.round(e+u*n))+bn(Math.round(r+o*n))+bn(Math.round(i+a*n))}}function dr(n,t){var e,r={},i={};for(e in n)e in t?r[e]=Mr(n[e],t[e]):i[e]=n[e];for(e in t)e in n||(i[e]=t[e]);return function(n){for(e in r)i[e]=r[e](n);return i}}function yr(n,t){return n=+n,t=+t,function(e){return n*(1-e)+t*e}}function mr(n,t){var e,r,i,u=hl.lastIndex=pl.lastIndex=0,o=-1,a=[],l=[];for(n+="",t+="";(e=hl.exec(n))&&(r=pl.exec(t));)(i=r.index)>u&&(i=t.slice(u,i),a[o]?a[o]+=i:a[++o]=i),(e=e[0])===(r=r[0])?a[o]?a[o]+=r:a[++o]=r:(a[++o]=null,l.push({i:o,x:yr(e,r)})),u=pl.lastIndex;return u<t.length&&(i=t.slice(u),a[o]?a[o]+=i:a[++o]=i),a.length<2?l[0]?(t=l[0].x,function(n){return t(n)+""}):function(){return t}:(t=l.length,function(n){for(var e,r=0;t>r;++r)a[(e=l[r]).i]=e.x(n);return a.join("")})}function Mr(n,t){for(var e,r=ao.interpolators.length;--r>=0&&!(e=ao.interpolators[r](n,t)););return e}function xr(n,t){var e,r=[],i=[],u=n.length,o=t.length,a=Math.min(n.length,t.length);for(e=0;a>e;++e)r.push(Mr(n[e],t[e]));for(;u>e;++e)i[e]=n[e];for(;o>e;++e)i[e]=t[e];return function(n){for(e=0;a>e;++e)i[e]=r[e](n);return i}}function br(n){return function(t){return 0>=t?0:t>=1?1:n(t)}}function _r(n){return function(t){return 1-n(1-t)}}function wr(n){return function(t){return.5*(.5>t?n(2*t):2-n(2-2*t))}}function Sr(n){return n*n}function kr(n){return n*n*n}function Nr(n){if(0>=n)return 0;if(n>=1)return 1;var t=n*n,e=t*n;return 4*(.5>n?e:3*(n-t)+e-.75)}function Er(n){return function(t){return Math.pow(t,n)}}function Ar(n){return 1-Math.cos(n*Io)}function Cr(n){return Math.pow(2,10*(n-1))}function zr(n){return 1-Math.sqrt(1-n*n)}function Lr(n,t){var e;return arguments.length<2&&(t=.45),arguments.length?e=t/Ho*Math.asin(1/n):(n=1,e=t/4),function(r){return 1+n*Math.pow(2,-10*r)*Math.sin((r-e)*Ho/t)}}function qr(n){return n||(n=1.70158),function(t){return t*t*((n+1)*t-n)}}function Tr(n){return 1/2.75>n?7.5625*n*n:2/2.75>n?7.5625*(n-=1.5/2.75)*n+.75:2.5/2.75>n?7.5625*(n-=2.25/2.75)*n+.9375:7.5625*(n-=2.625/2.75)*n+.984375}function Rr(n,t){n=ao.hcl(n),t=ao.hcl(t);var e=n.h,r=n.c,i=n.l,u=t.h-e,o=t.c-r,a=t.l-i;return isNaN(o)&&(o=0,r=isNaN(r)?t.c:r),isNaN(u)?(u=0,e=isNaN(e)?t.h:e):u>180?u-=360:-180>u&&(u+=360),function(n){return sn(e+u*n,r+o*n,i+a*n)+""}}function Dr(n,t){n=ao.hsl(n),t=ao.hsl(t);var e=n.h,r=n.s,i=n.l,u=t.h-e,o=t.s-r,a=t.l-i;return isNaN(o)&&(o=0,r=isNaN(r)?t.s:r),isNaN(u)?(u=0,e=isNaN(e)?t.h:e):u>180?u-=360:-180>u&&(u+=360),function(n){return cn(e+u*n,r+o*n,i+a*n)+""}}function Pr(n,t){n=ao.lab(n),t=ao.lab(t);var e=n.l,r=n.a,i=n.b,u=t.l-e,o=t.a-r,a=t.b-i;return function(n){return pn(e+u*n,r+o*n,i+a*n)+""}}function Ur(n,t){return t-=n,function(e){return Math.round(n+t*e)}}function jr(n){var t=[n.a,n.b],e=[n.c,n.d],r=Hr(t),i=Fr(t,e),u=Hr(Or(e,t,-i))||0;t[0]*e[1]<e[0]*t[1]&&(t[0]*=-1,t[1]*=-1,r*=-1,i*=-1),this.rotate=(r?Math.atan2(t[1],t[0]):Math.atan2(-e[0],e[1]))*Zo,this.translate=[n.e,n.f],this.scale=[r,u],this.skew=u?Math.atan2(i,u)*Zo:0}function Fr(n,t){return n[0]*t[0]+n[1]*t[1]}function Hr(n){var t=Math.sqrt(Fr(n,n));return t&&(n[0]/=t,n[1]/=t),t}function Or(n,t,e){return n[0]+=e*t[0],n[1]+=e*t[1],n}function Ir(n){return n.length?n.pop()+",":""}function Yr(n,t,e,r){if(n[0]!==t[0]||n[1]!==t[1]){var i=e.push("translate(",null,",",null,")");r.push({i:i-4,x:yr(n[0],t[0])},{i:i-2,x:yr(n[1],t[1])})}else(t[0]||t[1])&&e.push("translate("+t+")")}function Zr(n,t,e,r){n!==t?(n-t>180?t+=360:t-n>180&&(n+=360),r.push({i:e.push(Ir(e)+"rotate(",null,")")-2,x:yr(n,t)})):t&&e.push(Ir(e)+"rotate("+t+")")}function Vr(n,t,e,r){n!==t?r.push({i:e.push(Ir(e)+"skewX(",null,")")-2,x:yr(n,t)}):t&&e.push(Ir(e)+"skewX("+t+")")}function Xr(n,t,e,r){if(n[0]!==t[0]||n[1]!==t[1]){var i=e.push(Ir(e)+"scale(",null,",",null,")");r.push({i:i-4,x:yr(n[0],t[0])},{i:i-2,x:yr(n[1],t[1])})}else 1===t[0]&&1===t[1]||e.push(Ir(e)+"scale("+t+")")}function $r(n,t){var e=[],r=[];return n=ao.transform(n),t=ao.transform(t),Yr(n.translate,t.translate,e,r),Zr(n.rotate,t.rotate,e,r),Vr(n.skew,t.skew,e,r),Xr(n.scale,t.scale,e,r),n=t=null,function(n){for(var t,i=-1,u=r.length;++i<u;)e[(t=r[i]).i]=t.x(n);return e.join("")}}function Br(n,t){return t=(t-=n=+n)||1/t,function(e){return(e-n)/t}}function Wr(n,t){return t=(t-=n=+n)||1/t,function(e){return Math.max(0,Math.min(1,(e-n)/t))}}function Jr(n){for(var t=n.source,e=n.target,r=Kr(t,e),i=[t];t!==r;)t=t.parent,i.push(t);for(var u=i.length;e!==r;)i.splice(u,0,e),e=e.parent;return i}function Gr(n){for(var t=[],e=n.parent;null!=e;)t.push(n),n=e,e=e.parent;return t.push(n),t}function Kr(n,t){if(n===t)return n;for(var e=Gr(n),r=Gr(t),i=e.pop(),u=r.pop(),o=null;i===u;)o=i,i=e.pop(),u=r.pop();return o}function Qr(n){n.fixed|=2}function ni(n){n.fixed&=-7}function ti(n){n.fixed|=4,n.px=n.x,n.py=n.y}function ei(n){n.fixed&=-5}function ri(n,t,e){var r=0,i=0;if(n.charge=0,!n.leaf)for(var u,o=n.nodes,a=o.length,l=-1;++l<a;)u=o[l],null!=u&&(ri(u,t,e),n.charge+=u.charge,r+=u.charge*u.cx,i+=u.charge*u.cy);if(n.point){n.leaf||(n.point.x+=Math.random()-.5,n.point.y+=Math.random()-.5);var c=t*e[n.point.index];n.charge+=n.pointCharge=c,r+=c*n.point.x,i+=c*n.point.y}n.cx=r/n.charge,n.cy=i/n.charge}function ii(n,t){return ao.rebind(n,t,"sort","children","value"),n.nodes=n,n.links=fi,n}function ui(n,t){for(var e=[n];null!=(n=e.pop());)if(t(n),(i=n.children)&&(r=i.length))for(var r,i;--r>=0;)e.push(i[r])}function oi(n,t){for(var e=[n],r=[];null!=(n=e.pop());)if(r.push(n),(u=n.children)&&(i=u.length))for(var i,u,o=-1;++o<i;)e.push(u[o]);for(;null!=(n=r.pop());)t(n)}function ai(n){return n.children}function li(n){return n.value}function ci(n,t){return t.value-n.value}function fi(n){return ao.merge(n.map(function(n){return(n.children||[]).map(function(t){return{source:n,target:t}})}))}function si(n){return n.x}function hi(n){return n.y}function pi(n,t,e){n.y0=t,n.y=e}function gi(n){return ao.range(n.length)}function vi(n){for(var t=-1,e=n[0].length,r=[];++t<e;)r[t]=0;return r}function di(n){for(var t,e=1,r=0,i=n[0][1],u=n.length;u>e;++e)(t=n[e][1])>i&&(r=e,i=t);return r}function yi(n){return n.reduce(mi,0)}function mi(n,t){return n+t[1]}function Mi(n,t){return xi(n,Math.ceil(Math.log(t.length)/Math.LN2+1))}function xi(n,t){for(var e=-1,r=+n[0],i=(n[1]-r)/t,u=[];++e<=t;)u[e]=i*e+r;return u}function bi(n){return[ao.min(n),ao.max(n)]}function _i(n,t){return n.value-t.value}function wi(n,t){var e=n._pack_next;n._pack_next=t,t._pack_prev=n,t._pack_next=e,e._pack_prev=t}function Si(n,t){n._pack_next=t,t._pack_prev=n}function ki(n,t){var e=t.x-n.x,r=t.y-n.y,i=n.r+t.r;return.999*i*i>e*e+r*r}function Ni(n){function t(n){f=Math.min(n.x-n.r,f),s=Math.max(n.x+n.r,s),h=Math.min(n.y-n.r,h),p=Math.max(n.y+n.r,p)}if((e=n.children)&&(c=e.length)){var e,r,i,u,o,a,l,c,f=1/0,s=-(1/0),h=1/0,p=-(1/0);if(e.forEach(Ei),r=e[0],r.x=-r.r,r.y=0,t(r),c>1&&(i=e[1],i.x=i.r,i.y=0,t(i),c>2))for(u=e[2],zi(r,i,u),t(u),wi(r,u),r._pack_prev=u,wi(u,i),i=r._pack_next,o=3;c>o;o++){zi(r,i,u=e[o]);var g=0,v=1,d=1;for(a=i._pack_next;a!==i;a=a._pack_next,v++)if(ki(a,u)){g=1;break}if(1==g)for(l=r._pack_prev;l!==a._pack_prev&&!ki(l,u);l=l._pack_prev,d++);g?(d>v||v==d&&i.r<r.r?Si(r,i=a):Si(r=l,i),o--):(wi(r,u),i=u,t(u))}var y=(f+s)/2,m=(h+p)/2,M=0;for(o=0;c>o;o++)u=e[o],u.x-=y,u.y-=m,M=Math.max(M,u.r+Math.sqrt(u.x*u.x+u.y*u.y));n.r=M,e.forEach(Ai)}}function Ei(n){n._pack_next=n._pack_prev=n}function Ai(n){delete n._pack_next,delete n._pack_prev}function Ci(n,t,e,r){var i=n.children;if(n.x=t+=r*n.x,n.y=e+=r*n.y,n.r*=r,i)for(var u=-1,o=i.length;++u<o;)Ci(i[u],t,e,r)}function zi(n,t,e){var r=n.r+e.r,i=t.x-n.x,u=t.y-n.y;if(r&&(i||u)){var o=t.r+e.r,a=i*i+u*u;o*=o,r*=r;var l=.5+(r-o)/(2*a),c=Math.sqrt(Math.max(0,2*o*(r+a)-(r-=a)*r-o*o))/(2*a);e.x=n.x+l*i+c*u,e.y=n.y+l*u-c*i}else e.x=n.x+r,e.y=n.y}function Li(n,t){return n.parent==t.parent?1:2}function qi(n){var t=n.children;return t.length?t[0]:n.t}function Ti(n){var t,e=n.children;return(t=e.length)?e[t-1]:n.t}function Ri(n,t,e){var r=e/(t.i-n.i);t.c-=r,t.s+=e,n.c+=r,t.z+=e,t.m+=e}function Di(n){for(var t,e=0,r=0,i=n.children,u=i.length;--u>=0;)t=i[u],t.z+=e,t.m+=e,e+=t.s+(r+=t.c)}function Pi(n,t,e){return n.a.parent===t.parent?n.a:e}function Ui(n){return 1+ao.max(n,function(n){return n.y})}function ji(n){return n.reduce(function(n,t){return n+t.x},0)/n.length}function Fi(n){var t=n.children;return t&&t.length?Fi(t[0]):n}function Hi(n){var t,e=n.children;return e&&(t=e.length)?Hi(e[t-1]):n}function Oi(n){return{x:n.x,y:n.y,dx:n.dx,dy:n.dy}}function Ii(n,t){var e=n.x+t[3],r=n.y+t[0],i=n.dx-t[1]-t[3],u=n.dy-t[0]-t[2];return 0>i&&(e+=i/2,i=0),0>u&&(r+=u/2,u=0),{x:e,y:r,dx:i,dy:u}}function Yi(n){var t=n[0],e=n[n.length-1];return e>t?[t,e]:[e,t]}function Zi(n){return n.rangeExtent?n.rangeExtent():Yi(n.range())}function Vi(n,t,e,r){var i=e(n[0],n[1]),u=r(t[0],t[1]);return function(n){return u(i(n))}}function Xi(n,t){var e,r=0,i=n.length-1,u=n[r],o=n[i];return u>o&&(e=r,r=i,i=e,e=u,u=o,o=e),n[r]=t.floor(u),n[i]=t.ceil(o),n}function $i(n){return n?{floor:function(t){return Math.floor(t/n)*n},ceil:function(t){return Math.ceil(t/n)*n}}:Sl}function Bi(n,t,e,r){var i=[],u=[],o=0,a=Math.min(n.length,t.length)-1;for(n[a]<n[0]&&(n=n.slice().reverse(),t=t.slice().reverse());++o<=a;)i.push(e(n[o-1],n[o])),u.push(r(t[o-1],t[o]));return function(t){var e=ao.bisect(n,t,1,a)-1;return u[e](i[e](t))}}function Wi(n,t,e,r){function i(){var i=Math.min(n.length,t.length)>2?Bi:Vi,l=r?Wr:Br;return o=i(n,t,l,e),a=i(t,n,l,Mr),u}function u(n){return o(n)}var o,a;return u.invert=function(n){return a(n)},u.domain=function(t){return arguments.length?(n=t.map(Number),i()):n},u.range=function(n){return arguments.length?(t=n,i()):t},u.rangeRound=function(n){return u.range(n).interpolate(Ur)},u.clamp=function(n){return arguments.length?(r=n,i()):r},u.interpolate=function(n){return arguments.length?(e=n,i()):e},u.ticks=function(t){return Qi(n,t)},u.tickFormat=function(t,e){return nu(n,t,e)},u.nice=function(t){return Gi(n,t),i()},u.copy=function(){return Wi(n,t,e,r)},i()}function Ji(n,t){return ao.rebind(n,t,"range","rangeRound","interpolate","clamp")}function Gi(n,t){return Xi(n,$i(Ki(n,t)[2])),Xi(n,$i(Ki(n,t)[2])),n}function Ki(n,t){null==t&&(t=10);var e=Yi(n),r=e[1]-e[0],i=Math.pow(10,Math.floor(Math.log(r/t)/Math.LN10)),u=t/r*i;return.15>=u?i*=10:.35>=u?i*=5:.75>=u&&(i*=2),e[0]=Math.ceil(e[0]/i)*i,e[1]=Math.floor(e[1]/i)*i+.5*i,e[2]=i,e}function Qi(n,t){return ao.range.apply(ao,Ki(n,t))}function nu(n,t,e){var r=Ki(n,t);if(e){var i=ha.exec(e);if(i.shift(),"s"===i[8]){var u=ao.formatPrefix(Math.max(xo(r[0]),xo(r[1])));return i[7]||(i[7]="."+tu(u.scale(r[2]))),i[8]="f",e=ao.format(i.join("")),function(n){return e(u.scale(n))+u.symbol}}i[7]||(i[7]="."+eu(i[8],r)),e=i.join("")}else e=",."+tu(r[2])+"f";return ao.format(e)}function tu(n){return-Math.floor(Math.log(n)/Math.LN10+.01)}function eu(n,t){var e=tu(t[2]);return n in kl?Math.abs(e-tu(Math.max(xo(t[0]),xo(t[1]))))+ +("e"!==n):e-2*("%"===n)}function ru(n,t,e,r){function i(n){return(e?Math.log(0>n?0:n):-Math.log(n>0?0:-n))/Math.log(t)}function u(n){return e?Math.pow(t,n):-Math.pow(t,-n)}function o(t){return n(i(t))}return o.invert=function(t){return u(n.invert(t))},o.domain=function(t){return arguments.length?(e=t[0]>=0,n.domain((r=t.map(Number)).map(i)),o):r},o.base=function(e){return arguments.length?(t=+e,n.domain(r.map(i)),o):t},o.nice=function(){var t=Xi(r.map(i),e?Math:El);return n.domain(t),r=t.map(u),o},o.ticks=function(){var n=Yi(r),o=[],a=n[0],l=n[1],c=Math.floor(i(a)),f=Math.ceil(i(l)),s=t%1?2:t;if(isFinite(f-c)){if(e){for(;f>c;c++)for(var h=1;s>h;h++)o.push(u(c)*h);o.push(u(c))}else for(o.push(u(c));c++<f;)for(var h=s-1;h>0;h--)o.push(u(c)*h);for(c=0;o[c]<a;c++);for(f=o.length;o[f-1]>l;f--);o=o.slice(c,f)}return o},o.tickFormat=function(n,e){if(!arguments.length)return Nl;arguments.length<2?e=Nl:"function"!=typeof e&&(e=ao.format(e));var r=Math.max(1,t*n/o.ticks().length);return function(n){var o=n/u(Math.round(i(n)));return t-.5>o*t&&(o*=t),r>=o?e(n):""}},o.copy=function(){return ru(n.copy(),t,e,r)},Ji(o,n)}function iu(n,t,e){function r(t){return n(i(t))}var i=uu(t),u=uu(1/t);return r.invert=function(t){return u(n.invert(t))},r.domain=function(t){return arguments.length?(n.domain((e=t.map(Number)).map(i)),r):e},r.ticks=function(n){return Qi(e,n)},r.tickFormat=function(n,t){return nu(e,n,t)},r.nice=function(n){return r.domain(Gi(e,n))},r.exponent=function(o){return arguments.length?(i=uu(t=o),u=uu(1/t),n.domain(e.map(i)),r):t},r.copy=function(){return iu(n.copy(),t,e)},Ji(r,n)}function uu(n){return function(t){return 0>t?-Math.pow(-t,n):Math.pow(t,n)}}function ou(n,t){function e(e){return u[((i.get(e)||("range"===t.t?i.set(e,n.push(e)):NaN))-1)%u.length]}function r(t,e){return ao.range(n.length).map(function(n){return t+e*n})}var i,u,o;return e.domain=function(r){if(!arguments.length)return n;n=[],i=new c;for(var u,o=-1,a=r.length;++o<a;)i.has(u=r[o])||i.set(u,n.push(u));return e[t.t].apply(e,t.a)},e.range=function(n){return arguments.length?(u=n,o=0,t={t:"range",a:arguments},e):u},e.rangePoints=function(i,a){arguments.length<2&&(a=0);var l=i[0],c=i[1],f=n.length<2?(l=(l+c)/2,0):(c-l)/(n.length-1+a);return u=r(l+f*a/2,f),o=0,t={t:"rangePoints",a:arguments},e},e.rangeRoundPoints=function(i,a){arguments.length<2&&(a=0);var l=i[0],c=i[1],f=n.length<2?(l=c=Math.round((l+c)/2),0):(c-l)/(n.length-1+a)|0;return u=r(l+Math.round(f*a/2+(c-l-(n.length-1+a)*f)/2),f),o=0,t={t:"rangeRoundPoints",a:arguments},e},e.rangeBands=function(i,a,l){arguments.length<2&&(a=0),arguments.length<3&&(l=a);var c=i[1]<i[0],f=i[c-0],s=i[1-c],h=(s-f)/(n.length-a+2*l);return u=r(f+h*l,h),c&&u.reverse(),o=h*(1-a),t={t:"rangeBands",a:arguments},e},e.rangeRoundBands=function(i,a,l){arguments.length<2&&(a=0),arguments.length<3&&(l=a);var c=i[1]<i[0],f=i[c-0],s=i[1-c],h=Math.floor((s-f)/(n.length-a+2*l));return u=r(f+Math.round((s-f-(n.length-a)*h)/2),h),c&&u.reverse(),o=Math.round(h*(1-a)),t={t:"rangeRoundBands",a:arguments},e},e.rangeBand=function(){return o},e.rangeExtent=function(){return Yi(t.a[0])},e.copy=function(){return ou(n,t)},e.domain(n)}function au(n,t){function u(){var e=0,r=t.length;for(a=[];++e<r;)a[e-1]=ao.quantile(n,e/r);return o}function o(n){return isNaN(n=+n)?void 0:t[ao.bisect(a,n)]}var a;return o.domain=function(t){return arguments.length?(n=t.map(r).filter(i).sort(e),u()):n},o.range=function(n){return arguments.length?(t=n,u()):t},o.quantiles=function(){return a},o.invertExtent=function(e){return e=t.indexOf(e),0>e?[NaN,NaN]:[e>0?a[e-1]:n[0],e<a.length?a[e]:n[n.length-1]]},o.copy=function(){return au(n,t)},u()}function lu(n,t,e){function r(t){return e[Math.max(0,Math.min(o,Math.floor(u*(t-n))))]}function i(){return u=e.length/(t-n),o=e.length-1,r}var u,o;return r.domain=function(e){return arguments.length?(n=+e[0],t=+e[e.length-1],i()):[n,t]},r.range=function(n){return arguments.length?(e=n,i()):e},r.invertExtent=function(t){return t=e.indexOf(t),t=0>t?NaN:t/u+n,[t,t+1/u]},r.copy=function(){return lu(n,t,e)},i()}function cu(n,t){function e(e){return e>=e?t[ao.bisect(n,e)]:void 0}return e.domain=function(t){return arguments.length?(n=t,e):n},e.range=function(n){return arguments.length?(t=n,e):t},e.invertExtent=function(e){return e=t.indexOf(e),[n[e-1],n[e]]},e.copy=function(){return cu(n,t)},e}function fu(n){function t(n){return+n}return t.invert=t,t.domain=t.range=function(e){return arguments.length?(n=e.map(t),t):n},t.ticks=function(t){return Qi(n,t)},t.tickFormat=function(t,e){return nu(n,t,e)},t.copy=function(){return fu(n)},t}function su(){return 0}function hu(n){return n.innerRadius}function pu(n){return n.outerRadius}function gu(n){return n.startAngle}function vu(n){return n.endAngle}function du(n){return n&&n.padAngle}function yu(n,t,e,r){return(n-e)*t-(t-r)*n>0?0:1}function mu(n,t,e,r,i){var u=n[0]-t[0],o=n[1]-t[1],a=(i?r:-r)/Math.sqrt(u*u+o*o),l=a*o,c=-a*u,f=n[0]+l,s=n[1]+c,h=t[0]+l,p=t[1]+c,g=(f+h)/2,v=(s+p)/2,d=h-f,y=p-s,m=d*d+y*y,M=e-r,x=f*p-h*s,b=(0>y?-1:1)*Math.sqrt(Math.max(0,M*M*m-x*x)),_=(x*y-d*b)/m,w=(-x*d-y*b)/m,S=(x*y+d*b)/m,k=(-x*d+y*b)/m,N=_-g,E=w-v,A=S-g,C=k-v;return N*N+E*E>A*A+C*C&&(_=S,w=k),[[_-l,w-c],[_*e/M,w*e/M]]}function Mu(n){function t(t){function o(){c.push("M",u(n(f),a))}for(var l,c=[],f=[],s=-1,h=t.length,p=En(e),g=En(r);++s<h;)i.call(this,l=t[s],s)?f.push([+p.call(this,l,s),+g.call(this,l,s)]):f.length&&(o(),f=[]);return f.length&&o(),c.length?c.join(""):null}var e=Ce,r=ze,i=zt,u=xu,o=u.key,a=.7;return t.x=function(n){return arguments.length?(e=n,t):e},t.y=function(n){return arguments.length?(r=n,t):r},t.defined=function(n){return arguments.length?(i=n,t):i},t.interpolate=function(n){return arguments.length?(o="function"==typeof n?u=n:(u=Tl.get(n)||xu).key,t):o},t.tension=function(n){return arguments.length?(a=n,t):a},t}function xu(n){return n.length>1?n.join("L"):n+"Z"}function bu(n){return n.join("L")+"Z"}function _u(n){for(var t=0,e=n.length,r=n[0],i=[r[0],",",r[1]];++t<e;)i.push("H",(r[0]+(r=n[t])[0])/2,"V",r[1]);return e>1&&i.push("H",r[0]),i.join("")}function wu(n){for(var t=0,e=n.length,r=n[0],i=[r[0],",",r[1]];++t<e;)i.push("V",(r=n[t])[1],"H",r[0]);return i.join("")}function Su(n){for(var t=0,e=n.length,r=n[0],i=[r[0],",",r[1]];++t<e;)i.push("H",(r=n[t])[0],"V",r[1]);return i.join("")}function ku(n,t){return n.length<4?xu(n):n[1]+Au(n.slice(1,-1),Cu(n,t))}function Nu(n,t){return n.length<3?bu(n):n[0]+Au((n.push(n[0]),n),Cu([n[n.length-2]].concat(n,[n[1]]),t))}function Eu(n,t){return n.length<3?xu(n):n[0]+Au(n,Cu(n,t))}function Au(n,t){if(t.length<1||n.length!=t.length&&n.length!=t.length+2)return xu(n);var e=n.length!=t.length,r="",i=n[0],u=n[1],o=t[0],a=o,l=1;if(e&&(r+="Q"+(u[0]-2*o[0]/3)+","+(u[1]-2*o[1]/3)+","+u[0]+","+u[1],i=n[1],l=2),t.length>1){a=t[1],u=n[l],l++,r+="C"+(i[0]+o[0])+","+(i[1]+o[1])+","+(u[0]-a[0])+","+(u[1]-a[1])+","+u[0]+","+u[1];for(var c=2;c<t.length;c++,l++)u=n[l],a=t[c],r+="S"+(u[0]-a[0])+","+(u[1]-a[1])+","+u[0]+","+u[1]}if(e){var f=n[l];r+="Q"+(u[0]+2*a[0]/3)+","+(u[1]+2*a[1]/3)+","+f[0]+","+f[1]}return r}function Cu(n,t){for(var e,r=[],i=(1-t)/2,u=n[0],o=n[1],a=1,l=n.length;++a<l;)e=u,u=o,o=n[a],r.push([i*(o[0]-e[0]),i*(o[1]-e[1])]);return r}function zu(n){if(n.length<3)return xu(n);var t=1,e=n.length,r=n[0],i=r[0],u=r[1],o=[i,i,i,(r=n[1])[0]],a=[u,u,u,r[1]],l=[i,",",u,"L",Ru(Pl,o),",",Ru(Pl,a)];for(n.push(n[e-1]);++t<=e;)r=n[t],o.shift(),o.push(r[0]),a.shift(),a.push(r[1]),Du(l,o,a);return n.pop(),l.push("L",r),l.join("")}function Lu(n){if(n.length<4)return xu(n);for(var t,e=[],r=-1,i=n.length,u=[0],o=[0];++r<3;)t=n[r],u.push(t[0]),o.push(t[1]);for(e.push(Ru(Pl,u)+","+Ru(Pl,o)),--r;++r<i;)t=n[r],u.shift(),u.push(t[0]),o.shift(),o.push(t[1]),Du(e,u,o);return e.join("")}function qu(n){for(var t,e,r=-1,i=n.length,u=i+4,o=[],a=[];++r<4;)e=n[r%i],o.push(e[0]),a.push(e[1]);for(t=[Ru(Pl,o),",",Ru(Pl,a)],--r;++r<u;)e=n[r%i],o.shift(),o.push(e[0]),a.shift(),a.push(e[1]),Du(t,o,a);return t.join("")}function Tu(n,t){var e=n.length-1;if(e)for(var r,i,u=n[0][0],o=n[0][1],a=n[e][0]-u,l=n[e][1]-o,c=-1;++c<=e;)r=n[c],i=c/e,r[0]=t*r[0]+(1-t)*(u+i*a),r[1]=t*r[1]+(1-t)*(o+i*l);return zu(n)}function Ru(n,t){return n[0]*t[0]+n[1]*t[1]+n[2]*t[2]+n[3]*t[3]}function Du(n,t,e){n.push("C",Ru(Rl,t),",",Ru(Rl,e),",",Ru(Dl,t),",",Ru(Dl,e),",",Ru(Pl,t),",",Ru(Pl,e))}function Pu(n,t){return(t[1]-n[1])/(t[0]-n[0])}function Uu(n){for(var t=0,e=n.length-1,r=[],i=n[0],u=n[1],o=r[0]=Pu(i,u);++t<e;)r[t]=(o+(o=Pu(i=u,u=n[t+1])))/2;return r[t]=o,r}function ju(n){for(var t,e,r,i,u=[],o=Uu(n),a=-1,l=n.length-1;++a<l;)t=Pu(n[a],n[a+1]),xo(t)<Uo?o[a]=o[a+1]=0:(e=o[a]/t,r=o[a+1]/t,i=e*e+r*r,i>9&&(i=3*t/Math.sqrt(i),o[a]=i*e,o[a+1]=i*r));for(a=-1;++a<=l;)i=(n[Math.min(l,a+1)][0]-n[Math.max(0,a-1)][0])/(6*(1+o[a]*o[a])),u.push([i||0,o[a]*i||0]);return u}function Fu(n){return n.length<3?xu(n):n[0]+Au(n,ju(n))}function Hu(n){for(var t,e,r,i=-1,u=n.length;++i<u;)t=n[i],e=t[0],r=t[1]-Io,t[0]=e*Math.cos(r),t[1]=e*Math.sin(r);return n}function Ou(n){function t(t){function l(){v.push("M",a(n(y),s),f,c(n(d.reverse()),s),"Z")}for(var h,p,g,v=[],d=[],y=[],m=-1,M=t.length,x=En(e),b=En(i),_=e===r?function(){
return p}:En(r),w=i===u?function(){return g}:En(u);++m<M;)o.call(this,h=t[m],m)?(d.push([p=+x.call(this,h,m),g=+b.call(this,h,m)]),y.push([+_.call(this,h,m),+w.call(this,h,m)])):d.length&&(l(),d=[],y=[]);return d.length&&l(),v.length?v.join(""):null}var e=Ce,r=Ce,i=0,u=ze,o=zt,a=xu,l=a.key,c=a,f="L",s=.7;return t.x=function(n){return arguments.length?(e=r=n,t):r},t.x0=function(n){return arguments.length?(e=n,t):e},t.x1=function(n){return arguments.length?(r=n,t):r},t.y=function(n){return arguments.length?(i=u=n,t):u},t.y0=function(n){return arguments.length?(i=n,t):i},t.y1=function(n){return arguments.length?(u=n,t):u},t.defined=function(n){return arguments.length?(o=n,t):o},t.interpolate=function(n){return arguments.length?(l="function"==typeof n?a=n:(a=Tl.get(n)||xu).key,c=a.reverse||a,f=a.closed?"M":"L",t):l},t.tension=function(n){return arguments.length?(s=n,t):s},t}function Iu(n){return n.radius}function Yu(n){return[n.x,n.y]}function Zu(n){return function(){var t=n.apply(this,arguments),e=t[0],r=t[1]-Io;return[e*Math.cos(r),e*Math.sin(r)]}}function Vu(){return 64}function Xu(){return"circle"}function $u(n){var t=Math.sqrt(n/Fo);return"M0,"+t+"A"+t+","+t+" 0 1,1 0,"+-t+"A"+t+","+t+" 0 1,1 0,"+t+"Z"}function Bu(n){return function(){var t,e,r;(t=this[n])&&(r=t[e=t.active])&&(r.timer.c=null,r.timer.t=NaN,--t.count?delete t[e]:delete this[n],t.active+=.5,r.event&&r.event.interrupt.call(this,this.__data__,r.index))}}function Wu(n,t,e){return ko(n,Yl),n.namespace=t,n.id=e,n}function Ju(n,t,e,r){var i=n.id,u=n.namespace;return Y(n,"function"==typeof e?function(n,o,a){n[u][i].tween.set(t,r(e.call(n,n.__data__,o,a)))}:(e=r(e),function(n){n[u][i].tween.set(t,e)}))}function Gu(n){return null==n&&(n=""),function(){this.textContent=n}}function Ku(n){return null==n?"__transition__":"__transition_"+n+"__"}function Qu(n,t,e,r,i){function u(n){var t=v.delay;return f.t=t+l,n>=t?o(n-t):void(f.c=o)}function o(e){var i=g.active,u=g[i];u&&(u.timer.c=null,u.timer.t=NaN,--g.count,delete g[i],u.event&&u.event.interrupt.call(n,n.__data__,u.index));for(var o in g)if(r>+o){var c=g[o];c.timer.c=null,c.timer.t=NaN,--g.count,delete g[o]}f.c=a,qn(function(){return f.c&&a(e||1)&&(f.c=null,f.t=NaN),1},0,l),g.active=r,v.event&&v.event.start.call(n,n.__data__,t),p=[],v.tween.forEach(function(e,r){(r=r.call(n,n.__data__,t))&&p.push(r)}),h=v.ease,s=v.duration}function a(i){for(var u=i/s,o=h(u),a=p.length;a>0;)p[--a].call(n,o);return u>=1?(v.event&&v.event.end.call(n,n.__data__,t),--g.count?delete g[r]:delete n[e],1):void 0}var l,f,s,h,p,g=n[e]||(n[e]={active:0,count:0}),v=g[r];v||(l=i.time,f=qn(u,0,l),v=g[r]={tween:new c,time:l,timer:f,delay:i.delay,duration:i.duration,ease:i.ease,index:t},i=null,++g.count)}function no(n,t,e){n.attr("transform",function(n){var r=t(n);return"translate("+(isFinite(r)?r:e(n))+",0)"})}function to(n,t,e){n.attr("transform",function(n){var r=t(n);return"translate(0,"+(isFinite(r)?r:e(n))+")"})}function eo(n){return n.toISOString()}function ro(n,t,e){function r(t){return n(t)}function i(n,e){var r=n[1]-n[0],i=r/e,u=ao.bisect(Kl,i);return u==Kl.length?[t.year,Ki(n.map(function(n){return n/31536e6}),e)[2]]:u?t[i/Kl[u-1]<Kl[u]/i?u-1:u]:[tc,Ki(n,e)[2]]}return r.invert=function(t){return io(n.invert(t))},r.domain=function(t){return arguments.length?(n.domain(t),r):n.domain().map(io)},r.nice=function(n,t){function e(e){return!isNaN(e)&&!n.range(e,io(+e+1),t).length}var u=r.domain(),o=Yi(u),a=null==n?i(o,10):"number"==typeof n&&i(o,n);return a&&(n=a[0],t=a[1]),r.domain(Xi(u,t>1?{floor:function(t){for(;e(t=n.floor(t));)t=io(t-1);return t},ceil:function(t){for(;e(t=n.ceil(t));)t=io(+t+1);return t}}:n))},r.ticks=function(n,t){var e=Yi(r.domain()),u=null==n?i(e,10):"number"==typeof n?i(e,n):!n.range&&[{range:n},t];return u&&(n=u[0],t=u[1]),n.range(e[0],io(+e[1]+1),1>t?1:t)},r.tickFormat=function(){return e},r.copy=function(){return ro(n.copy(),t,e)},Ji(r,n)}function io(n){return new Date(n)}function uo(n){return JSON.parse(n.responseText)}function oo(n){var t=fo.createRange();return t.selectNode(fo.body),t.createContextualFragment(n.responseText)}var ao={version:"3.5.17"},lo=[].slice,co=function(n){return lo.call(n)},fo=this.document;if(fo)try{co(fo.documentElement.childNodes)[0].nodeType}catch(so){co=function(n){for(var t=n.length,e=new Array(t);t--;)e[t]=n[t];return e}}if(Date.now||(Date.now=function(){return+new Date}),fo)try{fo.createElement("DIV").style.setProperty("opacity",0,"")}catch(ho){var po=this.Element.prototype,go=po.setAttribute,vo=po.setAttributeNS,yo=this.CSSStyleDeclaration.prototype,mo=yo.setProperty;po.setAttribute=function(n,t){go.call(this,n,t+"")},po.setAttributeNS=function(n,t,e){vo.call(this,n,t,e+"")},yo.setProperty=function(n,t,e){mo.call(this,n,t+"",e)}}ao.ascending=e,ao.descending=function(n,t){return n>t?-1:t>n?1:t>=n?0:NaN},ao.min=function(n,t){var e,r,i=-1,u=n.length;if(1===arguments.length){for(;++i<u;)if(null!=(r=n[i])&&r>=r){e=r;break}for(;++i<u;)null!=(r=n[i])&&e>r&&(e=r)}else{for(;++i<u;)if(null!=(r=t.call(n,n[i],i))&&r>=r){e=r;break}for(;++i<u;)null!=(r=t.call(n,n[i],i))&&e>r&&(e=r)}return e},ao.max=function(n,t){var e,r,i=-1,u=n.length;if(1===arguments.length){for(;++i<u;)if(null!=(r=n[i])&&r>=r){e=r;break}for(;++i<u;)null!=(r=n[i])&&r>e&&(e=r)}else{for(;++i<u;)if(null!=(r=t.call(n,n[i],i))&&r>=r){e=r;break}for(;++i<u;)null!=(r=t.call(n,n[i],i))&&r>e&&(e=r)}return e},ao.extent=function(n,t){var e,r,i,u=-1,o=n.length;if(1===arguments.length){for(;++u<o;)if(null!=(r=n[u])&&r>=r){e=i=r;break}for(;++u<o;)null!=(r=n[u])&&(e>r&&(e=r),r>i&&(i=r))}else{for(;++u<o;)if(null!=(r=t.call(n,n[u],u))&&r>=r){e=i=r;break}for(;++u<o;)null!=(r=t.call(n,n[u],u))&&(e>r&&(e=r),r>i&&(i=r))}return[e,i]},ao.sum=function(n,t){var e,r=0,u=n.length,o=-1;if(1===arguments.length)for(;++o<u;)i(e=+n[o])&&(r+=e);else for(;++o<u;)i(e=+t.call(n,n[o],o))&&(r+=e);return r},ao.mean=function(n,t){var e,u=0,o=n.length,a=-1,l=o;if(1===arguments.length)for(;++a<o;)i(e=r(n[a]))?u+=e:--l;else for(;++a<o;)i(e=r(t.call(n,n[a],a)))?u+=e:--l;return l?u/l:void 0},ao.quantile=function(n,t){var e=(n.length-1)*t+1,r=Math.floor(e),i=+n[r-1],u=e-r;return u?i+u*(n[r]-i):i},ao.median=function(n,t){var u,o=[],a=n.length,l=-1;if(1===arguments.length)for(;++l<a;)i(u=r(n[l]))&&o.push(u);else for(;++l<a;)i(u=r(t.call(n,n[l],l)))&&o.push(u);return o.length?ao.quantile(o.sort(e),.5):void 0},ao.variance=function(n,t){var e,u,o=n.length,a=0,l=0,c=-1,f=0;if(1===arguments.length)for(;++c<o;)i(e=r(n[c]))&&(u=e-a,a+=u/++f,l+=u*(e-a));else for(;++c<o;)i(e=r(t.call(n,n[c],c)))&&(u=e-a,a+=u/++f,l+=u*(e-a));return f>1?l/(f-1):void 0},ao.deviation=function(){var n=ao.variance.apply(this,arguments);return n?Math.sqrt(n):n};var Mo=u(e);ao.bisectLeft=Mo.left,ao.bisect=ao.bisectRight=Mo.right,ao.bisector=function(n){return u(1===n.length?function(t,r){return e(n(t),r)}:n)},ao.shuffle=function(n,t,e){(u=arguments.length)<3&&(e=n.length,2>u&&(t=0));for(var r,i,u=e-t;u;)i=Math.random()*u--|0,r=n[u+t],n[u+t]=n[i+t],n[i+t]=r;return n},ao.permute=function(n,t){for(var e=t.length,r=new Array(e);e--;)r[e]=n[t[e]];return r},ao.pairs=function(n){for(var t,e=0,r=n.length-1,i=n[0],u=new Array(0>r?0:r);r>e;)u[e]=[t=i,i=n[++e]];return u},ao.transpose=function(n){if(!(i=n.length))return[];for(var t=-1,e=ao.min(n,o),r=new Array(e);++t<e;)for(var i,u=-1,a=r[t]=new Array(i);++u<i;)a[u]=n[u][t];return r},ao.zip=function(){return ao.transpose(arguments)},ao.keys=function(n){var t=[];for(var e in n)t.push(e);return t},ao.values=function(n){var t=[];for(var e in n)t.push(n[e]);return t},ao.entries=function(n){var t=[];for(var e in n)t.push({key:e,value:n[e]});return t},ao.merge=function(n){for(var t,e,r,i=n.length,u=-1,o=0;++u<i;)o+=n[u].length;for(e=new Array(o);--i>=0;)for(r=n[i],t=r.length;--t>=0;)e[--o]=r[t];return e};var xo=Math.abs;ao.range=function(n,t,e){if(arguments.length<3&&(e=1,arguments.length<2&&(t=n,n=0)),(t-n)/e===1/0)throw new Error("infinite range");var r,i=[],u=a(xo(e)),o=-1;if(n*=u,t*=u,e*=u,0>e)for(;(r=n+e*++o)>t;)i.push(r/u);else for(;(r=n+e*++o)<t;)i.push(r/u);return i},ao.map=function(n,t){var e=new c;if(n instanceof c)n.forEach(function(n,t){e.set(n,t)});else if(Array.isArray(n)){var r,i=-1,u=n.length;if(1===arguments.length)for(;++i<u;)e.set(i,n[i]);else for(;++i<u;)e.set(t.call(n,r=n[i],i),r)}else for(var o in n)e.set(o,n[o]);return e};var bo="__proto__",_o="\x00";l(c,{has:h,get:function(n){return this._[f(n)]},set:function(n,t){return this._[f(n)]=t},remove:p,keys:g,values:function(){var n=[];for(var t in this._)n.push(this._[t]);return n},entries:function(){var n=[];for(var t in this._)n.push({key:s(t),value:this._[t]});return n},size:v,empty:d,forEach:function(n){for(var t in this._)n.call(this,s(t),this._[t])}}),ao.nest=function(){function n(t,o,a){if(a>=u.length)return r?r.call(i,o):e?o.sort(e):o;for(var l,f,s,h,p=-1,g=o.length,v=u[a++],d=new c;++p<g;)(h=d.get(l=v(f=o[p])))?h.push(f):d.set(l,[f]);return t?(f=t(),s=function(e,r){f.set(e,n(t,r,a))}):(f={},s=function(e,r){f[e]=n(t,r,a)}),d.forEach(s),f}function t(n,e){if(e>=u.length)return n;var r=[],i=o[e++];return n.forEach(function(n,i){r.push({key:n,values:t(i,e)})}),i?r.sort(function(n,t){return i(n.key,t.key)}):r}var e,r,i={},u=[],o=[];return i.map=function(t,e){return n(e,t,0)},i.entries=function(e){return t(n(ao.map,e,0),0)},i.key=function(n){return u.push(n),i},i.sortKeys=function(n){return o[u.length-1]=n,i},i.sortValues=function(n){return e=n,i},i.rollup=function(n){return r=n,i},i},ao.set=function(n){var t=new y;if(n)for(var e=0,r=n.length;r>e;++e)t.add(n[e]);return t},l(y,{has:h,add:function(n){return this._[f(n+="")]=!0,n},remove:p,values:g,size:v,empty:d,forEach:function(n){for(var t in this._)n.call(this,s(t))}}),ao.behavior={},ao.rebind=function(n,t){for(var e,r=1,i=arguments.length;++r<i;)n[e=arguments[r]]=M(n,t,t[e]);return n};var wo=["webkit","ms","moz","Moz","o","O"];ao.dispatch=function(){for(var n=new _,t=-1,e=arguments.length;++t<e;)n[arguments[t]]=w(n);return n},_.prototype.on=function(n,t){var e=n.indexOf("."),r="";if(e>=0&&(r=n.slice(e+1),n=n.slice(0,e)),n)return arguments.length<2?this[n].on(r):this[n].on(r,t);if(2===arguments.length){if(null==t)for(n in this)this.hasOwnProperty(n)&&this[n].on(r,null);return this}},ao.event=null,ao.requote=function(n){return n.replace(So,"\\$&")};var So=/[\\\^\$\*\+\?\|\[\]\(\)\.\{\}]/g,ko={}.__proto__?function(n,t){n.__proto__=t}:function(n,t){for(var e in t)n[e]=t[e]},No=function(n,t){return t.querySelector(n)},Eo=function(n,t){return t.querySelectorAll(n)},Ao=function(n,t){var e=n.matches||n[x(n,"matchesSelector")];return(Ao=function(n,t){return e.call(n,t)})(n,t)};"function"==typeof Sizzle&&(No=function(n,t){return Sizzle(n,t)[0]||null},Eo=Sizzle,Ao=Sizzle.matchesSelector),ao.selection=function(){return ao.select(fo.documentElement)};var Co=ao.selection.prototype=[];Co.select=function(n){var t,e,r,i,u=[];n=A(n);for(var o=-1,a=this.length;++o<a;){u.push(t=[]),t.parentNode=(r=this[o]).parentNode;for(var l=-1,c=r.length;++l<c;)(i=r[l])?(t.push(e=n.call(i,i.__data__,l,o)),e&&"__data__"in i&&(e.__data__=i.__data__)):t.push(null)}return E(u)},Co.selectAll=function(n){var t,e,r=[];n=C(n);for(var i=-1,u=this.length;++i<u;)for(var o=this[i],a=-1,l=o.length;++a<l;)(e=o[a])&&(r.push(t=co(n.call(e,e.__data__,a,i))),t.parentNode=e);return E(r)};var zo="http://www.w3.org/1999/xhtml",Lo={svg:"http://www.w3.org/2000/svg",xhtml:zo,xlink:"http://www.w3.org/1999/xlink",xml:"http://www.w3.org/XML/1998/namespace",xmlns:"http://www.w3.org/2000/xmlns/"};ao.ns={prefix:Lo,qualify:function(n){var t=n.indexOf(":"),e=n;return t>=0&&"xmlns"!==(e=n.slice(0,t))&&(n=n.slice(t+1)),Lo.hasOwnProperty(e)?{space:Lo[e],local:n}:n}},Co.attr=function(n,t){if(arguments.length<2){if("string"==typeof n){var e=this.node();return n=ao.ns.qualify(n),n.local?e.getAttributeNS(n.space,n.local):e.getAttribute(n)}for(t in n)this.each(z(t,n[t]));return this}return this.each(z(n,t))},Co.classed=function(n,t){if(arguments.length<2){if("string"==typeof n){var e=this.node(),r=(n=T(n)).length,i=-1;if(t=e.classList){for(;++i<r;)if(!t.contains(n[i]))return!1}else for(t=e.getAttribute("class");++i<r;)if(!q(n[i]).test(t))return!1;return!0}for(t in n)this.each(R(t,n[t]));return this}return this.each(R(n,t))},Co.style=function(n,e,r){var i=arguments.length;if(3>i){if("string"!=typeof n){2>i&&(e="");for(r in n)this.each(P(r,n[r],e));return this}if(2>i){var u=this.node();return t(u).getComputedStyle(u,null).getPropertyValue(n)}r=""}return this.each(P(n,e,r))},Co.property=function(n,t){if(arguments.length<2){if("string"==typeof n)return this.node()[n];for(t in n)this.each(U(t,n[t]));return this}return this.each(U(n,t))},Co.text=function(n){return arguments.length?this.each("function"==typeof n?function(){var t=n.apply(this,arguments);this.textContent=null==t?"":t}:null==n?function(){this.textContent=""}:function(){this.textContent=n}):this.node().textContent},Co.html=function(n){return arguments.length?this.each("function"==typeof n?function(){var t=n.apply(this,arguments);this.innerHTML=null==t?"":t}:null==n?function(){this.innerHTML=""}:function(){this.innerHTML=n}):this.node().innerHTML},Co.append=function(n){return n=j(n),this.select(function(){return this.appendChild(n.apply(this,arguments))})},Co.insert=function(n,t){return n=j(n),t=A(t),this.select(function(){return this.insertBefore(n.apply(this,arguments),t.apply(this,arguments)||null)})},Co.remove=function(){return this.each(F)},Co.data=function(n,t){function e(n,e){var r,i,u,o=n.length,s=e.length,h=Math.min(o,s),p=new Array(s),g=new Array(s),v=new Array(o);if(t){var d,y=new c,m=new Array(o);for(r=-1;++r<o;)(i=n[r])&&(y.has(d=t.call(i,i.__data__,r))?v[r]=i:y.set(d,i),m[r]=d);for(r=-1;++r<s;)(i=y.get(d=t.call(e,u=e[r],r)))?i!==!0&&(p[r]=i,i.__data__=u):g[r]=H(u),y.set(d,!0);for(r=-1;++r<o;)r in m&&y.get(m[r])!==!0&&(v[r]=n[r])}else{for(r=-1;++r<h;)i=n[r],u=e[r],i?(i.__data__=u,p[r]=i):g[r]=H(u);for(;s>r;++r)g[r]=H(e[r]);for(;o>r;++r)v[r]=n[r]}g.update=p,g.parentNode=p.parentNode=v.parentNode=n.parentNode,a.push(g),l.push(p),f.push(v)}var r,i,u=-1,o=this.length;if(!arguments.length){for(n=new Array(o=(r=this[0]).length);++u<o;)(i=r[u])&&(n[u]=i.__data__);return n}var a=Z([]),l=E([]),f=E([]);if("function"==typeof n)for(;++u<o;)e(r=this[u],n.call(r,r.parentNode.__data__,u));else for(;++u<o;)e(r=this[u],n);return l.enter=function(){return a},l.exit=function(){return f},l},Co.datum=function(n){return arguments.length?this.property("__data__",n):this.property("__data__")},Co.filter=function(n){var t,e,r,i=[];"function"!=typeof n&&(n=O(n));for(var u=0,o=this.length;o>u;u++){i.push(t=[]),t.parentNode=(e=this[u]).parentNode;for(var a=0,l=e.length;l>a;a++)(r=e[a])&&n.call(r,r.__data__,a,u)&&t.push(r)}return E(i)},Co.order=function(){for(var n=-1,t=this.length;++n<t;)for(var e,r=this[n],i=r.length-1,u=r[i];--i>=0;)(e=r[i])&&(u&&u!==e.nextSibling&&u.parentNode.insertBefore(e,u),u=e);return this},Co.sort=function(n){n=I.apply(this,arguments);for(var t=-1,e=this.length;++t<e;)this[t].sort(n);return this.order()},Co.each=function(n){return Y(this,function(t,e,r){n.call(t,t.__data__,e,r)})},Co.call=function(n){var t=co(arguments);return n.apply(t[0]=this,t),this},Co.empty=function(){return!this.node()},Co.node=function(){for(var n=0,t=this.length;t>n;n++)for(var e=this[n],r=0,i=e.length;i>r;r++){var u=e[r];if(u)return u}return null},Co.size=function(){var n=0;return Y(this,function(){++n}),n};var qo=[];ao.selection.enter=Z,ao.selection.enter.prototype=qo,qo.append=Co.append,qo.empty=Co.empty,qo.node=Co.node,qo.call=Co.call,qo.size=Co.size,qo.select=function(n){for(var t,e,r,i,u,o=[],a=-1,l=this.length;++a<l;){r=(i=this[a]).update,o.push(t=[]),t.parentNode=i.parentNode;for(var c=-1,f=i.length;++c<f;)(u=i[c])?(t.push(r[c]=e=n.call(i.parentNode,u.__data__,c,a)),e.__data__=u.__data__):t.push(null)}return E(o)},qo.insert=function(n,t){return arguments.length<2&&(t=V(this)),Co.insert.call(this,n,t)},ao.select=function(t){var e;return"string"==typeof t?(e=[No(t,fo)],e.parentNode=fo.documentElement):(e=[t],e.parentNode=n(t)),E([e])},ao.selectAll=function(n){var t;return"string"==typeof n?(t=co(Eo(n,fo)),t.parentNode=fo.documentElement):(t=co(n),t.parentNode=null),E([t])},Co.on=function(n,t,e){var r=arguments.length;if(3>r){if("string"!=typeof n){2>r&&(t=!1);for(e in n)this.each(X(e,n[e],t));return this}if(2>r)return(r=this.node()["__on"+n])&&r._;e=!1}return this.each(X(n,t,e))};var To=ao.map({mouseenter:"mouseover",mouseleave:"mouseout"});fo&&To.forEach(function(n){"on"+n in fo&&To.remove(n)});var Ro,Do=0;ao.mouse=function(n){return J(n,k())};var Po=this.navigator&&/WebKit/.test(this.navigator.userAgent)?-1:0;ao.touch=function(n,t,e){if(arguments.length<3&&(e=t,t=k().changedTouches),t)for(var r,i=0,u=t.length;u>i;++i)if((r=t[i]).identifier===e)return J(n,r)},ao.behavior.drag=function(){function n(){this.on("mousedown.drag",u).on("touchstart.drag",o)}function e(n,t,e,u,o){return function(){function a(){var n,e,r=t(h,v);r&&(n=r[0]-M[0],e=r[1]-M[1],g|=n|e,M=r,p({type:"drag",x:r[0]+c[0],y:r[1]+c[1],dx:n,dy:e}))}function l(){t(h,v)&&(y.on(u+d,null).on(o+d,null),m(g),p({type:"dragend"}))}var c,f=this,s=ao.event.target.correspondingElement||ao.event.target,h=f.parentNode,p=r.of(f,arguments),g=0,v=n(),d=".drag"+(null==v?"":"-"+v),y=ao.select(e(s)).on(u+d,a).on(o+d,l),m=W(s),M=t(h,v);i?(c=i.apply(f,arguments),c=[c.x-M[0],c.y-M[1]]):c=[0,0],p({type:"dragstart"})}}var r=N(n,"drag","dragstart","dragend"),i=null,u=e(b,ao.mouse,t,"mousemove","mouseup"),o=e(G,ao.touch,m,"touchmove","touchend");return n.origin=function(t){return arguments.length?(i=t,n):i},ao.rebind(n,r,"on")},ao.touches=function(n,t){return arguments.length<2&&(t=k().touches),t?co(t).map(function(t){var e=J(n,t);return e.identifier=t.identifier,e}):[]};var Uo=1e-6,jo=Uo*Uo,Fo=Math.PI,Ho=2*Fo,Oo=Ho-Uo,Io=Fo/2,Yo=Fo/180,Zo=180/Fo,Vo=Math.SQRT2,Xo=2,$o=4;ao.interpolateZoom=function(n,t){var e,r,i=n[0],u=n[1],o=n[2],a=t[0],l=t[1],c=t[2],f=a-i,s=l-u,h=f*f+s*s;if(jo>h)r=Math.log(c/o)/Vo,e=function(n){return[i+n*f,u+n*s,o*Math.exp(Vo*n*r)]};else{var p=Math.sqrt(h),g=(c*c-o*o+$o*h)/(2*o*Xo*p),v=(c*c-o*o-$o*h)/(2*c*Xo*p),d=Math.log(Math.sqrt(g*g+1)-g),y=Math.log(Math.sqrt(v*v+1)-v);r=(y-d)/Vo,e=function(n){var t=n*r,e=rn(d),a=o/(Xo*p)*(e*un(Vo*t+d)-en(d));return[i+a*f,u+a*s,o*e/rn(Vo*t+d)]}}return e.duration=1e3*r,e},ao.behavior.zoom=function(){function n(n){n.on(L,s).on(Wo+".zoom",p).on("dblclick.zoom",g).on(R,h)}function e(n){return[(n[0]-k.x)/k.k,(n[1]-k.y)/k.k]}function r(n){return[n[0]*k.k+k.x,n[1]*k.k+k.y]}function i(n){k.k=Math.max(A[0],Math.min(A[1],n))}function u(n,t){t=r(t),k.x+=n[0]-t[0],k.y+=n[1]-t[1]}function o(t,e,r,o){t.__chart__={x:k.x,y:k.y,k:k.k},i(Math.pow(2,o)),u(d=e,r),t=ao.select(t),C>0&&(t=t.transition().duration(C)),t.call(n.event)}function a(){b&&b.domain(x.range().map(function(n){return(n-k.x)/k.k}).map(x.invert)),w&&w.domain(_.range().map(function(n){return(n-k.y)/k.k}).map(_.invert))}function l(n){z++||n({type:"zoomstart"})}function c(n){a(),n({type:"zoom",scale:k.k,translate:[k.x,k.y]})}function f(n){--z||(n({type:"zoomend"}),d=null)}function s(){function n(){a=1,u(ao.mouse(i),h),c(o)}function r(){s.on(q,null).on(T,null),p(a),f(o)}var i=this,o=D.of(i,arguments),a=0,s=ao.select(t(i)).on(q,n).on(T,r),h=e(ao.mouse(i)),p=W(i);Il.call(i),l(o)}function h(){function n(){var n=ao.touches(g);return p=k.k,n.forEach(function(n){n.identifier in d&&(d[n.identifier]=e(n))}),n}function t(){var t=ao.event.target;ao.select(t).on(x,r).on(b,a),_.push(t);for(var e=ao.event.changedTouches,i=0,u=e.length;u>i;++i)d[e[i].identifier]=null;var l=n(),c=Date.now();if(1===l.length){if(500>c-M){var f=l[0];o(g,f,d[f.identifier],Math.floor(Math.log(k.k)/Math.LN2)+1),S()}M=c}else if(l.length>1){var f=l[0],s=l[1],h=f[0]-s[0],p=f[1]-s[1];y=h*h+p*p}}function r(){var n,t,e,r,o=ao.touches(g);Il.call(g);for(var a=0,l=o.length;l>a;++a,r=null)if(e=o[a],r=d[e.identifier]){if(t)break;n=e,t=r}if(r){var f=(f=e[0]-n[0])*f+(f=e[1]-n[1])*f,s=y&&Math.sqrt(f/y);n=[(n[0]+e[0])/2,(n[1]+e[1])/2],t=[(t[0]+r[0])/2,(t[1]+r[1])/2],i(s*p)}M=null,u(n,t),c(v)}function a(){if(ao.event.touches.length){for(var t=ao.event.changedTouches,e=0,r=t.length;r>e;++e)delete d[t[e].identifier];for(var i in d)return void n()}ao.selectAll(_).on(m,null),w.on(L,s).on(R,h),N(),f(v)}var p,g=this,v=D.of(g,arguments),d={},y=0,m=".zoom-"+ao.event.changedTouches[0].identifier,x="touchmove"+m,b="touchend"+m,_=[],w=ao.select(g),N=W(g);t(),l(v),w.on(L,null).on(R,t)}function p(){var n=D.of(this,arguments);m?clearTimeout(m):(Il.call(this),v=e(d=y||ao.mouse(this)),l(n)),m=setTimeout(function(){m=null,f(n)},50),S(),i(Math.pow(2,.002*Bo())*k.k),u(d,v),c(n)}function g(){var n=ao.mouse(this),t=Math.log(k.k)/Math.LN2;o(this,n,e(n),ao.event.shiftKey?Math.ceil(t)-1:Math.floor(t)+1)}var v,d,y,m,M,x,b,_,w,k={x:0,y:0,k:1},E=[960,500],A=Jo,C=250,z=0,L="mousedown.zoom",q="mousemove.zoom",T="mouseup.zoom",R="touchstart.zoom",D=N(n,"zoomstart","zoom","zoomend");return Wo||(Wo="onwheel"in fo?(Bo=function(){return-ao.event.deltaY*(ao.event.deltaMode?120:1)},"wheel"):"onmousewheel"in fo?(Bo=function(){return ao.event.wheelDelta},"mousewheel"):(Bo=function(){return-ao.event.detail},"MozMousePixelScroll")),n.event=function(n){n.each(function(){var n=D.of(this,arguments),t=k;Hl?ao.select(this).transition().each("start.zoom",function(){k=this.__chart__||{x:0,y:0,k:1},l(n)}).tween("zoom:zoom",function(){var e=E[0],r=E[1],i=d?d[0]:e/2,u=d?d[1]:r/2,o=ao.interpolateZoom([(i-k.x)/k.k,(u-k.y)/k.k,e/k.k],[(i-t.x)/t.k,(u-t.y)/t.k,e/t.k]);return function(t){var r=o(t),a=e/r[2];this.__chart__=k={x:i-r[0]*a,y:u-r[1]*a,k:a},c(n)}}).each("interrupt.zoom",function(){f(n)}).each("end.zoom",function(){f(n)}):(this.__chart__=k,l(n),c(n),f(n))})},n.translate=function(t){return arguments.length?(k={x:+t[0],y:+t[1],k:k.k},a(),n):[k.x,k.y]},n.scale=function(t){return arguments.length?(k={x:k.x,y:k.y,k:null},i(+t),a(),n):k.k},n.scaleExtent=function(t){return arguments.length?(A=null==t?Jo:[+t[0],+t[1]],n):A},n.center=function(t){return arguments.length?(y=t&&[+t[0],+t[1]],n):y},n.size=function(t){return arguments.length?(E=t&&[+t[0],+t[1]],n):E},n.duration=function(t){return arguments.length?(C=+t,n):C},n.x=function(t){return arguments.length?(b=t,x=t.copy(),k={x:0,y:0,k:1},n):b},n.y=function(t){return arguments.length?(w=t,_=t.copy(),k={x:0,y:0,k:1},n):w},ao.rebind(n,D,"on")};var Bo,Wo,Jo=[0,1/0];ao.color=an,an.prototype.toString=function(){return this.rgb()+""},ao.hsl=ln;var Go=ln.prototype=new an;Go.brighter=function(n){return n=Math.pow(.7,arguments.length?n:1),new ln(this.h,this.s,this.l/n)},Go.darker=function(n){return n=Math.pow(.7,arguments.length?n:1),new ln(this.h,this.s,n*this.l)},Go.rgb=function(){return cn(this.h,this.s,this.l)},ao.hcl=fn;var Ko=fn.prototype=new an;Ko.brighter=function(n){return new fn(this.h,this.c,Math.min(100,this.l+Qo*(arguments.length?n:1)))},Ko.darker=function(n){return new fn(this.h,this.c,Math.max(0,this.l-Qo*(arguments.length?n:1)))},Ko.rgb=function(){return sn(this.h,this.c,this.l).rgb()},ao.lab=hn;var Qo=18,na=.95047,ta=1,ea=1.08883,ra=hn.prototype=new an;ra.brighter=function(n){return new hn(Math.min(100,this.l+Qo*(arguments.length?n:1)),this.a,this.b)},ra.darker=function(n){return new hn(Math.max(0,this.l-Qo*(arguments.length?n:1)),this.a,this.b)},ra.rgb=function(){return pn(this.l,this.a,this.b)},ao.rgb=mn;var ia=mn.prototype=new an;ia.brighter=function(n){n=Math.pow(.7,arguments.length?n:1);var t=this.r,e=this.g,r=this.b,i=30;return t||e||r?(t&&i>t&&(t=i),e&&i>e&&(e=i),r&&i>r&&(r=i),new mn(Math.min(255,t/n),Math.min(255,e/n),Math.min(255,r/n))):new mn(i,i,i)},ia.darker=function(n){return n=Math.pow(.7,arguments.length?n:1),new mn(n*this.r,n*this.g,n*this.b)},ia.hsl=function(){return wn(this.r,this.g,this.b)},ia.toString=function(){return"#"+bn(this.r)+bn(this.g)+bn(this.b)};var ua=ao.map({aliceblue:15792383,antiquewhite:16444375,aqua:65535,aquamarine:8388564,azure:15794175,beige:16119260,bisque:16770244,black:0,blanchedalmond:16772045,blue:255,blueviolet:9055202,brown:10824234,burlywood:14596231,cadetblue:6266528,chartreuse:8388352,chocolate:13789470,coral:16744272,cornflowerblue:6591981,cornsilk:16775388,crimson:14423100,cyan:65535,darkblue:139,darkcyan:35723,darkgoldenrod:12092939,darkgray:11119017,darkgreen:25600,darkgrey:11119017,darkkhaki:12433259,darkmagenta:9109643,darkolivegreen:5597999,darkorange:16747520,darkorchid:10040012,darkred:9109504,darksalmon:15308410,darkseagreen:9419919,darkslateblue:4734347,darkslategray:3100495,darkslategrey:3100495,darkturquoise:52945,darkviolet:9699539,deeppink:16716947,deepskyblue:49151,dimgray:6908265,dimgrey:6908265,dodgerblue:2003199,firebrick:11674146,floralwhite:16775920,forestgreen:2263842,fuchsia:16711935,gainsboro:14474460,ghostwhite:16316671,gold:16766720,goldenrod:14329120,gray:8421504,green:32768,greenyellow:11403055,grey:8421504,honeydew:15794160,hotpink:16738740,indianred:13458524,indigo:4915330,ivory:16777200,khaki:15787660,lavender:15132410,lavenderblush:16773365,lawngreen:8190976,lemonchiffon:16775885,lightblue:11393254,lightcoral:15761536,lightcyan:14745599,lightgoldenrodyellow:16448210,lightgray:13882323,lightgreen:9498256,lightgrey:13882323,lightpink:16758465,lightsalmon:16752762,lightseagreen:2142890,lightskyblue:8900346,lightslategray:7833753,lightslategrey:7833753,lightsteelblue:11584734,lightyellow:16777184,lime:65280,limegreen:3329330,linen:16445670,magenta:16711935,maroon:8388608,mediumaquamarine:6737322,mediumblue:205,mediumorchid:12211667,mediumpurple:9662683,mediumseagreen:3978097,mediumslateblue:8087790,mediumspringgreen:64154,mediumturquoise:4772300,mediumvioletred:13047173,midnightblue:1644912,mintcream:16121850,mistyrose:16770273,moccasin:16770229,navajowhite:16768685,navy:128,oldlace:16643558,olive:8421376,olivedrab:7048739,orange:16753920,orangered:16729344,orchid:14315734,palegoldenrod:15657130,palegreen:10025880,paleturquoise:11529966,palevioletred:14381203,papayawhip:16773077,peachpuff:16767673,peru:13468991,pink:16761035,plum:14524637,powderblue:11591910,purple:8388736,rebeccapurple:6697881,red:16711680,rosybrown:12357519,royalblue:4286945,saddlebrown:9127187,salmon:16416882,sandybrown:16032864,seagreen:3050327,seashell:16774638,sienna:10506797,silver:12632256,skyblue:8900331,slateblue:6970061,slategray:7372944,slategrey:7372944,snow:16775930,springgreen:65407,steelblue:4620980,tan:13808780,teal:32896,thistle:14204888,tomato:16737095,turquoise:4251856,violet:15631086,wheat:16113331,white:16777215,whitesmoke:16119285,yellow:16776960,yellowgreen:10145074});ua.forEach(function(n,t){ua.set(n,Mn(t))}),ao.functor=En,ao.xhr=An(m),ao.dsv=function(n,t){function e(n,e,u){arguments.length<3&&(u=e,e=null);var o=Cn(n,t,null==e?r:i(e),u);return o.row=function(n){return arguments.length?o.response(null==(e=n)?r:i(n)):e},o}function r(n){return e.parse(n.responseText)}function i(n){return function(t){return e.parse(t.responseText,n)}}function u(t){return t.map(o).join(n)}function o(n){return a.test(n)?'"'+n.replace(/\"/g,'""')+'"':n}var a=new RegExp('["'+n+"\n]"),l=n.charCodeAt(0);return e.parse=function(n,t){var r;return e.parseRows(n,function(n,e){if(r)return r(n,e-1);var i=new Function("d","return {"+n.map(function(n,t){return JSON.stringify(n)+": d["+t+"]"}).join(",")+"}");r=t?function(n,e){return t(i(n),e)}:i})},e.parseRows=function(n,t){function e(){if(f>=c)return o;if(i)return i=!1,u;var t=f;if(34===n.charCodeAt(t)){for(var e=t;e++<c;)if(34===n.charCodeAt(e)){if(34!==n.charCodeAt(e+1))break;++e}f=e+2;var r=n.charCodeAt(e+1);return 13===r?(i=!0,10===n.charCodeAt(e+2)&&++f):10===r&&(i=!0),n.slice(t+1,e).replace(/""/g,'"')}for(;c>f;){var r=n.charCodeAt(f++),a=1;if(10===r)i=!0;else if(13===r)i=!0,10===n.charCodeAt(f)&&(++f,++a);else if(r!==l)continue;return n.slice(t,f-a)}return n.slice(t)}for(var r,i,u={},o={},a=[],c=n.length,f=0,s=0;(r=e())!==o;){for(var h=[];r!==u&&r!==o;)h.push(r),r=e();t&&null==(h=t(h,s++))||a.push(h)}return a},e.format=function(t){if(Array.isArray(t[0]))return e.formatRows(t);var r=new y,i=[];return t.forEach(function(n){for(var t in n)r.has(t)||i.push(r.add(t))}),[i.map(o).join(n)].concat(t.map(function(t){return i.map(function(n){return o(t[n])}).join(n)})).join("\n")},e.formatRows=function(n){return n.map(u).join("\n")},e},ao.csv=ao.dsv(",","text/csv"),ao.tsv=ao.dsv("	","text/tab-separated-values");var oa,aa,la,ca,fa=this[x(this,"requestAnimationFrame")]||function(n){setTimeout(n,17)};ao.timer=function(){qn.apply(this,arguments)},ao.timer.flush=function(){Rn(),Dn()},ao.round=function(n,t){return t?Math.round(n*(t=Math.pow(10,t)))/t:Math.round(n)};var sa=["y","z","a","f","p","n","\xb5","m","","k","M","G","T","P","E","Z","Y"].map(Un);ao.formatPrefix=function(n,t){var e=0;return(n=+n)&&(0>n&&(n*=-1),t&&(n=ao.round(n,Pn(n,t))),e=1+Math.floor(1e-12+Math.log(n)/Math.LN10),e=Math.max(-24,Math.min(24,3*Math.floor((e-1)/3)))),sa[8+e/3]};var ha=/(?:([^{])?([<>=^]))?([+\- ])?([$#])?(0)?(\d+)?(,)?(\.-?\d+)?([a-z%])?/i,pa=ao.map({b:function(n){return n.toString(2)},c:function(n){return String.fromCharCode(n)},o:function(n){return n.toString(8)},x:function(n){return n.toString(16)},X:function(n){return n.toString(16).toUpperCase()},g:function(n,t){return n.toPrecision(t)},e:function(n,t){return n.toExponential(t)},f:function(n,t){return n.toFixed(t)},r:function(n,t){return(n=ao.round(n,Pn(n,t))).toFixed(Math.max(0,Math.min(20,Pn(n*(1+1e-15),t))))}}),ga=ao.time={},va=Date;Hn.prototype={getDate:function(){return this._.getUTCDate()},getDay:function(){return this._.getUTCDay()},getFullYear:function(){return this._.getUTCFullYear()},getHours:function(){return this._.getUTCHours()},getMilliseconds:function(){return this._.getUTCMilliseconds()},getMinutes:function(){return this._.getUTCMinutes()},getMonth:function(){return this._.getUTCMonth()},getSeconds:function(){return this._.getUTCSeconds()},getTime:function(){return this._.getTime()},getTimezoneOffset:function(){return 0},valueOf:function(){return this._.valueOf()},setDate:function(){da.setUTCDate.apply(this._,arguments)},setDay:function(){da.setUTCDay.apply(this._,arguments)},setFullYear:function(){da.setUTCFullYear.apply(this._,arguments)},setHours:function(){da.setUTCHours.apply(this._,arguments)},setMilliseconds:function(){da.setUTCMilliseconds.apply(this._,arguments)},setMinutes:function(){da.setUTCMinutes.apply(this._,arguments)},setMonth:function(){da.setUTCMonth.apply(this._,arguments)},setSeconds:function(){da.setUTCSeconds.apply(this._,arguments)},setTime:function(){da.setTime.apply(this._,arguments)}};var da=Date.prototype;ga.year=On(function(n){return n=ga.day(n),n.setMonth(0,1),n},function(n,t){n.setFullYear(n.getFullYear()+t)},function(n){return n.getFullYear()}),ga.years=ga.year.range,ga.years.utc=ga.year.utc.range,ga.day=On(function(n){var t=new va(2e3,0);return t.setFullYear(n.getFullYear(),n.getMonth(),n.getDate()),t},function(n,t){n.setDate(n.getDate()+t)},function(n){return n.getDate()-1}),ga.days=ga.day.range,ga.days.utc=ga.day.utc.range,ga.dayOfYear=function(n){var t=ga.year(n);return Math.floor((n-t-6e4*(n.getTimezoneOffset()-t.getTimezoneOffset()))/864e5)},["sunday","monday","tuesday","wednesday","thursday","friday","saturday"].forEach(function(n,t){t=7-t;var e=ga[n]=On(function(n){return(n=ga.day(n)).setDate(n.getDate()-(n.getDay()+t)%7),n},function(n,t){n.setDate(n.getDate()+7*Math.floor(t))},function(n){var e=ga.year(n).getDay();return Math.floor((ga.dayOfYear(n)+(e+t)%7)/7)-(e!==t)});ga[n+"s"]=e.range,ga[n+"s"].utc=e.utc.range,ga[n+"OfYear"]=function(n){var e=ga.year(n).getDay();return Math.floor((ga.dayOfYear(n)+(e+t)%7)/7)}}),ga.week=ga.sunday,ga.weeks=ga.sunday.range,ga.weeks.utc=ga.sunday.utc.range,ga.weekOfYear=ga.sundayOfYear;var ya={"-":"",_:" ",0:"0"},ma=/^\s*\d+/,Ma=/^%/;ao.locale=function(n){return{numberFormat:jn(n),timeFormat:Yn(n)}};var xa=ao.locale({decimal:".",thousands:",",grouping:[3],currency:["$",""],dateTime:"%a %b %e %X %Y",date:"%m/%d/%Y",time:"%H:%M:%S",periods:["AM","PM"],days:["Sunday","Monday","Tuesday","Wednesday","Thursday","Friday","Saturday"],
shortDays:["Sun","Mon","Tue","Wed","Thu","Fri","Sat"],months:["January","February","March","April","May","June","July","August","September","October","November","December"],shortMonths:["Jan","Feb","Mar","Apr","May","Jun","Jul","Aug","Sep","Oct","Nov","Dec"]});ao.format=xa.numberFormat,ao.geo={},ft.prototype={s:0,t:0,add:function(n){st(n,this.t,ba),st(ba.s,this.s,this),this.s?this.t+=ba.t:this.s=ba.t},reset:function(){this.s=this.t=0},valueOf:function(){return this.s}};var ba=new ft;ao.geo.stream=function(n,t){n&&_a.hasOwnProperty(n.type)?_a[n.type](n,t):ht(n,t)};var _a={Feature:function(n,t){ht(n.geometry,t)},FeatureCollection:function(n,t){for(var e=n.features,r=-1,i=e.length;++r<i;)ht(e[r].geometry,t)}},wa={Sphere:function(n,t){t.sphere()},Point:function(n,t){n=n.coordinates,t.point(n[0],n[1],n[2])},MultiPoint:function(n,t){for(var e=n.coordinates,r=-1,i=e.length;++r<i;)n=e[r],t.point(n[0],n[1],n[2])},LineString:function(n,t){pt(n.coordinates,t,0)},MultiLineString:function(n,t){for(var e=n.coordinates,r=-1,i=e.length;++r<i;)pt(e[r],t,0)},Polygon:function(n,t){gt(n.coordinates,t)},MultiPolygon:function(n,t){for(var e=n.coordinates,r=-1,i=e.length;++r<i;)gt(e[r],t)},GeometryCollection:function(n,t){for(var e=n.geometries,r=-1,i=e.length;++r<i;)ht(e[r],t)}};ao.geo.area=function(n){return Sa=0,ao.geo.stream(n,Na),Sa};var Sa,ka=new ft,Na={sphere:function(){Sa+=4*Fo},point:b,lineStart:b,lineEnd:b,polygonStart:function(){ka.reset(),Na.lineStart=vt},polygonEnd:function(){var n=2*ka;Sa+=0>n?4*Fo+n:n,Na.lineStart=Na.lineEnd=Na.point=b}};ao.geo.bounds=function(){function n(n,t){M.push(x=[f=n,h=n]),s>t&&(s=t),t>p&&(p=t)}function t(t,e){var r=dt([t*Yo,e*Yo]);if(y){var i=mt(y,r),u=[i[1],-i[0],0],o=mt(u,i);bt(o),o=_t(o);var l=t-g,c=l>0?1:-1,v=o[0]*Zo*c,d=xo(l)>180;if(d^(v>c*g&&c*t>v)){var m=o[1]*Zo;m>p&&(p=m)}else if(v=(v+360)%360-180,d^(v>c*g&&c*t>v)){var m=-o[1]*Zo;s>m&&(s=m)}else s>e&&(s=e),e>p&&(p=e);d?g>t?a(f,t)>a(f,h)&&(h=t):a(t,h)>a(f,h)&&(f=t):h>=f?(f>t&&(f=t),t>h&&(h=t)):t>g?a(f,t)>a(f,h)&&(h=t):a(t,h)>a(f,h)&&(f=t)}else n(t,e);y=r,g=t}function e(){b.point=t}function r(){x[0]=f,x[1]=h,b.point=n,y=null}function i(n,e){if(y){var r=n-g;m+=xo(r)>180?r+(r>0?360:-360):r}else v=n,d=e;Na.point(n,e),t(n,e)}function u(){Na.lineStart()}function o(){i(v,d),Na.lineEnd(),xo(m)>Uo&&(f=-(h=180)),x[0]=f,x[1]=h,y=null}function a(n,t){return(t-=n)<0?t+360:t}function l(n,t){return n[0]-t[0]}function c(n,t){return t[0]<=t[1]?t[0]<=n&&n<=t[1]:n<t[0]||t[1]<n}var f,s,h,p,g,v,d,y,m,M,x,b={point:n,lineStart:e,lineEnd:r,polygonStart:function(){b.point=i,b.lineStart=u,b.lineEnd=o,m=0,Na.polygonStart()},polygonEnd:function(){Na.polygonEnd(),b.point=n,b.lineStart=e,b.lineEnd=r,0>ka?(f=-(h=180),s=-(p=90)):m>Uo?p=90:-Uo>m&&(s=-90),x[0]=f,x[1]=h}};return function(n){p=h=-(f=s=1/0),M=[],ao.geo.stream(n,b);var t=M.length;if(t){M.sort(l);for(var e,r=1,i=M[0],u=[i];t>r;++r)e=M[r],c(e[0],i)||c(e[1],i)?(a(i[0],e[1])>a(i[0],i[1])&&(i[1]=e[1]),a(e[0],i[1])>a(i[0],i[1])&&(i[0]=e[0])):u.push(i=e);for(var o,e,g=-(1/0),t=u.length-1,r=0,i=u[t];t>=r;i=e,++r)e=u[r],(o=a(i[1],e[0]))>g&&(g=o,f=e[0],h=i[1])}return M=x=null,f===1/0||s===1/0?[[NaN,NaN],[NaN,NaN]]:[[f,s],[h,p]]}}(),ao.geo.centroid=function(n){Ea=Aa=Ca=za=La=qa=Ta=Ra=Da=Pa=Ua=0,ao.geo.stream(n,ja);var t=Da,e=Pa,r=Ua,i=t*t+e*e+r*r;return jo>i&&(t=qa,e=Ta,r=Ra,Uo>Aa&&(t=Ca,e=za,r=La),i=t*t+e*e+r*r,jo>i)?[NaN,NaN]:[Math.atan2(e,t)*Zo,tn(r/Math.sqrt(i))*Zo]};var Ea,Aa,Ca,za,La,qa,Ta,Ra,Da,Pa,Ua,ja={sphere:b,point:St,lineStart:Nt,lineEnd:Et,polygonStart:function(){ja.lineStart=At},polygonEnd:function(){ja.lineStart=Nt}},Fa=Rt(zt,jt,Ht,[-Fo,-Fo/2]),Ha=1e9;ao.geo.clipExtent=function(){var n,t,e,r,i,u,o={stream:function(n){return i&&(i.valid=!1),i=u(n),i.valid=!0,i},extent:function(a){return arguments.length?(u=Zt(n=+a[0][0],t=+a[0][1],e=+a[1][0],r=+a[1][1]),i&&(i.valid=!1,i=null),o):[[n,t],[e,r]]}};return o.extent([[0,0],[960,500]])},(ao.geo.conicEqualArea=function(){return Vt(Xt)}).raw=Xt,ao.geo.albers=function(){return ao.geo.conicEqualArea().rotate([96,0]).center([-.6,38.7]).parallels([29.5,45.5]).scale(1070)},ao.geo.albersUsa=function(){function n(n){var u=n[0],o=n[1];return t=null,e(u,o),t||(r(u,o),t)||i(u,o),t}var t,e,r,i,u=ao.geo.albers(),o=ao.geo.conicEqualArea().rotate([154,0]).center([-2,58.5]).parallels([55,65]),a=ao.geo.conicEqualArea().rotate([157,0]).center([-3,19.9]).parallels([8,18]),l={point:function(n,e){t=[n,e]}};return n.invert=function(n){var t=u.scale(),e=u.translate(),r=(n[0]-e[0])/t,i=(n[1]-e[1])/t;return(i>=.12&&.234>i&&r>=-.425&&-.214>r?o:i>=.166&&.234>i&&r>=-.214&&-.115>r?a:u).invert(n)},n.stream=function(n){var t=u.stream(n),e=o.stream(n),r=a.stream(n);return{point:function(n,i){t.point(n,i),e.point(n,i),r.point(n,i)},sphere:function(){t.sphere(),e.sphere(),r.sphere()},lineStart:function(){t.lineStart(),e.lineStart(),r.lineStart()},lineEnd:function(){t.lineEnd(),e.lineEnd(),r.lineEnd()},polygonStart:function(){t.polygonStart(),e.polygonStart(),r.polygonStart()},polygonEnd:function(){t.polygonEnd(),e.polygonEnd(),r.polygonEnd()}}},n.precision=function(t){return arguments.length?(u.precision(t),o.precision(t),a.precision(t),n):u.precision()},n.scale=function(t){return arguments.length?(u.scale(t),o.scale(.35*t),a.scale(t),n.translate(u.translate())):u.scale()},n.translate=function(t){if(!arguments.length)return u.translate();var c=u.scale(),f=+t[0],s=+t[1];return e=u.translate(t).clipExtent([[f-.455*c,s-.238*c],[f+.455*c,s+.238*c]]).stream(l).point,r=o.translate([f-.307*c,s+.201*c]).clipExtent([[f-.425*c+Uo,s+.12*c+Uo],[f-.214*c-Uo,s+.234*c-Uo]]).stream(l).point,i=a.translate([f-.205*c,s+.212*c]).clipExtent([[f-.214*c+Uo,s+.166*c+Uo],[f-.115*c-Uo,s+.234*c-Uo]]).stream(l).point,n},n.scale(1070)};var Oa,Ia,Ya,Za,Va,Xa,$a={point:b,lineStart:b,lineEnd:b,polygonStart:function(){Ia=0,$a.lineStart=$t},polygonEnd:function(){$a.lineStart=$a.lineEnd=$a.point=b,Oa+=xo(Ia/2)}},Ba={point:Bt,lineStart:b,lineEnd:b,polygonStart:b,polygonEnd:b},Wa={point:Gt,lineStart:Kt,lineEnd:Qt,polygonStart:function(){Wa.lineStart=ne},polygonEnd:function(){Wa.point=Gt,Wa.lineStart=Kt,Wa.lineEnd=Qt}};ao.geo.path=function(){function n(n){return n&&("function"==typeof a&&u.pointRadius(+a.apply(this,arguments)),o&&o.valid||(o=i(u)),ao.geo.stream(n,o)),u.result()}function t(){return o=null,n}var e,r,i,u,o,a=4.5;return n.area=function(n){return Oa=0,ao.geo.stream(n,i($a)),Oa},n.centroid=function(n){return Ca=za=La=qa=Ta=Ra=Da=Pa=Ua=0,ao.geo.stream(n,i(Wa)),Ua?[Da/Ua,Pa/Ua]:Ra?[qa/Ra,Ta/Ra]:La?[Ca/La,za/La]:[NaN,NaN]},n.bounds=function(n){return Va=Xa=-(Ya=Za=1/0),ao.geo.stream(n,i(Ba)),[[Ya,Za],[Va,Xa]]},n.projection=function(n){return arguments.length?(i=(e=n)?n.stream||re(n):m,t()):e},n.context=function(n){return arguments.length?(u=null==(r=n)?new Wt:new te(n),"function"!=typeof a&&u.pointRadius(a),t()):r},n.pointRadius=function(t){return arguments.length?(a="function"==typeof t?t:(u.pointRadius(+t),+t),n):a},n.projection(ao.geo.albersUsa()).context(null)},ao.geo.transform=function(n){return{stream:function(t){var e=new ie(t);for(var r in n)e[r]=n[r];return e}}},ie.prototype={point:function(n,t){this.stream.point(n,t)},sphere:function(){this.stream.sphere()},lineStart:function(){this.stream.lineStart()},lineEnd:function(){this.stream.lineEnd()},polygonStart:function(){this.stream.polygonStart()},polygonEnd:function(){this.stream.polygonEnd()}},ao.geo.projection=oe,ao.geo.projectionMutator=ae,(ao.geo.equirectangular=function(){return oe(ce)}).raw=ce.invert=ce,ao.geo.rotation=function(n){function t(t){return t=n(t[0]*Yo,t[1]*Yo),t[0]*=Zo,t[1]*=Zo,t}return n=se(n[0]%360*Yo,n[1]*Yo,n.length>2?n[2]*Yo:0),t.invert=function(t){return t=n.invert(t[0]*Yo,t[1]*Yo),t[0]*=Zo,t[1]*=Zo,t},t},fe.invert=ce,ao.geo.circle=function(){function n(){var n="function"==typeof r?r.apply(this,arguments):r,t=se(-n[0]*Yo,-n[1]*Yo,0).invert,i=[];return e(null,null,1,{point:function(n,e){i.push(n=t(n,e)),n[0]*=Zo,n[1]*=Zo}}),{type:"Polygon",coordinates:[i]}}var t,e,r=[0,0],i=6;return n.origin=function(t){return arguments.length?(r=t,n):r},n.angle=function(r){return arguments.length?(e=ve((t=+r)*Yo,i*Yo),n):t},n.precision=function(r){return arguments.length?(e=ve(t*Yo,(i=+r)*Yo),n):i},n.angle(90)},ao.geo.distance=function(n,t){var e,r=(t[0]-n[0])*Yo,i=n[1]*Yo,u=t[1]*Yo,o=Math.sin(r),a=Math.cos(r),l=Math.sin(i),c=Math.cos(i),f=Math.sin(u),s=Math.cos(u);return Math.atan2(Math.sqrt((e=s*o)*e+(e=c*f-l*s*a)*e),l*f+c*s*a)},ao.geo.graticule=function(){function n(){return{type:"MultiLineString",coordinates:t()}}function t(){return ao.range(Math.ceil(u/d)*d,i,d).map(h).concat(ao.range(Math.ceil(c/y)*y,l,y).map(p)).concat(ao.range(Math.ceil(r/g)*g,e,g).filter(function(n){return xo(n%d)>Uo}).map(f)).concat(ao.range(Math.ceil(a/v)*v,o,v).filter(function(n){return xo(n%y)>Uo}).map(s))}var e,r,i,u,o,a,l,c,f,s,h,p,g=10,v=g,d=90,y=360,m=2.5;return n.lines=function(){return t().map(function(n){return{type:"LineString",coordinates:n}})},n.outline=function(){return{type:"Polygon",coordinates:[h(u).concat(p(l).slice(1),h(i).reverse().slice(1),p(c).reverse().slice(1))]}},n.extent=function(t){return arguments.length?n.majorExtent(t).minorExtent(t):n.minorExtent()},n.majorExtent=function(t){return arguments.length?(u=+t[0][0],i=+t[1][0],c=+t[0][1],l=+t[1][1],u>i&&(t=u,u=i,i=t),c>l&&(t=c,c=l,l=t),n.precision(m)):[[u,c],[i,l]]},n.minorExtent=function(t){return arguments.length?(r=+t[0][0],e=+t[1][0],a=+t[0][1],o=+t[1][1],r>e&&(t=r,r=e,e=t),a>o&&(t=a,a=o,o=t),n.precision(m)):[[r,a],[e,o]]},n.step=function(t){return arguments.length?n.majorStep(t).minorStep(t):n.minorStep()},n.majorStep=function(t){return arguments.length?(d=+t[0],y=+t[1],n):[d,y]},n.minorStep=function(t){return arguments.length?(g=+t[0],v=+t[1],n):[g,v]},n.precision=function(t){return arguments.length?(m=+t,f=ye(a,o,90),s=me(r,e,m),h=ye(c,l,90),p=me(u,i,m),n):m},n.majorExtent([[-180,-90+Uo],[180,90-Uo]]).minorExtent([[-180,-80-Uo],[180,80+Uo]])},ao.geo.greatArc=function(){function n(){return{type:"LineString",coordinates:[t||r.apply(this,arguments),e||i.apply(this,arguments)]}}var t,e,r=Me,i=xe;return n.distance=function(){return ao.geo.distance(t||r.apply(this,arguments),e||i.apply(this,arguments))},n.source=function(e){return arguments.length?(r=e,t="function"==typeof e?null:e,n):r},n.target=function(t){return arguments.length?(i=t,e="function"==typeof t?null:t,n):i},n.precision=function(){return arguments.length?n:0},n},ao.geo.interpolate=function(n,t){return be(n[0]*Yo,n[1]*Yo,t[0]*Yo,t[1]*Yo)},ao.geo.length=function(n){return Ja=0,ao.geo.stream(n,Ga),Ja};var Ja,Ga={sphere:b,point:b,lineStart:_e,lineEnd:b,polygonStart:b,polygonEnd:b},Ka=we(function(n){return Math.sqrt(2/(1+n))},function(n){return 2*Math.asin(n/2)});(ao.geo.azimuthalEqualArea=function(){return oe(Ka)}).raw=Ka;var Qa=we(function(n){var t=Math.acos(n);return t&&t/Math.sin(t)},m);(ao.geo.azimuthalEquidistant=function(){return oe(Qa)}).raw=Qa,(ao.geo.conicConformal=function(){return Vt(Se)}).raw=Se,(ao.geo.conicEquidistant=function(){return Vt(ke)}).raw=ke;var nl=we(function(n){return 1/n},Math.atan);(ao.geo.gnomonic=function(){return oe(nl)}).raw=nl,Ne.invert=function(n,t){return[n,2*Math.atan(Math.exp(t))-Io]},(ao.geo.mercator=function(){return Ee(Ne)}).raw=Ne;var tl=we(function(){return 1},Math.asin);(ao.geo.orthographic=function(){return oe(tl)}).raw=tl;var el=we(function(n){return 1/(1+n)},function(n){return 2*Math.atan(n)});(ao.geo.stereographic=function(){return oe(el)}).raw=el,Ae.invert=function(n,t){return[-t,2*Math.atan(Math.exp(n))-Io]},(ao.geo.transverseMercator=function(){var n=Ee(Ae),t=n.center,e=n.rotate;return n.center=function(n){return n?t([-n[1],n[0]]):(n=t(),[n[1],-n[0]])},n.rotate=function(n){return n?e([n[0],n[1],n.length>2?n[2]+90:90]):(n=e(),[n[0],n[1],n[2]-90])},e([0,0,90])}).raw=Ae,ao.geom={},ao.geom.hull=function(n){function t(n){if(n.length<3)return[];var t,i=En(e),u=En(r),o=n.length,a=[],l=[];for(t=0;o>t;t++)a.push([+i.call(this,n[t],t),+u.call(this,n[t],t),t]);for(a.sort(qe),t=0;o>t;t++)l.push([a[t][0],-a[t][1]]);var c=Le(a),f=Le(l),s=f[0]===c[0],h=f[f.length-1]===c[c.length-1],p=[];for(t=c.length-1;t>=0;--t)p.push(n[a[c[t]][2]]);for(t=+s;t<f.length-h;++t)p.push(n[a[f[t]][2]]);return p}var e=Ce,r=ze;return arguments.length?t(n):(t.x=function(n){return arguments.length?(e=n,t):e},t.y=function(n){return arguments.length?(r=n,t):r},t)},ao.geom.polygon=function(n){return ko(n,rl),n};var rl=ao.geom.polygon.prototype=[];rl.area=function(){for(var n,t=-1,e=this.length,r=this[e-1],i=0;++t<e;)n=r,r=this[t],i+=n[1]*r[0]-n[0]*r[1];return.5*i},rl.centroid=function(n){var t,e,r=-1,i=this.length,u=0,o=0,a=this[i-1];for(arguments.length||(n=-1/(6*this.area()));++r<i;)t=a,a=this[r],e=t[0]*a[1]-a[0]*t[1],u+=(t[0]+a[0])*e,o+=(t[1]+a[1])*e;return[u*n,o*n]},rl.clip=function(n){for(var t,e,r,i,u,o,a=De(n),l=-1,c=this.length-De(this),f=this[c-1];++l<c;){for(t=n.slice(),n.length=0,i=this[l],u=t[(r=t.length-a)-1],e=-1;++e<r;)o=t[e],Te(o,f,i)?(Te(u,f,i)||n.push(Re(u,o,f,i)),n.push(o)):Te(u,f,i)&&n.push(Re(u,o,f,i)),u=o;a&&n.push(n[0]),f=i}return n};var il,ul,ol,al,ll,cl=[],fl=[];Ye.prototype.prepare=function(){for(var n,t=this.edges,e=t.length;e--;)n=t[e].edge,n.b&&n.a||t.splice(e,1);return t.sort(Ve),t.length},tr.prototype={start:function(){return this.edge.l===this.site?this.edge.a:this.edge.b},end:function(){return this.edge.l===this.site?this.edge.b:this.edge.a}},er.prototype={insert:function(n,t){var e,r,i;if(n){if(t.P=n,t.N=n.N,n.N&&(n.N.P=t),n.N=t,n.R){for(n=n.R;n.L;)n=n.L;n.L=t}else n.R=t;e=n}else this._?(n=or(this._),t.P=null,t.N=n,n.P=n.L=t,e=n):(t.P=t.N=null,this._=t,e=null);for(t.L=t.R=null,t.U=e,t.C=!0,n=t;e&&e.C;)r=e.U,e===r.L?(i=r.R,i&&i.C?(e.C=i.C=!1,r.C=!0,n=r):(n===e.R&&(ir(this,e),n=e,e=n.U),e.C=!1,r.C=!0,ur(this,r))):(i=r.L,i&&i.C?(e.C=i.C=!1,r.C=!0,n=r):(n===e.L&&(ur(this,e),n=e,e=n.U),e.C=!1,r.C=!0,ir(this,r))),e=n.U;this._.C=!1},remove:function(n){n.N&&(n.N.P=n.P),n.P&&(n.P.N=n.N),n.N=n.P=null;var t,e,r,i=n.U,u=n.L,o=n.R;if(e=u?o?or(o):u:o,i?i.L===n?i.L=e:i.R=e:this._=e,u&&o?(r=e.C,e.C=n.C,e.L=u,u.U=e,e!==o?(i=e.U,e.U=n.U,n=e.R,i.L=n,e.R=o,o.U=e):(e.U=i,i=e,n=e.R)):(r=n.C,n=e),n&&(n.U=i),!r){if(n&&n.C)return void(n.C=!1);do{if(n===this._)break;if(n===i.L){if(t=i.R,t.C&&(t.C=!1,i.C=!0,ir(this,i),t=i.R),t.L&&t.L.C||t.R&&t.R.C){t.R&&t.R.C||(t.L.C=!1,t.C=!0,ur(this,t),t=i.R),t.C=i.C,i.C=t.R.C=!1,ir(this,i),n=this._;break}}else if(t=i.L,t.C&&(t.C=!1,i.C=!0,ur(this,i),t=i.L),t.L&&t.L.C||t.R&&t.R.C){t.L&&t.L.C||(t.R.C=!1,t.C=!0,ir(this,t),t=i.L),t.C=i.C,i.C=t.L.C=!1,ur(this,i),n=this._;break}t.C=!0,n=i,i=i.U}while(!n.C);n&&(n.C=!1)}}},ao.geom.voronoi=function(n){function t(n){var t=new Array(n.length),r=a[0][0],i=a[0][1],u=a[1][0],o=a[1][1];return ar(e(n),a).cells.forEach(function(e,a){var l=e.edges,c=e.site,f=t[a]=l.length?l.map(function(n){var t=n.start();return[t.x,t.y]}):c.x>=r&&c.x<=u&&c.y>=i&&c.y<=o?[[r,o],[u,o],[u,i],[r,i]]:[];f.point=n[a]}),t}function e(n){return n.map(function(n,t){return{x:Math.round(u(n,t)/Uo)*Uo,y:Math.round(o(n,t)/Uo)*Uo,i:t}})}var r=Ce,i=ze,u=r,o=i,a=sl;return n?t(n):(t.links=function(n){return ar(e(n)).edges.filter(function(n){return n.l&&n.r}).map(function(t){return{source:n[t.l.i],target:n[t.r.i]}})},t.triangles=function(n){var t=[];return ar(e(n)).cells.forEach(function(e,r){for(var i,u,o=e.site,a=e.edges.sort(Ve),l=-1,c=a.length,f=a[c-1].edge,s=f.l===o?f.r:f.l;++l<c;)i=f,u=s,f=a[l].edge,s=f.l===o?f.r:f.l,r<u.i&&r<s.i&&cr(o,u,s)<0&&t.push([n[r],n[u.i],n[s.i]])}),t},t.x=function(n){return arguments.length?(u=En(r=n),t):r},t.y=function(n){return arguments.length?(o=En(i=n),t):i},t.clipExtent=function(n){return arguments.length?(a=null==n?sl:n,t):a===sl?null:a},t.size=function(n){return arguments.length?t.clipExtent(n&&[[0,0],n]):a===sl?null:a&&a[1]},t)};var sl=[[-1e6,-1e6],[1e6,1e6]];ao.geom.delaunay=function(n){return ao.geom.voronoi().triangles(n)},ao.geom.quadtree=function(n,t,e,r,i){function u(n){function u(n,t,e,r,i,u,o,a){if(!isNaN(e)&&!isNaN(r))if(n.leaf){var l=n.x,f=n.y;if(null!=l)if(xo(l-e)+xo(f-r)<.01)c(n,t,e,r,i,u,o,a);else{var s=n.point;n.x=n.y=n.point=null,c(n,s,l,f,i,u,o,a),c(n,t,e,r,i,u,o,a)}else n.x=e,n.y=r,n.point=t}else c(n,t,e,r,i,u,o,a)}function c(n,t,e,r,i,o,a,l){var c=.5*(i+a),f=.5*(o+l),s=e>=c,h=r>=f,p=h<<1|s;n.leaf=!1,n=n.nodes[p]||(n.nodes[p]=hr()),s?i=c:a=c,h?o=f:l=f,u(n,t,e,r,i,o,a,l)}var f,s,h,p,g,v,d,y,m,M=En(a),x=En(l);if(null!=t)v=t,d=e,y=r,m=i;else if(y=m=-(v=d=1/0),s=[],h=[],g=n.length,o)for(p=0;g>p;++p)f=n[p],f.x<v&&(v=f.x),f.y<d&&(d=f.y),f.x>y&&(y=f.x),f.y>m&&(m=f.y),s.push(f.x),h.push(f.y);else for(p=0;g>p;++p){var b=+M(f=n[p],p),_=+x(f,p);v>b&&(v=b),d>_&&(d=_),b>y&&(y=b),_>m&&(m=_),s.push(b),h.push(_)}var w=y-v,S=m-d;w>S?m=d+w:y=v+S;var k=hr();if(k.add=function(n){u(k,n,+M(n,++p),+x(n,p),v,d,y,m)},k.visit=function(n){pr(n,k,v,d,y,m)},k.find=function(n){return gr(k,n[0],n[1],v,d,y,m)},p=-1,null==t){for(;++p<g;)u(k,n[p],s[p],h[p],v,d,y,m);--p}else n.forEach(k.add);return s=h=n=f=null,k}var o,a=Ce,l=ze;return(o=arguments.length)?(a=fr,l=sr,3===o&&(i=e,r=t,e=t=0),u(n)):(u.x=function(n){return arguments.length?(a=n,u):a},u.y=function(n){return arguments.length?(l=n,u):l},u.extent=function(n){return arguments.length?(null==n?t=e=r=i=null:(t=+n[0][0],e=+n[0][1],r=+n[1][0],i=+n[1][1]),u):null==t?null:[[t,e],[r,i]]},u.size=function(n){return arguments.length?(null==n?t=e=r=i=null:(t=e=0,r=+n[0],i=+n[1]),u):null==t?null:[r-t,i-e]},u)},ao.interpolateRgb=vr,ao.interpolateObject=dr,ao.interpolateNumber=yr,ao.interpolateString=mr;var hl=/[-+]?(?:\d+\.?\d*|\.?\d+)(?:[eE][-+]?\d+)?/g,pl=new RegExp(hl.source,"g");ao.interpolate=Mr,ao.interpolators=[function(n,t){var e=typeof t;return("string"===e?ua.has(t.toLowerCase())||/^(#|rgb\(|hsl\()/i.test(t)?vr:mr:t instanceof an?vr:Array.isArray(t)?xr:"object"===e&&isNaN(t)?dr:yr)(n,t)}],ao.interpolateArray=xr;var gl=function(){return m},vl=ao.map({linear:gl,poly:Er,quad:function(){return Sr},cubic:function(){return kr},sin:function(){return Ar},exp:function(){return Cr},circle:function(){return zr},elastic:Lr,back:qr,bounce:function(){return Tr}}),dl=ao.map({"in":m,out:_r,"in-out":wr,"out-in":function(n){return wr(_r(n))}});ao.ease=function(n){var t=n.indexOf("-"),e=t>=0?n.slice(0,t):n,r=t>=0?n.slice(t+1):"in";return e=vl.get(e)||gl,r=dl.get(r)||m,br(r(e.apply(null,lo.call(arguments,1))))},ao.interpolateHcl=Rr,ao.interpolateHsl=Dr,ao.interpolateLab=Pr,ao.interpolateRound=Ur,ao.transform=function(n){var t=fo.createElementNS(ao.ns.prefix.svg,"g");return(ao.transform=function(n){if(null!=n){t.setAttribute("transform",n);var e=t.transform.baseVal.consolidate()}return new jr(e?e.matrix:yl)})(n)},jr.prototype.toString=function(){return"translate("+this.translate+")rotate("+this.rotate+")skewX("+this.skew+")scale("+this.scale+")"};var yl={a:1,b:0,c:0,d:1,e:0,f:0};ao.interpolateTransform=$r,ao.layout={},ao.layout.bundle=function(){return function(n){for(var t=[],e=-1,r=n.length;++e<r;)t.push(Jr(n[e]));return t}},ao.layout.chord=function(){function n(){var n,c,s,h,p,g={},v=[],d=ao.range(u),y=[];for(e=[],r=[],n=0,h=-1;++h<u;){for(c=0,p=-1;++p<u;)c+=i[h][p];v.push(c),y.push(ao.range(u)),n+=c}for(o&&d.sort(function(n,t){return o(v[n],v[t])}),a&&y.forEach(function(n,t){n.sort(function(n,e){return a(i[t][n],i[t][e])})}),n=(Ho-f*u)/n,c=0,h=-1;++h<u;){for(s=c,p=-1;++p<u;){var m=d[h],M=y[m][p],x=i[m][M],b=c,_=c+=x*n;g[m+"-"+M]={index:m,subindex:M,startAngle:b,endAngle:_,value:x}}r[m]={index:m,startAngle:s,endAngle:c,value:v[m]},c+=f}for(h=-1;++h<u;)for(p=h-1;++p<u;){var w=g[h+"-"+p],S=g[p+"-"+h];(w.value||S.value)&&e.push(w.value<S.value?{source:S,target:w}:{source:w,target:S})}l&&t()}function t(){e.sort(function(n,t){return l((n.source.value+n.target.value)/2,(t.source.value+t.target.value)/2)})}var e,r,i,u,o,a,l,c={},f=0;return c.matrix=function(n){return arguments.length?(u=(i=n)&&i.length,e=r=null,c):i},c.padding=function(n){return arguments.length?(f=n,e=r=null,c):f},c.sortGroups=function(n){return arguments.length?(o=n,e=r=null,c):o},c.sortSubgroups=function(n){return arguments.length?(a=n,e=null,c):a},c.sortChords=function(n){return arguments.length?(l=n,e&&t(),c):l},c.chords=function(){return e||n(),e},c.groups=function(){return r||n(),r},c},ao.layout.force=function(){function n(n){return function(t,e,r,i){if(t.point!==n){var u=t.cx-n.x,o=t.cy-n.y,a=i-e,l=u*u+o*o;if(l>a*a/y){if(v>l){var c=t.charge/l;n.px-=u*c,n.py-=o*c}return!0}if(t.point&&l&&v>l){var c=t.pointCharge/l;n.px-=u*c,n.py-=o*c}}return!t.charge}}function t(n){n.px=ao.event.x,n.py=ao.event.y,l.resume()}var e,r,i,u,o,a,l={},c=ao.dispatch("start","tick","end"),f=[1,1],s=.9,h=ml,p=Ml,g=-30,v=xl,d=.1,y=.64,M=[],x=[];return l.tick=function(){if((i*=.99)<.005)return e=null,c.end({type:"end",alpha:i=0}),!0;var t,r,l,h,p,v,y,m,b,_=M.length,w=x.length;for(r=0;w>r;++r)l=x[r],h=l.source,p=l.target,m=p.x-h.x,b=p.y-h.y,(v=m*m+b*b)&&(v=i*o[r]*((v=Math.sqrt(v))-u[r])/v,m*=v,b*=v,p.x-=m*(y=h.weight+p.weight?h.weight/(h.weight+p.weight):.5),p.y-=b*y,h.x+=m*(y=1-y),h.y+=b*y);if((y=i*d)&&(m=f[0]/2,b=f[1]/2,r=-1,y))for(;++r<_;)l=M[r],l.x+=(m-l.x)*y,l.y+=(b-l.y)*y;if(g)for(ri(t=ao.geom.quadtree(M),i,a),r=-1;++r<_;)(l=M[r]).fixed||t.visit(n(l));for(r=-1;++r<_;)l=M[r],l.fixed?(l.x=l.px,l.y=l.py):(l.x-=(l.px-(l.px=l.x))*s,l.y-=(l.py-(l.py=l.y))*s);c.tick({type:"tick",alpha:i})},l.nodes=function(n){return arguments.length?(M=n,l):M},l.links=function(n){return arguments.length?(x=n,l):x},l.size=function(n){return arguments.length?(f=n,l):f},l.linkDistance=function(n){return arguments.length?(h="function"==typeof n?n:+n,l):h},l.distance=l.linkDistance,l.linkStrength=function(n){return arguments.length?(p="function"==typeof n?n:+n,l):p},l.friction=function(n){return arguments.length?(s=+n,l):s},l.charge=function(n){return arguments.length?(g="function"==typeof n?n:+n,l):g},l.chargeDistance=function(n){return arguments.length?(v=n*n,l):Math.sqrt(v)},l.gravity=function(n){return arguments.length?(d=+n,l):d},l.theta=function(n){return arguments.length?(y=n*n,l):Math.sqrt(y)},l.alpha=function(n){return arguments.length?(n=+n,i?n>0?i=n:(e.c=null,e.t=NaN,e=null,c.end({type:"end",alpha:i=0})):n>0&&(c.start({type:"start",alpha:i=n}),e=qn(l.tick)),l):i},l.start=function(){function n(n,r){if(!e){for(e=new Array(i),l=0;i>l;++l)e[l]=[];for(l=0;c>l;++l){var u=x[l];e[u.source.index].push(u.target),e[u.target.index].push(u.source)}}for(var o,a=e[t],l=-1,f=a.length;++l<f;)if(!isNaN(o=a[l][n]))return o;return Math.random()*r}var t,e,r,i=M.length,c=x.length,s=f[0],v=f[1];for(t=0;i>t;++t)(r=M[t]).index=t,r.weight=0;for(t=0;c>t;++t)r=x[t],"number"==typeof r.source&&(r.source=M[r.source]),"number"==typeof r.target&&(r.target=M[r.target]),++r.source.weight,++r.target.weight;for(t=0;i>t;++t)r=M[t],isNaN(r.x)&&(r.x=n("x",s)),isNaN(r.y)&&(r.y=n("y",v)),isNaN(r.px)&&(r.px=r.x),isNaN(r.py)&&(r.py=r.y);if(u=[],"function"==typeof h)for(t=0;c>t;++t)u[t]=+h.call(this,x[t],t);else for(t=0;c>t;++t)u[t]=h;if(o=[],"function"==typeof p)for(t=0;c>t;++t)o[t]=+p.call(this,x[t],t);else for(t=0;c>t;++t)o[t]=p;if(a=[],"function"==typeof g)for(t=0;i>t;++t)a[t]=+g.call(this,M[t],t);else for(t=0;i>t;++t)a[t]=g;return l.resume()},l.resume=function(){return l.alpha(.1)},l.stop=function(){return l.alpha(0)},l.drag=function(){return r||(r=ao.behavior.drag().origin(m).on("dragstart.force",Qr).on("drag.force",t).on("dragend.force",ni)),arguments.length?void this.on("mouseover.force",ti).on("mouseout.force",ei).call(r):r},ao.rebind(l,c,"on")};var ml=20,Ml=1,xl=1/0;ao.layout.hierarchy=function(){function n(i){var u,o=[i],a=[];for(i.depth=0;null!=(u=o.pop());)if(a.push(u),(c=e.call(n,u,u.depth))&&(l=c.length)){for(var l,c,f;--l>=0;)o.push(f=c[l]),f.parent=u,f.depth=u.depth+1;r&&(u.value=0),u.children=c}else r&&(u.value=+r.call(n,u,u.depth)||0),delete u.children;return oi(i,function(n){var e,i;t&&(e=n.children)&&e.sort(t),r&&(i=n.parent)&&(i.value+=n.value)}),a}var t=ci,e=ai,r=li;return n.sort=function(e){return arguments.length?(t=e,n):t},n.children=function(t){return arguments.length?(e=t,n):e},n.value=function(t){return arguments.length?(r=t,n):r},n.revalue=function(t){return r&&(ui(t,function(n){n.children&&(n.value=0)}),oi(t,function(t){var e;t.children||(t.value=+r.call(n,t,t.depth)||0),(e=t.parent)&&(e.value+=t.value)})),t},n},ao.layout.partition=function(){function n(t,e,r,i){var u=t.children;if(t.x=e,t.y=t.depth*i,t.dx=r,t.dy=i,u&&(o=u.length)){var o,a,l,c=-1;for(r=t.value?r/t.value:0;++c<o;)n(a=u[c],e,l=a.value*r,i),e+=l}}function t(n){var e=n.children,r=0;if(e&&(i=e.length))for(var i,u=-1;++u<i;)r=Math.max(r,t(e[u]));return 1+r}function e(e,u){var o=r.call(this,e,u);return n(o[0],0,i[0],i[1]/t(o[0])),o}var r=ao.layout.hierarchy(),i=[1,1];return e.size=function(n){return arguments.length?(i=n,e):i},ii(e,r)},ao.layout.pie=function(){function n(o){var a,l=o.length,c=o.map(function(e,r){return+t.call(n,e,r)}),f=+("function"==typeof r?r.apply(this,arguments):r),s=("function"==typeof i?i.apply(this,arguments):i)-f,h=Math.min(Math.abs(s)/l,+("function"==typeof u?u.apply(this,arguments):u)),p=h*(0>s?-1:1),g=ao.sum(c),v=g?(s-l*p)/g:0,d=ao.range(l),y=[];return null!=e&&d.sort(e===bl?function(n,t){return c[t]-c[n]}:function(n,t){return e(o[n],o[t])}),d.forEach(function(n){y[n]={data:o[n],value:a=c[n],startAngle:f,endAngle:f+=a*v+p,padAngle:h}}),y}var t=Number,e=bl,r=0,i=Ho,u=0;return n.value=function(e){return arguments.length?(t=e,n):t},n.sort=function(t){return arguments.length?(e=t,n):e},n.startAngle=function(t){return arguments.length?(r=t,n):r},n.endAngle=function(t){return arguments.length?(i=t,n):i},n.padAngle=function(t){return arguments.length?(u=t,n):u},n};var bl={};ao.layout.stack=function(){function n(a,l){if(!(h=a.length))return a;var c=a.map(function(e,r){return t.call(n,e,r)}),f=c.map(function(t){return t.map(function(t,e){return[u.call(n,t,e),o.call(n,t,e)]})}),s=e.call(n,f,l);c=ao.permute(c,s),f=ao.permute(f,s);var h,p,g,v,d=r.call(n,f,l),y=c[0].length;for(g=0;y>g;++g)for(i.call(n,c[0][g],v=d[g],f[0][g][1]),p=1;h>p;++p)i.call(n,c[p][g],v+=f[p-1][g][1],f[p][g][1]);return a}var t=m,e=gi,r=vi,i=pi,u=si,o=hi;return n.values=function(e){return arguments.length?(t=e,n):t},n.order=function(t){return arguments.length?(e="function"==typeof t?t:_l.get(t)||gi,n):e},n.offset=function(t){return arguments.length?(r="function"==typeof t?t:wl.get(t)||vi,n):r},n.x=function(t){return arguments.length?(u=t,n):u},n.y=function(t){return arguments.length?(o=t,n):o},n.out=function(t){return arguments.length?(i=t,n):i},n};var _l=ao.map({"inside-out":function(n){var t,e,r=n.length,i=n.map(di),u=n.map(yi),o=ao.range(r).sort(function(n,t){return i[n]-i[t]}),a=0,l=0,c=[],f=[];for(t=0;r>t;++t)e=o[t],l>a?(a+=u[e],c.push(e)):(l+=u[e],f.push(e));return f.reverse().concat(c)},reverse:function(n){return ao.range(n.length).reverse()},"default":gi}),wl=ao.map({silhouette:function(n){var t,e,r,i=n.length,u=n[0].length,o=[],a=0,l=[];for(e=0;u>e;++e){for(t=0,r=0;i>t;t++)r+=n[t][e][1];r>a&&(a=r),o.push(r)}for(e=0;u>e;++e)l[e]=(a-o[e])/2;return l},wiggle:function(n){var t,e,r,i,u,o,a,l,c,f=n.length,s=n[0],h=s.length,p=[];for(p[0]=l=c=0,e=1;h>e;++e){for(t=0,i=0;f>t;++t)i+=n[t][e][1];for(t=0,u=0,a=s[e][0]-s[e-1][0];f>t;++t){for(r=0,o=(n[t][e][1]-n[t][e-1][1])/(2*a);t>r;++r)o+=(n[r][e][1]-n[r][e-1][1])/a;u+=o*n[t][e][1]}p[e]=l-=i?u/i*a:0,c>l&&(c=l)}for(e=0;h>e;++e)p[e]-=c;return p},expand:function(n){var t,e,r,i=n.length,u=n[0].length,o=1/i,a=[];for(e=0;u>e;++e){for(t=0,r=0;i>t;t++)r+=n[t][e][1];if(r)for(t=0;i>t;t++)n[t][e][1]/=r;else for(t=0;i>t;t++)n[t][e][1]=o}for(e=0;u>e;++e)a[e]=0;return a},zero:vi});ao.layout.histogram=function(){function n(n,u){for(var o,a,l=[],c=n.map(e,this),f=r.call(this,c,u),s=i.call(this,f,c,u),u=-1,h=c.length,p=s.length-1,g=t?1:1/h;++u<p;)o=l[u]=[],o.dx=s[u+1]-(o.x=s[u]),o.y=0;if(p>0)for(u=-1;++u<h;)a=c[u],a>=f[0]&&a<=f[1]&&(o=l[ao.bisect(s,a,1,p)-1],o.y+=g,o.push(n[u]));return l}var t=!0,e=Number,r=bi,i=Mi;return n.value=function(t){return arguments.length?(e=t,n):e},n.range=function(t){return arguments.length?(r=En(t),n):r},n.bins=function(t){return arguments.length?(i="number"==typeof t?function(n){return xi(n,t)}:En(t),n):i},n.frequency=function(e){return arguments.length?(t=!!e,n):t},n},ao.layout.pack=function(){function n(n,u){var o=e.call(this,n,u),a=o[0],l=i[0],c=i[1],f=null==t?Math.sqrt:"function"==typeof t?t:function(){return t};if(a.x=a.y=0,oi(a,function(n){n.r=+f(n.value)}),oi(a,Ni),r){var s=r*(t?1:Math.max(2*a.r/l,2*a.r/c))/2;oi(a,function(n){n.r+=s}),oi(a,Ni),oi(a,function(n){n.r-=s})}return Ci(a,l/2,c/2,t?1:1/Math.max(2*a.r/l,2*a.r/c)),o}var t,e=ao.layout.hierarchy().sort(_i),r=0,i=[1,1];return n.size=function(t){return arguments.length?(i=t,n):i},n.radius=function(e){return arguments.length?(t=null==e||"function"==typeof e?e:+e,n):t},n.padding=function(t){return arguments.length?(r=+t,n):r},ii(n,e)},ao.layout.tree=function(){function n(n,i){var f=o.call(this,n,i),s=f[0],h=t(s);if(oi(h,e),h.parent.m=-h.z,ui(h,r),c)ui(s,u);else{var p=s,g=s,v=s;ui(s,function(n){n.x<p.x&&(p=n),n.x>g.x&&(g=n),n.depth>v.depth&&(v=n)});var d=a(p,g)/2-p.x,y=l[0]/(g.x+a(g,p)/2+d),m=l[1]/(v.depth||1);ui(s,function(n){n.x=(n.x+d)*y,n.y=n.depth*m})}return f}function t(n){for(var t,e={A:null,children:[n]},r=[e];null!=(t=r.pop());)for(var i,u=t.children,o=0,a=u.length;a>o;++o)r.push((u[o]=i={_:u[o],parent:t,children:(i=u[o].children)&&i.slice()||[],A:null,a:null,z:0,m:0,c:0,s:0,t:null,i:o}).a=i);return e.children[0]}function e(n){var t=n.children,e=n.parent.children,r=n.i?e[n.i-1]:null;if(t.length){Di(n);var u=(t[0].z+t[t.length-1].z)/2;r?(n.z=r.z+a(n._,r._),n.m=n.z-u):n.z=u}else r&&(n.z=r.z+a(n._,r._));n.parent.A=i(n,r,n.parent.A||e[0])}function r(n){n._.x=n.z+n.parent.m,n.m+=n.parent.m}function i(n,t,e){if(t){for(var r,i=n,u=n,o=t,l=i.parent.children[0],c=i.m,f=u.m,s=o.m,h=l.m;o=Ti(o),i=qi(i),o&&i;)l=qi(l),u=Ti(u),u.a=n,r=o.z+s-i.z-c+a(o._,i._),r>0&&(Ri(Pi(o,n,e),n,r),c+=r,f+=r),s+=o.m,c+=i.m,h+=l.m,f+=u.m;o&&!Ti(u)&&(u.t=o,u.m+=s-f),i&&!qi(l)&&(l.t=i,l.m+=c-h,e=n)}return e}function u(n){n.x*=l[0],n.y=n.depth*l[1]}var o=ao.layout.hierarchy().sort(null).value(null),a=Li,l=[1,1],c=null;return n.separation=function(t){return arguments.length?(a=t,n):a},n.size=function(t){return arguments.length?(c=null==(l=t)?u:null,n):c?null:l},n.nodeSize=function(t){return arguments.length?(c=null==(l=t)?null:u,n):c?l:null},ii(n,o)},ao.layout.cluster=function(){function n(n,u){var o,a=t.call(this,n,u),l=a[0],c=0;oi(l,function(n){var t=n.children;t&&t.length?(n.x=ji(t),n.y=Ui(t)):(n.x=o?c+=e(n,o):0,n.y=0,o=n)});var f=Fi(l),s=Hi(l),h=f.x-e(f,s)/2,p=s.x+e(s,f)/2;return oi(l,i?function(n){n.x=(n.x-l.x)*r[0],n.y=(l.y-n.y)*r[1]}:function(n){n.x=(n.x-h)/(p-h)*r[0],n.y=(1-(l.y?n.y/l.y:1))*r[1]}),a}var t=ao.layout.hierarchy().sort(null).value(null),e=Li,r=[1,1],i=!1;return n.separation=function(t){return arguments.length?(e=t,n):e},n.size=function(t){return arguments.length?(i=null==(r=t),n):i?null:r},n.nodeSize=function(t){return arguments.length?(i=null!=(r=t),n):i?r:null},ii(n,t)},ao.layout.treemap=function(){function n(n,t){for(var e,r,i=-1,u=n.length;++i<u;)r=(e=n[i]).value*(0>t?0:t),e.area=isNaN(r)||0>=r?0:r}function t(e){var u=e.children;if(u&&u.length){var o,a,l,c=s(e),f=[],h=u.slice(),g=1/0,v="slice"===p?c.dx:"dice"===p?c.dy:"slice-dice"===p?1&e.depth?c.dy:c.dx:Math.min(c.dx,c.dy);for(n(h,c.dx*c.dy/e.value),f.area=0;(l=h.length)>0;)f.push(o=h[l-1]),f.area+=o.area,"squarify"!==p||(a=r(f,v))<=g?(h.pop(),g=a):(f.area-=f.pop().area,i(f,v,c,!1),v=Math.min(c.dx,c.dy),f.length=f.area=0,g=1/0);f.length&&(i(f,v,c,!0),f.length=f.area=0),u.forEach(t)}}function e(t){var r=t.children;if(r&&r.length){var u,o=s(t),a=r.slice(),l=[];for(n(a,o.dx*o.dy/t.value),l.area=0;u=a.pop();)l.push(u),l.area+=u.area,null!=u.z&&(i(l,u.z?o.dx:o.dy,o,!a.length),l.length=l.area=0);r.forEach(e)}}function r(n,t){for(var e,r=n.area,i=0,u=1/0,o=-1,a=n.length;++o<a;)(e=n[o].area)&&(u>e&&(u=e),e>i&&(i=e));return r*=r,t*=t,r?Math.max(t*i*g/r,r/(t*u*g)):1/0}function i(n,t,e,r){var i,u=-1,o=n.length,a=e.x,c=e.y,f=t?l(n.area/t):0;
if(t==e.dx){for((r||f>e.dy)&&(f=e.dy);++u<o;)i=n[u],i.x=a,i.y=c,i.dy=f,a+=i.dx=Math.min(e.x+e.dx-a,f?l(i.area/f):0);i.z=!0,i.dx+=e.x+e.dx-a,e.y+=f,e.dy-=f}else{for((r||f>e.dx)&&(f=e.dx);++u<o;)i=n[u],i.x=a,i.y=c,i.dx=f,c+=i.dy=Math.min(e.y+e.dy-c,f?l(i.area/f):0);i.z=!1,i.dy+=e.y+e.dy-c,e.x+=f,e.dx-=f}}function u(r){var i=o||a(r),u=i[0];return u.x=u.y=0,u.value?(u.dx=c[0],u.dy=c[1]):u.dx=u.dy=0,o&&a.revalue(u),n([u],u.dx*u.dy/u.value),(o?e:t)(u),h&&(o=i),i}var o,a=ao.layout.hierarchy(),l=Math.round,c=[1,1],f=null,s=Oi,h=!1,p="squarify",g=.5*(1+Math.sqrt(5));return u.size=function(n){return arguments.length?(c=n,u):c},u.padding=function(n){function t(t){var e=n.call(u,t,t.depth);return null==e?Oi(t):Ii(t,"number"==typeof e?[e,e,e,e]:e)}function e(t){return Ii(t,n)}if(!arguments.length)return f;var r;return s=null==(f=n)?Oi:"function"==(r=typeof n)?t:"number"===r?(n=[n,n,n,n],e):e,u},u.round=function(n){return arguments.length?(l=n?Math.round:Number,u):l!=Number},u.sticky=function(n){return arguments.length?(h=n,o=null,u):h},u.ratio=function(n){return arguments.length?(g=n,u):g},u.mode=function(n){return arguments.length?(p=n+"",u):p},ii(u,a)},ao.random={normal:function(n,t){var e=arguments.length;return 2>e&&(t=1),1>e&&(n=0),function(){var e,r,i;do e=2*Math.random()-1,r=2*Math.random()-1,i=e*e+r*r;while(!i||i>1);return n+t*e*Math.sqrt(-2*Math.log(i)/i)}},logNormal:function(){var n=ao.random.normal.apply(ao,arguments);return function(){return Math.exp(n())}},bates:function(n){var t=ao.random.irwinHall(n);return function(){return t()/n}},irwinHall:function(n){return function(){for(var t=0,e=0;n>e;e++)t+=Math.random();return t}}},ao.scale={};var Sl={floor:m,ceil:m};ao.scale.linear=function(){return Wi([0,1],[0,1],Mr,!1)};var kl={s:1,g:1,p:1,r:1,e:1};ao.scale.log=function(){return ru(ao.scale.linear().domain([0,1]),10,!0,[1,10])};var Nl=ao.format(".0e"),El={floor:function(n){return-Math.ceil(-n)},ceil:function(n){return-Math.floor(-n)}};ao.scale.pow=function(){return iu(ao.scale.linear(),1,[0,1])},ao.scale.sqrt=function(){return ao.scale.pow().exponent(.5)},ao.scale.ordinal=function(){return ou([],{t:"range",a:[[]]})},ao.scale.category10=function(){return ao.scale.ordinal().range(Al)},ao.scale.category20=function(){return ao.scale.ordinal().range(Cl)},ao.scale.category20b=function(){return ao.scale.ordinal().range(zl)},ao.scale.category20c=function(){return ao.scale.ordinal().range(Ll)};var Al=[2062260,16744206,2924588,14034728,9725885,9197131,14907330,8355711,12369186,1556175].map(xn),Cl=[2062260,11454440,16744206,16759672,2924588,10018698,14034728,16750742,9725885,12955861,9197131,12885140,14907330,16234194,8355711,13092807,12369186,14408589,1556175,10410725].map(xn),zl=[3750777,5395619,7040719,10264286,6519097,9216594,11915115,13556636,9202993,12426809,15186514,15190932,8666169,11356490,14049643,15177372,8077683,10834324,13528509,14589654].map(xn),Ll=[3244733,7057110,10406625,13032431,15095053,16616764,16625259,16634018,3253076,7652470,10607003,13101504,7695281,10394312,12369372,14342891,6513507,9868950,12434877,14277081].map(xn);ao.scale.quantile=function(){return au([],[])},ao.scale.quantize=function(){return lu(0,1,[0,1])},ao.scale.threshold=function(){return cu([.5],[0,1])},ao.scale.identity=function(){return fu([0,1])},ao.svg={},ao.svg.arc=function(){function n(){var n=Math.max(0,+e.apply(this,arguments)),c=Math.max(0,+r.apply(this,arguments)),f=o.apply(this,arguments)-Io,s=a.apply(this,arguments)-Io,h=Math.abs(s-f),p=f>s?0:1;if(n>c&&(g=c,c=n,n=g),h>=Oo)return t(c,p)+(n?t(n,1-p):"")+"Z";var g,v,d,y,m,M,x,b,_,w,S,k,N=0,E=0,A=[];if((y=(+l.apply(this,arguments)||0)/2)&&(d=u===ql?Math.sqrt(n*n+c*c):+u.apply(this,arguments),p||(E*=-1),c&&(E=tn(d/c*Math.sin(y))),n&&(N=tn(d/n*Math.sin(y)))),c){m=c*Math.cos(f+E),M=c*Math.sin(f+E),x=c*Math.cos(s-E),b=c*Math.sin(s-E);var C=Math.abs(s-f-2*E)<=Fo?0:1;if(E&&yu(m,M,x,b)===p^C){var z=(f+s)/2;m=c*Math.cos(z),M=c*Math.sin(z),x=b=null}}else m=M=0;if(n){_=n*Math.cos(s-N),w=n*Math.sin(s-N),S=n*Math.cos(f+N),k=n*Math.sin(f+N);var L=Math.abs(f-s+2*N)<=Fo?0:1;if(N&&yu(_,w,S,k)===1-p^L){var q=(f+s)/2;_=n*Math.cos(q),w=n*Math.sin(q),S=k=null}}else _=w=0;if(h>Uo&&(g=Math.min(Math.abs(c-n)/2,+i.apply(this,arguments)))>.001){v=c>n^p?0:1;var T=g,R=g;if(Fo>h){var D=null==S?[_,w]:null==x?[m,M]:Re([m,M],[S,k],[x,b],[_,w]),P=m-D[0],U=M-D[1],j=x-D[0],F=b-D[1],H=1/Math.sin(Math.acos((P*j+U*F)/(Math.sqrt(P*P+U*U)*Math.sqrt(j*j+F*F)))/2),O=Math.sqrt(D[0]*D[0]+D[1]*D[1]);R=Math.min(g,(n-O)/(H-1)),T=Math.min(g,(c-O)/(H+1))}if(null!=x){var I=mu(null==S?[_,w]:[S,k],[m,M],c,T,p),Y=mu([x,b],[_,w],c,T,p);g===T?A.push("M",I[0],"A",T,",",T," 0 0,",v," ",I[1],"A",c,",",c," 0 ",1-p^yu(I[1][0],I[1][1],Y[1][0],Y[1][1]),",",p," ",Y[1],"A",T,",",T," 0 0,",v," ",Y[0]):A.push("M",I[0],"A",T,",",T," 0 1,",v," ",Y[0])}else A.push("M",m,",",M);if(null!=S){var Z=mu([m,M],[S,k],n,-R,p),V=mu([_,w],null==x?[m,M]:[x,b],n,-R,p);g===R?A.push("L",V[0],"A",R,",",R," 0 0,",v," ",V[1],"A",n,",",n," 0 ",p^yu(V[1][0],V[1][1],Z[1][0],Z[1][1]),",",1-p," ",Z[1],"A",R,",",R," 0 0,",v," ",Z[0]):A.push("L",V[0],"A",R,",",R," 0 0,",v," ",Z[0])}else A.push("L",_,",",w)}else A.push("M",m,",",M),null!=x&&A.push("A",c,",",c," 0 ",C,",",p," ",x,",",b),A.push("L",_,",",w),null!=S&&A.push("A",n,",",n," 0 ",L,",",1-p," ",S,",",k);return A.push("Z"),A.join("")}function t(n,t){return"M0,"+n+"A"+n+","+n+" 0 1,"+t+" 0,"+-n+"A"+n+","+n+" 0 1,"+t+" 0,"+n}var e=hu,r=pu,i=su,u=ql,o=gu,a=vu,l=du;return n.innerRadius=function(t){return arguments.length?(e=En(t),n):e},n.outerRadius=function(t){return arguments.length?(r=En(t),n):r},n.cornerRadius=function(t){return arguments.length?(i=En(t),n):i},n.padRadius=function(t){return arguments.length?(u=t==ql?ql:En(t),n):u},n.startAngle=function(t){return arguments.length?(o=En(t),n):o},n.endAngle=function(t){return arguments.length?(a=En(t),n):a},n.padAngle=function(t){return arguments.length?(l=En(t),n):l},n.centroid=function(){var n=(+e.apply(this,arguments)+ +r.apply(this,arguments))/2,t=(+o.apply(this,arguments)+ +a.apply(this,arguments))/2-Io;return[Math.cos(t)*n,Math.sin(t)*n]},n};var ql="auto";ao.svg.line=function(){return Mu(m)};var Tl=ao.map({linear:xu,"linear-closed":bu,step:_u,"step-before":wu,"step-after":Su,basis:zu,"basis-open":Lu,"basis-closed":qu,bundle:Tu,cardinal:Eu,"cardinal-open":ku,"cardinal-closed":Nu,monotone:Fu});Tl.forEach(function(n,t){t.key=n,t.closed=/-closed$/.test(n)});var Rl=[0,2/3,1/3,0],Dl=[0,1/3,2/3,0],Pl=[0,1/6,2/3,1/6];ao.svg.line.radial=function(){var n=Mu(Hu);return n.radius=n.x,delete n.x,n.angle=n.y,delete n.y,n},wu.reverse=Su,Su.reverse=wu,ao.svg.area=function(){return Ou(m)},ao.svg.area.radial=function(){var n=Ou(Hu);return n.radius=n.x,delete n.x,n.innerRadius=n.x0,delete n.x0,n.outerRadius=n.x1,delete n.x1,n.angle=n.y,delete n.y,n.startAngle=n.y0,delete n.y0,n.endAngle=n.y1,delete n.y1,n},ao.svg.chord=function(){function n(n,a){var l=t(this,u,n,a),c=t(this,o,n,a);return"M"+l.p0+r(l.r,l.p1,l.a1-l.a0)+(e(l,c)?i(l.r,l.p1,l.r,l.p0):i(l.r,l.p1,c.r,c.p0)+r(c.r,c.p1,c.a1-c.a0)+i(c.r,c.p1,l.r,l.p0))+"Z"}function t(n,t,e,r){var i=t.call(n,e,r),u=a.call(n,i,r),o=l.call(n,i,r)-Io,f=c.call(n,i,r)-Io;return{r:u,a0:o,a1:f,p0:[u*Math.cos(o),u*Math.sin(o)],p1:[u*Math.cos(f),u*Math.sin(f)]}}function e(n,t){return n.a0==t.a0&&n.a1==t.a1}function r(n,t,e){return"A"+n+","+n+" 0 "+ +(e>Fo)+",1 "+t}function i(n,t,e,r){return"Q 0,0 "+r}var u=Me,o=xe,a=Iu,l=gu,c=vu;return n.radius=function(t){return arguments.length?(a=En(t),n):a},n.source=function(t){return arguments.length?(u=En(t),n):u},n.target=function(t){return arguments.length?(o=En(t),n):o},n.startAngle=function(t){return arguments.length?(l=En(t),n):l},n.endAngle=function(t){return arguments.length?(c=En(t),n):c},n},ao.svg.diagonal=function(){function n(n,i){var u=t.call(this,n,i),o=e.call(this,n,i),a=(u.y+o.y)/2,l=[u,{x:u.x,y:a},{x:o.x,y:a},o];return l=l.map(r),"M"+l[0]+"C"+l[1]+" "+l[2]+" "+l[3]}var t=Me,e=xe,r=Yu;return n.source=function(e){return arguments.length?(t=En(e),n):t},n.target=function(t){return arguments.length?(e=En(t),n):e},n.projection=function(t){return arguments.length?(r=t,n):r},n},ao.svg.diagonal.radial=function(){var n=ao.svg.diagonal(),t=Yu,e=n.projection;return n.projection=function(n){return arguments.length?e(Zu(t=n)):t},n},ao.svg.symbol=function(){function n(n,r){return(Ul.get(t.call(this,n,r))||$u)(e.call(this,n,r))}var t=Xu,e=Vu;return n.type=function(e){return arguments.length?(t=En(e),n):t},n.size=function(t){return arguments.length?(e=En(t),n):e},n};var Ul=ao.map({circle:$u,cross:function(n){var t=Math.sqrt(n/5)/2;return"M"+-3*t+","+-t+"H"+-t+"V"+-3*t+"H"+t+"V"+-t+"H"+3*t+"V"+t+"H"+t+"V"+3*t+"H"+-t+"V"+t+"H"+-3*t+"Z"},diamond:function(n){var t=Math.sqrt(n/(2*Fl)),e=t*Fl;return"M0,"+-t+"L"+e+",0 0,"+t+" "+-e+",0Z"},square:function(n){var t=Math.sqrt(n)/2;return"M"+-t+","+-t+"L"+t+","+-t+" "+t+","+t+" "+-t+","+t+"Z"},"triangle-down":function(n){var t=Math.sqrt(n/jl),e=t*jl/2;return"M0,"+e+"L"+t+","+-e+" "+-t+","+-e+"Z"},"triangle-up":function(n){var t=Math.sqrt(n/jl),e=t*jl/2;return"M0,"+-e+"L"+t+","+e+" "+-t+","+e+"Z"}});ao.svg.symbolTypes=Ul.keys();var jl=Math.sqrt(3),Fl=Math.tan(30*Yo);Co.transition=function(n){for(var t,e,r=Hl||++Zl,i=Ku(n),u=[],o=Ol||{time:Date.now(),ease:Nr,delay:0,duration:250},a=-1,l=this.length;++a<l;){u.push(t=[]);for(var c=this[a],f=-1,s=c.length;++f<s;)(e=c[f])&&Qu(e,f,i,r,o),t.push(e)}return Wu(u,i,r)},Co.interrupt=function(n){return this.each(null==n?Il:Bu(Ku(n)))};var Hl,Ol,Il=Bu(Ku()),Yl=[],Zl=0;Yl.call=Co.call,Yl.empty=Co.empty,Yl.node=Co.node,Yl.size=Co.size,ao.transition=function(n,t){return n&&n.transition?Hl?n.transition(t):n:ao.selection().transition(n)},ao.transition.prototype=Yl,Yl.select=function(n){var t,e,r,i=this.id,u=this.namespace,o=[];n=A(n);for(var a=-1,l=this.length;++a<l;){o.push(t=[]);for(var c=this[a],f=-1,s=c.length;++f<s;)(r=c[f])&&(e=n.call(r,r.__data__,f,a))?("__data__"in r&&(e.__data__=r.__data__),Qu(e,f,u,i,r[u][i]),t.push(e)):t.push(null)}return Wu(o,u,i)},Yl.selectAll=function(n){var t,e,r,i,u,o=this.id,a=this.namespace,l=[];n=C(n);for(var c=-1,f=this.length;++c<f;)for(var s=this[c],h=-1,p=s.length;++h<p;)if(r=s[h]){u=r[a][o],e=n.call(r,r.__data__,h,c),l.push(t=[]);for(var g=-1,v=e.length;++g<v;)(i=e[g])&&Qu(i,g,a,o,u),t.push(i)}return Wu(l,a,o)},Yl.filter=function(n){var t,e,r,i=[];"function"!=typeof n&&(n=O(n));for(var u=0,o=this.length;o>u;u++){i.push(t=[]);for(var e=this[u],a=0,l=e.length;l>a;a++)(r=e[a])&&n.call(r,r.__data__,a,u)&&t.push(r)}return Wu(i,this.namespace,this.id)},Yl.tween=function(n,t){var e=this.id,r=this.namespace;return arguments.length<2?this.node()[r][e].tween.get(n):Y(this,null==t?function(t){t[r][e].tween.remove(n)}:function(i){i[r][e].tween.set(n,t)})},Yl.attr=function(n,t){function e(){this.removeAttribute(a)}function r(){this.removeAttributeNS(a.space,a.local)}function i(n){return null==n?e:(n+="",function(){var t,e=this.getAttribute(a);return e!==n&&(t=o(e,n),function(n){this.setAttribute(a,t(n))})})}function u(n){return null==n?r:(n+="",function(){var t,e=this.getAttributeNS(a.space,a.local);return e!==n&&(t=o(e,n),function(n){this.setAttributeNS(a.space,a.local,t(n))})})}if(arguments.length<2){for(t in n)this.attr(t,n[t]);return this}var o="transform"==n?$r:Mr,a=ao.ns.qualify(n);return Ju(this,"attr."+n,t,a.local?u:i)},Yl.attrTween=function(n,t){function e(n,e){var r=t.call(this,n,e,this.getAttribute(i));return r&&function(n){this.setAttribute(i,r(n))}}function r(n,e){var r=t.call(this,n,e,this.getAttributeNS(i.space,i.local));return r&&function(n){this.setAttributeNS(i.space,i.local,r(n))}}var i=ao.ns.qualify(n);return this.tween("attr."+n,i.local?r:e)},Yl.style=function(n,e,r){function i(){this.style.removeProperty(n)}function u(e){return null==e?i:(e+="",function(){var i,u=t(this).getComputedStyle(this,null).getPropertyValue(n);return u!==e&&(i=Mr(u,e),function(t){this.style.setProperty(n,i(t),r)})})}var o=arguments.length;if(3>o){if("string"!=typeof n){2>o&&(e="");for(r in n)this.style(r,n[r],e);return this}r=""}return Ju(this,"style."+n,e,u)},Yl.styleTween=function(n,e,r){function i(i,u){var o=e.call(this,i,u,t(this).getComputedStyle(this,null).getPropertyValue(n));return o&&function(t){this.style.setProperty(n,o(t),r)}}return arguments.length<3&&(r=""),this.tween("style."+n,i)},Yl.text=function(n){return Ju(this,"text",n,Gu)},Yl.remove=function(){var n=this.namespace;return this.each("end.transition",function(){var t;this[n].count<2&&(t=this.parentNode)&&t.removeChild(this)})},Yl.ease=function(n){var t=this.id,e=this.namespace;return arguments.length<1?this.node()[e][t].ease:("function"!=typeof n&&(n=ao.ease.apply(ao,arguments)),Y(this,function(r){r[e][t].ease=n}))},Yl.delay=function(n){var t=this.id,e=this.namespace;return arguments.length<1?this.node()[e][t].delay:Y(this,"function"==typeof n?function(r,i,u){r[e][t].delay=+n.call(r,r.__data__,i,u)}:(n=+n,function(r){r[e][t].delay=n}))},Yl.duration=function(n){var t=this.id,e=this.namespace;return arguments.length<1?this.node()[e][t].duration:Y(this,"function"==typeof n?function(r,i,u){r[e][t].duration=Math.max(1,n.call(r,r.__data__,i,u))}:(n=Math.max(1,n),function(r){r[e][t].duration=n}))},Yl.each=function(n,t){var e=this.id,r=this.namespace;if(arguments.length<2){var i=Ol,u=Hl;try{Hl=e,Y(this,function(t,i,u){Ol=t[r][e],n.call(t,t.__data__,i,u)})}finally{Ol=i,Hl=u}}else Y(this,function(i){var u=i[r][e];(u.event||(u.event=ao.dispatch("start","end","interrupt"))).on(n,t)});return this},Yl.transition=function(){for(var n,t,e,r,i=this.id,u=++Zl,o=this.namespace,a=[],l=0,c=this.length;c>l;l++){a.push(n=[]);for(var t=this[l],f=0,s=t.length;s>f;f++)(e=t[f])&&(r=e[o][i],Qu(e,f,o,u,{time:r.time,ease:r.ease,delay:r.delay+r.duration,duration:r.duration})),n.push(e)}return Wu(a,o,u)},ao.svg.axis=function(){function n(n){n.each(function(){var n,c=ao.select(this),f=this.__chart__||e,s=this.__chart__=e.copy(),h=null==l?s.ticks?s.ticks.apply(s,a):s.domain():l,p=null==t?s.tickFormat?s.tickFormat.apply(s,a):m:t,g=c.selectAll(".tick").data(h,s),v=g.enter().insert("g",".domain").attr("class","tick").style("opacity",Uo),d=ao.transition(g.exit()).style("opacity",Uo).remove(),y=ao.transition(g.order()).style("opacity",1),M=Math.max(i,0)+o,x=Zi(s),b=c.selectAll(".domain").data([0]),_=(b.enter().append("path").attr("class","domain"),ao.transition(b));v.append("line"),v.append("text");var w,S,k,N,E=v.select("line"),A=y.select("line"),C=g.select("text").text(p),z=v.select("text"),L=y.select("text"),q="top"===r||"left"===r?-1:1;if("bottom"===r||"top"===r?(n=no,w="x",k="y",S="x2",N="y2",C.attr("dy",0>q?"0em":".71em").style("text-anchor","middle"),_.attr("d","M"+x[0]+","+q*u+"V0H"+x[1]+"V"+q*u)):(n=to,w="y",k="x",S="y2",N="x2",C.attr("dy",".32em").style("text-anchor",0>q?"end":"start"),_.attr("d","M"+q*u+","+x[0]+"H0V"+x[1]+"H"+q*u)),E.attr(N,q*i),z.attr(k,q*M),A.attr(S,0).attr(N,q*i),L.attr(w,0).attr(k,q*M),s.rangeBand){var T=s,R=T.rangeBand()/2;f=s=function(n){return T(n)+R}}else f.rangeBand?f=s:d.call(n,s,f);v.call(n,f,s),y.call(n,s,s)})}var t,e=ao.scale.linear(),r=Vl,i=6,u=6,o=3,a=[10],l=null;return n.scale=function(t){return arguments.length?(e=t,n):e},n.orient=function(t){return arguments.length?(r=t in Xl?t+"":Vl,n):r},n.ticks=function(){return arguments.length?(a=co(arguments),n):a},n.tickValues=function(t){return arguments.length?(l=t,n):l},n.tickFormat=function(e){return arguments.length?(t=e,n):t},n.tickSize=function(t){var e=arguments.length;return e?(i=+t,u=+arguments[e-1],n):i},n.innerTickSize=function(t){return arguments.length?(i=+t,n):i},n.outerTickSize=function(t){return arguments.length?(u=+t,n):u},n.tickPadding=function(t){return arguments.length?(o=+t,n):o},n.tickSubdivide=function(){return arguments.length&&n},n};var Vl="bottom",Xl={top:1,right:1,bottom:1,left:1};ao.svg.brush=function(){function n(t){t.each(function(){var t=ao.select(this).style("pointer-events","all").style("-webkit-tap-highlight-color","rgba(0,0,0,0)").on("mousedown.brush",u).on("touchstart.brush",u),o=t.selectAll(".background").data([0]);o.enter().append("rect").attr("class","background").style("visibility","hidden").style("cursor","crosshair"),t.selectAll(".extent").data([0]).enter().append("rect").attr("class","extent").style("cursor","move");var a=t.selectAll(".resize").data(v,m);a.exit().remove(),a.enter().append("g").attr("class",function(n){return"resize "+n}).style("cursor",function(n){return $l[n]}).append("rect").attr("x",function(n){return/[ew]$/.test(n)?-3:null}).attr("y",function(n){return/^[ns]/.test(n)?-3:null}).attr("width",6).attr("height",6).style("visibility","hidden"),a.style("display",n.empty()?"none":null);var l,s=ao.transition(t),h=ao.transition(o);c&&(l=Zi(c),h.attr("x",l[0]).attr("width",l[1]-l[0]),r(s)),f&&(l=Zi(f),h.attr("y",l[0]).attr("height",l[1]-l[0]),i(s)),e(s)})}function e(n){n.selectAll(".resize").attr("transform",function(n){return"translate("+s[+/e$/.test(n)]+","+h[+/^s/.test(n)]+")"})}function r(n){n.select(".extent").attr("x",s[0]),n.selectAll(".extent,.n>rect,.s>rect").attr("width",s[1]-s[0])}function i(n){n.select(".extent").attr("y",h[0]),n.selectAll(".extent,.e>rect,.w>rect").attr("height",h[1]-h[0])}function u(){function u(){32==ao.event.keyCode&&(C||(M=null,L[0]-=s[1],L[1]-=h[1],C=2),S())}function v(){32==ao.event.keyCode&&2==C&&(L[0]+=s[1],L[1]+=h[1],C=0,S())}function d(){var n=ao.mouse(b),t=!1;x&&(n[0]+=x[0],n[1]+=x[1]),C||(ao.event.altKey?(M||(M=[(s[0]+s[1])/2,(h[0]+h[1])/2]),L[0]=s[+(n[0]<M[0])],L[1]=h[+(n[1]<M[1])]):M=null),E&&y(n,c,0)&&(r(k),t=!0),A&&y(n,f,1)&&(i(k),t=!0),t&&(e(k),w({type:"brush",mode:C?"move":"resize"}))}function y(n,t,e){var r,i,u=Zi(t),l=u[0],c=u[1],f=L[e],v=e?h:s,d=v[1]-v[0];return C&&(l-=f,c-=d+f),r=(e?g:p)?Math.max(l,Math.min(c,n[e])):n[e],C?i=(r+=f)+d:(M&&(f=Math.max(l,Math.min(c,2*M[e]-r))),r>f?(i=r,r=f):i=f),v[0]!=r||v[1]!=i?(e?a=null:o=null,v[0]=r,v[1]=i,!0):void 0}function m(){d(),k.style("pointer-events","all").selectAll(".resize").style("display",n.empty()?"none":null),ao.select("body").style("cursor",null),q.on("mousemove.brush",null).on("mouseup.brush",null).on("touchmove.brush",null).on("touchend.brush",null).on("keydown.brush",null).on("keyup.brush",null),z(),w({type:"brushend"})}var M,x,b=this,_=ao.select(ao.event.target),w=l.of(b,arguments),k=ao.select(b),N=_.datum(),E=!/^(n|s)$/.test(N)&&c,A=!/^(e|w)$/.test(N)&&f,C=_.classed("extent"),z=W(b),L=ao.mouse(b),q=ao.select(t(b)).on("keydown.brush",u).on("keyup.brush",v);if(ao.event.changedTouches?q.on("touchmove.brush",d).on("touchend.brush",m):q.on("mousemove.brush",d).on("mouseup.brush",m),k.interrupt().selectAll("*").interrupt(),C)L[0]=s[0]-L[0],L[1]=h[0]-L[1];else if(N){var T=+/w$/.test(N),R=+/^n/.test(N);x=[s[1-T]-L[0],h[1-R]-L[1]],L[0]=s[T],L[1]=h[R]}else ao.event.altKey&&(M=L.slice());k.style("pointer-events","none").selectAll(".resize").style("display",null),ao.select("body").style("cursor",_.style("cursor")),w({type:"brushstart"}),d()}var o,a,l=N(n,"brushstart","brush","brushend"),c=null,f=null,s=[0,0],h=[0,0],p=!0,g=!0,v=Bl[0];return n.event=function(n){n.each(function(){var n=l.of(this,arguments),t={x:s,y:h,i:o,j:a},e=this.__chart__||t;this.__chart__=t,Hl?ao.select(this).transition().each("start.brush",function(){o=e.i,a=e.j,s=e.x,h=e.y,n({type:"brushstart"})}).tween("brush:brush",function(){var e=xr(s,t.x),r=xr(h,t.y);return o=a=null,function(i){s=t.x=e(i),h=t.y=r(i),n({type:"brush",mode:"resize"})}}).each("end.brush",function(){o=t.i,a=t.j,n({type:"brush",mode:"resize"}),n({type:"brushend"})}):(n({type:"brushstart"}),n({type:"brush",mode:"resize"}),n({type:"brushend"}))})},n.x=function(t){return arguments.length?(c=t,v=Bl[!c<<1|!f],n):c},n.y=function(t){return arguments.length?(f=t,v=Bl[!c<<1|!f],n):f},n.clamp=function(t){return arguments.length?(c&&f?(p=!!t[0],g=!!t[1]):c?p=!!t:f&&(g=!!t),n):c&&f?[p,g]:c?p:f?g:null},n.extent=function(t){var e,r,i,u,l;return arguments.length?(c&&(e=t[0],r=t[1],f&&(e=e[0],r=r[0]),o=[e,r],c.invert&&(e=c(e),r=c(r)),e>r&&(l=e,e=r,r=l),e==s[0]&&r==s[1]||(s=[e,r])),f&&(i=t[0],u=t[1],c&&(i=i[1],u=u[1]),a=[i,u],f.invert&&(i=f(i),u=f(u)),i>u&&(l=i,i=u,u=l),i==h[0]&&u==h[1]||(h=[i,u])),n):(c&&(o?(e=o[0],r=o[1]):(e=s[0],r=s[1],c.invert&&(e=c.invert(e),r=c.invert(r)),e>r&&(l=e,e=r,r=l))),f&&(a?(i=a[0],u=a[1]):(i=h[0],u=h[1],f.invert&&(i=f.invert(i),u=f.invert(u)),i>u&&(l=i,i=u,u=l))),c&&f?[[e,i],[r,u]]:c?[e,r]:f&&[i,u])},n.clear=function(){return n.empty()||(s=[0,0],h=[0,0],o=a=null),n},n.empty=function(){return!!c&&s[0]==s[1]||!!f&&h[0]==h[1]},ao.rebind(n,l,"on")};var $l={n:"ns-resize",e:"ew-resize",s:"ns-resize",w:"ew-resize",nw:"nwse-resize",ne:"nesw-resize",se:"nwse-resize",sw:"nesw-resize"},Bl=[["n","e","s","w","nw","ne","se","sw"],["e","w"],["n","s"],[]],Wl=ga.format=xa.timeFormat,Jl=Wl.utc,Gl=Jl("%Y-%m-%dT%H:%M:%S.%LZ");Wl.iso=Date.prototype.toISOString&&+new Date("2000-01-01T00:00:00.000Z")?eo:Gl,eo.parse=function(n){var t=new Date(n);return isNaN(t)?null:t},eo.toString=Gl.toString,ga.second=On(function(n){return new va(1e3*Math.floor(n/1e3))},function(n,t){n.setTime(n.getTime()+1e3*Math.floor(t))},function(n){return n.getSeconds()}),ga.seconds=ga.second.range,ga.seconds.utc=ga.second.utc.range,ga.minute=On(function(n){return new va(6e4*Math.floor(n/6e4))},function(n,t){n.setTime(n.getTime()+6e4*Math.floor(t))},function(n){return n.getMinutes()}),ga.minutes=ga.minute.range,ga.minutes.utc=ga.minute.utc.range,ga.hour=On(function(n){var t=n.getTimezoneOffset()/60;return new va(36e5*(Math.floor(n/36e5-t)+t))},function(n,t){n.setTime(n.getTime()+36e5*Math.floor(t))},function(n){return n.getHours()}),ga.hours=ga.hour.range,ga.hours.utc=ga.hour.utc.range,ga.month=On(function(n){return n=ga.day(n),n.setDate(1),n},function(n,t){n.setMonth(n.getMonth()+t)},function(n){return n.getMonth()}),ga.months=ga.month.range,ga.months.utc=ga.month.utc.range;var Kl=[1e3,5e3,15e3,3e4,6e4,3e5,9e5,18e5,36e5,108e5,216e5,432e5,864e5,1728e5,6048e5,2592e6,7776e6,31536e6],Ql=[[ga.second,1],[ga.second,5],[ga.second,15],[ga.second,30],[ga.minute,1],[ga.minute,5],[ga.minute,15],[ga.minute,30],[ga.hour,1],[ga.hour,3],[ga.hour,6],[ga.hour,12],[ga.day,1],[ga.day,2],[ga.week,1],[ga.month,1],[ga.month,3],[ga.year,1]],nc=Wl.multi([[".%L",function(n){return n.getMilliseconds()}],[":%S",function(n){return n.getSeconds()}],["%I:%M",function(n){return n.getMinutes()}],["%I %p",function(n){return n.getHours()}],["%a %d",function(n){return n.getDay()&&1!=n.getDate()}],["%b %d",function(n){return 1!=n.getDate()}],["%B",function(n){return n.getMonth()}],["%Y",zt]]),tc={range:function(n,t,e){return ao.range(Math.ceil(n/e)*e,+t,e).map(io)},floor:m,ceil:m};Ql.year=ga.year,ga.scale=function(){return ro(ao.scale.linear(),Ql,nc)};var ec=Ql.map(function(n){return[n[0].utc,n[1]]}),rc=Jl.multi([[".%L",function(n){return n.getUTCMilliseconds()}],[":%S",function(n){return n.getUTCSeconds()}],["%I:%M",function(n){return n.getUTCMinutes()}],["%I %p",function(n){return n.getUTCHours()}],["%a %d",function(n){return n.getUTCDay()&&1!=n.getUTCDate()}],["%b %d",function(n){return 1!=n.getUTCDate()}],["%B",function(n){return n.getUTCMonth()}],["%Y",zt]]);ec.year=ga.year.utc,ga.scale.utc=function(){return ro(ao.scale.linear(),ec,rc)},ao.text=An(function(n){return n.responseText}),ao.json=function(n,t){return Cn(n,"application/json",uo,t)},ao.html=function(n,t){return Cn(n,"text/html",oo,t)},ao.xml=An(function(n){return n.responseXML}),"function"==typeof define&&define.amd?(this.d3=ao,define(ao)):"object"==typeof module&&module.exports?module.exports=ao:this.d3=ao}();
```
