## Supplementary material for "Method for inferring migration routes from phylogenetic trees": Some route-trees: diff.js

```
/**
 * This library was modified by Harrison Liddiard. The source code to this
 * modified version can be found at https://github.com/liddiard/google-diff/.
 * The original source code can be found at
 * http://code.google.com/p/google-diff-match-patch/. This unofficial fork is
 * not maintained by or affiliated with Google Inc. The original attribution
 * and licensing information follows.
 */

/**
 * Diff Match and Patch
 *
 * Copyright 2006 Google Inc.
 * http://code.google.com/p/google-diff-match-patch/
 *
 * Licensed under the Apache License, Version 2.0 (the "License");
 * you may not use this file except in compliance with the License.
 * You may obtain a copy of the License at
 *
 *   http://www.apache.org/licenses/LICENSE-2.0
 *
 * Unless required by applicable law or agreed to in writing, software
 * distributed under the License is distributed on an "AS IS" BASIS,
 * WITHOUT WARRANTIES OR CONDITIONS OF ANY KIND, either express or implied.
 * See the License for the specific language governing permissions and
 * limitations under the License.
 */

/**
 * @fileoverview Computes the difference between two texts to create a patch.
 * Applies the patch onto another text, allowing for errors.
 * @author (Neil Fraser)
 */

/**
 * Class containing the diff.
 * @constructor
 */
function diff(options) {
  var options = options || {};

  // Defaults.
  // Redefine these in your program to override the defaults.

  // Number of seconds to map a diff before giving up (0 for infinity).
  this.Timeout = options.timeout || 1.0;
  // Cost of an empty edit operation in terms of edit characters.
  this.EditCost = options.editCost || 4;
}


//  DIFF FUNCTIONS


/**
 * The data structure representing a diff is an array of tuples:
 * [[DIFF_DELETE, 'Hello'], [DIFF_INSERT, 'Goodbye'], [DIFF_EQUAL, ' world.']]
 * which means: delete 'Hello', add 'Goodbye' and keep ' world.'
 */
var DIFF_DELETE = -1;
var DIFF_INSERT = 1;
var DIFF_EQUAL = 0;

/** @typedef {{0: number, 1: string}} */
diff.Diff;


/**
 * Find the differences between two texts.  Simplifies the problem by stripping
 * any common prefix or suffix off the texts before diffing.
 * @param {string} text1 Old string to be diffed.
 * @param {string} text2 New string to be diffed.
 * @param {boolean=} opt_checklines Optional speedup flag. If present and false,
 *     then don't run a line-level diff first to identify the changed areas.
 *     Defaults to true, which does a faster, slightly less optimal diff.
 * @param {number} opt_deadline Optional time when the diff should be complete
 *     by.  Used internally for recursive calls.  Users should set DiffTimeout
 *     instead.
 * @return {!Array.<!diff.Diff>} Array of diff tuples.
 */
diff.prototype.main = function(text1, text2, opt_checklines,
    opt_deadline) {
  // Set a deadline by which time the diff must be complete.
  if (typeof opt_deadline == 'undefined') {
    if (this.Timeout <= 0) {
      opt_deadline = Number.MAX_VALUE;
    } else {
      opt_deadline = (new Date).getTime() + this.Timeout * 1000;
    }
  }
  var deadline = opt_deadline;

  // Check for null inputs.
  if (text1 == null || text2 == null) {
    throw new Error('Null input. (diff_main)');
  }

  // Check for equality (speedup).
  if (text1 == text2) {
    if (text1) {
      return [[DIFF_EQUAL, text1]];
    }
    return [];
  }

  if (typeof opt_checklines == 'undefined') {
    opt_checklines = true;
  }
  var checklines = opt_checklines;

  // Trim off common prefix (speedup).
  var commonlength = this.commonPrefix(text1, text2);
  var commonprefix = text1.substring(0, commonlength);
  text1 = text1.substring(commonlength);
  text2 = text2.substring(commonlength);

  // Trim off common suffix (speedup).
  commonlength = this.commonSuffix(text1, text2);
  var commonsuffix = text1.substring(text1.length - commonlength);
  text1 = text1.substring(0, text1.length - commonlength);
  text2 = text2.substring(0, text2.length - commonlength);

  // Compute the diff on the middle block.
  var diffs = this.compute_(text1, text2, checklines, deadline);

  // Restore the prefix and suffix.
  if (commonprefix) {
    diffs.unshift([DIFF_EQUAL, commonprefix]);
  }
  if (commonsuffix) {
    diffs.push([DIFF_EQUAL, commonsuffix]);
  }
  this.cleanupMerge(diffs);
  return diffs;
};


/**
 * Find the differences between two texts.  Assumes that the texts do not
 * have any common prefix or suffix.
 * @param {string} text1 Old string to be diffed.
 * @param {string} text2 New string to be diffed.
 * @param {boolean} checklines Speedup flag.  If false, then don't run a
 *     line-level diff first to identify the changed areas.
 *     If true, then run a faster, slightly less optimal diff.
 * @param {number} deadline Time when the diff should be complete by.
 * @return {!Array.<!diff.Diff>} Array of diff tuples.
 * @private
 */
diff.prototype.compute_ = function(text1, text2, checklines,
    deadline) {
  var diffs;

  if (!text1) {
    // Just add some text (speedup).
    return [[DIFF_INSERT, text2]];
  }

  if (!text2) {
    // Just delete some text (speedup).
    return [[DIFF_DELETE, text1]];
  }

  var longtext = text1.length > text2.length ? text1 : text2;
  var shorttext = text1.length > text2.length ? text2 : text1;
  var i = longtext.indexOf(shorttext);
  if (i != -1) {
    // Shorter text is inside the longer text (speedup).
    diffs = [[DIFF_INSERT, longtext.substring(0, i)],
             [DIFF_EQUAL, shorttext],
             [DIFF_INSERT, longtext.substring(i + shorttext.length)]];
    // Swap insertions for deletions if diff is reversed.
    if (text1.length > text2.length) {
      diffs[0][0] = diffs[2][0] = DIFF_DELETE;
    }
    return diffs;
  }

  if (shorttext.length == 1) {
    // Single character string.
    // After the previous speedup, the character can't be an equality.
    return [[DIFF_DELETE, text1], [DIFF_INSERT, text2]];
  }

  // Check to see if the problem can be split in two.
  var hm = this.halfMatch_(text1, text2);
  if (hm) {
    // A half-match was found, sort out the return data.
    var text1_a = hm[0];
    var text1_b = hm[1];
    var text2_a = hm[2];
    var text2_b = hm[3];
    var mid_common = hm[4];
    // Send both pairs off for separate processing.
    var diffs_a = this.main(text1_a, text2_a, checklines, deadline);
    var diffs_b = this.main(text1_b, text2_b, checklines, deadline);
    // Merge the results.
    return diffs_a.concat([[DIFF_EQUAL, mid_common]], diffs_b);
  }

  if (checklines && text1.length > 100 && text2.length > 100) {
    return this.lineMode_(text1, text2, deadline);
  }

  return this.bisect_(text1, text2, deadline);
};


/**
 * Do a quick line-level diff on both strings, then rediff the parts for
 * greater accuracy.
 * This speedup can produce non-minimal diffs.
 * @param {string} text1 Old string to be diffed.
 * @param {string} text2 New string to be diffed.
 * @param {number} deadline Time when the diff should be complete by.
 * @return {!Array.<!diff.Diff>} Array of diff tuples.
 * @private
 */
diff.prototype.lineMode_ = function(text1, text2, deadline) {
  // Scan the text on a line-by-line basis first.
  var a = this.linesToChars_(text1, text2);
  text1 = a.chars1;
  text2 = a.chars2;
  var linearray = a.lineArray;

  var diffs = this.main(text1, text2, false, deadline);

  // Convert the diff back to original text.
  this.charsToLines_(diffs, linearray);
  // Eliminate freak matches (e.g. blank lines)
  this.cleanupSemantic(diffs);

  // Rediff any replacement blocks, this time character-by-character.
  // Add a dummy entry at the end.
  diffs.push([DIFF_EQUAL, '']);
  var pointer = 0;
  var count_delete = 0;
  var count_insert = 0;
  var text_delete = '';
  var text_insert = '';
  while (pointer < diffs.length) {
    switch (diffs[pointer][0]) {
      case DIFF_INSERT:
        count_insert++;
        text_insert += diffs[pointer][1];
        break;
      case DIFF_DELETE:
        count_delete++;
        text_delete += diffs[pointer][1];
        break;
      case DIFF_EQUAL:
        // Upon reaching an equality, check for prior redundancies.
        if (count_delete >= 1 && count_insert >= 1) {
          // Delete the offending records and add the merged ones.
          diffs.splice(pointer - count_delete - count_insert,
                       count_delete + count_insert);
          pointer = pointer - count_delete - count_insert;
          var a = this.main(text_delete, text_insert, false, deadline);
          for (var j = a.length - 1; j >= 0; j--) {
            diffs.splice(pointer, 0, a[j]);
          }
          pointer = pointer + a.length;
        }
        count_insert = 0;
        count_delete = 0;
        text_delete = '';
        text_insert = '';
        break;
    }
    pointer++;
  }
  diffs.pop();  // Remove the dummy entry at the end.

  return diffs;
};


/**
 * Find the 'middle snake' of a diff, split the problem in two
 * and return the recursively constructed diff.
 * See Myers 1986 paper: An O(ND) Difference Algorithm and Its Variations.
 * @param {string} text1 Old string to be diffed.
 * @param {string} text2 New string to be diffed.
 * @param {number} deadline Time at which to bail if not yet complete.
 * @return {!Array.<!diff.Diff>} Array of diff tuples.
 * @private
 */
diff.prototype.bisect_ = function(text1, text2, deadline) {
  // Cache the text lengths to prevent multiple calls.
  var text1_length = text1.length;
  var text2_length = text2.length;
  var max_d = Math.ceil((text1_length + text2_length) / 2);
  var v_offset = max_d;
  var v_length = 2 * max_d;
  var v1 = new Array(v_length);
  var v2 = new Array(v_length);
  // Setting all elements to -1 is faster in Chrome & Firefox than mixing
  // integers and undefined.
  for (var x = 0; x < v_length; x++) {
    v1[x] = -1;
    v2[x] = -1;
  }
  v1[v_offset + 1] = 0;
  v2[v_offset + 1] = 0;
  var delta = text1_length - text2_length;
  // If the total number of characters is odd, then the front path will collide
  // with the reverse path.
  var front = (delta % 2 != 0);
  // Offsets for start and end of k loop.
  // Prevents mapping of space beyond the grid.
  var k1start = 0;
  var k1end = 0;
  var k2start = 0;
  var k2end = 0;
  for (var d = 0; d < max_d; d++) {
    // Bail out if deadline is reached.
    if ((new Date()).getTime() > deadline) {
      break;
    }

    // Walk the front path one step.
    for (var k1 = -d + k1start; k1 <= d - k1end; k1 += 2) {
      var k1_offset = v_offset + k1;
      var x1;
      if (k1 == -d || (k1 != d && v1[k1_offset - 1] < v1[k1_offset + 1])) {
        x1 = v1[k1_offset + 1];
      } else {
        x1 = v1[k1_offset - 1] + 1;
      }
      var y1 = x1 - k1;
      while (x1 < text1_length && y1 < text2_length &&
             text1.charAt(x1) == text2.charAt(y1)) {
        x1++;
        y1++;
      }
      v1[k1_offset] = x1;
      if (x1 > text1_length) {
        // Ran off the right of the graph.
        k1end += 2;
      } else if (y1 > text2_length) {
        // Ran off the bottom of the graph.
        k1start += 2;
      } else if (front) {
        var k2_offset = v_offset + delta - k1;
        if (k2_offset >= 0 && k2_offset < v_length && v2[k2_offset] != -1) {
          // Mirror x2 onto top-left coordinate system.
          var x2 = text1_length - v2[k2_offset];
          if (x1 >= x2) {
            // Overlap detected.
            return this.bisectSplit_(text1, text2, x1, y1, deadline);
          }
        }
      }
    }

    // Walk the reverse path one step.
    for (var k2 = -d + k2start; k2 <= d - k2end; k2 += 2) {
      var k2_offset = v_offset + k2;
      var x2;
      if (k2 == -d || (k2 != d && v2[k2_offset - 1] < v2[k2_offset + 1])) {
        x2 = v2[k2_offset + 1];
      } else {
        x2 = v2[k2_offset - 1] + 1;
      }
      var y2 = x2 - k2;
      while (x2 < text1_length && y2 < text2_length &&
             text1.charAt(text1_length - x2 - 1) ==
             text2.charAt(text2_length - y2 - 1)) {
        x2++;
        y2++;
      }
      v2[k2_offset] = x2;
      if (x2 > text1_length) {
        // Ran off the left of the graph.
        k2end += 2;
      } else if (y2 > text2_length) {
        // Ran off the top of the graph.
        k2start += 2;
      } else if (!front) {
        var k1_offset = v_offset + delta - k2;
        if (k1_offset >= 0 && k1_offset < v_length && v1[k1_offset] != -1) {
          var x1 = v1[k1_offset];
          var y1 = v_offset + x1 - k1_offset;
          // Mirror x2 onto top-left coordinate system.
          x2 = text1_length - x2;
          if (x1 >= x2) {
            // Overlap detected.
            return this.bisectSplit_(text1, text2, x1, y1, deadline);
          }
        }
      }
    }
  }
  // Diff took too long and hit the deadline or
  // number of diffs equals number of characters, no commonality at all.
  return [[DIFF_DELETE, text1], [DIFF_INSERT, text2]];
};


/**
 * Given the location of the 'middle snake', split the diff in two parts
 * and recurse.
 * @param {string} text1 Old string to be diffed.
 * @param {string} text2 New string to be diffed.
 * @param {number} x Index of split point in text1.
 * @param {number} y Index of split point in text2.
 * @param {number} deadline Time at which to bail if not yet complete.
 * @return {!Array.<!diff.Diff>} Array of diff tuples.
 * @private
 */
diff.prototype.bisectSplit_ = function(text1, text2, x, y,
    deadline) {
  var text1a = text1.substring(0, x);
  var text2a = text2.substring(0, y);
  var text1b = text1.substring(x);
  var text2b = text2.substring(y);

  // Compute both diffs serially.
  var diffs = this.main(text1a, text2a, false, deadline);
  var diffsb = this.main(text1b, text2b, false, deadline);

  return diffs.concat(diffsb);
};


/**
 * Split two texts into an array of strings.  Reduce the texts to a string of
 * hashes where each Unicode character represents one line.
 * @param {string} text1 First string.
 * @param {string} text2 Second string.
 * @return {{chars1: string, chars2: string, lineArray: !Array.<string>}}
 *     An object containing the encoded text1, the encoded text2 and
 *     the array of unique strings.
 *     The zeroth element of the array of unique strings is intentionally blank.
 * @private
 */
diff.prototype.linesToChars_ = function(text1, text2) {
  var lineArray = [];  // e.g. lineArray[4] == 'Hello\n'
  var lineHash = {};   // e.g. lineHash['Hello\n'] == 4

  // '\x00' is a valid character, but various debuggers don't like it.
  // So we'll insert a junk entry to avoid generating a null character.
  lineArray[0] = '';

  /**
   * Split a text into an array of strings.  Reduce the texts to a string of
   * hashes where each Unicode character represents one line.
   * Modifies linearray and linehash through being a closure.
   * @param {string} text String to encode.
   * @return {string} Encoded string.
   * @private
   */
  function diff_linesToCharsMunge_(text) {
    var chars = '';
    // Walk the text, pulling out a substring for each line.
    // text.split('\n') would would temporarily double our memory footprint.
    // Modifying text would create many large strings to garbage collect.
    var lineStart = 0;
    var lineEnd = -1;
    // Keeping our own length variable is faster than looking it up.
    var lineArrayLength = lineArray.length;
    while (lineEnd < text.length - 1) {
      lineEnd = text.indexOf('\n', lineStart);
      if (lineEnd == -1) {
        lineEnd = text.length - 1;
      }
      var line = text.substring(lineStart, lineEnd + 1);
      lineStart = lineEnd + 1;

      if (lineHash.hasOwnProperty ? lineHash.hasOwnProperty(line) :
          (lineHash[line] !== undefined)) {
        chars += String.fromCharCode(lineHash[line]);
      } else {
        chars += String.fromCharCode(lineArrayLength);
        lineHash[line] = lineArrayLength;
        lineArray[lineArrayLength++] = line;
      }
    }
    return chars;
  }

  var chars1 = diff_linesToCharsMunge_(text1);
  var chars2 = diff_linesToCharsMunge_(text2);
  return {chars1: chars1, chars2: chars2, lineArray: lineArray};
};


/**
 * Rehydrate the text in a diff from a string of line hashes to real lines of
 * text.
 * @param {!Array.<!diff.Diff>} diffs Array of diff tuples.
 * @param {!Array.<string>} lineArray Array of unique strings.
 * @private
 */
diff.prototype.charsToLines_ = function(diffs, lineArray) {
  for (var x = 0; x < diffs.length; x++) {
    var chars = diffs[x][1];
    var text = [];
    for (var y = 0; y < chars.length; y++) {
      text[y] = lineArray[chars.charCodeAt(y)];
    }
    diffs[x][1] = text.join('');
  }
};


/**
 * Determine the common prefix of two strings.
 * @param {string} text1 First string.
 * @param {string} text2 Second string.
 * @return {number} The number of characters common to the start of each
 *     string.
 */
diff.prototype.commonPrefix = function(text1, text2) {
  // Quick check for common null cases.
  if (!text1 || !text2 || text1.charAt(0) != text2.charAt(0)) {
    return 0;
  }
  // Binary search.
  // Performance analysis: http://neil.fraser.name/news/2007/10/09/
  var pointermin = 0;
  var pointermax = Math.min(text1.length, text2.length);
  var pointermid = pointermax;
  var pointerstart = 0;
  while (pointermin < pointermid) {
    if (text1.substring(pointerstart, pointermid) ==
        text2.substring(pointerstart, pointermid)) {
      pointermin = pointermid;
      pointerstart = pointermin;
    } else {
      pointermax = pointermid;
    }
    pointermid = Math.floor((pointermax - pointermin) / 2 + pointermin);
  }
  return pointermid;
};


/**
 * Determine the common suffix of two strings.
 * @param {string} text1 First string.
 * @param {string} text2 Second string.
 * @return {number} The number of characters common to the end of each string.
 */
diff.prototype.commonSuffix = function(text1, text2) {
  // Quick check for common null cases.
  if (!text1 || !text2 ||
      text1.charAt(text1.length - 1) != text2.charAt(text2.length - 1)) {
    return 0;
  }
  // Binary search.
  // Performance analysis: http://neil.fraser.name/news/2007/10/09/
  var pointermin = 0;
  var pointermax = Math.min(text1.length, text2.length);
  var pointermid = pointermax;
  var pointerend = 0;
  while (pointermin < pointermid) {
    if (text1.substring(text1.length - pointermid, text1.length - pointerend) ==
        text2.substring(text2.length - pointermid, text2.length - pointerend)) {
      pointermin = pointermid;
      pointerend = pointermin;
    } else {
      pointermax = pointermid;
    }
    pointermid = Math.floor((pointermax - pointermin) / 2 + pointermin);
  }
  return pointermid;
};


/**
 * Determine if the suffix of one string is the prefix of another.
 * @param {string} text1 First string.
 * @param {string} text2 Second string.
 * @return {number} The number of characters common to the end of the first
 *     string and the start of the second string.
 * @private
 */
diff.prototype.commonOverlap_ = function(text1, text2) {
  // Cache the text lengths to prevent multiple calls.
  var text1_length = text1.length;
  var text2_length = text2.length;
  // Eliminate the null case.
  if (text1_length == 0 || text2_length == 0) {
    return 0;
  }
  // Truncate the longer string.
  if (text1_length > text2_length) {
    text1 = text1.substring(text1_length - text2_length);
  } else if (text1_length < text2_length) {
    text2 = text2.substring(0, text1_length);
  }
  var text_length = Math.min(text1_length, text2_length);
  // Quick check for the worst case.
  if (text1 == text2) {
    return text_length;
  }

  // Start by looking for a single character match
  // and increase length until no match is found.
  // Performance analysis: http://neil.fraser.name/news/2010/11/04/
  var best = 0;
  var length = 1;
  while (true) {
    var pattern = text1.substring(text_length - length);
    var found = text2.indexOf(pattern);
    if (found == -1) {
      return best;
    }
    length += found;
    if (found == 0 || text1.substring(text_length - length) ==
        text2.substring(0, length)) {
      best = length;
      length++;
    }
  }
};


/**
 * Do the two texts share a substring which is at least half the length of the
 * longer text?
 * This speedup can produce non-minimal diffs.
 * @param {string} text1 First string.
 * @param {string} text2 Second string.
 * @return {Array.<string>} Five element Array, containing the prefix of
 *     text1, the suffix of text1, the prefix of text2, the suffix of
 *     text2 and the common middle.  Or null if there was no match.
 * @private
 */
diff.prototype.halfMatch_ = function(text1, text2) {
  if (this.Timeout <= 0) {
    // Don't risk returning a non-optimal diff if we have unlimited time.
    return null;
  }
  var longtext = text1.length > text2.length ? text1 : text2;
  var shorttext = text1.length > text2.length ? text2 : text1;
  if (longtext.length < 4 || shorttext.length * 2 < longtext.length) {
    return null;  // Pointless.
  }
  var dmp = this;  // 'this' becomes 'window' in a closure.

  /**
   * Does a substring of shorttext exist within longtext such that the substring
   * is at least half the length of longtext?
   * Closure, but does not reference any external variables.
   * @param {string} longtext Longer string.
   * @param {string} shorttext Shorter string.
   * @param {number} i Start index of quarter length substring within longtext.
   * @return {Array.<string>} Five element Array, containing the prefix of
   *     longtext, the suffix of longtext, the prefix of shorttext, the suffix
   *     of shorttext and the common middle.  Or null if there was no match.
   * @private
   */
  function diff_halfMatchI_(longtext, shorttext, i) {
    // Start with a 1/4 length substring at position i as a seed.
    var seed = longtext.substring(i, i + Math.floor(longtext.length / 4));
    var j = -1;
    var best_common = '';
    var best_longtext_a, best_longtext_b, best_shorttext_a, best_shorttext_b;
    while ((j = shorttext.indexOf(seed, j + 1)) != -1) {
      var prefixLength = dmp.commonPrefix(longtext.substring(i),
                                          shorttext.substring(j));
      var suffixLength = dmp.commonSuffix(longtext.substring(0, i),
                                          shorttext.substring(0, j));
      if (best_common.length < suffixLength + prefixLength) {
        best_common = shorttext.substring(j - suffixLength, j) +
            shorttext.substring(j, j + prefixLength);
        best_longtext_a = longtext.substring(0, i - suffixLength);
        best_longtext_b = longtext.substring(i + prefixLength);
        best_shorttext_a = shorttext.substring(0, j - suffixLength);
        best_shorttext_b = shorttext.substring(j + prefixLength);
      }
    }
    if (best_common.length * 2 >= longtext.length) {
      return [best_longtext_a, best_longtext_b,
              best_shorttext_a, best_shorttext_b, best_common];
    } else {
      return null;
    }
  }

  // First check if the second quarter is the seed for a half-match.
  var hm1 = diff_halfMatchI_(longtext, shorttext,
                             Math.ceil(longtext.length / 4));
  // Check again based on the third quarter.
  var hm2 = diff_halfMatchI_(longtext, shorttext,
                             Math.ceil(longtext.length / 2));
  var hm;
  if (!hm1 && !hm2) {
    return null;
  } else if (!hm2) {
    hm = hm1;
  } else if (!hm1) {
    hm = hm2;
  } else {
    // Both matched.  Select the longest.
    hm = hm1[4].length > hm2[4].length ? hm1 : hm2;
  }

  // A half-match was found, sort out the return data.
  var text1_a, text1_b, text2_a, text2_b;
  if (text1.length > text2.length) {
    text1_a = hm[0];
    text1_b = hm[1];
    text2_a = hm[2];
    text2_b = hm[3];
  } else {
    text2_a = hm[0];
    text2_b = hm[1];
    text1_a = hm[2];
    text1_b = hm[3];
  }
  var mid_common = hm[4];
  return [text1_a, text1_b, text2_a, text2_b, mid_common];
};


/**
 * Reduce the number of edits by eliminating semantically trivial equalities.
 * @param {!Array.<!diff.Diff>} diffs Array of diff tuples.
 */
diff.prototype.cleanupSemantic = function(diffs) {
  var changes = false;
  var equalities = [];  // Stack of indices where equalities are found.
  var equalitiesLength = 0;  // Keeping our own length var is faster in JS.
  /** @type {?string} */
  var lastequality = null;
  // Always equal to diffs[equalities[equalitiesLength - 1]][1]
  var pointer = 0;  // Index of current position.
  // Number of characters that changed prior to the equality.
  var length_insertions1 = 0;
  var length_deletions1 = 0;
  // Number of characters that changed after the equality.
  var length_insertions2 = 0;
  var length_deletions2 = 0;
  while (pointer < diffs.length) {
    if (diffs[pointer][0] == DIFF_EQUAL) {  // Equality found.
      equalities[equalitiesLength++] = pointer;
      length_insertions1 = length_insertions2;
      length_deletions1 = length_deletions2;
      length_insertions2 = 0;
      length_deletions2 = 0;
      lastequality = diffs[pointer][1];
    } else {  // An insertion or deletion.
      if (diffs[pointer][0] == DIFF_INSERT) {
        length_insertions2 += diffs[pointer][1].length;
      } else {
        length_deletions2 += diffs[pointer][1].length;
      }
      // Eliminate an equality that is smaller or equal to the edits on both
      // sides of it.
      if (lastequality && (lastequality.length <=
          Math.max(length_insertions1, length_deletions1)) &&
          (lastequality.length <= Math.max(length_insertions2,
                                           length_deletions2))) {
        // Duplicate record.
        diffs.splice(equalities[equalitiesLength - 1], 0,
                     [DIFF_DELETE, lastequality]);
        // Change second copy to insert.
        diffs[equalities[equalitiesLength - 1] + 1][0] = DIFF_INSERT;
        // Throw away the equality we just deleted.
        equalitiesLength--;
        // Throw away the previous equality (it needs to be reevaluated).
        equalitiesLength--;
        pointer = equalitiesLength > 0 ? equalities[equalitiesLength - 1] : -1;
        length_insertions1 = 0;  // Reset the counters.
        length_deletions1 = 0;
        length_insertions2 = 0;
        length_deletions2 = 0;
        lastequality = null;
        changes = true;
      }
    }
    pointer++;
  }

  // Normalize the diff.
  if (changes) {
    this.cleanupMerge(diffs);
  }
  this.cleanupSemanticLossless(diffs);

  // Find any overlaps between deletions and insertions.
  // e.g: <del>abcxxx</del><ins>xxxdef</ins>
  //   -> <del>abc</del>xxx<ins>def</ins>
  // e.g: <del>xxxabc</del><ins>defxxx</ins>
  //   -> <ins>def</ins>xxx<del>abc</del>
  // Only extract an overlap if it is as big as the edit ahead or behind it.
  pointer = 1;
  while (pointer < diffs.length) {
    if (diffs[pointer - 1][0] == DIFF_DELETE &&
        diffs[pointer][0] == DIFF_INSERT) {
      var deletion = diffs[pointer - 1][1];
      var insertion = diffs[pointer][1];
      var overlap_length1 = this.commonOverlap_(deletion, insertion);
      var overlap_length2 = this.commonOverlap_(insertion, deletion);
      if (overlap_length1 >= overlap_length2) {
        if (overlap_length1 >= deletion.length / 2 ||
            overlap_length1 >= insertion.length / 2) {
          // Overlap found.  Insert an equality and trim the surrounding edits.
          diffs.splice(pointer, 0,
              [DIFF_EQUAL, insertion.substring(0, overlap_length1)]);
          diffs[pointer - 1][1] =
              deletion.substring(0, deletion.length - overlap_length1);
          diffs[pointer + 1][1] = insertion.substring(overlap_length1);
          pointer++;
        }
      } else {
        if (overlap_length2 >= deletion.length / 2 ||
            overlap_length2 >= insertion.length / 2) {
          // Reverse overlap found.
          // Insert an equality and swap and trim the surrounding edits.
          diffs.splice(pointer, 0,
              [DIFF_EQUAL, deletion.substring(0, overlap_length2)]);
          diffs[pointer - 1][0] = DIFF_INSERT;
          diffs[pointer - 1][1] =
              insertion.substring(0, insertion.length - overlap_length2);
          diffs[pointer + 1][0] = DIFF_DELETE;
          diffs[pointer + 1][1] =
              deletion.substring(overlap_length2);
          pointer++;
        }
      }
      pointer++;
    }
    pointer++;
  }
};


/**
 * Look for single edits surrounded on both sides by equalities
 * which can be shifted sideways to align the edit to a word boundary.
 * e.g: The c<ins>at c</ins>ame. -> The <ins>cat </ins>came.
 * @param {!Array.<!diff.Diff>} diffs Array of diff tuples.
 */
diff.prototype.cleanupSemanticLossless = function(diffs) {
  /**
   * Given two strings, compute a score representing whether the internal
   * boundary falls on logical boundaries.
   * Scores range from 6 (best) to 0 (worst).
   * Closure, but does not reference any external variables.
   * @param {string} one First string.
   * @param {string} two Second string.
   * @return {number} The score.
   * @private
   */
  function diff_cleanupSemanticScore_(one, two) {
    if (!one || !two) {
      // Edges are the best.
      return 6;
    }

    // Each port of this function behaves slightly differently due to
    // subtle differences in each language's definition of things like
    // 'whitespace'.  Since this function's purpose is largely cosmetic,
    // the choice has been made to use each language's native features
    // rather than force total conformity.
    var char1 = one.charAt(one.length - 1);
    var char2 = two.charAt(0);
    var nonAlphaNumeric1 = char1.match(diff.nonAlphaNumericRegex_);
    var nonAlphaNumeric2 = char2.match(diff.nonAlphaNumericRegex_);
    var whitespace1 = nonAlphaNumeric1 &&
        char1.match(diff.whitespaceRegex_);
    var whitespace2 = nonAlphaNumeric2 &&
        char2.match(diff.whitespaceRegex_);
    var lineBreak1 = whitespace1 &&
        char1.match(diff.linebreakRegex_);
    var lineBreak2 = whitespace2 &&
        char2.match(diff.linebreakRegex_);
    var blankLine1 = lineBreak1 &&
        one.match(diff.blanklineEndRegex_);
    var blankLine2 = lineBreak2 &&
        two.match(diff.blanklineStartRegex_);

    if (blankLine1 || blankLine2) {
      // Five points for blank lines.
      return 5;
    } else if (lineBreak1 || lineBreak2) {
      // Four points for line breaks.
      return 4;
    } else if (nonAlphaNumeric1 && !whitespace1 && whitespace2) {
      // Three points for end of sentences.
      return 3;
    } else if (whitespace1 || whitespace2) {
      // Two points for whitespace.
      return 2;
    } else if (nonAlphaNumeric1 || nonAlphaNumeric2) {
      // One point for non-alphanumeric.
      return 1;
    }
    return 0;
  }

  var pointer = 1;
  // Intentionally ignore the first and last element (don't need checking).
  while (pointer < diffs.length - 1) {
    if (diffs[pointer - 1][0] == DIFF_EQUAL &&
        diffs[pointer + 1][0] == DIFF_EQUAL) {
      // This is a single edit surrounded by equalities.
      var equality1 = diffs[pointer - 1][1];
      var edit = diffs[pointer][1];
      var equality2 = diffs[pointer + 1][1];

      // First, shift the edit as far left as possible.
      var commonOffset = this.commonSuffix(equality1, edit);
      if (commonOffset) {
        var commonString = edit.substring(edit.length - commonOffset);
        equality1 = equality1.substring(0, equality1.length - commonOffset);
        edit = commonString + edit.substring(0, edit.length - commonOffset);
        equality2 = commonString + equality2;
      }

      // Second, step character by character right, looking for the best fit.
      var bestEquality1 = equality1;
      var bestEdit = edit;
      var bestEquality2 = equality2;
      var bestScore = diff_cleanupSemanticScore_(equality1, edit) +
          diff_cleanupSemanticScore_(edit, equality2);
      while (edit.charAt(0) === equality2.charAt(0)) {
        equality1 += edit.charAt(0);
        edit = edit.substring(1) + equality2.charAt(0);
        equality2 = equality2.substring(1);
        var score = diff_cleanupSemanticScore_(equality1, edit) +
            diff_cleanupSemanticScore_(edit, equality2);
        // The >= encourages trailing rather than leading whitespace on edits.
        if (score >= bestScore) {
          bestScore = score;
          bestEquality1 = equality1;
          bestEdit = edit;
          bestEquality2 = equality2;
        }
      }

      if (diffs[pointer - 1][1] != bestEquality1) {
        // We have an improvement, save it back to the diff.
        if (bestEquality1) {
          diffs[pointer - 1][1] = bestEquality1;
        } else {
          diffs.splice(pointer - 1, 1);
          pointer--;
        }
        diffs[pointer][1] = bestEdit;
        if (bestEquality2) {
          diffs[pointer + 1][1] = bestEquality2;
        } else {
          diffs.splice(pointer + 1, 1);
          pointer--;
        }
      }
    }
    pointer++;
  }
};

// Define some regex patterns for matching boundaries.
diff.nonAlphaNumericRegex_ = /[^a-zA-Z0-9]/;
diff.whitespaceRegex_ = /\s/;
diff.linebreakRegex_ = /[\r\n]/;
diff.blanklineEndRegex_ = /\n\r?\n$/;
diff.blanklineStartRegex_ = /^\r?\n\r?\n/;

/**
 * Reduce the number of edits by eliminating operationally trivial equalities.
 * @param {!Array.<!diff.Diff>} diffs Array of diff tuples.
 */
diff.prototype.cleanupEfficiency = function(diffs) {
  var changes = false;
  var equalities = [];  // Stack of indices where equalities are found.
  var equalitiesLength = 0;  // Keeping our own length var is faster in JS.
  /** @type {?string} */
  var lastequality = null;
  // Always equal to diffs[equalities[equalitiesLength - 1]][1]
  var pointer = 0;  // Index of current position.
  // Is there an insertion operation before the last equality.
  var pre_ins = false;
  // Is there a deletion operation before the last equality.
  var pre_del = false;
  // Is there an insertion operation after the last equality.
  var post_ins = false;
  // Is there a deletion operation after the last equality.
  var post_del = false;
  while (pointer < diffs.length) {
    if (diffs[pointer][0] == DIFF_EQUAL) {  // Equality found.
      if (diffs[pointer][1].length < this.EditCost &&
          (post_ins || post_del)) {
        // Candidate found.
        equalities[equalitiesLength++] = pointer;
        pre_ins = post_ins;
        pre_del = post_del;
        lastequality = diffs[pointer][1];
      } else {
        // Not a candidate, and can never become one.
        equalitiesLength = 0;
        lastequality = null;
      }
      post_ins = post_del = false;
    } else {  // An insertion or deletion.
      if (diffs[pointer][0] == DIFF_DELETE) {
        post_del = true;
      } else {
        post_ins = true;
      }
      /*
       * Five types to be split:
       * <ins>A</ins><del>B</del>XY<ins>C</ins><del>D</del>
       * <ins>A</ins>X<ins>C</ins><del>D</del>
       * <ins>A</ins><del>B</del>X<ins>C</ins>
       * <ins>A</del>X<ins>C</ins><del>D</del>
       * <ins>A</ins><del>B</del>X<del>C</del>
       */
      if (lastequality && ((pre_ins && pre_del && post_ins && post_del) ||
                           ((lastequality.length < this.EditCost / 2) &&
                            (pre_ins + pre_del + post_ins + post_del) == 3))) {
        // Duplicate record.
        diffs.splice(equalities[equalitiesLength - 1], 0,
                     [DIFF_DELETE, lastequality]);
        // Change second copy to insert.
        diffs[equalities[equalitiesLength - 1] + 1][0] = DIFF_INSERT;
        equalitiesLength--;  // Throw away the equality we just deleted;
        lastequality = null;
        if (pre_ins && pre_del) {
          // No changes made which could affect previous entry, keep going.
          post_ins = post_del = true;
          equalitiesLength = 0;
        } else {
          equalitiesLength--;  // Throw away the previous equality.
          pointer = equalitiesLength > 0 ?
              equalities[equalitiesLength - 1] : -1;
          post_ins = post_del = false;
        }
        changes = true;
      }
    }
    pointer++;
  }

  if (changes) {
    this.cleanupMerge(diffs);
  }
};


/**
 * Reorder and merge like edit sections.  Merge equalities.
 * Any edit section can move as long as it doesn't cross an equality.
 * @param {!Array.<!diff.Diff>} diffs Array of diff tuples.
 */
diff.prototype.cleanupMerge = function(diffs) {
  diffs.push([DIFF_EQUAL, '']);  // Add a dummy entry at the end.
  var pointer = 0;
  var count_delete = 0;
  var count_insert = 0;
  var text_delete = '';
  var text_insert = '';
  var commonlength;
  while (pointer < diffs.length) {
    switch (diffs[pointer][0]) {
      case DIFF_INSERT:
        count_insert++;
        text_insert += diffs[pointer][1];
        pointer++;
        break;
      case DIFF_DELETE:
        count_delete++;
        text_delete += diffs[pointer][1];
        pointer++;
        break;
      case DIFF_EQUAL:
        // Upon reaching an equality, check for prior redundancies.
        if (count_delete + count_insert > 1) {
          if (count_delete !== 0 && count_insert !== 0) {
            // Factor out any common prefixies.
            commonlength = this.commonPrefix(text_insert, text_delete);
            if (commonlength !== 0) {
              if ((pointer - count_delete - count_insert) > 0 &&
                  diffs[pointer - count_delete - count_insert - 1][0] ==
                  DIFF_EQUAL) {
                diffs[pointer - count_delete - count_insert - 1][1] +=
                    text_insert.substring(0, commonlength);
              } else {
                diffs.splice(0, 0, [DIFF_EQUAL,
                                    text_insert.substring(0, commonlength)]);
                pointer++;
              }
              text_insert = text_insert.substring(commonlength);
              text_delete = text_delete.substring(commonlength);
            }
            // Factor out any common suffixies.
            commonlength = this.commonSuffix(text_insert, text_delete);
            if (commonlength !== 0) {
              diffs[pointer][1] = text_insert.substring(text_insert.length -
                  commonlength) + diffs[pointer][1];
              text_insert = text_insert.substring(0, text_insert.length -
                  commonlength);
              text_delete = text_delete.substring(0, text_delete.length -
                  commonlength);
            }
          }
          // Delete the offending records and add the merged ones.
          if (count_delete === 0) {
            diffs.splice(pointer - count_insert,
                count_delete + count_insert, [DIFF_INSERT, text_insert]);
          } else if (count_insert === 0) {
            diffs.splice(pointer - count_delete,
                count_delete + count_insert, [DIFF_DELETE, text_delete]);
          } else {
            diffs.splice(pointer - count_delete - count_insert,
                count_delete + count_insert, [DIFF_DELETE, text_delete],
                [DIFF_INSERT, text_insert]);
          }
          pointer = pointer - count_delete - count_insert +
                    (count_delete ? 1 : 0) + (count_insert ? 1 : 0) + 1;
        } else if (pointer !== 0 && diffs[pointer - 1][0] == DIFF_EQUAL) {
          // Merge this equality with the previous one.
          diffs[pointer - 1][1] += diffs[pointer][1];
          diffs.splice(pointer, 1);
        } else {
          pointer++;
        }
        count_insert = 0;
        count_delete = 0;
        text_delete = '';
        text_insert = '';
        break;
    }
  }
  if (diffs[diffs.length - 1][1] === '') {
    diffs.pop();  // Remove the dummy entry at the end.
  }

  // Second pass: look for single edits surrounded on both sides by equalities
  // which can be shifted sideways to eliminate an equality.
  // e.g: A<ins>BA</ins>C -> <ins>AB</ins>AC
  var changes = false;
  pointer = 1;
  // Intentionally ignore the first and last element (don't need checking).
  while (pointer < diffs.length - 1) {
    if (diffs[pointer - 1][0] == DIFF_EQUAL &&
        diffs[pointer + 1][0] == DIFF_EQUAL) {
      // This is a single edit surrounded by equalities.
      if (diffs[pointer][1].substring(diffs[pointer][1].length -
          diffs[pointer - 1][1].length) == diffs[pointer - 1][1]) {
        // Shift the edit over the previous equality.
        diffs[pointer][1] = diffs[pointer - 1][1] +
            diffs[pointer][1].substring(0, diffs[pointer][1].length -
                                        diffs[pointer - 1][1].length);
        diffs[pointer + 1][1] = diffs[pointer - 1][1] + diffs[pointer + 1][1];
        diffs.splice(pointer - 1, 1);
        changes = true;
      } else if (diffs[pointer][1].substring(0, diffs[pointer + 1][1].length) ==
          diffs[pointer + 1][1]) {
        // Shift the edit over the next equality.
        diffs[pointer - 1][1] += diffs[pointer + 1][1];
        diffs[pointer][1] =
            diffs[pointer][1].substring(diffs[pointer + 1][1].length) +
            diffs[pointer + 1][1];
        diffs.splice(pointer + 1, 1);
        changes = true;
      }
    }
    pointer++;
  }
  // If shifts were made, the diff needs reordering and another shift sweep.
  if (changes) {
    this.cleanupMerge(diffs);
  }
};


/**
 * loc is a location in text1, compute and return the equivalent location in
 * text2.
 * e.g. 'The cat' vs 'The big cat', 1->1, 5->8
 * @param {!Array.<!diff.Diff>} diffs Array of diff tuples.
 * @param {number} loc Location within text1.
 * @return {number} Location within text2.
 */
diff.prototype.xIndex = function(diffs, loc) {
  var chars1 = 0;
  var chars2 = 0;
  var last_chars1 = 0;
  var last_chars2 = 0;
  var x;
  for (x = 0; x < diffs.length; x++) {
    if (diffs[x][0] !== DIFF_INSERT) {  // Equality or deletion.
      chars1 += diffs[x][1].length;
    }
    if (diffs[x][0] !== DIFF_DELETE) {  // Equality or insertion.
      chars2 += diffs[x][1].length;
    }
    if (chars1 > loc) {  // Overshot the location.
      break;
    }
    last_chars1 = chars1;
    last_chars2 = chars2;
  }
  // Was the location was deleted?
  if (diffs.length != x && diffs[x][0] === DIFF_DELETE) {
    return last_chars2;
  }
  // Add the remaining character length.
  return last_chars2 + (loc - last_chars1);
};


/**
 * Convert a diff array into a pretty HTML report.
 * @param {!Array.<!diff.Diff>} diffs Array of diff tuples.
 * @return {string} HTML representation.
 */
diff.prototype.prettyHtml = function(diffs) {
  var html = [];
  var pattern_amp = /&/g;
  var pattern_lt = /</g;
  var pattern_gt = />/g;
  var pattern_br = /\n/g;
  for (var x = 0; x < diffs.length; x++) {
    var op = diffs[x][0];    // Operation (insert, delete, equal)
    var data = diffs[x][1];  // Text of change.
    var text = data.replace(pattern_amp, '&amp;').replace(pattern_lt, '&lt;')
        .replace(pattern_gt, '&gt;').replace(pattern_br, '<br/>');
    switch (op) {
      case DIFF_INSERT:
        html[x] = '<ins>' + text + '</ins>';
        break;
      case DIFF_DELETE:
        html[x] = '<del>' + text + '</del>';
        break;
      case DIFF_EQUAL:
        html[x] = '<span>' + text + '</span>';
        break;
    }
  }
  return html.join('');
};


/**
 * Compute and return the source text (all equalities and deletions).
 * @param {!Array.<!diff.Diff>} diffs Array of diff tuples.
 * @return {string} Source text.
 */
diff.prototype.text1 = function(diffs) {
  var text = [];
  for (var x = 0; x < diffs.length; x++) {
    if (diffs[x][0] !== DIFF_INSERT) {
      text[x] = diffs[x][1];
    }
  }
  return text.join('');
};


/**
 * Compute and return the destination text (all equalities and insertions).
 * @param {!Array.<!diff.Diff>} diffs Array of diff tuples.
 * @return {string} Destination text.
 */
diff.prototype.text2 = function(diffs) {
  var text = [];
  for (var x = 0; x < diffs.length; x++) {
    if (diffs[x][0] !== DIFF_DELETE) {
      text[x] = diffs[x][1];
    }
  }
  return text.join('');
};


/**
 * Compute the Levenshtein distance; the number of inserted, deleted or
 * substituted characters.
 * @param {!Array.<!diff.Diff>} diffs Array of diff tuples.
 * @return {number} Number of changes.
 */
diff.prototype.levenshtein = function(diffs) {
  var levenshtein = 0;
  var insertions = 0;
  var deletions = 0;
  for (var x = 0; x < diffs.length; x++) {
    var op = diffs[x][0];
    var data = diffs[x][1];
    switch (op) {
      case DIFF_INSERT:
        insertions += data.length;
        break;
      case DIFF_DELETE:
        deletions += data.length;
        break;
      case DIFF_EQUAL:
        // A deletion and an insertion is one substitution.
        levenshtein += Math.max(insertions, deletions);
        insertions = 0;
        deletions = 0;
        break;
    }
  }
  levenshtein += Math.max(insertions, deletions);
  return levenshtein;
};


/**
 * Crush the diff into an encoded string which describes the operations
 * required to transform text1 into text2.
 * E.g. =3\t-2\t+ing  -> Keep 3 chars, delete 2 chars, insert 'ing'.
 * Operations are tab-separated.  Inserted text is escaped using %xx notation.
 * @param {!Array.<!diff.Diff>} diffs Array of diff tuples.
 * @return {string} Delta text.
 */
diff.prototype.toDelta = function(diffs) {
  var text = [];
  for (var x = 0; x < diffs.length; x++) {
    switch (diffs[x][0]) {
      case DIFF_INSERT:
        text[x] = '+' + encodeURI(diffs[x][1]);
        break;
      case DIFF_DELETE:
        text[x] = '-' + diffs[x][1].length;
        break;
      case DIFF_EQUAL:
        text[x] = '=' + diffs[x][1].length;
        break;
    }
  }
  return text.join('\t').replace(/%20/g, ' ');
};


/**
 * Given the original text1, and an encoded string which describes the
 * operations required to transform text1 into text2, compute the full diff.
 * @param {string} text1 Source string for the diff.
 * @param {string} delta Delta text.
 * @return {!Array.<!diff.Diff>} Array of diff tuples.
 * @throws {!Error} If invalid input.
 */
diff.prototype.fromDelta = function(text1, delta) {
  var diffs = [];
  var diffsLength = 0;  // Keeping our own length var is faster in JS.
  var pointer = 0;  // Cursor in text1
  var tokens = delta.split(/\t/g);
  for (var x = 0; x < tokens.length; x++) {
    // Each token begins with a one character parameter which specifies the
    // operation of this token (delete, insert, equality).
    var param = tokens[x].substring(1);
    switch (tokens[x].charAt(0)) {
      case '+':
        try {
          diffs[diffsLength++] = [DIFF_INSERT, decodeURI(param)];
        } catch (ex) {
          // Malformed URI sequence.
          throw new Error('Illegal escape in diff_fromDelta: ' + param);
        }
        break;
      case '-':
        // Fall through.
      case '=':
        var n = parseInt(param, 10);
        if (isNaN(n) || n < 0) {
          throw new Error('Invalid number in diff_fromDelta: ' + param);
        }
        var text = text1.substring(pointer, pointer += n);
        if (tokens[x].charAt(0) == '=') {
          diffs[diffsLength++] = [DIFF_EQUAL, text];
        } else {
          diffs[diffsLength++] = [DIFF_DELETE, text];
        }
        break;
      default:
        // Blank tokens are ok (from a trailing \t).
        // Anything else is an error.
        if (tokens[x]) {
          throw new Error('Invalid diff operation in diff_fromDelta: ' +
                          tokens[x]);
        }
    }
  }
  if (pointer != text1.length) {
    throw new Error('Delta length (' + pointer +
        ') does not equal source text length (' + text1.length + ').');
  }
  return diffs;
};


// Export these global variables so that they survive Google's JS compiler.
// In a browser, 'this' will be 'window'.
// Users of node.js should 'require' the uncompressed version since Google's
// JS compiler may break the following exports for non-browser environments.
this['diff'] = diff;
this['DIFF_DELETE'] = DIFF_DELETE;
this['DIFF_INSERT'] = DIFF_INSERT;
this['DIFF_EQUAL'] = DIFF_EQUAL;
```
