## Supplementary material for "Method for inferring migration routes from phylogenetic trees": Some route-trees: FileSaver.js

```
(function (global, factory) {
  if (typeof define === "function" && define.amd) {
    define([], factory);
  } else if (typeof exports !== "undefined") {
    factory();
  } else {
    var mod = {
      exports: {}
    };
    factory();
    global.FileSaver = mod.exports;
  }
})(this, function () {
  "use strict";

  /*
  * FileSaver.js
  * A saveAs() FileSaver implementation.
  *
  * By Eli Grey, http://eligrey.com
  *
  * License : https://github.com/eligrey/FileSaver.js/blob/master/LICENSE.md (MIT)
  * source  : http://purl.eligrey.com/github/FileSaver.js
  */
  // The one and only way of getting global scope in all environments
  // https://stackoverflow.com/q/3277182/1008999
  var _global = typeof window === 'object' && window.window === window ? window : typeof self === 'object' && self.self === self ? self : typeof global === 'object' && global.global === global ? global : void 0;

  function bom(blob, opts) {
    if (typeof opts === 'undefined') opts = {
      autoBom: false
    };else if (typeof opts !== 'object') {
      console.warn('Deprecated: Expected third argument to be a object');
      opts = {
        autoBom: !opts
      };
    } // prepend BOM for UTF-8 XML and text/* types (including HTML)
    // note: your browser will automatically convert UTF-16 U+FEFF to EF BB BF

    if (opts.autoBom && /^\s*(?:text\/\S*|application\/xml|\S*\/\S*\+xml)\s*;.*charset\s*=\s*utf-8/i.test(blob.type)) {
      return new Blob([String.fromCharCode(0xFEFF), blob], {
        type: blob.type
      });
    }

    return blob;
  }

  function download(url, name, opts) {
    var xhr = new XMLHttpRequest();
    xhr.open('GET', url);
    xhr.responseType = 'blob';

    xhr.onload = function () {
      saveAs(xhr.response, name, opts);
    };

    xhr.onerror = function () {
      console.error('could not download file');
    };

    xhr.send();
  }

  function corsEnabled(url) {
    var xhr = new XMLHttpRequest(); // use sync to avoid popup blocker

    xhr.open('HEAD', url, false);

    try {
      xhr.send();
    } catch (e) {}

    return xhr.status >= 200 && xhr.status <= 299;
  } // `a.click()` doesn't work for all browsers (#465)


  function click(node) {
    try {
      node.dispatchEvent(new MouseEvent('click'));
    } catch (e) {
      var evt = document.createEvent('MouseEvents');
      evt.initMouseEvent('click', true, true, window, 0, 0, 0, 80, 20, false, false, false, false, 0, null);
      node.dispatchEvent(evt);
    }
  }

  var saveAs = _global.saveAs || ( // probably in some web worker
  typeof window !== 'object' || window !== _global ? function saveAs() {}
  /* noop */
  // Use download attribute first if possible (#193 Lumia mobile)
  : 'download' in HTMLAnchorElement.prototype ? function saveAs(blob, name, opts) {
    var URL = _global.URL || _global.webkitURL;
    var a = document.createElement('a');
    name = name || blob.name || 'download';
    a.download = name;
    a.rel = 'noopener'; // tabnabbing
    // TODO: detect chrome extensions & packaged apps
    // a.target = '_blank'

    if (typeof blob === 'string') {
      // Support regular links
      a.href = blob;

      if (a.origin !== location.origin) {
        corsEnabled(a.href) ? download(blob, name, opts) : click(a, a.target = '_blank');
      } else {
        click(a);
      }
    } else {
      // Support blobs
      a.href = URL.createObjectURL(blob);
      setTimeout(function () {
        URL.revokeObjectURL(a.href);
      }, 4E4); // 40s

      setTimeout(function () {
        click(a);
      }, 0);
    }
  } // Use msSaveOrOpenBlob as a second approach
  : 'msSaveOrOpenBlob' in navigator ? function saveAs(blob, name, opts) {
    name = name || blob.name || 'download';

    if (typeof blob === 'string') {
      if (corsEnabled(blob)) {
        download(blob, name, opts);
      } else {
        var a = document.createElement('a');
        a.href = blob;
        a.target = '_blank';
        setTimeout(function () {
          click(a);
        });
      }
    } else {
      navigator.msSaveOrOpenBlob(bom(blob, opts), name);
    }
  } // Fallback to using FileReader and a popup
  : function saveAs(blob, name, opts, popup) {
    // Open a popup immediately do go around popup blocker
    // Mostly only available on user interaction and the fileReader is async so...
    popup = popup || open('', '_blank');

    if (popup) {
      popup.document.title = popup.document.body.innerText = 'downloading...';
    }

    if (typeof blob === 'string') return download(blob, name, opts);
    var force = blob.type === 'application/octet-stream';

    var isSafari = /constructor/i.test(_global.HTMLElement) || _global.safari;

    var isChromeIOS = /CriOS\/[\d]+/.test(navigator.userAgent);

    if ((isChromeIOS || force && isSafari) && typeof FileReader !== 'undefined') {
      // Safari doesn't allow downloading of blob URLs
      var reader = new FileReader();

      reader.onloadend = function () {
        var url = reader.result;
        url = isChromeIOS ? url : url.replace(/^data:[^;]*;/, 'data:attachment/file;');
        if (popup) popup.location.href = url;else location = url;
        popup = null; // reverse-tabnabbing #460
      };

      reader.readAsDataURL(blob);
    } else {
      var URL = _global.URL || _global.webkitURL;
      var url = URL.createObjectURL(blob);
      if (popup) popup.location = url;else location.href = url;
      popup = null; // reverse-tabnabbing #460

      setTimeout(function () {
        URL.revokeObjectURL(url);
      }, 4E4); // 40s
    }
  });
  _global.saveAs = saveAs.saveAs = saveAs;

  if (typeof module !== 'undefined') {
    module.exports = saveAs;
  }
});
```
