## Supplementary material for "Method for inferring migration routes from phylogenetic trees": Some route-trees: heatmap.js

```
function factorial(numero) {
	resultado = 1;
	for (var i = 1; i <= numero; i++) {
		resultado *= i;
	}
	return resultado;
}

function hamming_distance(sA, sB) {
  let result = 0

  if (sA.length == sB.length) {

      for (let i = 0; i < sA.length; i++) {
          if (sA[i].toLowerCase() != sB[i].toLowerCase()) {
              result++
          }
      }

      return result

  } else {
      throw new Error('Strings do not have equal length')
  }
}

function get_array_colors(dna1,dna2,genome_length,L0,ini)
{
    var colors=Array();
    for(var i=0;i<ini;i++)
    {
        colors[i]=-1;
    }
    for(var i=0;i<(dna1.length-ini)/L0;i++)
    {
        var w1=dna1.substring(ini+i*L0,ini+i*L0+L0);
        var w2=dna2.substring(ini+i*L0,ini+i*L0+L0);
        var L=w1.length;
        var hd=hamming_distance(w1,w2);
        if(hd>=L-1)
            hd-=10;
        var pos=Math.pow(4,L);
        var fav=Math.pow(3,hd)*factorial(L)/(factorial(hd)*factorial(L-hd));
        var p=fav/pos;
        var pazar=1;
        try{
            pazar=1-Math.pow(1-p,genome_length/L);
        }
        catch(e){}
        var color=511*pazar;
        for(var j=0;j<L;j++)
            colors.push(color);
    }

    return colors;
}

function get_heatmap_randomness(dna1,dna2,species,L0)
{
    var res="";
    var genome_length=3000000000;
    switch(species)
    {
        case 'mouse':
        case 'dog':
        case 'chimpanzee':
            genome_length=2500000000;
            break;
    }

    var acolors=Array();
    

    acolors.push(get_array_colors(dna1,dna2,genome_length,L0,0));
    acolors.push(get_array_colors(dna1,dna2,genome_length,L0,L0/8));
    acolors.push(get_array_colors(dna1,dna2,genome_length,L0,2*L0/8));
    acolors.push(get_array_colors(dna1,dna2,genome_length,L0,3*L0/8));
    acolors.push(get_array_colors(dna1,dna2,genome_length,L0,4*L0/8));
    acolors.push(get_array_colors(dna1,dna2,genome_length,L0,5*L0/8));
    acolors.push(get_array_colors(dna1,dna2,genome_length,L0,6*L0/8));
    acolors.push(get_array_colors(dna1,dna2,genome_length,L0,7*L0/8));

    
    
    for(var i=0;i<dna1.length;i++)
    {
        var spaces=' ';
        var color=0;
        var n=0;
        for(var j=0;j<acolors.length;j++)
        {
            if(acolors[j][i]!=-1)
            {
                if(color=='')
                    color=parseFloat(acolors[j][i]);
                else
                    color=parseFloat(color)+parseFloat(acolors[j][i]);
                n++;
            }
        }
        color=color/n;

        if(color<256)
        {
            colorg=255-parseInt(color);
            var hex = ('00' + colorg.toString(16).toUpperCase()).slice(-2);
            res+="<span style=\"background-color:#00"+hex+"00\">"+spaces+"</span>";
        }
        else            
        {
            colorg=parseInt(color)-256;
            var hex = ('00' + colorg.toString(16).toUpperCase()).slice(-2);            
            res+="<span style=\"background-color:#"+hex+"0000\">"+spaces+"</span>";
        }

    }
    return res;
}

function get_match_map(dna1,dna2)
{
    var res='';
    for(var i=0;i<dna1.length;i++)
    {
        if(dna1[i]==dna2[i])
            res+='|';
        else
            res+=' ';
    }
    return res;
}
```
