## Supplementary material for "Method for inferring migration routes from phylogenetic trees": Some route-trees: highlight.min.js

````
/*! highlight.js v9.12.0 | BSD3 License | git.io/hljslicense */
!function(e){var t="object"==typeof window&&window||"object"==typeof self&&self;"undefined"!=typeof exports?e(exports):t&&(t.hljs=e({}),"function"==typeof define&&define.amd&&define([],function(){return t.hljs}))}(function(e){function t(e){return e.replace(/&/g,"&amp;").replace(/</g,"&lt;").replace(/>/g,"&gt;")}function r(e){return e.nodeName.toLowerCase()}function a(e,t){var r=e&&e.exec(t);return r&&0===r.index}function n(e){return E.test(e)}function i(e){var t,r,a,i,s=e.className+" ";if(s+=e.parentNode?e.parentNode.className:"",r=M.exec(s))return w(r[1])?r[1]:"no-highlight";for(s=s.split(/\s+/),t=0,a=s.length;a>t;t++)if(i=s[t],n(i)||w(i))return i}function s(e){var t,r={},a=Array.prototype.slice.call(arguments,1);for(t in e)r[t]=e[t];return a.forEach(function(e){for(t in e)r[t]=e[t]}),r}function c(e){var t=[];return function a(e,n){for(var i=e.firstChild;i;i=i.nextSibling)3===i.nodeType?n+=i.nodeValue.length:1===i.nodeType&&(t.push({event:"start",offset:n,node:i}),n=a(i,n),r(i).match(/br|hr|img|input/)||t.push({event:"stop",offset:n,node:i}));return n}(e,0),t}function o(e,a,n){function i(){return e.length&&a.length?e[0].offset!==a[0].offset?e[0].offset<a[0].offset?e:a:"start"===a[0].event?e:a:e.length?e:a}function s(e){function a(e){return" "+e.nodeName+'="'+t(e.value).replace('"',"&quot;")+'"'}u+="<"+r(e)+N.map.call(e.attributes,a).join("")+">"}function c(e){u+="</"+r(e)+">"}function o(e){("start"===e.event?s:c)(e.node)}for(var l=0,u="",d=[];e.length||a.length;){var b=i();if(u+=t(n.substring(l,b[0].offset)),l=b[0].offset,b===e){d.reverse().forEach(c);do o(b.splice(0,1)[0]),b=i();while(b===e&&b.length&&b[0].offset===l);d.reverse().forEach(s)}else"start"===b[0].event?d.push(b[0].node):d.pop(),o(b.splice(0,1)[0])}return u+t(n.substr(l))}function l(e){return e.v&&!e.cached_variants&&(e.cached_variants=e.v.map(function(t){return s(e,{v:null},t)})),e.cached_variants||e.eW&&[s(e)]||[e]}function u(e){function t(e){return e&&e.source||e}function r(r,a){return new RegExp(t(r),"m"+(e.cI?"i":"")+(a?"g":""))}function a(n,i){if(!n.compiled){if(n.compiled=!0,n.k=n.k||n.bK,n.k){var s={},c=function(t,r){e.cI&&(r=r.toLowerCase()),r.split(" ").forEach(function(e){var r=e.split("|");s[r[0]]=[t,r[1]?Number(r[1]):1]})};"string"==typeof n.k?c("keyword",n.k):k(n.k).forEach(function(e){c(e,n.k[e])}),n.k=s}n.lR=r(n.l||/\w+/,!0),i&&(n.bK&&(n.b="\\b("+n.bK.split(" ").join("|")+")\\b"),n.b||(n.b=/\B|\b/),n.bR=r(n.b),n.e||n.eW||(n.e=/\B|\b/),n.e&&(n.eR=r(n.e)),n.tE=t(n.e)||"",n.eW&&i.tE&&(n.tE+=(n.e?"|":"")+i.tE)),n.i&&(n.iR=r(n.i)),null==n.r&&(n.r=1),n.c||(n.c=[]),n.c=Array.prototype.concat.apply([],n.c.map(function(e){return l("self"===e?n:e)})),n.c.forEach(function(e){a(e,n)}),n.starts&&a(n.starts,i);var o=n.c.map(function(e){return e.bK?"\\.?("+e.b+")\\.?":e.b}).concat([n.tE,n.i]).map(t).filter(Boolean);n.t=o.length?r(o.join("|"),!0):{exec:function(){return null}}}}a(e)}function d(e,r,n,i){function s(e,t){var r,n;for(r=0,n=t.c.length;n>r;r++)if(a(t.c[r].bR,e))return t.c[r]}function c(e,t){if(a(e.eR,t)){for(;e.endsParent&&e.parent;)e=e.parent;return e}return e.eW?c(e.parent,t):void 0}function o(e,t){return!n&&a(t.iR,e)}function l(e,t){var r=v.cI?t[0].toLowerCase():t[0];return e.k.hasOwnProperty(r)&&e.k[r]}function p(e,t,r,a){var n=a?"":L.classPrefix,i='<span class="'+n,s=r?"":R;return i+=e+'">',i+t+s}function m(){var e,r,a,n;if(!N.k)return t(E);for(n="",r=0,N.lR.lastIndex=0,a=N.lR.exec(E);a;)n+=t(E.substring(r,a.index)),e=l(N,a),e?(M+=e[1],n+=p(e[0],t(a[0]))):n+=t(a[0]),r=N.lR.lastIndex,a=N.lR.exec(E);return n+t(E.substr(r))}function f(){var e="string"==typeof N.sL;if(e&&!x[N.sL])return t(E);var r=e?d(N.sL,E,!0,k[N.sL]):b(E,N.sL.length?N.sL:void 0);return N.r>0&&(M+=r.r),e&&(k[N.sL]=r.top),p(r.language,r.value,!1,!0)}function g(){C+=null!=N.sL?f():m(),E=""}function _(e){C+=e.cN?p(e.cN,"",!0):"",N=Object.create(e,{parent:{value:N}})}function h(e,t){if(E+=e,null==t)return g(),0;var r=s(t,N);if(r)return r.skip?E+=t:(r.eB&&(E+=t),g(),r.rB||r.eB||(E=t)),_(r,t),r.rB?0:t.length;var a=c(N,t);if(a){var n=N;n.skip?E+=t:(n.rE||n.eE||(E+=t),g(),n.eE&&(E=t));do N.cN&&(C+=R),N.skip||(M+=N.r),N=N.parent;while(N!==a.parent);return a.starts&&_(a.starts,""),n.rE?0:t.length}if(o(t,N))throw new Error('Illegal lexeme "'+t+'" for mode "'+(N.cN||"<unnamed>")+'"');return E+=t,t.length||1}var v=w(e);if(!v)throw new Error('Unknown language: "'+e+'"');u(v);var y,N=i||v,k={},C="";for(y=N;y!==v;y=y.parent)y.cN&&(C=p(y.cN,"",!0)+C);var E="",M=0;try{for(var B,S,$=0;;){if(N.t.lastIndex=$,B=N.t.exec(r),!B)break;S=h(r.substring($,B.index),B[0]),$=B.index+S}for(h(r.substr($)),y=N;y.parent;y=y.parent)y.cN&&(C+=R);return{r:M,value:C,language:e,top:N}}catch(A){if(A.message&&-1!==A.message.indexOf("Illegal"))return{r:0,value:t(r)};throw A}}function b(e,r){r=r||L.languages||k(x);var a={r:0,value:t(e)},n=a;return r.filter(w).forEach(function(t){var r=d(t,e,!1);r.language=t,r.r>n.r&&(n=r),r.r>a.r&&(n=a,a=r)}),n.language&&(a.second_best=n),a}function p(e){return L.tabReplace||L.useBR?e.replace(B,function(e,t){return L.useBR&&"\n"===e?"<br>":L.tabReplace?t.replace(/\t/g,L.tabReplace):""}):e}function m(e,t,r){var a=t?C[t]:r,n=[e.trim()];return e.match(/\bhljs\b/)||n.push("hljs"),-1===e.indexOf(a)&&n.push(a),n.join(" ").trim()}function f(e){var t,r,a,s,l,u=i(e);n(u)||(L.useBR?(t=document.createElementNS("http://www.w3.org/1999/xhtml","div"),t.innerHTML=e.innerHTML.replace(/\n/g,"").replace(/<br[ \/]*>/g,"\n")):t=e,l=t.textContent,a=u?d(u,l,!0):b(l),r=c(t),r.length&&(s=document.createElementNS("http://www.w3.org/1999/xhtml","div"),s.innerHTML=a.value,a.value=o(r,c(s),l)),a.value=p(a.value),e.innerHTML=a.value,e.className=m(e.className,u,a.language),e.result={language:a.language,re:a.r},a.second_best&&(e.second_best={language:a.second_best.language,re:a.second_best.r}))}function g(e){L=s(L,e)}function _(){if(!_.called){_.called=!0;var e=document.querySelectorAll("pre code");N.forEach.call(e,f)}}function h(){addEventListener("DOMContentLoaded",_,!1),addEventListener("load",_,!1)}function v(t,r){var a=x[t]=r(e);a.aliases&&a.aliases.forEach(function(e){C[e]=t})}function y(){return k(x)}function w(e){return e=(e||"").toLowerCase(),x[e]||x[C[e]]}var N=[],k=Object.keys,x={},C={},E=/^(no-?highlight|plain|text)$/i,M=/\blang(?:uage)?-([\w-]+)\b/i,B=/((^(<[^>]+>|\t|)+|(?:\n)))/gm,R="</span>",L={classPrefix:"hljs-",tabReplace:null,useBR:!1,languages:void 0};return e.highlight=d,e.highlightAuto=b,e.fixMarkup=p,e.highlightBlock=f,e.configure=g,e.initHighlighting=_,e.initHighlightingOnLoad=h,e.registerLanguage=v,e.listLanguages=y,e.getLanguage=w,e.inherit=s,e.IR="[a-zA-Z]\\w*",e.UIR="[a-zA-Z_]\\w*",e.NR="\\b\\d+(\\.\\d+)?",e.CNR="(-?)(\\b0[xX][a-fA-F0-9]+|(\\b\\d+(\\.\\d*)?|\\.\\d+)([eE][-+]?\\d+)?)",e.BNR="\\b(0b[01]+)",e.RSR="!|!=|!==|%|%=|&|&&|&=|\\*|\\*=|\\+|\\+=|,|-|-=|/=|/|:|;|<<|<<=|<=|<|===|==|=|>>>=|>>=|>=|>>>|>>|>|\\?|\\[|\\{|\\(|\\^|\\^=|\\||\\|=|\\|\\||~",e.BE={b:"\\\\[\\s\\S]",r:0},e.ASM={cN:"string",b:"'",e:"'",i:"\\n",c:[e.BE]},e.QSM={cN:"string",b:'"',e:'"',i:"\\n",c:[e.BE]},e.PWM={b:/\b(a|an|the|are|I'm|isn't|don't|doesn't|won't|but|just|should|pretty|simply|enough|gonna|going|wtf|so|such|will|you|your|they|like|more)\b/},e.C=function(t,r,a){var n=e.inherit({cN:"comment",b:t,e:r,c:[]},a||{});return n.c.push(e.PWM),n.c.push({cN:"doctag",b:"(?:TODO|FIXME|NOTE|BUG|XXX):",r:0}),n},e.CLCM=e.C("//","$"),e.CBCM=e.C("/\\*","\\*/"),e.HCM=e.C("#","$"),e.NM={cN:"number",b:e.NR,r:0},e.CNM={cN:"number",b:e.CNR,r:0},e.BNM={cN:"number",b:e.BNR,r:0},e.CSSNM={cN:"number",b:e.NR+"(%|em|ex|ch|rem|vw|vh|vmin|vmax|cm|mm|in|pt|pc|px|deg|grad|rad|turn|s|ms|Hz|kHz|dpi|dpcm|dppx)?",r:0},e.RM={cN:"regexp",b:/\//,e:/\/[gimuy]*/,i:/\n/,c:[e.BE,{b:/\[/,e:/\]/,r:0,c:[e.BE]}]},e.TM={cN:"title",b:e.IR,r:0},e.UTM={cN:"title",b:e.UIR,r:0},e.METHOD_GUARD={b:"\\.\\s*"+e.UIR,r:0},e.registerLanguage("apache",function(e){var t={cN:"number",b:"[\\$%]\\d+"};return{aliases:["apacheconf"],cI:!0,c:[e.HCM,{cN:"section",b:"</?",e:">"},{cN:"attribute",b:/\w+/,r:0,k:{nomarkup:"order deny allow setenv rewriterule rewriteengine rewritecond documentroot sethandler errordocument loadmodule options header listen serverroot servername"},starts:{e:/$/,r:0,k:{literal:"on off all"},c:[{cN:"meta",b:"\\s\\[",e:"\\]$"},{cN:"variable",b:"[\\$%]\\{",e:"\\}",c:["self",t]},t,e.QSM]}}],i:/\S/}}),e.registerLanguage("bash",function(e){var t={cN:"variable",v:[{b:/\$[\w\d#@][\w\d_]*/},{b:/\$\{(.*?)}/}]},r={cN:"string",b:/"/,e:/"/,c:[e.BE,t,{cN:"variable",b:/\$\(/,e:/\)/,c:[e.BE]}]},a={cN:"string",b:/'/,e:/'/};return{aliases:["sh","zsh"],l:/\b-?[a-z\._]+\b/,k:{keyword:"if then else elif fi for while in do done case esac function",literal:"true false",built_in:"break cd continue eval exec exit export getopts hash pwd readonly return shift test times trap umask unset alias bind builtin caller command declare echo enable help let local logout mapfile printf read readarray source type typeset ulimit unalias set shopt autoload bg bindkey bye cap chdir clone comparguments compcall compctl compdescribe compfiles compgroups compquote comptags comptry compvalues dirs disable disown echotc echoti emulate fc fg float functions getcap getln history integer jobs kill limit log noglob popd print pushd pushln rehash sched setcap setopt stat suspend ttyctl unfunction unhash unlimit unsetopt vared wait whence where which zcompile zformat zftp zle zmodload zparseopts zprof zpty zregexparse zsocket zstyle ztcp",_:"-ne -eq -lt -gt -f -d -e -s -l -a"},c:[{cN:"meta",b:/^#![^\n]+sh\s*$/,r:10},{cN:"function",b:/\w[\w\d_]*\s*\(\s*\)\s*\{/,rB:!0,c:[e.inherit(e.TM,{b:/\w[\w\d_]*/})],r:0},e.HCM,r,a,t]}}),e.registerLanguage("coffeescript",function(e){var t={keyword:"in if for while finally new do return else break catch instanceof throw try this switch continue typeof delete debugger super yield import export from as default await then unless until loop of by when and or is isnt not",literal:"true false null undefined yes no on off",built_in:"npm require console print module global window document"},r="[A-Za-z$_][0-9A-Za-z$_]*",a={cN:"subst",b:/#\{/,e:/}/,k:t},n=[e.BNM,e.inherit(e.CNM,{starts:{e:"(\\s*/)?",r:0}}),{cN:"string",v:[{b:/'''/,e:/'''/,c:[e.BE]},{b:/'/,e:/'/,c:[e.BE]},{b:/"""/,e:/"""/,c:[e.BE,a]},{b:/"/,e:/"/,c:[e.BE,a]}]},{cN:"regexp",v:[{b:"///",e:"///",c:[a,e.HCM]},{b:"//[gim]*",r:0},{b:/\/(?![ *])(\\\/|.)*?\/[gim]*(?=\W|$)/}]},{b:"@"+r},{sL:"javascript",eB:!0,eE:!0,v:[{b:"```",e:"```"},{b:"`",e:"`"}]}];a.c=n;var i=e.inherit(e.TM,{b:r}),s="(\\(.*\\))?\\s*\\B[-=]>",c={cN:"params",b:"\\([^\\(]",rB:!0,c:[{b:/\(/,e:/\)/,k:t,c:["self"].concat(n)}]};return{aliases:["coffee","cson","iced"],k:t,i:/\/\*/,c:n.concat([e.C("###","###"),e.HCM,{cN:"function",b:"^\\s*"+r+"\\s*=\\s*"+s,e:"[-=]>",rB:!0,c:[i,c]},{b:/[:\(,=]\s*/,r:0,c:[{cN:"function",b:s,e:"[-=]>",rB:!0,c:[c]}]},{cN:"class",bK:"class",e:"$",i:/[:="\[\]]/,c:[{bK:"extends",eW:!0,i:/[:="\[\]]/,c:[i]},i]},{b:r+":",e:":",rB:!0,rE:!0,r:0}])}}),e.registerLanguage("cpp",function(e){var t={cN:"keyword",b:"\\b[a-z\\d_]*_t\\b"},r={cN:"string",v:[{b:'(u8?|U)?L?"',e:'"',i:"\\n",c:[e.BE]},{b:'(u8?|U)?R"',e:'"',c:[e.BE]},{b:"'\\\\?.",e:"'",i:"."}]},a={cN:"number",v:[{b:"\\b(0b[01']+)"},{b:"(-?)\\b([\\d']+(\\.[\\d']*)?|\\.[\\d']+)(u|U|l|L|ul|UL|f|F|b|B)"},{b:"(-?)(\\b0[xX][a-fA-F0-9']+|(\\b[\\d']+(\\.[\\d']*)?|\\.[\\d']+)([eE][-+]?[\\d']+)?)"}],r:0},n={cN:"meta",b:/#\s*[a-z]+\b/,e:/$/,k:{"meta-keyword":"if else elif endif define undef warning error line pragma ifdef ifndef include"},c:[{b:/\\\n/,r:0},e.inherit(r,{cN:"meta-string"}),{cN:"meta-string",b:/<[^\n>]*>/,e:/$/,i:"\\n"},e.CLCM,e.CBCM]},i=e.IR+"\\s*\\(",s={keyword:"int float while private char catch import module export virtual operator sizeof dynamic_cast|10 typedef const_cast|10 const for static_cast|10 union namespace unsigned long volatile static protected bool template mutable if public friend do goto auto void enum else break extern using asm case typeid short reinterpret_cast|10 default double register explicit signed typename try this switch continue inline delete alignof constexpr decltype noexcept static_assert thread_local restrict _Bool complex _Complex _Imaginary atomic_bool atomic_char atomic_schar atomic_uchar atomic_short atomic_ushort atomic_int atomic_uint atomic_long atomic_ulong atomic_llong atomic_ullong new throw return and or not",built_in:"std string cin cout cerr clog stdin stdout stderr stringstream istringstream ostringstream auto_ptr deque list queue stack vector map set bitset multiset multimap unordered_set unordered_map unordered_multiset unordered_multimap array shared_ptr abort abs acos asin atan2 atan calloc ceil cosh cos exit exp fabs floor fmod fprintf fputs free frexp fscanf isalnum isalpha iscntrl isdigit isgraph islower isprint ispunct isspace isupper isxdigit tolower toupper labs ldexp log10 log malloc realloc memchr memcmp memcpy memset modf pow printf putchar puts scanf sinh sin snprintf sprintf sqrt sscanf strcat strchr strcmp strcpy strcspn strlen strncat strncmp strncpy strpbrk strrchr strspn strstr tanh tan vfprintf vprintf vsprintf endl initializer_list unique_ptr",literal:"true false nullptr NULL"},c=[t,e.CLCM,e.CBCM,a,r];return{aliases:["c","cc","h","c++","h++","hpp"],k:s,i:"</",c:c.concat([n,{b:"\\b(deque|list|queue|stack|vector|map|set|bitset|multiset|multimap|unordered_map|unordered_set|unordered_multiset|unordered_multimap|array)\\s*<",e:">",k:s,c:["self",t]},{b:e.IR+"::",k:s},{v:[{b:/=/,e:/;/},{b:/\(/,e:/\)/},{bK:"new throw return else",e:/;/}],k:s,c:c.concat([{b:/\(/,e:/\)/,k:s,c:c.concat(["self"]),r:0}]),r:0},{cN:"function",b:"("+e.IR+"[\\*&\\s]+)+"+i,rB:!0,e:/[{;=]/,eE:!0,k:s,i:/[^\w\s\*&]/,c:[{b:i,rB:!0,c:[e.TM],r:0},{cN:"params",b:/\(/,e:/\)/,k:s,r:0,c:[e.CLCM,e.CBCM,r,a,t]},e.CLCM,e.CBCM,n]},{cN:"class",bK:"class struct",e:/[{;:]/,c:[{b:/</,e:/>/,c:["self"]},e.TM]}]),exports:{preprocessor:n,strings:r,k:s}}}),e.registerLanguage("cs",function(e){var t={keyword:"abstract as base bool break byte case catch char checked const continue decimal default delegate do double enum event explicit extern finally fixed float for foreach goto if implicit in int interface internal is lock long nameof object operator out override params private protected public readonly ref sbyte sealed short sizeof stackalloc static string struct switch this try typeof uint ulong unchecked unsafe ushort using virtual void volatile while add alias ascending async await by descending dynamic equals from get global group into join let on orderby partial remove select set value var where yield",literal:"null false true"},r={cN:"string",b:'@"',e:'"',c:[{b:'""'}]},a=e.inherit(r,{i:/\n/}),n={cN:"subst",b:"{",e:"}",k:t},i=e.inherit(n,{i:/\n/}),s={cN:"string",b:/\$"/,e:'"',i:/\n/,c:[{b:"{{"},{b:"}}"},e.BE,i]},c={cN:"string",b:/\$@"/,e:'"',c:[{b:"{{"},{b:"}}"},{b:'""'},n]},o=e.inherit(c,{i:/\n/,c:[{b:"{{"},{b:"}}"},{b:'""'},i]});n.c=[c,s,r,e.ASM,e.QSM,e.CNM,e.CBCM],i.c=[o,s,a,e.ASM,e.QSM,e.CNM,e.inherit(e.CBCM,{i:/\n/})];var l={v:[c,s,r,e.ASM,e.QSM]},u=e.IR+"(<"+e.IR+"(\\s*,\\s*"+e.IR+")*>)?(\\[\\])?";return{aliases:["csharp"],k:t,i:/::/,c:[e.C("///","$",{rB:!0,c:[{cN:"doctag",v:[{b:"///",r:0},{b:"<!--|-->"},{b:"</?",e:">"}]}]}),e.CLCM,e.CBCM,{cN:"meta",b:"#",e:"$",k:{"meta-keyword":"if else elif endif define undef warning error line region endregion pragma checksum"}},l,e.CNM,{bK:"class interface",e:/[{;=]/,i:/[^\s:]/,c:[e.TM,e.CLCM,e.CBCM]},{bK:"namespace",e:/[{;=]/,i:/[^\s:]/,c:[e.inherit(e.TM,{b:"[a-zA-Z](\\.?\\w)*"}),e.CLCM,e.CBCM]},{cN:"meta",b:"^\\s*\\[",eB:!0,e:"\\]",eE:!0,c:[{cN:"meta-string",b:/"/,e:/"/}]},{bK:"new return throw await else",r:0},{cN:"function",b:"("+u+"\\s+)+"+e.IR+"\\s*\\(",rB:!0,e:/[{;=]/,eE:!0,k:t,c:[{b:e.IR+"\\s*\\(",rB:!0,c:[e.TM],r:0},{cN:"params",b:/\(/,e:/\)/,eB:!0,eE:!0,k:t,r:0,c:[l,e.CNM,e.CBCM]},e.CLCM,e.CBCM]}]}}),e.registerLanguage("css",function(e){var t="[a-zA-Z-][a-zA-Z0-9_-]*",r={b:/[A-Z\_\.\-]+\s*:/,rB:!0,e:";",eW:!0,c:[{cN:"attribute",b:/\S/,e:":",eE:!0,starts:{eW:!0,eE:!0,c:[{b:/[\w-]+\(/,rB:!0,c:[{cN:"built_in",b:/[\w-]+/},{b:/\(/,e:/\)/,c:[e.ASM,e.QSM]}]},e.CSSNM,e.QSM,e.ASM,e.CBCM,{cN:"number",b:"#[0-9A-Fa-f]+"},{cN:"meta",b:"!important"}]}}]};return{cI:!0,i:/[=\/|'\$]/,c:[e.CBCM,{cN:"selector-id",b:/#[A-Za-z0-9_-]+/},{cN:"selector-class",b:/\.[A-Za-z0-9_-]+/},{cN:"selector-attr",b:/\[/,e:/\]/,i:"$"},{cN:"selector-pseudo",b:/:(:)?[a-zA-Z0-9\_\-\+\(\)"'.]+/},{b:"@(font-face|page)",l:"[a-z-]+",k:"font-face page"},{b:"@",e:"[{;]",i:/:/,c:[{cN:"keyword",b:/\w+/},{b:/\s/,eW:!0,eE:!0,r:0,c:[e.ASM,e.QSM,e.CSSNM]}]},{cN:"selector-tag",b:t,r:0},{b:"{",e:"}",i:/\S/,c:[e.CBCM,r]}]}}),e.registerLanguage("diff",function(e){return{aliases:["patch"],c:[{cN:"meta",r:10,v:[{b:/^@@ +\-\d+,\d+ +\+\d+,\d+ +@@$/},{b:/^\*\*\* +\d+,\d+ +\*\*\*\*$/},{b:/^\-\-\- +\d+,\d+ +\-\-\-\-$/}]},{cN:"comment",v:[{b:/Index: /,e:/$/},{b:/={3,}/,e:/$/},{b:/^\-{3}/,e:/$/},{b:/^\*{3} /,e:/$/},{b:/^\+{3}/,e:/$/},{b:/\*{5}/,e:/\*{5}$/}]},{cN:"addition",b:"^\\+",e:"$"},{cN:"deletion",b:"^\\-",e:"$"},{cN:"addition",b:"^\\!",e:"$"}]}}),e.registerLanguage("http",function(e){var t="HTTP/[0-9\\.]+";return{aliases:["https"],i:"\\S",c:[{b:"^"+t,e:"$",c:[{cN:"number",b:"\\b\\d{3}\\b"}]},{b:"^[A-Z]+ (.*?) "+t+"$",rB:!0,e:"$",c:[{cN:"string",b:" ",e:" ",eB:!0,eE:!0},{b:t},{cN:"keyword",b:"[A-Z]+"}]},{cN:"attribute",b:"^\\w",e:": ",eE:!0,i:"\\n|\\s|=",starts:{e:"$",r:0}},{b:"\\n\\n",starts:{sL:[],eW:!0}}]}}),e.registerLanguage("ini",function(e){var t={cN:"string",c:[e.BE],v:[{b:"'''",e:"'''",r:10},{b:'"""',e:'"""',r:10},{b:'"',e:'"'},{b:"'",e:"'"}]};return{aliases:["toml"],cI:!0,i:/\S/,c:[e.C(";","$"),e.HCM,{cN:"section",b:/^\s*\[+/,e:/\]+/},{b:/^[a-z0-9\[\]_-]+\s*=\s*/,e:"$",rB:!0,c:[{cN:"attr",b:/[a-z0-9\[\]_-]+/},{b:/=/,eW:!0,r:0,c:[{cN:"literal",b:/\bon|off|true|false|yes|no\b/},{cN:"variable",v:[{b:/\$[\w\d"][\w\d_]*/},{b:/\$\{(.*?)}/}]},t,{cN:"number",b:/([\+\-]+)?[\d]+_[\d_]+/},e.NM]}]}]}}),e.registerLanguage("java",function(e){var t="[À-ʸa-zA-Z_$][À-ʸa-zA-Z_$0-9]*",r=t+"(<"+t+"(\\s*,\\s*"+t+")*>)?",a="false synchronized int abstract float private char boolean static null if const for true while long strictfp finally protected import native final void enum else break transient catch instanceof byte super volatile case assert short package default double public try this switch continue throws protected public private module requires exports do",n="\\b(0[bB]([01]+[01_]+[01]+|[01]+)|0[xX]([a-fA-F0-9]+[a-fA-F0-9_]+[a-fA-F0-9]+|[a-fA-F0-9]+)|(([\\d]+[\\d_]+[\\d]+|[\\d]+)(\\.([\\d]+[\\d_]+[\\d]+|[\\d]+))?|\\.([\\d]+[\\d_]+[\\d]+|[\\d]+))([eE][-+]?\\d+)?)[lLfF]?",i={cN:"number",b:n,r:0};return{aliases:["jsp"],k:a,i:/<\/|#/,c:[e.C("/\\*\\*","\\*/",{r:0,c:[{b:/\w+@/,r:0},{cN:"doctag",b:"@[A-Za-z]+"}]}),e.CLCM,e.CBCM,e.ASM,e.QSM,{cN:"class",bK:"class interface",e:/[{;=]/,eE:!0,k:"class interface",i:/[:"\[\]]/,c:[{bK:"extends implements"},e.UTM]},{bK:"new throw return else",r:0},{cN:"function",b:"("+r+"\\s+)+"+e.UIR+"\\s*\\(",rB:!0,e:/[{;=]/,eE:!0,k:a,c:[{b:e.UIR+"\\s*\\(",rB:!0,r:0,c:[e.UTM]},{cN:"params",b:/\(/,e:/\)/,k:a,r:0,c:[e.ASM,e.QSM,e.CNM,e.CBCM]},e.CLCM,e.CBCM]},i,{cN:"meta",b:"@[A-Za-z]+"}]}}),e.registerLanguage("javascript",function(e){var t="[A-Za-z$_][0-9A-Za-z$_]*",r={keyword:"in of if for while finally var new function do return void else break catch instanceof with throw case default try this switch continue typeof delete let yield const export super debugger as async await static import from as",literal:"true false null undefined NaN Infinity",built_in:"eval isFinite isNaN parseFloat parseInt decodeURI decodeURIComponent encodeURI encodeURIComponent escape unescape Object Function Boolean Error EvalError InternalError RangeError ReferenceError StopIteration SyntaxError TypeError URIError Number Math Date String RegExp Array Float32Array Float64Array Int16Array Int32Array Int8Array Uint16Array Uint32Array Uint8Array Uint8ClampedArray ArrayBuffer DataView JSON Intl arguments require module console window document Symbol Set Map WeakSet WeakMap Proxy Reflect Promise"},a={cN:"number",v:[{b:"\\b(0[bB][01]+)"},{b:"\\b(0[oO][0-7]+)"},{b:e.CNR}],r:0},n={cN:"subst",b:"\\$\\{",e:"\\}",k:r,c:[]},i={cN:"string",b:"`",e:"`",c:[e.BE,n]};n.c=[e.ASM,e.QSM,i,a,e.RM];var s=n.c.concat([e.CBCM,e.CLCM]);return{aliases:["js","jsx"],k:r,c:[{cN:"meta",r:10,b:/^\s*['"]use (strict|asm)['"]/},{cN:"meta",b:/^#!/,e:/$/},e.ASM,e.QSM,i,e.CLCM,e.CBCM,a,{b:/[{,]\s*/,r:0,c:[{b:t+"\\s*:",rB:!0,r:0,c:[{cN:"attr",b:t,r:0}]}]},{b:"("+e.RSR+"|\\b(case|return|throw)\\b)\\s*",k:"return throw case",c:[e.CLCM,e.CBCM,e.RM,{cN:"function",b:"(\\(.*?\\)|"+t+")\\s*=>",rB:!0,e:"\\s*=>",c:[{cN:"params",v:[{b:t},{b:/\(\s*\)/},{b:/\(/,e:/\)/,eB:!0,eE:!0,k:r,c:s}]}]},{b:/</,e:/(\/\w+|\w+\/)>/,sL:"xml",c:[{b:/<\w+\s*\/>/,skip:!0},{b:/<\w+/,e:/(\/\w+|\w+\/)>/,skip:!0,c:[{b:/<\w+\s*\/>/,skip:!0},"self"]}]}],r:0},{cN:"function",bK:"function",e:/\{/,eE:!0,c:[e.inherit(e.TM,{b:t}),{cN:"params",b:/\(/,e:/\)/,eB:!0,eE:!0,c:s}],i:/\[|%/},{b:/\$[(.]/},e.METHOD_GUARD,{cN:"class",bK:"class",e:/[{;=]/,eE:!0,i:/[:"\[\]]/,c:[{bK:"extends"},e.UTM]},{bK:"constructor",e:/\{/,eE:!0}],i:/#(?!!)/}}),e.registerLanguage("json",function(e){var t={literal:"true false null"},r=[e.QSM,e.CNM],a={e:",",eW:!0,eE:!0,c:r,k:t},n={b:"{",e:"}",c:[{cN:"attr",b:/"/,e:/"/,c:[e.BE],i:"\\n"},e.inherit(a,{b:/:/})],i:"\\S"},i={b:"\\[",e:"\\]",c:[e.inherit(a)],i:"\\S"};return r.splice(r.length,0,n,i),{c:r,k:t,i:"\\S"}}),e.registerLanguage("makefile",function(e){var t={cN:"variable",v:[{b:"\\$\\("+e.UIR+"\\)",c:[e.BE]},{b:/\$[@%<?\^\+\*]/}]},r={cN:"string",b:/"/,e:/"/,c:[e.BE,t]},a={cN:"variable",b:/\$\([\w-]+\s/,e:/\)/,k:{built_in:"subst patsubst strip findstring filter filter-out sort word wordlist firstword lastword dir notdir suffix basename addsuffix addprefix join wildcard realpath abspath error warning shell origin flavor foreach if or and call eval file value"},c:[t]},n={b:"^"+e.UIR+"\\s*[:+?]?=",i:"\\n",rB:!0,c:[{b:"^"+e.UIR,e:"[:+?]?=",eE:!0}]},i={cN:"meta",b:/^\.PHONY:/,e:/$/,k:{"meta-keyword":".PHONY"},l:/[\.\w]+/},s={cN:"section",b:/^[^\s]+:/,e:/$/,c:[t]};return{aliases:["mk","mak"],k:"define endef undefine ifdef ifndef ifeq ifneq else endif include -include sinclude override export unexport private vpath",l:/[\w-]+/,c:[e.HCM,t,r,a,n,i,s]}}),e.registerLanguage("xml",function(e){var t="[A-Za-z0-9\\._:-]+",r={eW:!0,i:/</,r:0,c:[{cN:"attr",b:t,r:0},{b:/=\s*/,r:0,c:[{cN:"string",endsParent:!0,v:[{b:/"/,e:/"/},{b:/'/,e:/'/},{b:/[^\s"'=<>`]+/}]}]}]};return{aliases:["html","xhtml","rss","atom","xjb","xsd","xsl","plist"],cI:!0,c:[{cN:"meta",b:"<!DOCTYPE",e:">",r:10,c:[{b:"\\[",e:"\\]"}]},e.C("<!--","-->",{r:10}),{b:"<\\!\\[CDATA\\[",e:"\\]\\]>",r:10},{b:/<\?(php)?/,e:/\?>/,sL:"php",c:[{b:"/\\*",e:"\\*/",skip:!0}]},{cN:"tag",b:"<style(?=\\s|>|$)",e:">",k:{name:"style"},c:[r],starts:{e:"</style>",rE:!0,sL:["css","xml"]}},{cN:"tag",b:"<script(?=\\s|>|$)",e:">",k:{name:"script"},c:[r],starts:{e:"</script>",rE:!0,sL:["actionscript","javascript","handlebars","xml"]}},{cN:"meta",v:[{b:/<\?xml/,e:/\?>/,r:10},{b:/<\?\w+/,e:/\?>/}]},{cN:"tag",b:"</?",e:"/?>",c:[{cN:"name",b:/[^\/><\s]+/,r:0},r]}]}}),e.registerLanguage("markdown",function(e){return{aliases:["md","mkdown","mkd"],c:[{cN:"section",v:[{b:"^#{1,6}",e:"$"},{b:"^.+?\\n[=-]{2,}$"}]},{b:"<",e:">",sL:"xml",r:0},{cN:"bullet",b:"^([*+-]|(\\d+\\.))\\s+"},{cN:"strong",b:"[*_]{2}.+?[*_]{2}"},{cN:"emphasis",v:[{b:"\\*.+?\\*"},{b:"_.+?_",r:0}]},{cN:"quote",b:"^>\\s+",e:"$"},{cN:"code",v:[{b:"^```w*s*$",e:"^```s*$"},{b:"`.+?`"},{b:"^( {4}|	)",e:"$",r:0}]},{b:"^[-\\*]{3,}",e:"$"},{b:"\\[.+?\\][\\(\\[].*?[\\)\\]]",rB:!0,c:[{cN:"string",b:"\\[",e:"\\]",eB:!0,rE:!0,r:0},{cN:"link",b:"\\]\\(",e:"\\)",eB:!0,eE:!0},{cN:"symbol",b:"\\]\\[",e:"\\]",eB:!0,eE:!0}],r:10},{b:/^\[[^\n]+\]:/,rB:!0,c:[{cN:"symbol",b:/\[/,e:/\]/,eB:!0,eE:!0},{cN:"link",b:/:\s*/,e:/$/,eB:!0}]}]}}),e.registerLanguage("nginx",function(e){var t={cN:"variable",v:[{b:/\$\d+/},{b:/\$\{/,e:/}/},{b:"[\\$\\@]"+e.UIR}]},r={eW:!0,l:"[a-z/_]+",k:{literal:"on off yes no true false none blocked debug info notice warn error crit select break last permanent redirect kqueue rtsig epoll poll /dev/poll"},r:0,i:"=>",c:[e.HCM,{cN:"string",c:[e.BE,t],v:[{b:/"/,e:/"/},{b:/'/,e:/'/}]},{b:"([a-z]+):/",e:"\\s",eW:!0,eE:!0,c:[t]},{cN:"regexp",c:[e.BE,t],v:[{b:"\\s\\^",e:"\\s|{|;",rE:!0},{b:"~\\*?\\s+",e:"\\s|{|;",rE:!0},{b:"\\*(\\.[a-z\\-]+)+"},{b:"([a-z\\-]+\\.)+\\*"}]},{cN:"number",b:"\\b\\d{1,3}\\.\\d{1,3}\\.\\d{1,3}\\.\\d{1,3}(:\\d{1,5})?\\b"},{cN:"number",b:"\\b\\d+[kKmMgGdshdwy]*\\b",r:0},t]};return{aliases:["nginxconf"],c:[e.HCM,{b:e.UIR+"\\s+{",rB:!0,e:"{",c:[{cN:"section",b:e.UIR}],r:0},{b:e.UIR+"\\s",e:";|{",rB:!0,c:[{cN:"attribute",b:e.UIR,starts:r}],r:0}],i:"[^\\s\\}]"}}),e.registerLanguage("objectivec",function(e){var t={cN:"built_in",b:"\\b(AV|CA|CF|CG|CI|CL|CM|CN|CT|MK|MP|MTK|MTL|NS|SCN|SK|UI|WK|XC)\\w+"},r={keyword:"int float while char export sizeof typedef const struct for union unsigned long volatile static bool mutable if do return goto void enum else break extern asm case short default double register explicit signed typename this switch continue wchar_t inline readonly assign readwrite self @synchronized id typeof nonatomic super unichar IBOutlet IBAction strong weak copy in out inout bycopy byref oneway __strong __weak __block __autoreleasing @private @protected @public @try @property @end @throw @catch @finally @autoreleasepool @synthesize @dynamic @selector @optional @required @encode @package @import @defs @compatibility_alias __bridge __bridge_transfer __bridge_retained __bridge_retain __covariant __contravariant __kindof _Nonnull _Nullable _Null_unspecified __FUNCTION__ __PRETTY_FUNCTION__ __attribute__ getter setter retain unsafe_unretained nonnull nullable null_unspecified null_resettable class instancetype NS_DESIGNATED_INITIALIZER NS_UNAVAILABLE NS_REQUIRES_SUPER NS_RETURNS_INNER_POINTER NS_INLINE NS_AVAILABLE NS_DEPRECATED NS_ENUM NS_OPTIONS NS_SWIFT_UNAVAILABLE NS_ASSUME_NONNULL_BEGIN NS_ASSUME_NONNULL_END NS_REFINED_FOR_SWIFT NS_SWIFT_NAME NS_SWIFT_NOTHROW NS_DURING NS_HANDLER NS_ENDHANDLER NS_VALUERETURN NS_VOIDRETURN",literal:"false true FALSE TRUE nil YES NO NULL",built_in:"BOOL dispatch_once_t dispatch_queue_t dispatch_sync dispatch_async dispatch_once"},a=/[a-zA-Z@][a-zA-Z0-9_]*/,n="@interface @class @protocol @implementation";return{aliases:["mm","objc","obj-c"],k:r,l:a,i:"</",c:[t,e.CLCM,e.CBCM,e.CNM,e.QSM,{cN:"string",v:[{b:'@"',e:'"',i:"\\n",c:[e.BE]},{b:"'",e:"[^\\\\]'",i:"[^\\\\][^']"}]},{cN:"meta",b:"#",e:"$",c:[{cN:"meta-string",v:[{b:'"',e:'"'},{b:"<",e:">"}]}]},{cN:"class",b:"("+n.split(" ").join("|")+")\\b",e:"({|$)",eE:!0,k:n,l:a,c:[e.UTM]},{b:"\\."+e.UIR,r:0}]}}),e.registerLanguage("perl",function(e){var t="getpwent getservent quotemeta msgrcv scalar kill dbmclose undef lc ma syswrite tr send umask sysopen shmwrite vec qx utime local oct semctl localtime readpipe do return format read sprintf dbmopen pop getpgrp not getpwnam rewinddir qqfileno qw endprotoent wait sethostent bless s|0 opendir continue each sleep endgrent shutdown dump chomp connect getsockname die socketpair close flock exists index shmgetsub for endpwent redo lstat msgctl setpgrp abs exit select print ref gethostbyaddr unshift fcntl syscall goto getnetbyaddr join gmtime symlink semget splice x|0 getpeername recv log setsockopt cos last reverse gethostbyname getgrnam study formline endhostent times chop length gethostent getnetent pack getprotoent getservbyname rand mkdir pos chmod y|0 substr endnetent printf next open msgsnd readdir use unlink getsockopt getpriority rindex wantarray hex system getservbyport endservent int chr untie rmdir prototype tell listen fork shmread ucfirst setprotoent else sysseek link getgrgid shmctl waitpid unpack getnetbyname reset chdir grep split require caller lcfirst until warn while values shift telldir getpwuid my getprotobynumber delete and sort uc defined srand accept package seekdir getprotobyname semop our rename seek if q|0 chroot sysread setpwent no crypt getc chown sqrt write setnetent setpriority foreach tie sin msgget map stat getlogin unless elsif truncate exec keys glob tied closedirioctl socket readlink eval xor readline binmode setservent eof ord bind alarm pipe atan2 getgrent exp time push setgrent gt lt or ne m|0 break given say state when",r={cN:"subst",b:"[$@]\\{",e:"\\}",k:t},a={b:"->{",e:"}"},n={v:[{b:/\$\d/},{b:/[\$%@](\^\w\b|#\w+(::\w+)*|{\w+}|\w+(::\w*)*)/},{b:/[\$%@][^\s\w{]/,r:0}]},i=[e.BE,r,n],s=[n,e.HCM,e.C("^\\=\\w","\\=cut",{eW:!0}),a,{cN:"string",c:i,v:[{b:"q[qwxr]?\\s*\\(",e:"\\)",r:5},{b:"q[qwxr]?\\s*\\[",e:"\\]",r:5},{b:"q[qwxr]?\\s*\\{",e:"\\}",r:5},{b:"q[qwxr]?\\s*\\|",e:"\\|",r:5},{b:"q[qwxr]?\\s*\\<",e:"\\>",r:5},{b:"qw\\s+q",e:"q",r:5},{b:"'",e:"'",c:[e.BE]},{b:'"',e:'"'},{b:"`",e:"`",c:[e.BE]},{b:"{\\w+}",c:[],r:0},{b:"-?\\w+\\s*\\=\\>",c:[],r:0}]},{cN:"number",b:"(\\b0[0-7_]+)|(\\b0x[0-9a-fA-F_]+)|(\\b[1-9][0-9_]*(\\.[0-9_]+)?)|[0_]\\b",r:0},{b:"(\\/\\/|"+e.RSR+"|\\b(split|return|print|reverse|grep)\\b)\\s*",k:"split return print reverse grep",r:0,c:[e.HCM,{cN:"regexp",b:"(s|tr|y)/(\\\\.|[^/])*/(\\\\.|[^/])*/[a-z]*",r:10},{cN:"regexp",b:"(m|qr)?/",e:"/[a-z]*",c:[e.BE],r:0}]},{cN:"function",bK:"sub",e:"(\\s*\\(.*?\\))?[;{]",eE:!0,r:5,c:[e.TM]},{b:"-\\w\\b",r:0},{b:"^__DATA__$",e:"^__END__$",sL:"mojolicious",c:[{b:"^@@.*",e:"$",cN:"comment"}]}];return r.c=s,a.c=s,{aliases:["pl","pm"],l:/[\w\.]+/,k:t,c:s}}),e.registerLanguage("php",function(e){var t={b:"\\$+[a-zA-Z_-ÿ][a-zA-Z0-9_-ÿ]*"},r={cN:"meta",b:/<\?(php)?|\?>/},a={cN:"string",c:[e.BE,r],v:[{b:'b"',e:'"'},{b:"b'",e:"'"},e.inherit(e.ASM,{i:null}),e.inherit(e.QSM,{i:null})]},n={v:[e.BNM,e.CNM]};return{aliases:["php3","php4","php5","php6"],cI:!0,k:"and include_once list abstract global private echo interface as static endswitch array null if endwhile or const for endforeach self var while isset public protected exit foreach throw elseif include __FILE__ empty require_once do xor return parent clone use __CLASS__ __LINE__ else break print eval new catch __METHOD__ case exception default die require __FUNCTION__ enddeclare final try switch continue endfor endif declare unset true false trait goto instanceof insteadof __DIR__ __NAMESPACE__ yield finally",c:[e.HCM,e.C("//","$",{c:[r]}),e.C("/\\*","\\*/",{c:[{cN:"doctag",b:"@[A-Za-z]+"}]}),e.C("__halt_compiler.+?;",!1,{eW:!0,k:"__halt_compiler",l:e.UIR}),{cN:"string",b:/<<<['"]?\w+['"]?$/,e:/^\w+;?$/,c:[e.BE,{cN:"subst",v:[{b:/\$\w+/},{b:/\{\$/,e:/\}/}]}]},r,{cN:"keyword",b:/\$this\b/},t,{b:/(::|->)+[a-zA-Z_\x7f-\xff][a-zA-Z0-9_\x7f-\xff]*/},{cN:"function",bK:"function",e:/[;{]/,eE:!0,i:"\\$|\\[|%",c:[e.UTM,{cN:"params",b:"\\(",e:"\\)",c:["self",t,e.CBCM,a,n]}]},{cN:"class",bK:"class interface",e:"{",eE:!0,i:/[:\(\$"]/,c:[{bK:"extends implements"},e.UTM]},{bK:"namespace",e:";",i:/[\.']/,c:[e.UTM]},{bK:"use",e:";",c:[e.UTM]},{b:"=>"},a,n]}}),e.registerLanguage("python",function(e){var t={keyword:"and elif is global as in if from raise for except finally print import pass return exec else break not with class assert yield try while continue del or def lambda async await nonlocal|10 None True False",built_in:"Ellipsis NotImplemented"},r={cN:"meta",b:/^(>>>|\.\.\.) /},a={cN:"subst",b:/\{/,e:/\}/,k:t,i:/#/},n={cN:"string",c:[e.BE],v:[{b:/(u|b)?r?'''/,e:/'''/,c:[r],r:10},{b:/(u|b)?r?"""/,e:/"""/,c:[r],r:10},{b:/(fr|rf|f)'''/,e:/'''/,c:[r,a]},{b:/(fr|rf|f)"""/,e:/"""/,c:[r,a]},{b:/(u|r|ur)'/,e:/'/,r:10},{b:/(u|r|ur)"/,e:/"/,r:10},{b:/(b|br)'/,e:/'/},{b:/(b|br)"/,e:/"/},{b:/(fr|rf|f)'/,e:/'/,c:[a]},{b:/(fr|rf|f)"/,e:/"/,c:[a]},e.ASM,e.QSM]},i={cN:"number",r:0,v:[{b:e.BNR+"[lLjJ]?"},{b:"\\b(0o[0-7]+)[lLjJ]?"},{b:e.CNR+"[lLjJ]?"}]},s={cN:"params",b:/\(/,e:/\)/,c:["self",r,i,n]};return a.c=[n,i,r],{aliases:["py","gyp"],k:t,i:/(<\/|->|\?)|=>/,c:[r,i,n,e.HCM,{v:[{cN:"function",bK:"def"},{cN:"class",bK:"class"}],e:/:/,i:/[${=;\n,]/,c:[e.UTM,s,{b:/->/,eW:!0,k:"None"}]},{cN:"meta",b:/^[\t ]*@/,e:/$/},{b:/\b(print|exec)\(/}]}}),e.registerLanguage("ruby",function(e){
var t="[a-zA-Z_]\\w*[!?=]?|[-+~]\\@|<<|>>|=~|===?|<=>|[<>]=?|\\*\\*|[-/+%^&*~`|]|\\[\\]=?",r={keyword:"and then defined module in return redo if BEGIN retry end for self when next until do begin unless END rescue else break undef not super class case require yield alias while ensure elsif or include attr_reader attr_writer attr_accessor",literal:"true false nil"},a={cN:"doctag",b:"@[A-Za-z]+"},n={b:"#<",e:">"},i=[e.C("#","$",{c:[a]}),e.C("^\\=begin","^\\=end",{c:[a],r:10}),e.C("^__END__","\\n$")],s={cN:"subst",b:"#\\{",e:"}",k:r},c={cN:"string",c:[e.BE,s],v:[{b:/'/,e:/'/},{b:/"/,e:/"/},{b:/`/,e:/`/},{b:"%[qQwWx]?\\(",e:"\\)"},{b:"%[qQwWx]?\\[",e:"\\]"},{b:"%[qQwWx]?{",e:"}"},{b:"%[qQwWx]?<",e:">"},{b:"%[qQwWx]?/",e:"/"},{b:"%[qQwWx]?%",e:"%"},{b:"%[qQwWx]?-",e:"-"},{b:"%[qQwWx]?\\|",e:"\\|"},{b:/\B\?(\\\d{1,3}|\\x[A-Fa-f0-9]{1,2}|\\u[A-Fa-f0-9]{4}|\\?\S)\b/},{b:/<<(-?)\w+$/,e:/^\s*\w+$/}]},o={cN:"params",b:"\\(",e:"\\)",endsParent:!0,k:r},l=[c,n,{cN:"class",bK:"class module",e:"$|;",i:/=/,c:[e.inherit(e.TM,{b:"[A-Za-z_]\\w*(::\\w+)*(\\?|\\!)?"}),{b:"<\\s*",c:[{b:"("+e.IR+"::)?"+e.IR}]}].concat(i)},{cN:"function",bK:"def",e:"$|;",c:[e.inherit(e.TM,{b:t}),o].concat(i)},{b:e.IR+"::"},{cN:"symbol",b:e.UIR+"(\\!|\\?)?:",r:0},{cN:"symbol",b:":(?!\\s)",c:[c,{b:t}],r:0},{cN:"number",b:"(\\b0[0-7_]+)|(\\b0x[0-9a-fA-F_]+)|(\\b[1-9][0-9_]*(\\.[0-9_]+)?)|[0_]\\b",r:0},{b:"(\\$\\W)|((\\$|\\@\\@?)(\\w+))"},{cN:"params",b:/\|/,e:/\|/,k:r},{b:"("+e.RSR+"|unless)\\s*",k:"unless",c:[n,{cN:"regexp",c:[e.BE,s],i:/\n/,v:[{b:"/",e:"/[a-z]*"},{b:"%r{",e:"}[a-z]*"},{b:"%r\\(",e:"\\)[a-z]*"},{b:"%r!",e:"![a-z]*"},{b:"%r\\[",e:"\\][a-z]*"}]}].concat(i),r:0}].concat(i);s.c=l,o.c=l;var u="[>?]>",d="[\\w#]+\\(\\w+\\):\\d+:\\d+>",b="(\\w+-)?\\d+\\.\\d+\\.\\d(p\\d+)?[^>]+>",p=[{b:/^\s*=>/,starts:{e:"$",c:l}},{cN:"meta",b:"^("+u+"|"+d+"|"+b+")",starts:{e:"$",c:l}}];return{aliases:["rb","gemspec","podspec","thor","irb"],k:r,i:/\/\*/,c:i.concat(p).concat(l)}}),e.registerLanguage("shell",function(e){return{aliases:["console"],c:[{cN:"meta",b:"^\\s{0,3}[\\w\\d\\[\\]()@-]*[>%$#]",starts:{e:"$",sL:"bash"}}]}}),e.registerLanguage("sql",function(e){var t=e.C("--","$");return{cI:!0,i:/[<>{}*#]/,c:[{bK:"begin end start commit rollback savepoint lock alter create drop rename call delete do handler insert load replace select truncate update set show pragma grant merge describe use explain help declare prepare execute deallocate release unlock purge reset change stop analyze cache flush optimize repair kill install uninstall checksum restore check backup revoke comment",e:/;/,eW:!0,l:/[\w\.]+/,k:{keyword:"abort abs absolute acc acce accep accept access accessed accessible account acos action activate add addtime admin administer advanced advise aes_decrypt aes_encrypt after agent aggregate ali alia alias allocate allow alter always analyze ancillary and any anydata anydataset anyschema anytype apply archive archived archivelog are as asc ascii asin assembly assertion associate asynchronous at atan atn2 attr attri attrib attribu attribut attribute attributes audit authenticated authentication authid authors auto autoallocate autodblink autoextend automatic availability avg backup badfile basicfile before begin beginning benchmark between bfile bfile_base big bigfile bin binary_double binary_float binlog bit_and bit_count bit_length bit_or bit_xor bitmap blob_base block blocksize body both bound buffer_cache buffer_pool build bulk by byte byteordermark bytes cache caching call calling cancel capacity cascade cascaded case cast catalog category ceil ceiling chain change changed char_base char_length character_length characters characterset charindex charset charsetform charsetid check checksum checksum_agg child choose chr chunk class cleanup clear client clob clob_base clone close cluster_id cluster_probability cluster_set clustering coalesce coercibility col collate collation collect colu colum column column_value columns columns_updated comment commit compact compatibility compiled complete composite_limit compound compress compute concat concat_ws concurrent confirm conn connec connect connect_by_iscycle connect_by_isleaf connect_by_root connect_time connection consider consistent constant constraint constraints constructor container content contents context contributors controlfile conv convert convert_tz corr corr_k corr_s corresponding corruption cos cost count count_big counted covar_pop covar_samp cpu_per_call cpu_per_session crc32 create creation critical cross cube cume_dist curdate current current_date current_time current_timestamp current_user cursor curtime customdatum cycle data database databases datafile datafiles datalength date_add date_cache date_format date_sub dateadd datediff datefromparts datename datepart datetime2fromparts day day_to_second dayname dayofmonth dayofweek dayofyear days db_role_change dbtimezone ddl deallocate declare decode decompose decrement decrypt deduplicate def defa defau defaul default defaults deferred defi defin define degrees delayed delegate delete delete_all delimited demand dense_rank depth dequeue des_decrypt des_encrypt des_key_file desc descr descri describ describe descriptor deterministic diagnostics difference dimension direct_load directory disable disable_all disallow disassociate discardfile disconnect diskgroup distinct distinctrow distribute distributed div do document domain dotnet double downgrade drop dumpfile duplicate duration each edition editionable editions element ellipsis else elsif elt empty enable enable_all enclosed encode encoding encrypt end end-exec endian enforced engine engines enqueue enterprise entityescaping eomonth error errors escaped evalname evaluate event eventdata events except exception exceptions exchange exclude excluding execu execut execute exempt exists exit exp expire explain export export_set extended extent external external_1 external_2 externally extract failed failed_login_attempts failover failure far fast feature_set feature_value fetch field fields file file_name_convert filesystem_like_logging final finish first first_value fixed flash_cache flashback floor flush following follows for forall force form forma format found found_rows freelist freelists freepools fresh from from_base64 from_days ftp full function general generated get get_format get_lock getdate getutcdate global global_name globally go goto grant grants greatest group group_concat group_id grouping grouping_id groups gtid_subtract guarantee guard handler hash hashkeys having hea head headi headin heading heap help hex hierarchy high high_priority hosts hour http id ident_current ident_incr ident_seed identified identity idle_time if ifnull ignore iif ilike ilm immediate import in include including increment index indexes indexing indextype indicator indices inet6_aton inet6_ntoa inet_aton inet_ntoa infile initial initialized initially initrans inmemory inner innodb input insert install instance instantiable instr interface interleaved intersect into invalidate invisible is is_free_lock is_ipv4 is_ipv4_compat is_not is_not_null is_used_lock isdate isnull isolation iterate java join json json_exists keep keep_duplicates key keys kill language large last last_day last_insert_id last_value lax lcase lead leading least leaves left len lenght length less level levels library like like2 like4 likec limit lines link list listagg little ln load load_file lob lobs local localtime localtimestamp locate locator lock locked log log10 log2 logfile logfiles logging logical logical_reads_per_call logoff logon logs long loop low low_priority lower lpad lrtrim ltrim main make_set makedate maketime managed management manual map mapping mask master master_pos_wait match matched materialized max maxextents maximize maxinstances maxlen maxlogfiles maxloghistory maxlogmembers maxsize maxtrans md5 measures median medium member memcompress memory merge microsecond mid migration min minextents minimum mining minus minute minvalue missing mod mode model modification modify module monitoring month months mount move movement multiset mutex name name_const names nan national native natural nav nchar nclob nested never new newline next nextval no no_write_to_binlog noarchivelog noaudit nobadfile nocheck nocompress nocopy nocycle nodelay nodiscardfile noentityescaping noguarantee nokeep nologfile nomapping nomaxvalue nominimize nominvalue nomonitoring none noneditionable nonschema noorder nopr nopro noprom nopromp noprompt norely noresetlogs noreverse normal norowdependencies noschemacheck noswitch not nothing notice notrim novalidate now nowait nth_value nullif nulls num numb numbe nvarchar nvarchar2 object ocicoll ocidate ocidatetime ociduration ociinterval ociloblocator ocinumber ociref ocirefcursor ocirowid ocistring ocitype oct octet_length of off offline offset oid oidindex old on online only opaque open operations operator optimal optimize option optionally or oracle oracle_date oradata ord ordaudio orddicom orddoc order ordimage ordinality ordvideo organization orlany orlvary out outer outfile outline output over overflow overriding package pad parallel parallel_enable parameters parent parse partial partition partitions pascal passing password password_grace_time password_lock_time password_reuse_max password_reuse_time password_verify_function patch path patindex pctincrease pctthreshold pctused pctversion percent percent_rank percentile_cont percentile_disc performance period period_add period_diff permanent physical pi pipe pipelined pivot pluggable plugin policy position post_transaction pow power pragma prebuilt precedes preceding precision prediction prediction_cost prediction_details prediction_probability prediction_set prepare present preserve prior priority private private_sga privileges procedural procedure procedure_analyze processlist profiles project prompt protection public publishingservername purge quarter query quick quiesce quota quotename radians raise rand range rank raw read reads readsize rebuild record records recover recovery recursive recycle redo reduced ref reference referenced references referencing refresh regexp_like register regr_avgx regr_avgy regr_count regr_intercept regr_r2 regr_slope regr_sxx regr_sxy reject rekey relational relative relaylog release release_lock relies_on relocate rely rem remainder rename repair repeat replace replicate replication required reset resetlogs resize resource respect restore restricted result result_cache resumable resume retention return returning returns reuse reverse revoke right rlike role roles rollback rolling rollup round row row_count rowdependencies rowid rownum rows rtrim rules safe salt sample save savepoint sb1 sb2 sb4 scan schema schemacheck scn scope scroll sdo_georaster sdo_topo_geometry search sec_to_time second section securefile security seed segment select self sequence sequential serializable server servererror session session_user sessions_per_user set sets settings sha sha1 sha2 share shared shared_pool short show shrink shutdown si_averagecolor si_colorhistogram si_featurelist si_positionalcolor si_stillimage si_texture siblings sid sign sin size size_t sizes skip slave sleep smalldatetimefromparts smallfile snapshot some soname sort soundex source space sparse spfile split sql sql_big_result sql_buffer_result sql_cache sql_calc_found_rows sql_small_result sql_variant_property sqlcode sqldata sqlerror sqlname sqlstate sqrt square standalone standby start starting startup statement static statistics stats_binomial_test stats_crosstab stats_ks_test stats_mode stats_mw_test stats_one_way_anova stats_t_test_ stats_t_test_indep stats_t_test_one stats_t_test_paired stats_wsr_test status std stddev stddev_pop stddev_samp stdev stop storage store stored str str_to_date straight_join strcmp strict string struct stuff style subdate subpartition subpartitions substitutable substr substring subtime subtring_index subtype success sum suspend switch switchoffset switchover sync synchronous synonym sys sys_xmlagg sysasm sysaux sysdate sysdatetimeoffset sysdba sysoper system system_user sysutcdatetime table tables tablespace tan tdo template temporary terminated tertiary_weights test than then thread through tier ties time time_format time_zone timediff timefromparts timeout timestamp timestampadd timestampdiff timezone_abbr timezone_minute timezone_region to to_base64 to_date to_days to_seconds todatetimeoffset trace tracking transaction transactional translate translation treat trigger trigger_nestlevel triggers trim truncate try_cast try_convert try_parse type ub1 ub2 ub4 ucase unarchived unbounded uncompress under undo unhex unicode uniform uninstall union unique unix_timestamp unknown unlimited unlock unpivot unrecoverable unsafe unsigned until untrusted unusable unused update updated upgrade upped upper upsert url urowid usable usage use use_stored_outlines user user_data user_resources users using utc_date utc_timestamp uuid uuid_short validate validate_password_strength validation valist value values var var_samp varcharc vari varia variab variabl variable variables variance varp varraw varrawc varray verify version versions view virtual visible void wait wallet warning warnings week weekday weekofyear wellformed when whene whenev wheneve whenever where while whitespace with within without work wrapped xdb xml xmlagg xmlattributes xmlcast xmlcolattval xmlelement xmlexists xmlforest xmlindex xmlnamespaces xmlpi xmlquery xmlroot xmlschema xmlserialize xmltable xmltype xor year year_to_month years yearweek",literal:"true false null",built_in:"array bigint binary bit blob boolean char character date dec decimal float int int8 integer interval number numeric real record serial serial8 smallint text varchar varying void"},c:[{cN:"string",b:"'",e:"'",c:[e.BE,{b:"''"}]},{cN:"string",b:'"',e:'"',c:[e.BE,{b:'""'}]},{cN:"string",b:"`",e:"`",c:[e.BE]},e.CNM,e.CBCM,t]},e.CBCM,t]}}),e});
````
