## Supplementary material for "Method for inferring migration routes from phylogenetic trees": Some route-trees: html2canvas.js

```
/*!
 * html2canvas 1.0.0-rc.5 <https://html2canvas.hertzen.com>
 * Copyright (c) 2019 Niklas von Hertzen <https://hertzen.com>
 * Released under MIT License
 */
(function (global, factory) {
    typeof exports === 'object' && typeof module !== 'undefined' ? module.exports = factory() :
    typeof define === 'function' && define.amd ? define(factory) :
    (global = global || self, global.html2canvas = factory());
}(this, function () { 'use strict';

    /*! *****************************************************************************
    Copyright (c) Microsoft Corporation. All rights reserved.
    Licensed under the Apache License, Version 2.0 (the "License"); you may not use
    this file except in compliance with the License. You may obtain a copy of the
    License at http://www.apache.org/licenses/LICENSE-2.0

    THIS CODE IS PROVIDED ON AN *AS IS* BASIS, WITHOUT WARRANTIES OR CONDITIONS OF ANY
    KIND, EITHER EXPRESS OR IMPLIED, INCLUDING WITHOUT LIMITATION ANY IMPLIED
    WARRANTIES OR CONDITIONS OF TITLE, FITNESS FOR A PARTICULAR PURPOSE,
    MERCHANTABLITY OR NON-INFRINGEMENT.

    See the Apache Version 2.0 License for specific language governing permissions
    and limitations under the License.
    ***************************************************************************** */
    /* global Reflect, Promise */

    var extendStatics = function(d, b) {
        extendStatics = Object.setPrototypeOf ||
            ({ __proto__: [] } instanceof Array && function (d, b) { d.__proto__ = b; }) ||
            function (d, b) { for (var p in b) if (b.hasOwnProperty(p)) d[p] = b[p]; };
        return extendStatics(d, b);
    };

    function __extends(d, b) {
        extendStatics(d, b);
        function __() { this.constructor = d; }
        d.prototype = b === null ? Object.create(b) : (__.prototype = b.prototype, new __());
    }

    var __assign = function() {
        __assign = Object.assign || function __assign(t) {
            for (var s, i = 1, n = arguments.length; i < n; i++) {
                s = arguments[i];
                for (var p in s) if (Object.prototype.hasOwnProperty.call(s, p)) t[p] = s[p];
            }
            return t;
        };
        return __assign.apply(this, arguments);
    };

    function __awaiter(thisArg, _arguments, P, generator) {
        return new (P || (P = Promise))(function (resolve, reject) {
            function fulfilled(value) { try { step(generator.next(value)); } catch (e) { reject(e); } }
            function rejected(value) { try { step(generator["throw"](value)); } catch (e) { reject(e); } }
            function step(result) { result.done ? resolve(result.value) : new P(function (resolve) { resolve(result.value); }).then(fulfilled, rejected); }
            step((generator = generator.apply(thisArg, _arguments || [])).next());
        });
    }

    function __generator(thisArg, body) {
        var _ = { label: 0, sent: function() { if (t[0] & 1) throw t[1]; return t[1]; }, trys: [], ops: [] }, f, y, t, g;
        return g = { next: verb(0), "throw": verb(1), "return": verb(2) }, typeof Symbol === "function" && (g[Symbol.iterator] = function() { return this; }), g;
        function verb(n) { return function (v) { return step([n, v]); }; }
        function step(op) {
            if (f) throw new TypeError("Generator is already executing.");
            while (_) try {
                if (f = 1, y && (t = op[0] & 2 ? y["return"] : op[0] ? y["throw"] || ((t = y["return"]) && t.call(y), 0) : y.next) && !(t = t.call(y, op[1])).done) return t;
                if (y = 0, t) op = [op[0] & 2, t.value];
                switch (op[0]) {
                    case 0: case 1: t = op; break;
                    case 4: _.label++; return { value: op[1], done: false };
                    case 5: _.label++; y = op[1]; op = [0]; continue;
                    case 7: op = _.ops.pop(); _.trys.pop(); continue;
                    default:
                        if (!(t = _.trys, t = t.length > 0 && t[t.length - 1]) && (op[0] === 6 || op[0] === 2)) { _ = 0; continue; }
                        if (op[0] === 3 && (!t || (op[1] > t[0] && op[1] < t[3]))) { _.label = op[1]; break; }
                        if (op[0] === 6 && _.label < t[1]) { _.label = t[1]; t = op; break; }
                        if (t && _.label < t[2]) { _.label = t[2]; _.ops.push(op); break; }
                        if (t[2]) _.ops.pop();
                        _.trys.pop(); continue;
                }
                op = body.call(thisArg, _);
            } catch (e) { op = [6, e]; y = 0; } finally { f = t = 0; }
            if (op[0] & 5) throw op[1]; return { value: op[0] ? op[1] : void 0, done: true };
        }
    }

    var Bounds = /** @class */ (function () {
        function Bounds(x, y, w, h) {
            this.left = x;
            this.top = y;
            this.width = w;
            this.height = h;
        }
        Bounds.prototype.add = function (x, y, w, h) {
            return new Bounds(this.left + x, this.top + y, this.width + w, this.height + h);
        };
        Bounds.fromClientRect = function (clientRect) {
            return new Bounds(clientRect.left, clientRect.top, clientRect.width, clientRect.height);
        };
        return Bounds;
    }());
    var parseBounds = function (node) {
        return Bounds.fromClientRect(node.getBoundingClientRect());
    };
    var parseDocumentSize = function (document) {
        var body = document.body;
        var documentElement = document.documentElement;
        if (!body || !documentElement) {
            throw new Error("Unable to get document size");
        }
        var width = Math.max(Math.max(body.scrollWidth, documentElement.scrollWidth), Math.max(body.offsetWidth, documentElement.offsetWidth), Math.max(body.clientWidth, documentElement.clientWidth));
        var height = Math.max(Math.max(body.scrollHeight, documentElement.scrollHeight), Math.max(body.offsetHeight, documentElement.offsetHeight), Math.max(body.clientHeight, documentElement.clientHeight));
        return new Bounds(0, 0, width, height);
    };

    /*
     * css-line-break 1.1.1 <https://github.com/niklasvh/css-line-break#readme>
     * Copyright (c) 2019 Niklas von Hertzen <https://hertzen.com>
     * Released under MIT License
     */
    var toCodePoints = function (str) {
        var codePoints = [];
        var i = 0;
        var length = str.length;
        while (i < length) {
            var value = str.charCodeAt(i++);
            if (value >= 0xd800 && value <= 0xdbff && i < length) {
                var extra = str.charCodeAt(i++);
                if ((extra & 0xfc00) === 0xdc00) {
                    codePoints.push(((value & 0x3ff) << 10) + (extra & 0x3ff) + 0x10000);
                }
                else {
                    codePoints.push(value);
                    i--;
                }
            }
            else {
                codePoints.push(value);
            }
        }
        return codePoints;
    };
    var fromCodePoint = function () {
        var codePoints = [];
        for (var _i = 0; _i < arguments.length; _i++) {
            codePoints[_i] = arguments[_i];
        }
        if (String.fromCodePoint) {
            return String.fromCodePoint.apply(String, codePoints);
        }
        var length = codePoints.length;
        if (!length) {
            return '';
        }
        var codeUnits = [];
        var index = -1;
        var result = '';
        while (++index < length) {
            var codePoint = codePoints[index];
            if (codePoint <= 0xffff) {
                codeUnits.push(codePoint);
            }
            else {
                codePoint -= 0x10000;
                codeUnits.push((codePoint >> 10) + 0xd800, codePoint % 0x400 + 0xdc00);
            }
            if (index + 1 === length || codeUnits.length > 0x4000) {
                result += String.fromCharCode.apply(String, codeUnits);
                codeUnits.length = 0;
            }
        }
        return result;
    };
    var chars = 'ABCDEFGHIJKLMNOPQRSTUVWXYZabcdefghijklmnopqrstuvwxyz0123456789+/';
    // Use a lookup table to find the index.
    var lookup = typeof Uint8Array === 'undefined' ? [] : new Uint8Array(256);
    for (var i = 0; i < chars.length; i++) {
        lookup[chars.charCodeAt(i)] = i;
    }
    var decode = function (base64) {
        var bufferLength = base64.length * 0.75, len = base64.length, i, p = 0, encoded1, encoded2, encoded3, encoded4;
        if (base64[base64.length - 1] === '=') {
            bufferLength--;
            if (base64[base64.length - 2] === '=') {
                bufferLength--;
            }
        }
        var buffer = typeof ArrayBuffer !== 'undefined' &&
            typeof Uint8Array !== 'undefined' &&
            typeof Uint8Array.prototype.slice !== 'undefined'
            ? new ArrayBuffer(bufferLength)
            : new Array(bufferLength);
        var bytes = Array.isArray(buffer) ? buffer : new Uint8Array(buffer);
        for (i = 0; i < len; i += 4) {
            encoded1 = lookup[base64.charCodeAt(i)];
            encoded2 = lookup[base64.charCodeAt(i + 1)];
            encoded3 = lookup[base64.charCodeAt(i + 2)];
            encoded4 = lookup[base64.charCodeAt(i + 3)];
            bytes[p++] = (encoded1 << 2) | (encoded2 >> 4);
            bytes[p++] = ((encoded2 & 15) << 4) | (encoded3 >> 2);
            bytes[p++] = ((encoded3 & 3) << 6) | (encoded4 & 63);
        }
        return buffer;
    };
    var polyUint16Array = function (buffer) {
        var length = buffer.length;
        var bytes = [];
        for (var i = 0; i < length; i += 2) {
            bytes.push((buffer[i + 1] << 8) | buffer[i]);
        }
        return bytes;
    };
    var polyUint32Array = function (buffer) {
        var length = buffer.length;
        var bytes = [];
        for (var i = 0; i < length; i += 4) {
            bytes.push((buffer[i + 3] << 24) | (buffer[i + 2] << 16) | (buffer[i + 1] << 8) | buffer[i]);
        }
        return bytes;
    };

    /** Shift size for getting the index-2 table offset. */
    var UTRIE2_SHIFT_2 = 5;
    /** Shift size for getting the index-1 table offset. */
    var UTRIE2_SHIFT_1 = 6 + 5;
    /**
     * Shift size for shifting left the index array values.
     * Increases possible data size with 16-bit index values at the cost
     * of compactability.
     * This requires data blocks to be aligned by UTRIE2_DATA_GRANULARITY.
     */
    var UTRIE2_INDEX_SHIFT = 2;
    /**
     * Difference between the two shift sizes,
     * for getting an index-1 offset from an index-2 offset. 6=11-5
     */
    var UTRIE2_SHIFT_1_2 = UTRIE2_SHIFT_1 - UTRIE2_SHIFT_2;
    /**
     * The part of the index-2 table for U+D800..U+DBFF stores values for
     * lead surrogate code _units_ not code _points_.
     * Values for lead surrogate code _points_ are indexed with this portion of the table.
     * Length=32=0x20=0x400>>UTRIE2_SHIFT_2. (There are 1024=0x400 lead surrogates.)
     */
    var UTRIE2_LSCP_INDEX_2_OFFSET = 0x10000 >> UTRIE2_SHIFT_2;
    /** Number of entries in a data block. 32=0x20 */
    var UTRIE2_DATA_BLOCK_LENGTH = 1 << UTRIE2_SHIFT_2;
    /** Mask for getting the lower bits for the in-data-block offset. */
    var UTRIE2_DATA_MASK = UTRIE2_DATA_BLOCK_LENGTH - 1;
    var UTRIE2_LSCP_INDEX_2_LENGTH = 0x400 >> UTRIE2_SHIFT_2;
    /** Count the lengths of both BMP pieces. 2080=0x820 */
    var UTRIE2_INDEX_2_BMP_LENGTH = UTRIE2_LSCP_INDEX_2_OFFSET + UTRIE2_LSCP_INDEX_2_LENGTH;
    /**
     * The 2-byte UTF-8 version of the index-2 table follows at offset 2080=0x820.
     * Length 32=0x20 for lead bytes C0..DF, regardless of UTRIE2_SHIFT_2.
     */
    var UTRIE2_UTF8_2B_INDEX_2_OFFSET = UTRIE2_INDEX_2_BMP_LENGTH;
    var UTRIE2_UTF8_2B_INDEX_2_LENGTH = 0x800 >> 6; /* U+0800 is the first code point after 2-byte UTF-8 */
    /**
     * The index-1 table, only used for supplementary code points, at offset 2112=0x840.
     * Variable length, for code points up to highStart, where the last single-value range starts.
     * Maximum length 512=0x200=0x100000>>UTRIE2_SHIFT_1.
     * (For 0x100000 supplementary code points U+10000..U+10ffff.)
     *
     * The part of the index-2 table for supplementary code points starts
     * after this index-1 table.
     *
     * Both the index-1 table and the following part of the index-2 table
     * are omitted completely if there is only BMP data.
     */
    var UTRIE2_INDEX_1_OFFSET = UTRIE2_UTF8_2B_INDEX_2_OFFSET + UTRIE2_UTF8_2B_INDEX_2_LENGTH;
    /**
     * Number of index-1 entries for the BMP. 32=0x20
     * This part of the index-1 table is omitted from the serialized form.
     */
    var UTRIE2_OMITTED_BMP_INDEX_1_LENGTH = 0x10000 >> UTRIE2_SHIFT_1;
    /** Number of entries in an index-2 block. 64=0x40 */
    var UTRIE2_INDEX_2_BLOCK_LENGTH = 1 << UTRIE2_SHIFT_1_2;
    /** Mask for getting the lower bits for the in-index-2-block offset. */
    var UTRIE2_INDEX_2_MASK = UTRIE2_INDEX_2_BLOCK_LENGTH - 1;
    var slice16 = function (view, start, end) {
        if (view.slice) {
            return view.slice(start, end);
        }
        return new Uint16Array(Array.prototype.slice.call(view, start, end));
    };
    var slice32 = function (view, start, end) {
        if (view.slice) {
            return view.slice(start, end);
        }
        return new Uint32Array(Array.prototype.slice.call(view, start, end));
    };
    var createTrieFromBase64 = function (base64) {
        var buffer = decode(base64);
        var view32 = Array.isArray(buffer) ? polyUint32Array(buffer) : new Uint32Array(buffer);
        var view16 = Array.isArray(buffer) ? polyUint16Array(buffer) : new Uint16Array(buffer);
        var headerLength = 24;
        var index = slice16(view16, headerLength / 2, view32[4] / 2);
        var data = view32[5] === 2
            ? slice16(view16, (headerLength + view32[4]) / 2)
            : slice32(view32, Math.ceil((headerLength + view32[4]) / 4));
        return new Trie(view32[0], view32[1], view32[2], view32[3], index, data);
    };
    var Trie = /** @class */ (function () {
        function Trie(initialValue, errorValue, highStart, highValueIndex, index, data) {
            this.initialValue = initialValue;
            this.errorValue = errorValue;
            this.highStart = highStart;
            this.highValueIndex = highValueIndex;
            this.index = index;
            this.data = data;
        }
        /**
         * Get the value for a code point as stored in the Trie.
         *
         * @param codePoint the code point
         * @return the value
         */
        Trie.prototype.get = function (codePoint) {
            var ix;
            if (codePoint >= 0) {
                if (codePoint < 0x0d800 || (codePoint > 0x0dbff && codePoint <= 0x0ffff)) {
                    // Ordinary BMP code point, excluding leading surrogates.
                    // BMP uses a single level lookup.  BMP index starts at offset 0 in the Trie2 index.
                    // 16 bit data is stored in the index array itself.
                    ix = this.index[codePoint >> UTRIE2_SHIFT_2];
                    ix = (ix << UTRIE2_INDEX_SHIFT) + (codePoint & UTRIE2_DATA_MASK);
                    return this.data[ix];
                }
                if (codePoint <= 0xffff) {
                    // Lead Surrogate Code Point.  A Separate index section is stored for
                    // lead surrogate code units and code points.
                    //   The main index has the code unit data.
                    //   For this function, we need the code point data.
                    // Note: this expression could be refactored for slightly improved efficiency, but
                    //       surrogate code points will be so rare in practice that it's not worth it.
                    ix = this.index[UTRIE2_LSCP_INDEX_2_OFFSET + ((codePoint - 0xd800) >> UTRIE2_SHIFT_2)];
                    ix = (ix << UTRIE2_INDEX_SHIFT) + (codePoint & UTRIE2_DATA_MASK);
                    return this.data[ix];
                }
                if (codePoint < this.highStart) {
                    // Supplemental code point, use two-level lookup.
                    ix = UTRIE2_INDEX_1_OFFSET - UTRIE2_OMITTED_BMP_INDEX_1_LENGTH + (codePoint >> UTRIE2_SHIFT_1);
                    ix = this.index[ix];
                    ix += (codePoint >> UTRIE2_SHIFT_2) & UTRIE2_INDEX_2_MASK;
                    ix = this.index[ix];
                    ix = (ix << UTRIE2_INDEX_SHIFT) + (codePoint & UTRIE2_DATA_MASK);
                    return this.data[ix];
                }
                if (codePoint <= 0x10ffff) {
                    return this.data[this.highValueIndex];
                }
            }
            // Fall through.  The code point is outside of the legal range of 0..0x10ffff.
            return this.errorValue;
        };
        return Trie;
    }());

    var base64 = 'KwAAAAAAAAAACA4AIDoAAPAfAAACAAAAAAAIABAAGABAAEgAUABYAF4AZgBeAGYAYABoAHAAeABeAGYAfACEAIAAiACQAJgAoACoAK0AtQC9AMUAXgBmAF4AZgBeAGYAzQDVAF4AZgDRANkA3gDmAOwA9AD8AAQBDAEUARoBIgGAAIgAJwEvATcBPwFFAU0BTAFUAVwBZAFsAXMBewGDATAAiwGTAZsBogGkAawBtAG8AcIBygHSAdoB4AHoAfAB+AH+AQYCDgIWAv4BHgImAi4CNgI+AkUCTQJTAlsCYwJrAnECeQKBAk0CiQKRApkCoQKoArACuALAAsQCzAIwANQC3ALkAjAA7AL0AvwCAQMJAxADGAMwACADJgMuAzYDPgOAAEYDSgNSA1IDUgNaA1oDYANiA2IDgACAAGoDgAByA3YDfgOAAIQDgACKA5IDmgOAAIAAogOqA4AAgACAAIAAgACAAIAAgACAAIAAgACAAIAAgACAAIAAgACAAK8DtwOAAIAAvwPHA88D1wPfAyAD5wPsA/QD/AOAAIAABAQMBBIEgAAWBB4EJgQuBDMEIAM7BEEEXgBJBCADUQRZBGEEaQQwADAAcQQ+AXkEgQSJBJEEgACYBIAAoASoBK8EtwQwAL8ExQSAAIAAgACAAIAAgACgAM0EXgBeAF4AXgBeAF4AXgBeANUEXgDZBOEEXgDpBPEE+QQBBQkFEQUZBSEFKQUxBTUFPQVFBUwFVAVcBV4AYwVeAGsFcwV7BYMFiwWSBV4AmgWgBacFXgBeAF4AXgBeAKsFXgCyBbEFugW7BcIFwgXIBcIFwgXQBdQF3AXkBesF8wX7BQMGCwYTBhsGIwYrBjMGOwZeAD8GRwZNBl4AVAZbBl4AXgBeAF4AXgBeAF4AXgBeAF4AXgBeAGMGXgBqBnEGXgBeAF4AXgBeAF4AXgBeAF4AXgB5BoAG4wSGBo4GkwaAAIADHgR5AF4AXgBeAJsGgABGA4AAowarBrMGswagALsGwwbLBjAA0wbaBtoG3QbaBtoG2gbaBtoG2gblBusG8wb7BgMHCwcTBxsHCwcjBysHMAc1BzUHOgdCB9oGSgdSB1oHYAfaBloHaAfaBlIH2gbaBtoG2gbaBtoG2gbaBjUHNQc1BzUHNQc1BzUHNQc1BzUHNQc1BzUHNQc1BzUHNQc1BzUHNQc1BzUHNQc1BzUHNQc1BzUHNQc1BzUHNQc1BzUHNQc1BzUHNQc1BzUHNQc1BzUHNQc1BzUHNQc1BzUHNQc1BzUHNQc1BzUHNQc1BzUHNQc1BzUHNQc1BzUHNQc1BzUHNQc1BzUHNQc1BzUHNQc1BzUHNQc1BzUHNQc1BzUHNQc1BzUHNQc1BzUHNQc1BzUHNQc1BzUHNQc1BzUHNQc1BzUHNQc1BzUHNQc1BzUHNQc1BzUHNQc1BzUHNQc1BzUHNQc1BzUHNQc1BzUHNQc1BzUHNQc1BzUHNQc1BzUHNQc1BzUHNQc1BzUHNQc1BzUHNQc1BzUHNQc1BzUHNQc1BzUHNQc1BzUHNQc1BzUHNQc1BzUHNQc1BzUHNQc1BzUHNQc1BzUHNQc1BzUHNQc1BzUHNQc1BzUHNQc1BzUHNQc1BzUHNQc1BzUHNQc1BzUHNQc1BzUHNQc1BzUHNQc1BzUHNQc1BzUHNQc1BzUHNQc1BzUHbQdeAF4ANQc1BzUHNQc1BzUHNQc1BzUHNQc1BzUHNQc1BzUHNQc1BzUHNQc1BzUHNQc1BzUHNQc1BzUHNQc1BzUHNQc1BzUHNQc1BzUHNQc1BzUHNQc1BzUHNQc1BzUHNQc1BzUHNQc1BzUHNQc1BzUHNQc1BzUHNQc1BzUHNQc1BzUHNQc1BzUHNQc1BzUHNQc1BzUHNQc1BzUHNQc1BzUHNQc1BzUHNQc1BzUHNQc1BzUHNQc1BzUHNQc1BzUHNQc1BzUHNQc1BzUHNQc1BzUHNQc1BzUHNQc1BzUHNQc1BzUHNQc1BzUHNQc1BzUHNQc1BzUHNQc1BzUHNQc1BzUHNQc1BzUHNQc1BzUHNQc1BzUHNQc1BzUHNQc1BzUHNQc1BzUHNQc1BzUHNQc1BzUHNQc1BzUHNQc1BzUHNQc1BzUHNQc1BzUHNQc1BzUHNQc1BzUHNQc1BzUHNQc1BzUHNQc1BzUHNQc1BzUHNQc1BzUHNQc1BzUHNQc1BzUHNQc1BzUHNQc1BzUHNQc1BzUHNQc1BzUHNQc1BzUHNQc1BzUHNQc1BzUHNQc1BzUHNQc1BzUHNQc1BzUHNQc1BzUHNQc1BzUHNQc1BzUHNQc1BzUHNQc1BzUHNQc1BzUHNQc1BzUHNQc1BzUHNQc1BzUHNQc1BzUHNQc1BzUHNQc1BzUHNQc1BzUHNQc1BzUHNQc1BzUHNQc1BzUHNQc1BzUHNQc1BzUHNQc1BzUHNQc1BzUHNQc1BzUHNQc1BzUHNQc1BzUHNQc1BzUHNQc1BzUHNQc1BzUHNQc1BzUHNQc1BzUHNQc1BzUHNQc1BzUHNQc1BzUHNQc1BzUHNQc1BzUHNQc1BzUHNQc1BzUHNQc1BzUHNQc1BzUHNQc1BzUHNQc1BzUHNQc1BzUHNQc1BzUHNQc1BzUHNQc1BzUHNQc1BzUHNQc1BzUHNQc1BzUHNQc1BzUHNQc1BzUHNQc1BzUHNQc1BzUHNQc1BzUHNQc1BzUHNQc1BzUHNQc1BzUHNQc1BzUHNQc1BzUHNQc1BzUHNQc1BzUHNQc1BzUHNQc1BzUHNQc1BzUHNQc1BzUHNQc1BzUHNQc1BzUHNQc1BzUHNQc1BzUHNQc1BzUHNQc1BzUHNQc1BzUHNQc1BzUHNQc1BzUHNQc1BzUHNQc1BzUHNQc1BzUHNQc1BzUHNQc1BzUHNQc1BzUHNQc1BzUHNQc1BzUHNQc1BzUHNQc1BzUHNQc1BzUHNQc1BzUHNQc1BzUHNQc1BzUHNQc1BzUHNQc1BzUHNQc1BzUHNQc1BzUHNQc1BzUHNQc1BzUHNQc1BzUHNQc1BzUHNQc1BzUHNQc1BzUHNQc1BzUHNQc1BzUHNQc1BzUHNQc1BzUHNQc1BzUHNQc1BzUHNQc1BzUHNQc1BzUHNQc1BzUHNQc1BzUHNQc1BzUHNQc1BzUHNQc1BzUHNQc1BzUHNQc1BzUHNQc1BzUHNQc1BzUHNQc1BzUHNQc1BzUHNQc1BzUHNQc1BzUHNQc1BzUHNQc1BzUHNQc1BzUHNQc1BzUHNQc1BzUHNQc1BzUHNQc1BzUHNQc1BzUHNQc1BzUHNQc1BzUHNQc1BzUHNQc1BzUHNQc1BzUHNQc1BzUHNQc1BzUHNQc1BzUHNQc1BzUHNQc1BzUHNQc1BzUHNQc1BzUHNQc1BzUHNQc1BzUHNQc1BzUHNQc1BzUHNQc1BzUHNQc1BzUHNQc1BzUHNQc1BzUHNQc1BzUHNQd1B30HNQc1BzUHNQc1BzUHNQc1BzUHNQc1BzUHNQc1BzUHNQc1BzUHNQc1BzUHNQc1BzUHNQc1BzUHNQc1BzUHNQc1BzUHNQc1B4MH2gaKB68EgACAAIAAgACAAIAAgACAAI8HlwdeAJ8HpweAAIAArwe3B14AXgC/B8UHygcwANAH2AfgB4AA6AfwBz4B+AcACFwBCAgPCBcIogEYAR8IJwiAAC8INwg/CCADRwhPCFcIXwhnCEoDGgSAAIAAgABvCHcIeAh5CHoIewh8CH0Idwh4CHkIegh7CHwIfQh3CHgIeQh6CHsIfAh9CHcIeAh5CHoIewh8CH0Idwh4CHkIegh7CHwIfQh3CHgIeQh6CHsIfAh9CHcIeAh5CHoIewh8CH0Idwh4CHkIegh7CHwIfQh3CHgIeQh6CHsIfAh9CHcIeAh5CHoIewh8CH0Idwh4CHkIegh7CHwIfQh3CHgIeQh6CHsIfAh9CHcIeAh5CHoIewh8CH0Idwh4CHkIegh7CHwIfQh3CHgIeQh6CHsIfAh9CHcIeAh5CHoIewh8CH0Idwh4CHkIegh7CHwIfQh3CHgIeQh6CHsIfAh9CHcIeAh5CHoIewh8CH0Idwh4CHkIegh7CHwIfQh3CHgIeQh6CHsIfAh9CHcIeAh5CHoIewh8CH0Idwh4CHkIegh7CHwIfQh3CHgIeQh6CHsIfAh9CHcIeAh5CHoIewh8CH0Idwh4CHkIegh7CHwIfQh3CHgIeQh6CHsIfAh9CHcIeAh5CHoIewh8CH0Idwh4CHkIegh7CHwIfQh3CHgIeQh6CHsIfAh9CHcIeAh5CHoIewh8CH0Idwh4CHkIegh7CHwIfQh3CHgIeQh6CHsIfAh9CHcIeAh5CHoIewh8CH0Idwh4CHkIegh7CHwIfQh3CHgIeQh6CHsIfAh9CHcIeAh5CHoIewh8CH0Idwh4CHkIegh7CHwIfQh3CHgIeQh6CHsIfAh9CHcIeAh5CHoIewh8CH0Idwh4CHkIegh7CHwIfQh3CHgIeQh6CHsIfAh9CHcIeAh5CHoIewh8CH0Idwh4CHkIegh7CHwIfQh3CHgIeQh6CHsIfAh9CHcIeAh5CHoIewh8CH0Idwh4CHkIegh7CHwIfQh3CHgIeQh6CHsIfAh9CHcIeAh5CHoIewh8CH0Idwh4CHkIegh7CHwIhAiLCI4IMAAwADAAMAAwADAAMAAwADAAMAAwADAAMAAwADAAMAAwADAAMAAwADAAMAAwADAAMAAwADAAMAAwADAAMAAwAJYIlgiWCJYIlgiWCJYIlgiWCJYIlgiWCJYIlgiWCJYIlgiWCJYIlgiWCJYIlgiWCJYIlgiWCJYIlgiWCJYIlggwADAAMAAwADAAMAAwADAAMAAwADAAMAAwADAAMAAwADAAMAAwADAAMAAwADAAMAAwADAAMAAwADAAMAAwADAAMAAwADAAMAAwADAAMAAwADAAMAAwADAAMAAwADAAMAAwADAAMAAwADAAMAAwADAAMAAwADAAMAAwADAAMAAwADAAMAAwADAAMAAwADAAMAAwADAAMAAwADAAMAAwADAAMAAwADAAMAAwADAAMAAwADAAMAAwADAAMAAwADAAMAAwADAAMAAwADAAMAAwADAAMAAwADAAMAAwADAAMAAwADAAMAAwADAAMAAwADAAMAAwADAAMAAwADAAMAAwADAAMAAwADAAMAAwADAAMAAwADAAMAAwADAAMAAwADAAMAAwADAAMAAwADAAMAAwADAAMAAwADAAMAAwADAAMAAwADAAMAAwADAAMAAwADAAMAAwADAAMAAwADAAMAAwADAAMAAwADAAMAAwADAAMAAwADAAMAAwADAAMAAwADAAMAAwADAAMAAwADAAMAAwADAANQc1BzUHNQc1BzUHNQc1BzUHNQc1B54INQc1B6II2gaqCLIIugiAAIAAvgjGCIAAgACAAIAAgACAAIAAgACAAIAAywiHAYAA0wiAANkI3QjlCO0I9Aj8CIAAgACAAAIJCgkSCRoJIgknCTYHLwk3CZYIlgiWCJYIlgiWCJYIlgiWCJYIlgiWCJYIlgiWCJYIlgiWCJYIlgiWCJYIlgiWCJYIlgiWCJYIlgiWCJYIlgiAAIAAAAFAAXgBeAGAAcABeAHwAQACQAKAArQC9AJ4AXgBeAE0A3gBRAN4A7AD8AMwBGgEAAKcBNwEFAUwBXAF4QkhCmEKnArcCgAHHAsABz4LAAcABwAHAAd+C6ABoAG+C/4LAAcABwAHAAc+DF4MAAcAB54M3gweDV4Nng3eDaABoAGgAaABoAGgAaABoAGgAaABoAGgAaABoAGgAaABoAGgAaABoAEeDqABVg6WDqABoQ6gAaABoAHXDvcONw/3DvcO9w73DvcO9w73DvcO9w73DvcO9w73DvcO9w73DvcO9w73DvcO9w73DvcO9w73DvcO9w73DvcO9w73DncPAAcABwAHAAcABwAHAAcABwAHAAcABwAHAAcABwAHAAcABwAHAAcABwAHAAcABwAHAAcABwAHAAcABwAHAAcABwAHAAcABwAHAAcABwAHAAcABwAHAAcABwAHAAcABwAHAAcABwAHAAcABwAHAAcABwAHAAcABwAHAAcABwAHAAcABwAHAAcABwAHAAcABwAHAAcABwAHAAcABwAHAAcABwAHAAcABwAHAAcABwAHAAcABwAHAAcABwAHAAcABwAHAAcABwAHAAcABwAHAAcABwAHAAcABwAHAAcABwAHAAcABwAHAAcABwAHAAcABwAHAAcABwAHAAcABwAHAAcABwAHAAcABwAHAAcABwAHAAcABwAHAAcABwAHAAcABwAHAAcABwAHAAcABwAHAAcABwAHAAcABwAHAAcABwAHAAcABwAHAAcABwAHAAcABwAHAAcABwAHAAcABwAHAAcABwAHAAcABwAHAAcABwAHAAcABwAHAAcABwAHAAcABwAHAAcABwAHAAcABwAHAAcABwAHAAcABwAHAAcABwAHAAcABwAHAAcABwAHAAcABwAHAAcABwAHAAcABwAHAAcABwAHAAcABwAHAAcABwAHAAcABwAHAAcABwAHAAcABwAHAAcABwAHAAcABwAHAAcABwAHAAcABwAHAAcABwAHAAcABwAHAAcABwAHAAcABwAHAAcABwAHAAcABwAHAAcABwAHAAcABwAHAAcABwAHAAcABwAHAAcABwAHAAcABwAHAAcABwAHAAcABwAHAAcABwAHAAcABwAHAAcABwAHAAcABwAHAAcABwAHAAcABwAHAAcABwAHAAcABwAHAAcAB7cPPwlGCU4JMACAAIAAgABWCV4JYQmAAGkJcAl4CXwJgAkwADAAMAAwAIgJgACLCZMJgACZCZ8JowmrCYAAswkwAF4AXgB8AIAAuwkABMMJyQmAAM4JgADVCTAAMAAwADAAgACAAIAAgACAAIAAgACAAIAAqwYWBNkIMAAwADAAMADdCeAJ6AnuCR4E9gkwAP4JBQoNCjAAMACAABUK0wiAAB0KJAosCjQKgAAwADwKQwqAAEsKvQmdCVMKWwowADAAgACAALcEMACAAGMKgABrCjAAMAAwADAAMAAwADAAMAAwADAAMAAeBDAAMAAwADAAMAAwADAAMAAwADAAMAAwAIkEPQFzCnoKiQSCCooKkAqJBJgKoAqkCokEGAGsCrQKvArBCjAAMADJCtEKFQHZCuEK/gHpCvEKMAAwADAAMACAAIwE+QowAIAAPwEBCzAAMAAwADAAMACAAAkLEQswAIAAPwEZCyELgAAOCCkLMAAxCzkLMAAwADAAMAAwADAAXgBeAEELMAAwADAAMAAwADAAMAAwAEkLTQtVC4AAXAtkC4AAiQkwADAAMAAwADAAMAAwADAAbAtxC3kLgAuFC4sLMAAwAJMLlwufCzAAMAAwADAAMAAwADAAMAAwADAAMAAwADAAMAAwADAAMAAwADAAMAAwAIAAgACAAIAAgACAAIAAgACAAIAAgACAAIAAgACAAIAAgACAAIAAgACAAIAAgACAAIAAgACAAIAApwswADAAMACAAIAAgACvC4AAgACAAIAAgACAALcLMAAwADAAMAAwADAAMAAwADAAMAAwADAAMAAwADAAMAAwADAAMAAwADAAgACAAIAAgACAAIAAgACAAIAAgACAAIAAgACAAIAAgACAAIAAvwuAAMcLgACAAIAAgACAAIAAyguAAIAAgACAAIAA0QswADAAMAAwADAAMAAwADAAMAAwADAAMAAwADAAMAAwADAAMAAwADAAMAAwADAAMAAwADAAMAAwADAAMAAwADAAgACAAIAAgACAAIAAgACAAIAAgACAAIAAgACAANkLgACAAIAA4AswADAAMAAwADAAMAAwADAAMAAwADAAMAAwAIAAgACAAIAAgACAAIAAgACAAIAAgACAAIAAgACAAIAAgACJCR4E6AswADAAhwHwC4AA+AsADAgMEAwwADAAMAAwADAAMAAwADAAMAAwADAAMAAwADAAMAAwADAAMAAwADAAMAAwADAAMAAwADAAMACAAIAAGAwdDCUMMAAwAC0MNQc1BzUHNQc1BzUHNQc1BzUHNQc1BzUHNQc1BzUHNQc1BzUHNQc1BzUHNQc1BzUHNQc1BzUHNQc1BzUHNQc1BzUHNQc1BzUHNQc1BzUHNQc1BzUHNQc1BzUHNQc1BzUHNQc1BzUHNQc1BzUHNQc1BzUHNQc1BzUHNQc1BzUHNQw1BzUHNQc1BzUHNQc1BzUHNQc1BzUHNQc1BzUHNQc1BzUHNQc1BzUHNQc1BzUHPQwwADAAMAAwADAAMAAwADAAMAAwADAAMAAwADAAMAAwADAAMAAwADAAMAAwADAAMAAwADAAMAAwADAAMAAwADAAMAAwADAAMAAwADAAMAAwADUHNQc1BzUHNQc1BzUHNQc2BzAAMAA5DDUHNQc1BzUHNQc1BzUHNQc1BzUHNQdFDDAAMAAwADAAMAAwADAAMAAwADAAMAAwADAAMAAwADAAMAAwADAAMAAwADAAMAAwADAAMAAwADAAMAAwADAAMAAwADAAMAAwADAAMAAwADAAgACAAIAATQxSDFoMMAAwADAAMAAwADAAMAAwADAAMAAwADAAMAAwADAAMAAwADAAMAAwADAAMAAwADAAMAAwAF4AXgBeAF4AXgBeAF4AYgxeAGoMXgBxDHkMfwxeAIUMXgBeAI0MMAAwADAAMAAwAF4AXgCVDJ0MMAAwADAAMABeAF4ApQxeAKsMswy7DF4Awgy9DMoMXgBeAF4AXgBeAF4AXgBeAF4AXgDRDNkMeQBqCeAM3Ax8AOYM7Az0DPgMXgBeAF4AXgBeAF4AXgBeAF4AXgBeAF4AXgBeAF4AXgCgAAANoAAHDQ4NFg0wADAAMAAwADAAMAAwADAAMAAwADAAMAAwADAAMAAwADAAMAAwADAAMAAwADAAMAAwADAAMAAwADAAMAAwADAAMAAwADAAMAAwADAAMAAwADAAMAAeDSYNMAAwADAAMAAwADAAMAAwADAAMAAwADAAMAAwADAAMAAwADAAMAAwADAAMAAwADAAMAAwADAAMAAwADAAMAAwADAAMAAwADAAMAAwADAAMAAwADAAMAAwADAAMAAwADAAMAAwADAAMAAwADAAMAAwADAAMAAwADAAMAAwAIAAgACAAIAAgACAAC4NMABeAF4ANg0wADAAMAAwADAAMAAwADAAMAAwADAAMAAwADAAMAAwADAAMAAwADAAMAAwADAAMAAwADAAMAAwADAAMAAwADAAMAAwADAAMAAwAD4NRg1ODVYNXg1mDTAAbQ0wADAAMAAwADAAMAAwADAA2gbaBtoG2gbaBtoG2gbaBnUNeg3CBYANwgWFDdoGjA3aBtoG2gbaBtoG2gbaBtoG2gbaBtoG2gaUDZwNpA2oDdoG2gawDbcNvw3HDdoG2gbPDdYN3A3fDeYN2gbsDfMN2gbaBvoN/g3aBgYODg7aBl4AXgBeABYOXgBeACUG2gYeDl4AJA5eACwO2w3aBtoGMQ45DtoG2gbaBtoGQQ7aBtoG2gbaBtoG2gbaBtoG2gbaBtoG2gbaBtoG2gbaBtoG2gbaBtoG2gbaBtoG2gbaBtoG2gbaBtoG2gbaBtoG2gbaBtoG2gbaBtoG2gbaBtoG2gbaBtoG2gbaBtoG2gZJDjUHNQc1BzUHNQc1BzUHNQc1BzUHNQc1BzUHNQc1BzUHNQc1BzUHNQc1BzUHNQc1BzUHNQc1BzUHNQc1BzUHNQc1BzUHNQc1BzUHNQc1BzUHNQc1BzUHNQc1BzUHNQc1BzUHNQc1BzUHNQc1B1EO2gY1BzUHNQc1BzUHNQc1BzUHNQc1BzUHNQc1BzUHNQc1BzUHNQc1BzUHNQc1BzUHNQc1BzUHNQc1BzUHNQc1BzUHNQc1BzUHNQc1BzUHNQc1BzUHNQc1BzUHNQc1BzUHNQc1BzUHNQc1BzUHNQc1BzUHNQdZDjUHNQc1BzUHNQc1B2EONQc1BzUHNQc1BzUHNQc1BzUHNQc1BzUHNQc1BzUHNQc1BzUHNQc1BzUHNQc1BzUHNQc1BzUHNQc1BzUHNQc1BzUHNQc1BzUHNQc1BzUHNQc1BzUHNQc1BzUHNQc1BzUHNQc1BzUHNQc1BzUHNQc1BzUHNQc1BzUHNQc1BzUHaA41BzUHNQc1BzUHNQc1BzUHNQc1BzUHNQc1BzUHNQc1BzUHNQc1BzUHNQc1BzUHNQc1BzUHNQc1BzUHNQc1B3AO2gbaBtoG2gbaBtoG2gbaBtoG2gbaBtoG2gbaBtoG2gbaBtoG2gbaBtoG2gbaBtoG2gbaBtoG2gbaBtoG2gbaBtoG2gbaBtoG2gbaBtoG2gbaBtoG2gbaBtoG2gbaBtoG2gbaBtoG2gbaBtoG2gbaBtoG2gbaBtoG2gbaBtoG2gY1BzUHNQc1BzUHNQc1BzUHNQc1BzUHNQc1BzUHNQc1B2EO2gbaBtoG2gbaBtoG2gbaBtoG2gbaBtoG2gbaBtoG2gbaBtoG2gbaBtoG2gbaBtoG2gbaBtoG2gbaBtoG2gbaBtoG2gbaBtoG2gbaBtoG2gbaBtoG2gbaBtoG2gZJDtoG2gbaBtoG2gbaBtoG2gbaBtoG2gbaBtoG2gbaBtoG2gbaBtoG2gbaBtoG2gbaBtoG2gbaBtoG2gbaBtoG2gbaBtoG2gbaBtoG2gbaBtoG2gbaBtoG2gbaBtoG2gbaBtoG2gbaBtoG2gbaBtoG2gbaBtoG2gbaBtoG2gbaBkkOeA6gAKAAoAAwADAAMAAwAKAAoACgAKAAoACgAKAAgA4wADAAMAAwADAAMAAwADAAMAAwADAAMAAwADAAMAAwADAAMAAwADAAMAAwADAAMAAwADAAMAAwADAAMAAwADAAMAAwADAAMAAwADAAMAAwADAAMAAwADAAMAAwADAAMAD//wQABAAEAAQABAAEAAQABAAEAA0AAwABAAEAAgAEAAQABAAEAAQABAAEAAQABAAEAAQABAAEAAQABAAEAAQABAAKABMAFwAeABsAGgAeABcAFgASAB4AGwAYAA8AGAAcAEsASwBLAEsASwBLAEsASwBLAEsAGAAYAB4AHgAeABMAHgBQAFAAUABQAFAAUABQAFAAUABQAFAAUABQAFAAUABQAFAAUABQAFAAUABQAFAAUABQAFAAFgAbABIAHgAeAB4AUABQAFAAUABQAFAAUABQAFAAUABQAFAAUABQAFAAUABQAFAAUABQAFAAUABQAFAAUABQABYADQARAB4ABAAAAAAAAAAAAAAAAAAAAAAAAAAAAAAAAAAAAAAAAAAAAAAAAAAAAAAAAAAAAAAAAAAAAAAAAAAAAAAAAAAAAAAAAAAAAAAAAAAAAAAAAAAAAAAAAAAAAAAAAAAAAAAAAAAAAAAAAAAAAAAAAAAAAAAAAAAAAAAAAAAAAAAAAAAAACsAKwArACsAKwArACsAKwArACsAKwArACsAKwArACsAKwArACsAKwArACsAKwArACsAKwArACsAKwArACsAKwArACsAKwArACsAKwArACsAKwArACsAKwArACsAKwArACsAKwArACsAKwArACsAKwArACsAKwArACsAKwArACsABAAEAAQABAAEAAUABAAEAAQABAAEAAQABAAEAAQABAAEAAQABAAEAAQABAAEAAQABAAEAAQABAAEAAQABAAEAAkAFgAaABsAGwAbAB4AHQAdAB4ATwAXAB4ADQAeAB4AGgAbAE8ATwAOAFAAHQAdAB0ATwBPABcATwBPAE8AFgBQAFAAUABQAFAAUABQAFAAUABQAFAAUABQAFAAUABQAFAAUABQAFAAUABQAFAAHQAeAB4AHgAeAB4AHgAeAB4AHgAeAB4AHgAeAB4AHgAeAB4AHgAeAB4AHgAeAB4AHgAeAB4AHgAeAB4AHgAeAB0AHgAeAB4AHgAeAB4AHgAeAB4AHgAeAB4AHgAeAB4AHgAeAB4AHgAeAB4AHgAeAB4AHgAeAB4AHgAeAB4AHgAeAB4AHgAeAB4AHgAeAB4AHgAeAB4AHgAeAB4AHgAeAB4AHgAeAB4AHgAeAB4AHgAeAB4AHgAeAB4AHgAeAB4AHgAeAB4AHgBQAB4AHgAeAB4AUABQAFAAUAAeAB4AHgAeAB4AHgAeAB4AHgAeAB4AHgAeAB4AHgAeAB4AHgAeAB4AHgAeAB4AHgAeAB4AHgAeAB4AHgAeAB4AHgAeAB4AHgAeAB4AHgAeAB4AHgAeAB4AHgAeAB4AHgAeAB4AHgAeAB4AHgAeAB4AHgAeAB4AHgBQAFAAUABQAFAAUABQAFAAUABQAFAAUABQAFAAUABQAFAAUABQAFAAUABQAFAAUABQAFAAUABQAFAAUABQAFAAUABQAFAAUABQAFAAUABQAFAAUABQAFAAUABQAFAAUABQAFAAUABQAFAAUABQAFAAUABQAFAAUABQAFAAUABQAFAAUAAeAB4AHgAeAFAATwBAAE8ATwBPAEAATwBQAFAATwBQAB4AHgAeAB4AHgAeAB0AHQAdAB0AHgAdAB4ADgBQAFAAUABQAFAAHgAeAB4AHgAeAB4AHgBQAB4AUAAeAB4AHgAeAB4AHgAeAB4AHgAeAB4AHgAeAB4AHgAeAB4ABAAEAAQABAAEAAQABAAEAAQABAAEAAQABAAEAAQABAAEAAQABAAEAAQABAAEAAQABAAEAAQABAAEAAQABAAEAAQABAAEAAQABAAEAAQABAAEAAQABAAEAAQABAAEAAQABAAEAAQABAAEAAQABAAEAAQABAAEAAQABAAEAAQABAAEAAQABAAJAAQABAAEAAQABAAEAAQABAAEAAQABAAEAAkACQAJAAkACQAJAAkABAAEAAQABAAEAAQABAAEAAQABAAEAAQABAAeAB4AHgAeAFAAHgAeAB4AKwArAFAAUABQAFAAGABQACsAKwArACsAHgAeAFAAHgBQAFAAUAArAFAAKwAeAB4AHgAeAB4AHgAeAB4AHgAeAB4AHgAeAB4AHgAeAB4AHgAeAB4AKwAeAB4AHgAeAB4AHgAeAB4AHgAeAB4AHgAeAB4AHgAeAB4AHgAeAB4AHgAeAB4AHgAeAB4AHgAeAB4AHgAeAB4ABAAEAAQABAAEAAQABAAeAB4AHgAeAB4AHgAeAB4AHgAeAB4AHgAeAB4AHgAeAB4AHgAeAB4AHgAeAB4AHgAeAB4AHgAeAB4AHgAeAB4AHgAeAB4AHgAeAB4AHgAeAB4AHgAeAB4AHgAeAB4AHgAeAB4AHgAeAB4AHgArAFAAUABQAFAAUABQAFAAUABQAFAAUABQAFAAUABQAFAAUABQAFAAUABQAFAAUABQAFAAUAArACsAUAAeAB4AHgAeAB4AHgArAFAAUABQAFAAUABQAFAAUABQAFAAUABQAFAAUABQAFAAUABQAFAAUABQAFAAUABQAFAAUABQAFAAUABQAFAAKwAYAA0AKwArAB4AHgAbACsABAAEAAQABAAEAAQABAAEAAQABAAEAAQABAAEAAQABAAEAAQABAAEAAQABAAEAAQABAAEAAQABAAEAAQABAAEAAQABAAEAAQABAAEAAQABAAEAAQABAAEAAQADQAEAB4ABAAEAB4ABAAEABMABAArACsAKwArACsAKwArACsAVgBWAFYAVgBWAFYAVgBWAFYAVgBWAFYAVgBWAFYAVgBWAFYAVgBWAFYAVgBWAFYAVgBWAFYAKwArACsAKwArAFYAVgBWAB4AHgArACsAKwArACsAKwArACsAKwArACsAHgAeAB4AHgAeAB4AHgAeAB4AGgAaABoAGAAYAB4AHgAEAAQABAAEAAQABAAEAAQABAAEAAQAEwAEACsAEwATAFAAUABQAFAAUABQAFAAUABQAFAAUABQAFAAUABQAFAAUABQAFAAUABQAFAAUABQAFAAUABQAFAAUABQAFAAUABQAFAAUAAEAAQABAAEAAQABAAEAAQABAAEAAQABAAEAAQABAAEAAQABAAEAAQABABLAEsASwBLAEsASwBLAEsASwBLABoAGQAZAB4AUABQAAQAUABQAFAAUABQAFAAUABQAFAAUABQAFAAUABQAFAAUABQAFAAUABQAFAAUABQABMAUAAEAAQABAAEAAQABAAEAB4AHgAEAAQABAAEAAQABABQAFAABAAEAB4ABAAEAAQABABQAFAASwBLAEsASwBLAEsASwBLAEsASwBQAFAAUAAeAB4AUAAeAB4AHgAeAB4AHgAeAB4AHgAeAB4AHgAeAB4AKwAeAFAABABQAFAAUABQAFAAUABQAFAAUABQAFAAUABQAFAAUABQAFAAUABQAFAAUABQAFAAUABQAFAAUABQAFAAUAAEAAQABAAEAAQABAAEAAQABAAEAAQABAAEAAQABAAEAAQABAAEACsAKwBQAFAAUABQAFAAUABQAFAAUABQAFAAUABQAFAAUABQAFAAUABQAFAAUABQAFAAUABQAFAAUABQAFAAUABQAFAAUABQAFAAUABQAFAAUABQAFAAUABQAFAAUABQAFAAUABQAFAAUABQAFAABAAEAAQABAAEAAQABAAEAAQABAAEAFAAKwArACsAKwArACsAKwArACsAKwArACsAKwArAEsASwBLAEsASwBLAEsASwBLAEsAUABQAFAAUABQAFAAUABQAFAAUABQAFAAUABQAFAAUABQAFAAUABQAFAAUABQAFAAUABQAFAAUABQAFAAUABQAFAABAAEAAQABAAEAAQABAAEAAQAUABQAB4AHgAYABMAUAArACsAKwArACsAUABQAFAAUABQAFAAUABQAFAAUABQAFAAUABQAFAAUABQAFAAUABQAFAAUAAEAAQABAAEAFAABAAEAAQABAAEAFAABAAEAAQAUAAEAAQABAAEAAQAKwArAB4AHgAeAB4AHgAeAB4AHgAeAB4AHgAeAB4AHgAeACsAUABQAFAAUABQAFAAUABQAFAAUABQAFAAUABQAFAAUABQAFAAUABQAFAAUABQAFAAUAAEAAQABAArACsAHgArAFAAUABQAFAAUABQAFAAUABQAFAAUAArACsAKwArACsAKwArACsAKwArACsAKwArACsAKwArACsAKwArACsAKwBQAFAAUABQAFAAUABQAFAAUABQAFAAUABQAFAAUABQAFAAUABQAFAAUAArAFAAUABQAFAAUABQAFAAUAArACsAKwArACsAKwArACsAKwArACsAKwArACsAKwArACsAKwArACsAKwArAAQABAAEAAQABAAEAAQABAAEAAQABAAEAAQABAAeAAQABAAEAAQABAAEAAQABAAEAAQABAAEAAQABAAEAAQABAAEAAQABAAEAAQABAAEAAQABAAEAAQABABQAFAAUABQAFAAUABQAFAAUABQAFAAUABQAFAAUABQAFAAUABQAFAAUABQAFAAUABQAFAAUABQAFAAUAAEAAQABABQAAQABAAEAAQABAAEAAQABAAEAAQABAAEAAQABAAEAAQABAAEAFAABAAEAAQABAAEAAQABABQAFAAUABQAFAAUABQAFAAUABQAAQABAANAA0ASwBLAEsASwBLAEsASwBLAEsASwAeAFAAUABQAFAAUABQAFAAUABQAFAAUABQAFAAUABQAFAABAAEAAQAKwBQAFAAUABQAFAAUABQAFAAKwArAFAAUAArACsAUABQAFAAUABQAFAAUABQAFAAUABQAFAAUABQACsAUABQAFAAUABQAFAAUAArAFAAKwArACsAUABQAFAAUAArACsABABQAAQABAAEAAQABAAEAAQAKwArAAQABAArACsABAAEAAQAUAArACsAKwArACsAKwArACsABAArACsAKwArAFAAUAArAFAAUABQAAQABAArACsASwBLAEsASwBLAEsASwBLAEsASwBQAFAAGgAaAFAAUABQAFAAUABMAB4AGwBQAB4AKwArACsABAAEAAQAKwBQAFAAUABQAFAAUAArACsAKwArAFAAUAArACsAUABQAFAAUABQAFAAUABQAFAAUABQAFAAUABQACsAUABQAFAAUABQAFAAUAArAFAAUAArAFAAUAArAFAAUAArACsABAArAAQABAAEAAQABAArACsAKwArAAQABAArACsABAAEAAQAKwArACsABAArACsAKwArACsAKwArAFAAUABQAFAAKwBQACsAKwArACsAKwArACsASwBLAEsASwBLAEsASwBLAEsASwAEAAQAUABQAFAABAArACsAKwArACsAKwArACsAKwArACsABAAEAAQAKwBQAFAAUABQAFAAUABQAFAAUAArAFAAUABQACsAUABQAFAAUABQAFAAUABQAFAAUABQAFAAUABQACsAUABQAFAAUABQAFAAUAArAFAAUAArAFAAUABQAFAAUAArACsABABQAAQABAAEAAQABAAEAAQABAArAAQABAAEACsABAAEAAQAKwArAFAAKwArACsAKwArACsAKwArACsAKwArACsAKwArACsAUABQAAQABAArACsASwBLAEsASwBLAEsASwBLAEsASwAeABsAKwArACsAKwArACsAKwBQAAQABAAEAAQABAAEACsABAAEAAQAKwBQAFAAUABQAFAAUABQAFAAKwArAFAAUAArACsAUABQAFAAUABQAFAAUABQAFAAUABQAFAAUAAEAAQABAAEAAQAKwArAAQABAArACsABAAEAAQAKwArACsAKwArACsAKwArAAQABAArACsAKwArAFAAUAArAFAAUABQAAQABAArACsASwBLAEsASwBLAEsASwBLAEsASwAeAFAAUABQAFAAUABQAFAAKwArACsAKwArACsAKwArACsAKwAEAFAAKwBQAFAAUABQAFAAUAArACsAKwBQAFAAUAArAFAAUABQAFAAKwArACsAUABQACsAUAArAFAAUAArACsAKwBQAFAAKwArACsAUABQAFAAKwArACsAUABQAFAAUABQAFAAUABQAFAAUABQAFAAKwArACsAKwAEAAQABAAEAAQAKwArACsABAAEAAQAKwAEAAQABAAEACsAKwBQACsAKwArACsAKwArAAQAKwArACsAKwArACsAKwArACsAKwBLAEsASwBLAEsASwBLAEsASwBLAFAAUABQAB4AHgAeAB4AHgAeABsAHgArACsAKwArACsABAAEAAQABAArAFAAUABQAFAAUABQAFAAUAArAFAAUABQACsAUABQAFAAUABQAFAAUABQAFAAUABQAFAAUABQAFAAKwBQAFAAUABQAFAAUABQAFAAUABQAFAAUABQAFAAUABQACsAKwArAFAABAAEAAQABAAEAAQABAArAAQABAAEACsABAAEAAQABAArACsAKwArACsAKwArAAQABAArAFAAUABQACsAKwArACsAKwBQAFAABAAEACsAKwBLAEsASwBLAEsASwBLAEsASwBLACsAKwArACsAKwArACsAKwBQAFAAUABQAFAAUABQAB4AUAAEAAQABAArAFAAUABQAFAAUABQAFAAUAArAFAAUABQACsAUABQAFAAUABQAFAAUABQAFAAUABQAFAAUABQAFAAKwBQAFAAUABQAFAAUABQAFAAUABQACsAUABQAFAAUABQACsAKwAEAFAABAAEAAQABAAEAAQABAArAAQABAAEACsABAAEAAQABAArACsAKwArACsAKwArAAQABAArACsAKwArACsAKwArAFAAKwBQAFAABAAEACsAKwBLAEsASwBLAEsASwBLAEsASwBLACsAUABQACsAKwArACsAKwArACsAKwArACsAKwArACsAUABQAFAAUABQAFAAUABQAFAAUABQAFAAUABQAFAAUABQAFAAUABQAFAAUABQAFAAUABQAFAABAAEAFAABAAEAAQABAAEAAQABAArAAQABAAEACsABAAEAAQABABQAB4AKwArACsAKwBQAFAAUAAEAFAAUABQAFAAUABQAFAAUABQAFAABAAEACsAKwBLAEsASwBLAEsASwBLAEsASwBLAFAAUABQAFAAUABQAFAAUABQABoAUABQAFAAUABQAFAAKwArAAQABAArAFAAUABQAFAAUABQAFAAUABQAFAAUABQAFAAUABQAFAAUABQACsAKwArAFAAUABQAFAAUABQAFAAUABQAFAAUABQAFAAUABQAFAAUABQAFAAUAArAFAAUABQAFAAUABQAFAAUABQACsAUAArACsAUABQAFAAUABQAFAAUAArACsAKwAEACsAKwArACsABAAEAAQABAAEAAQAKwAEACsABAAEAAQABAAEAAQABAAEACsAKwArACsAKwArAEsASwBLAEsASwBLAEsASwBLAEsAKwArAAQABAAeACsAKwArACsAKwArACsAKwArACsAKwArAFwAXABcAFwAXABcAFwAXABcAFwAXABcAFwAXABcAFwAXABcAFwAXABcAFwAXABcAFwAXABcAFwAXABcAFwAXAAqAFwAXAAqACoAKgAqACoAKgAqACsAKwArACsAGwBcAFwAXABcAFwAXABcACoAKgAqACoAKgAqACoAKgAeAEsASwBLAEsASwBLAEsASwBLAEsADQANACsAKwArACsAKwBcAFwAKwBcACsAKwBcAFwAKwBcACsAKwBcACsAKwArACsAKwArAFwAXABcAFwAKwBcAFwAXABcAFwAXABcACsAXABcAFwAKwBcACsAXAArACsAXABcACsAXABcAFwAXAAqAFwAXAAqACoAKgAqACoAKgArACoAKgBcACsAKwBcAFwAXABcAFwAKwBcACsAKgAqACoAKgAqACoAKwArAEsASwBLAEsASwBLAEsASwBLAEsAKwArAFwAXABcAFwAUAAOAA4ADgAOAB4ADgAOAAkADgAOAA0ACQATABMAEwATABMACQAeABMAHgAeAB4ABAAEAB4AHgAeAB4AHgAeAEsASwBLAEsASwBLAEsASwBLAEsAUABQAFAAUABQAFAAUABQAFAAUAANAAQAHgAEAB4ABAAWABEAFgARAAQABABQAFAAUABQAFAAUABQAFAAKwBQAFAAUABQAFAAUABQAFAAUABQAFAAUABQAFAAUABQAFAAUABQAFAAUABQAFAAUAArACsAKwArAAQABAAEAAQABAAEAAQABAAEAAQABAAEAAQABAANAAQABAAEAAQABAANAAQABABQAFAAUABQAFAABAAEAAQABAAEAAQABAAEAAQABAAEACsABAAEAAQABAAEAAQABAAEAAQABAAEAAQABAAEAAQABAAEAAQABAAEAAQABAAEAAQABAAEAAQABAAEAAQABAAEACsADQANAB4AHgAeAB4AHgAeAAQAHgAeAB4AHgAeAB4AKwAeAB4ADgAOAA0ADgAeAB4AHgAeAB4ACQAJACsAKwArACsAKwBcAFwAXABcAFwAXABcAFwAXABcAFwAXABcAFwAXABcAFwAXABcAFwAXABcAFwAXABcAFwAXABcAFwAXABcAFwAXABcAFwAKgAqACoAKgAqACoAKgAqACoAKgAqACoAKgAqACoAKgAqACoAKgAqAFwASwBLAEsASwBLAEsASwBLAEsASwANAA0AHgAeAB4AHgBcAFwAXABcAFwAXAAqACoAKgAqAFwAXABcAFwAKgAqACoAXAAqACoAKgBcAFwAKgAqACoAKgAqACoAKgBcAFwAXAAqACoAKgAqAFwAXABcAFwAXABcAFwAXABcAFwAXABcAFwAKgAqACoAKgAqACoAKgAqACoAKgAqACoAXAAqAEsASwBLAEsASwBLAEsASwBLAEsAKgAqACoAKgAqACoAUABQAFAAUABQAFAAKwBQACsAKwArACsAKwBQACsAKwBQAFAAUABQAFAAUABQAFAAUABQAFAAUABQAFAAUABQAFAAUABQAFAAUABQAFAAUABQAFAAUAAeAFAAUABQAFAAWABYAFgAWABYAFgAWABYAFgAWABYAFgAWABYAFgAWABYAFgAWABYAFgAWABYAFgAWABYAFgAWABYAFgAWABYAFkAWQBZAFkAWQBZAFkAWQBZAFkAWQBZAFkAWQBZAFkAWQBZAFkAWQBZAFkAWQBZAFkAWQBZAFkAWQBZAFkAWQBaAFoAWgBaAFoAWgBaAFoAWgBaAFoAWgBaAFoAWgBaAFoAWgBaAFoAWgBaAFoAWgBaAFoAWgBaAFoAWgBaAFoAUABQAFAAUABQAFAAUABQAFAAKwBQAFAAUABQACsAKwBQAFAAUABQAFAAUABQACsAUAArAFAAUABQAFAAKwArAFAAUABQAFAAUABQAFAAUABQACsAUABQAFAAUAArACsAUABQAFAAUABQAFAAUABQAFAAUABQAFAAUABQAFAAUABQACsAUABQAFAAUAArACsAUABQAFAAUABQAFAAUAArAFAAKwBQAFAAUABQACsAKwBQAFAAUABQAFAAUABQAFAAUABQAFAAUABQAFAAUAArAFAAUABQAFAAUABQAFAAUABQAFAAUABQAFAAUABQAFAAUAArAFAAUABQAFAAKwArAFAAUABQAFAAUABQAFAAUABQAFAAUABQAFAAUABQAFAAUABQAFAAUABQAFAAUABQAFAAUABQACsAKwAEAAQABAAeAA0AHgAeAB4AHgAeAB4AHgBQAFAAUABQAFAAUABQAFAAUABQAFAAUABQAFAAUABQAFAAUABQAFAAKwArACsAUABQAFAAUABQAFAAUABQAFAAUABQAFAAUABQAFAAUAAeAB4AHgAeAB4AHgAeAB4AHgAeACsAKwArACsAKwArAFAAUABQAFAAUABQAFAAUABQAFAAUABQAFAAUABQAFAAUABQAFAAUABQAFAAKwArAFAAUABQAFAAUABQACsAKwANAFAAUABQAFAAUABQAFAAUABQAFAAUABQAFAAUABQAFAAUABQAFAAUABQAFAAUABQAFAAUABQAFAAUABQAFAAUAAeAB4AUABQAFAAUABQAFAAUABQAFAAUABQAFAAUABQAFAAUABQAA0AUABQAFAAUABQAFAAUABQAFAAUABQAFAAUABQAFAAUABQAFAAUABQAFAAUABQAFAAUABQABYAEQArACsAKwBQAFAAUABQAFAAUABQAFAAUABQAFAADQANAA0AUABQAFAAUABQAFAAUABQAFAAUABQACsAKwArACsAKwArACsAUABQAFAAUABQAFAAUABQAFAAUABQAFAAUAArAFAAUABQAFAABAAEAAQAKwArACsAKwArACsAKwArACsAKwArAFAAUABQAFAAUABQAFAAUABQAFAAUABQAFAAUABQAFAAUABQAAQABAAEAA0ADQArACsAKwArACsAKwArACsAKwBQAFAAUABQAFAAUABQAFAAUABQAFAAUABQAFAAUABQAFAAUAAEAAQAKwArACsAKwArACsAKwArACsAKwArACsAUABQAFAAUABQAFAAUABQAFAAUABQAFAAUAArAFAAUABQACsABAAEACsAKwArACsAKwArACsAKwArACsAKwArAFwAXABcAFwAXABcAFwAXABcAFwAXABcAFwAXABcAFwAXABcAFwAXAAqACoAKgAqACoAKgAqACoAKgAqACoAKgAqACoAKgAqACoAKgAqACoADQANABUAXAANAB4ADQAbAFwAKgArACsASwBLAEsASwBLAEsASwBLAEsASwArACsAKwArACsAKwBQAFAAUABQAFAAUABQAFAAUABQACsAKwArACsAKwArAB4AHgATABMADQANAA4AHgATABMAHgAEAAQABAAJACsASwBLAEsASwBLAEsASwBLAEsASwArACsAKwArACsAKwBQAFAAUABQAFAAUABQAFAAUABQAFAAUABQAFAAUABQAFAAUABQAFAAUABQAFAAUAArACsAKwArACsAKwArACsAUABQAFAAUABQAAQABABQAFAAUABQAFAAUABQAFAAUABQAFAAUABQAFAAUABQAFAAUABQAFAAUABQAFAAUABQAFAABABQACsAKwArACsAKwBQAFAAUABQAFAAUABQAFAAUABQAFAAUABQAFAAUABQAFAAUABQAFAAUABQACsAKwArACsAKwArACsAKwArACsAUABQAFAAUABQAFAAUABQAFAAUABQAFAAUABQAFAAUABQAFAAUABQAFAAUABQAFAAUABQAFAAUABQAFAAUAArAAQABAAEAAQABAAEAAQABAAEAAQABAAEACsAKwArACsABAAEAAQABAAEAAQABAAEAAQABAAEAAQAKwArACsAKwAeACsAKwArABMAEwBLAEsASwBLAEsASwBLAEsASwBLAFwAXABcAFwAXABcAFwAXABcAFwAXABcAFwAXABcAFwAXABcACsAKwBcAFwAXABcAFwAKwArACsAKwArACsAKwArACsAKwArAFwAXABcAFwAXABcAFwAXABcAFwAXABcACsAKwArACsAXABcAFwAXABcAFwAXABcAFwAXABcAFwAXABcAFwAXABcAFwAKwArACsAKwArACsASwBLAEsASwBLAEsASwBLAEsASwBcACsAKwArACoAKgBQAFAAUABQAFAAUABQAFAAUABQAFAAUABQAFAAUABQAFAAUABQAFAAUABQAFAABAAEAAQABAAEACsAKwAeAB4AXABcAFwAXABcAFwAXABcAFwAXABcAFwAXABcAFwAXABcAFwAXABcAFwAKgAqACoAKgAqACoAKgAqACoAKgArACoAKgAqACoAKgAqACoAKgAqACoAKgAqACoAKgAqACoAKgAqACoAKgAqACoAKgAqACoAKgAqACoAKgArACsABABLAEsASwBLAEsASwBLAEsASwBLACsAKwArACsAKwArAEsASwBLAEsASwBLAEsASwBLAEsAKwArACsAKwArACsAKgAqACoAKgAqACoAKgBcACoAKgAqACoAKgAqACsAKwAEAAQABAAEAAQABAAEAAQABAAEAAQABAAEAAQABAArAAQABAAEAAQABABQAFAAUABQAFAAUABQAFAAUABQAFAAUABQAFAAUABQAFAAUABQAFAAUABQAFAAUABQAFAAUAAEAAQABAAEAAQAUABQAFAAUABQAFAAUAArACsAKwArAEsASwBLAEsASwBLAEsASwBLAEsADQANAB4ADQANAA0ADQAeAB4AHgAeAB4AHgAeAB4AHgAeAAQABAAEAAQABAAEAAQABAAEAB4AHgAeAB4AHgAeAB4AHgAeACsAKwArAAQABAAEAFAAUABQAFAAUABQAFAAUABQAFAAUABQAFAAUABQAFAAUABQAFAAUABQAFAAUABQAFAAUABQAFAAUABQAAQABAAEAAQABAAEAAQABAAEAAQABAAEAAQAUABQAEsASwBLAEsASwBLAEsASwBLAEsAUABQAFAAUABQAFAAUABQAAQABAAEAAQABAAEAAQABAAEAAQABAAEAAQABAArACsAKwArACsAKwArACsAHgAeAB4AHgBQAFAAUABQAAQABAAEAAQABAAEAAQABAAEAAQABAAEAAQABAAEAAQABAAEAAQABAArACsAKwANAA0ADQANAA0ASwBLAEsASwBLAEsASwBLAEsASwArACsAKwBQAFAAUABLAEsASwBLAEsASwBLAEsASwBLAFAAUABQAFAAUABQAFAAUABQAFAAUABQAFAAUABQAFAAUABQAFAAUABQAFAAUABQAFAAUABQAFAAUABQAFAAUAANAA0AUABQAFAAUABQAFAAUABQAFAAKwArACsAKwArACsAKwArACsAKwArACsAKwArACsAKwArACsAKwArACsAKwArAB4AHgAeAB4AHgAeAB4AHgArACsAKwArACsAKwArACsABAAEAAQAHgAEAAQABAAEAAQABAAEAAQABAAEAAQABAAEAFAAUABQAFAABABQAFAAUABQAAQABAAEAFAAUAAEAAQABAArACsAKwArACsAKwAEAAQABAAEAAQABAAEAAQABAAEAAQABAAEAAQABAAEAAQABAAEAAQABAAEAAQABAAEAAQAKwAEAAQABAAEAAQAHgAeAB4AHgAeAB4AHgAeAB4AHgAeAB4AHgAeAB4AHgAeAB4AHgAeAB4AHgArACsAUABQAFAAUABQAFAAKwArAFAAUABQAFAAUABQAFAAUAArAFAAKwBQACsAUAArAB4AHgAeAB4AHgAeAB4AHgAeAB4AHgAeAB4AHgAeAB4AHgAeAB4AHgAeAB4AHgAeAB4AHgAeAB4AHgAeAB4AKwArAB4AHgAeAB4AHgAeAB4AHgAeAB4AHgAeAB4AHgAeAB4AHgAeAB4AHgAeACsAHgAeAB4AHgAeAB4AHgAeAFAAHgAeAB4AUABQAFAAKwAeAB4AHgAeAB4AHgAeAB4AHgAeAFAAUABQAFAAKwArAB4AHgAeAB4AHgAeACsAHgAeAB4AHgAeAB4AHgAeAB4AHgAeAB4AHgAeAB4AHgAeAB4AHgArACsAUABQAFAAKwAeAB4AHgAeAB4AHgAeAA4AHgArAA0ADQANAA0ADQANAA0ACQANAA0ADQAIAAQACwAEAAQADQAJAA0ADQAMAB0AHQAeABcAFwAWABcAFwAXABYAFwAdAB0AHgAeABQAFAAUAA0AAQABAAQABAAEAAQABAAJABoAGgAaABoAGgAaABoAGgAeABcAFwAdABUAFQAeAB4AHgAeAB4AHgAYABYAEQAVABUAFQAeAB4AHgAeAB4AHgAeAB4AHgAeAB4AHgANAB4ADQANAA0ADQAeAA0ADQANAAcAHgAeAB4AHgArAAQABAAEAAQABAAEAAQABAAEAAQAUABQACsAKwBPAFAAUABQAFAAUAAeAB4AHgAWABEATwBQAE8ATwBPAE8AUABQAFAAUABQAB4AHgAeABYAEQArAFAAUABQAFAAUABQAFAAUABQAFAAUABQAFAAKwArACsAGwAbABsAGwAbABsAGwAaABsAGwAbABsAGwAbABsAGwAbABsAGwAbABsAGwAaABsAGwAbABsAGgAbABsAGgAbABsAGwAbABsAGwAbABsAGwAbABsAGwAbABsAGwAbABsABAAEAAQABAAEAAQABAAEAAQABAAEAAQABAAEAAQABAAEACsAKwArACsAKwArACsAKwArACsAKwArACsAKwArAB4AHgBQABoAHgAdAB4AUAAeABoAHgAeAB4AHgAeAB4AHgAeAB4ATwAeAFAAGwAeAB4AUABQAFAAUABQAB4AHgAeAB0AHQAeAFAAHgBQAB4AUAAeAFAATwBQAFAAHgAeAB4AHgAeAB4AHgBQAFAAUABQAFAAHgAeAB4AHgAeAB4AHgAeAB4AHgAeAB4AHgAeAB4AHgBQAB4AUABQAFAAUABPAE8AUABQAFAAUABQAE8AUABQAE8AUABPAE8ATwBPAE8ATwBPAE8ATwBPAE8ATwBQAFAAUABQAE8ATwBPAE8ATwBPAE8ATwBPAE8AUABQAFAAUABQAFAAUABQAFAAHgAeAFAAUABQAFAATwAeAB4AKwArACsAKwAdAB0AHQAdAB0AHQAdAB0AHQAdAB4AHgAeAB4AHgAeAB4AHgAeAB4AHgAeAB4AHgAeAB4AHgAeAB4AHgAdAB4AHQAeAB4AHgAeAB4AHgAeAB4AHgAeAB4AHQAeAB0AHQAeAB4AHgAdAB0AHgAeAB0AHgAeAB4AHQAeAB0AGwAbAB4AHQAeAB4AHgAeAB0AHgAeAB0AHQAdAB0AHgAeAB0AHgAdAB4AHQAdAB0AHQAdAB0AHgAdAB4AHgAeAB4AHgAdAB0AHQAdAB4AHgAeAB4AHQAdAB4AHgAeAB4AHgAeAB4AHgAeAB4AHQAeAB4AHgAdAB4AHgAeAB4AHgAdAB4AHgAeAB4AHgAeAB4AHgAeAB4AHgAeAB4AHQAdAB4AHgAdAB0AHQAdAB4AHgAdAB0AHgAeAB0AHQAeAB4AHgAeAB4AHgAeAB4AHgAeAB4AHgAeAB4AHgAeAB4AHgAdAB0AHgAeAB0AHQAeAB4AHgAeAB4AHgAeAB4AHgAeAB4AHgAeAB0AHgAeAB4AHQAeAB4AHgAeAB4AHgAeAB0AHgAeAB4AHgAeAB4AHgAeAB4AHgAeAB4AHgAeAB4AHgAeAB4AHgAeAB4AHgAeAB4AHgAdAB4AHgAeAB4AHgAeAB4AHgAeAB4AHgAeAB4AHgAeABQAHgAeAB4AHgAeAB4AHgAeAB4AHgAeAB4AHgAeAB4AHgAWABEAFgARAB4AHgAeAB4AHgAeAB0AHgAeAB4AHgAeAB4AHgAlACUAHgAeAB4AHgAeAB4AHgAeAB4AFgARAB4AHgAeAB4AHgAeAB4AHgAeAB4AHgAeAB4AHgAeAB4AHgAeAB4AHgAeACUAJQAlACUAHgAeAB4AHgAeAB4AHgAeAB4AHgAeAB4AHgAeAB4AKwArACsAKwArACsAKwArACsAKwArACsAKwArACsAKwArACsAKwArACsAKwArACsAKwBPAE8ATwBPAE8ATwBPAE8ATwBPAE8ATwBPAE8ATwBPAE8ATwBPAE8ATwBPAE8ATwBPAE8ATwBPAE8ATwBPAE8AHQAdAB0AHQAdAB0AHQAdAB0AHQAdAB0AHQAdAB0AHQAdAB0AHQAdAB0AHQAdAB0AHQAdAB0AHQAdAB0AHQAdAB0AHQBPAE8ATwBPAE8ATwBPAE8ATwBPAE8ATwBPAE8ATwBPAE8ATwBPAE8ATwBQAB0AHQAdAB0AHQAdAB0AHQAdAB0AHQAdAB4AHgAeAB4AHQAdAB0AHQAdAB0AHQAdAB0AHQAdAB0AHQAdAB0AHQAdAB0AHQAdAB0AHgAeAB4AHgAeAB4AHgAeAB4AHgAeAB0AHQAdAB0AHQAdAB0AHQAdAB0AHQAdAB0AHQAdAB0AHgAeAB0AHQAdAB0AHgAeAB4AHgAeAB4AHgAeAB4AHgAdAB0AHgAdAB0AHQAdAB0AHQAdAB4AHgAeAB4AHgAeAB4AHgAdAB0AHgAeAB0AHQAeAB4AHgAeAB0AHQAeAB4AHgAeAB0AHQAdAB4AHgAdAB4AHgAdAB0AHQAdAB4AHgAeAB4AHgAeAB4AHgAeAB4AHgAeAB4AHgAeAB4AHQAdAB0AHQAeAB4AHgAeAB4AHgAeAB4AHgAdAB4AHgAeAB4AHgAeAB4AHgAeAB4AHgAeAB4AHgAeAB4AJQAlACUAJQAeAB0AHQAeAB4AHQAeAB4AHgAeAB0AHQAeAB4AHgAeACUAJQAdAB0AJQAeACUAJQAlACAAJQAlAB4AHgAeAB4AHgAeAB4AHgAeAB4AHgAeAB4AHgAeAB4AHgAeAB4AHgAeAB4AHgAeAB4AJQAlACUAHgAeAB4AHgAdAB4AHQAeAB4AHgAeAB4AHgAeAB4AHgAeAB4AHgAeAB4AHgAeAB4AHgAeAB4AHgAeAB4AHgAeAB4AHgAeAB4AHQAdAB4AHQAdAB0AHgAdACUAHQAdAB4AHQAdAB4AHQAeAB4AHgAeAB4AHgAeAB4AHgAeAB4AHgAeAB4AHgAlAB4AHgAeAB4AHgAeAB4AHgAeAB4AHgAeAB4AHgAeAB4AHgAeAB4AHgAeAB4AHgAeAB4AHgAeAB4AHgAeAB0AHQAeAB4AHgAeAB4AHgAeAB4AHgAeAB4AHgAeAB4AHgAeAB4AHgAeAB4AHgAeAB4AHgAeAB4AHgAeAB4AJQAlACUAJQAlACUAJQAlACUAJQAlACUAHQAdAB0AHQAlAB4AJQAlACUAHQAlACUAHQAdAB0AJQAlAB0AHQAlAB0AHQAlACUAJQAeAB0AHgAeAB4AHgAdAB0AJQAdAB0AHQAdAB0AHQAlACUAJQAlACUAHQAlACUAIAAlAB0AHQAlACUAJQAlACUAJQAlACUAHgAeAB4AJQAlACAAIAAgACAAHgAeAB4AHgAeAB4AHgAeAB4AHgAeAB4AHgAeAB4AHgAeAB4AHgAeAB4AHgAeAB4AHgAdAB4AHgAeABcAFwAXABcAFwAXAB4AEwATACUAHgAeAB4AFgARABYAEQAWABEAFgARABYAEQAWABEAFgARAE8ATwBPAE8ATwBPAE8ATwBPAE8ATwBPAE8ATwBPAE8ATwBPAE8ATwBPAE8AHgAeAB4AHgAeAB4AHgAeAB4AHgAeAB4AHgAWABEAHgAeAB4AHgAeAB4AHgAeAB4AHgAeAB4AHgAeAB4AHgAeAB4AHgAeAB4AHgAeAB4AHgAeAB4AFgARABYAEQAWABEAFgARABYAEQAeAB4AHgAeAB4AHgAeAB4AHgAeAB4AHgAeAB4AHgAeAB4AHgAeABYAEQAWABEAFgARABYAEQAWABEAFgARABYAEQAWABEAFgARABYAEQAWABEAHgAeAB4AHgAeAB4AHgAeAB4AHgAeAB4AHgAeAB4AHgAeAB4AHgAeAB4AHgAeAB4AHgAeAB4AFgARABYAEQAeAB4AHgAeAB4AHgAeAB4AHgAeAB4AHgAeAB4AHgAeAB4AHgAeAB4AHgAeAB4AHgAeAB4AHgAeABYAEQAeAB4AHgAeAB4AHgAeAB4AHgAeAB4AHgAeAB4AHgAeAB4AHgAeAB4AHgAeAB4AHQAdAB0AHQAdAB4AHgAeAB4AHgAeAB4AHgAeAB4AHgAeAB4AHgAeAB4AHgAeAB4AHgAeAB4AKwArAB4AHgAeAB4AHgAeAB4AHgAeAB4AHgAeAB4AHgAeAB4AHgAeAB4AHgAeAB4AHgAeAB4AHgAeAB4AKwArACsAHgAeAB4AHgAeAB4AHgAeAB4AHgAeAB4AKwAeAB4AHgAeAB4AHgAeAB4AHgArACsAKwArACsAKwArACsAKwArACsAKwArAB4AHgAeAB4AKwArACsAKwArACsAKwArACsAKwArACsAKwArACsAKwAeAB4AHgAeAB4AHgAeAB4AHgAeAB4AHgAeAB4AHgAEAAQABAAeAB4AKwArACsAKwArABMADQANAA0AUAATAA0AUABQAFAAUABQAFAAUABQACsAKwArACsAKwArACsAUAANACsAKwArACsAKwArACsAKwArACsAKwArACsAKwAEAFAAUABQAFAAUABQAFAAUABQAFAAUABQAFAAUABQAFAAUABQAFAAUABQAFAAUAArACsAKwArACsAKwArACsAKwBQAFAAUABQAFAAUABQACsAUABQAFAAUABQAFAAUAArAFAAUABQAFAAUABQAFAAKwBQAFAAUABQAFAAUABQACsAFwAXABcAFwAXABcAFwAXABcAFwAXABcAFwAXAA0ADQANAA0ADQANAA0ADQAeAA0AFgANAB4AHgAXABcAHgAeABcAFwAWABEAFgARABYAEQAWABEADQANAA0ADQATAFAADQANAB4ADQANAB4AHgAeAB4AHgAMAAwADQANAA0AHgANAA0AFgANAA0ADQANAA0ADQANACsAKwArACsAKwArACsAKwArACsAKwArACsAKwArACsAKwArACsAKwArACsAJQAlACUAJQAlACUAJQAlACUAJQAlACUAJQAlACUAJQAlACUAJQAlACUAJQAlACUAJQAlACsAJQAlACUAJQAlACUAJQAlACUAJQAlACUAJQAlACUAJQAlACUAJQAlACUAJQAlACUAJQAlACUAJQAlACUAJQAlACUAKwArACsAKwArACsAKwArACsAKwArACsAJQAlACUAJQAlACUAJQAlACUAJQAlACUAJQAlACUAJQAlACUAJQAlACUAJQArACsAKwArACsAKwArACsAKwArACsAKwArACsAKwArACsAKwAlACUAJQAlACUAJQAlACUAJQAlACUAJQArACsAKwArAA0AEQARACUAJQBHAFcAVwAWABEAFgARABYAEQAWABEAFgARACUAJQAWABEAFgARABYAEQAWABEAFQAWABEAEQAlAFcAVwBXAFcAVwBXAFcAVwBXAAQABAAEAAQABAAEACUAVwBXAFcAVwA2ACUAJQBXAFcAVwBHAEcAJQAlACUAKwBRAFcAUQBXAFEAVwBRAFcAUQBXAFcAVwBXAFcAVwBXAFcAVwBXAFcAVwBXAFcAVwBXAFcAVwBXAFcAVwBXAFcAVwBXAFEAVwBXAFcAVwBXAFcAVwBXAFcAVwBXAFcAVwBXAFcAVwBXAFcAVwBXAFcAVwBXAFcAVwBXAFcAVwBXAFcAVwBRAFcAUQBXAFEAVwBXAFcAVwBXAFcAUQBXAFcAVwBXAFcAVwBRAFEAKwArAAQABAAVABUARwBHAFcAFQBRAFcAUQBXAFEAVwBRAFcAUQBXAFcAVwBXAFcAVwBXAFcAVwBXAFcAVwBXAFcAVwBXAFcAVwBXAFcAVwBXAFcAVwBXAFEAVwBRAFcAUQBXAFcAVwBXAFcAVwBRAFcAVwBXAFcAVwBXAFEAUQBXAFcAVwBXABUAUQBHAEcAVwArACsAKwArACsAVwBXAFcAVwBXAFcAVwBXAFcAVwBXAFcAVwBXAFcAVwBXAFcAVwBXAFcAVwBXAFcAVwBXAFcAVwBXAFcAKwArAFcAVwBXAFcAVwBXAFcAVwBXAFcAVwBXAFcAVwBXAFcAVwBXAFcAVwBXAFcAVwBXAFcAVwBXAFcAVwBXAFcAVwBXAFcAVwBXAFcAVwArACUAJQBXAFcAVwBXACUAJQAlACUAJQAlACUAJQAlACUAVwBXAFcAVwBXAFcAVwBXAFcAVwBXAFcAVwBXAFcAVwBXAFcAVwBXAFcAVwBXAFcAVwBXAFcAKwArACsAKwArACUAJQAlACUAKwArACsAKwArACsAKwArACsAKwArACsAUQBRAFEAUQBRAFEAUQBRAFEAUQBRAFEAUQBRAFEAUQAlACUAJQAlACUAJQAlACUAJQAlACUAJQAlACUAJQAlACUAJQAlACUAJQAlACUAJQAlACUAJQAlACUAJQAlACsAVwBXAFcAVwBXAFcAVwBXAFcAVwAlACUAJQAlACUAJQAlACUAJQAlACUAJQAlACUAJQAlACUAJQAlACUAJQAlAE8ATwBPAE8ATwBPAE8ATwAlAFcAVwBXAFcAVwBXAFcAVwBXAFcAVwBXAFcAVwBXACUAJQAlACUAJQAlACUAJQAlACUAJQAlACUAJQAlACUAJQBXAFcAVwBXAFcAVwBXAFcAVwBXAFcAVwBXAFcAVwBXAFcAVwBXAFcAVwBXAFcAVwBXACUAJQAlACUAJQAlACUAJQAlACUAVwBXAFcAVwBXAFcAVwBXAFcAVwBXACUAJQAlACUAJQAlACUAJQAlACUAJQAlACUAJQAlACUAJQAlACUAJQAlAFcAVwBXAFcAVwBXAFcAVwBXAFcAVwBXAFcAVwBXAFcAVwBXAFcAVwBXAEcAVwBXAFcAVwBXAFcAVwBXAFcAVwBXAFcAVwBXAFcAKwArACsAJQAlACUAJQAlACUAJQAlACUAJQAlACUAJQAlACUAJQAlACUAJQArACsAKwArACsAKwArACsAKwBQAFAAUABQAFAAUABQAFAAUABQAFAAUABQAFAAUABQAFAADQATAA0AUABQAFAAUABQAFAAUABQAFAAUABQAFAAUABQAFAAUABLAEsASwBLAEsASwBLAEsASwBLAFAAUAArACsAKwArACsAKwArACsAKwArACsAKwArACsAKwArACsAKwArACsAHgAeAB4AHgAeAB4AHgAeAB4AHgAeAB4AHgAeAFAABAAEAAQABAAeAAQABAAEAAQABAAEAAQABAAEAAQAHgBQAB4AHgAeAB4AHgAeAB4AHgAeAB4AHgAeAB4AHgAeAB4AHgAeAB4AHgAeAB4AHgAeAB4AHgAeAB4AUABQAAQABABQAFAAUABQAFAAUABQAFAAUABQAFAAUABQAFAAUABQAAQABAAeAA0ADQANAA0ADQArACsAKwArACsAKwArACsAHgAeAB4AHgAeAB4AHgAeAB4AHgAeAB4AHgAeAB4AHgAeAB4AHgAeAB4AHgAeAFAAUABQAFAAUABQAFAAUABQAB4AHgAeAB4AHgAeAB4AHgAeAB4AHgAeAB4AHgAeAB4AUAAeAB4AHgAeAB4AHgAeAB4AHgAeAB4AHgAeAB4AHgBQAB4AHgAeAB4AHgAeAFAAHgAeAB4AHgAeAB4AHgAeAB4AHgAeAB4AHgAeAB4AHgAeAB4AHgArAB4AHgAeAB4AHgAeAB4AHgArACsAKwArACsAKwArACsAKwArACsAKwArACsAKwArACsAKwArACsAKwArACsAUABQAFAAUABQAFAAUABQAFAAUABQAAQAUABQAFAABABQAFAAUABQAAQAUABQAFAAUABQAFAAUABQAFAAUABQAFAAUABQAFAAUABQAFAAUABQAFAAUABQAAQABAAEAAQABAAeAB4AHgAeACsAKwArACsAUABQAFAAUABQAFAAHgAeABoAHgArACsAKwArACsAKwBQAFAAUABQAFAAUABQAFAAUABQAFAAUABQAFAAUABQAFAAUABQAFAADgAOABMAEwArACsAKwArACsAKwArACsABAAEAFAAUABQAFAAUABQAFAAUABQAFAAUABQAFAAUABQAFAAUABQAFAAUABQAFAAUABQAFAAUABQAFAAUABQAAQABAAEAAQABAAEACsAKwArACsAKwArACsAKwANAA0ASwBLAEsASwBLAEsASwBLAEsASwArACsAKwArACsAKwAEAAQABAAEAAQABAAEAAQABAAEAAQABAAEAAQABAAEAAQABABQAFAAUABQAFAAUAAeAB4AHgBQAA4AUAArACsAUABQAFAAUABQAFAABAAEAAQABAAEAAQABAAEAA0ADQBQAFAAUABQAFAAUABQAFAAUABQAFAAUABQAFAAUABQAFAAUABQAAQABAAEAAQABAAEAAQABAAEAAQABAAEAAQAKwArACsAKwArACsAKwArACsAKwArAB4AWABYAFgAWABYAFgAWABYAFgAWABYAFgAWABYAFgAWABYAFgAWABYAFgAWABYAFgAWABYAFgAWABYACsAKwArAAQAHgAeAB4AHgAeAB4ADQANAA0AHgAeAB4AHgArAFAASwBLAEsASwBLAEsASwBLAEsASwArACsAKwArAB4AHgBcAFwAXABcAFwAKgBcAFwAXABcAFwAXABcAFwAXABcAEsASwBLAEsASwBLAEsASwBLAEsAXABcAFwAXABcACsAUABQAFAAUABQAFAAUABQAFAABAAEAAQABAAEAAQABAAEAAQABAAEAAQABAAEACsAKwArACsAKwArACsAKwArAFAAUABQAAQAUABQAFAAUABQAFAAUABQAAQABAArACsASwBLAEsASwBLAEsASwBLAEsASwArACsAHgANAA0ADQBcAFwAXABcAFwAXABcAFwAXABcAFwAXABcAFwAXABcAFwAXABcAFwAXABcAFwAKgAqACoAXAAqACoAKgBcAFwAXABcAFwAXABcAFwAXABcAFwAXABcAFwAXABcAFwAXAAqAFwAKgAqACoAXABcACoAKgBcAFwAXABcAFwAKgAqAFwAKgBcACsAKwArACsAKwArACsAKwArACsAKwArACsAKwArACsAKwArACsAKwArACsAKwArAFwAXABcACoAKgBQAFAAUABQAFAAUABQAFAAUABQAFAABAAEAAQABAAEAA0ADQBQAFAAUAAEAAQAKwArACsAKwArACsAKwArACsAKwBQAFAAUABQAFAAUAArACsAUABQAFAAUABQAFAAKwArAFAAUABQAFAAUABQACsAKwArACsAKwArACsAKwArAFAAUABQAFAAUABQAFAAKwBQAFAAUABQAFAAUABQACsAUABQAFAAUABQAFAAUABQAFAAUABQAFAAUABQAFAAUABQAFAAUAAEAAQABAAEAAQABAAEAAQADQAEAAQAKwArAEsASwBLAEsASwBLAEsASwBLAEsAKwArACsAKwArACsAVABVAFUAVQBVAFUAVQBVAFUAVQBVAFUAVQBVAFUAVQBVAFUAVQBVAFUAVQBVAFUAVQBVAFUAVQBUAFUAVQBVAFUAVQBVAFUAVQBVAFUAVQBVAFUAVQBVAFUAVQBVAFUAVQBVAFUAVQBVAFUAVQBVACsAKwArACsAKwArACsAKwArACsAKwArAFkAWQBZAFkAWQBZAFkAWQBZAFkAWQBZAFkAWQBZAFkAWQBZAFkAKwArACsAKwBaAFoAWgBaAFoAWgBaAFoAWgBaAFoAWgBaAFoAWgBaAFoAWgBaAFoAWgBaAFoAWgBaAFoAWgBaAFoAKwArACsAKwAGAAYABgAGAAYABgAGAAYABgAGAAYABgAGAAYABgAGAAYABgAGAAYABgAGAAYABgAGAAYABgAGAAYABgAGAAYAVwBXAFcAVwBXAFcAVwBXAFcAVwBXAFcAVwBXACUAJQBXAFcAVwBXAFcAVwBXAFcAVwBXAFcAVwBXAFcAVwBXAFcAVwBXAFcAVwBXAFcAVwBXAFcAJQAlACUAJQAlACUAUABQAFAAUABQAFAAUAArACsAKwArACsAKwArACsAKwArACsAKwBQAFAAUABQAFAAKwArACsAKwArAFYABABWAFYAVgBWAFYAVgBWAFYAVgBWAB4AVgBWAFYAVgBWAFYAVgBWAFYAVgBWAFYAVgArAFYAVgBWAFYAVgArAFYAKwBWAFYAKwBWAFYAKwBWAFYAVgBWAFYAVgBWAFYAVgBWAFAAUABQAFAAUABQAFAAUABQAFAAUABQAFAAUABQAFAAUABQAB4AHgAeAB4AHgAeAB4AHgAeAB4AHgAeAB4AHgAeAB4AKwArACsAKwArACsAKwArACsAKwArACsAKwArACsAKwArAFAAUABQAFAAUABQAFAAUABQAFAAUABQAFAAUABQAFAAUABQAFAAUABQAFAAUABQAFAAUABQAFAAUABQAFAAEQAWAFAAUABQAFAAUABQAFAAUABQAFAAUABQAFAAUABQAFAAKwArAFAAUABQAFAAUABQAFAAUABQAFAAUABQAFAAUAArACsAKwArACsAKwArACsAKwArACsAKwArACsAKwArACsAKwArACsAKwArACsAKwBQAFAAUABQAFAAUABQAFAAUABQAFAAUAAaAB4AKwArAAQABAAEAAQABAAEAAQABAAEAAQABAAEAAQABAAEAAQAGAARABEAGAAYABMAEwAWABEAFAArACsAKwArACsAKwAEAAQABAAEAAQABAAEAAQABAAEAAQABAAEAAQABAAEACUAJQAlACUAJQAWABEAFgARABYAEQAWABEAFgARABYAEQAlACUAFgARACUAJQAlACUAJQAlACUAEQAlABEAKwAVABUAEwATACUAFgARABYAEQAWABEAJQAlACUAJQAlACUAJQAlACsAJQAbABoAJQArACsAKwArAFAAUABQAFAAUAArAFAAUABQAFAAUABQAFAAUABQAFAAUABQAFAAUABQAFAAUABQAFAAUABQAFAAUABQAFAAUABQAFAAUABQAFAAKwArAAcAKwATACUAJQAbABoAJQAlABYAEQAlACUAEQAlABEAJQBXAFcAVwBXAFcAVwBXAFcAVwBXABUAFQAlACUAJQATACUAVwBXAFcAVwBXAFcAVwBXAFcAVwBXAFcAVwBXAFcAVwBXAFcAVwBXAFcAVwBXAFcAVwBXABYAJQARACUAJQAlAFcAVwBXAFcAVwBXAFcAVwBXAFcAVwBXAFcAVwBXAFcAVwBXAFcAVwBXAFcAVwBXAFcAVwAWACUAEQAlABYAEQARABYAEQARABUAVwBRAFEAUQBRAFEAUQBRAFEAUQBRAFcAVwBXAFcAVwBXAFcAVwBXAFcAVwBXAFcAVwBXAFcAVwBXAFcAVwBXAFcAVwBXAFcAVwBXAFcAVwBXAFcAVwBXAEcARwArACsAVwBXAFcAVwBXAFcAKwArAFcAVwBXAFcAVwBXACsAKwBXAFcAVwBXAFcAVwArACsAVwBXAFcAKwArACsAGgAbACUAJQAlABsAGwArAB4AHgAeAB4AHgAeAB4AKwArACsAKwArACsAKwArACsAKwAEAAQABAAQAB0AKwArAFAAUABQAFAAUABQAFAAUABQAFAAUABQACsAUABQAFAAUABQAFAAUABQAFAAUABQAFAAUABQAFAAUABQAFAAUABQAFAAUAArAFAAUABQAFAAUABQAFAAUABQAFAAUABQAFAAUABQAFAAUABQAFAAKwBQAFAAKwBQAFAAUABQAFAAUABQAFAAUABQAFAAUABQAFAAUAArACsAUABQAFAAUABQAFAAUABQAFAAUABQAFAAUABQACsAKwBQAFAAUABQAFAAUABQAFAAUABQAFAAUABQAFAAUABQAFAAUABQAFAAUABQAFAAUABQAFAAUAArACsAKwArACsADQANAA0AKwArACsAKwBQAFAAUABQAFAAUABQAFAAUABQAFAAUABQAFAAUABQAFAAUABQAFAAUABQAFAAUABQACsAKwArAB4AHgAeAB4AHgAeAB4AHgAeAFAAUABQAFAAUABQAFAAUABQAFAAUABQAFAAUABQAFAAUABQAFAAUABQAFAAUABQAFAAHgAeAB4AHgAeAB4AHgAeAB4AHgAeAB4AHgBQAFAAHgAeAB4AKwAeAB4AHgAeAB4AHgAeAB4AHgAeAB4AHgArACsAKwArAB4AKwArACsAKwArACsAKwArACsAKwArACsAKwArACsAKwArACsAKwArACsAKwArACsAKwArACsAKwArACsAKwAeAB4AHgAeAB4AHgAeAB4AHgAeAB4AHgAeAB4AHgAeAB4AHgAeAB4AHgAeAB4AHgAeAB4AHgAeAB4ABAArACsAUABQAFAAUABQAFAAUABQAFAAUABQAFAAUABQAFAAUABQAFAAUABQAFAAUABQAFAAUABQAFAAUABQACsAKwArACsAKwArACsAKwArACsAKwArACsAKwArAAQAUABQAFAAUABQAFAAUABQAFAAUABQAFAAUABQAFAAUABQAFAAUABQAFAAUABQAFAAUABQAFAAKwArACsAKwArACsAKwArACsAUABQAFAAUABQAFAAUABQAFAAUABQAFAAUABQAFAAUABQAFAAUABQAFAAUAArACsAKwArACsAUABQAFAAUABQAFAAUABQAFAAUABQAFAAUABQAFAAUABQAFAAUABQAFAAUAAEAAQABAAEAAQAKwArACsAKwArAFAAUABQAFAAUABQAFAAUABQAFAAUABQAFAAUABQAFAAUABQAFAAUABQAFAAUABQAFAAUABQAFAAUABQACsADQBQAFAAUABQACsAKwArACsAUABQAFAAUABQAFAAUABQAA0AUABQAFAAUABQACsAKwArACsAKwArACsAKwArACsAUABQAFAAUABQAFAAUABQAFAAUABQAFAAUABQAFAAUABQAFAAUABQAFAAUABQAFAAUABQAFAAUABQAFAAKwArAFAAUABQAFAAUABQAFAAUABQAFAAUABQAFAAUABQAFAAUABQAFAAUAArACsAKwArAFAAUABQAFAAUABQAFAAUABQAFAAUABQAFAAUABQAFAAUABQAFAAUABQAFAAUABQAFAAUABQAFAAKwArACsAKwArACsAKwArAFAAUABQAFAAUABQAFAAUABQAFAAUABQAFAAUABQAFAAKwArACsAKwArACsAKwArACsAKwArAB4AKwArACsAKwArACsAKwArACsAKwArACsAKwArACsAKwBQAFAAUABQAFAAUAArACsAUAArAFAAUABQAFAAUABQAFAAUABQAFAAUABQAFAAUABQAFAAUABQAFAAUABQAFAAUABQACsAUABQACsAKwArAFAAKwArAFAAUABQAFAAUABQAFAAUABQAFAAUABQAFAAUABQAFAAUABQAFAAUABQAFAAUAArAA0AUABQAFAAUABQAFAAUABQAFAAUABQAFAAUABQAFAAUABQAFAAUABQAFAAUABQAB4AHgBQAFAAUABQAFAAUABQACsAKwArACsAKwArACsAUABQAFAAUABQAFAAUABQAFAAKwArACsAKwArACsAKwArACsAKwArACsAKwArACsAKwBQAFAAUABQAFAAUABQAFAAUABQAFAAUABQAFAAUABQAFAAUABQACsAUABQACsAKwArACsAKwBQAFAAUABQAFAAUABQAFAAUABQAFAAUABQAFAAUABQAFAAUABQAFAAUABQAFAAUABQAFAAUABQAFAAKwArACsADQBQAFAAUABQAFAAUABQAFAAUABQAFAAUABQAFAAUABQAFAAUABQAFAAUABQAFAAUABQAFAAKwArACsAKwArAB4AUABQAFAAUABQAFAAUABQAFAAUABQAFAAUABQAFAAUABQAFAAUABQAFAAUABQAFAAKwArACsAKwBQAFAAUABQAFAABAAEAAQAKwAEAAQAKwArACsAKwArAAQABAAEAAQAUABQAFAAUAArAFAAUABQACsAUABQAFAAUABQAFAAUABQAFAAUABQAFAAUABQAFAAUABQAFAAUABQAFAAUABQACsAKwArACsABAAEAAQAKwArACsAKwAEAFAAUABQAFAAUABQAFAAUAArACsAKwArACsAKwArACsADQANAA0ADQANAA0ADQANAB4AKwArACsAKwArACsAKwBQAFAAUABQAFAAUABQAFAAUABQAFAAUABQAFAAUABQAFAAUABQAFAAUABQAFAAUABQAFAAUABQAFAAUABQAB4AUABQAFAAUABQAFAAUABQAB4AUABQAFAAUABQAFAAUABQAFAAUABQAFAAUABQAFAAUABQAFAAUABQAFAAUABQAFAABAAEACsAKwArACsAUABQAFAAUABQAA0ADQANAA0ADQANABQAKwArACsAKwArACsAKwArACsAUABQAFAAUABQAFAAUABQAFAAUABQAFAAUABQAFAAUABQAFAAUABQAFAAUAArACsAKwANAA0ADQANAA0ADQANAFAAUABQAFAAUABQAFAAUABQAFAAUABQAFAAUABQAFAAUABQACsAKwArACsAKwArACsAHgAeAB4AHgArACsAKwArACsAKwArACsAKwArACsAKwBQAFAAUABQAFAAUABQACsAKwArACsAKwArACsAKwArACsAKwArACsAKwArACsAUABQAFAAUABQAFAAUABQAFAAUABQAFAAUABQAFAAUABQAFAAUAArACsAKwArACsAKwArACsAKwArACsAKwArAFAAUABQAFAAUABQAFAAUABQAFAAUABQAFAAUABQAFAAUABQAFAAKwArACsAKwArACsAKwBQAFAAUABQAFAAUAAEAAQABAAEAAQABAAEAA0ADQAeAB4AHgAeAB4AKwArACsAKwBQAFAAUABQAFAAUABQAFAAUABQAFAAUABQAFAAUABQAEsASwBLAEsASwBLAEsASwBLAEsAKwArACsAKwArACsAKwArACsAKwArACsAKwArACsABABQAFAAUABQAFAAUABQAFAAUABQAFAAUABQAFAAUABQAAQABAAEAAQABAAEAAQABAAEAAQABAAeAB4AHgANAA0ADQANACsAKwArACsAKwArACsAKwArACsAKwArACsAKwBQAFAAUABQAFAAUABQAFAAUABQAFAAUABQAFAAUABQAFAAKwArACsAKwArACsAKwBLAEsASwBLAEsASwBLAEsASwBLACsAKwArACsAKwArAFAAUABQAFAAUABQAFAABAAEAAQABAAEAAQABAAEAAQABAAEAAQABAAEACsASwBLAEsASwBLAEsASwBLAEsASwANAA0ADQANACsAKwArACsAKwArACsAKwArACsAKwArAFAAUABQAFAAUABQAFAAUABQAFAAUABQAFAAUABQAFAAUABQAFAABAAeAA4AUAArACsAKwArACsAKwArACsAKwAEAFAAUABQAFAADQANAB4ADQAeAAQABAAEAB4AKwArAEsASwBLAEsASwBLAEsASwBLAEsAUAAOAFAADQANAA0AKwBQAFAAUABQAFAAUABQAFAAUABQAFAAUABQAFAAUABQAFAAUABQAFAAKwArACsAKwArACsAKwArACsAKwArAFAAUABQAFAAUABQAFAAUABQAFAAUABQAFAAUABQAFAAUABQACsAUABQAFAAUABQAFAAUABQAFAAUABQAFAAUAAEAAQABAAEAAQABAAEAAQABAAEAAQABAANAA0AHgANAA0AHgAEACsAUABQAFAAUABQAFAAUAArAFAAKwBQAFAAUABQACsAUABQAFAAUABQAFAAUABQAFAAUABQAFAAUABQAFAAKwBQAFAAUABQAFAAUABQAFAAUABQAA0AKwArACsAKwArACsAUABQAFAAUABQAFAAUABQAFAAUABQAFAAUABQAFAAUAAEAAQABAAEAAQABAAEAAQABAAEAAQAKwArACsAKwArAEsASwBLAEsASwBLAEsASwBLAEsAKwArACsAKwArACsABAAEAAQABAArAFAAUABQAFAAUABQAFAAUAArACsAUABQACsAKwBQAFAAUABQAFAAUABQAFAAUABQAFAAUABQAAQABAAEAAQABAArACsABAAEACsAKwAEAAQABAArACsAUAArACsAKwArACsAKwAEACsAKwArACsAKwBQAFAAUABQAFAABAAEACsAKwAEAAQABAAEAAQABAAEACsAKwArAAQABAAEAAQABAArACsAKwArACsAKwArACsAKwArACsABAAEAAQABAAEAAQABABQAFAAUABQAA0ADQANAA0AHgBLAEsASwBLAEsASwBLAEsASwBLACsADQArAB4AKwArAAQABAAEAAQAUABQAB4AUAArACsAKwArACsAKwArACsASwBLAEsASwBLAEsASwBLAEsASwArACsAKwArACsAKwBQAFAAUABQAFAAUABQAFAAUABQAFAAUABQAFAAUAAEAAQABAAEAAQABAAEACsAKwAEAAQABAAEAAQABAAEAAQABAAOAA0ADQATABMAHgAeAB4ADQANAA0ADQANAA0ADQANAA0ADQANAA0ADQANAA0AUABQAFAAUAAEAAQAKwArAAQADQANAB4AUAArACsAKwArACsAKwArACsAKwArACsASwBLAEsASwBLAEsASwBLAEsASwArACsAKwArACsAKwAOAA4ADgAOAA4ADgAOAA4ADgAOAA4ADgAOACsAKwArACsAKwArACsAKwArACsAKwArACsAKwArACsAKwArACsASwBLAEsASwBLAEsASwBLAEsASwArACsAKwArACsAKwArACsAKwArACsAKwArACsAKwArACsAKwArACsAKwArAFwAXABcAFwAXABcAFwAXABcAFwAXABcAFwAXABcAFwAXABcAFwAXABcAFwAXABcAFwAXAArACsAKwAqACoAKgAqACoAKgAqACoAKgAqACoAKgAqACoAKgArACsAKwArAEsASwBLAEsASwBLAEsASwBLAEsAXABcAA0ADQANACoASwBLAEsASwBLAEsASwBLAEsASwBQAFAAUABQAFAAUABQAFAAUAArACsAKwArACsAKwArACsAKwArACsAKwBQAFAABAAEAAQABAAEAAQABAAEAAQABABQAFAAUABQAFAAUABQAFAAUABQAFAAUABQAFAAUABQAFAAUABQAFAAUABQAFAAUAAEAAQABAAEAAQABAAEAFAABAAEAAQABAAOAB4ADQANAA0ADQAOAB4ABAArACsAKwArACsAKwArACsAUAAEAAQABAAEAAQABAAEAAQABAAEAAQAUABQAFAAUAArACsAUABQAFAAUAAEAAQABAAEAAQABAAEAAQABAAEAAQABAAEAAQABAAEAA0ADQANACsADgAOAA4ADQANACsAKwArACsAKwArACsAKwArACsAKwArACsAKwArACsAKwArACsAKwArACsAKwArACsAKwArACsAKwBQAFAAUABQAFAAUABQAFAAUAArAFAAUABQAFAAUABQAFAAUABQAFAAUABQAFAAUABQAFAAUABQAFAAUABQAFAAUABQAFAABAAEAAQABAAEAAQABAAEACsABAAEAAQABAAEAAQABAAEAFAADQANAA0ADQANACsAKwArACsAKwArACsAKwArACsASwBLAEsASwBLAEsASwBLAEsASwBQAFAAUABQAFAAUABQAFAAUABQAFAAUABQAFAAUAArACsAKwAOABMAUABQAFAAUABQAFAAUABQAFAAUABQAFAAUABQAFAAUABQAFAAKwArAAQABAAEAAQABAAEAAQABAAEAAQABAAEAAQABAArAAQABAAEAAQABAAEAAQABAAEAAQABAAEAAQABAArACsAKwArACsAKwArACsAKwBQAFAAUABQAFAAUABQACsAUABQACsAUABQAFAAUABQAFAAUABQAFAAUABQAFAAUABQAFAAUABQAFAAUABQAFAAUAAEAAQABAAEAAQABAArACsAKwAEACsABAAEACsABAAEAAQABAAEAAQABABQAAQAKwArACsAKwArACsAKwArAEsASwBLAEsASwBLAEsASwBLAEsAKwArACsAKwArACsAUABQAFAAUABQAFAAUABQAFAAUABQAFAAUABQAFAAUABQAFAAUABQAFAAUABQAFAAUABQACsAKwArACsAKwArAFAAUABQAFAAUABQAFAAUABQAFAAUABQAFAAUABQACsADQANAA0ADQANACsAKwArACsAKwArACsAKwArACsAKwBQAFAAUABQACsAKwArACsAKwArACsAKwArACsAKwArACsAKwArACsAKwArACsAKwArACsAKwArACsAKwArACsAUABQAFAAUABQAFAAUABQAFAAUABQAFAAUABQAFAAUABQAFAAUABQAFAAUABQAFAASABIAEgAQwBDAEMAUABQAFAAUABDAFAAUABQAEgAQwBIAEMAUABQAFAAUABQAFAAUABQAFAAUABQAFAAUABQAFAAUABQAFAAUABQAFAAUABQAFAAUABQAFAASABDAEMAUABQAFAAUABQAFAAUABQAFAAUABQAFAAUABQAFAAKwArACsAKwArACsAKwArACsAKwArACsAKwArACsAKwArAFAAUABQAFAAUABQAFAAUABQAFAAUABQAFAAUABIAEMAUABQAFAAUABQAFAAUABQAFAAUABQAFAAUABQAFAAUABQAFAAUAArACsAKwArACsAKwArACsAKwArACsAKwArACsAKwArACsAKwArACsAKwArACsAKwArAEsASwBLAEsASwBLAEsASwBLAEsAKwArACsAKwANAA0AKwArACsAKwArACsAKwArACsAKwArACsAKwArACsAKwBQAFAAUABQAFAAUABQAFAAUABQAFAAUABQAFAAKwArAAQABAAEAAQABAANACsAKwArACsAKwArACsAKwArACsAUABQAFAAUABQAFAAUABQAFAAUABQAFAAUABQAFAAUAAEAAQABAAEAAQABAAEAA0ADQANAB4AHgAeAB4AHgAeAFAAUABQAFAADQAeACsAKwArACsAKwArACsAKwArACsASwBLAEsASwBLAEsASwBLAEsASwArAFAAUABQAFAAUABQAFAAKwBQAFAAUABQAFAAUABQAFAAUABQAFAAUABQAFAAUABQAFAAUABQAFAAUAArACsAKwArACsAUABQAFAAUABQAFAAUABQAFAAUABQAFAAUABQAFAAUABQAFAAUAArACsAKwArACsAKwArACsAKwArACsAKwArACsAKwArAFAAUABQAFAAUAArACsAKwArACsAKwArACsAKwArACsAUAAEAAQABAAEAAQABAAEAAQABAAEAAQABAAEAAQABAAEAAQABAAEAAQABAAEAAQABAAEAAQABAAEAAQABAAEAAQABAAEACsAKwArACsAKwArACsAKwArACsAKwArACsAKwArACsABAAEAAQABABQAFAAUABQAFAAUABQAFAAUABQAFAAUABQAEcARwArACsAKwArACsAKwArACsAKwArACsAKwArACsAKwArACsAKwArACsAKwArACsAKwArACsAKwArACsAKwBXAFcAVwBXAFcAVwBXAFcAVwBXAFcAVwBXACsAKwArACsAKwArACsAKwArACsAKwArACsAKwArACsAKwArACsAVwBXAFcAVwBXAFcAVwBXAFcAVwBXAFcAVwBXAFcAVwBXAFcAVwArACsAKwArACsAKwArACsAKwArACsAKwArAFcAVwBXAFcAVwBXAFcAVwBXAFcAVwBXAFcAVwBXAFcAVwBXAFcAVwBXAFcAVwBXAFcAVwBXAFcAKwArACsAKwBQAFAAUABQAFAAUABQAFAAUABQAFAAKwArACsAKwArAFAAUABQAFAAUABQAFAAUABQAFAAUABQAFAAKwArACsAKwArACsAKwBQAFAAUABQAFAAUABQAFAAUABQACsAKwAeAAQABAANAAQABAAEAAQAKwArACsAKwArACsAKwArACsAKwArACsAKwArACsAKwArACsAKwArACsAKwArACsAKwArACsAKwAeAB4AHgAeAB4AHgAeAB4AHgAeAB4AHgAeAB4AHgAeAB4AHgAeAB4AHgAeACsAKwArACsAKwArACsAKwArACsAHgAeAB4AHgAeAB4AHgArACsAHgAeAB4AHgAeAB4AHgAeAB4AHgAeAB4AHgAeAB4AHgAeAB4AHgAeAB4AHgAeAB4ABAAEAAQABAAEAB4AHgAeAAQABAAEAAQABAAEAAQABAAEAAQABAAEAAQABAAEAAQABAAEAAQABAAEAAQAHgAeAAQABAAEAAQABAAEAAQAHgAeAB4AHgAeAB4AHgAeAB4AHgAeAB4AHgAeAB4AHgAeAB4AHgAeAB4AHgAEAAQABAAEAB4AHgAeAB4AHgAeAB4AHgAeAB4AHgAeAB4AHgAeAB4AHgAeAB4AKwArACsAKwArACsAKwArACsAKwArACsAKwArACsAKwArACsAKwArACsAKwArAB4AHgAEAAQABAAeACsAKwArACsAKwArACsAKwArACsAKwArACsAKwArACsAKwArACsAKwArACsAKwArACsAKwAeAB4AHgAeAB4AHgAeAB4AHgAeAB4AHgAeAB4AHgAeAB4AHgAeAB4AHgAeAB4AKwArACsAKwArACsAKwArACsAUABQAFAAUABQAFAAUABQAFAAUABQAFAAUABQAFAAUABQAFAAKwArACsAKwArACsAKwArACsAKwArACsAKwArAB4AHgAeAB4AHgAeAB4AHgAeAB4AHgAeAB4AHgAeAB4AHgAeAB4AHgAeACsAHgAeAB4AHgAeAB4AHgAeAB4AHgAeAB4AHgAeAB4AHgAeAB4AHgAeAB4AHgAeAB4AHgAeAB4AHgAeAB4AHgArAFAAUAArACsAUAArACsAUABQACsAKwBQAFAAUABQACsAHgAeAB4AHgAeAB4AHgAeAB4AHgAeAB4AKwBQACsAUABQAFAAUABQAFAAUAArAB4AHgAeAB4AHgAeAB4AHgAeAB4AHgAeAB4AHgAeAB4AHgAeAB4AHgAeAB4AHgAeAB4AHgAeAB4AHgArAFAAUABQAFAAKwArAFAAUABQAFAAUABQAFAAUAArAFAAUABQAFAAUABQAFAAKwAeAB4AUABQAFAAUABQACsAUAArACsAKwBQAFAAUABQAFAAUABQACsAHgAeAB4AHgAeAB4AHgAeAB4AHgAeAB4AHgAeAB4AHgArACsAUABQAFAAUABQAFAAUABQAFAAUABQAFAAUABQAFAAUABQAFAAUABQAFAAUABQAFAAUAAeAFAAUABQAFAAUABQAFAAUABQAFAAUABQAFAAUABQAFAAUABQAFAAUABQAFAAUABQAFAAHgAeAB4AHgAeAB4AHgAeAB4AHgAeAB4AHgAeAB4AHgAeAB4AHgAeAB4AHgAeAFAAUABQAFAAUABQAFAAUABQAFAAUAAeAB4AHgAeAB4AHgAeAB4AHgAeAB4AHgAeAB4AHgAeAB4AHgAeAB4AHgAeAB4AHgAeAFAAUABQAFAAUABQAFAAUABQAFAAUABQAFAAUABQAFAAUABQAFAAUABQAFAAUABQAFAAHgAeAB4AHgAeAB4AHgAeAB4AKwArAEsASwBLAEsASwBLAEsASwBLAEsASwBLAEsASwBLAEsASwBLAEsASwBLAEsASwBLAEsASwBLAEsASwBLAEsASwBLAEsABAAEAAQABAAEAAQABAAEAAQABAAEAAQABAAEAAQABAAEAAQABAAEAAQABAAEAB4AHgAeAB4ABAAEAAQABAAEAAQABAAEAAQABAAEAAQABAAEAB4AHgAeAB4AHgAeAB4AHgAEAB4AHgAeAB4AHgAeAB4AHgAeAB4ABAAeAB4ADQANAA0ADQAeACsAKwArACsAKwArACsAKwArACsAKwArACsAKwArAAQABAAEAAQABAArAAQABAAEAAQABAAEAAQABAAEAAQABAAEAAQABAAEACsAKwArACsAKwArACsAKwArACsAKwArACsAKwArACsABAAEAAQABAAEAAQABAArAAQABAAEAAQABAAEAAQABAAEAAQABAAEAAQABAAEAAQABAArACsABAAEAAQABAAEAAQABAArAAQABAArAAQABAAEAAQABAArACsAKwArACsAKwArACsAKwArACsAKwArACsAKwArACsAKwArACsAKwBQAFAAUABQAFAAKwArAFAAUABQAFAAUABQAFAAUABQAAQABAAEAAQABAAEAAQAKwArACsAKwArACsAKwArACsAHgAeAB4AHgAEAAQABAAEAAQABAAEACsAKwArACsAKwBLAEsASwBLAEsASwBLAEsASwBLACsAKwArACsAFgAWAFAAUABQAFAAKwBQAFAAUABQAFAAUABQAFAAUABQAFAAUABQAFAAUABQAFAAUABQAFAAUABQAFAAUABQAFAAUAArAFAAUAArAFAAKwArAFAAKwBQAFAAUABQAFAAUABQAFAAUABQACsAUABQAFAAUAArAFAAKwBQACsAKwArACsAKwArAFAAKwArACsAKwBQACsAUAArAFAAKwBQAFAAUAArAFAAUAArAFAAKwArAFAAKwBQACsAUAArAFAAKwBQACsAUABQACsAUAArACsAUABQAFAAUAArAFAAUABQAFAAUABQAFAAKwBQAFAAUABQACsAUABQAFAAUAArAFAAKwBQAFAAUABQAFAAUABQAFAAUABQACsAUABQAFAAUABQAFAAUABQAFAAUABQAFAAUABQAFAAUABQACsAKwArACsAKwBQAFAAUAArAFAAUABQAFAAUAArAFAAUABQAFAAUABQAFAAUABQAFAAUABQAFAAUABQAFAAUAArACsAKwArACsAKwArACsAKwArACsAKwArACsAKwArAB4AHgArACsAKwArACsAKwArACsAKwArACsAKwArACsATwBPAE8ATwBPAE8ATwBPAE8ATwBPAE8ATwAlACUAJQAdAB0AHQAdAB0AHQAdAB0AHQAdAB0AHQAdAB0AHQAdAB0AHQAeACUAHQAdAB0AHQAdAB0AHQAdAB0AHQAdAB0AHQAdAB0AHQAdAB0AHgAeACUAJQAlACUAHQAdAB0AHQAdAB0AHQAdAB0AHQAdAB0AHQAdAB0AHQAdACUAJQAlACUAJQAlACUAJQAlACUAJQAlACUAJQAlACUAJQAlACUAJQAlACkAKQApACkAKQApACkAKQApACkAKQApACkAKQApACkAKQApACkAKQApACkAKQApACkAKQAlACUAJQAlACUAIAAlACUAJQAlACUAJQAlACUAJQAlACUAJQAlACUAJQAlACUAJQAlACUAJQAlAB4AHgAlACUAJQAlACUAJQAlACUAJQAlACUAJQAlACUAJQAlACUAJQAlACUAJQAlACUAHgAeACUAJQAlACUAJQAeACUAJQAlACUAJQAgACAAIAAlACUAIAAlACUAIAAgACAAJQAlACUAJQAlACUAJQAlACUAJQAlACUAJQAlACUAJQAlACUAJQAlACUAJQAlACUAJQAlACUAJQAlACUAIQAhACEAIQAhACUAJQAgACAAJQAlACAAIAAgACAAIAAgACAAIAAgACAAIAAlACUAJQAlACUAJQAlACUAJQAlACUAJQAlACUAJQAlACUAIAAgACAAIAAlACUAJQAlACAAJQAgACAAIAAgACAAIAAgACAAIAAlACUAJQAgACUAJQAlACUAIAAgACAAJQAgACAAIAAlACUAJQAlACUAJQAlACUAJQAlACUAJQAlACUAJQAlACUAJQAlACUAJQAlACUAJQAeACUAHgAlAB4AJQAlACUAJQAlACAAJQAlACUAJQAeACUAHgAeACUAJQAlACUAJQAlACUAJQAlACUAJQAlACUAHgAeAB4AHgAeAB4AHgAlACUAJQAlACUAJQAlACUAJQAlACUAJQAlACUAJQAlAB4AHgAeAB4AHgAeAB4AHgAeAB4AJQAlACUAJQAlACUAJQAlACUAJQAlACUAJQAeAB4AHgAeAB4AHgAeAB4AHgAeAB4AHgAeAB4AHgAeACUAJQAlACUAJQAlACUAJQAlACUAJQAlACUAJQAlACUAJQAlACUAJQAlACUAIAAgACUAJQAlACUAIAAlACUAJQAlACUAJQAlACUAJQAlACUAJQAlACUAJQAlACUAIAAlACUAJQAlACAAIAAlACUAJQAlACUAJQAlACUAJQAlACUAJQAlACUAJQAlACUAJQAlACUAJQAeAB4AHgAeAB4AHgAeAB4AJQAlACUAJQAlACUAJQAlACUAJQAlACUAJQAlACUAJQAlACUAJQAlAB4AHgAeAB4AHgAeACUAJQAlACUAJQAlACUAIAAgACAAJQAlACUAIAAgACAAIAAgAB4AHgAeAB4AHgAeAB4AHgAeAB4AHgAeAB4AHgAeAB4AHgAeAB4AHgAeAB4AFwAXABcAFQAVABUAHgAeAB4AHgAlACUAJQAgACUAJQAlACUAJQAlACUAJQAlACUAJQAlACUAJQAlACUAIAAgACAAJQAlACUAJQAlACUAJQAlACUAIAAlACUAJQAlACUAJQAlACUAJQAlACUAIAAlACUAJQAlACUAJQAlACUAJQAlACUAJQAlACUAJQAlACUAJQAlAB4AHgAeAB4AHgAeAB4AHgAeAB4AHgAeAB4AHgAeAB4AHgAeAB4AHgAlACUAJQAlACUAJQAlACUAJQAlACUAJQAeAB4AHgAeAB4AHgAeAB4AHgAeAB4AHgAlACUAJQAlAB4AHgAeAB4AHgAeAB4AHgAeAB4AHgAeAB4AHgAeAB4AJQAlACUAJQAlACUAJQAlAB4AHgAeAB4AHgAeAB4AHgAeAB4AJQAlACUAJQAlACUAHgAeAB4AHgAeAB4AHgAeACUAJQAlACUAJQAlACUAJQAeAB4AHgAeAB4AHgAeAB4AHgAeAB4AHgAeAB4AHgAeACUAJQAlACUAJQAlACUAJQAlACUAJQAlACAAIAAgACAAIAAlACAAIAAlACUAJQAlACUAJQAgACUAJQAlACUAJQAlACUAJQAlACAAIAAgACAAIAAgACAAIAAgACAAJQAlACUAIAAgACUAJQAlACUAJQAlACUAJQAlACUAJQAlACUAJQAlACUAJQAlACAAIAAgACAAIAAgACAAIAAgACAAIAAgACAAJQAlACUAJQAlACUAJQAlACUAJQAlACUAJQAlACUAJQAlACUAJQAlACUAJQAlACUAJQAlACUAJQAlACUAJQAlACsAKwBXAFcAVwBXAFcAVwBXAFcAVwBXAFcAVwBXAFcAVwBXAFcAVwBXAFcAVwBXAFcAJQAlACUAJQAlACUAJQAlACUAVwBXAFcAVwBXAFcAVwBXAFcAVwBXAFcAVwBXAFcAVwBXAFcAVwBXAFcAJQAlACUAJQAlACUAJQAlACUAJQAlAFcAVwBXAFcAVwBXAFcAVwBXAFcAVwBXAFcAVwBXAFcAVwBXAFcAVwBXAFcAVwBXAFcAVwBXAFcAVwBXACUAJQAlACUAJQAlACUAJQAlACUAJQAlACUAJQBXAFcAVwBXAFcAVwBXAFcAVwBXAFcAVwBXAFcAVwBXAFcAJQAlACUAJQAlACUAJQAlACUAJQAlACUAJQAlACUAJQAlACUAJQAlACUAJQAlACUAJQAlACUAJQAlACUAJQArAAQAKwArACsAKwArACsAKwArACsAKwArACsAKwArACsAKwArACsAKwArACsAKwArACsAKwArACsAKwArACsABAAEAAQABAAEAAQABAAEAAQABAAEAAQABAAEAAQABAArACsAKwArACsAKwArACsAKwArACsAKwArACsAKwArACsAKwArACsA';

    /* @flow */
    var LETTER_NUMBER_MODIFIER = 50;
    // Non-tailorable Line Breaking Classes
    var BK = 1; //  Cause a line break (after)
    var CR = 2; //  Cause a line break (after), except between CR and LF
    var LF = 3; //  Cause a line break (after)
    var CM = 4; //  Prohibit a line break between the character and the preceding character
    var NL = 5; //  Cause a line break (after)
    var WJ = 7; //  Prohibit line breaks before and after
    var ZW = 8; //  Provide a break opportunity
    var GL = 9; //  Prohibit line breaks before and after
    var SP = 10; // Enable indirect line breaks
    var ZWJ = 11; // Prohibit line breaks within joiner sequences
    // Break Opportunities
    var B2 = 12; //  Provide a line break opportunity before and after the character
    var BA = 13; //  Generally provide a line break opportunity after the character
    var BB = 14; //  Generally provide a line break opportunity before the character
    var HY = 15; //  Provide a line break opportunity after the character, except in numeric context
    var CB = 16; //   Provide a line break opportunity contingent on additional information
    // Characters Prohibiting Certain Breaks
    var CL = 17; //  Prohibit line breaks before
    var CP = 18; //  Prohibit line breaks before
    var EX = 19; //  Prohibit line breaks before
    var IN = 20; //  Allow only indirect line breaks between pairs
    var NS = 21; //  Allow only indirect line breaks before
    var OP = 22; //  Prohibit line breaks after
    var QU = 23; //  Act like they are both opening and closing
    // Numeric Context
    var IS = 24; //  Prevent breaks after any and before numeric
    var NU = 25; //  Form numeric expressions for line breaking purposes
    var PO = 26; //  Do not break following a numeric expression
    var PR = 27; //  Do not break in front of a numeric expression
    var SY = 28; //  Prevent a break before; and allow a break after
    // Other Characters
    var AI = 29; //  Act like AL when the resolvedEAW is N; otherwise; act as ID
    var AL = 30; //  Are alphabetic characters or symbols that are used with alphabetic characters
    var CJ = 31; //  Treat as NS or ID for strict or normal breaking.
    var EB = 32; //  Do not break from following Emoji Modifier
    var EM = 33; //  Do not break from preceding Emoji Base
    var H2 = 34; //  Form Korean syllable blocks
    var H3 = 35; //  Form Korean syllable blocks
    var HL = 36; //  Do not break around a following hyphen; otherwise act as Alphabetic
    var ID = 37; //  Break before or after; except in some numeric context
    var JL = 38; //  Form Korean syllable blocks
    var JV = 39; //  Form Korean syllable blocks
    var JT = 40; //  Form Korean syllable blocks
    var RI = 41; //  Keep pairs together. For pairs; break before and after other classes
    var SA = 42; //  Provide a line break opportunity contingent on additional, language-specific context analysis
    var XX = 43; //  Have as yet unknown line breaking behavior or unassigned code positions
    var BREAK_MANDATORY = '!';
    var BREAK_NOT_ALLOWED = '×';
    var BREAK_ALLOWED = '÷';
    var UnicodeTrie = createTrieFromBase64(base64);
    var ALPHABETICS = [AL, HL];
    var HARD_LINE_BREAKS = [BK, CR, LF, NL];
    var SPACE = [SP, ZW];
    var PREFIX_POSTFIX = [PR, PO];
    var LINE_BREAKS = HARD_LINE_BREAKS.concat(SPACE);
    var KOREAN_SYLLABLE_BLOCK = [JL, JV, JT, H2, H3];
    var HYPHEN = [HY, BA];
    var codePointsToCharacterClasses = function (codePoints, lineBreak) {
        if (lineBreak === void 0) { lineBreak = 'strict'; }
        var types = [];
        var indicies = [];
        var categories = [];
        codePoints.forEach(function (codePoint, index) {
            var classType = UnicodeTrie.get(codePoint);
            if (classType > LETTER_NUMBER_MODIFIER) {
                categories.push(true);
                classType -= LETTER_NUMBER_MODIFIER;
            }
            else {
                categories.push(false);
            }
            if (['normal', 'auto', 'loose'].indexOf(lineBreak) !== -1) {
                // U+2010, – U+2013, 〜 U+301C, ゠ U+30A0
                if ([0x2010, 0x2013, 0x301c, 0x30a0].indexOf(codePoint) !== -1) {
                    indicies.push(index);
                    return types.push(CB);
                }
            }
            if (classType === CM || classType === ZWJ) {
                // LB10 Treat any remaining combining mark or ZWJ as AL.
                if (index === 0) {
                    indicies.push(index);
                    return types.push(AL);
                }
                // LB9 Do not break a combining character sequence; treat it as if it has the line breaking class of
                // the base character in all of the following rules. Treat ZWJ as if it were CM.
                var prev = types[index - 1];
                if (LINE_BREAKS.indexOf(prev) === -1) {
                    indicies.push(indicies[index - 1]);
                    return types.push(prev);
                }
                indicies.push(index);
                return types.push(AL);
            }
            indicies.push(index);
            if (classType === CJ) {
                return types.push(lineBreak === 'strict' ? NS : ID);
            }
            if (classType === SA) {
                return types.push(AL);
            }
            if (classType === AI) {
                return types.push(AL);
            }
            // For supplementary characters, a useful default is to treat characters in the range 10000..1FFFD as AL
            // and characters in the ranges 20000..2FFFD and 30000..3FFFD as ID, until the implementation can be revised
            // to take into account the actual line breaking properties for these characters.
            if (classType === XX) {
                if ((codePoint >= 0x20000 && codePoint <= 0x2fffd) || (codePoint >= 0x30000 && codePoint <= 0x3fffd)) {
                    return types.push(ID);
                }
                else {
                    return types.push(AL);
                }
            }
            types.push(classType);
        });
        return [indicies, types, categories];
    };
    var isAdjacentWithSpaceIgnored = function (a, b, currentIndex, classTypes) {
        var current = classTypes[currentIndex];
        if (Array.isArray(a) ? a.indexOf(current) !== -1 : a === current) {
            var i = currentIndex;
            while (i <= classTypes.length) {
                i++;
                var next = classTypes[i];
                if (next === b) {
                    return true;
                }
                if (next !== SP) {
                    break;
                }
            }
        }
        if (current === SP) {
            var i = currentIndex;
            while (i > 0) {
                i--;
                var prev = classTypes[i];
                if (Array.isArray(a) ? a.indexOf(prev) !== -1 : a === prev) {
                    var n = currentIndex;
                    while (n <= classTypes.length) {
                        n++;
                        var next = classTypes[n];
                        if (next === b) {
                            return true;
                        }
                        if (next !== SP) {
                            break;
                        }
                    }
                }
                if (prev !== SP) {
                    break;
                }
            }
        }
        return false;
    };
    var previousNonSpaceClassType = function (currentIndex, classTypes) {
        var i = currentIndex;
        while (i >= 0) {
            var type = classTypes[i];
            if (type === SP) {
                i--;
            }
            else {
                return type;
            }
        }
        return 0;
    };
    var _lineBreakAtIndex = function (codePoints, classTypes, indicies, index, forbiddenBreaks) {
        if (indicies[index] === 0) {
            return BREAK_NOT_ALLOWED;
        }
        var currentIndex = index - 1;
        if (Array.isArray(forbiddenBreaks) && forbiddenBreaks[currentIndex] === true) {
            return BREAK_NOT_ALLOWED;
        }
        var beforeIndex = currentIndex - 1;
        var afterIndex = currentIndex + 1;
        var current = classTypes[currentIndex];
        // LB4 Always break after hard line breaks.
        // LB5 Treat CR followed by LF, as well as CR, LF, and NL as hard line breaks.
        var before = beforeIndex >= 0 ? classTypes[beforeIndex] : 0;
        var next = classTypes[afterIndex];
        if (current === CR && next === LF) {
            return BREAK_NOT_ALLOWED;
        }
        if (HARD_LINE_BREAKS.indexOf(current) !== -1) {
            return BREAK_MANDATORY;
        }
        // LB6 Do not break before hard line breaks.
        if (HARD_LINE_BREAKS.indexOf(next) !== -1) {
            return BREAK_NOT_ALLOWED;
        }
        // LB7 Do not break before spaces or zero width space.
        if (SPACE.indexOf(next) !== -1) {
            return BREAK_NOT_ALLOWED;
        }
        // LB8 Break before any character following a zero-width space, even if one or more spaces intervene.
        if (previousNonSpaceClassType(currentIndex, classTypes) === ZW) {
            return BREAK_ALLOWED;
        }
        // LB8a Do not break between a zero width joiner and an ideograph, emoji base or emoji modifier.
        if (UnicodeTrie.get(codePoints[currentIndex]) === ZWJ && (next === ID || next === EB || next === EM)) {
            return BREAK_NOT_ALLOWED;
        }
        // LB11 Do not break before or after Word joiner and related characters.
        if (current === WJ || next === WJ) {
            return BREAK_NOT_ALLOWED;
        }
        // LB12 Do not break after NBSP and related characters.
        if (current === GL) {
            return BREAK_NOT_ALLOWED;
        }
        // LB12a Do not break before NBSP and related characters, except after spaces and hyphens.
        if ([SP, BA, HY].indexOf(current) === -1 && next === GL) {
            return BREAK_NOT_ALLOWED;
        }
        // LB13 Do not break before ‘]’ or ‘!’ or ‘;’ or ‘/’, even after spaces.
        if ([CL, CP, EX, IS, SY].indexOf(next) !== -1) {
            return BREAK_NOT_ALLOWED;
        }
        // LB14 Do not break after ‘[’, even after spaces.
        if (previousNonSpaceClassType(currentIndex, classTypes) === OP) {
            return BREAK_NOT_ALLOWED;
        }
        // LB15 Do not break within ‘”[’, even with intervening spaces.
        if (isAdjacentWithSpaceIgnored(QU, OP, currentIndex, classTypes)) {
            return BREAK_NOT_ALLOWED;
        }
        // LB16 Do not break between closing punctuation and a nonstarter (lb=NS), even with intervening spaces.
        if (isAdjacentWithSpaceIgnored([CL, CP], NS, currentIndex, classTypes)) {
            return BREAK_NOT_ALLOWED;
        }
        // LB17 Do not break within ‘——’, even with intervening spaces.
        if (isAdjacentWithSpaceIgnored(B2, B2, currentIndex, classTypes)) {
            return BREAK_NOT_ALLOWED;
        }
        // LB18 Break after spaces.
        if (current === SP) {
            return BREAK_ALLOWED;
        }
        // LB19 Do not break before or after quotation marks, such as ‘ ” ’.
        if (current === QU || next === QU) {
            return BREAK_NOT_ALLOWED;
        }
        // LB20 Break before and after unresolved CB.
        if (next === CB || current === CB) {
            return BREAK_ALLOWED;
        }
        // LB21 Do not break before hyphen-minus, other hyphens, fixed-width spaces, small kana, and other non-starters, or after acute accents.
        if ([BA, HY, NS].indexOf(next) !== -1 || current === BB) {
            return BREAK_NOT_ALLOWED;
        }
        // LB21a Don't break after Hebrew + Hyphen.
        if (before === HL && HYPHEN.indexOf(current) !== -1) {
            return BREAK_NOT_ALLOWED;
        }
        // LB21b Don’t break between Solidus and Hebrew letters.
        if (current === SY && next === HL) {
            return BREAK_NOT_ALLOWED;
        }
        // LB22 Do not break between two ellipses, or between letters, numbers or exclamations and ellipsis.
        if (next === IN && ALPHABETICS.concat(IN, EX, NU, ID, EB, EM).indexOf(current) !== -1) {
            return BREAK_NOT_ALLOWED;
        }
        // LB23 Do not break between digits and letters.
        if ((ALPHABETICS.indexOf(next) !== -1 && current === NU) || (ALPHABETICS.indexOf(current) !== -1 && next === NU)) {
            return BREAK_NOT_ALLOWED;
        }
        // LB23a Do not break between numeric prefixes and ideographs, or between ideographs and numeric postfixes.
        if ((current === PR && [ID, EB, EM].indexOf(next) !== -1) ||
            ([ID, EB, EM].indexOf(current) !== -1 && next === PO)) {
            return BREAK_NOT_ALLOWED;
        }
        // LB24 Do not break between numeric prefix/postfix and letters, or between letters and prefix/postfix.
        if ((ALPHABETICS.indexOf(current) !== -1 && PREFIX_POSTFIX.indexOf(next) !== -1) ||
            (PREFIX_POSTFIX.indexOf(current) !== -1 && ALPHABETICS.indexOf(next) !== -1)) {
            return BREAK_NOT_ALLOWED;
        }
        // LB25 Do not break between the following pairs of classes relevant to numbers:
        if (
        // (PR | PO) × ( OP | HY )? NU
        ([PR, PO].indexOf(current) !== -1 &&
            (next === NU || ([OP, HY].indexOf(next) !== -1 && classTypes[afterIndex + 1] === NU))) ||
            // ( OP | HY ) × NU
            ([OP, HY].indexOf(current) !== -1 && next === NU) ||
            // NU ×	(NU | SY | IS)
            (current === NU && [NU, SY, IS].indexOf(next) !== -1)) {
            return BREAK_NOT_ALLOWED;
        }
        // NU (NU | SY | IS)* × (NU | SY | IS | CL | CP)
        if ([NU, SY, IS, CL, CP].indexOf(next) !== -1) {
            var prevIndex = currentIndex;
            while (prevIndex >= 0) {
                var type = classTypes[prevIndex];
                if (type === NU) {
                    return BREAK_NOT_ALLOWED;
                }
                else if ([SY, IS].indexOf(type) !== -1) {
                    prevIndex--;
                }
                else {
                    break;
                }
            }
        }
        // NU (NU | SY | IS)* (CL | CP)? × (PO | PR))
        if ([PR, PO].indexOf(next) !== -1) {
            var prevIndex = [CL, CP].indexOf(current) !== -1 ? beforeIndex : currentIndex;
            while (prevIndex >= 0) {
                var type = classTypes[prevIndex];
                if (type === NU) {
                    return BREAK_NOT_ALLOWED;
                }
                else if ([SY, IS].indexOf(type) !== -1) {
                    prevIndex--;
                }
                else {
                    break;
                }
            }
        }
        // LB26 Do not break a Korean syllable.
        if ((JL === current && [JL, JV, H2, H3].indexOf(next) !== -1) ||
            ([JV, H2].indexOf(current) !== -1 && [JV, JT].indexOf(next) !== -1) ||
            ([JT, H3].indexOf(current) !== -1 && next === JT)) {
            return BREAK_NOT_ALLOWED;
        }
        // LB27 Treat a Korean Syllable Block the same as ID.
        if ((KOREAN_SYLLABLE_BLOCK.indexOf(current) !== -1 && [IN, PO].indexOf(next) !== -1) ||
            (KOREAN_SYLLABLE_BLOCK.indexOf(next) !== -1 && current === PR)) {
            return BREAK_NOT_ALLOWED;
        }
        // LB28 Do not break between alphabetics (“at”).
        if (ALPHABETICS.indexOf(current) !== -1 && ALPHABETICS.indexOf(next) !== -1) {
            return BREAK_NOT_ALLOWED;
        }
        // LB29 Do not break between numeric punctuation and alphabetics (“e.g.”).
        if (current === IS && ALPHABETICS.indexOf(next) !== -1) {
            return BREAK_NOT_ALLOWED;
        }
        // LB30 Do not break between letters, numbers, or ordinary symbols and opening or closing parentheses.
        if ((ALPHABETICS.concat(NU).indexOf(current) !== -1 && next === OP) ||
            (ALPHABETICS.concat(NU).indexOf(next) !== -1 && current === CP)) {
            return BREAK_NOT_ALLOWED;
        }
        // LB30a Break between two regional indicator symbols if and only if there are an even number of regional
        // indicators preceding the position of the break.
        if (current === RI && next === RI) {
            var i = indicies[currentIndex];
            var count = 1;
            while (i > 0) {
                i--;
                if (classTypes[i] === RI) {
                    count++;
                }
                else {
                    break;
                }
            }
            if (count % 2 !== 0) {
                return BREAK_NOT_ALLOWED;
            }
        }
        // LB30b Do not break between an emoji base and an emoji modifier.
        if (current === EB && next === EM) {
            return BREAK_NOT_ALLOWED;
        }
        return BREAK_ALLOWED;
    };
    var cssFormattedClasses = function (codePoints, options) {
        if (!options) {
            options = { lineBreak: 'normal', wordBreak: 'normal' };
        }
        var _a = codePointsToCharacterClasses(codePoints, options.lineBreak), indicies = _a[0], classTypes = _a[1], isLetterNumber = _a[2];
        if (options.wordBreak === 'break-all' || options.wordBreak === 'break-word') {
            classTypes = classTypes.map(function (type) { return ([NU, AL, SA].indexOf(type) !== -1 ? ID : type); });
        }
        var forbiddenBreakpoints = options.wordBreak === 'keep-all'
            ? isLetterNumber.map(function (letterNumber, i) {
                return letterNumber && codePoints[i] >= 0x4e00 && codePoints[i] <= 0x9fff;
            })
            : undefined;
        return [indicies, classTypes, forbiddenBreakpoints];
    };
    var Break = /** @class */ (function () {
        function Break(codePoints, lineBreak, start, end) {
            this.codePoints = codePoints;
            this.required = lineBreak === BREAK_MANDATORY;
            this.start = start;
            this.end = end;
        }
        Break.prototype.slice = function () {
            return fromCodePoint.apply(void 0, this.codePoints.slice(this.start, this.end));
        };
        return Break;
    }());
    var LineBreaker = function (str, options) {
        var codePoints = toCodePoints(str);
        var _a = cssFormattedClasses(codePoints, options), indicies = _a[0], classTypes = _a[1], forbiddenBreakpoints = _a[2];
        var length = codePoints.length;
        var lastEnd = 0;
        var nextIndex = 0;
        return {
            next: function () {
                if (nextIndex >= length) {
                    return { done: true, value: null };
                }
                var lineBreak = BREAK_NOT_ALLOWED;
                while (nextIndex < length &&
                    (lineBreak = _lineBreakAtIndex(codePoints, classTypes, indicies, ++nextIndex, forbiddenBreakpoints)) ===
                        BREAK_NOT_ALLOWED) { }
                if (lineBreak !== BREAK_NOT_ALLOWED || nextIndex === length) {
                    var value = new Break(codePoints, lineBreak, lastEnd, nextIndex);
                    lastEnd = nextIndex;
                    return { value: value, done: false };
                }
                return { done: true, value: null };
            },
        };
    };

    // https://www.w3.org/TR/css-syntax-3
    var TokenType;
    (function (TokenType) {
        TokenType[TokenType["STRING_TOKEN"] = 0] = "STRING_TOKEN";
        TokenType[TokenType["BAD_STRING_TOKEN"] = 1] = "BAD_STRING_TOKEN";
        TokenType[TokenType["LEFT_PARENTHESIS_TOKEN"] = 2] = "LEFT_PARENTHESIS_TOKEN";
        TokenType[TokenType["RIGHT_PARENTHESIS_TOKEN"] = 3] = "RIGHT_PARENTHESIS_TOKEN";
        TokenType[TokenType["COMMA_TOKEN"] = 4] = "COMMA_TOKEN";
        TokenType[TokenType["HASH_TOKEN"] = 5] = "HASH_TOKEN";
        TokenType[TokenType["DELIM_TOKEN"] = 6] = "DELIM_TOKEN";
        TokenType[TokenType["AT_KEYWORD_TOKEN"] = 7] = "AT_KEYWORD_TOKEN";
        TokenType[TokenType["PREFIX_MATCH_TOKEN"] = 8] = "PREFIX_MATCH_TOKEN";
        TokenType[TokenType["DASH_MATCH_TOKEN"] = 9] = "DASH_MATCH_TOKEN";
        TokenType[TokenType["INCLUDE_MATCH_TOKEN"] = 10] = "INCLUDE_MATCH_TOKEN";
        TokenType[TokenType["LEFT_CURLY_BRACKET_TOKEN"] = 11] = "LEFT_CURLY_BRACKET_TOKEN";
        TokenType[TokenType["RIGHT_CURLY_BRACKET_TOKEN"] = 12] = "RIGHT_CURLY_BRACKET_TOKEN";
        TokenType[TokenType["SUFFIX_MATCH_TOKEN"] = 13] = "SUFFIX_MATCH_TOKEN";
        TokenType[TokenType["SUBSTRING_MATCH_TOKEN"] = 14] = "SUBSTRING_MATCH_TOKEN";
        TokenType[TokenType["DIMENSION_TOKEN"] = 15] = "DIMENSION_TOKEN";
        TokenType[TokenType["PERCENTAGE_TOKEN"] = 16] = "PERCENTAGE_TOKEN";
        TokenType[TokenType["NUMBER_TOKEN"] = 17] = "NUMBER_TOKEN";
        TokenType[TokenType["FUNCTION"] = 18] = "FUNCTION";
        TokenType[TokenType["FUNCTION_TOKEN"] = 19] = "FUNCTION_TOKEN";
        TokenType[TokenType["IDENT_TOKEN"] = 20] = "IDENT_TOKEN";
        TokenType[TokenType["COLUMN_TOKEN"] = 21] = "COLUMN_TOKEN";
        TokenType[TokenType["URL_TOKEN"] = 22] = "URL_TOKEN";
        TokenType[TokenType["BAD_URL_TOKEN"] = 23] = "BAD_URL_TOKEN";
        TokenType[TokenType["CDC_TOKEN"] = 24] = "CDC_TOKEN";
        TokenType[TokenType["CDO_TOKEN"] = 25] = "CDO_TOKEN";
        TokenType[TokenType["COLON_TOKEN"] = 26] = "COLON_TOKEN";
        TokenType[TokenType["SEMICOLON_TOKEN"] = 27] = "SEMICOLON_TOKEN";
        TokenType[TokenType["LEFT_SQUARE_BRACKET_TOKEN"] = 28] = "LEFT_SQUARE_BRACKET_TOKEN";
        TokenType[TokenType["RIGHT_SQUARE_BRACKET_TOKEN"] = 29] = "RIGHT_SQUARE_BRACKET_TOKEN";
        TokenType[TokenType["UNICODE_RANGE_TOKEN"] = 30] = "UNICODE_RANGE_TOKEN";
        TokenType[TokenType["WHITESPACE_TOKEN"] = 31] = "WHITESPACE_TOKEN";
        TokenType[TokenType["EOF_TOKEN"] = 32] = "EOF_TOKEN";
    })(TokenType || (TokenType = {}));
    var FLAG_UNRESTRICTED = 1 << 0;
    var FLAG_ID = 1 << 1;
    var FLAG_INTEGER = 1 << 2;
    var FLAG_NUMBER = 1 << 3;
    var LINE_FEED = 0x000a;
    var SOLIDUS = 0x002f;
    var REVERSE_SOLIDUS = 0x005c;
    var CHARACTER_TABULATION = 0x0009;
    var SPACE$1 = 0x0020;
    var QUOTATION_MARK = 0x0022;
    var EQUALS_SIGN = 0x003d;
    var NUMBER_SIGN = 0x0023;
    var DOLLAR_SIGN = 0x0024;
    var PERCENTAGE_SIGN = 0x0025;
    var APOSTROPHE = 0x0027;
    var LEFT_PARENTHESIS = 0x0028;
    var RIGHT_PARENTHESIS = 0x0029;
    var LOW_LINE = 0x005f;
    var HYPHEN_MINUS = 0x002d;
    var EXCLAMATION_MARK = 0x0021;
    var LESS_THAN_SIGN = 0x003c;
    var GREATER_THAN_SIGN = 0x003e;
    var COMMERCIAL_AT = 0x0040;
    var LEFT_SQUARE_BRACKET = 0x005b;
    var RIGHT_SQUARE_BRACKET = 0x005d;
    var CIRCUMFLEX_ACCENT = 0x003d;
    var LEFT_CURLY_BRACKET = 0x007b;
    var QUESTION_MARK = 0x003f;
    var RIGHT_CURLY_BRACKET = 0x007d;
    var VERTICAL_LINE = 0x007c;
    var TILDE = 0x007e;
    var CONTROL = 0x0080;
    var REPLACEMENT_CHARACTER = 0xfffd;
    var ASTERISK = 0x002a;
    var PLUS_SIGN = 0x002b;
    var COMMA = 0x002c;
    var COLON = 0x003a;
    var SEMICOLON = 0x003b;
    var FULL_STOP = 0x002e;
    var NULL = 0x0000;
    var BACKSPACE = 0x0008;
    var LINE_TABULATION = 0x000b;
    var SHIFT_OUT = 0x000e;
    var INFORMATION_SEPARATOR_ONE = 0x001f;
    var DELETE = 0x007f;
    var EOF = -1;
    var ZERO = 0x0030;
    var a = 0x0061;
    var e = 0x0065;
    var f = 0x0066;
    var u = 0x0075;
    var z = 0x007a;
    var A = 0x0041;
    var E = 0x0045;
    var F = 0x0046;
    var U = 0x0055;
    var Z = 0x005a;
    var isDigit = function (codePoint) { return codePoint >= ZERO && codePoint <= 0x0039; };
    var isSurrogateCodePoint = function (codePoint) { return codePoint >= 0xd800 && codePoint <= 0xdfff; };
    var isHex = function (codePoint) {
        return isDigit(codePoint) || (codePoint >= A && codePoint <= F) || (codePoint >= a && codePoint <= f);
    };
    var isLowerCaseLetter = function (codePoint) { return codePoint >= a && codePoint <= z; };
    var isUpperCaseLetter = function (codePoint) { return codePoint >= A && codePoint <= Z; };
    var isLetter = function (codePoint) { return isLowerCaseLetter(codePoint) || isUpperCaseLetter(codePoint); };
    var isNonASCIICodePoint = function (codePoint) { return codePoint >= CONTROL; };
    var isWhiteSpace = function (codePoint) {
        return codePoint === LINE_FEED || codePoint === CHARACTER_TABULATION || codePoint === SPACE$1;
    };
    var isNameStartCodePoint = function (codePoint) {
        return isLetter(codePoint) || isNonASCIICodePoint(codePoint) || codePoint === LOW_LINE;
    };
    var isNameCodePoint = function (codePoint) {
        return isNameStartCodePoint(codePoint) || isDigit(codePoint) || codePoint === HYPHEN_MINUS;
    };
    var isNonPrintableCodePoint = function (codePoint) {
        return ((codePoint >= NULL && codePoint <= BACKSPACE) ||
            codePoint === LINE_TABULATION ||
            (codePoint >= SHIFT_OUT && codePoint <= INFORMATION_SEPARATOR_ONE) ||
            codePoint === DELETE);
    };
    var isValidEscape = function (c1, c2) {
        if (c1 !== REVERSE_SOLIDUS) {
            return false;
        }
        return c2 !== LINE_FEED;
    };
    var isIdentifierStart = function (c1, c2, c3) {
        if (c1 === HYPHEN_MINUS) {
            return isNameStartCodePoint(c2) || isValidEscape(c2, c3);
        }
        else if (isNameStartCodePoint(c1)) {
            return true;
        }
        else if (c1 === REVERSE_SOLIDUS && isValidEscape(c1, c2)) {
            return true;
        }
        return false;
    };
    var isNumberStart = function (c1, c2, c3) {
        if (c1 === PLUS_SIGN || c1 === HYPHEN_MINUS) {
            if (isDigit(c2)) {
                return true;
            }
            return c2 === FULL_STOP && isDigit(c3);
        }
        if (c1 === FULL_STOP) {
            return isDigit(c2);
        }
        return isDigit(c1);
    };
    var stringToNumber = function (codePoints) {
        var c = 0;
        var sign = 1;
        if (codePoints[c] === PLUS_SIGN || codePoints[c] === HYPHEN_MINUS) {
            if (codePoints[c] === HYPHEN_MINUS) {
                sign = -1;
            }
            c++;
        }
        var integers = [];
        while (isDigit(codePoints[c])) {
            integers.push(codePoints[c++]);
        }
        var int = integers.length ? parseInt(fromCodePoint.apply(void 0, integers), 10) : 0;
        if (codePoints[c] === FULL_STOP) {
            c++;
        }
        var fraction = [];
        while (isDigit(codePoints[c])) {
            fraction.push(codePoints[c++]);
        }
        var fracd = fraction.length;
        var frac = fracd ? parseInt(fromCodePoint.apply(void 0, fraction), 10) : 0;
        if (codePoints[c] === E || codePoints[c] === e) {
            c++;
        }
        var expsign = 1;
        if (codePoints[c] === PLUS_SIGN || codePoints[c] === HYPHEN_MINUS) {
            if (codePoints[c] === HYPHEN_MINUS) {
                expsign = -1;
            }
            c++;
        }
        var exponent = [];
        while (isDigit(codePoints[c])) {
            exponent.push(codePoints[c++]);
        }
        var exp = exponent.length ? parseInt(fromCodePoint.apply(void 0, exponent), 10) : 0;
        return sign * (int + frac * Math.pow(10, -fracd)) * Math.pow(10, expsign * exp);
    };
    var LEFT_PARENTHESIS_TOKEN = {
        type: TokenType.LEFT_PARENTHESIS_TOKEN
    };
    var RIGHT_PARENTHESIS_TOKEN = {
        type: TokenType.RIGHT_PARENTHESIS_TOKEN
    };
    var COMMA_TOKEN = { type: TokenType.COMMA_TOKEN };
    var SUFFIX_MATCH_TOKEN = { type: TokenType.SUFFIX_MATCH_TOKEN };
    var PREFIX_MATCH_TOKEN = { type: TokenType.PREFIX_MATCH_TOKEN };
    var COLUMN_TOKEN = { type: TokenType.COLUMN_TOKEN };
    var DASH_MATCH_TOKEN = { type: TokenType.DASH_MATCH_TOKEN };
    var INCLUDE_MATCH_TOKEN = { type: TokenType.INCLUDE_MATCH_TOKEN };
    var LEFT_CURLY_BRACKET_TOKEN = {
        type: TokenType.LEFT_CURLY_BRACKET_TOKEN
    };
    var RIGHT_CURLY_BRACKET_TOKEN = {
        type: TokenType.RIGHT_CURLY_BRACKET_TOKEN
    };
    var SUBSTRING_MATCH_TOKEN = { type: TokenType.SUBSTRING_MATCH_TOKEN };
    var BAD_URL_TOKEN = { type: TokenType.BAD_URL_TOKEN };
    var BAD_STRING_TOKEN = { type: TokenType.BAD_STRING_TOKEN };
    var CDO_TOKEN = { type: TokenType.CDO_TOKEN };
    var CDC_TOKEN = { type: TokenType.CDC_TOKEN };
    var COLON_TOKEN = { type: TokenType.COLON_TOKEN };
    var SEMICOLON_TOKEN = { type: TokenType.SEMICOLON_TOKEN };
    var LEFT_SQUARE_BRACKET_TOKEN = {
        type: TokenType.LEFT_SQUARE_BRACKET_TOKEN
    };
    var RIGHT_SQUARE_BRACKET_TOKEN = {
        type: TokenType.RIGHT_SQUARE_BRACKET_TOKEN
    };
    var WHITESPACE_TOKEN = { type: TokenType.WHITESPACE_TOKEN };
    var EOF_TOKEN = { type: TokenType.EOF_TOKEN };
    var Tokenizer = /** @class */ (function () {
        function Tokenizer() {
            this._value = [];
        }
        Tokenizer.prototype.write = function (chunk) {
            this._value = this._value.concat(toCodePoints(chunk));
        };
        Tokenizer.prototype.read = function () {
            var tokens = [];
            var token = this.consumeToken();
            while (token !== EOF_TOKEN) {
                tokens.push(token);
                token = this.consumeToken();
            }
            return tokens;
        };
        Tokenizer.prototype.consumeToken = function () {
            var codePoint = this.consumeCodePoint();
            switch (codePoint) {
                case QUOTATION_MARK:
                    return this.consumeStringToken(QUOTATION_MARK);
                case NUMBER_SIGN:
                    var c1 = this.peekCodePoint(0);
                    var c2 = this.peekCodePoint(1);
                    var c3 = this.peekCodePoint(2);
                    if (isNameCodePoint(c1) || isValidEscape(c2, c3)) {
                        var flags = isIdentifierStart(c1, c2, c3) ? FLAG_ID : FLAG_UNRESTRICTED;
                        var value = this.consumeName();
                        return { type: TokenType.HASH_TOKEN, value: value, flags: flags };
                    }
                    break;
                case DOLLAR_SIGN:
                    if (this.peekCodePoint(0) === EQUALS_SIGN) {
                        this.consumeCodePoint();
                        return SUFFIX_MATCH_TOKEN;
                    }
                    break;
                case APOSTROPHE:
                    return this.consumeStringToken(APOSTROPHE);
                case LEFT_PARENTHESIS:
                    return LEFT_PARENTHESIS_TOKEN;
                case RIGHT_PARENTHESIS:
                    return RIGHT_PARENTHESIS_TOKEN;
                case ASTERISK:
                    if (this.peekCodePoint(0) === EQUALS_SIGN) {
                        this.consumeCodePoint();
                        return SUBSTRING_MATCH_TOKEN;
                    }
                    break;
                case PLUS_SIGN:
                    if (isNumberStart(codePoint, this.peekCodePoint(0), this.peekCodePoint(1))) {
                        this.reconsumeCodePoint(codePoint);
                        return this.consumeNumericToken();
                    }
                    break;
                case COMMA:
                    return COMMA_TOKEN;
                case HYPHEN_MINUS:
                    var e1 = codePoint;
                    var e2 = this.peekCodePoint(0);
                    var e3 = this.peekCodePoint(1);
                    if (isNumberStart(e1, e2, e3)) {
                        this.reconsumeCodePoint(codePoint);
                        return this.consumeNumericToken();
                    }
                    if (isIdentifierStart(e1, e2, e3)) {
                        this.reconsumeCodePoint(codePoint);
                        return this.consumeIdentLikeToken();
                    }
                    if (e2 === HYPHEN_MINUS && e3 === GREATER_THAN_SIGN) {
                        this.consumeCodePoint();
                        this.consumeCodePoint();
                        return CDC_TOKEN;
                    }
                    break;
                case FULL_STOP:
                    if (isNumberStart(codePoint, this.peekCodePoint(0), this.peekCodePoint(1))) {
                        this.reconsumeCodePoint(codePoint);
                        return this.consumeNumericToken();
                    }
                    break;
                case SOLIDUS:
                    if (this.peekCodePoint(0) === ASTERISK) {
                        this.consumeCodePoint();
                        while (true) {
                            var c = this.consumeCodePoint();
                            if (c === ASTERISK) {
                                c = this.consumeCodePoint();
                                if (c === SOLIDUS) {
                                    return this.consumeToken();
                                }
                            }
                            if (c === EOF) {
                                return this.consumeToken();
                            }
                        }
                    }
                    break;
                case COLON:
                    return COLON_TOKEN;
                case SEMICOLON:
                    return SEMICOLON_TOKEN;
                case LESS_THAN_SIGN:
                    if (this.peekCodePoint(0) === EXCLAMATION_MARK &&
                        this.peekCodePoint(1) === HYPHEN_MINUS &&
                        this.peekCodePoint(2) === HYPHEN_MINUS) {
                        this.consumeCodePoint();
                        this.consumeCodePoint();
                        return CDO_TOKEN;
                    }
                    break;
                case COMMERCIAL_AT:
                    var a1 = this.peekCodePoint(0);
                    var a2 = this.peekCodePoint(1);
                    var a3 = this.peekCodePoint(2);
                    if (isIdentifierStart(a1, a2, a3)) {
                        var value = this.consumeName();
                        return { type: TokenType.AT_KEYWORD_TOKEN, value: value };
                    }
                    break;
                case LEFT_SQUARE_BRACKET:
                    return LEFT_SQUARE_BRACKET_TOKEN;
                case REVERSE_SOLIDUS:
                    if (isValidEscape(codePoint, this.peekCodePoint(0))) {
                        this.reconsumeCodePoint(codePoint);
                        return this.consumeIdentLikeToken();
                    }
                    break;
                case RIGHT_SQUARE_BRACKET:
                    return RIGHT_SQUARE_BRACKET_TOKEN;
                case CIRCUMFLEX_ACCENT:
                    if (this.peekCodePoint(0) === EQUALS_SIGN) {
                        this.consumeCodePoint();
                        return PREFIX_MATCH_TOKEN;
                    }
                    break;
                case LEFT_CURLY_BRACKET:
                    return LEFT_CURLY_BRACKET_TOKEN;
                case RIGHT_CURLY_BRACKET:
                    return RIGHT_CURLY_BRACKET_TOKEN;
                case u:
                case U:
                    var u1 = this.peekCodePoint(0);
                    var u2 = this.peekCodePoint(1);
                    if (u1 === PLUS_SIGN && (isHex(u2) || u2 === QUESTION_MARK)) {
                        this.consumeCodePoint();
                        this.consumeUnicodeRangeToken();
                    }
                    this.reconsumeCodePoint(codePoint);
                    return this.consumeIdentLikeToken();
                case VERTICAL_LINE:
                    if (this.peekCodePoint(0) === EQUALS_SIGN) {
                        this.consumeCodePoint();
                        return DASH_MATCH_TOKEN;
                    }
                    if (this.peekCodePoint(0) === VERTICAL_LINE) {
                        this.consumeCodePoint();
                        return COLUMN_TOKEN;
                    }
                    break;
                case TILDE:
                    if (this.peekCodePoint(0) === EQUALS_SIGN) {
                        this.consumeCodePoint();
                        return INCLUDE_MATCH_TOKEN;
                    }
                    break;
                case EOF:
                    return EOF_TOKEN;
            }
            if (isWhiteSpace(codePoint)) {
                this.consumeWhiteSpace();
                return WHITESPACE_TOKEN;
            }
            if (isDigit(codePoint)) {
                this.reconsumeCodePoint(codePoint);
                return this.consumeNumericToken();
            }
            if (isNameStartCodePoint(codePoint)) {
                this.reconsumeCodePoint(codePoint);
                return this.consumeIdentLikeToken();
            }
            return { type: TokenType.DELIM_TOKEN, value: fromCodePoint(codePoint) };
        };
        Tokenizer.prototype.consumeCodePoint = function () {
            var value = this._value.shift();
            return typeof value === 'undefined' ? -1 : value;
        };
        Tokenizer.prototype.reconsumeCodePoint = function (codePoint) {
            this._value.unshift(codePoint);
        };
        Tokenizer.prototype.peekCodePoint = function (delta) {
            if (delta >= this._value.length) {
                return -1;
            }
            return this._value[delta];
        };
        Tokenizer.prototype.consumeUnicodeRangeToken = function () {
            var digits = [];
            var codePoint = this.consumeCodePoint();
            while (isHex(codePoint) && digits.length < 6) {
                digits.push(codePoint);
                codePoint = this.consumeCodePoint();
            }
            var questionMarks = false;
            while (codePoint === QUESTION_MARK && digits.length < 6) {
                digits.push(codePoint);
                codePoint = this.consumeCodePoint();
                questionMarks = true;
            }
            if (questionMarks) {
                var start_1 = parseInt(fromCodePoint.apply(void 0, digits.map(function (digit) { return (digit === QUESTION_MARK ? ZERO : digit); })), 16);
                var end = parseInt(fromCodePoint.apply(void 0, digits.map(function (digit) { return (digit === QUESTION_MARK ? F : digit); })), 16);
                return { type: TokenType.UNICODE_RANGE_TOKEN, start: start_1, end: end };
            }
            var start = parseInt(fromCodePoint.apply(void 0, digits), 16);
            if (this.peekCodePoint(0) === HYPHEN_MINUS && isHex(this.peekCodePoint(1))) {
                this.consumeCodePoint();
                codePoint = this.consumeCodePoint();
                var endDigits = [];
                while (isHex(codePoint) && endDigits.length < 6) {
                    endDigits.push(codePoint);
                    codePoint = this.consumeCodePoint();
                }
                var end = parseInt(fromCodePoint.apply(void 0, endDigits), 16);
                return { type: TokenType.UNICODE_RANGE_TOKEN, start: start, end: end };
            }
            else {
                return { type: TokenType.UNICODE_RANGE_TOKEN, start: start, end: start };
            }
        };
        Tokenizer.prototype.consumeIdentLikeToken = function () {
            var value = this.consumeName();
            if (value.toLowerCase() === 'url' && this.peekCodePoint(0) === LEFT_PARENTHESIS) {
                this.consumeCodePoint();
                return this.consumeUrlToken();
            }
            else if (this.peekCodePoint(0) === LEFT_PARENTHESIS) {
                this.consumeCodePoint();
                return { type: TokenType.FUNCTION_TOKEN, value: value };
            }
            return { type: TokenType.IDENT_TOKEN, value: value };
        };
        Tokenizer.prototype.consumeUrlToken = function () {
            var value = [];
            this.consumeWhiteSpace();
            if (this.peekCodePoint(0) === EOF) {
                return { type: TokenType.URL_TOKEN, value: '' };
            }
            var next = this.peekCodePoint(0);
            if (next === APOSTROPHE || next === QUOTATION_MARK) {
                var stringToken = this.consumeStringToken(this.consumeCodePoint());
                if (stringToken.type === TokenType.STRING_TOKEN) {
                    this.consumeWhiteSpace();
                    if (this.peekCodePoint(0) === EOF || this.peekCodePoint(0) === RIGHT_PARENTHESIS) {
                        this.consumeCodePoint();
                        return { type: TokenType.URL_TOKEN, value: stringToken.value };
                    }
                }
                this.consumeBadUrlRemnants();
                return BAD_URL_TOKEN;
            }
            while (true) {
                var codePoint = this.consumeCodePoint();
                if (codePoint === EOF || codePoint === RIGHT_PARENTHESIS) {
                    return { type: TokenType.URL_TOKEN, value: fromCodePoint.apply(void 0, value) };
                }
                else if (isWhiteSpace(codePoint)) {
                    this.consumeWhiteSpace();
                    if (this.peekCodePoint(0) === EOF || this.peekCodePoint(0) === RIGHT_PARENTHESIS) {
                        this.consumeCodePoint();
                        return { type: TokenType.URL_TOKEN, value: fromCodePoint.apply(void 0, value) };
                    }
                    this.consumeBadUrlRemnants();
                    return BAD_URL_TOKEN;
                }
                else if (codePoint === QUOTATION_MARK ||
                    codePoint === APOSTROPHE ||
                    codePoint === LEFT_PARENTHESIS ||
                    isNonPrintableCodePoint(codePoint)) {
                    this.consumeBadUrlRemnants();
                    return BAD_URL_TOKEN;
                }
                else if (codePoint === REVERSE_SOLIDUS) {
                    if (isValidEscape(codePoint, this.peekCodePoint(0))) {
                        value.push(this.consumeEscapedCodePoint());
                    }
                    else {
                        this.consumeBadUrlRemnants();
                        return BAD_URL_TOKEN;
                    }
                }
                else {
                    value.push(codePoint);
                }
            }
        };
        Tokenizer.prototype.consumeWhiteSpace = function () {
            while (isWhiteSpace(this.peekCodePoint(0))) {
                this.consumeCodePoint();
            }
        };
        Tokenizer.prototype.consumeBadUrlRemnants = function () {
            while (true) {
                var codePoint = this.consumeCodePoint();
                if (codePoint === RIGHT_PARENTHESIS || codePoint === EOF) {
                    return;
                }
                if (isValidEscape(codePoint, this.peekCodePoint(0))) {
                    this.consumeEscapedCodePoint();
                }
            }
        };
        Tokenizer.prototype.consumeStringSlice = function (count) {
            var SLICE_STACK_SIZE = 60000;
            var value = '';
            while (count > 0) {
                var amount = Math.min(SLICE_STACK_SIZE, count);
                value += fromCodePoint.apply(void 0, this._value.splice(0, amount));
                count -= amount;
            }
            this._value.shift();
            return value;
        };
        Tokenizer.prototype.consumeStringToken = function (endingCodePoint) {
            var value = '';
            var i = 0;
            do {
                var codePoint = this._value[i];
                if (codePoint === EOF || codePoint === undefined || codePoint === endingCodePoint) {
                    value += this.consumeStringSlice(i);
                    return { type: TokenType.STRING_TOKEN, value: value };
                }
                if (codePoint === LINE_FEED) {
                    this._value.splice(0, i);
                    return BAD_STRING_TOKEN;
                }
                if (codePoint === REVERSE_SOLIDUS) {
                    var next = this._value[i + 1];
                    if (next !== EOF && next !== undefined) {
                        if (next === LINE_FEED) {
                            value += this.consumeStringSlice(i);
                            i = -1;
                            this._value.shift();
                        }
                        else if (isValidEscape(codePoint, next)) {
                            value += this.consumeStringSlice(i);
                            value += fromCodePoint(this.consumeEscapedCodePoint());
                            i = -1;
                        }
                    }
                }
                i++;
            } while (true);
        };
        Tokenizer.prototype.consumeNumber = function () {
            var repr = [];
            var type = FLAG_INTEGER;
            var c1 = this.peekCodePoint(0);
            if (c1 === PLUS_SIGN || c1 === HYPHEN_MINUS) {
                repr.push(this.consumeCodePoint());
            }
            while (isDigit(this.peekCodePoint(0))) {
                repr.push(this.consumeCodePoint());
            }
            c1 = this.peekCodePoint(0);
            var c2 = this.peekCodePoint(1);
            if (c1 === FULL_STOP && isDigit(c2)) {
                repr.push(this.consumeCodePoint(), this.consumeCodePoint());
                type = FLAG_NUMBER;
                while (isDigit(this.peekCodePoint(0))) {
                    repr.push(this.consumeCodePoint());
                }
            }
            c1 = this.peekCodePoint(0);
            c2 = this.peekCodePoint(1);
            var c3 = this.peekCodePoint(2);
            if ((c1 === E || c1 === e) && (((c2 === PLUS_SIGN || c2 === HYPHEN_MINUS) && isDigit(c3)) || isDigit(c2))) {
                repr.push(this.consumeCodePoint(), this.consumeCodePoint());
                type = FLAG_NUMBER;
                while (isDigit(this.peekCodePoint(0))) {
                    repr.push(this.consumeCodePoint());
                }
            }
            return [stringToNumber(repr), type];
        };
        Tokenizer.prototype.consumeNumericToken = function () {
            var _a = this.consumeNumber(), number = _a[0], flags = _a[1];
            var c1 = this.peekCodePoint(0);
            var c2 = this.peekCodePoint(1);
            var c3 = this.peekCodePoint(2);
            if (isIdentifierStart(c1, c2, c3)) {
                var unit = this.consumeName();
                return { type: TokenType.DIMENSION_TOKEN, number: number, flags: flags, unit: unit };
            }
            if (c1 === PERCENTAGE_SIGN) {
                this.consumeCodePoint();
                return { type: TokenType.PERCENTAGE_TOKEN, number: number, flags: flags };
            }
            return { type: TokenType.NUMBER_TOKEN, number: number, flags: flags };
        };
        Tokenizer.prototype.consumeEscapedCodePoint = function () {
            var codePoint = this.consumeCodePoint();
            if (isHex(codePoint)) {
                var hex = fromCodePoint(codePoint);
                while (isHex(this.peekCodePoint(0)) && hex.length < 6) {
                    hex += fromCodePoint(this.consumeCodePoint());
                }
                if (isWhiteSpace(this.peekCodePoint(0))) {
                    this.consumeCodePoint();
                }
                var hexCodePoint = parseInt(hex, 16);
                if (hexCodePoint === 0 || isSurrogateCodePoint(hexCodePoint) || hexCodePoint > 0x10ffff) {
                    return REPLACEMENT_CHARACTER;
                }
                return hexCodePoint;
            }
            if (codePoint === EOF) {
                return REPLACEMENT_CHARACTER;
            }
            return codePoint;
        };
        Tokenizer.prototype.consumeName = function () {
            var result = '';
            while (true) {
                var codePoint = this.consumeCodePoint();
                if (isNameCodePoint(codePoint)) {
                    result += fromCodePoint(codePoint);
                }
                else if (isValidEscape(codePoint, this.peekCodePoint(0))) {
                    result += fromCodePoint(this.consumeEscapedCodePoint());
                }
                else {
                    this.reconsumeCodePoint(codePoint);
                    return result;
                }
            }
        };
        return Tokenizer;
    }());

    var Parser = /** @class */ (function () {
        function Parser(tokens) {
            this._tokens = tokens;
        }
        Parser.create = function (value) {
            var tokenizer = new Tokenizer();
            tokenizer.write(value);
            return new Parser(tokenizer.read());
        };
        Parser.parseValue = function (value) {
            return Parser.create(value).parseComponentValue();
        };
        Parser.parseValues = function (value) {
            return Parser.create(value).parseComponentValues();
        };
        Parser.prototype.parseComponentValue = function () {
            var token = this.consumeToken();
            while (token.type === TokenType.WHITESPACE_TOKEN) {
                token = this.consumeToken();
            }
            if (token.type === TokenType.EOF_TOKEN) {
                throw new SyntaxError("Error parsing CSS component value, unexpected EOF");
            }
            this.reconsumeToken(token);
            var value = this.consumeComponentValue();
            do {
                token = this.consumeToken();
            } while (token.type === TokenType.WHITESPACE_TOKEN);
            if (token.type === TokenType.EOF_TOKEN) {
                return value;
            }
            throw new SyntaxError("Error parsing CSS component value, multiple values found when expecting only one");
        };
        Parser.prototype.parseComponentValues = function () {
            var values = [];
            while (true) {
                var value = this.consumeComponentValue();
                if (value.type === TokenType.EOF_TOKEN) {
                    return values;
                }
                values.push(value);
                values.push();
            }
        };
        Parser.prototype.consumeComponentValue = function () {
            var token = this.consumeToken();
            switch (token.type) {
                case TokenType.LEFT_CURLY_BRACKET_TOKEN:
                case TokenType.LEFT_SQUARE_BRACKET_TOKEN:
                case TokenType.LEFT_PARENTHESIS_TOKEN:
                    return this.consumeSimpleBlock(token.type);
                case TokenType.FUNCTION_TOKEN:
                    return this.consumeFunction(token);
            }
            return token;
        };
        Parser.prototype.consumeSimpleBlock = function (type) {
            var block = { type: type, values: [] };
            var token = this.consumeToken();
            while (true) {
                if (token.type === TokenType.EOF_TOKEN || isEndingTokenFor(token, type)) {
                    return block;
                }
                this.reconsumeToken(token);
                block.values.push(this.consumeComponentValue());
                token = this.consumeToken();
            }
        };
        Parser.prototype.consumeFunction = function (functionToken) {
            var cssFunction = {
                name: functionToken.value,
                values: [],
                type: TokenType.FUNCTION
            };
            while (true) {
                var token = this.consumeToken();
                if (token.type === TokenType.EOF_TOKEN || token.type === TokenType.RIGHT_PARENTHESIS_TOKEN) {
                    return cssFunction;
                }
                this.reconsumeToken(token);
                cssFunction.values.push(this.consumeComponentValue());
            }
        };
        Parser.prototype.consumeToken = function () {
            var token = this._tokens.shift();
            return typeof token === 'undefined' ? EOF_TOKEN : token;
        };
        Parser.prototype.reconsumeToken = function (token) {
            this._tokens.unshift(token);
        };
        return Parser;
    }());
    var isDimensionToken = function (token) { return token.type === TokenType.DIMENSION_TOKEN; };
    var isNumberToken = function (token) { return token.type === TokenType.NUMBER_TOKEN; };
    var isIdentToken = function (token) { return token.type === TokenType.IDENT_TOKEN; };
    var isStringToken = function (token) { return token.type === TokenType.STRING_TOKEN; };
    var isIdentWithValue = function (token, value) {
        return isIdentToken(token) && token.value === value;
    };
    var nonWhiteSpace = function (token) { return token.type !== TokenType.WHITESPACE_TOKEN; };
    var nonFunctionArgSeparator = function (token) {
        return token.type !== TokenType.WHITESPACE_TOKEN && token.type !== TokenType.COMMA_TOKEN;
    };
    var parseFunctionArgs = function (tokens) {
        var args = [];
        var arg = [];
        tokens.forEach(function (token) {
            if (token.type === TokenType.COMMA_TOKEN) {
                if (arg.length === 0) {
                    throw new Error("Error parsing function args, zero tokens for arg");
                }
                args.push(arg);
                arg = [];
                return;
            }
            if (token.type !== TokenType.WHITESPACE_TOKEN) {
                arg.push(token);
            }
        });
        if (arg.length) {
            args.push(arg);
        }
        return args;
    };
    var isEndingTokenFor = function (token, type) {
        if (type === TokenType.LEFT_CURLY_BRACKET_TOKEN && token.type === TokenType.RIGHT_CURLY_BRACKET_TOKEN) {
            return true;
        }
        if (type === TokenType.LEFT_SQUARE_BRACKET_TOKEN && token.type === TokenType.RIGHT_SQUARE_BRACKET_TOKEN) {
            return true;
        }
        return type === TokenType.LEFT_PARENTHESIS_TOKEN && token.type === TokenType.RIGHT_PARENTHESIS_TOKEN;
    };

    var isLength = function (token) {
        return token.type === TokenType.NUMBER_TOKEN || token.type === TokenType.DIMENSION_TOKEN;
    };

    var isLengthPercentage = function (token) {
        return token.type === TokenType.PERCENTAGE_TOKEN || isLength(token);
    };
    var parseLengthPercentageTuple = function (tokens) {
        return tokens.length > 1 ? [tokens[0], tokens[1]] : [tokens[0]];
    };
    var ZERO_LENGTH = {
        type: TokenType.NUMBER_TOKEN,
        number: 0,
        flags: FLAG_INTEGER
    };
    var FIFTY_PERCENT = {
        type: TokenType.PERCENTAGE_TOKEN,
        number: 50,
        flags: FLAG_INTEGER
    };
    var HUNDRED_PERCENT = {
        type: TokenType.PERCENTAGE_TOKEN,
        number: 100,
        flags: FLAG_INTEGER
    };
    var getAbsoluteValueForTuple = function (tuple, width, height) {
        var x = tuple[0], y = tuple[1];
        return [getAbsoluteValue(x, width), getAbsoluteValue(typeof y !== 'undefined' ? y : x, height)];
    };
    var getAbsoluteValue = function (token, parent) {
        if (token.type === TokenType.PERCENTAGE_TOKEN) {
            return (token.number / 100) * parent;
        }
        if (isDimensionToken(token)) {
            switch (token.unit) {
                case 'rem':
                case 'em':
                    return 16 * token.number; // TODO use correct font-size
                case 'px':
                default:
                    return token.number;
            }
        }
        return token.number;
    };

    var DEG = 'deg';
    var GRAD = 'grad';
    var RAD = 'rad';
    var TURN = 'turn';
    var angle = {
        name: 'angle',
        parse: function (value) {
            if (value.type === TokenType.DIMENSION_TOKEN) {
                switch (value.unit) {
                    case DEG:
                        return (Math.PI * value.number) / 180;
                    case GRAD:
                        return (Math.PI / 200) * value.number;
                    case RAD:
                        return value.number;
                    case TURN:
                        return Math.PI * 2 * value.number;
                }
            }
            throw new Error("Unsupported angle type");
        }
    };
    var isAngle = function (value) {
        if (value.type === TokenType.DIMENSION_TOKEN) {
            if (value.unit === DEG || value.unit === GRAD || value.unit === RAD || value.unit === TURN) {
                return true;
            }
        }
        return false;
    };
    var parseNamedSide = function (tokens) {
        var sideOrCorner = tokens
            .filter(isIdentToken)
            .map(function (ident) { return ident.value; })
            .join(' ');
        switch (sideOrCorner) {
            case 'to bottom right':
            case 'to right bottom':
            case 'left top':
            case 'top left':
                return [ZERO_LENGTH, ZERO_LENGTH];
            case 'to top':
            case 'bottom':
                return deg(0);
            case 'to bottom left':
            case 'to left bottom':
            case 'right top':
            case 'top right':
                return [ZERO_LENGTH, HUNDRED_PERCENT];
            case 'to right':
            case 'left':
                return deg(90);
            case 'to top left':
            case 'to left top':
            case 'right bottom':
            case 'bottom right':
                return [HUNDRED_PERCENT, HUNDRED_PERCENT];
            case 'to bottom':
            case 'top':
                return deg(180);
            case 'to top right':
            case 'to right top':
            case 'left bottom':
            case 'bottom left':
                return [HUNDRED_PERCENT, ZERO_LENGTH];
            case 'to left':
            case 'right':
                return deg(270);
        }
        return 0;
    };
    var deg = function (deg) { return (Math.PI * deg) / 180; };

    var color = {
        name: 'color',
        parse: function (value) {
            if (value.type === TokenType.FUNCTION) {
                var colorFunction = SUPPORTED_COLOR_FUNCTIONS[value.name];
                if (typeof colorFunction === 'undefined') {
                    throw new Error("Attempting to parse an unsupported color function \"" + value.name + "\"");
                }
                return colorFunction(value.values);
            }
            if (value.type === TokenType.HASH_TOKEN) {
                if (value.value.length === 3) {
                    var r = value.value.substring(0, 1);
                    var g = value.value.substring(1, 2);
                    var b = value.value.substring(2, 3);
                    return pack(parseInt(r + r, 16), parseInt(g + g, 16), parseInt(b + b, 16), 1);
                }
                if (value.value.length === 4) {
                    var r = value.value.substring(0, 1);
                    var g = value.value.substring(1, 2);
                    var b = value.value.substring(2, 3);
                    var a = value.value.substring(3, 4);
                    return pack(parseInt(r + r, 16), parseInt(g + g, 16), parseInt(b + b, 16), parseInt(a + a, 16) / 255);
                }
                if (value.value.length === 6) {
                    var r = value.value.substring(0, 2);
                    var g = value.value.substring(2, 4);
                    var b = value.value.substring(4, 6);
                    return pack(parseInt(r, 16), parseInt(g, 16), parseInt(b, 16), 1);
                }
                if (value.value.length === 8) {
                    var r = value.value.substring(0, 2);
                    var g = value.value.substring(2, 4);
                    var b = value.value.substring(4, 6);
                    var a = value.value.substring(6, 8);
                    return pack(parseInt(r, 16), parseInt(g, 16), parseInt(b, 16), parseInt(a, 16) / 255);
                }
            }
            if (value.type === TokenType.IDENT_TOKEN) {
                var namedColor = COLORS[value.value.toUpperCase()];
                if (typeof namedColor !== 'undefined') {
                    return namedColor;
                }
            }
            return COLORS.TRANSPARENT;
        }
    };
    var isTransparent = function (color) { return (0xff & color) === 0; };
    var asString = function (color) {
        var alpha = 0xff & color;
        var blue = 0xff & (color >> 8);
        var green = 0xff & (color >> 16);
        var red = 0xff & (color >> 24);
        return alpha < 255 ? "rgba(" + red + "," + green + "," + blue + "," + alpha / 255 + ")" : "rgb(" + red + "," + green + "," + blue + ")";
    };
    var pack = function (r, g, b, a) {
        return ((r << 24) | (g << 16) | (b << 8) | (Math.round(a * 255) << 0)) >>> 0;
    };
    var getTokenColorValue = function (token, i) {
        if (token.type === TokenType.NUMBER_TOKEN) {
            return token.number;
        }
        if (token.type === TokenType.PERCENTAGE_TOKEN) {
            var max = i === 3 ? 1 : 255;
            return i === 3 ? (token.number / 100) * max : Math.round((token.number / 100) * max);
        }
        return 0;
    };
    var rgb = function (args) {
        var tokens = args.filter(nonFunctionArgSeparator);
        if (tokens.length === 3) {
            var _a = tokens.map(getTokenColorValue), r = _a[0], g = _a[1], b = _a[2];
            return pack(r, g, b, 1);
        }
        if (tokens.length === 4) {
            var _b = tokens.map(getTokenColorValue), r = _b[0], g = _b[1], b = _b[2], a = _b[3];
            return pack(r, g, b, a);
        }
        return 0;
    };
    function hue2rgb(t1, t2, hue) {
        if (hue < 0) {
            hue += 1;
        }
        if (hue >= 1) {
            hue -= 1;
        }
        if (hue < 1 / 6) {
            return (t2 - t1) * hue * 6 + t1;
        }
        else if (hue < 1 / 2) {
            return t2;
        }
        else if (hue < 2 / 3) {
            return (t2 - t1) * 6 * (2 / 3 - hue) + t1;
        }
        else {
            return t1;
        }
    }
    var hsl = function (args) {
        var tokens = args.filter(nonFunctionArgSeparator);
        var hue = tokens[0], saturation = tokens[1], lightness = tokens[2], alpha = tokens[3];
        var h = (hue.type === TokenType.NUMBER_TOKEN ? deg(hue.number) : angle.parse(hue)) / (Math.PI * 2);
        var s = isLengthPercentage(saturation) ? saturation.number / 100 : 0;
        var l = isLengthPercentage(lightness) ? lightness.number / 100 : 0;
        var a = typeof alpha !== 'undefined' && isLengthPercentage(alpha) ? getAbsoluteValue(alpha, 1) : 1;
        if (s === 0) {
            return pack(l * 255, l * 255, l * 255, 1);
        }
        var t2 = l <= 0.5 ? l * (s + 1) : l + s - l * s;
        var t1 = l * 2 - t2;
        var r = hue2rgb(t1, t2, h + 1 / 3);
        var g = hue2rgb(t1, t2, h);
        var b = hue2rgb(t1, t2, h - 1 / 3);
        return pack(r * 255, g * 255, b * 255, a);
    };
    var SUPPORTED_COLOR_FUNCTIONS = {
        hsl: hsl,
        hsla: hsl,
        rgb: rgb,
        rgba: rgb
    };
    var COLORS = {
        ALICEBLUE: 0xf0f8ffff,
        ANTIQUEWHITE: 0xfaebd7ff,
        AQUA: 0x00ffffff,
        AQUAMARINE: 0x7fffd4ff,
        AZURE: 0xf0ffffff,
        BEIGE: 0xf5f5dcff,
        BISQUE: 0xffe4c4ff,
        BLACK: 0x000000ff,
        BLANCHEDALMOND: 0xffebcdff,
        BLUE: 0x0000ffff,
        BLUEVIOLET: 0x8a2be2ff,
        BROWN: 0xa52a2aff,
        BURLYWOOD: 0xdeb887ff,
        CADETBLUE: 0x5f9ea0ff,
        CHARTREUSE: 0x7fff00ff,
        CHOCOLATE: 0xd2691eff,
        CORAL: 0xff7f50ff,
        CORNFLOWERBLUE: 0x6495edff,
        CORNSILK: 0xfff8dcff,
        CRIMSON: 0xdc143cff,
        CYAN: 0x00ffffff,
        DARKBLUE: 0x00008bff,
        DARKCYAN: 0x008b8bff,
        DARKGOLDENROD: 0xb886bbff,
        DARKGRAY: 0xa9a9a9ff,
        DARKGREEN: 0x006400ff,
        DARKGREY: 0xa9a9a9ff,
        DARKKHAKI: 0xbdb76bff,
        DARKMAGENTA: 0x8b008bff,
        DARKOLIVEGREEN: 0x556b2fff,
        DARKORANGE: 0xff8c00ff,
        DARKORCHID: 0x9932ccff,
        DARKRED: 0x8b0000ff,
        DARKSALMON: 0xe9967aff,
        DARKSEAGREEN: 0x8fbc8fff,
        DARKSLATEBLUE: 0x483d8bff,
        DARKSLATEGRAY: 0x2f4f4fff,
        DARKSLATEGREY: 0x2f4f4fff,
        DARKTURQUOISE: 0x00ced1ff,
        DARKVIOLET: 0x9400d3ff,
        DEEPPINK: 0xff1493ff,
        DEEPSKYBLUE: 0x00bfffff,
        DIMGRAY: 0x696969ff,
        DIMGREY: 0x696969ff,
        DODGERBLUE: 0x1e90ffff,
        FIREBRICK: 0xb22222ff,
        FLORALWHITE: 0xfffaf0ff,
        FORESTGREEN: 0x228b22ff,
        FUCHSIA: 0xff00ffff,
        GAINSBORO: 0xdcdcdcff,
        GHOSTWHITE: 0xf8f8ffff,
        GOLD: 0xffd700ff,
        GOLDENROD: 0xdaa520ff,
        GRAY: 0x808080ff,
        GREEN: 0x008000ff,
        GREENYELLOW: 0xadff2fff,
        GREY: 0x808080ff,
        HONEYDEW: 0xf0fff0ff,
        HOTPINK: 0xff69b4ff,
        INDIANRED: 0xcd5c5cff,
        INDIGO: 0x4b0082ff,
        IVORY: 0xfffff0ff,
        KHAKI: 0xf0e68cff,
        LAVENDER: 0xe6e6faff,
        LAVENDERBLUSH: 0xfff0f5ff,
        LAWNGREEN: 0x7cfc00ff,
        LEMONCHIFFON: 0xfffacdff,
        LIGHTBLUE: 0xadd8e6ff,
        LIGHTCORAL: 0xf08080ff,
        LIGHTCYAN: 0xe0ffffff,
        LIGHTGOLDENRODYELLOW: 0xfafad2ff,
        LIGHTGRAY: 0xd3d3d3ff,
        LIGHTGREEN: 0x90ee90ff,
        LIGHTGREY: 0xd3d3d3ff,
        LIGHTPINK: 0xffb6c1ff,
        LIGHTSALMON: 0xffa07aff,
        LIGHTSEAGREEN: 0x20b2aaff,
        LIGHTSKYBLUE: 0x87cefaff,
        LIGHTSLATEGRAY: 0x778899ff,
        LIGHTSLATEGREY: 0x778899ff,
        LIGHTSTEELBLUE: 0xb0c4deff,
        LIGHTYELLOW: 0xffffe0ff,
        LIME: 0x00ff00ff,
        LIMEGREEN: 0x32cd32ff,
        LINEN: 0xfaf0e6ff,
        MAGENTA: 0xff00ffff,
        MAROON: 0x800000ff,
        MEDIUMAQUAMARINE: 0x66cdaaff,
        MEDIUMBLUE: 0x0000cdff,
        MEDIUMORCHID: 0xba55d3ff,
        MEDIUMPURPLE: 0x9370dbff,
        MEDIUMSEAGREEN: 0x3cb371ff,
        MEDIUMSLATEBLUE: 0x7b68eeff,
        MEDIUMSPRINGGREEN: 0x00fa9aff,
        MEDIUMTURQUOISE: 0x48d1ccff,
        MEDIUMVIOLETRED: 0xc71585ff,
        MIDNIGHTBLUE: 0x191970ff,
        MINTCREAM: 0xf5fffaff,
        MISTYROSE: 0xffe4e1ff,
        MOCCASIN: 0xffe4b5ff,
        NAVAJOWHITE: 0xffdeadff,
        NAVY: 0x000080ff,
        OLDLACE: 0xfdf5e6ff,
        OLIVE: 0x808000ff,
        OLIVEDRAB: 0x6b8e23ff,
        ORANGE: 0xffa500ff,
        ORANGERED: 0xff4500ff,
        ORCHID: 0xda70d6ff,
        PALEGOLDENROD: 0xeee8aaff,
        PALEGREEN: 0x98fb98ff,
        PALETURQUOISE: 0xafeeeeff,
        PALEVIOLETRED: 0xdb7093ff,
        PAPAYAWHIP: 0xffefd5ff,
        PEACHPUFF: 0xffdab9ff,
        PERU: 0xcd853fff,
        PINK: 0xffc0cbff,
        PLUM: 0xdda0ddff,
        POWDERBLUE: 0xb0e0e6ff,
        PURPLE: 0x800080ff,
        REBECCAPURPLE: 0x663399ff,
        RED: 0xff0000ff,
        ROSYBROWN: 0xbc8f8fff,
        ROYALBLUE: 0x4169e1ff,
        SADDLEBROWN: 0x8b4513ff,
        SALMON: 0xfa8072ff,
        SANDYBROWN: 0xf4a460ff,
        SEAGREEN: 0x2e8b57ff,
        SEASHELL: 0xfff5eeff,
        SIENNA: 0xa0522dff,
        SILVER: 0xc0c0c0ff,
        SKYBLUE: 0x87ceebff,
        SLATEBLUE: 0x6a5acdff,
        SLATEGRAY: 0x708090ff,
        SLATEGREY: 0x708090ff,
        SNOW: 0xfffafaff,
        SPRINGGREEN: 0x00ff7fff,
        STEELBLUE: 0x4682b4ff,
        TAN: 0xd2b48cff,
        TEAL: 0x008080ff,
        THISTLE: 0xd8bfd8ff,
        TOMATO: 0xff6347ff,
        TRANSPARENT: 0x00000000,
        TURQUOISE: 0x40e0d0ff,
        VIOLET: 0xee82eeff,
        WHEAT: 0xf5deb3ff,
        WHITE: 0xffffffff,
        WHITESMOKE: 0xf5f5f5ff,
        YELLOW: 0xffff00ff,
        YELLOWGREEN: 0x9acd32ff
    };

    var PropertyDescriptorParsingType;
    (function (PropertyDescriptorParsingType) {
        PropertyDescriptorParsingType[PropertyDescriptorParsingType["VALUE"] = 0] = "VALUE";
        PropertyDescriptorParsingType[PropertyDescriptorParsingType["LIST"] = 1] = "LIST";
        PropertyDescriptorParsingType[PropertyDescriptorParsingType["IDENT_VALUE"] = 2] = "IDENT_VALUE";
        PropertyDescriptorParsingType[PropertyDescriptorParsingType["TYPE_VALUE"] = 3] = "TYPE_VALUE";
        PropertyDescriptorParsingType[PropertyDescriptorParsingType["TOKEN_VALUE"] = 4] = "TOKEN_VALUE";
    })(PropertyDescriptorParsingType || (PropertyDescriptorParsingType = {}));

    var BACKGROUND_CLIP;
    (function (BACKGROUND_CLIP) {
        BACKGROUND_CLIP[BACKGROUND_CLIP["BORDER_BOX"] = 0] = "BORDER_BOX";
        BACKGROUND_CLIP[BACKGROUND_CLIP["PADDING_BOX"] = 1] = "PADDING_BOX";
        BACKGROUND_CLIP[BACKGROUND_CLIP["CONTENT_BOX"] = 2] = "CONTENT_BOX";
    })(BACKGROUND_CLIP || (BACKGROUND_CLIP = {}));
    var backgroundClip = {
        name: 'background-clip',
        initialValue: 'border-box',
        prefix: false,
        type: PropertyDescriptorParsingType.LIST,
        parse: function (tokens) {
            return tokens.map(function (token) {
                if (isIdentToken(token)) {
                    switch (token.value) {
                        case 'padding-box':
                            return BACKGROUND_CLIP.PADDING_BOX;
                        case 'content-box':
                            return BACKGROUND_CLIP.CONTENT_BOX;
                    }
                }
                return BACKGROUND_CLIP.BORDER_BOX;
            });
        }
    };

    var backgroundColor = {
        name: "background-color",
        initialValue: 'transparent',
        prefix: false,
        type: PropertyDescriptorParsingType.TYPE_VALUE,
        format: 'color'
    };

    var parseColorStop = function (args) {
        var color$1 = color.parse(args[0]);
        var stop = args[1];
        return stop && isLengthPercentage(stop) ? { color: color$1, stop: stop } : { color: color$1, stop: null };
    };
    var processColorStops = function (stops, lineLength) {
        var first = stops[0];
        var last = stops[stops.length - 1];
        if (first.stop === null) {
            first.stop = ZERO_LENGTH;
        }
        if (last.stop === null) {
            last.stop = HUNDRED_PERCENT;
        }
        var processStops = [];
        var previous = 0;
        for (var i = 0; i < stops.length; i++) {
            var stop_1 = stops[i].stop;
            if (stop_1 !== null) {
                var absoluteValue = getAbsoluteValue(stop_1, lineLength);
                if (absoluteValue > previous) {
                    processStops.push(absoluteValue);
                }
                else {
                    processStops.push(previous);
                }
                previous = absoluteValue;
            }
            else {
                processStops.push(null);
            }
        }
        var gapBegin = null;
        for (var i = 0; i < processStops.length; i++) {
            var stop_2 = processStops[i];
            if (stop_2 === null) {
                if (gapBegin === null) {
                    gapBegin = i;
                }
            }
            else if (gapBegin !== null) {
                var gapLength = i - gapBegin;
                var beforeGap = processStops[gapBegin - 1];
                var gapValue = (stop_2 - beforeGap) / (gapLength + 1);
                for (var g = 1; g <= gapLength; g++) {
                    processStops[gapBegin + g - 1] = gapValue * g;
                }
                gapBegin = null;
            }
        }
        return stops.map(function (_a, i) {
            var color = _a.color;
            return { color: color, stop: Math.max(Math.min(1, processStops[i] / lineLength), 0) };
        });
    };
    var getAngleFromCorner = function (corner, width, height) {
        var centerX = width / 2;
        var centerY = height / 2;
        var x = getAbsoluteValue(corner[0], width) - centerX;
        var y = centerY - getAbsoluteValue(corner[1], height);
        return (Math.atan2(y, x) + Math.PI * 2) % (Math.PI * 2);
    };
    var calculateGradientDirection = function (angle, width, height) {
        var radian = typeof angle === 'number' ? angle : getAngleFromCorner(angle, width, height);
        var lineLength = Math.abs(width * Math.sin(radian)) + Math.abs(height * Math.cos(radian));
        var halfWidth = width / 2;
        var halfHeight = height / 2;
        var halfLineLength = lineLength / 2;
        var yDiff = Math.sin(radian - Math.PI / 2) * halfLineLength;
        var xDiff = Math.cos(radian - Math.PI / 2) * halfLineLength;
        return [lineLength, halfWidth - xDiff, halfWidth + xDiff, halfHeight - yDiff, halfHeight + yDiff];
    };
    var distance = function (a, b) { return Math.sqrt(a * a + b * b); };
    var findCorner = function (width, height, x, y, closest) {
        var corners = [[0, 0], [0, height], [width, 0], [width, height]];
        return corners.reduce(function (stat, corner) {
            var cx = corner[0], cy = corner[1];
            var d = distance(x - cx, y - cy);
            if (closest ? d < stat.optimumDistance : d > stat.optimumDistance) {
                return {
                    optimumCorner: corner,
                    optimumDistance: d
                };
            }
            return stat;
        }, {
            optimumDistance: closest ? Infinity : -Infinity,
            optimumCorner: null
        }).optimumCorner;
    };
    var calculateRadius = function (gradient, x, y, width, height) {
        var rx = 0;
        var ry = 0;
        switch (gradient.size) {
            case CSSRadialExtent.CLOSEST_SIDE:
                // The ending shape is sized so that that it exactly meets the side of the gradient box closest to the gradient’s center.
                // If the shape is an ellipse, it exactly meets the closest side in each dimension.
                if (gradient.shape === CSSRadialShape.CIRCLE) {
                    rx = ry = Math.min(Math.abs(x), Math.abs(x - width), Math.abs(y), Math.abs(y - height));
                }
                else if (gradient.shape === CSSRadialShape.ELLIPSE) {
                    rx = Math.min(Math.abs(x), Math.abs(x - width));
                    ry = Math.min(Math.abs(y), Math.abs(y - height));
                }
                break;
            case CSSRadialExtent.CLOSEST_CORNER:
                // The ending shape is sized so that that it passes through the corner of the gradient box closest to the gradient’s center.
                // If the shape is an ellipse, the ending shape is given the same aspect-ratio it would have if closest-side were specified.
                if (gradient.shape === CSSRadialShape.CIRCLE) {
                    rx = ry = Math.min(distance(x, y), distance(x, y - height), distance(x - width, y), distance(x - width, y - height));
                }
                else if (gradient.shape === CSSRadialShape.ELLIPSE) {
                    // Compute the ratio ry/rx (which is to be the same as for "closest-side")
                    var c = Math.min(Math.abs(y), Math.abs(y - height)) / Math.min(Math.abs(x), Math.abs(x - width));
                    var _a = findCorner(width, height, x, y, true), cx = _a[0], cy = _a[1];
                    rx = distance(cx - x, (cy - y) / c);
                    ry = c * rx;
                }
                break;
            case CSSRadialExtent.FARTHEST_SIDE:
                // Same as closest-side, except the ending shape is sized based on the farthest side(s)
                if (gradient.shape === CSSRadialShape.CIRCLE) {
                    rx = ry = Math.max(Math.abs(x), Math.abs(x - width), Math.abs(y), Math.abs(y - height));
                }
                else if (gradient.shape === CSSRadialShape.ELLIPSE) {
                    rx = Math.max(Math.abs(x), Math.abs(x - width));
                    ry = Math.max(Math.abs(y), Math.abs(y - height));
                }
                break;
            case CSSRadialExtent.FARTHEST_CORNER:
                // Same as closest-corner, except the ending shape is sized based on the farthest corner.
                // If the shape is an ellipse, the ending shape is given the same aspect ratio it would have if farthest-side were specified.
                if (gradient.shape === CSSRadialShape.CIRCLE) {
                    rx = ry = Math.max(distance(x, y), distance(x, y - height), distance(x - width, y), distance(x - width, y - height));
                }
                else if (gradient.shape === CSSRadialShape.ELLIPSE) {
                    // Compute the ratio ry/rx (which is to be the same as for "farthest-side")
                    var c = Math.max(Math.abs(y), Math.abs(y - height)) / Math.max(Math.abs(x), Math.abs(x - width));
                    var _b = findCorner(width, height, x, y, false), cx = _b[0], cy = _b[1];
                    rx = distance(cx - x, (cy - y) / c);
                    ry = c * rx;
                }
                break;
        }
        if (Array.isArray(gradient.size)) {
            rx = getAbsoluteValue(gradient.size[0], width);
            ry = gradient.size.length === 2 ? getAbsoluteValue(gradient.size[1], height) : rx;
        }
        return [rx, ry];
    };

    var linearGradient = function (tokens) {
        var angle$1 = deg(180);
        var stops = [];
        parseFunctionArgs(tokens).forEach(function (arg, i) {
            if (i === 0) {
                var firstToken = arg[0];
                if (firstToken.type === TokenType.IDENT_TOKEN && firstToken.value === 'to') {
                    angle$1 = parseNamedSide(arg);
                    return;
                }
                else if (isAngle(firstToken)) {
                    angle$1 = angle.parse(firstToken);
                    return;
                }
            }
            var colorStop = parseColorStop(arg);
            stops.push(colorStop);
        });
        return { angle: angle$1, stops: stops, type: CSSImageType.LINEAR_GRADIENT };
    };

    var prefixLinearGradient = function (tokens) {
        var angle$1 = deg(180);
        var stops = [];
        parseFunctionArgs(tokens).forEach(function (arg, i) {
            if (i === 0) {
                var firstToken = arg[0];
                if (firstToken.type === TokenType.IDENT_TOKEN &&
                    ['top', 'left', 'right', 'bottom'].indexOf(firstToken.value) !== -1) {
                    angle$1 = parseNamedSide(arg);
                    return;
                }
                else if (isAngle(firstToken)) {
                    angle$1 = (angle.parse(firstToken) + deg(270)) % deg(360);
                    return;
                }
            }
            var colorStop = parseColorStop(arg);
            stops.push(colorStop);
        });
        return {
            angle: angle$1,
            stops: stops,
            type: CSSImageType.LINEAR_GRADIENT
        };
    };

    var testRangeBounds = function (document) {
        var TEST_HEIGHT = 123;
        if (document.createRange) {
            var range = document.createRange();
            if (range.getBoundingClientRect) {
                var testElement = document.createElement('boundtest');
                testElement.style.height = TEST_HEIGHT + "px";
                testElement.style.display = 'block';
                document.body.appendChild(testElement);
                range.selectNode(testElement);
                var rangeBounds = range.getBoundingClientRect();
                var rangeHeight = Math.round(rangeBounds.height);
                document.body.removeChild(testElement);
                if (rangeHeight === TEST_HEIGHT) {
                    return true;
                }
            }
        }
        return false;
    };
    var testCORS = function () { return typeof new Image().crossOrigin !== 'undefined'; };
    var testResponseType = function () { return typeof new XMLHttpRequest().responseType === 'string'; };
    var testSVG = function (document) {
        var img = new Image();
        var canvas = document.createElement('canvas');
        var ctx = canvas.getContext('2d');
        if (!ctx) {
            return false;
        }
        img.src = "data:image/svg+xml,<svg xmlns='http://www.w3.org/2000/svg'></svg>";
        try {
            ctx.drawImage(img, 0, 0);
            canvas.toDataURL();
        }
        catch (e) {
            return false;
        }
        return true;
    };
    var isGreenPixel = function (data) {
        return data[0] === 0 && data[1] === 255 && data[2] === 0 && data[3] === 255;
    };
    var testForeignObject = function (document) {
        var canvas = document.createElement('canvas');
        var size = 100;
        canvas.width = size;
        canvas.height = size;
        var ctx = canvas.getContext('2d');
        if (!ctx) {
            return Promise.reject(false);
        }
        ctx.fillStyle = 'rgb(0, 255, 0)';
        ctx.fillRect(0, 0, size, size);
        var img = new Image();
        var greenImageSrc = canvas.toDataURL();
        img.src = greenImageSrc;
        var svg = createForeignObjectSVG(size, size, 0, 0, img);
        ctx.fillStyle = 'red';
        ctx.fillRect(0, 0, size, size);
        return loadSerializedSVG(svg)
            .then(function (img) {
            ctx.drawImage(img, 0, 0);
            var data = ctx.getImageData(0, 0, size, size).data;
            ctx.fillStyle = 'red';
            ctx.fillRect(0, 0, size, size);
            var node = document.createElement('div');
            node.style.backgroundImage = "url(" + greenImageSrc + ")";
            node.style.height = size + "px";
            // Firefox 55 does not render inline <img /> tags
            return isGreenPixel(data)
                ? loadSerializedSVG(createForeignObjectSVG(size, size, 0, 0, node))
                : Promise.reject(false);
        })
            .then(function (img) {
            ctx.drawImage(img, 0, 0);
            // Edge does not render background-images
            return isGreenPixel(ctx.getImageData(0, 0, size, size).data);
        })
            .catch(function () { return false; });
    };
    var createForeignObjectSVG = function (width, height, x, y, node) {
        var xmlns = 'http://www.w3.org/2000/svg';
        var svg = document.createElementNS(xmlns, 'svg');
        var foreignObject = document.createElementNS(xmlns, 'foreignObject');
        svg.setAttributeNS(null, 'width', width.toString());
        svg.setAttributeNS(null, 'height', height.toString());
        foreignObject.setAttributeNS(null, 'width', '100%');
        foreignObject.setAttributeNS(null, 'height', '100%');
        foreignObject.setAttributeNS(null, 'x', x.toString());
        foreignObject.setAttributeNS(null, 'y', y.toString());
        foreignObject.setAttributeNS(null, 'externalResourcesRequired', 'true');
        svg.appendChild(foreignObject);
        foreignObject.appendChild(node);
        return svg;
    };
    var loadSerializedSVG = function (svg) {
        return new Promise(function (resolve, reject) {
            var img = new Image();
            img.onload = function () { return resolve(img); };
            img.onerror = reject;
            img.src = "data:image/svg+xml;charset=utf-8," + encodeURIComponent(new XMLSerializer().serializeToString(svg));
        });
    };
    var FEATURES = {
        get SUPPORT_RANGE_BOUNDS() {
            var value = testRangeBounds(document);
            Object.defineProperty(FEATURES, 'SUPPORT_RANGE_BOUNDS', { value: value });
            return value;
        },
        get SUPPORT_SVG_DRAWING() {
            var value = testSVG(document);
            Object.defineProperty(FEATURES, 'SUPPORT_SVG_DRAWING', { value: value });
            return value;
        },
        get SUPPORT_FOREIGNOBJECT_DRAWING() {
            var value = typeof Array.from === 'function' && typeof window.fetch === 'function'
                ? testForeignObject(document)
                : Promise.resolve(false);
            Object.defineProperty(FEATURES, 'SUPPORT_FOREIGNOBJECT_DRAWING', { value: value });
            return value;
        },
        get SUPPORT_CORS_IMAGES() {
            var value = testCORS();
            Object.defineProperty(FEATURES, 'SUPPORT_CORS_IMAGES', { value: value });
            return value;
        },
        get SUPPORT_RESPONSE_TYPE() {
            var value = testResponseType();
            Object.defineProperty(FEATURES, 'SUPPORT_RESPONSE_TYPE', { value: value });
            return value;
        },
        get SUPPORT_CORS_XHR() {
            var value = 'withCredentials' in new XMLHttpRequest();
            Object.defineProperty(FEATURES, 'SUPPORT_CORS_XHR', { value: value });
            return value;
        }
    };

    var Logger = /** @class */ (function () {
        function Logger(_a) {
            var id = _a.id, enabled = _a.enabled;
            this.id = id;
            this.enabled = enabled;
            this.start = Date.now();
        }
        // eslint-disable-next-line @typescript-eslint/no-explicit-any
        Logger.prototype.debug = function () {
            var args = [];
            for (var _i = 0; _i < arguments.length; _i++) {
                args[_i] = arguments[_i];
            }
            if (this.enabled) {
                // eslint-disable-next-line no-console
                if (typeof window !== 'undefined' && window.console && typeof console.debug === 'function') {
                    // eslint-disable-next-line no-console
                    console.debug.apply(console, [this.id, this.getTime() + "ms"].concat(args));
                }
                else {
                    this.info.apply(this, args);
                }
            }
        };
        Logger.prototype.getTime = function () {
            return Date.now() - this.start;
        };
        Logger.create = function (options) {
            Logger.instances[options.id] = new Logger(options);
        };
        Logger.destroy = function (id) {
            delete Logger.instances[id];
        };
        Logger.getInstance = function (id) {
            var instance = Logger.instances[id];
            if (typeof instance === 'undefined') {
                throw new Error("No logger instance found with id " + id);
            }
            return instance;
        };
        // eslint-disable-next-line @typescript-eslint/no-explicit-any
        Logger.prototype.info = function () {
            var args = [];
            for (var _i = 0; _i < arguments.length; _i++) {
                args[_i] = arguments[_i];
            }
            if (this.enabled) {
                // eslint-disable-next-line no-console
                if (typeof window !== 'undefined' && window.console && typeof console.info === 'function') {
                    // eslint-disable-next-line no-console
                    console.info.apply(console, [this.id, this.getTime() + "ms"].concat(args));
                }
            }
        };
        // eslint-disable-next-line @typescript-eslint/no-explicit-any
        Logger.prototype.error = function () {
            var args = [];
            for (var _i = 0; _i < arguments.length; _i++) {
                args[_i] = arguments[_i];
            }
            if (this.enabled) {
                // eslint-disable-next-line no-console
                if (typeof window !== 'undefined' && window.console && typeof console.error === 'function') {
                    // eslint-disable-next-line no-console
                    console.error.apply(console, [this.id, this.getTime() + "ms"].concat(args));
                }
                else {
                    this.info.apply(this, args);
                }
            }
        };
        Logger.instances = {};
        return Logger;
    }());

    var CacheStorage = /** @class */ (function () {
        function CacheStorage() {
        }
        CacheStorage.create = function (name, options) {
            return (CacheStorage._caches[name] = new Cache(name, options));
        };
        CacheStorage.destroy = function (name) {
            delete CacheStorage._caches[name];
        };
        CacheStorage.open = function (name) {
            var cache = CacheStorage._caches[name];
            if (typeof cache !== 'undefined') {
                return cache;
            }
            throw new Error("Cache with key \"" + name + "\" not found");
        };
        CacheStorage.getOrigin = function (url) {
            var link = CacheStorage._link;
            if (!link) {
                return 'about:blank';
            }
            link.href = url;
            link.href = link.href; // IE9, LOL! - http://jsfiddle.net/niklasvh/2e48b/
            return link.protocol + link.hostname + link.port;
        };
        CacheStorage.isSameOrigin = function (src) {
            return CacheStorage.getOrigin(src) === CacheStorage._origin;
        };
        CacheStorage.setContext = function (window) {
            CacheStorage._link = window.document.createElement('a');
            CacheStorage._origin = CacheStorage.getOrigin(window.location.href);
        };
        CacheStorage.getInstance = function () {
            var current = CacheStorage._current;
            if (current === null) {
                throw new Error("No cache instance attached");
            }
            return current;
        };
        CacheStorage.attachInstance = function (cache) {
            CacheStorage._current = cache;
        };
        CacheStorage.detachInstance = function () {
            CacheStorage._current = null;
        };
        CacheStorage._caches = {};
        CacheStorage._origin = 'about:blank';
        CacheStorage._current = null;
        return CacheStorage;
    }());
    var Cache = /** @class */ (function () {
        function Cache(id, options) {
            this.id = id;
            this._options = options;
            this._cache = {};
        }
        Cache.prototype.addImage = function (src) {
            var result = Promise.resolve();
            if (this.has(src)) {
                return result;
            }
            if (isBlobImage(src) || isRenderable(src)) {
                this._cache[src] = this.loadImage(src);
                return result;
            }
            return result;
        };
        // eslint-disable-next-line @typescript-eslint/no-explicit-any
        Cache.prototype.match = function (src) {
            return this._cache[src];
        };
        Cache.prototype.loadImage = function (key) {
            return __awaiter(this, void 0, void 0, function () {
                var isSameOrigin, useCORS, useProxy, src;
                var _this = this;
                return __generator(this, function (_a) {
                    switch (_a.label) {
                        case 0:
                            isSameOrigin = CacheStorage.isSameOrigin(key);
                            useCORS = !isInlineImage(key) && this._options.useCORS === true && FEATURES.SUPPORT_CORS_IMAGES && !isSameOrigin;
                            useProxy = !isInlineImage(key) &&
                                !isSameOrigin &&
                                typeof this._options.proxy === 'string' &&
                                FEATURES.SUPPORT_CORS_XHR &&
                                !useCORS;
                            if (!isSameOrigin && this._options.allowTaint === false && !isInlineImage(key) && !useProxy && !useCORS) {
                                return [2 /*return*/];
                            }
                            src = key;
                            if (!useProxy) return [3 /*break*/, 2];
                            return [4 /*yield*/, this.proxy(src)];
                        case 1:
                            src = _a.sent();
                            _a.label = 2;
                        case 2:
                            Logger.getInstance(this.id).debug("Added image " + key.substring(0, 256));
                            return [4 /*yield*/, new Promise(function (resolve, reject) {
                                    var img = new Image();
                                    img.onload = function () { return resolve(img); };
                                    img.onerror = reject;
                                    //ios safari 10.3 taints canvas with data urls unless crossOrigin is set to anonymous
                                    if (isInlineBase64Image(src) || useCORS) {
                                        img.crossOrigin = 'anonymous';
                                    }
                                    img.src = src;
                                    if (img.complete === true) {
                                        // Inline XML images may fail to parse, throwing an Error later on
                                        setTimeout(function () { return resolve(img); }, 500);
                                    }
                                    if (_this._options.imageTimeout > 0) {
                                        setTimeout(function () { return reject("Timed out (" + _this._options.imageTimeout + "ms) loading image"); }, _this._options.imageTimeout);
                                    }
                                })];
                        case 3: return [2 /*return*/, _a.sent()];
                    }
                });
            });
        };
        Cache.prototype.has = function (key) {
            return typeof this._cache[key] !== 'undefined';
        };
        Cache.prototype.keys = function () {
            return Promise.resolve(Object.keys(this._cache));
        };
        Cache.prototype.proxy = function (src) {
            var _this = this;
            var proxy = this._options.proxy;
            if (!proxy) {
                throw new Error('No proxy defined');
            }
            var key = src.substring(0, 256);
            return new Promise(function (resolve, reject) {
                var responseType = FEATURES.SUPPORT_RESPONSE_TYPE ? 'blob' : 'text';
                var xhr = new XMLHttpRequest();
                xhr.onload = function () {
                    if (xhr.status === 200) {
                        if (responseType === 'text') {
                            resolve(xhr.response);
                        }
                        else {
                            var reader_1 = new FileReader();
                            reader_1.addEventListener('load', function () { return resolve(reader_1.result); }, false);
                            reader_1.addEventListener('error', function (e) { return reject(e); }, false);
                            reader_1.readAsDataURL(xhr.response);
                        }
                    }
                    else {
                        reject("Failed to proxy resource " + key + " with status code " + xhr.status);
                    }
                };
                xhr.onerror = reject;
                xhr.open('GET', proxy + "?url=" + encodeURIComponent(src) + "&responseType=" + responseType);
                if (responseType !== 'text' && xhr instanceof XMLHttpRequest) {
                    xhr.responseType = responseType;
                }
                if (_this._options.imageTimeout) {
                    var timeout_1 = _this._options.imageTimeout;
                    xhr.timeout = timeout_1;
                    xhr.ontimeout = function () { return reject("Timed out (" + timeout_1 + "ms) proxying " + key); };
                }
                xhr.send();
            });
        };
        return Cache;
    }());
    var INLINE_SVG = /^data:image\/svg\+xml/i;
    var INLINE_BASE64 = /^data:image\/.*;base64,/i;
    var INLINE_IMG = /^data:image\/.*/i;
    var isRenderable = function (src) { return FEATURES.SUPPORT_SVG_DRAWING || !isSVG(src); };
    var isInlineImage = function (src) { return INLINE_IMG.test(src); };
    var isInlineBase64Image = function (src) { return INLINE_BASE64.test(src); };
    var isBlobImage = function (src) { return src.substr(0, 4) === 'blob'; };
    var isSVG = function (src) { return src.substr(-3).toLowerCase() === 'svg' || INLINE_SVG.test(src); };

    var webkitGradient = function (tokens) {
        var angle = deg(180);
        var stops = [];
        var type = CSSImageType.LINEAR_GRADIENT;
        var shape = CSSRadialShape.CIRCLE;
        var size = CSSRadialExtent.FARTHEST_CORNER;
        var position = [];
        parseFunctionArgs(tokens).forEach(function (arg, i) {
            var firstToken = arg[0];
            if (i === 0) {
                if (isIdentToken(firstToken) && firstToken.value === 'linear') {
                    type = CSSImageType.LINEAR_GRADIENT;
                    return;
                }
                else if (isIdentToken(firstToken) && firstToken.value === 'radial') {
                    type = CSSImageType.RADIAL_GRADIENT;
                    return;
                }
            }
            if (firstToken.type === TokenType.FUNCTION) {
                if (firstToken.name === 'from') {
                    var color$1 = color.parse(firstToken.values[0]);
                    stops.push({ stop: ZERO_LENGTH, color: color$1 });
                }
                else if (firstToken.name === 'to') {
                    var color$1 = color.parse(firstToken.values[0]);
                    stops.push({ stop: HUNDRED_PERCENT, color: color$1 });
                }
                else if (firstToken.name === 'color-stop') {
                    var values = firstToken.values.filter(nonFunctionArgSeparator);
                    if (values.length === 2) {
                        var color$1 = color.parse(values[1]);
                        var stop_1 = values[0];
                        if (isNumberToken(stop_1)) {
                            stops.push({
                                stop: { type: TokenType.PERCENTAGE_TOKEN, number: stop_1.number * 100, flags: stop_1.flags },
                                color: color$1
                            });
                        }
                    }
                }
            }
        });
        return type === CSSImageType.LINEAR_GRADIENT
            ? {
                angle: (angle + deg(180)) % deg(360),
                stops: stops,
                type: type
            }
            : { size: size, shape: shape, stops: stops, position: position, type: type };
    };

    var CLOSEST_SIDE = 'closest-side';
    var FARTHEST_SIDE = 'farthest-side';
    var CLOSEST_CORNER = 'closest-corner';
    var FARTHEST_CORNER = 'farthest-corner';
    var CIRCLE = 'circle';
    var ELLIPSE = 'ellipse';
    var COVER = 'cover';
    var CONTAIN = 'contain';
    var radialGradient = function (tokens) {
        var shape = CSSRadialShape.CIRCLE;
        var size = CSSRadialExtent.FARTHEST_CORNER;
        var stops = [];
        var position = [];
        parseFunctionArgs(tokens).forEach(function (arg, i) {
            var isColorStop = true;
            if (i === 0) {
                var isAtPosition_1 = false;
                isColorStop = arg.reduce(function (acc, token) {
                    if (isAtPosition_1) {
                        if (isIdentToken(token)) {
                            switch (token.value) {
                                case 'center':
                                    position.push(FIFTY_PERCENT);
                                    return acc;
                                case 'top':
                                case 'left':
                                    position.push(ZERO_LENGTH);
                                    return acc;
                                case 'right':
                                case 'bottom':
                                    position.push(HUNDRED_PERCENT);
                                    return acc;
                            }
                        }
                        else if (isLengthPercentage(token) || isLength(token)) {
                            position.push(token);
                        }
                    }
                    else if (isIdentToken(token)) {
                        switch (token.value) {
                            case CIRCLE:
                                shape = CSSRadialShape.CIRCLE;
                                return false;
                            case ELLIPSE:
                                shape = CSSRadialShape.ELLIPSE;
                                return false;
                            case 'at':
                                isAtPosition_1 = true;
                                return false;
                            case CLOSEST_SIDE:
                                size = CSSRadialExtent.CLOSEST_SIDE;
                                return false;
                            case COVER:
                            case FARTHEST_SIDE:
                                size = CSSRadialExtent.FARTHEST_SIDE;
                                return false;
                            case CONTAIN:
                            case CLOSEST_CORNER:
                                size = CSSRadialExtent.CLOSEST_CORNER;
                                return false;
                            case FARTHEST_CORNER:
                                size = CSSRadialExtent.FARTHEST_CORNER;
                                return false;
                        }
                    }
                    else if (isLength(token) || isLengthPercentage(token)) {
                        if (!Array.isArray(size)) {
                            size = [];
                        }
                        size.push(token);
                        return false;
                    }
                    return acc;
                }, isColorStop);
            }
            if (isColorStop) {
                var colorStop = parseColorStop(arg);
                stops.push(colorStop);
            }
        });
        return { size: size, shape: shape, stops: stops, position: position, type: CSSImageType.RADIAL_GRADIENT };
    };

    var prefixRadialGradient = function (tokens) {
        var shape = CSSRadialShape.CIRCLE;
        var size = CSSRadialExtent.FARTHEST_CORNER;
        var stops = [];
        var position = [];
        parseFunctionArgs(tokens).forEach(function (arg, i) {
            var isColorStop = true;
            if (i === 0) {
                isColorStop = arg.reduce(function (acc, token) {
                    if (isIdentToken(token)) {
                        switch (token.value) {
                            case 'center':
                                position.push(FIFTY_PERCENT);
                                return false;
                            case 'top':
                            case 'left':
                                position.push(ZERO_LENGTH);
                                return false;
                            case 'right':
                            case 'bottom':
                                position.push(HUNDRED_PERCENT);
                                return false;
                        }
                    }
                    else if (isLengthPercentage(token) || isLength(token)) {
                        position.push(token);
                        return false;
                    }
                    return acc;
                }, isColorStop);
            }
            else if (i === 1) {
                isColorStop = arg.reduce(function (acc, token) {
                    if (isIdentToken(token)) {
                        switch (token.value) {
                            case CIRCLE:
                                shape = CSSRadialShape.CIRCLE;
                                return false;
                            case ELLIPSE:
                                shape = CSSRadialShape.ELLIPSE;
                                return false;
                            case CONTAIN:
                            case CLOSEST_SIDE:
                                size = CSSRadialExtent.CLOSEST_SIDE;
                                return false;
                            case FARTHEST_SIDE:
                                size = CSSRadialExtent.FARTHEST_SIDE;
                                return false;
                            case CLOSEST_CORNER:
                                size = CSSRadialExtent.CLOSEST_CORNER;
                                return false;
                            case COVER:
                            case FARTHEST_CORNER:
                                size = CSSRadialExtent.FARTHEST_CORNER;
                                return false;
                        }
                    }
                    else if (isLength(token) || isLengthPercentage(token)) {
                        if (!Array.isArray(size)) {
                            size = [];
                        }
                        size.push(token);
                        return false;
                    }
                    return acc;
                }, isColorStop);
            }
            if (isColorStop) {
                var colorStop = parseColorStop(arg);
                stops.push(colorStop);
            }
        });
        return { size: size, shape: shape, stops: stops, position: position, type: CSSImageType.RADIAL_GRADIENT };
    };

    var CSSImageType;
    (function (CSSImageType) {
        CSSImageType[CSSImageType["URL"] = 0] = "URL";
        CSSImageType[CSSImageType["LINEAR_GRADIENT"] = 1] = "LINEAR_GRADIENT";
        CSSImageType[CSSImageType["RADIAL_GRADIENT"] = 2] = "RADIAL_GRADIENT";
    })(CSSImageType || (CSSImageType = {}));
    var isLinearGradient = function (background) {
        return background.type === CSSImageType.LINEAR_GRADIENT;
    };
    var isRadialGradient = function (background) {
        return background.type === CSSImageType.RADIAL_GRADIENT;
    };
    var CSSRadialShape;
    (function (CSSRadialShape) {
        CSSRadialShape[CSSRadialShape["CIRCLE"] = 0] = "CIRCLE";
        CSSRadialShape[CSSRadialShape["ELLIPSE"] = 1] = "ELLIPSE";
    })(CSSRadialShape || (CSSRadialShape = {}));
    var CSSRadialExtent;
    (function (CSSRadialExtent) {
        CSSRadialExtent[CSSRadialExtent["CLOSEST_SIDE"] = 0] = "CLOSEST_SIDE";
        CSSRadialExtent[CSSRadialExtent["FARTHEST_SIDE"] = 1] = "FARTHEST_SIDE";
        CSSRadialExtent[CSSRadialExtent["CLOSEST_CORNER"] = 2] = "CLOSEST_CORNER";
        CSSRadialExtent[CSSRadialExtent["FARTHEST_CORNER"] = 3] = "FARTHEST_CORNER";
    })(CSSRadialExtent || (CSSRadialExtent = {}));
    var image = {
        name: 'image',
        parse: function (value) {
            if (value.type === TokenType.URL_TOKEN) {
                var image_1 = { url: value.value, type: CSSImageType.URL };
                CacheStorage.getInstance().addImage(value.value);
                return image_1;
            }
            if (value.type === TokenType.FUNCTION) {
                var imageFunction = SUPPORTED_IMAGE_FUNCTIONS[value.name];
                if (typeof imageFunction === 'undefined') {
                    throw new Error("Attempting to parse an unsupported image function \"" + value.name + "\"");
                }
                return imageFunction(value.values);
            }
            throw new Error("Unsupported image type");
        }
    };
    function isSupportedImage(value) {
        return value.type !== TokenType.FUNCTION || SUPPORTED_IMAGE_FUNCTIONS[value.name];
    }
    var SUPPORTED_IMAGE_FUNCTIONS = {
        'linear-gradient': linearGradient,
        '-moz-linear-gradient': prefixLinearGradient,
        '-ms-linear-gradient': prefixLinearGradient,
        '-o-linear-gradient': prefixLinearGradient,
        '-webkit-linear-gradient': prefixLinearGradient,
        'radial-gradient': radialGradient,
        '-moz-radial-gradient': prefixRadialGradient,
        '-ms-radial-gradient': prefixRadialGradient,
        '-o-radial-gradient': prefixRadialGradient,
        '-webkit-radial-gradient': prefixRadialGradient,
        '-webkit-gradient': webkitGradient
    };

    var backgroundImage = {
        name: 'background-image',
        initialValue: 'none',
        type: PropertyDescriptorParsingType.LIST,
        prefix: false,
        parse: function (tokens) {
            if (tokens.length === 0) {
                return [];
            }
            var first = tokens[0];
            if (first.type === TokenType.IDENT_TOKEN && first.value === 'none') {
                return [];
            }
            return tokens.filter(function (value) { return nonFunctionArgSeparator(value) && isSupportedImage(value); }).map(image.parse);
        }
    };

    var backgroundOrigin = {
        name: 'background-origin',
        initialValue: 'border-box',
        prefix: false,
        type: PropertyDescriptorParsingType.LIST,
        parse: function (tokens) {
            return tokens.map(function (token) {
                if (isIdentToken(token)) {
                    switch (token.value) {
                        case 'padding-box':
                            return 1 /* PADDING_BOX */;
                        case 'content-box':
                            return 2 /* CONTENT_BOX */;
                    }
                }
                return 0 /* BORDER_BOX */;
            });
        }
    };

    var backgroundPosition = {
        name: 'background-position',
        initialValue: '0% 0%',
        type: PropertyDescriptorParsingType.LIST,
        prefix: false,
        parse: function (tokens) {
            return parseFunctionArgs(tokens)
                .map(function (values) { return values.filter(isLengthPercentage); })
                .map(parseLengthPercentageTuple);
        }
    };

    var BACKGROUND_REPEAT;
    (function (BACKGROUND_REPEAT) {
        BACKGROUND_REPEAT[BACKGROUND_REPEAT["REPEAT"] = 0] = "REPEAT";
        BACKGROUND_REPEAT[BACKGROUND_REPEAT["NO_REPEAT"] = 1] = "NO_REPEAT";
        BACKGROUND_REPEAT[BACKGROUND_REPEAT["REPEAT_X"] = 2] = "REPEAT_X";
        BACKGROUND_REPEAT[BACKGROUND_REPEAT["REPEAT_Y"] = 3] = "REPEAT_Y";
    })(BACKGROUND_REPEAT || (BACKGROUND_REPEAT = {}));
    var backgroundRepeat = {
        name: 'background-repeat',
        initialValue: 'repeat',
        prefix: false,
        type: PropertyDescriptorParsingType.LIST,
        parse: function (tokens) {
            return parseFunctionArgs(tokens)
                .map(function (values) {
                return values
                    .filter(isIdentToken)
                    .map(function (token) { return token.value; })
                    .join(' ');
            })
                .map(parseBackgroundRepeat);
        }
    };
    var parseBackgroundRepeat = function (value) {
        switch (value) {
            case 'no-repeat':
                return BACKGROUND_REPEAT.NO_REPEAT;
            case 'repeat-x':
            case 'repeat no-repeat':
                return BACKGROUND_REPEAT.REPEAT_X;
            case 'repeat-y':
            case 'no-repeat repeat':
                return BACKGROUND_REPEAT.REPEAT_Y;
            case 'repeat':
            default:
                return BACKGROUND_REPEAT.REPEAT;
        }
    };

    var BACKGROUND_SIZE;
    (function (BACKGROUND_SIZE) {
        BACKGROUND_SIZE["AUTO"] = "auto";
        BACKGROUND_SIZE["CONTAIN"] = "contain";
        BACKGROUND_SIZE["COVER"] = "cover";
    })(BACKGROUND_SIZE || (BACKGROUND_SIZE = {}));
    var backgroundSize = {
        name: 'background-size',
        initialValue: '0',
        prefix: false,
        type: PropertyDescriptorParsingType.LIST,
        parse: function (tokens) {
            return parseFunctionArgs(tokens).map(function (values) { return values.filter(isBackgroundSizeInfoToken); });
        }
    };
    var isBackgroundSizeInfoToken = function (value) {
        return isIdentToken(value) || isLengthPercentage(value);
    };

    var borderColorForSide = function (side) { return ({
        name: "border-" + side + "-color",
        initialValue: 'transparent',
        prefix: false,
        type: PropertyDescriptorParsingType.TYPE_VALUE,
        format: 'color'
    }); };
    var borderTopColor = borderColorForSide('top');
    var borderRightColor = borderColorForSide('right');
    var borderBottomColor = borderColorForSide('bottom');
    var borderLeftColor = borderColorForSide('left');

    var borderRadiusForSide = function (side) { return ({
        name: "border-radius-" + side,
        initialValue: '0 0',
        prefix: false,
        type: PropertyDescriptorParsingType.LIST,
        parse: function (tokens) { return parseLengthPercentageTuple(tokens.filter(isLengthPercentage)); }
    }); };
    var borderTopLeftRadius = borderRadiusForSide('top-left');
    var borderTopRightRadius = borderRadiusForSide('top-right');
    var borderBottomRightRadius = borderRadiusForSide('bottom-right');
    var borderBottomLeftRadius = borderRadiusForSide('bottom-left');

    var BORDER_STYLE;
    (function (BORDER_STYLE) {
        BORDER_STYLE[BORDER_STYLE["NONE"] = 0] = "NONE";
        BORDER_STYLE[BORDER_STYLE["SOLID"] = 1] = "SOLID";
    })(BORDER_STYLE || (BORDER_STYLE = {}));
    var borderStyleForSide = function (side) { return ({
        name: "border-" + side + "-style",
        initialValue: 'solid',
        prefix: false,
        type: PropertyDescriptorParsingType.IDENT_VALUE,
        parse: function (style) {
            switch (style) {
                case 'none':
                    return BORDER_STYLE.NONE;
            }
            return BORDER_STYLE.SOLID;
        }
    }); };
    var borderTopStyle = borderStyleForSide('top');
    var borderRightStyle = borderStyleForSide('right');
    var borderBottomStyle = borderStyleForSide('bottom');
    var borderLeftStyle = borderStyleForSide('left');

    var borderWidthForSide = function (side) { return ({
        name: "border-" + side + "-width",
        initialValue: '0',
        type: PropertyDescriptorParsingType.VALUE,
        prefix: false,
        parse: function (token) {
            if (isDimensionToken(token)) {
                return token.number;
            }
            return 0;
        }
    }); };
    var borderTopWidth = borderWidthForSide('top');
    var borderRightWidth = borderWidthForSide('right');
    var borderBottomWidth = borderWidthForSide('bottom');
    var borderLeftWidth = borderWidthForSide('left');

    var color$1 = {
        name: "color",
        initialValue: 'transparent',
        prefix: false,
        type: PropertyDescriptorParsingType.TYPE_VALUE,
        format: 'color'
    };

    var display = {
        name: 'display',
        initialValue: 'inline-block',
        prefix: false,
        type: PropertyDescriptorParsingType.LIST,
        parse: function (tokens) {
            return tokens.filter(isIdentToken).reduce(function (bit, token) {
                return bit | parseDisplayValue(token.value);
            }, 0 /* NONE */);
        }
    };
    var parseDisplayValue = function (display) {
        switch (display) {
            case 'block':
                return 2 /* BLOCK */;
            case 'inline':
                return 4 /* INLINE */;
            case 'run-in':
                return 8 /* RUN_IN */;
            case 'flow':
                return 16 /* FLOW */;
            case 'flow-root':
                return 32 /* FLOW_ROOT */;
            case 'table':
                return 64 /* TABLE */;
            case 'flex':
            case '-webkit-flex':
                return 128 /* FLEX */;
            case 'grid':
            case '-ms-grid':
                return 256 /* GRID */;
            case 'ruby':
                return 512 /* RUBY */;
            case 'subgrid':
                return 1024 /* SUBGRID */;
            case 'list-item':
                return 2048 /* LIST_ITEM */;
            case 'table-row-group':
                return 4096 /* TABLE_ROW_GROUP */;
            case 'table-header-group':
                return 8192 /* TABLE_HEADER_GROUP */;
            case 'table-footer-group':
                return 16384 /* TABLE_FOOTER_GROUP */;
            case 'table-row':
                return 32768 /* TABLE_ROW */;
            case 'table-cell':
                return 65536 /* TABLE_CELL */;
            case 'table-column-group':
                return 131072 /* TABLE_COLUMN_GROUP */;
            case 'table-column':
                return 262144 /* TABLE_COLUMN */;
            case 'table-caption':
                return 524288 /* TABLE_CAPTION */;
            case 'ruby-base':
                return 1048576 /* RUBY_BASE */;
            case 'ruby-text':
                return 2097152 /* RUBY_TEXT */;
            case 'ruby-base-container':
                return 4194304 /* RUBY_BASE_CONTAINER */;
            case 'ruby-text-container':
                return 8388608 /* RUBY_TEXT_CONTAINER */;
            case 'contents':
                return 16777216 /* CONTENTS */;
            case 'inline-block':
                return 33554432 /* INLINE_BLOCK */;
            case 'inline-list-item':
                return 67108864 /* INLINE_LIST_ITEM */;
            case 'inline-table':
                return 134217728 /* INLINE_TABLE */;
            case 'inline-flex':
                return 268435456 /* INLINE_FLEX */;
            case 'inline-grid':
                return 536870912 /* INLINE_GRID */;
        }
        return 0 /* NONE */;
    };

    var FLOAT;
    (function (FLOAT) {
        FLOAT[FLOAT["NONE"] = 0] = "NONE";
        FLOAT[FLOAT["LEFT"] = 1] = "LEFT";
        FLOAT[FLOAT["RIGHT"] = 2] = "RIGHT";
        FLOAT[FLOAT["INLINE_START"] = 3] = "INLINE_START";
        FLOAT[FLOAT["INLINE_END"] = 4] = "INLINE_END";
    })(FLOAT || (FLOAT = {}));
    var float = {
        name: 'float',
        initialValue: 'none',
        prefix: false,
        type: PropertyDescriptorParsingType.IDENT_VALUE,
        parse: function (float) {
            switch (float) {
                case 'left':
                    return FLOAT.LEFT;
                case 'right':
                    return FLOAT.RIGHT;
                case 'inline-start':
                    return FLOAT.INLINE_START;
                case 'inline-end':
                    return FLOAT.INLINE_END;
            }
            return FLOAT.NONE;
        }
    };

    var letterSpacing = {
        name: 'letter-spacing',
        initialValue: '0',
        prefix: false,
        type: PropertyDescriptorParsingType.VALUE,
        parse: function (token) {
            if (token.type === TokenType.IDENT_TOKEN && token.value === 'normal') {
                return 0;
            }
            if (token.type === TokenType.NUMBER_TOKEN) {
                return token.number;
            }
            if (token.type === TokenType.DIMENSION_TOKEN) {
                return token.number;
            }
            return 0;
        }
    };

    var LINE_BREAK;
    (function (LINE_BREAK) {
        LINE_BREAK["NORMAL"] = "normal";
        LINE_BREAK["STRICT"] = "strict";
    })(LINE_BREAK || (LINE_BREAK = {}));
    var lineBreak = {
        name: 'line-break',
        initialValue: 'normal',
        prefix: false,
        type: PropertyDescriptorParsingType.IDENT_VALUE,
        parse: function (lineBreak) {
            switch (lineBreak) {
                case 'strict':
                    return LINE_BREAK.STRICT;
                case 'normal':
                default:
                    return LINE_BREAK.NORMAL;
            }
        }
    };

    var lineHeight = {
        name: 'line-height',
        initialValue: 'normal',
        prefix: false,
        type: PropertyDescriptorParsingType.TOKEN_VALUE
    };
    var computeLineHeight = function (token, fontSize) {
        if (isIdentToken(token) && token.value === 'normal') {
            return 1.2 * fontSize;
        }
        else if (token.type === TokenType.NUMBER_TOKEN) {
            return fontSize * token.number;
        }
        else if (isLengthPercentage(token)) {
            return getAbsoluteValue(token, fontSize);
        }
        return fontSize;
    };

    var listStyleImage = {
        name: 'list-style-image',
        initialValue: 'none',
        type: PropertyDescriptorParsingType.VALUE,
        prefix: false,
        parse: function (token) {
            if (token.type === TokenType.IDENT_TOKEN && token.value === 'none') {
                return null;
            }
            return image.parse(token);
        }
    };

    var LIST_STYLE_POSITION;
    (function (LIST_STYLE_POSITION) {
        LIST_STYLE_POSITION[LIST_STYLE_POSITION["INSIDE"] = 0] = "INSIDE";
        LIST_STYLE_POSITION[LIST_STYLE_POSITION["OUTSIDE"] = 1] = "OUTSIDE";
    })(LIST_STYLE_POSITION || (LIST_STYLE_POSITION = {}));
    var listStylePosition = {
        name: 'list-style-position',
        initialValue: 'outside',
        prefix: false,
        type: PropertyDescriptorParsingType.IDENT_VALUE,
        parse: function (position) {
            switch (position) {
                case 'inside':
                    return LIST_STYLE_POSITION.INSIDE;
                case 'outside':
                default:
                    return LIST_STYLE_POSITION.OUTSIDE;
            }
        }
    };

    var LIST_STYLE_TYPE;
    (function (LIST_STYLE_TYPE) {
        LIST_STYLE_TYPE[LIST_STYLE_TYPE["NONE"] = -1] = "NONE";
        LIST_STYLE_TYPE[LIST_STYLE_TYPE["DISC"] = 0] = "DISC";
        LIST_STYLE_TYPE[LIST_STYLE_TYPE["CIRCLE"] = 1] = "CIRCLE";
        LIST_STYLE_TYPE[LIST_STYLE_TYPE["SQUARE"] = 2] = "SQUARE";
        LIST_STYLE_TYPE[LIST_STYLE_TYPE["DECIMAL"] = 3] = "DECIMAL";
        LIST_STYLE_TYPE[LIST_STYLE_TYPE["CJK_DECIMAL"] = 4] = "CJK_DECIMAL";
        LIST_STYLE_TYPE[LIST_STYLE_TYPE["DECIMAL_LEADING_ZERO"] = 5] = "DECIMAL_LEADING_ZERO";
        LIST_STYLE_TYPE[LIST_STYLE_TYPE["LOWER_ROMAN"] = 6] = "LOWER_ROMAN";
        LIST_STYLE_TYPE[LIST_STYLE_TYPE["UPPER_ROMAN"] = 7] = "UPPER_ROMAN";
        LIST_STYLE_TYPE[LIST_STYLE_TYPE["LOWER_GREEK"] = 8] = "LOWER_GREEK";
        LIST_STYLE_TYPE[LIST_STYLE_TYPE["LOWER_ALPHA"] = 9] = "LOWER_ALPHA";
        LIST_STYLE_TYPE[LIST_STYLE_TYPE["UPPER_ALPHA"] = 10] = "UPPER_ALPHA";
        LIST_STYLE_TYPE[LIST_STYLE_TYPE["ARABIC_INDIC"] = 11] = "ARABIC_INDIC";
        LIST_STYLE_TYPE[LIST_STYLE_TYPE["ARMENIAN"] = 12] = "ARMENIAN";
        LIST_STYLE_TYPE[LIST_STYLE_TYPE["BENGALI"] = 13] = "BENGALI";
        LIST_STYLE_TYPE[LIST_STYLE_TYPE["CAMBODIAN"] = 14] = "CAMBODIAN";
        LIST_STYLE_TYPE[LIST_STYLE_TYPE["CJK_EARTHLY_BRANCH"] = 15] = "CJK_EARTHLY_BRANCH";
        LIST_STYLE_TYPE[LIST_STYLE_TYPE["CJK_HEAVENLY_STEM"] = 16] = "CJK_HEAVENLY_STEM";
        LIST_STYLE_TYPE[LIST_STYLE_TYPE["CJK_IDEOGRAPHIC"] = 17] = "CJK_IDEOGRAPHIC";
        LIST_STYLE_TYPE[LIST_STYLE_TYPE["DEVANAGARI"] = 18] = "DEVANAGARI";
        LIST_STYLE_TYPE[LIST_STYLE_TYPE["ETHIOPIC_NUMERIC"] = 19] = "ETHIOPIC_NUMERIC";
        LIST_STYLE_TYPE[LIST_STYLE_TYPE["GEORGIAN"] = 20] = "GEORGIAN";
        LIST_STYLE_TYPE[LIST_STYLE_TYPE["GUJARATI"] = 21] = "GUJARATI";
        LIST_STYLE_TYPE[LIST_STYLE_TYPE["GURMUKHI"] = 22] = "GURMUKHI";
        LIST_STYLE_TYPE[LIST_STYLE_TYPE["HEBREW"] = 22] = "HEBREW";
        LIST_STYLE_TYPE[LIST_STYLE_TYPE["HIRAGANA"] = 23] = "HIRAGANA";
        LIST_STYLE_TYPE[LIST_STYLE_TYPE["HIRAGANA_IROHA"] = 24] = "HIRAGANA_IROHA";
        LIST_STYLE_TYPE[LIST_STYLE_TYPE["JAPANESE_FORMAL"] = 25] = "JAPANESE_FORMAL";
        LIST_STYLE_TYPE[LIST_STYLE_TYPE["JAPANESE_INFORMAL"] = 26] = "JAPANESE_INFORMAL";
        LIST_STYLE_TYPE[LIST_STYLE_TYPE["KANNADA"] = 27] = "KANNADA";
        LIST_STYLE_TYPE[LIST_STYLE_TYPE["KATAKANA"] = 28] = "KATAKANA";
        LIST_STYLE_TYPE[LIST_STYLE_TYPE["KATAKANA_IROHA"] = 29] = "KATAKANA_IROHA";
        LIST_STYLE_TYPE[LIST_STYLE_TYPE["KHMER"] = 30] = "KHMER";
        LIST_STYLE_TYPE[LIST_STYLE_TYPE["KOREAN_HANGUL_FORMAL"] = 31] = "KOREAN_HANGUL_FORMAL";
        LIST_STYLE_TYPE[LIST_STYLE_TYPE["KOREAN_HANJA_FORMAL"] = 32] = "KOREAN_HANJA_FORMAL";
        LIST_STYLE_TYPE[LIST_STYLE_TYPE["KOREAN_HANJA_INFORMAL"] = 33] = "KOREAN_HANJA_INFORMAL";
        LIST_STYLE_TYPE[LIST_STYLE_TYPE["LAO"] = 34] = "LAO";
        LIST_STYLE_TYPE[LIST_STYLE_TYPE["LOWER_ARMENIAN"] = 35] = "LOWER_ARMENIAN";
        LIST_STYLE_TYPE[LIST_STYLE_TYPE["MALAYALAM"] = 36] = "MALAYALAM";
        LIST_STYLE_TYPE[LIST_STYLE_TYPE["MONGOLIAN"] = 37] = "MONGOLIAN";
        LIST_STYLE_TYPE[LIST_STYLE_TYPE["MYANMAR"] = 38] = "MYANMAR";
        LIST_STYLE_TYPE[LIST_STYLE_TYPE["ORIYA"] = 39] = "ORIYA";
        LIST_STYLE_TYPE[LIST_STYLE_TYPE["PERSIAN"] = 40] = "PERSIAN";
        LIST_STYLE_TYPE[LIST_STYLE_TYPE["SIMP_CHINESE_FORMAL"] = 41] = "SIMP_CHINESE_FORMAL";
        LIST_STYLE_TYPE[LIST_STYLE_TYPE["SIMP_CHINESE_INFORMAL"] = 42] = "SIMP_CHINESE_INFORMAL";
        LIST_STYLE_TYPE[LIST_STYLE_TYPE["TAMIL"] = 43] = "TAMIL";
        LIST_STYLE_TYPE[LIST_STYLE_TYPE["TELUGU"] = 44] = "TELUGU";
        LIST_STYLE_TYPE[LIST_STYLE_TYPE["THAI"] = 45] = "THAI";
        LIST_STYLE_TYPE[LIST_STYLE_TYPE["TIBETAN"] = 46] = "TIBETAN";
        LIST_STYLE_TYPE[LIST_STYLE_TYPE["TRAD_CHINESE_FORMAL"] = 47] = "TRAD_CHINESE_FORMAL";
        LIST_STYLE_TYPE[LIST_STYLE_TYPE["TRAD_CHINESE_INFORMAL"] = 48] = "TRAD_CHINESE_INFORMAL";
        LIST_STYLE_TYPE[LIST_STYLE_TYPE["UPPER_ARMENIAN"] = 49] = "UPPER_ARMENIAN";
        LIST_STYLE_TYPE[LIST_STYLE_TYPE["DISCLOSURE_OPEN"] = 50] = "DISCLOSURE_OPEN";
        LIST_STYLE_TYPE[LIST_STYLE_TYPE["DISCLOSURE_CLOSED"] = 51] = "DISCLOSURE_CLOSED";
    })(LIST_STYLE_TYPE || (LIST_STYLE_TYPE = {}));
    var listStyleType = {
        name: 'list-style-type',
        initialValue: 'none',
        prefix: false,
        type: PropertyDescriptorParsingType.IDENT_VALUE,
        parse: function (type) {
            switch (type) {
                case 'disc':
                    return LIST_STYLE_TYPE.DISC;
                case 'circle':
                    return LIST_STYLE_TYPE.CIRCLE;
                case 'square':
                    return LIST_STYLE_TYPE.SQUARE;
                case 'decimal':
                    return LIST_STYLE_TYPE.DECIMAL;
                case 'cjk-decimal':
                    return LIST_STYLE_TYPE.CJK_DECIMAL;
                case 'decimal-leading-zero':
                    return LIST_STYLE_TYPE.DECIMAL_LEADING_ZERO;
                case 'lower-roman':
                    return LIST_STYLE_TYPE.LOWER_ROMAN;
                case 'upper-roman':
                    return LIST_STYLE_TYPE.UPPER_ROMAN;
                case 'lower-greek':
                    return LIST_STYLE_TYPE.LOWER_GREEK;
                case 'lower-alpha':
                    return LIST_STYLE_TYPE.LOWER_ALPHA;
                case 'upper-alpha':
                    return LIST_STYLE_TYPE.UPPER_ALPHA;
                case 'arabic-indic':
                    return LIST_STYLE_TYPE.ARABIC_INDIC;
                case 'armenian':
                    return LIST_STYLE_TYPE.ARMENIAN;
                case 'bengali':
                    return LIST_STYLE_TYPE.BENGALI;
                case 'cambodian':
                    return LIST_STYLE_TYPE.CAMBODIAN;
                case 'cjk-earthly-branch':
                    return LIST_STYLE_TYPE.CJK_EARTHLY_BRANCH;
                case 'cjk-heavenly-stem':
                    return LIST_STYLE_TYPE.CJK_HEAVENLY_STEM;
                case 'cjk-ideographic':
                    return LIST_STYLE_TYPE.CJK_IDEOGRAPHIC;
                case 'devanagari':
                    return LIST_STYLE_TYPE.DEVANAGARI;
                case 'ethiopic-numeric':
                    return LIST_STYLE_TYPE.ETHIOPIC_NUMERIC;
                case 'georgian':
                    return LIST_STYLE_TYPE.GEORGIAN;
                case 'gujarati':
                    return LIST_STYLE_TYPE.GUJARATI;
                case 'gurmukhi':
                    return LIST_STYLE_TYPE.GURMUKHI;
                case 'hebrew':
                    return LIST_STYLE_TYPE.HEBREW;
                case 'hiragana':
                    return LIST_STYLE_TYPE.HIRAGANA;
                case 'hiragana-iroha':
                    return LIST_STYLE_TYPE.HIRAGANA_IROHA;
                case 'japanese-formal':
                    return LIST_STYLE_TYPE.JAPANESE_FORMAL;
                case 'japanese-informal':
                    return LIST_STYLE_TYPE.JAPANESE_INFORMAL;
                case 'kannada':
                    return LIST_STYLE_TYPE.KANNADA;
                case 'katakana':
                    return LIST_STYLE_TYPE.KATAKANA;
                case 'katakana-iroha':
                    return LIST_STYLE_TYPE.KATAKANA_IROHA;
                case 'khmer':
                    return LIST_STYLE_TYPE.KHMER;
                case 'korean-hangul-formal':
                    return LIST_STYLE_TYPE.KOREAN_HANGUL_FORMAL;
                case 'korean-hanja-formal':
                    return LIST_STYLE_TYPE.KOREAN_HANJA_FORMAL;
                case 'korean-hanja-informal':
                    return LIST_STYLE_TYPE.KOREAN_HANJA_INFORMAL;
                case 'lao':
                    return LIST_STYLE_TYPE.LAO;
                case 'lower-armenian':
                    return LIST_STYLE_TYPE.LOWER_ARMENIAN;
                case 'malayalam':
                    return LIST_STYLE_TYPE.MALAYALAM;
                case 'mongolian':
                    return LIST_STYLE_TYPE.MONGOLIAN;
                case 'myanmar':
                    return LIST_STYLE_TYPE.MYANMAR;
                case 'oriya':
                    return LIST_STYLE_TYPE.ORIYA;
                case 'persian':
                    return LIST_STYLE_TYPE.PERSIAN;
                case 'simp-chinese-formal':
                    return LIST_STYLE_TYPE.SIMP_CHINESE_FORMAL;
                case 'simp-chinese-informal':
                    return LIST_STYLE_TYPE.SIMP_CHINESE_INFORMAL;
                case 'tamil':
                    return LIST_STYLE_TYPE.TAMIL;
                case 'telugu':
                    return LIST_STYLE_TYPE.TELUGU;
                case 'thai':
                    return LIST_STYLE_TYPE.THAI;
                case 'tibetan':
                    return LIST_STYLE_TYPE.TIBETAN;
                case 'trad-chinese-formal':
                    return LIST_STYLE_TYPE.TRAD_CHINESE_FORMAL;
                case 'trad-chinese-informal':
                    return LIST_STYLE_TYPE.TRAD_CHINESE_INFORMAL;
                case 'upper-armenian':
                    return LIST_STYLE_TYPE.UPPER_ARMENIAN;
                case 'disclosure-open':
                    return LIST_STYLE_TYPE.DISCLOSURE_OPEN;
                case 'disclosure-closed':
                    return LIST_STYLE_TYPE.DISCLOSURE_CLOSED;
                case 'none':
                default:
                    return LIST_STYLE_TYPE.NONE;
            }
        }
    };

    var marginForSide = function (side) { return ({
        name: "margin-" + side,
        initialValue: '0',
        prefix: false,
        type: PropertyDescriptorParsingType.TOKEN_VALUE
    }); };
    var marginTop = marginForSide('top');
    var marginRight = marginForSide('right');
    var marginBottom = marginForSide('bottom');
    var marginLeft = marginForSide('left');

    var OVERFLOW;
    (function (OVERFLOW) {
        OVERFLOW[OVERFLOW["VISIBLE"] = 0] = "VISIBLE";
        OVERFLOW[OVERFLOW["HIDDEN"] = 1] = "HIDDEN";
        OVERFLOW[OVERFLOW["SCROLL"] = 2] = "SCROLL";
        OVERFLOW[OVERFLOW["AUTO"] = 3] = "AUTO";
    })(OVERFLOW || (OVERFLOW = {}));
    var overflow = {
        name: 'overflow',
        initialValue: 'visible',
        prefix: false,
        type: PropertyDescriptorParsingType.LIST,
        parse: function (tokens) {
            return tokens.filter(isIdentToken).map(function (overflow) {
                switch (overflow.value) {
                    case 'hidden':
                        return OVERFLOW.HIDDEN;
                    case 'scroll':
                        return OVERFLOW.SCROLL;
                    case 'auto':
                        return OVERFLOW.AUTO;
                    case 'visible':
                    default:
                        return OVERFLOW.VISIBLE;
                }
            });
        }
    };

    var OVERFLOW_WRAP;
    (function (OVERFLOW_WRAP) {
        OVERFLOW_WRAP["NORMAL"] = "normal";
        OVERFLOW_WRAP["BREAK_WORD"] = "break-word";
    })(OVERFLOW_WRAP || (OVERFLOW_WRAP = {}));
    var overflowWrap = {
        name: 'overflow-wrap',
        initialValue: 'normal',
        prefix: false,
        type: PropertyDescriptorParsingType.IDENT_VALUE,
        parse: function (overflow) {
            switch (overflow) {
                case 'break-word':
                    return OVERFLOW_WRAP.BREAK_WORD;
                case 'normal':
                default:
                    return OVERFLOW_WRAP.NORMAL;
            }
        }
    };

    var paddingForSide = function (side) { return ({
        name: "padding-" + side,
        initialValue: '0',
        prefix: false,
        type: PropertyDescriptorParsingType.TYPE_VALUE,
        format: 'length-percentage'
    }); };
    var paddingTop = paddingForSide('top');
    var paddingRight = paddingForSide('right');
    var paddingBottom = paddingForSide('bottom');
    var paddingLeft = paddingForSide('left');

    var TEXT_ALIGN;
    (function (TEXT_ALIGN) {
        TEXT_ALIGN[TEXT_ALIGN["LEFT"] = 0] = "LEFT";
        TEXT_ALIGN[TEXT_ALIGN["CENTER"] = 1] = "CENTER";
        TEXT_ALIGN[TEXT_ALIGN["RIGHT"] = 2] = "RIGHT";
    })(TEXT_ALIGN || (TEXT_ALIGN = {}));
    var textAlign = {
        name: 'text-align',
        initialValue: 'left',
        prefix: false,
        type: PropertyDescriptorParsingType.IDENT_VALUE,
        parse: function (textAlign) {
            switch (textAlign) {
                case 'right':
                    return TEXT_ALIGN.RIGHT;
                case 'center':
                case 'justify':
                    return TEXT_ALIGN.CENTER;
                case 'left':
                default:
                    return TEXT_ALIGN.LEFT;
            }
        }
    };

    var POSITION;
    (function (POSITION) {
        POSITION[POSITION["STATIC"] = 0] = "STATIC";
        POSITION[POSITION["RELATIVE"] = 1] = "RELATIVE";
        POSITION[POSITION["ABSOLUTE"] = 2] = "ABSOLUTE";
        POSITION[POSITION["FIXED"] = 3] = "FIXED";
        POSITION[POSITION["STICKY"] = 4] = "STICKY";
    })(POSITION || (POSITION = {}));
    var position = {
        name: 'position',
        initialValue: 'static',
        prefix: false,
        type: PropertyDescriptorParsingType.IDENT_VALUE,
        parse: function (position) {
            switch (position) {
                case 'relative':
                    return POSITION.RELATIVE;
                case 'absolute':
                    return POSITION.ABSOLUTE;
                case 'fixed':
                    return POSITION.FIXED;
                case 'sticky':
                    return POSITION.STICKY;
            }
            return POSITION.STATIC;
        }
    };

    var textShadow = {
        name: 'text-shadow',
        initialValue: 'none',
        type: PropertyDescriptorParsingType.LIST,
        prefix: false,
        parse: function (tokens) {
            if (tokens.length === 1 && isIdentWithValue(tokens[0], 'none')) {
                return [];
            }
            return parseFunctionArgs(tokens).map(function (values) {
                var shadow = {
                    color: COLORS.TRANSPARENT,
                    offsetX: ZERO_LENGTH,
                    offsetY: ZERO_LENGTH,
                    blur: ZERO_LENGTH
                };
                var c = 0;
                for (var i = 0; i < values.length; i++) {
                    var token = values[i];
                    if (isLength(token)) {
                        if (c === 0) {
                            shadow.offsetX = token;
                        }
                        else if (c === 1) {
                            shadow.offsetY = token;
                        }
                        else {
                            shadow.blur = token;
                        }
                        c++;
                    }
                    else {
                        shadow.color = color.parse(token);
                    }
                }
                return shadow;
            });
        }
    };

    var TEXT_TRANSFORM;
    (function (TEXT_TRANSFORM) {
        TEXT_TRANSFORM[TEXT_TRANSFORM["NONE"] = 0] = "NONE";
        TEXT_TRANSFORM[TEXT_TRANSFORM["LOWERCASE"] = 1] = "LOWERCASE";
        TEXT_TRANSFORM[TEXT_TRANSFORM["UPPERCASE"] = 2] = "UPPERCASE";
        TEXT_TRANSFORM[TEXT_TRANSFORM["CAPITALIZE"] = 3] = "CAPITALIZE";
    })(TEXT_TRANSFORM || (TEXT_TRANSFORM = {}));
    var textTransform = {
        name: 'text-transform',
        initialValue: 'none',
        prefix: false,
        type: PropertyDescriptorParsingType.IDENT_VALUE,
        parse: function (textTransform) {
            switch (textTransform) {
                case 'uppercase':
                    return TEXT_TRANSFORM.UPPERCASE;
                case 'lowercase':
                    return TEXT_TRANSFORM.LOWERCASE;
                case 'capitalize':
                    return TEXT_TRANSFORM.CAPITALIZE;
            }
            return TEXT_TRANSFORM.NONE;
        }
    };

    var transform = {
        name: 'transform',
        initialValue: 'none',
        prefix: true,
        type: PropertyDescriptorParsingType.VALUE,
        parse: function (token) {
            if (token.type === TokenType.IDENT_TOKEN && token.value === 'none') {
                return null;
            }
            if (token.type === TokenType.FUNCTION) {
                var transformFunction = SUPPORTED_TRANSFORM_FUNCTIONS[token.name];
                if (typeof transformFunction === 'undefined') {
                    throw new Error("Attempting to parse an unsupported transform function \"" + token.name + "\"");
                }
                return transformFunction(token.values);
            }
            return null;
        }
    };
    var matrix = function (args) {
        var values = args.filter(function (arg) { return arg.type === TokenType.NUMBER_TOKEN; }).map(function (arg) { return arg.number; });
        return values.length === 6 ? values : null;
    };
    // doesn't support 3D transforms at the moment
    var matrix3d = function (args) {
        var values = args.filter(function (arg) { return arg.type === TokenType.NUMBER_TOKEN; }).map(function (arg) { return arg.number; });
        var a1 = values[0], b1 = values[1], _a = values[2], _b = values[3], a2 = values[4], b2 = values[5], _c = values[6], _d = values[7], _e = values[8], _f = values[9], _g = values[10], _h = values[11], a4 = values[12], b4 = values[13], _j = values[14], _k = values[15];
        return values.length === 16 ? [a1, b1, a2, b2, a4, b4] : null;
    };
    var SUPPORTED_TRANSFORM_FUNCTIONS = {
        matrix: matrix,
        matrix3d: matrix3d
    };

    var DEFAULT_VALUE = {
        type: TokenType.PERCENTAGE_TOKEN,
        number: 50,
        flags: FLAG_INTEGER
    };
    var DEFAULT = [DEFAULT_VALUE, DEFAULT_VALUE];
    var transformOrigin = {
        name: 'transform-origin',
        initialValue: '50% 50%',
        prefix: true,
        type: PropertyDescriptorParsingType.LIST,
        parse: function (tokens) {
            var origins = tokens.filter(isLengthPercentage);
            if (origins.length !== 2) {
                return DEFAULT;
            }
            return [origins[0], origins[1]];
        }
    };

    var VISIBILITY;
    (function (VISIBILITY) {
        VISIBILITY[VISIBILITY["VISIBLE"] = 0] = "VISIBLE";
        VISIBILITY[VISIBILITY["HIDDEN"] = 1] = "HIDDEN";
        VISIBILITY[VISIBILITY["COLLAPSE"] = 2] = "COLLAPSE";
    })(VISIBILITY || (VISIBILITY = {}));
    var visibility = {
        name: 'visible',
        initialValue: 'none',
        prefix: false,
        type: PropertyDescriptorParsingType.IDENT_VALUE,
        parse: function (visibility) {
            switch (visibility) {
                case 'hidden':
                    return VISIBILITY.HIDDEN;
                case 'collapse':
                    return VISIBILITY.COLLAPSE;
                case 'visible':
                default:
                    return VISIBILITY.VISIBLE;
            }
        }
    };

    var WORD_BREAK;
    (function (WORD_BREAK) {
        WORD_BREAK["NORMAL"] = "normal";
        WORD_BREAK["BREAK_ALL"] = "break-all";
        WORD_BREAK["KEEP_ALL"] = "keep-all";
    })(WORD_BREAK || (WORD_BREAK = {}));
    var wordBreak = {
        name: 'word-break',
        initialValue: 'normal',
        prefix: false,
        type: PropertyDescriptorParsingType.IDENT_VALUE,
        parse: function (wordBreak) {
            switch (wordBreak) {
                case 'break-all':
                    return WORD_BREAK.BREAK_ALL;
                case 'keep-all':
                    return WORD_BREAK.KEEP_ALL;
                case 'normal':
                default:
                    return WORD_BREAK.NORMAL;
            }
        }
    };

    var zIndex = {
        name: 'z-index',
        initialValue: 'auto',
        prefix: false,
        type: PropertyDescriptorParsingType.VALUE,
        parse: function (token) {
            if (token.type === TokenType.IDENT_TOKEN) {
                return { auto: true, order: 0 };
            }
            if (isNumberToken(token)) {
                return { auto: false, order: token.number };
            }
            throw new Error("Invalid z-index number parsed");
        }
    };

    var opacity = {
        name: 'opacity',
        initialValue: '1',
        type: PropertyDescriptorParsingType.VALUE,
        prefix: false,
        parse: function (token) {
            if (isNumberToken(token)) {
                return token.number;
            }
            return 1;
        }
    };

    var textDecorationColor = {
        name: "text-decoration-color",
        initialValue: 'transparent',
        prefix: false,
        type: PropertyDescriptorParsingType.TYPE_VALUE,
        format: 'color'
    };

    var textDecorationLine = {
        name: 'text-decoration-line',
        initialValue: 'none',
        prefix: false,
        type: PropertyDescriptorParsingType.LIST,
        parse: function (tokens) {
            return tokens
                .filter(isIdentToken)
                .map(function (token) {
                switch (token.value) {
                    case 'underline':
                        return 1 /* UNDERLINE */;
                    case 'overline':
                        return 2 /* OVERLINE */;
                    case 'line-through':
                        return 3 /* LINE_THROUGH */;
                    case 'none':
                        return 4 /* BLINK */;
                }
                return 0 /* NONE */;
            })
                .filter(function (line) { return line !== 0 /* NONE */; });
        }
    };

    var fontFamily = {
        name: "font-family",
        initialValue: '',
        prefix: false,
        type: PropertyDescriptorParsingType.LIST,
        parse: function (tokens) {
            return tokens.filter(isStringToken$1).map(function (token) { return token.value; });
        }
    };
    var isStringToken$1 = function (token) {
        return token.type === TokenType.STRING_TOKEN || token.type === TokenType.IDENT_TOKEN;
    };

    var fontSize = {
        name: "font-size",
        initialValue: '0',
        prefix: false,
        type: PropertyDescriptorParsingType.TYPE_VALUE,
        format: 'length'
    };

    var fontWeight = {
        name: 'font-weight',
        initialValue: 'normal',
        type: PropertyDescriptorParsingType.VALUE,
        prefix: false,
        parse: function (token) {
            if (isNumberToken(token)) {
                return token.number;
            }
            if (isIdentToken(token)) {
                switch (token.value) {
                    case 'bold':
                        return 700;
                    case 'normal':
                    default:
                        return 400;
                }
            }
            return 400;
        }
    };

    var fontVariant = {
        name: 'font-variant',
        initialValue: 'none',
        type: PropertyDescriptorParsingType.LIST,
        prefix: false,
        parse: function (tokens) {
            return tokens.filter(isIdentToken).map(function (token) { return token.value; });
        }
    };

    var FONT_STYLE;
    (function (FONT_STYLE) {
        FONT_STYLE["NORMAL"] = "normal";
        FONT_STYLE["ITALIC"] = "italic";
        FONT_STYLE["OBLIQUE"] = "oblique";
    })(FONT_STYLE || (FONT_STYLE = {}));
    var fontStyle = {
        name: 'font-style',
        initialValue: 'normal',
        prefix: false,
        type: PropertyDescriptorParsingType.IDENT_VALUE,
        parse: function (overflow) {
            switch (overflow) {
                case 'oblique':
                    return FONT_STYLE.OBLIQUE;
                case 'italic':
                    return FONT_STYLE.ITALIC;
                case 'normal':
                default:
                    return FONT_STYLE.NORMAL;
            }
        }
    };

    var contains = function (bit, value) { return (bit & value) !== 0; };

    var content = {
        name: 'content',
        initialValue: 'none',
        type: PropertyDescriptorParsingType.LIST,
        prefix: false,
        parse: function (tokens) {
            if (tokens.length === 0) {
                return [];
            }
            var first = tokens[0];
            if (first.type === TokenType.IDENT_TOKEN && first.value === 'none') {
                return [];
            }
            return tokens;
        }
    };

    var counterIncrement = {
        name: 'counter-increment',
        initialValue: 'none',
        prefix: true,
        type: PropertyDescriptorParsingType.LIST,
        parse: function (tokens) {
            if (tokens.length === 0) {
                return null;
            }
            var first = tokens[0];
            if (first.type === TokenType.IDENT_TOKEN && first.value === 'none') {
                return null;
            }
            var increments = [];
            var filtered = tokens.filter(nonWhiteSpace);
            for (var i = 0; i < filtered.length; i++) {
                var counter = filtered[i];
                var next = filtered[i + 1];
                if (counter.type === TokenType.IDENT_TOKEN) {
                    var increment = next && isNumberToken(next) ? next.number : 1;
                    increments.push({ counter: counter.value, increment: increment });
                }
            }
            return increments;
        }
    };

    var counterReset = {
        name: 'counter-reset',
        initialValue: 'none',
        prefix: true,
        type: PropertyDescriptorParsingType.LIST,
        parse: function (tokens) {
            if (tokens.length === 0) {
                return [];
            }
            var resets = [];
            var filtered = tokens.filter(nonWhiteSpace);
            for (var i = 0; i < filtered.length; i++) {
                var counter = filtered[i];
                var next = filtered[i + 1];
                if (isIdentToken(counter) && counter.value !== 'none') {
                    var reset = next && isNumberToken(next) ? next.number : 0;
                    resets.push({ counter: counter.value, reset: reset });
                }
            }
            return resets;
        }
    };

    var quotes = {
        name: 'quotes',
        initialValue: 'none',
        prefix: true,
        type: PropertyDescriptorParsingType.LIST,
        parse: function (tokens) {
            if (tokens.length === 0) {
                return null;
            }
            var first = tokens[0];
            if (first.type === TokenType.IDENT_TOKEN && first.value === 'none') {
                return null;
            }
            var quotes = [];
            var filtered = tokens.filter(isStringToken);
            if (filtered.length % 2 !== 0) {
                return null;
            }
            for (var i = 0; i < filtered.length; i += 2) {
                var open_1 = filtered[i].value;
                var close_1 = filtered[i + 1].value;
                quotes.push({ open: open_1, close: close_1 });
            }
            return quotes;
        }
    };
    var getQuote = function (quotes, depth, open) {
        if (!quotes) {
            return '';
        }
        var quote = quotes[Math.min(depth, quotes.length - 1)];
        if (!quote) {
            return '';
        }
        return open ? quote.open : quote.close;
    };

    var boxShadow = {
        name: 'box-shadow',
        initialValue: 'none',
        type: PropertyDescriptorParsingType.LIST,
        prefix: false,
        parse: function (tokens) {
            if (tokens.length === 1 && isIdentWithValue(tokens[0], 'none')) {
                return [];
            }
            return parseFunctionArgs(tokens).map(function (values) {
                var shadow = {
                    color: 0x000000ff,
                    offsetX: ZERO_LENGTH,
                    offsetY: ZERO_LENGTH,
                    blur: ZERO_LENGTH,
                    spread: ZERO_LENGTH,
                    inset: false
                };
                var c = 0;
                for (var i = 0; i < values.length; i++) {
                    var token = values[i];
                    if (isIdentWithValue(token, 'inset')) {
                        shadow.inset = true;
                    }
                    else if (isLength(token)) {
                        if (c === 0) {
                            shadow.offsetX = token;
                        }
                        else if (c === 1) {
                            shadow.offsetY = token;
                        }
                        else if (c === 2) {
                            shadow.blur = token;
                        }
                        else {
                            shadow.spread = token;
                        }
                        c++;
                    }
                    else {
                        shadow.color = color.parse(token);
                    }
                }
                return shadow;
            });
        }
    };

    var CSSParsedDeclaration = /** @class */ (function () {
        function CSSParsedDeclaration(declaration) {
            this.backgroundClip = parse(backgroundClip, declaration.backgroundClip);
            this.backgroundColor = parse(backgroundColor, declaration.backgroundColor);
            this.backgroundImage = parse(backgroundImage, declaration.backgroundImage);
            this.backgroundOrigin = parse(backgroundOrigin, declaration.backgroundOrigin);
            this.backgroundPosition = parse(backgroundPosition, declaration.backgroundPosition);
            this.backgroundRepeat = parse(backgroundRepeat, declaration.backgroundRepeat);
            this.backgroundSize = parse(backgroundSize, declaration.backgroundSize);
            this.borderTopColor = parse(borderTopColor, declaration.borderTopColor);
            this.borderRightColor = parse(borderRightColor, declaration.borderRightColor);
            this.borderBottomColor = parse(borderBottomColor, declaration.borderBottomColor);
            this.borderLeftColor = parse(borderLeftColor, declaration.borderLeftColor);
            this.borderTopLeftRadius = parse(borderTopLeftRadius, declaration.borderTopLeftRadius);
            this.borderTopRightRadius = parse(borderTopRightRadius, declaration.borderTopRightRadius);
            this.borderBottomRightRadius = parse(borderBottomRightRadius, declaration.borderBottomRightRadius);
            this.borderBottomLeftRadius = parse(borderBottomLeftRadius, declaration.borderBottomLeftRadius);
            this.borderTopStyle = parse(borderTopStyle, declaration.borderTopStyle);
            this.borderRightStyle = parse(borderRightStyle, declaration.borderRightStyle);
            this.borderBottomStyle = parse(borderBottomStyle, declaration.borderBottomStyle);
            this.borderLeftStyle = parse(borderLeftStyle, declaration.borderLeftStyle);
            this.borderTopWidth = parse(borderTopWidth, declaration.borderTopWidth);
            this.borderRightWidth = parse(borderRightWidth, declaration.borderRightWidth);
            this.borderBottomWidth = parse(borderBottomWidth, declaration.borderBottomWidth);
            this.borderLeftWidth = parse(borderLeftWidth, declaration.borderLeftWidth);
            this.boxShadow = parse(boxShadow, declaration.boxShadow);
            this.color = parse(color$1, declaration.color);
            this.display = parse(display, declaration.display);
            this.float = parse(float, declaration.cssFloat);
            this.fontFamily = parse(fontFamily, declaration.fontFamily);
            this.fontSize = parse(fontSize, declaration.fontSize);
            this.fontStyle = parse(fontStyle, declaration.fontStyle);
            this.fontVariant = parse(fontVariant, declaration.fontVariant);
            this.fontWeight = parse(fontWeight, declaration.fontWeight);
            this.letterSpacing = parse(letterSpacing, declaration.letterSpacing);
            this.lineBreak = parse(lineBreak, declaration.lineBreak);
            this.lineHeight = parse(lineHeight, declaration.lineHeight);
            this.listStyleImage = parse(listStyleImage, declaration.listStyleImage);
            this.listStylePosition = parse(listStylePosition, declaration.listStylePosition);
            this.listStyleType = parse(listStyleType, declaration.listStyleType);
            this.marginTop = parse(marginTop, declaration.marginTop);
            this.marginRight = parse(marginRight, declaration.marginRight);
            this.marginBottom = parse(marginBottom, declaration.marginBottom);
            this.marginLeft = parse(marginLeft, declaration.marginLeft);
            this.opacity = parse(opacity, declaration.opacity);
            var overflowTuple = parse(overflow, declaration.overflow);
            this.overflowX = overflowTuple[0];
            this.overflowY = overflowTuple[overflowTuple.length > 1 ? 1 : 0];
            this.overflowWrap = parse(overflowWrap, declaration.overflowWrap);
            this.paddingTop = parse(paddingTop, declaration.paddingTop);
            this.paddingRight = parse(paddingRight, declaration.paddingRight);
            this.paddingBottom = parse(paddingBottom, declaration.paddingBottom);
            this.paddingLeft = parse(paddingLeft, declaration.paddingLeft);
            this.position = parse(position, declaration.position);
            this.textAlign = parse(textAlign, declaration.textAlign);
            this.textDecorationColor = parse(textDecorationColor, declaration.textDecorationColor || declaration.color);
            this.textDecorationLine = parse(textDecorationLine, declaration.textDecorationLine);
            this.textShadow = parse(textShadow, declaration.textShadow);
            this.textTransform = parse(textTransform, declaration.textTransform);
            this.transform = parse(transform, declaration.transform);
            this.transformOrigin = parse(transformOrigin, declaration.transformOrigin);
            this.visibility = parse(visibility, declaration.visibility);
            this.wordBreak = parse(wordBreak, declaration.wordBreak);
            this.zIndex = parse(zIndex, declaration.zIndex);
        }
        CSSParsedDeclaration.prototype.isVisible = function () {
            return this.display > 0 && this.opacity > 0 && this.visibility === VISIBILITY.VISIBLE;
        };
        CSSParsedDeclaration.prototype.isTransparent = function () {
            return isTransparent(this.backgroundColor);
        };
        CSSParsedDeclaration.prototype.isTransformed = function () {
            return this.transform !== null;
        };
        CSSParsedDeclaration.prototype.isPositioned = function () {
            return this.position !== POSITION.STATIC;
        };
        CSSParsedDeclaration.prototype.isPositionedWithZIndex = function () {
            return this.isPositioned() && !this.zIndex.auto;
        };
        CSSParsedDeclaration.prototype.isFloating = function () {
            return this.float !== FLOAT.NONE;
        };
        CSSParsedDeclaration.prototype.isInlineLevel = function () {
            return (contains(this.display, 4 /* INLINE */) ||
                contains(this.display, 33554432 /* INLINE_BLOCK */) ||
                contains(this.display, 268435456 /* INLINE_FLEX */) ||
                contains(this.display, 536870912 /* INLINE_GRID */) ||
                contains(this.display, 67108864 /* INLINE_LIST_ITEM */) ||
                contains(this.display, 134217728 /* INLINE_TABLE */));
        };
        return CSSParsedDeclaration;
    }());
    var CSSParsedPseudoDeclaration = /** @class */ (function () {
        function CSSParsedPseudoDeclaration(declaration) {
            this.content = parse(content, declaration.content);
            this.quotes = parse(quotes, declaration.quotes);
        }
        return CSSParsedPseudoDeclaration;
    }());
    var CSSParsedCounterDeclaration = /** @class */ (function () {
        function CSSParsedCounterDeclaration(declaration) {
            this.counterIncrement = parse(counterIncrement, declaration.counterIncrement);
            this.counterReset = parse(counterReset, declaration.counterReset);
        }
        return CSSParsedCounterDeclaration;
    }());
    // eslint-disable-next-line @typescript-eslint/no-explicit-any
    var parse = function (descriptor, style) {
        var tokenizer = new Tokenizer();
        var value = style !== null && typeof style !== 'undefined' ? style.toString() : descriptor.initialValue;
        tokenizer.write(value);
        var parser = new Parser(tokenizer.read());
        switch (descriptor.type) {
            case PropertyDescriptorParsingType.IDENT_VALUE:
                var token = parser.parseComponentValue();
                return descriptor.parse(isIdentToken(token) ? token.value : descriptor.initialValue);
            case PropertyDescriptorParsingType.VALUE:
                return descriptor.parse(parser.parseComponentValue());
            case PropertyDescriptorParsingType.LIST:
                return descriptor.parse(parser.parseComponentValues());
            case PropertyDescriptorParsingType.TOKEN_VALUE:
                return parser.parseComponentValue();
            case PropertyDescriptorParsingType.TYPE_VALUE:
                switch (descriptor.format) {
                    case 'angle':
                        return angle.parse(parser.parseComponentValue());
                    case 'color':
                        return color.parse(parser.parseComponentValue());
                    case 'image':
                        return image.parse(parser.parseComponentValue());
                    case 'length':
                        var length_1 = parser.parseComponentValue();
                        return isLength(length_1) ? length_1 : ZERO_LENGTH;
                    case 'length-percentage':
                        var value_1 = parser.parseComponentValue();
                        return isLengthPercentage(value_1) ? value_1 : ZERO_LENGTH;
                }
        }
        throw new Error("Attempting to parse unsupported css format type " + descriptor.format);
    };

    var ElementContainer = /** @class */ (function () {
        function ElementContainer(element) {
            this.styles = new CSSParsedDeclaration(window.getComputedStyle(element, null));
            this.textNodes = [];
            this.elements = [];
            if (this.styles.transform !== null && isHTMLElementNode(element)) {
                // getBoundingClientRect takes transforms into account
                element.style.transform = 'none';
            }
            this.bounds = parseBounds(element);
            this.flags = 0;
        }
        return ElementContainer;
    }());

    var TextBounds = /** @class */ (function () {
        function TextBounds(text, bounds) {
            this.text = text;
            this.bounds = bounds;
        }
        return TextBounds;
    }());
    var parseTextBounds = function (value, styles, node) {
        var textList = breakText(value, styles);
        var textBounds = [];
        var offset = 0;
        textList.forEach(function (text) {
            if (styles.textDecorationLine.length || text.trim().length > 0) {
                if (FEATURES.SUPPORT_RANGE_BOUNDS) {
                    textBounds.push(new TextBounds(text, getRangeBounds(node, offset, text.length)));
                }
                else {
                    var replacementNode = node.splitText(text.length);
                    textBounds.push(new TextBounds(text, getWrapperBounds(node)));
                    node = replacementNode;
                }
            }
            else if (!FEATURES.SUPPORT_RANGE_BOUNDS) {
                node = node.splitText(text.length);
            }
            offset += text.length;
        });
        return textBounds;
    };
    var getWrapperBounds = function (node) {
        var ownerDocument = node.ownerDocument;
        if (ownerDocument) {
            var wrapper = ownerDocument.createElement('html2canvaswrapper');
            wrapper.appendChild(node.cloneNode(true));
            var parentNode = node.parentNode;
            if (parentNode) {
                parentNode.replaceChild(wrapper, node);
                var bounds = parseBounds(wrapper);
                if (wrapper.firstChild) {
                    parentNode.replaceChild(wrapper.firstChild, wrapper);
                }
                return bounds;
            }
        }
        return new Bounds(0, 0, 0, 0);
    };
    var getRangeBounds = function (node, offset, length) {
        var ownerDocument = node.ownerDocument;
        if (!ownerDocument) {
            throw new Error('Node has no owner document');
        }
        var range = ownerDocument.createRange();
        range.setStart(node, offset);
        range.setEnd(node, offset + length);
        return Bounds.fromClientRect(range.getBoundingClientRect());
    };
    var breakText = function (value, styles) {
        return styles.letterSpacing !== 0 ? toCodePoints(value).map(function (i) { return fromCodePoint(i); }) : breakWords(value, styles);
    };
    var breakWords = function (str, styles) {
        var breaker = LineBreaker(str, {
            lineBreak: styles.lineBreak,
            wordBreak: styles.overflowWrap === OVERFLOW_WRAP.BREAK_WORD ? 'break-word' : styles.wordBreak
        });
        var words = [];
        var bk;
        while (!(bk = breaker.next()).done) {
            if (bk.value) {
                words.push(bk.value.slice());
            }
        }
        return words;
    };

    var TextContainer = /** @class */ (function () {
        function TextContainer(node, styles) {
            this.text = transform$1(node.data, styles.textTransform);
            this.textBounds = parseTextBounds(this.text, styles, node);
        }
        return TextContainer;
    }());
    var transform$1 = function (text, transform) {
        switch (transform) {
            case TEXT_TRANSFORM.LOWERCASE:
                return text.toLowerCase();
            case TEXT_TRANSFORM.CAPITALIZE:
                return text.replace(CAPITALIZE, capitalize);
            case TEXT_TRANSFORM.UPPERCASE:
                return text.toUpperCase();
            default:
                return text;
        }
    };
    var CAPITALIZE = /(^|\s|:|-|\(|\))([a-z])/g;
    var capitalize = function (m, p1, p2) {
        if (m.length > 0) {
            return p1 + p2.toUpperCase();
        }
        return m;
    };

    var ImageElementContainer = /** @class */ (function (_super) {
        __extends(ImageElementContainer, _super);
        function ImageElementContainer(img) {
            var _this = _super.call(this, img) || this;
            _this.src = img.currentSrc || img.src;
            _this.intrinsicWidth = img.naturalWidth;
            _this.intrinsicHeight = img.naturalHeight;
            CacheStorage.getInstance().addImage(_this.src);
            return _this;
        }
        return ImageElementContainer;
    }(ElementContainer));

    var CanvasElementContainer = /** @class */ (function (_super) {
        __extends(CanvasElementContainer, _super);
        function CanvasElementContainer(canvas) {
            var _this = _super.call(this, canvas) || this;
            _this.canvas = canvas;
            _this.intrinsicWidth = canvas.width;
            _this.intrinsicHeight = canvas.height;
            return _this;
        }
        return CanvasElementContainer;
    }(ElementContainer));

    var SVGElementContainer = /** @class */ (function (_super) {
        __extends(SVGElementContainer, _super);
        function SVGElementContainer(img) {
            var _this = _super.call(this, img) || this;
            var s = new XMLSerializer();
            _this.svg = "data:image/svg+xml," + encodeURIComponent(s.serializeToString(img));
            _this.intrinsicWidth = img.width.baseVal.value;
            _this.intrinsicHeight = img.height.baseVal.value;
            CacheStorage.getInstance().addImage(_this.svg);
            return _this;
        }
        return SVGElementContainer;
    }(ElementContainer));

    var LIElementContainer = /** @class */ (function (_super) {
        __extends(LIElementContainer, _super);
        function LIElementContainer(element) {
            var _this = _super.call(this, element) || this;
            _this.value = element.value;
            return _this;
        }
        return LIElementContainer;
    }(ElementContainer));

    var OLElementContainer = /** @class */ (function (_super) {
        __extends(OLElementContainer, _super);
        function OLElementContainer(element) {
            var _this = _super.call(this, element) || this;
            _this.start = element.start;
            _this.reversed = typeof element.reversed === 'boolean' && element.reversed === true;
            return _this;
        }
        return OLElementContainer;
    }(ElementContainer));

    var CHECKBOX_BORDER_RADIUS = [
        {
            type: TokenType.DIMENSION_TOKEN,
            flags: 0,
            unit: 'px',
            number: 3
        }
    ];
    var RADIO_BORDER_RADIUS = [
        {
            type: TokenType.PERCENTAGE_TOKEN,
            flags: 0,
            number: 50
        }
    ];
    var reformatInputBounds = function (bounds) {
        if (bounds.width > bounds.height) {
            return new Bounds(bounds.left + (bounds.width - bounds.height) / 2, bounds.top, bounds.height, bounds.height);
        }
        else if (bounds.width < bounds.height) {
            return new Bounds(bounds.left, bounds.top + (bounds.height - bounds.width) / 2, bounds.width, bounds.width);
        }
        return bounds;
    };
    var getInputValue = function (node) {
        var value = node.type === PASSWORD ? new Array(node.value.length + 1).join('\u2022') : node.value;
        return value.length === 0 ? node.placeholder || '' : value;
    };
    var CHECKBOX = 'checkbox';
    var RADIO = 'radio';
    var PASSWORD = 'password';
    var INPUT_COLOR = 0x2a2a2aff;
    var InputElementContainer = /** @class */ (function (_super) {
        __extends(InputElementContainer, _super);
        function InputElementContainer(input) {
            var _this = _super.call(this, input) || this;
            _this.type = input.type.toLowerCase();
            _this.checked = input.checked;
            _this.value = getInputValue(input);
            if (_this.type === CHECKBOX || _this.type === RADIO) {
                _this.styles.backgroundColor = 0xdededeff;
                _this.styles.borderTopColor = _this.styles.borderRightColor = _this.styles.borderBottomColor = _this.styles.borderLeftColor = 0xa5a5a5ff;
                _this.styles.borderTopWidth = _this.styles.borderRightWidth = _this.styles.borderBottomWidth = _this.styles.borderLeftWidth = 1;
                _this.styles.borderTopStyle = _this.styles.borderRightStyle = _this.styles.borderBottomStyle = _this.styles.borderLeftStyle =
                    BORDER_STYLE.SOLID;
                _this.styles.backgroundClip = [BACKGROUND_CLIP.BORDER_BOX];
                _this.styles.backgroundOrigin = [0 /* BORDER_BOX */];
                _this.bounds = reformatInputBounds(_this.bounds);
            }
            switch (_this.type) {
                case CHECKBOX:
                    _this.styles.borderTopRightRadius = _this.styles.borderTopLeftRadius = _this.styles.borderBottomRightRadius = _this.styles.borderBottomLeftRadius = CHECKBOX_BORDER_RADIUS;
                    break;
                case RADIO:
                    _this.styles.borderTopRightRadius = _this.styles.borderTopLeftRadius = _this.styles.borderBottomRightRadius = _this.styles.borderBottomLeftRadius = RADIO_BORDER_RADIUS;
                    break;
            }
            return _this;
        }
        return InputElementContainer;
    }(ElementContainer));

    var SelectElementContainer = /** @class */ (function (_super) {
        __extends(SelectElementContainer, _super);
        function SelectElementContainer(element) {
            var _this = _super.call(this, element) || this;
            var option = element.options[element.selectedIndex || 0];
            _this.value = option ? option.text || '' : '';
            return _this;
        }
        return SelectElementContainer;
    }(ElementContainer));

    var TextareaElementContainer = /** @class */ (function (_super) {
        __extends(TextareaElementContainer, _super);
        function TextareaElementContainer(element) {
            var _this = _super.call(this, element) || this;
            _this.value = element.value;
            return _this;
        }
        return TextareaElementContainer;
    }(ElementContainer));

    var parseColor = function (value) { return color.parse(Parser.create(value).parseComponentValue()); };
    var IFrameElementContainer = /** @class */ (function (_super) {
        __extends(IFrameElementContainer, _super);
        function IFrameElementContainer(iframe) {
            var _this = _super.call(this, iframe) || this;
            _this.src = iframe.src;
            _this.width = parseInt(iframe.width, 10) || 0;
            _this.height = parseInt(iframe.height, 10) || 0;
            _this.backgroundColor = _this.styles.backgroundColor;
            try {
                if (iframe.contentWindow &&
                    iframe.contentWindow.document &&
                    iframe.contentWindow.document.documentElement) {
                    _this.tree = parseTree(iframe.contentWindow.document.documentElement);
                    // http://www.w3.org/TR/css3-background/#special-backgrounds
                    var documentBackgroundColor = iframe.contentWindow.document.documentElement
                        ? parseColor(getComputedStyle(iframe.contentWindow.document.documentElement)
                            .backgroundColor)
                        : COLORS.TRANSPARENT;
                    var bodyBackgroundColor = iframe.contentWindow.document.body
                        ? parseColor(getComputedStyle(iframe.contentWindow.document.body).backgroundColor)
                        : COLORS.TRANSPARENT;
                    _this.backgroundColor = isTransparent(documentBackgroundColor)
                        ? isTransparent(bodyBackgroundColor)
                            ? _this.styles.backgroundColor
                            : bodyBackgroundColor
                        : documentBackgroundColor;
                }
            }
            catch (e) { }
            return _this;
        }
        return IFrameElementContainer;
    }(ElementContainer));

    var LIST_OWNERS = ['OL', 'UL', 'MENU'];
    var parseNodeTree = function (node, parent, root) {
        for (var childNode = node.firstChild, nextNode = void 0; childNode; childNode = nextNode) {
            nextNode = childNode.nextSibling;
            if (isTextNode(childNode) && childNode.data.trim().length > 0) {
                parent.textNodes.push(new TextContainer(childNode, parent.styles));
            }
            else if (isElementNode(childNode)) {
                var container = createContainer(childNode);
                if (container.styles.isVisible()) {
                    if (createsRealStackingContext(childNode, container, root)) {
                        container.flags |= 4 /* CREATES_REAL_STACKING_CONTEXT */;
                    }
                    else if (createsStackingContext(container.styles)) {
                        container.flags |= 2 /* CREATES_STACKING_CONTEXT */;
                    }
                    if (LIST_OWNERS.indexOf(childNode.tagName) !== -1) {
                        container.flags |= 8 /* IS_LIST_OWNER */;
                    }
                    parent.elements.push(container);
                    if (!isTextareaElement(childNode) && !isSVGElement(childNode) && !isSelectElement(childNode)) {
                        parseNodeTree(childNode, container, root);
                    }
                }
            }
        }
    };
    var createContainer = function (element) {
        if (isImageElement(element)) {
            return new ImageElementContainer(element);
        }
        if (isCanvasElement(element)) {
            return new CanvasElementContainer(element);
        }
        if (isSVGElement(element)) {
            return new SVGElementContainer(element);
        }
        if (isLIElement(element)) {
            return new LIElementContainer(element);
        }
        if (isOLElement(element)) {
            return new OLElementContainer(element);
        }
        if (isInputElement(element)) {
            return new InputElementContainer(element);
        }
        if (isSelectElement(element)) {
            return new SelectElementContainer(element);
        }
        if (isTextareaElement(element)) {
            return new TextareaElementContainer(element);
        }
        if (isIFrameElement(element)) {
            return new IFrameElementContainer(element);
        }
        return new ElementContainer(element);
    };
    var parseTree = function (element) {
        var container = createContainer(element);
        container.flags |= 4 /* CREATES_REAL_STACKING_CONTEXT */;
        parseNodeTree(element, container, container);
        return container;
    };
    var createsRealStackingContext = function (node, container, root) {
        return (container.styles.isPositionedWithZIndex() ||
            container.styles.opacity < 1 ||
            container.styles.isTransformed() ||
            (isBodyElement(node) && root.styles.isTransparent()));
    };
    var createsStackingContext = function (styles) { return styles.isPositioned() || styles.isFloating(); };
    var isTextNode = function (node) { return node.nodeType === Node.TEXT_NODE; };
    var isElementNode = function (node) { return node.nodeType === Node.ELEMENT_NODE; };
    var isHTMLElementNode = function (node) {
        return typeof node.style !== 'undefined';
    };
    var isSVGElementNode = function (element) {
        return typeof element.className === 'object';
    };
    var isLIElement = function (node) { return node.tagName === 'LI'; };
    var isOLElement = function (node) { return node.tagName === 'OL'; };
    var isInputElement = function (node) { return node.tagName === 'INPUT'; };
    var isHTMLElement = function (node) { return node.tagName === 'HTML'; };
    var isSVGElement = function (node) { return node.tagName === 'svg'; };
    var isBodyElement = function (node) { return node.tagName === 'BODY'; };
    var isCanvasElement = function (node) { return node.tagName === 'CANVAS'; };
    var isImageElement = function (node) { return node.tagName === 'IMG'; };
    var isIFrameElement = function (node) { return node.tagName === 'IFRAME'; };
    var isStyleElement = function (node) { return node.tagName === 'STYLE'; };
    var isScriptElement = function (node) { return node.tagName === 'SCRIPT'; };
    var isTextareaElement = function (node) { return node.tagName === 'TEXTAREA'; };
    var isSelectElement = function (node) { return node.tagName === 'SELECT'; };

    var CounterState = /** @class */ (function () {
        function CounterState() {
            this.counters = {};
        }
        CounterState.prototype.getCounterValue = function (name) {
            var counter = this.counters[name];
            if (counter && counter.length) {
                return counter[counter.length - 1];
            }
            return 1;
        };
        CounterState.prototype.getCounterValues = function (name) {
            var counter = this.counters[name];
            return counter ? counter : [];
        };
        CounterState.prototype.pop = function (counters) {
            var _this = this;
            counters.forEach(function (counter) { return _this.counters[counter].pop(); });
        };
        CounterState.prototype.parse = function (style) {
            var _this = this;
            var counterIncrement = style.counterIncrement;
            var counterReset = style.counterReset;
            var canReset = true;
            if (counterIncrement !== null) {
                counterIncrement.forEach(function (entry) {
                    var counter = _this.counters[entry.counter];
                    if (counter && entry.increment !== 0) {
                        canReset = false;
                        counter[Math.max(0, counter.length - 1)] += entry.increment;
                    }
                });
            }
            var counterNames = [];
            if (canReset) {
                counterReset.forEach(function (entry) {
                    var counter = _this.counters[entry.counter];
                    counterNames.push(entry.counter);
                    if (!counter) {
                        counter = _this.counters[entry.counter] = [];
                    }
                    counter.push(entry.reset);
                });
            }
            return counterNames;
        };
        return CounterState;
    }());
    var ROMAN_UPPER = {
        integers: [1000, 900, 500, 400, 100, 90, 50, 40, 10, 9, 5, 4, 1],
        values: ['M', 'CM', 'D', 'CD', 'C', 'XC', 'L', 'XL', 'X', 'IX', 'V', 'IV', 'I']
    };
    var ARMENIAN = {
        integers: [
            9000,
            8000,
            7000,
            6000,
            5000,
            4000,
            3000,
            2000,
            1000,
            900,
            800,
            700,
            600,
            500,
            400,
            300,
            200,
            100,
            90,
            80,
            70,
            60,
            50,
            40,
            30,
            20,
            10,
            9,
            8,
            7,
            6,
            5,
            4,
            3,
            2,
            1
        ],
        values: [
            'Ք',
            'Փ',
            'Ւ',
            'Ց',
            'Ր',
            'Տ',
            'Վ',
            'Ս',
            'Ռ',
            'Ջ',
            'Պ',
            'Չ',
            'Ո',
            'Շ',
            'Ն',
            'Յ',
            'Մ',
            'Ճ',
            'Ղ',
            'Ձ',
            'Հ',
            'Կ',
            'Ծ',
            'Խ',
            'Լ',
            'Ի',
            'Ժ',
            'Թ',
            'Ը',
            'Է',
            'Զ',
            'Ե',
            'Դ',
            'Գ',
            'Բ',
            'Ա'
        ]
    };
    var HEBREW = {
        integers: [
            10000,
            9000,
            8000,
            7000,
            6000,
            5000,
            4000,
            3000,
            2000,
            1000,
            400,
            300,
            200,
            100,
            90,
            80,
            70,
            60,
            50,
            40,
            30,
            20,
            19,
            18,
            17,
            16,
            15,
            10,
            9,
            8,
            7,
            6,
            5,
            4,
            3,
            2,
            1
        ],
        values: [
            'י׳',
            'ט׳',
            'ח׳',
            'ז׳',
            'ו׳',
            'ה׳',
            'ד׳',
            'ג׳',
            'ב׳',
            'א׳',
            'ת',
            'ש',
            'ר',
            'ק',
            'צ',
            'פ',
            'ע',
            'ס',
            'נ',
            'מ',
            'ל',
            'כ',
            'יט',
            'יח',
            'יז',
            'טז',
            'טו',
            'י',
            'ט',
            'ח',
            'ז',
            'ו',
            'ה',
            'ד',
            'ג',
            'ב',
            'א'
        ]
    };
    var GEORGIAN = {
        integers: [
            10000,
            9000,
            8000,
            7000,
            6000,
            5000,
            4000,
            3000,
            2000,
            1000,
            900,
            800,
            700,
            600,
            500,
            400,
            300,
            200,
            100,
            90,
            80,
            70,
            60,
            50,
            40,
            30,
            20,
            10,
            9,
            8,
            7,
            6,
            5,
            4,
            3,
            2,
            1
        ],
        values: [
            'ჵ',
            'ჰ',
            'ჯ',
            'ჴ',
            'ხ',
            'ჭ',
            'წ',
            'ძ',
            'ც',
            'ჩ',
            'შ',
            'ყ',
            'ღ',
            'ქ',
            'ფ',
            'ჳ',
            'ტ',
            'ს',
            'რ',
            'ჟ',
            'პ',
            'ო',
            'ჲ',
            'ნ',
            'მ',
            'ლ',
            'კ',
            'ი',
            'თ',
            'ჱ',
            'ზ',
            'ვ',
            'ე',
            'დ',
            'გ',
            'ბ',
            'ა'
        ]
    };
    var createAdditiveCounter = function (value, min, max, symbols, fallback, suffix) {
        if (value < min || value > max) {
            return createCounterText(value, fallback, suffix.length > 0);
        }
        return (symbols.integers.reduce(function (string, integer, index) {
            while (value >= integer) {
                value -= integer;
                string += symbols.values[index];
            }
            return string;
        }, '') + suffix);
    };
    var createCounterStyleWithSymbolResolver = function (value, codePointRangeLength, isNumeric, resolver) {
        var string = '';
        do {
            if (!isNumeric) {
                value--;
            }
            string = resolver(value) + string;
            value /= codePointRangeLength;
        } while (value * codePointRangeLength >= codePointRangeLength);
        return string;
    };
    var createCounterStyleFromRange = function (value, codePointRangeStart, codePointRangeEnd, isNumeric, suffix) {
        var codePointRangeLength = codePointRangeEnd - codePointRangeStart + 1;
        return ((value < 0 ? '-' : '') +
            (createCounterStyleWithSymbolResolver(Math.abs(value), codePointRangeLength, isNumeric, function (codePoint) {
                return fromCodePoint(Math.floor(codePoint % codePointRangeLength) + codePointRangeStart);
            }) +
                suffix));
    };
    var createCounterStyleFromSymbols = function (value, symbols, suffix) {
        if (suffix === void 0) { suffix = '. '; }
        var codePointRangeLength = symbols.length;
        return (createCounterStyleWithSymbolResolver(Math.abs(value), codePointRangeLength, false, function (codePoint) { return symbols[Math.floor(codePoint % codePointRangeLength)]; }) + suffix);
    };
    var CJK_ZEROS = 1 << 0;
    var CJK_TEN_COEFFICIENTS = 1 << 1;
    var CJK_TEN_HIGH_COEFFICIENTS = 1 << 2;
    var CJK_HUNDRED_COEFFICIENTS = 1 << 3;
    var createCJKCounter = function (value, numbers, multipliers, negativeSign, suffix, flags) {
        if (value < -9999 || value > 9999) {
            return createCounterText(value, LIST_STYLE_TYPE.CJK_DECIMAL, suffix.length > 0);
        }
        var tmp = Math.abs(value);
        var string = suffix;
        if (tmp === 0) {
            return numbers[0] + string;
        }
        for (var digit = 0; tmp > 0 && digit <= 4; digit++) {
            var coefficient = tmp % 10;
            if (coefficient === 0 && contains(flags, CJK_ZEROS) && string !== '') {
                string = numbers[coefficient] + string;
            }
            else if (coefficient > 1 ||
                (coefficient === 1 && digit === 0) ||
                (coefficient === 1 && digit === 1 && contains(flags, CJK_TEN_COEFFICIENTS)) ||
                (coefficient === 1 && digit === 1 && contains(flags, CJK_TEN_HIGH_COEFFICIENTS) && value > 100) ||
                (coefficient === 1 && digit > 1 && contains(flags, CJK_HUNDRED_COEFFICIENTS))) {
                string = numbers[coefficient] + (digit > 0 ? multipliers[digit - 1] : '') + string;
            }
            else if (coefficient === 1 && digit > 0) {
                string = multipliers[digit - 1] + string;
            }
            tmp = Math.floor(tmp / 10);
        }
        return (value < 0 ? negativeSign : '') + string;
    };
    var CHINESE_INFORMAL_MULTIPLIERS = '十百千萬';
    var CHINESE_FORMAL_MULTIPLIERS = '拾佰仟萬';
    var JAPANESE_NEGATIVE = 'マイナス';
    var KOREAN_NEGATIVE = '마이너스';
    var createCounterText = function (value, type, appendSuffix) {
        var defaultSuffix = appendSuffix ? '. ' : '';
        var cjkSuffix = appendSuffix ? '、' : '';
        var koreanSuffix = appendSuffix ? ', ' : '';
        var spaceSuffix = appendSuffix ? ' ' : '';
        switch (type) {
            case LIST_STYLE_TYPE.DISC:
                return '•' + spaceSuffix;
            case LIST_STYLE_TYPE.CIRCLE:
                return '◦' + spaceSuffix;
            case LIST_STYLE_TYPE.SQUARE:
                return '◾' + spaceSuffix;
            case LIST_STYLE_TYPE.DECIMAL_LEADING_ZERO:
                var string = createCounterStyleFromRange(value, 48, 57, true, defaultSuffix);
                return string.length < 4 ? "0" + string : string;
            case LIST_STYLE_TYPE.CJK_DECIMAL:
                return createCounterStyleFromSymbols(value, '〇一二三四五六七八九', cjkSuffix);
            case LIST_STYLE_TYPE.LOWER_ROMAN:
                return createAdditiveCounter(value, 1, 3999, ROMAN_UPPER, LIST_STYLE_TYPE.DECIMAL, defaultSuffix).toLowerCase();
            case LIST_STYLE_TYPE.UPPER_ROMAN:
                return createAdditiveCounter(value, 1, 3999, ROMAN_UPPER, LIST_STYLE_TYPE.DECIMAL, defaultSuffix);
            case LIST_STYLE_TYPE.LOWER_GREEK:
                return createCounterStyleFromRange(value, 945, 969, false, defaultSuffix);
            case LIST_STYLE_TYPE.LOWER_ALPHA:
                return createCounterStyleFromRange(value, 97, 122, false, defaultSuffix);
            case LIST_STYLE_TYPE.UPPER_ALPHA:
                return createCounterStyleFromRange(value, 65, 90, false, defaultSuffix);
            case LIST_STYLE_TYPE.ARABIC_INDIC:
                return createCounterStyleFromRange(value, 1632, 1641, true, defaultSuffix);
            case LIST_STYLE_TYPE.ARMENIAN:
            case LIST_STYLE_TYPE.UPPER_ARMENIAN:
                return createAdditiveCounter(value, 1, 9999, ARMENIAN, LIST_STYLE_TYPE.DECIMAL, defaultSuffix);
            case LIST_STYLE_TYPE.LOWER_ARMENIAN:
                return createAdditiveCounter(value, 1, 9999, ARMENIAN, LIST_STYLE_TYPE.DECIMAL, defaultSuffix).toLowerCase();
            case LIST_STYLE_TYPE.BENGALI:
                return createCounterStyleFromRange(value, 2534, 2543, true, defaultSuffix);
            case LIST_STYLE_TYPE.CAMBODIAN:
            case LIST_STYLE_TYPE.KHMER:
                return createCounterStyleFromRange(value, 6112, 6121, true, defaultSuffix);
            case LIST_STYLE_TYPE.CJK_EARTHLY_BRANCH:
                return createCounterStyleFromSymbols(value, '子丑寅卯辰巳午未申酉戌亥', cjkSuffix);
            case LIST_STYLE_TYPE.CJK_HEAVENLY_STEM:
                return createCounterStyleFromSymbols(value, '甲乙丙丁戊己庚辛壬癸', cjkSuffix);
            case LIST_STYLE_TYPE.CJK_IDEOGRAPHIC:
            case LIST_STYLE_TYPE.TRAD_CHINESE_INFORMAL:
                return createCJKCounter(value, '零一二三四五六七八九', CHINESE_INFORMAL_MULTIPLIERS, '負', cjkSuffix, CJK_TEN_COEFFICIENTS | CJK_TEN_HIGH_COEFFICIENTS | CJK_HUNDRED_COEFFICIENTS);
            case LIST_STYLE_TYPE.TRAD_CHINESE_FORMAL:
                return createCJKCounter(value, '零壹貳參肆伍陸柒捌玖', CHINESE_FORMAL_MULTIPLIERS, '負', cjkSuffix, CJK_ZEROS | CJK_TEN_COEFFICIENTS | CJK_TEN_HIGH_COEFFICIENTS | CJK_HUNDRED_COEFFICIENTS);
            case LIST_STYLE_TYPE.SIMP_CHINESE_INFORMAL:
                return createCJKCounter(value, '零一二三四五六七八九', CHINESE_INFORMAL_MULTIPLIERS, '负', cjkSuffix, CJK_TEN_COEFFICIENTS | CJK_TEN_HIGH_COEFFICIENTS | CJK_HUNDRED_COEFFICIENTS);
            case LIST_STYLE_TYPE.SIMP_CHINESE_FORMAL:
                return createCJKCounter(value, '零壹贰叁肆伍陆柒捌玖', CHINESE_FORMAL_MULTIPLIERS, '负', cjkSuffix, CJK_ZEROS | CJK_TEN_COEFFICIENTS | CJK_TEN_HIGH_COEFFICIENTS | CJK_HUNDRED_COEFFICIENTS);
            case LIST_STYLE_TYPE.JAPANESE_INFORMAL:
                return createCJKCounter(value, '〇一二三四五六七八九', '十百千万', JAPANESE_NEGATIVE, cjkSuffix, 0);
            case LIST_STYLE_TYPE.JAPANESE_FORMAL:
                return createCJKCounter(value, '零壱弐参四伍六七八九', '拾百千万', JAPANESE_NEGATIVE, cjkSuffix, CJK_ZEROS | CJK_TEN_COEFFICIENTS | CJK_TEN_HIGH_COEFFICIENTS);
            case LIST_STYLE_TYPE.KOREAN_HANGUL_FORMAL:
                return createCJKCounter(value, '영일이삼사오육칠팔구', '십백천만', KOREAN_NEGATIVE, koreanSuffix, CJK_ZEROS | CJK_TEN_COEFFICIENTS | CJK_TEN_HIGH_COEFFICIENTS);
            case LIST_STYLE_TYPE.KOREAN_HANJA_INFORMAL:
                return createCJKCounter(value, '零一二三四五六七八九', '十百千萬', KOREAN_NEGATIVE, koreanSuffix, 0);
            case LIST_STYLE_TYPE.KOREAN_HANJA_FORMAL:
                return createCJKCounter(value, '零壹貳參四五六七八九', '拾百千', KOREAN_NEGATIVE, koreanSuffix, CJK_ZEROS | CJK_TEN_COEFFICIENTS | CJK_TEN_HIGH_COEFFICIENTS);
            case LIST_STYLE_TYPE.DEVANAGARI:
                return createCounterStyleFromRange(value, 0x966, 0x96f, true, defaultSuffix);
            case LIST_STYLE_TYPE.GEORGIAN:
                return createAdditiveCounter(value, 1, 19999, GEORGIAN, LIST_STYLE_TYPE.DECIMAL, defaultSuffix);
            case LIST_STYLE_TYPE.GUJARATI:
                return createCounterStyleFromRange(value, 0xae6, 0xaef, true, defaultSuffix);
            case LIST_STYLE_TYPE.GURMUKHI:
                return createCounterStyleFromRange(value, 0xa66, 0xa6f, true, defaultSuffix);
            case LIST_STYLE_TYPE.HEBREW:
                return createAdditiveCounter(value, 1, 10999, HEBREW, LIST_STYLE_TYPE.DECIMAL, defaultSuffix);
            case LIST_STYLE_TYPE.HIRAGANA:
                return createCounterStyleFromSymbols(value, 'あいうえおかきくけこさしすせそたちつてとなにぬねのはひふへほまみむめもやゆよらりるれろわゐゑをん');
            case LIST_STYLE_TYPE.HIRAGANA_IROHA:
                return createCounterStyleFromSymbols(value, 'いろはにほへとちりぬるをわかよたれそつねならむうゐのおくやまけふこえてあさきゆめみしゑひもせす');
            case LIST_STYLE_TYPE.KANNADA:
                return createCounterStyleFromRange(value, 0xce6, 0xcef, true, defaultSuffix);
            case LIST_STYLE_TYPE.KATAKANA:
                return createCounterStyleFromSymbols(value, 'アイウエオカキクケコサシスセソタチツテトナニヌネノハヒフヘホマミムメモヤユヨラリルレロワヰヱヲン', cjkSuffix);
            case LIST_STYLE_TYPE.KATAKANA_IROHA:
                return createCounterStyleFromSymbols(value, 'イロハニホヘトチリヌルヲワカヨタレソツネナラムウヰノオクヤマケフコエテアサキユメミシヱヒモセス', cjkSuffix);
            case LIST_STYLE_TYPE.LAO:
                return createCounterStyleFromRange(value, 0xed0, 0xed9, true, defaultSuffix);
            case LIST_STYLE_TYPE.MONGOLIAN:
                return createCounterStyleFromRange(value, 0x1810, 0x1819, true, defaultSuffix);
            case LIST_STYLE_TYPE.MYANMAR:
                return createCounterStyleFromRange(value, 0x1040, 0x1049, true, defaultSuffix);
            case LIST_STYLE_TYPE.ORIYA:
                return createCounterStyleFromRange(value, 0xb66, 0xb6f, true, defaultSuffix);
            case LIST_STYLE_TYPE.PERSIAN:
                return createCounterStyleFromRange(value, 0x6f0, 0x6f9, true, defaultSuffix);
            case LIST_STYLE_TYPE.TAMIL:
                return createCounterStyleFromRange(value, 0xbe6, 0xbef, true, defaultSuffix);
            case LIST_STYLE_TYPE.TELUGU:
                return createCounterStyleFromRange(value, 0xc66, 0xc6f, true, defaultSuffix);
            case LIST_STYLE_TYPE.THAI:
                return createCounterStyleFromRange(value, 0xe50, 0xe59, true, defaultSuffix);
            case LIST_STYLE_TYPE.TIBETAN:
                return createCounterStyleFromRange(value, 0xf20, 0xf29, true, defaultSuffix);
            case LIST_STYLE_TYPE.DECIMAL:
            default:
                return createCounterStyleFromRange(value, 48, 57, true, defaultSuffix);
        }
    };

    var IGNORE_ATTRIBUTE = 'data-html2canvas-ignore';
    var DocumentCloner = /** @class */ (function () {
        function DocumentCloner(element, options) {
            this.options = options;
            this.scrolledElements = [];
            this.referenceElement = element;
            this.counters = new CounterState();
            this.quoteDepth = 0;
            if (!element.ownerDocument) {
                throw new Error('Cloned element does not have an owner document');
            }
            this.documentElement = this.cloneNode(element.ownerDocument.documentElement);
        }
        DocumentCloner.prototype.toIFrame = function (ownerDocument, windowSize) {
            var _this = this;
            var iframe = createIFrameContainer(ownerDocument, windowSize);
            if (!iframe.contentWindow) {
                return Promise.reject("Unable to find iframe window");
            }
            var scrollX = ownerDocument.defaultView.pageXOffset;
            var scrollY = ownerDocument.defaultView.pageYOffset;
            var cloneWindow = iframe.contentWindow;
            var documentClone = cloneWindow.document;
            /* Chrome doesn't detect relative background-images assigned in inline <style> sheets when fetched through getComputedStyle
             if window url is about:blank, we can assign the url to current by writing onto the document
             */
            var iframeLoad = iframeLoader(iframe).then(function () { return __awaiter(_this, void 0, void 0, function () {
                var onclone;
                return __generator(this, function (_a) {
                    switch (_a.label) {
                        case 0:
                            this.scrolledElements.forEach(restoreNodeScroll);
                            if (cloneWindow) {
                                cloneWindow.scrollTo(windowSize.left, windowSize.top);
                                if (/(iPad|iPhone|iPod)/g.test(navigator.userAgent) &&
                                    (cloneWindow.scrollY !== windowSize.top || cloneWindow.scrollX !== windowSize.left)) {
                                    documentClone.documentElement.style.top = -windowSize.top + 'px';
                                    documentClone.documentElement.style.left = -windowSize.left + 'px';
                                    documentClone.documentElement.style.position = 'absolute';
                                }
                            }
                            onclone = this.options.onclone;
                            if (typeof this.clonedReferenceElement === 'undefined') {
                                return [2 /*return*/, Promise.reject("Error finding the " + this.referenceElement.nodeName + " in the cloned document")];
                            }
                            if (!(documentClone.fonts && documentClone.fonts.ready)) return [3 /*break*/, 2];
                            return [4 /*yield*/, documentClone.fonts.ready];
                        case 1:
                            _a.sent();
                            _a.label = 2;
                        case 2:
                            if (typeof onclone === 'function') {
                                return [2 /*return*/, Promise.resolve()
                                        .then(function () { return onclone(documentClone); })
                                        .then(function () { return iframe; })];
                            }
                            return [2 /*return*/, iframe];
                    }
                });
            }); });
            documentClone.open();
            documentClone.write(serializeDoctype(document.doctype) + "<html></html>");
            // Chrome scrolls the parent document for some reason after the write to the cloned window???
            restoreOwnerScroll(this.referenceElement.ownerDocument, scrollX, scrollY);
            documentClone.replaceChild(documentClone.adoptNode(this.documentElement), documentClone.documentElement);
            documentClone.close();
            return iframeLoad;
        };
        DocumentCloner.prototype.createElementClone = function (node) {
            if (isCanvasElement(node)) {
                return this.createCanvasClone(node);
            }
            /*
            if (isIFrameElement(node)) {
                return this.createIFrameClone(node);
            }
    */
            if (isStyleElement(node)) {
                return this.createStyleClone(node);
            }
            return node.cloneNode(false);
        };
        DocumentCloner.prototype.createStyleClone = function (node) {
            try {
                var sheet = node.sheet;
                if (sheet && sheet.cssRules) {
                    var css = [].slice.call(sheet.cssRules, 0).reduce(function (css, rule) {
                        if (rule && typeof rule.cssText === 'string') {
                            return css + rule.cssText;
                        }
                        return css;
                    }, '');
                    var style = node.cloneNode(false);
                    style.textContent = css;
                    return style;
                }
            }
            catch (e) {
                // accessing node.sheet.cssRules throws a DOMException
                Logger.getInstance(this.options.id).error('Unable to access cssRules property', e);
                if (e.name !== 'SecurityError') {
                    throw e;
                }
            }
            return node.cloneNode(false);
        };
        DocumentCloner.prototype.createCanvasClone = function (canvas) {
            if (this.options.inlineImages && canvas.ownerDocument) {
                var img = canvas.ownerDocument.createElement('img');
                try {
                    img.src = canvas.toDataURL();
                    return img;
                }
                catch (e) {
                    Logger.getInstance(this.options.id).info("Unable to clone canvas contents, canvas is tainted");
                }
            }
            var clonedCanvas = canvas.cloneNode(false);
            try {
                clonedCanvas.width = canvas.width;
                clonedCanvas.height = canvas.height;
                var ctx = canvas.getContext('2d');
                var clonedCtx = clonedCanvas.getContext('2d');
                if (clonedCtx) {
                    if (ctx) {
                        clonedCtx.putImageData(ctx.getImageData(0, 0, canvas.width, canvas.height), 0, 0);
                    }
                    else {
                        clonedCtx.drawImage(canvas, 0, 0);
                    }
                }
                return clonedCanvas;
            }
            catch (e) { }
            return clonedCanvas;
        };
        /*
        createIFrameClone(iframe: HTMLIFrameElement) {
            const tempIframe = <HTMLIFrameElement>iframe.cloneNode(false);
            const iframeKey = generateIframeKey();
            tempIframe.setAttribute('data-html2canvas-internal-iframe-key', iframeKey);

            const {width, height} = parseBounds(iframe);

            this.resourceLoader.cache[iframeKey] = getIframeDocumentElement(iframe, this.options)
                .then(documentElement => {
                    return this.renderer(
                        documentElement,
                        {
                            allowTaint: this.options.allowTaint,
                            backgroundColor: '#ffffff',
                            canvas: null,
                            imageTimeout: this.options.imageTimeout,
                            logging: this.options.logging,
                            proxy: this.options.proxy,
                            removeContainer: this.options.removeContainer,
                            scale: this.options.scale,
                            foreignObjectRendering: this.options.foreignObjectRendering,
                            useCORS: this.options.useCORS,
                            target: new CanvasRenderer(),
                            width,
                            height,
                            x: 0,
                            y: 0,
                            windowWidth: documentElement.ownerDocument.defaultView.innerWidth,
                            windowHeight: documentElement.ownerDocument.defaultView.innerHeight,
                            scrollX: documentElement.ownerDocument.defaultView.pageXOffset,
                            scrollY: documentElement.ownerDocument.defaultView.pageYOffset
                        },
                    );
                })
                .then(
                    (canvas: HTMLCanvasElement) =>
                        new Promise((resolve, reject) => {
                            const iframeCanvas = document.createElement('img');
                            iframeCanvas.onload = () => resolve(canvas);
                            iframeCanvas.onerror = (event) => {
                                // Empty iframes may result in empty "data:," URLs, which are invalid from the <img>'s point of view
                                // and instead of `onload` cause `onerror` and unhandled rejection warnings
                                // https://github.com/niklasvh/html2canvas/issues/1502
                                iframeCanvas.src == 'data:,' ? resolve(canvas) : reject(event);
                            };
                            iframeCanvas.src = canvas.toDataURL();
                            if (tempIframe.parentNode && iframe.ownerDocument && iframe.ownerDocument.defaultView) {
                                tempIframe.parentNode.replaceChild(
                                    copyCSSStyles(
                                        iframe.ownerDocument.defaultView.getComputedStyle(iframe),
                                        iframeCanvas
                                    ),
                                    tempIframe
                                );
                            }
                        })
                );
            return tempIframe;
        }
    */
        DocumentCloner.prototype.cloneNode = function (node) {
            if (isTextNode(node)) {
                return document.createTextNode(node.data);
            }
            if (!node.ownerDocument) {
                return node.cloneNode(false);
            }
            var window = node.ownerDocument.defaultView;
            if (isHTMLElementNode(node) && window) {
                var clone = this.createElementClone(node);
                var style = window.getComputedStyle(node);
                var styleBefore = window.getComputedStyle(node, ':before');
                var styleAfter = window.getComputedStyle(node, ':after');
                if (this.referenceElement === node) {
                    this.clonedReferenceElement = clone;
                }
                if (isBodyElement(clone)) {
                    createPseudoHideStyles(clone);
                }
                var counters = this.counters.parse(new CSSParsedCounterDeclaration(style));
                var before = this.resolvePseudoContent(node, clone, styleBefore, PseudoElementType.BEFORE);
                for (var child = node.firstChild; child; child = child.nextSibling) {
                    if (!isElementNode(child) ||
                        (!isScriptElement(child) &&
                            !child.hasAttribute(IGNORE_ATTRIBUTE) &&
                            (typeof this.options.ignoreElements !== 'function' || !this.options.ignoreElements(child)))) {
                        if (!this.options.copyStyles || !isElementNode(child) || !isStyleElement(child)) {
                            clone.appendChild(this.cloneNode(child));
                        }
                    }
                }
                if (before) {
                    clone.insertBefore(before, clone.firstChild);
                }
                var after = this.resolvePseudoContent(node, clone, styleAfter, PseudoElementType.AFTER);
                if (after) {
                    clone.appendChild(after);
                }
                this.counters.pop(counters);
                if (style && this.options.copyStyles && !isIFrameElement(node)) {
                    copyCSSStyles(style, clone);
                }
                //this.inlineAllImages(clone);
                if (node.scrollTop !== 0 || node.scrollLeft !== 0) {
                    this.scrolledElements.push([clone, node.scrollLeft, node.scrollTop]);
                }
                if ((isTextareaElement(node) || isSelectElement(node)) &&
                    (isTextareaElement(clone) || isSelectElement(clone))) {
                    clone.value = node.value;
                }
                return clone;
            }
            return node.cloneNode(false);
        };
        DocumentCloner.prototype.resolvePseudoContent = function (node, clone, style, pseudoElt) {
            var _this = this;
            if (!style) {
                return;
            }
            var value = style.content;
            var document = clone.ownerDocument;
            if (!document || !value || value === 'none' || value === '-moz-alt-content' || style.display === 'none') {
                return;
            }
            this.counters.parse(new CSSParsedCounterDeclaration(style));
            var declaration = new CSSParsedPseudoDeclaration(style);
            var anonymousReplacedElement = document.createElement('html2canvaspseudoelement');
            copyCSSStyles(style, anonymousReplacedElement);
            declaration.content.forEach(function (token) {
                if (token.type === TokenType.STRING_TOKEN) {
                    anonymousReplacedElement.appendChild(document.createTextNode(token.value));
                }
                else if (token.type === TokenType.URL_TOKEN) {
                    var img = document.createElement('img');
                    img.src = token.value;
                    img.style.opacity = '1';
                    anonymousReplacedElement.appendChild(img);
                }
                else if (token.type === TokenType.FUNCTION) {
                    if (token.name === 'attr') {
                        var attr = token.values.filter(isIdentToken);
                        if (attr.length) {
                            anonymousReplacedElement.appendChild(document.createTextNode(node.getAttribute(attr[0].value) || ''));
                        }
                    }
                    else if (token.name === 'counter') {
                        var _a = token.values.filter(nonFunctionArgSeparator), counter = _a[0], counterStyle = _a[1];
                        if (counter && isIdentToken(counter)) {
                            var counterState = _this.counters.getCounterValue(counter.value);
                            var counterType = counterStyle && isIdentToken(counterStyle)
                                ? listStyleType.parse(counterStyle.value)
                                : LIST_STYLE_TYPE.DECIMAL;
                            anonymousReplacedElement.appendChild(document.createTextNode(createCounterText(counterState, counterType, false)));
                        }
                    }
                    else if (token.name === 'counters') {
                        var _b = token.values.filter(nonFunctionArgSeparator), counter = _b[0], delim = _b[1], counterStyle = _b[2];
                        if (counter && isIdentToken(counter)) {
                            var counterStates = _this.counters.getCounterValues(counter.value);
                            var counterType_1 = counterStyle && isIdentToken(counterStyle)
                                ? listStyleType.parse(counterStyle.value)
                                : LIST_STYLE_TYPE.DECIMAL;
                            var separator = delim && delim.type === TokenType.STRING_TOKEN ? delim.value : '';
                            var text = counterStates
                                .map(function (value) { return createCounterText(value, counterType_1, false); })
                                .join(separator);
                            anonymousReplacedElement.appendChild(document.createTextNode(text));
                        }
                    }
                }
                else if (token.type === TokenType.IDENT_TOKEN) {
                    switch (token.value) {
                        case 'open-quote':
                            anonymousReplacedElement.appendChild(document.createTextNode(getQuote(declaration.quotes, _this.quoteDepth++, true)));
                            break;
                        case 'close-quote':
                            anonymousReplacedElement.appendChild(document.createTextNode(getQuote(declaration.quotes, --_this.quoteDepth, false)));
                            break;
                        default:
                            // safari doesn't parse string tokens correctly because of lack of quotes
                            anonymousReplacedElement.appendChild(document.createTextNode(token.value));
                    }
                }
            });
            anonymousReplacedElement.className = PSEUDO_HIDE_ELEMENT_CLASS_BEFORE + " " + PSEUDO_HIDE_ELEMENT_CLASS_AFTER;
            var newClassName = pseudoElt === PseudoElementType.BEFORE
                ? " " + PSEUDO_HIDE_ELEMENT_CLASS_BEFORE
                : " " + PSEUDO_HIDE_ELEMENT_CLASS_AFTER;
            if (isSVGElementNode(clone)) {
                clone.className.baseValue += newClassName;
            }
            else {
                clone.className += newClassName;
            }
            return anonymousReplacedElement;
        };
        DocumentCloner.destroy = function (container) {
            if (container.parentNode) {
                container.parentNode.removeChild(container);
                return true;
            }
            return false;
        };
        return DocumentCloner;
    }());
    var PseudoElementType;
    (function (PseudoElementType) {
        PseudoElementType[PseudoElementType["BEFORE"] = 0] = "BEFORE";
        PseudoElementType[PseudoElementType["AFTER"] = 1] = "AFTER";
    })(PseudoElementType || (PseudoElementType = {}));
    var createIFrameContainer = function (ownerDocument, bounds) {
        var cloneIframeContainer = ownerDocument.createElement('iframe');
        cloneIframeContainer.className = 'html2canvas-container';
        cloneIframeContainer.style.visibility = 'hidden';
        cloneIframeContainer.style.position = 'fixed';
        cloneIframeContainer.style.left = '-10000px';
        cloneIframeContainer.style.top = '0px';
        cloneIframeContainer.style.border = '0';
        cloneIframeContainer.width = bounds.width.toString();
        cloneIframeContainer.height = bounds.height.toString();
        cloneIframeContainer.scrolling = 'no'; // ios won't scroll without it
        cloneIframeContainer.setAttribute(IGNORE_ATTRIBUTE, 'true');
        ownerDocument.body.appendChild(cloneIframeContainer);
        return cloneIframeContainer;
    };
    var iframeLoader = function (iframe) {
        return new Promise(function (resolve, reject) {
            var cloneWindow = iframe.contentWindow;
            if (!cloneWindow) {
                return reject("No window assigned for iframe");
            }
            var documentClone = cloneWindow.document;
            cloneWindow.onload = iframe.onload = documentClone.onreadystatechange = function () {
                cloneWindow.onload = iframe.onload = documentClone.onreadystatechange = null;
                var interval = setInterval(function () {
                    if (documentClone.body.childNodes.length > 0 && documentClone.readyState === 'complete') {
                        clearInterval(interval);
                        resolve(iframe);
                    }
                }, 50);
            };
        });
    };
    var copyCSSStyles = function (style, target) {
        // Edge does not provide value for cssText
        for (var i = style.length - 1; i >= 0; i--) {
            var property = style.item(i);
            // Safari shows pseudoelements if content is set
            if (property !== 'content') {
                target.style.setProperty(property, style.getPropertyValue(property));
            }
        }
        return target;
    };
    var serializeDoctype = function (doctype) {
        var str = '';
        if (doctype) {
            str += '<!DOCTYPE ';
            if (doctype.name) {
                str += doctype.name;
            }
            if (doctype.internalSubset) {
                str += doctype.internalSubset;
            }
            if (doctype.publicId) {
                str += "\"" + doctype.publicId + "\"";
            }
            if (doctype.systemId) {
                str += "\"" + doctype.systemId + "\"";
            }
            str += '>';
        }
        return str;
    };
    var restoreOwnerScroll = function (ownerDocument, x, y) {
        if (ownerDocument &&
            ownerDocument.defaultView &&
            (x !== ownerDocument.defaultView.pageXOffset || y !== ownerDocument.defaultView.pageYOffset)) {
            ownerDocument.defaultView.scrollTo(x, y);
        }
    };
    var restoreNodeScroll = function (_a) {
        var element = _a[0], x = _a[1], y = _a[2];
        element.scrollLeft = x;
        element.scrollTop = y;
    };
    var PSEUDO_BEFORE = ':before';
    var PSEUDO_AFTER = ':after';
    var PSEUDO_HIDE_ELEMENT_CLASS_BEFORE = '___html2canvas___pseudoelement_before';
    var PSEUDO_HIDE_ELEMENT_CLASS_AFTER = '___html2canvas___pseudoelement_after';
    var PSEUDO_HIDE_ELEMENT_STYLE = "{\n    content: \"\" !important;\n    display: none !important;\n}";
    var createPseudoHideStyles = function (body) {
        createStyles(body, "." + PSEUDO_HIDE_ELEMENT_CLASS_BEFORE + PSEUDO_BEFORE + PSEUDO_HIDE_ELEMENT_STYLE + "\n         ." + PSEUDO_HIDE_ELEMENT_CLASS_AFTER + PSEUDO_AFTER + PSEUDO_HIDE_ELEMENT_STYLE);
    };
    var createStyles = function (body, styles) {
        var document = body.ownerDocument;
        if (document) {
            var style = document.createElement('style');
            style.textContent = styles;
            body.appendChild(style);
        }
    };

    var PathType;
    (function (PathType) {
        PathType[PathType["VECTOR"] = 0] = "VECTOR";
        PathType[PathType["BEZIER_CURVE"] = 1] = "BEZIER_CURVE";
    })(PathType || (PathType = {}));
    var equalPath = function (a, b) {
        if (a.length === b.length) {
            return a.some(function (v, i) { return v === b[i]; });
        }
        return false;
    };
    var transformPath = function (path, deltaX, deltaY, deltaW, deltaH) {
        return path.map(function (point, index) {
            switch (index) {
                case 0:
                    return point.add(deltaX, deltaY);
                case 1:
                    return point.add(deltaX + deltaW, deltaY);
                case 2:
                    return point.add(deltaX + deltaW, deltaY + deltaH);
                case 3:
                    return point.add(deltaX, deltaY + deltaH);
            }
            return point;
        });
    };

    var Vector = /** @class */ (function () {
        function Vector(x, y) {
            this.type = PathType.VECTOR;
            this.x = x;
            this.y = y;
        }
        Vector.prototype.add = function (deltaX, deltaY) {
            return new Vector(this.x + deltaX, this.y + deltaY);
        };
        return Vector;
    }());

    var lerp = function (a, b, t) {
        return new Vector(a.x + (b.x - a.x) * t, a.y + (b.y - a.y) * t);
    };
    var BezierCurve = /** @class */ (function () {
        function BezierCurve(start, startControl, endControl, end) {
            this.type = PathType.BEZIER_CURVE;
            this.start = start;
            this.startControl = startControl;
            this.endControl = endControl;
            this.end = end;
        }
        BezierCurve.prototype.subdivide = function (t, firstHalf) {
            var ab = lerp(this.start, this.startControl, t);
            var bc = lerp(this.startControl, this.endControl, t);
            var cd = lerp(this.endControl, this.end, t);
            var abbc = lerp(ab, bc, t);
            var bccd = lerp(bc, cd, t);
            var dest = lerp(abbc, bccd, t);
            return firstHalf ? new BezierCurve(this.start, ab, abbc, dest) : new BezierCurve(dest, bccd, cd, this.end);
        };
        BezierCurve.prototype.add = function (deltaX, deltaY) {
            return new BezierCurve(this.start.add(deltaX, deltaY), this.startControl.add(deltaX, deltaY), this.endControl.add(deltaX, deltaY), this.end.add(deltaX, deltaY));
        };
        BezierCurve.prototype.reverse = function () {
            return new BezierCurve(this.end, this.endControl, this.startControl, this.start);
        };
        return BezierCurve;
    }());
    var isBezierCurve = function (path) { return path.type === PathType.BEZIER_CURVE; };

    var BoundCurves = /** @class */ (function () {
        function BoundCurves(element) {
            var styles = element.styles;
            var bounds = element.bounds;
            var _a = getAbsoluteValueForTuple(styles.borderTopLeftRadius, bounds.width, bounds.height), tlh = _a[0], tlv = _a[1];
            var _b = getAbsoluteValueForTuple(styles.borderTopRightRadius, bounds.width, bounds.height), trh = _b[0], trv = _b[1];
            var _c = getAbsoluteValueForTuple(styles.borderBottomRightRadius, bounds.width, bounds.height), brh = _c[0], brv = _c[1];
            var _d = getAbsoluteValueForTuple(styles.borderBottomLeftRadius, bounds.width, bounds.height), blh = _d[0], blv = _d[1];
            var factors = [];
            factors.push((tlh + trh) / bounds.width);
            factors.push((blh + brh) / bounds.width);
            factors.push((tlv + blv) / bounds.height);
            factors.push((trv + brv) / bounds.height);
            var maxFactor = Math.max.apply(Math, factors);
            if (maxFactor > 1) {
                tlh /= maxFactor;
                tlv /= maxFactor;
                trh /= maxFactor;
                trv /= maxFactor;
                brh /= maxFactor;
                brv /= maxFactor;
                blh /= maxFactor;
                blv /= maxFactor;
            }
            var topWidth = bounds.width - trh;
            var rightHeight = bounds.height - brv;
            var bottomWidth = bounds.width - brh;
            var leftHeight = bounds.height - blv;
            var borderTopWidth = styles.borderTopWidth;
            var borderRightWidth = styles.borderRightWidth;
            var borderBottomWidth = styles.borderBottomWidth;
            var borderLeftWidth = styles.borderLeftWidth;
            var paddingTop = getAbsoluteValue(styles.paddingTop, element.bounds.width);
            var paddingRight = getAbsoluteValue(styles.paddingRight, element.bounds.width);
            var paddingBottom = getAbsoluteValue(styles.paddingBottom, element.bounds.width);
            var paddingLeft = getAbsoluteValue(styles.paddingLeft, element.bounds.width);
            this.topLeftBorderBox =
                tlh > 0 || tlv > 0
                    ? getCurvePoints(bounds.left, bounds.top, tlh, tlv, CORNER.TOP_LEFT)
                    : new Vector(bounds.left, bounds.top);
            this.topRightBorderBox =
                trh > 0 || trv > 0
                    ? getCurvePoints(bounds.left + topWidth, bounds.top, trh, trv, CORNER.TOP_RIGHT)
                    : new Vector(bounds.left + bounds.width, bounds.top);
            this.bottomRightBorderBox =
                brh > 0 || brv > 0
                    ? getCurvePoints(bounds.left + bottomWidth, bounds.top + rightHeight, brh, brv, CORNER.BOTTOM_RIGHT)
                    : new Vector(bounds.left + bounds.width, bounds.top + bounds.height);
            this.bottomLeftBorderBox =
                blh > 0 || blv > 0
                    ? getCurvePoints(bounds.left, bounds.top + leftHeight, blh, blv, CORNER.BOTTOM_LEFT)
                    : new Vector(bounds.left, bounds.top + bounds.height);
            this.topLeftPaddingBox =
                tlh > 0 || tlv > 0
                    ? getCurvePoints(bounds.left + borderLeftWidth, bounds.top + borderTopWidth, Math.max(0, tlh - borderLeftWidth), Math.max(0, tlv - borderTopWidth), CORNER.TOP_LEFT)
                    : new Vector(bounds.left + borderLeftWidth, bounds.top + borderTopWidth);
            this.topRightPaddingBox =
                trh > 0 || trv > 0
                    ? getCurvePoints(bounds.left + Math.min(topWidth, bounds.width + borderLeftWidth), bounds.top + borderTopWidth, topWidth > bounds.width + borderLeftWidth ? 0 : trh - borderLeftWidth, trv - borderTopWidth, CORNER.TOP_RIGHT)
                    : new Vector(bounds.left + bounds.width - borderRightWidth, bounds.top + borderTopWidth);
            this.bottomRightPaddingBox =
                brh > 0 || brv > 0
                    ? getCurvePoints(bounds.left + Math.min(bottomWidth, bounds.width - borderLeftWidth), bounds.top + Math.min(rightHeight, bounds.height + borderTopWidth), Math.max(0, brh - borderRightWidth), brv - borderBottomWidth, CORNER.BOTTOM_RIGHT)
                    : new Vector(bounds.left + bounds.width - borderRightWidth, bounds.top + bounds.height - borderBottomWidth);
            this.bottomLeftPaddingBox =
                blh > 0 || blv > 0
                    ? getCurvePoints(bounds.left + borderLeftWidth, bounds.top + leftHeight, Math.max(0, blh - borderLeftWidth), blv - borderBottomWidth, CORNER.BOTTOM_LEFT)
                    : new Vector(bounds.left + borderLeftWidth, bounds.top + bounds.height - borderBottomWidth);
            this.topLeftContentBox =
                tlh > 0 || tlv > 0
                    ? getCurvePoints(bounds.left + borderLeftWidth + paddingLeft, bounds.top + borderTopWidth + paddingTop, Math.max(0, tlh - (borderLeftWidth + paddingLeft)), Math.max(0, tlv - (borderTopWidth + paddingTop)), CORNER.TOP_LEFT)
                    : new Vector(bounds.left + borderLeftWidth + paddingLeft, bounds.top + borderTopWidth + paddingTop);
            this.topRightContentBox =
                trh > 0 || trv > 0
                    ? getCurvePoints(bounds.left + Math.min(topWidth, bounds.width + borderLeftWidth + paddingLeft), bounds.top + borderTopWidth + paddingTop, topWidth > bounds.width + borderLeftWidth + paddingLeft ? 0 : trh - borderLeftWidth + paddingLeft, trv - (borderTopWidth + paddingTop), CORNER.TOP_RIGHT)
                    : new Vector(bounds.left + bounds.width - (borderRightWidth + paddingRight), bounds.top + borderTopWidth + paddingTop);
            this.bottomRightContentBox =
                brh > 0 || brv > 0
                    ? getCurvePoints(bounds.left + Math.min(bottomWidth, bounds.width - (borderLeftWidth + paddingLeft)), bounds.top + Math.min(rightHeight, bounds.height + borderTopWidth + paddingTop), Math.max(0, brh - (borderRightWidth + paddingRight)), brv - (borderBottomWidth + paddingBottom), CORNER.BOTTOM_RIGHT)
                    : new Vector(bounds.left + bounds.width - (borderRightWidth + paddingRight), bounds.top + bounds.height - (borderBottomWidth + paddingBottom));
            this.bottomLeftContentBox =
                blh > 0 || blv > 0
                    ? getCurvePoints(bounds.left + borderLeftWidth + paddingLeft, bounds.top + leftHeight, Math.max(0, blh - (borderLeftWidth + paddingLeft)), blv - (borderBottomWidth + paddingBottom), CORNER.BOTTOM_LEFT)
                    : new Vector(bounds.left + borderLeftWidth + paddingLeft, bounds.top + bounds.height - (borderBottomWidth + paddingBottom));
        }
        return BoundCurves;
    }());
    var CORNER;
    (function (CORNER) {
        CORNER[CORNER["TOP_LEFT"] = 0] = "TOP_LEFT";
        CORNER[CORNER["TOP_RIGHT"] = 1] = "TOP_RIGHT";
        CORNER[CORNER["BOTTOM_RIGHT"] = 2] = "BOTTOM_RIGHT";
        CORNER[CORNER["BOTTOM_LEFT"] = 3] = "BOTTOM_LEFT";
    })(CORNER || (CORNER = {}));
    var getCurvePoints = function (x, y, r1, r2, position) {
        var kappa = 4 * ((Math.sqrt(2) - 1) / 3);
        var ox = r1 * kappa; // control point offset horizontal
        var oy = r2 * kappa; // control point offset vertical
        var xm = x + r1; // x-middle
        var ym = y + r2; // y-middle
        switch (position) {
            case CORNER.TOP_LEFT:
                return new BezierCurve(new Vector(x, ym), new Vector(x, ym - oy), new Vector(xm - ox, y), new Vector(xm, y));
            case CORNER.TOP_RIGHT:
                return new BezierCurve(new Vector(x, y), new Vector(x + ox, y), new Vector(xm, ym - oy), new Vector(xm, ym));
            case CORNER.BOTTOM_RIGHT:
                return new BezierCurve(new Vector(xm, y), new Vector(xm, y + oy), new Vector(x + ox, ym), new Vector(x, ym));
            case CORNER.BOTTOM_LEFT:
            default:
                return new BezierCurve(new Vector(xm, ym), new Vector(xm - ox, ym), new Vector(x, y + oy), new Vector(x, y));
        }
    };
    var calculateBorderBoxPath = function (curves) {
        return [curves.topLeftBorderBox, curves.topRightBorderBox, curves.bottomRightBorderBox, curves.bottomLeftBorderBox];
    };
    var calculateContentBoxPath = function (curves) {
        return [
            curves.topLeftContentBox,
            curves.topRightContentBox,
            curves.bottomRightContentBox,
            curves.bottomLeftContentBox
        ];
    };
    var calculatePaddingBoxPath = function (curves) {
        return [
            curves.topLeftPaddingBox,
            curves.topRightPaddingBox,
            curves.bottomRightPaddingBox,
            curves.bottomLeftPaddingBox
        ];
    };

    var TransformEffect = /** @class */ (function () {
        function TransformEffect(offsetX, offsetY, matrix) {
            this.type = 0 /* TRANSFORM */;
            this.offsetX = offsetX;
            this.offsetY = offsetY;
            this.matrix = matrix;
            this.target = 2 /* BACKGROUND_BORDERS */ | 4 /* CONTENT */;
        }
        return TransformEffect;
    }());
    var ClipEffect = /** @class */ (function () {
        function ClipEffect(path, target) {
            this.type = 1 /* CLIP */;
            this.target = target;
            this.path = path;
        }
        return ClipEffect;
    }());
    var isTransformEffect = function (effect) {
        return effect.type === 0 /* TRANSFORM */;
    };
    var isClipEffect = function (effect) { return effect.type === 1 /* CLIP */; };

    var StackingContext = /** @class */ (function () {
        function StackingContext(container) {
            this.element = container;
            this.inlineLevel = [];
            this.nonInlineLevel = [];
            this.negativeZIndex = [];
            this.zeroOrAutoZIndexOrTransformedOrOpacity = [];
            this.positiveZIndex = [];
            this.nonPositionedFloats = [];
            this.nonPositionedInlineLevel = [];
        }
        return StackingContext;
    }());
    var ElementPaint = /** @class */ (function () {
        function ElementPaint(element, parentStack) {
            this.container = element;
            this.effects = parentStack.slice(0);
            this.curves = new BoundCurves(element);
            if (element.styles.transform !== null) {
                var offsetX = element.bounds.left + element.styles.transformOrigin[0].number;
                var offsetY = element.bounds.top + element.styles.transformOrigin[1].number;
                var matrix = element.styles.transform;
                this.effects.push(new TransformEffect(offsetX, offsetY, matrix));
            }
            if (element.styles.overflowX !== OVERFLOW.VISIBLE) {
                var borderBox = calculateBorderBoxPath(this.curves);
                var paddingBox = calculatePaddingBoxPath(this.curves);
                if (equalPath(borderBox, paddingBox)) {
                    this.effects.push(new ClipEffect(borderBox, 2 /* BACKGROUND_BORDERS */ | 4 /* CONTENT */));
                }
                else {
                    this.effects.push(new ClipEffect(borderBox, 2 /* BACKGROUND_BORDERS */));
                    this.effects.push(new ClipEffect(paddingBox, 4 /* CONTENT */));
                }
            }
        }
        ElementPaint.prototype.getParentEffects = function () {
            var effects = this.effects.slice(0);
            if (this.container.styles.overflowX !== OVERFLOW.VISIBLE) {
                var borderBox = calculateBorderBoxPath(this.curves);
                var paddingBox = calculatePaddingBoxPath(this.curves);
                if (!equalPath(borderBox, paddingBox)) {
                    effects.push(new ClipEffect(paddingBox, 2 /* BACKGROUND_BORDERS */ | 4 /* CONTENT */));
                }
            }
            return effects;
        };
        return ElementPaint;
    }());
    var parseStackTree = function (parent, stackingContext, realStackingContext, listItems) {
        parent.container.elements.forEach(function (child) {
            var treatAsRealStackingContext = contains(child.flags, 4 /* CREATES_REAL_STACKING_CONTEXT */);
            var createsStackingContext = contains(child.flags, 2 /* CREATES_STACKING_CONTEXT */);
            var paintContainer = new ElementPaint(child, parent.getParentEffects());
            if (contains(child.styles.display, 2048 /* LIST_ITEM */)) {
                listItems.push(paintContainer);
            }
            var listOwnerItems = contains(child.flags, 8 /* IS_LIST_OWNER */) ? [] : listItems;
            if (treatAsRealStackingContext || createsStackingContext) {
                var parentStack = treatAsRealStackingContext || child.styles.isPositioned() ? realStackingContext : stackingContext;
                var stack = new StackingContext(paintContainer);
                if (child.styles.isPositioned() || child.styles.opacity < 1 || child.styles.isTransformed()) {
                    var order_1 = child.styles.zIndex.order;
                    if (order_1 < 0) {
                        var index_1 = 0;
                        parentStack.negativeZIndex.some(function (current, i) {
                            if (order_1 > current.element.container.styles.zIndex.order) {
                                index_1 = i;
                                return false;
                            }
                            else if (index_1 > 0) {
                                return true;
                            }
                            return false;
                        });
                        parentStack.negativeZIndex.splice(index_1, 0, stack);
                    }
                    else if (order_1 > 0) {
                        var index_2 = 0;
                        parentStack.positiveZIndex.some(function (current, i) {
                            if (order_1 > current.element.container.styles.zIndex.order) {
                                index_2 = i + 1;
                                return false;
                            }
                            else if (index_2 > 0) {
                                return true;
                            }
                            return false;
                        });
                        parentStack.positiveZIndex.splice(index_2, 0, stack);
                    }
                    else {
                        parentStack.zeroOrAutoZIndexOrTransformedOrOpacity.push(stack);
                    }
                }
                else {
                    if (child.styles.isFloating()) {
                        parentStack.nonPositionedFloats.push(stack);
                    }
                    else {
                        parentStack.nonPositionedInlineLevel.push(stack);
                    }
                }
                parseStackTree(paintContainer, stack, treatAsRealStackingContext ? stack : realStackingContext, listOwnerItems);
            }
            else {
                if (child.styles.isInlineLevel()) {
                    stackingContext.inlineLevel.push(paintContainer);
                }
                else {
                    stackingContext.nonInlineLevel.push(paintContainer);
                }
                parseStackTree(paintContainer, stackingContext, realStackingContext, listOwnerItems);
            }
            if (contains(child.flags, 8 /* IS_LIST_OWNER */)) {
                processListItems(child, listOwnerItems);
            }
        });
    };
    var processListItems = function (owner, elements) {
        var numbering = owner instanceof OLElementContainer ? owner.start : 1;
        var reversed = owner instanceof OLElementContainer ? owner.reversed : false;
        for (var i = 0; i < elements.length; i++) {
            var item = elements[i];
            if (item.container instanceof LIElementContainer &&
                typeof item.container.value === 'number' &&
                item.container.value !== 0) {
                numbering = item.container.value;
            }
            item.listValue = createCounterText(numbering, item.container.styles.listStyleType, true);
            numbering += reversed ? -1 : 1;
        }
    };
    var parseStackingContexts = function (container) {
        var paintContainer = new ElementPaint(container, []);
        var root = new StackingContext(paintContainer);
        var listItems = [];
        parseStackTree(paintContainer, root, root, listItems);
        processListItems(paintContainer.container, listItems);
        return root;
    };

    var parsePathForBorder = function (curves, borderSide) {
        switch (borderSide) {
            case 0:
                return createPathFromCurves(curves.topLeftBorderBox, curves.topLeftPaddingBox, curves.topRightBorderBox, curves.topRightPaddingBox);
            case 1:
                return createPathFromCurves(curves.topRightBorderBox, curves.topRightPaddingBox, curves.bottomRightBorderBox, curves.bottomRightPaddingBox);
            case 2:
                return createPathFromCurves(curves.bottomRightBorderBox, curves.bottomRightPaddingBox, curves.bottomLeftBorderBox, curves.bottomLeftPaddingBox);
            case 3:
            default:
                return createPathFromCurves(curves.bottomLeftBorderBox, curves.bottomLeftPaddingBox, curves.topLeftBorderBox, curves.topLeftPaddingBox);
        }
    };
    var createPathFromCurves = function (outer1, inner1, outer2, inner2) {
        var path = [];
        if (isBezierCurve(outer1)) {
            path.push(outer1.subdivide(0.5, false));
        }
        else {
            path.push(outer1);
        }
        if (isBezierCurve(outer2)) {
            path.push(outer2.subdivide(0.5, true));
        }
        else {
            path.push(outer2);
        }
        if (isBezierCurve(inner2)) {
            path.push(inner2.subdivide(0.5, true).reverse());
        }
        else {
            path.push(inner2);
        }
        if (isBezierCurve(inner1)) {
            path.push(inner1.subdivide(0.5, false).reverse());
        }
        else {
            path.push(inner1);
        }
        return path;
    };

    var paddingBox = function (element) {
        var bounds = element.bounds;
        var styles = element.styles;
        return bounds.add(styles.borderLeftWidth, styles.borderTopWidth, -(styles.borderRightWidth + styles.borderLeftWidth), -(styles.borderTopWidth + styles.borderBottomWidth));
    };
    var contentBox = function (element) {
        var styles = element.styles;
        var bounds = element.bounds;
        var paddingLeft = getAbsoluteValue(styles.paddingLeft, bounds.width);
        var paddingRight = getAbsoluteValue(styles.paddingRight, bounds.width);
        var paddingTop = getAbsoluteValue(styles.paddingTop, bounds.width);
        var paddingBottom = getAbsoluteValue(styles.paddingBottom, bounds.width);
        return bounds.add(paddingLeft + styles.borderLeftWidth, paddingTop + styles.borderTopWidth, -(styles.borderRightWidth + styles.borderLeftWidth + paddingLeft + paddingRight), -(styles.borderTopWidth + styles.borderBottomWidth + paddingTop + paddingBottom));
    };

    var calculateBackgroundPositioningArea = function (backgroundOrigin, element) {
        if (backgroundOrigin === 0 /* BORDER_BOX */) {
            return element.bounds;
        }
        if (backgroundOrigin === 2 /* CONTENT_BOX */) {
            return contentBox(element);
        }
        return paddingBox(element);
    };
    var calculateBackgroundPaintingArea = function (backgroundClip, element) {
        if (backgroundClip === BACKGROUND_CLIP.BORDER_BOX) {
            return element.bounds;
        }
        if (backgroundClip === BACKGROUND_CLIP.CONTENT_BOX) {
            return contentBox(element);
        }
        return paddingBox(element);
    };
    var calculateBackgroundRendering = function (container, index, intrinsicSize) {
        var backgroundPositioningArea = calculateBackgroundPositioningArea(getBackgroundValueForIndex(container.styles.backgroundOrigin, index), container);
        var backgroundPaintingArea = calculateBackgroundPaintingArea(getBackgroundValueForIndex(container.styles.backgroundClip, index), container);
        var backgroundImageSize = calculateBackgroundSize(getBackgroundValueForIndex(container.styles.backgroundSize, index), intrinsicSize, backgroundPositioningArea);
        var sizeWidth = backgroundImageSize[0], sizeHeight = backgroundImageSize[1];
        var position = getAbsoluteValueForTuple(getBackgroundValueForIndex(container.styles.backgroundPosition, index), backgroundPositioningArea.width - sizeWidth, backgroundPositioningArea.height - sizeHeight);
        var path = calculateBackgroundRepeatPath(getBackgroundValueForIndex(container.styles.backgroundRepeat, index), position, backgroundImageSize, backgroundPositioningArea, backgroundPaintingArea);
        var offsetX = Math.round(backgroundPositioningArea.left + position[0]);
        var offsetY = Math.round(backgroundPositioningArea.top + position[1]);
        return [path, offsetX, offsetY, sizeWidth, sizeHeight];
    };
    var isAuto = function (token) { return isIdentToken(token) && token.value === BACKGROUND_SIZE.AUTO; };
    var hasIntrinsicValue = function (value) { return typeof value === 'number'; };
    var calculateBackgroundSize = function (size, _a, bounds) {
        var intrinsicWidth = _a[0], intrinsicHeight = _a[1], intrinsicProportion = _a[2];
        var first = size[0], second = size[1];
        if (isLengthPercentage(first) && second && isLengthPercentage(second)) {
            return [getAbsoluteValue(first, bounds.width), getAbsoluteValue(second, bounds.height)];
        }
        var hasIntrinsicProportion = hasIntrinsicValue(intrinsicProportion);
        if (isIdentToken(first) && (first.value === BACKGROUND_SIZE.CONTAIN || first.value === BACKGROUND_SIZE.COVER)) {
            if (hasIntrinsicValue(intrinsicProportion)) {
                var targetRatio = bounds.width / bounds.height;
                return targetRatio < intrinsicProportion !== (first.value === BACKGROUND_SIZE.COVER)
                    ? [bounds.width, bounds.width / intrinsicProportion]
                    : [bounds.height * intrinsicProportion, bounds.height];
            }
            return [bounds.width, bounds.height];
        }
        var hasIntrinsicWidth = hasIntrinsicValue(intrinsicWidth);
        var hasIntrinsicHeight = hasIntrinsicValue(intrinsicHeight);
        var hasIntrinsicDimensions = hasIntrinsicWidth || hasIntrinsicHeight;
        // If the background-size is auto or auto auto:
        if (isAuto(first) && (!second || isAuto(second))) {
            // If the image has both horizontal and vertical intrinsic dimensions, it's rendered at that size.
            if (hasIntrinsicWidth && hasIntrinsicHeight) {
                return [intrinsicWidth, intrinsicHeight];
            }
            // If the image has no intrinsic dimensions and has no intrinsic proportions,
            // it's rendered at the size of the background positioning area.
            if (!hasIntrinsicProportion && !hasIntrinsicDimensions) {
                return [bounds.width, bounds.height];
            }
            // TODO If the image has no intrinsic dimensions but has intrinsic proportions, it's rendered as if contain had been specified instead.
            // If the image has only one intrinsic dimension and has intrinsic proportions, it's rendered at the size corresponding to that one dimension.
            // The other dimension is computed using the specified dimension and the intrinsic proportions.
            if (hasIntrinsicDimensions && hasIntrinsicProportion) {
                var width_1 = hasIntrinsicWidth
                    ? intrinsicWidth
                    : intrinsicHeight * intrinsicProportion;
                var height_1 = hasIntrinsicHeight
                    ? intrinsicHeight
                    : intrinsicWidth / intrinsicProportion;
                return [width_1, height_1];
            }
            // If the image has only one intrinsic dimension but has no intrinsic proportions,
            // it's rendered using the specified dimension and the other dimension of the background positioning area.
            var width_2 = hasIntrinsicWidth ? intrinsicWidth : bounds.width;
            var height_2 = hasIntrinsicHeight ? intrinsicHeight : bounds.height;
            return [width_2, height_2];
        }
        // If the image has intrinsic proportions, it's stretched to the specified dimension.
        // The unspecified dimension is computed using the specified dimension and the intrinsic proportions.
        if (hasIntrinsicProportion) {
            var width_3 = 0;
            var height_3 = 0;
            if (isLengthPercentage(first)) {
                width_3 = getAbsoluteValue(first, bounds.width);
            }
            else if (isLengthPercentage(second)) {
                height_3 = getAbsoluteValue(second, bounds.height);
            }
            if (isAuto(first)) {
                width_3 = height_3 * intrinsicProportion;
            }
            else if (!second || isAuto(second)) {
                height_3 = width_3 / intrinsicProportion;
            }
            return [width_3, height_3];
        }
        // If the image has no intrinsic proportions, it's stretched to the specified dimension.
        // The unspecified dimension is computed using the image's corresponding intrinsic dimension,
        // if there is one. If there is no such intrinsic dimension,
        // it becomes the corresponding dimension of the background positioning area.
        var width = null;
        var height = null;
        if (isLengthPercentage(first)) {
            width = getAbsoluteValue(first, bounds.width);
        }
        else if (second && isLengthPercentage(second)) {
            height = getAbsoluteValue(second, bounds.height);
        }
        if (width !== null && (!second || isAuto(second))) {
            height =
                hasIntrinsicWidth && hasIntrinsicHeight
                    ? (width / intrinsicWidth) * intrinsicHeight
                    : bounds.height;
        }
        if (height !== null && isAuto(first)) {
            width =
                hasIntrinsicWidth && hasIntrinsicHeight
                    ? (height / intrinsicHeight) * intrinsicWidth
                    : bounds.width;
        }
        if (width !== null && height !== null) {
            return [width, height];
        }
        throw new Error("Unable to calculate background-size for element");
    };
    var getBackgroundValueForIndex = function (values, index) {
        var value = values[index];
        if (typeof value === 'undefined') {
            return values[0];
        }
        return value;
    };
    var calculateBackgroundRepeatPath = function (repeat, _a, _b, backgroundPositioningArea, backgroundPaintingArea) {
        var x = _a[0], y = _a[1];
        var width = _b[0], height = _b[1];
        switch (repeat) {
            case BACKGROUND_REPEAT.REPEAT_X:
                return [
                    new Vector(Math.round(backgroundPositioningArea.left), Math.round(backgroundPositioningArea.top + y)),
                    new Vector(Math.round(backgroundPositioningArea.left + backgroundPositioningArea.width), Math.round(backgroundPositioningArea.top + y)),
                    new Vector(Math.round(backgroundPositioningArea.left + backgroundPositioningArea.width), Math.round(height + backgroundPositioningArea.top + y)),
                    new Vector(Math.round(backgroundPositioningArea.left), Math.round(height + backgroundPositioningArea.top + y))
                ];
            case BACKGROUND_REPEAT.REPEAT_Y:
                return [
                    new Vector(Math.round(backgroundPositioningArea.left + x), Math.round(backgroundPositioningArea.top)),
                    new Vector(Math.round(backgroundPositioningArea.left + x + width), Math.round(backgroundPositioningArea.top)),
                    new Vector(Math.round(backgroundPositioningArea.left + x + width), Math.round(backgroundPositioningArea.height + backgroundPositioningArea.top)),
                    new Vector(Math.round(backgroundPositioningArea.left + x), Math.round(backgroundPositioningArea.height + backgroundPositioningArea.top))
                ];
            case BACKGROUND_REPEAT.NO_REPEAT:
                return [
                    new Vector(Math.round(backgroundPositioningArea.left + x), Math.round(backgroundPositioningArea.top + y)),
                    new Vector(Math.round(backgroundPositioningArea.left + x + width), Math.round(backgroundPositioningArea.top + y)),
                    new Vector(Math.round(backgroundPositioningArea.left + x + width), Math.round(backgroundPositioningArea.top + y + height)),
                    new Vector(Math.round(backgroundPositioningArea.left + x), Math.round(backgroundPositioningArea.top + y + height))
                ];
            default:
                return [
                    new Vector(Math.round(backgroundPaintingArea.left), Math.round(backgroundPaintingArea.top)),
                    new Vector(Math.round(backgroundPaintingArea.left + backgroundPaintingArea.width), Math.round(backgroundPaintingArea.top)),
                    new Vector(Math.round(backgroundPaintingArea.left + backgroundPaintingArea.width), Math.round(backgroundPaintingArea.height + backgroundPaintingArea.top)),
                    new Vector(Math.round(backgroundPaintingArea.left), Math.round(backgroundPaintingArea.height + backgroundPaintingArea.top))
                ];
        }
    };

    var SMALL_IMAGE = 'data:image/gif;base64,R0lGODlhAQABAIAAAAAAAP///yH5BAEAAAAALAAAAAABAAEAAAIBRAA7';

    var SAMPLE_TEXT = 'Hidden Text';
    var FontMetrics = /** @class */ (function () {
        function FontMetrics(document) {
            this._data = {};
            this._document = document;
        }
        FontMetrics.prototype.parseMetrics = function (fontFamily, fontSize) {
            var container = this._document.createElement('div');
            var img = this._document.createElement('img');
            var span = this._document.createElement('span');
            var body = this._document.body;
            container.style.visibility = 'hidden';
            container.style.fontFamily = fontFamily;
            container.style.fontSize = fontSize;
            container.style.margin = '0';
            container.style.padding = '0';
            body.appendChild(container);
            img.src = SMALL_IMAGE;
            img.width = 1;
            img.height = 1;
            img.style.margin = '0';
            img.style.padding = '0';
            img.style.verticalAlign = 'baseline';
            span.style.fontFamily = fontFamily;
            span.style.fontSize = fontSize;
            span.style.margin = '0';
            span.style.padding = '0';
            span.appendChild(this._document.createTextNode(SAMPLE_TEXT));
            container.appendChild(span);
            container.appendChild(img);
            var baseline = img.offsetTop - span.offsetTop + 2;
            container.removeChild(span);
            container.appendChild(this._document.createTextNode(SAMPLE_TEXT));
            container.style.lineHeight = 'normal';
            img.style.verticalAlign = 'super';
            var middle = img.offsetTop - container.offsetTop + 2;
            body.removeChild(container);
            return { baseline: baseline, middle: middle };
        };
        FontMetrics.prototype.getMetrics = function (fontFamily, fontSize) {
            var key = fontFamily + " " + fontSize;
            if (typeof this._data[key] === 'undefined') {
                this._data[key] = this.parseMetrics(fontFamily, fontSize);
            }
            return this._data[key];
        };
        return FontMetrics;
    }());

    var MASK_OFFSET = 10000;
    var CanvasRenderer = /** @class */ (function () {
        function CanvasRenderer(options) {
            this._activeEffects = [];
            this.canvas = options.canvas ? options.canvas : document.createElement('canvas');
            this.ctx = this.canvas.getContext('2d');
            this.options = options;
            if (!options.canvas) {
                this.canvas.width = Math.floor(options.width * options.scale);
                this.canvas.height = Math.floor(options.height * options.scale);
                this.canvas.style.width = options.width + "px";
                this.canvas.style.height = options.height + "px";
            }
            this.fontMetrics = new FontMetrics(document);
            this.ctx.scale(this.options.scale, this.options.scale);
            this.ctx.translate(-options.x + options.scrollX, -options.y + options.scrollY);
            this.ctx.textBaseline = 'bottom';
            this._activeEffects = [];
            Logger.getInstance(options.id).debug("Canvas renderer initialized (" + options.width + "x" + options.height + " at " + options.x + "," + options.y + ") with scale " + options.scale);
        }
        CanvasRenderer.prototype.applyEffects = function (effects, target) {
            var _this = this;
            while (this._activeEffects.length) {
                this.popEffect();
            }
            effects.filter(function (effect) { return contains(effect.target, target); }).forEach(function (effect) { return _this.applyEffect(effect); });
        };
        CanvasRenderer.prototype.applyEffect = function (effect) {
            this.ctx.save();
            if (isTransformEffect(effect)) {
                this.ctx.translate(effect.offsetX, effect.offsetY);
                this.ctx.transform(effect.matrix[0], effect.matrix[1], effect.matrix[2], effect.matrix[3], effect.matrix[4], effect.matrix[5]);
                this.ctx.translate(-effect.offsetX, -effect.offsetY);
            }
            if (isClipEffect(effect)) {
                this.path(effect.path);
                this.ctx.clip();
            }
            this._activeEffects.push(effect);
        };
        CanvasRenderer.prototype.popEffect = function () {
            this._activeEffects.pop();
            this.ctx.restore();
        };
        CanvasRenderer.prototype.renderStack = function (stack) {
            return __awaiter(this, void 0, void 0, function () {
                var styles;
                return __generator(this, function (_a) {
                    switch (_a.label) {
                        case 0:
                            styles = stack.element.container.styles;
                            if (!styles.isVisible()) return [3 /*break*/, 2];
                            this.ctx.globalAlpha = styles.opacity;
                            return [4 /*yield*/, this.renderStackContent(stack)];
                        case 1:
                            _a.sent();
                            _a.label = 2;
                        case 2: return [2 /*return*/];
                    }
                });
            });
        };
        CanvasRenderer.prototype.renderNode = function (paint) {
            return __awaiter(this, void 0, void 0, function () {
                return __generator(this, function (_a) {
                    switch (_a.label) {
                        case 0:
                            if (!paint.container.styles.isVisible()) return [3 /*break*/, 3];
                            return [4 /*yield*/, this.renderNodeBackgroundAndBorders(paint)];
                        case 1:
                            _a.sent();
                            return [4 /*yield*/, this.renderNodeContent(paint)];
                        case 2:
                            _a.sent();
                            _a.label = 3;
                        case 3: return [2 /*return*/];
                    }
                });
            });
        };
        CanvasRenderer.prototype.renderTextWithLetterSpacing = function (text, letterSpacing) {
            var _this = this;
            if (letterSpacing === 0) {
                this.ctx.fillText(text.text, text.bounds.left, text.bounds.top + text.bounds.height);
            }
            else {
                var letters = toCodePoints(text.text).map(function (i) { return fromCodePoint(i); });
                letters.reduce(function (left, letter) {
                    _this.ctx.fillText(letter, left, text.bounds.top + text.bounds.height);
                    return left + _this.ctx.measureText(letter).width;
                }, text.bounds.left);
            }
        };
        CanvasRenderer.prototype.createFontStyle = function (styles) {
            var fontVariant = styles.fontVariant
                .filter(function (variant) { return variant === 'normal' || variant === 'small-caps'; })
                .join('');
            var fontFamily = styles.fontFamily.join(', ');
            var fontSize = isDimensionToken(styles.fontSize)
                ? "" + styles.fontSize.number + styles.fontSize.unit
                : styles.fontSize.number + "px";
            return [
                [styles.fontStyle, fontVariant, styles.fontWeight, fontSize, fontFamily].join(' '),
                fontFamily,
                fontSize
            ];
        };
        CanvasRenderer.prototype.renderTextNode = function (text, styles) {
            return __awaiter(this, void 0, void 0, function () {
                var _a, font, fontFamily, fontSize;
                var _this = this;
                return __generator(this, function (_b) {
                    _a = this.createFontStyle(styles), font = _a[0], fontFamily = _a[1], fontSize = _a[2];
                    this.ctx.font = font;
                    text.textBounds.forEach(function (text) {
                        _this.ctx.fillStyle = asString(styles.color);
                        _this.renderTextWithLetterSpacing(text, styles.letterSpacing);
                        var textShadows = styles.textShadow;
                        if (textShadows.length && text.text.trim().length) {
                            textShadows
                                .slice(0)
                                .reverse()
                                .forEach(function (textShadow) {
                                _this.ctx.shadowColor = asString(textShadow.color);
                                _this.ctx.shadowOffsetX = textShadow.offsetX.number * _this.options.scale;
                                _this.ctx.shadowOffsetY = textShadow.offsetY.number * _this.options.scale;
                                _this.ctx.shadowBlur = textShadow.blur.number;
                                _this.ctx.fillText(text.text, text.bounds.left, text.bounds.top + text.bounds.height);
                            });
                            _this.ctx.shadowColor = '';
                            _this.ctx.shadowOffsetX = 0;
                            _this.ctx.shadowOffsetY = 0;
                            _this.ctx.shadowBlur = 0;
                        }
                        if (styles.textDecorationLine.length) {
                            _this.ctx.fillStyle = asString(styles.textDecorationColor || styles.color);
                            styles.textDecorationLine.forEach(function (textDecorationLine) {
                                switch (textDecorationLine) {
                                    case 1 /* UNDERLINE */:
                                        // Draws a line at the baseline of the font
                                        // TODO As some browsers display the line as more than 1px if the font-size is big,
                                        // need to take that into account both in position and size
                                        var baseline = _this.fontMetrics.getMetrics(fontFamily, fontSize).baseline;
                                        _this.ctx.fillRect(text.bounds.left, Math.round(text.bounds.top + baseline), text.bounds.width, 1);
                                        break;
                                    case 2 /* OVERLINE */:
                                        _this.ctx.fillRect(text.bounds.left, Math.round(text.bounds.top), text.bounds.width, 1);
                                        break;
                                    case 3 /* LINE_THROUGH */:
                                        // TODO try and find exact position for line-through
                                        var middle = _this.fontMetrics.getMetrics(fontFamily, fontSize).middle;
                                        _this.ctx.fillRect(text.bounds.left, Math.ceil(text.bounds.top + middle), text.bounds.width, 1);
                                        break;
                                }
                            });
                        }
                    });
                    return [2 /*return*/];
                });
            });
        };
        CanvasRenderer.prototype.renderReplacedElement = function (container, curves, image) {
            if (image && container.intrinsicWidth > 0 && container.intrinsicHeight > 0) {
                var box = contentBox(container);
                var path = calculatePaddingBoxPath(curves);
                this.path(path);
                this.ctx.save();
                this.ctx.clip();
                this.ctx.drawImage(image, 0, 0, container.intrinsicWidth, container.intrinsicHeight, box.left, box.top, box.width, box.height);
                this.ctx.restore();
            }
        };
        CanvasRenderer.prototype.renderNodeContent = function (paint) {
            return __awaiter(this, void 0, void 0, function () {
                var container, curves, styles, _i, _a, child, image, e_1, image, e_2, iframeRenderer, canvas, size, bounds, x, textBounds, img, image, url, e_3, bounds;
                return __generator(this, function (_b) {
                    switch (_b.label) {
                        case 0:
                            this.applyEffects(paint.effects, 4 /* CONTENT */);
                            container = paint.container;
                            curves = paint.curves;
                            styles = container.styles;
                            _i = 0, _a = container.textNodes;
                            _b.label = 1;
                        case 1:
                            if (!(_i < _a.length)) return [3 /*break*/, 4];
                            child = _a[_i];
                            return [4 /*yield*/, this.renderTextNode(child, styles)];
                        case 2:
                            _b.sent();
                            _b.label = 3;
                        case 3:
                            _i++;
                            return [3 /*break*/, 1];
                        case 4:
                            if (!(container instanceof ImageElementContainer)) return [3 /*break*/, 8];
                            _b.label = 5;
                        case 5:
                            _b.trys.push([5, 7, , 8]);
                            return [4 /*yield*/, this.options.cache.match(container.src)];
                        case 6:
                            image = _b.sent();
                            this.renderReplacedElement(container, curves, image);
                            return [3 /*break*/, 8];
                        case 7:
                            e_1 = _b.sent();
                            Logger.getInstance(this.options.id).error("Error loading image " + container.src);
                            return [3 /*break*/, 8];
                        case 8:
                            if (container instanceof CanvasElementContainer) {
                                this.renderReplacedElement(container, curves, container.canvas);
                            }
                            if (!(container instanceof SVGElementContainer)) return [3 /*break*/, 12];
                            _b.label = 9;
                        case 9:
                            _b.trys.push([9, 11, , 12]);
                            return [4 /*yield*/, this.options.cache.match(container.svg)];
                        case 10:
                            image = _b.sent();
                            this.renderReplacedElement(container, curves, image);
                            return [3 /*break*/, 12];
                        case 11:
                            e_2 = _b.sent();
                            Logger.getInstance(this.options.id).error("Error loading svg " + container.svg.substring(0, 255));
                            return [3 /*break*/, 12];
                        case 12:
                            if (!(container instanceof IFrameElementContainer && container.tree)) return [3 /*break*/, 14];
                            iframeRenderer = new CanvasRenderer({
                                id: this.options.id,
                                scale: this.options.scale,
                                backgroundColor: container.backgroundColor,
                                x: 0,
                                y: 0,
                                scrollX: 0,
                                scrollY: 0,
                                width: container.width,
                                height: container.height,
                                cache: this.options.cache,
                                windowWidth: container.width,
                                windowHeight: container.height
                            });
                            return [4 /*yield*/, iframeRenderer.render(container.tree)];
                        case 13:
                            canvas = _b.sent();
                            if (container.width && container.height) {
                                this.ctx.drawImage(canvas, 0, 0, container.width, container.height, container.bounds.left, container.bounds.top, container.bounds.width, container.bounds.height);
                            }
                            _b.label = 14;
                        case 14:
                            if (container instanceof InputElementContainer) {
                                size = Math.min(container.bounds.width, container.bounds.height);
                                if (container.type === CHECKBOX) {
                                    if (container.checked) {
                                        this.ctx.save();
                                        this.path([
                                            new Vector(container.bounds.left + size * 0.39363, container.bounds.top + size * 0.79),
                                            new Vector(container.bounds.left + size * 0.16, container.bounds.top + size * 0.5549),
                                            new Vector(container.bounds.left + size * 0.27347, container.bounds.top + size * 0.44071),
                                            new Vector(container.bounds.left + size * 0.39694, container.bounds.top + size * 0.5649),
                                            new Vector(container.bounds.left + size * 0.72983, container.bounds.top + size * 0.23),
                                            new Vector(container.bounds.left + size * 0.84, container.bounds.top + size * 0.34085),
                                            new Vector(container.bounds.left + size * 0.39363, container.bounds.top + size * 0.79)
                                        ]);
                                        this.ctx.fillStyle = asString(INPUT_COLOR);
                                        this.ctx.fill();
                                        this.ctx.restore();
                                    }
                                }
                                else if (container.type === RADIO) {
                                    if (container.checked) {
                                        this.ctx.save();
                                        this.ctx.beginPath();
                                        this.ctx.arc(container.bounds.left + size / 2, container.bounds.top + size / 2, size / 4, 0, Math.PI * 2, true);
                                        this.ctx.fillStyle = asString(INPUT_COLOR);
                                        this.ctx.fill();
                                        this.ctx.restore();
                                    }
                                }
                            }
                            if (isTextInputElement(container) && container.value.length) {
                                this.ctx.font = this.createFontStyle(styles)[0];
                                this.ctx.fillStyle = asString(styles.color);
                                this.ctx.textBaseline = 'middle';
                                this.ctx.textAlign = canvasTextAlign(container.styles.textAlign);
                                bounds = contentBox(container);
                                x = 0;
                                switch (container.styles.textAlign) {
                                    case TEXT_ALIGN.CENTER:
                                        x += bounds.width / 2;
                                        break;
                                    case TEXT_ALIGN.RIGHT:
                                        x += bounds.width;
                                        break;
                                }
                                textBounds = bounds.add(x, 0, 0, -bounds.height / 2 + 1);
                                this.ctx.save();
                                this.path([
                                    new Vector(bounds.left, bounds.top),
                                    new Vector(bounds.left + bounds.width, bounds.top),
                                    new Vector(bounds.left + bounds.width, bounds.top + bounds.height),
                                    new Vector(bounds.left, bounds.top + bounds.height)
                                ]);
                                this.ctx.clip();
                                this.renderTextWithLetterSpacing(new TextBounds(container.value, textBounds), styles.letterSpacing);
                                this.ctx.restore();
                                this.ctx.textBaseline = 'bottom';
                                this.ctx.textAlign = 'left';
                            }
                            if (!contains(container.styles.display, 2048 /* LIST_ITEM */)) return [3 /*break*/, 20];
                            if (!(container.styles.listStyleImage !== null)) return [3 /*break*/, 19];
                            img = container.styles.listStyleImage;
                            if (!(img.type === CSSImageType.URL)) return [3 /*break*/, 18];
                            image = void 0;
                            url = img.url;
                            _b.label = 15;
                        case 15:
                            _b.trys.push([15, 17, , 18]);
                            return [4 /*yield*/, this.options.cache.match(url)];
                        case 16:
                            image = _b.sent();
                            this.ctx.drawImage(image, container.bounds.left - (image.width + 10), container.bounds.top);
                            return [3 /*break*/, 18];
                        case 17:
                            e_3 = _b.sent();
                            Logger.getInstance(this.options.id).error("Error loading list-style-image " + url);
                            return [3 /*break*/, 18];
                        case 18: return [3 /*break*/, 20];
                        case 19:
                            if (paint.listValue && container.styles.listStyleType !== LIST_STYLE_TYPE.NONE) {
                                this.ctx.font = this.createFontStyle(styles)[0];
                                this.ctx.fillStyle = asString(styles.color);
                                this.ctx.textBaseline = 'middle';
                                this.ctx.textAlign = 'right';
                                bounds = new Bounds(container.bounds.left, container.bounds.top + getAbsoluteValue(container.styles.paddingTop, container.bounds.width), container.bounds.width, computeLineHeight(styles.lineHeight, styles.fontSize.number) / 2 + 1);
                                this.renderTextWithLetterSpacing(new TextBounds(paint.listValue, bounds), styles.letterSpacing);
                                this.ctx.textBaseline = 'bottom';
                                this.ctx.textAlign = 'left';
                            }
                            _b.label = 20;
                        case 20: return [2 /*return*/];
                    }
                });
            });
        };
        CanvasRenderer.prototype.renderStackContent = function (stack) {
            return __awaiter(this, void 0, void 0, function () {
                var _i, _a, child, _b, _c, child, _d, _e, child, _f, _g, child, _h, _j, child, _k, _l, child, _m, _o, child;
                return __generator(this, function (_p) {
                    switch (_p.label) {
                        case 0: 
                        // https://www.w3.org/TR/css-position-3/#painting-order
                        // 1. the background and borders of the element forming the stacking context.
                        return [4 /*yield*/, this.renderNodeBackgroundAndBorders(stack.element)];
                        case 1:
                            // https://www.w3.org/TR/css-position-3/#painting-order
                            // 1. the background and borders of the element forming the stacking context.
                            _p.sent();
                            _i = 0, _a = stack.negativeZIndex;
                            _p.label = 2;
                        case 2:
                            if (!(_i < _a.length)) return [3 /*break*/, 5];
                            child = _a[_i];
                            return [4 /*yield*/, this.renderStack(child)];
                        case 3:
                            _p.sent();
                            _p.label = 4;
                        case 4:
                            _i++;
                            return [3 /*break*/, 2];
                        case 5: 
                        // 3. For all its in-flow, non-positioned, block-level descendants in tree order:
                        return [4 /*yield*/, this.renderNodeContent(stack.element)];
                        case 6:
                            // 3. For all its in-flow, non-positioned, block-level descendants in tree order:
                            _p.sent();
                            _b = 0, _c = stack.nonInlineLevel;
                            _p.label = 7;
                        case 7:
                            if (!(_b < _c.length)) return [3 /*break*/, 10];
                            child = _c[_b];
                            return [4 /*yield*/, this.renderNode(child)];
                        case 8:
                            _p.sent();
                            _p.label = 9;
                        case 9:
                            _b++;
                            return [3 /*break*/, 7];
                        case 10:
                            _d = 0, _e = stack.nonPositionedFloats;
                            _p.label = 11;
                        case 11:
                            if (!(_d < _e.length)) return [3 /*break*/, 14];
                            child = _e[_d];
                            return [4 /*yield*/, this.renderStack(child)];
                        case 12:
                            _p.sent();
                            _p.label = 13;
                        case 13:
                            _d++;
                            return [3 /*break*/, 11];
                        case 14:
                            _f = 0, _g = stack.nonPositionedInlineLevel;
                            _p.label = 15;
                        case 15:
                            if (!(_f < _g.length)) return [3 /*break*/, 18];
                            child = _g[_f];
                            return [4 /*yield*/, this.renderStack(child)];
                        case 16:
                            _p.sent();
                            _p.label = 17;
                        case 17:
                            _f++;
                            return [3 /*break*/, 15];
                        case 18:
                            _h = 0, _j = stack.inlineLevel;
                            _p.label = 19;
                        case 19:
                            if (!(_h < _j.length)) return [3 /*break*/, 22];
                            child = _j[_h];
                            return [4 /*yield*/, this.renderNode(child)];
                        case 20:
                            _p.sent();
                            _p.label = 21;
                        case 21:
                            _h++;
                            return [3 /*break*/, 19];
                        case 22:
                            _k = 0, _l = stack.zeroOrAutoZIndexOrTransformedOrOpacity;
                            _p.label = 23;
                        case 23:
                            if (!(_k < _l.length)) return [3 /*break*/, 26];
                            child = _l[_k];
                            return [4 /*yield*/, this.renderStack(child)];
                        case 24:
                            _p.sent();
                            _p.label = 25;
                        case 25:
                            _k++;
                            return [3 /*break*/, 23];
                        case 26:
                            _m = 0, _o = stack.positiveZIndex;
                            _p.label = 27;
                        case 27:
                            if (!(_m < _o.length)) return [3 /*break*/, 30];
                            child = _o[_m];
                            return [4 /*yield*/, this.renderStack(child)];
                        case 28:
                            _p.sent();
                            _p.label = 29;
                        case 29:
                            _m++;
                            return [3 /*break*/, 27];
                        case 30: return [2 /*return*/];
                    }
                });
            });
        };
        CanvasRenderer.prototype.mask = function (paths) {
            this.ctx.beginPath();
            this.ctx.moveTo(0, 0);
            this.ctx.lineTo(this.canvas.width, 0);
            this.ctx.lineTo(this.canvas.width, this.canvas.height);
            this.ctx.lineTo(0, this.canvas.height);
            this.ctx.lineTo(0, 0);
            this.formatPath(paths.slice(0).reverse());
            this.ctx.closePath();
        };
        CanvasRenderer.prototype.path = function (paths) {
            this.ctx.beginPath();
            this.formatPath(paths);
            this.ctx.closePath();
        };
        CanvasRenderer.prototype.formatPath = function (paths) {
            var _this = this;
            paths.forEach(function (point, index) {
                var start = isBezierCurve(point) ? point.start : point;
                if (index === 0) {
                    _this.ctx.moveTo(start.x, start.y);
                }
                else {
                    _this.ctx.lineTo(start.x, start.y);
                }
                if (isBezierCurve(point)) {
                    _this.ctx.bezierCurveTo(point.startControl.x, point.startControl.y, point.endControl.x, point.endControl.y, point.end.x, point.end.y);
                }
            });
        };
        CanvasRenderer.prototype.renderRepeat = function (path, pattern, offsetX, offsetY) {
            this.path(path);
            this.ctx.fillStyle = pattern;
            this.ctx.translate(offsetX, offsetY);
            this.ctx.fill();
            this.ctx.translate(-offsetX, -offsetY);
        };
        CanvasRenderer.prototype.resizeImage = function (image, width, height) {
            if (image.width === width && image.height === height) {
                return image;
            }
            var canvas = this.canvas.ownerDocument.createElement('canvas');
            canvas.width = width;
            canvas.height = height;
            var ctx = canvas.getContext('2d');
            ctx.drawImage(image, 0, 0, image.width, image.height, 0, 0, width, height);
            return canvas;
        };
        CanvasRenderer.prototype.renderBackgroundImage = function (container) {
            return __awaiter(this, void 0, void 0, function () {
                var index, _loop_1, this_1, _i, _a, backgroundImage;
                return __generator(this, function (_b) {
                    switch (_b.label) {
                        case 0:
                            index = container.styles.backgroundImage.length - 1;
                            _loop_1 = function (backgroundImage) {
                                var image, url, e_4, _a, path, x, y, width, height, pattern, _b, path, x, y, width, height, _c, lineLength, x0, x1, y0, y1, canvas, ctx, gradient_1, pattern, _d, path, left, top_1, width, height, position, x, y, _e, rx, ry, radialGradient_1, midX, midY, f, invF;
                                return __generator(this, function (_f) {
                                    switch (_f.label) {
                                        case 0:
                                            if (!(backgroundImage.type === CSSImageType.URL)) return [3 /*break*/, 5];
                                            image = void 0;
                                            url = backgroundImage.url;
                                            _f.label = 1;
                                        case 1:
                                            _f.trys.push([1, 3, , 4]);
                                            return [4 /*yield*/, this_1.options.cache.match(url)];
                                        case 2:
                                            image = _f.sent();
                                            return [3 /*break*/, 4];
                                        case 3:
                                            e_4 = _f.sent();
                                            Logger.getInstance(this_1.options.id).error("Error loading background-image " + url);
                                            return [3 /*break*/, 4];
                                        case 4:
                                            if (image) {
                                                _a = calculateBackgroundRendering(container, index, [
                                                    image.width,
                                                    image.height,
                                                    image.width / image.height
                                                ]), path = _a[0], x = _a[1], y = _a[2], width = _a[3], height = _a[4];
                                                pattern = this_1.ctx.createPattern(this_1.resizeImage(image, width, height), 'repeat');
                                                this_1.renderRepeat(path, pattern, x, y);
                                            }
                                            return [3 /*break*/, 6];
                                        case 5:
                                            if (isLinearGradient(backgroundImage)) {
                                                _b = calculateBackgroundRendering(container, index, [null, null, null]), path = _b[0], x = _b[1], y = _b[2], width = _b[3], height = _b[4];
                                                _c = calculateGradientDirection(backgroundImage.angle, width, height), lineLength = _c[0], x0 = _c[1], x1 = _c[2], y0 = _c[3], y1 = _c[4];
                                                canvas = document.createElement('canvas');
                                                canvas.width = width;
                                                canvas.height = height;
                                                ctx = canvas.getContext('2d');
                                                gradient_1 = ctx.createLinearGradient(x0, y0, x1, y1);
                                                processColorStops(backgroundImage.stops, lineLength).forEach(function (colorStop) {
                                                    return gradient_1.addColorStop(colorStop.stop, asString(colorStop.color));
                                                });
                                                ctx.fillStyle = gradient_1;
                                                ctx.fillRect(0, 0, width, height);
                                                if (width > 0 && height > 0) {
                                                    pattern = this_1.ctx.createPattern(canvas, 'repeat');
                                                    this_1.renderRepeat(path, pattern, x, y);
                                                }
                                            }
                                            else if (isRadialGradient(backgroundImage)) {
                                                _d = calculateBackgroundRendering(container, index, [
                                                    null,
                                                    null,
                                                    null
                                                ]), path = _d[0], left = _d[1], top_1 = _d[2], width = _d[3], height = _d[4];
                                                position = backgroundImage.position.length === 0 ? [FIFTY_PERCENT] : backgroundImage.position;
                                                x = getAbsoluteValue(position[0], width);
                                                y = getAbsoluteValue(position[position.length - 1], height);
                                                _e = calculateRadius(backgroundImage, x, y, width, height), rx = _e[0], ry = _e[1];
                                                if (rx > 0 && rx > 0) {
                                                    radialGradient_1 = this_1.ctx.createRadialGradient(left + x, top_1 + y, 0, left + x, top_1 + y, rx);
                                                    processColorStops(backgroundImage.stops, rx * 2).forEach(function (colorStop) {
                                                        return radialGradient_1.addColorStop(colorStop.stop, asString(colorStop.color));
                                                    });
                                                    this_1.path(path);
                                                    this_1.ctx.fillStyle = radialGradient_1;
                                                    if (rx !== ry) {
                                                        midX = container.bounds.left + 0.5 * container.bounds.width;
                                                        midY = container.bounds.top + 0.5 * container.bounds.height;
                                                        f = ry / rx;
                                                        invF = 1 / f;
                                                        this_1.ctx.save();
                                                        this_1.ctx.translate(midX, midY);
                                                        this_1.ctx.transform(1, 0, 0, f, 0, 0);
                                                        this_1.ctx.translate(-midX, -midY);
                                                        this_1.ctx.fillRect(left, invF * (top_1 - midY) + midY, width, height * invF);
                                                        this_1.ctx.restore();
                                                    }
                                                    else {
                                                        this_1.ctx.fill();
                                                    }
                                                }
                                            }
                                            _f.label = 6;
                                        case 6:
                                            index--;
                                            return [2 /*return*/];
                                    }
                                });
                            };
                            this_1 = this;
                            _i = 0, _a = container.styles.backgroundImage.slice(0).reverse();
                            _b.label = 1;
                        case 1:
                            if (!(_i < _a.length)) return [3 /*break*/, 4];
                            backgroundImage = _a[_i];
                            return [5 /*yield**/, _loop_1(backgroundImage)];
                        case 2:
                            _b.sent();
                            _b.label = 3;
                        case 3:
                            _i++;
                            return [3 /*break*/, 1];
                        case 4: return [2 /*return*/];
                    }
                });
            });
        };
        CanvasRenderer.prototype.renderBorder = function (color, side, curvePoints) {
            return __awaiter(this, void 0, void 0, function () {
                return __generator(this, function (_a) {
                    this.path(parsePathForBorder(curvePoints, side));
                    this.ctx.fillStyle = asString(color);
                    this.ctx.fill();
                    return [2 /*return*/];
                });
            });
        };
        CanvasRenderer.prototype.renderNodeBackgroundAndBorders = function (paint) {
            return __awaiter(this, void 0, void 0, function () {
                var styles, hasBackground, borders, backgroundPaintingArea, side, _i, borders_1, border;
                var _this = this;
                return __generator(this, function (_a) {
                    switch (_a.label) {
                        case 0:
                            this.applyEffects(paint.effects, 2 /* BACKGROUND_BORDERS */);
                            styles = paint.container.styles;
                            hasBackground = !isTransparent(styles.backgroundColor) || styles.backgroundImage.length;
                            borders = [
                                { style: styles.borderTopStyle, color: styles.borderTopColor },
                                { style: styles.borderRightStyle, color: styles.borderRightColor },
                                { style: styles.borderBottomStyle, color: styles.borderBottomColor },
                                { style: styles.borderLeftStyle, color: styles.borderLeftColor }
                            ];
                            backgroundPaintingArea = calculateBackgroundCurvedPaintingArea(getBackgroundValueForIndex(styles.backgroundClip, 0), paint.curves);
                            if (!(hasBackground || styles.boxShadow.length)) return [3 /*break*/, 2];
                            this.ctx.save();
                            this.path(backgroundPaintingArea);
                            this.ctx.clip();
                            if (!isTransparent(styles.backgroundColor)) {
                                this.ctx.fillStyle = asString(styles.backgroundColor);
                                this.ctx.fill();
                            }
                            return [4 /*yield*/, this.renderBackgroundImage(paint.container)];
                        case 1:
                            _a.sent();
                            this.ctx.restore();
                            styles.boxShadow
                                .slice(0)
                                .reverse()
                                .forEach(function (shadow) {
                                _this.ctx.save();
                                var borderBoxArea = calculateBorderBoxPath(paint.curves);
                                var maskOffset = shadow.inset ? 0 : MASK_OFFSET;
                                var shadowPaintingArea = transformPath(borderBoxArea, -maskOffset + (shadow.inset ? 1 : -1) * shadow.spread.number, (shadow.inset ? 1 : -1) * shadow.spread.number, shadow.spread.number * (shadow.inset ? -2 : 2), shadow.spread.number * (shadow.inset ? -2 : 2));
                                if (shadow.inset) {
                                    _this.path(borderBoxArea);
                                    _this.ctx.clip();
                                    _this.mask(shadowPaintingArea);
                                }
                                else {
                                    _this.mask(borderBoxArea);
                                    _this.ctx.clip();
                                    _this.path(shadowPaintingArea);
                                }
                                _this.ctx.shadowOffsetX = shadow.offsetX.number + maskOffset;
                                _this.ctx.shadowOffsetY = shadow.offsetY.number;
                                _this.ctx.shadowColor = asString(shadow.color);
                                _this.ctx.shadowBlur = shadow.blur.number;
                                _this.ctx.fillStyle = shadow.inset ? asString(shadow.color) : 'rgba(0,0,0,1)';
                                _this.ctx.fill();
                                _this.ctx.restore();
                            });
                            _a.label = 2;
                        case 2:
                            side = 0;
                            _i = 0, borders_1 = borders;
                            _a.label = 3;
                        case 3:
                            if (!(_i < borders_1.length)) return [3 /*break*/, 7];
                            border = borders_1[_i];
                            if (!(border.style !== BORDER_STYLE.NONE && !isTransparent(border.color))) return [3 /*break*/, 5];
                            return [4 /*yield*/, this.renderBorder(border.color, side, paint.curves)];
                        case 4:
                            _a.sent();
                            _a.label = 5;
                        case 5:
                            side++;
                            _a.label = 6;
                        case 6:
                            _i++;
                            return [3 /*break*/, 3];
                        case 7: return [2 /*return*/];
                    }
                });
            });
        };
        CanvasRenderer.prototype.render = function (element) {
            return __awaiter(this, void 0, void 0, function () {
                var stack;
                return __generator(this, function (_a) {
                    switch (_a.label) {
                        case 0:
                            if (this.options.backgroundColor) {
                                this.ctx.fillStyle = asString(this.options.backgroundColor);
                                this.ctx.fillRect(this.options.x - this.options.scrollX, this.options.y - this.options.scrollY, this.options.width, this.options.height);
                            }
                            stack = parseStackingContexts(element);
                            return [4 /*yield*/, this.renderStack(stack)];
                        case 1:
                            _a.sent();
                            this.applyEffects([], 2 /* BACKGROUND_BORDERS */);
                            return [2 /*return*/, this.canvas];
                    }
                });
            });
        };
        return CanvasRenderer;
    }());
    var isTextInputElement = function (container) {
        if (container instanceof TextareaElementContainer) {
            return true;
        }
        else if (container instanceof SelectElementContainer) {
            return true;
        }
        else if (container instanceof InputElementContainer && container.type !== RADIO && container.type !== CHECKBOX) {
            return true;
        }
        return false;
    };
    var calculateBackgroundCurvedPaintingArea = function (clip, curves) {
        switch (clip) {
            case BACKGROUND_CLIP.BORDER_BOX:
                return calculateBorderBoxPath(curves);
            case BACKGROUND_CLIP.CONTENT_BOX:
                return calculateContentBoxPath(curves);
            case BACKGROUND_CLIP.PADDING_BOX:
            default:
                return calculatePaddingBoxPath(curves);
        }
    };
    var canvasTextAlign = function (textAlign) {
        switch (textAlign) {
            case TEXT_ALIGN.CENTER:
                return 'center';
            case TEXT_ALIGN.RIGHT:
                return 'right';
            case TEXT_ALIGN.LEFT:
            default:
                return 'left';
        }
    };

    var ForeignObjectRenderer = /** @class */ (function () {
        function ForeignObjectRenderer(options) {
            this.canvas = options.canvas ? options.canvas : document.createElement('canvas');
            this.ctx = this.canvas.getContext('2d');
            this.options = options;
            this.canvas.width = Math.floor(options.width * options.scale);
            this.canvas.height = Math.floor(options.height * options.scale);
            this.canvas.style.width = options.width + "px";
            this.canvas.style.height = options.height + "px";
            this.ctx.scale(this.options.scale, this.options.scale);
            this.ctx.translate(-options.x + options.scrollX, -options.y + options.scrollY);
            Logger.getInstance(options.id).debug("EXPERIMENTAL ForeignObject renderer initialized (" + options.width + "x" + options.height + " at " + options.x + "," + options.y + ") with scale " + options.scale);
        }
        ForeignObjectRenderer.prototype.render = function (element) {
            return __awaiter(this, void 0, void 0, function () {
                var svg, img;
                return __generator(this, function (_a) {
                    switch (_a.label) {
                        case 0:
                            svg = createForeignObjectSVG(Math.max(this.options.windowWidth, this.options.width) * this.options.scale, Math.max(this.options.windowHeight, this.options.height) * this.options.scale, this.options.scrollX * this.options.scale, this.options.scrollY * this.options.scale, element);
                            return [4 /*yield*/, loadSerializedSVG$1(svg)];
                        case 1:
                            img = _a.sent();
                            if (this.options.backgroundColor) {
                                this.ctx.fillStyle = asString(this.options.backgroundColor);
                                this.ctx.fillRect(0, 0, this.options.width * this.options.scale, this.options.height * this.options.scale);
                            }
                            this.ctx.drawImage(img, -this.options.x * this.options.scale, -this.options.y * this.options.scale);
                            return [2 /*return*/, this.canvas];
                    }
                });
            });
        };
        return ForeignObjectRenderer;
    }());
    var loadSerializedSVG$1 = function (svg) {
        return new Promise(function (resolve, reject) {
            var img = new Image();
            img.onload = function () {
                resolve(img);
            };
            img.onerror = reject;
            img.src = "data:image/svg+xml;charset=utf-8," + encodeURIComponent(new XMLSerializer().serializeToString(svg));
        });
    };

    var _this = undefined;
    var parseColor$1 = function (value) { return color.parse(Parser.create(value).parseComponentValue()); };
    var html2canvas = function (element, options) {
        if (options === void 0) { options = {}; }
        return renderElement(element, options);
    };
    if (typeof window !== "undefined") {
        CacheStorage.setContext(window);
    }
    var renderElement = function (element, opts) { return __awaiter(_this, void 0, void 0, function () {
        var ownerDocument, defaultView, instanceName, _a, width, height, left, top, defaultResourceOptions, resourceOptions, defaultOptions, options, windowBounds, documentCloner, clonedElement, container, documentBackgroundColor, bodyBackgroundColor, bgColor, defaultBackgroundColor, backgroundColor, renderOptions, canvas, renderer, root, renderer;
        return __generator(this, function (_b) {
            switch (_b.label) {
                case 0:
                    ownerDocument = element.ownerDocument;
                    if (!ownerDocument) {
                        throw new Error("Element is not attached to a Document");
                    }
                    defaultView = ownerDocument.defaultView;
                    if (!defaultView) {
                        throw new Error("Document is not attached to a Window");
                    }
                    instanceName = (Math.round(Math.random() * 1000) + Date.now()).toString(16);
                    _a = isBodyElement(element) || isHTMLElement(element) ? parseDocumentSize(ownerDocument) : parseBounds(element), width = _a.width, height = _a.height, left = _a.left, top = _a.top;
                    defaultResourceOptions = {
                        allowTaint: false,
                        imageTimeout: 15000,
                        proxy: undefined,
                        useCORS: false
                    };
                    resourceOptions = __assign({}, defaultResourceOptions, opts);
                    defaultOptions = {
                        backgroundColor: '#ffffff',
                        cache: opts.cache ? opts.cache : CacheStorage.create(instanceName, resourceOptions),
                        logging: true,
                        removeContainer: true,
                        foreignObjectRendering: false,
                        scale: defaultView.devicePixelRatio || 1,
                        windowWidth: defaultView.innerWidth,
                        windowHeight: defaultView.innerHeight,
                        scrollX: defaultView.pageXOffset,
                        scrollY: defaultView.pageYOffset,
                        x: left,
                        y: top,
                        width: Math.ceil(width),
                        height: Math.ceil(height),
                        id: instanceName
                    };
                    options = __assign({}, defaultOptions, resourceOptions, opts);
                    windowBounds = new Bounds(options.scrollX, options.scrollY, options.windowWidth, options.windowHeight);
                    Logger.create({ id: instanceName, enabled: options.logging });
                    Logger.getInstance(instanceName).debug("Starting document clone");
                    documentCloner = new DocumentCloner(element, {
                        id: instanceName,
                        onclone: options.onclone,
                        ignoreElements: options.ignoreElements,
                        inlineImages: options.foreignObjectRendering,
                        copyStyles: options.foreignObjectRendering
                    });
                    clonedElement = documentCloner.clonedReferenceElement;
                    if (!clonedElement) {
                        return [2 /*return*/, Promise.reject("Unable to find element in cloned iframe")];
                    }
                    return [4 /*yield*/, documentCloner.toIFrame(ownerDocument, windowBounds)];
                case 1:
                    container = _b.sent();
                    documentBackgroundColor = ownerDocument.documentElement
                        ? parseColor$1(getComputedStyle(ownerDocument.documentElement).backgroundColor)
                        : COLORS.TRANSPARENT;
                    bodyBackgroundColor = ownerDocument.body
                        ? parseColor$1(getComputedStyle(ownerDocument.body).backgroundColor)
                        : COLORS.TRANSPARENT;
                    bgColor = opts.backgroundColor;
                    defaultBackgroundColor = typeof bgColor === 'string' ? parseColor$1(bgColor) : bgColor === null ? COLORS.TRANSPARENT : 0xffffffff;
                    backgroundColor = element === ownerDocument.documentElement
                        ? isTransparent(documentBackgroundColor)
                            ? isTransparent(bodyBackgroundColor)
                                ? defaultBackgroundColor
                                : bodyBackgroundColor
                            : documentBackgroundColor
                        : defaultBackgroundColor;
                    renderOptions = {
                        id: instanceName,
                        cache: options.cache,
                        canvas: options.canvas,
                        backgroundColor: backgroundColor,
                        scale: options.scale,
                        x: options.x,
                        y: options.y,
                        scrollX: options.scrollX,
                        scrollY: options.scrollY,
                        width: options.width,
                        height: options.height,
                        windowWidth: options.windowWidth,
                        windowHeight: options.windowHeight
                    };
                    if (!options.foreignObjectRendering) return [3 /*break*/, 3];
                    Logger.getInstance(instanceName).debug("Document cloned, using foreign object rendering");
                    renderer = new ForeignObjectRenderer(renderOptions);
                    return [4 /*yield*/, renderer.render(clonedElement)];
                case 2:
                    canvas = _b.sent();
                    return [3 /*break*/, 5];
                case 3:
                    Logger.getInstance(instanceName).debug("Document cloned, using computed rendering");
                    CacheStorage.attachInstance(options.cache);
                    Logger.getInstance(instanceName).debug("Starting DOM parsing");
                    root = parseTree(clonedElement);
                    CacheStorage.detachInstance();
                    if (backgroundColor === root.styles.backgroundColor) {
                        root.styles.backgroundColor = COLORS.TRANSPARENT;
                    }
                    Logger.getInstance(instanceName).debug("Starting renderer");
                    renderer = new CanvasRenderer(renderOptions);
                    return [4 /*yield*/, renderer.render(root)];
                case 4:
                    canvas = _b.sent();
                    _b.label = 5;
                case 5:
                    if (options.removeContainer === true) {
                        if (!DocumentCloner.destroy(container)) {
                            Logger.getInstance(instanceName).error("Cannot detach cloned iframe as it is not in the DOM anymore");
                        }
                    }
                    Logger.getInstance(instanceName).debug("Finished rendering");
                    Logger.destroy(instanceName);
                    CacheStorage.destroy(instanceName);
                    return [2 /*return*/, canvas];
            }
        });
    }); };

    return html2canvas;

}));
//# sourceMappingURL=html2canvas.js.map
```
