## Supplementary material for "Method for inferring migration routes from phylogenetic trees": Some route-trees: ntseq.js

```
var Nt = function() {

  'use strict';

  function makeArray(length, val) {
    if (val === undefined) { val = 0|0; }
    if (val < 0) { val = 0; }
    length |= 0;
    var max = 0;
    for (var i = length; i !== 0; i >>>= 1) { max++; }
      var n = Array(max);
      n[0] = [val];
      for (i = 1; i < max; i++) {
      n[i] = n[i-1].concat(n[i-1]);
    }
    var a = [];
    for (var i = 0, l = length; l !== 0; l >>>= 1, i++) {
      if (l&1) {
        a = a.concat(n[i]);
      }
    }
    return a;
  };

  var __bitCount = (function() {
    var a = new Uint8Array(256);
    var bin;
    for (var i = 0; i < 256; i++) {
      bin = i;
      bin = bin - ((bin >> 1) & 0x55555555);
      bin = (bin & 0x33333333) + ((bin >> 2) & 0x33333333);
      a[i] = (((bin + (bin >> 4)) & 0x0F0F0F0F) * 0x01010101) >> 24;
    }
    return a;
  })();

  var __nucleotideTo4Bit = Object.create(null);
  __nucleotideTo4Bit['A'] = 8; 			// 0b1000
  __nucleotideTo4Bit['T'] = 4; 			// 0b0100
  __nucleotideTo4Bit['G'] = 2; 			// 0b0010
  __nucleotideTo4Bit['C'] = 1; 			// 0b0001

  function setNucleotide() {
    var n = arguments[0];
    __nucleotideTo4Bit[n] = 0;
    for (var i = 1; i < arguments.length; i++) {
      __nucleotideTo4Bit[n] |= __nucleotideTo4Bit[arguments[i]];
    }
  };

  setNucleotide('-');
  setNucleotide('W', 'A', 'T');
  setNucleotide('S', 'G', 'C');
  setNucleotide('M', 'A', 'C');
  setNucleotide('K', 'G', 'T');
  setNucleotide('R', 'A', 'G');
  setNucleotide('Y', 'C', 'T');
  setNucleotide('B', 'C', 'G', 'T');
  setNucleotide('D', 'A', 'G', 'T');
  setNucleotide('H', 'A', 'C', 'T');
  setNucleotide('V', 'A', 'C', 'G');
  setNucleotide('N', 'A', 'T', 'G', 'C');

  var __4BitToNucleotide = (
    function() {
      var a = makeArray(16);
      var keys = Object.keys(__nucleotideTo4Bit);
      for (var i = 0, len = keys.length; i < len; i++) {
        a[__nucleotideTo4Bit[keys[i]]] = keys[i];
      }
      return a;
    }
  )();

  var __nucleotideList = Object.keys(__nucleotideTo4Bit);

  var __complementNucleotide = (
    function() {
      var a = Object.create(null);
      a['A'] = 'T';
      a['G'] = 'C';
      a['B'] = 'V';
      a['H'] = 'D';
      a['M'] = 'K';
      a['R'] = 'Y';
        // S, W, N, - not included
      var keys = Object.keys(a);
      for (var i = 0, len = keys.length; i < len; i++) {
        a[a[keys[i]]] = keys[i];
      }
      a['S'] = 'S';
      a['W'] = 'W';
      a['N'] = 'N';
      a['-'] = '-';
      return a;
    }
  )();

  var __complement4Bit = (
    function() {
      var a = new Uint8Array(16);
      for (var i = 0, len = a.length; i < len; i++) {
        a[i] = __nucleotideTo4Bit[__complementNucleotide[__4BitToNucleotide[i]]];
      }
      return a;
    }
  )();

  function nucleotideToBin(s) {
    return __nucleotideTo4Bit[s] | 0;
  }

  function binToNucleotide(b) {
    return __4BitToNucleotide[b] || '-';
  }

  function complementNucleotide(s) {
    return __complementNucleotide[s] || '-';
  }

  function complementBin(b) {
    return __complement4Bit[b] | 0;
  }

  /* Double up to form bytes */
  var __byteComplement;
  var __nucleotidesToByte;
  var __byteToNucleotides;
  var __byteNucleotideContent;

  void function() {

    var a = Object.create(null);
    var b = new Uint8Array(256);
    var c = new makeArray(256);
    var d = new makeArray(256);

    var keys = Object.keys(__nucleotideTo4Bit);
    var len = keys.length;
    var ki;
    var kj;
    var byte;

    for (var i = 0; i < len; i++) {
      ki = keys[i];
      for (var j = 0; j < len; j++) {
        kj = keys[j];
        byte = __nucleotideTo4Bit[ki] | (__nucleotideTo4Bit[kj] << 4);
        a[ki + kj] = byte;
        b[byte] = __nucleotideTo4Bit[complementNucleotide(kj)] | (__nucleotideTo4Bit[complementNucleotide(ki)] << 4);
        c[byte] = ki + kj;
        d[byte] = [ki, kj];
      }
    }

    __nucleotidesToByte = a;
    __byteComplement = b;
    __byteToNucleotides = c;
    __byteNucleotideContent = d;

  }();

  function nucleotidesToByte(ss) {
    return __nucleotidesToByte[ss] | 0;
  }

  /* amino acids */

  var __codonToAminoAcid = Object.create(null);
  __codonToAminoAcid['AAA'] = 'K';
  __codonToAminoAcid['AAT'] = 'N';
  __codonToAminoAcid['AAG'] = 'K';
  __codonToAminoAcid['AAC'] = 'N';
  __codonToAminoAcid['ATA'] = 'I';
  __codonToAminoAcid['ATT'] = 'I';
  __codonToAminoAcid['ATG'] = 'M';
  __codonToAminoAcid['ATC'] = 'I';
  __codonToAminoAcid['AGA'] = 'R';
  __codonToAminoAcid['AGT'] = 'S';
  __codonToAminoAcid['AGG'] = 'R';
  __codonToAminoAcid['AGC'] = 'S';
  __codonToAminoAcid['ACA'] = 'T';
  __codonToAminoAcid['ACT'] = 'T';
  __codonToAminoAcid['ACG'] = 'T';
  __codonToAminoAcid['ACC'] = 'T';
  __codonToAminoAcid['TAA'] = '*';
  __codonToAminoAcid['TAT'] = 'Y';
  __codonToAminoAcid['TAG'] = '&';
  __codonToAminoAcid['TAC'] = 'Y';
  __codonToAminoAcid['TTA'] = 'L';
  __codonToAminoAcid['TTT'] = 'F';
  __codonToAminoAcid['TTG'] = 'L';
  __codonToAminoAcid['TTC'] = 'F';
  __codonToAminoAcid['TGA'] = '$';
  __codonToAminoAcid['TGT'] = 'C';
  __codonToAminoAcid['TGG'] = 'W';
  __codonToAminoAcid['TGC'] = 'C';
  __codonToAminoAcid['TCA'] = 'S';
  __codonToAminoAcid['TCT'] = 'S';
  __codonToAminoAcid['TCG'] = 'S';
  __codonToAminoAcid['TCC'] = 'S';
  __codonToAminoAcid['GAA'] = 'E';
  __codonToAminoAcid['GAT'] = 'D';
  __codonToAminoAcid['GAG'] = 'E';
  __codonToAminoAcid['GAC'] = 'D';
  __codonToAminoAcid['GTA'] = 'V';
  __codonToAminoAcid['GTT'] = 'V';
  __codonToAminoAcid['GTG'] = 'V';
  __codonToAminoAcid['GTC'] = 'V';
  __codonToAminoAcid['GGA'] = 'G';
  __codonToAminoAcid['GGT'] = 'G';
  __codonToAminoAcid['GGG'] = 'G';
  __codonToAminoAcid['GGC'] = 'G';
  __codonToAminoAcid['GCA'] = 'A';
  __codonToAminoAcid['GCT'] = 'A';
  __codonToAminoAcid['GCG'] = 'A';
  __codonToAminoAcid['GCC'] = 'A';
  __codonToAminoAcid['CAA'] = 'Q';
  __codonToAminoAcid['CAT'] = 'H';
  __codonToAminoAcid['CAG'] = 'Q';
  __codonToAminoAcid['CAC'] = 'H';
  __codonToAminoAcid['CTA'] = 'L';
  __codonToAminoAcid['CTT'] = 'L';
  __codonToAminoAcid['CTG'] = 'L';
  __codonToAminoAcid['CTC'] = 'L';
  __codonToAminoAcid['CGA'] = 'R';
  __codonToAminoAcid['CGT'] = 'R';
  __codonToAminoAcid['CGG'] = 'R';
  __codonToAminoAcid['CGC'] = 'R';
  __codonToAminoAcid['CCA'] = 'P';
  __codonToAminoAcid['CCT'] = 'P';
  __codonToAminoAcid['CCG'] = 'P';
  __codonToAminoAcid['CCC'] = 'P';

  var __12BitToAminoAcid = (function() {

    var a = makeArray(4096, '?');
    var codons = Object.keys(__codonToAminoAcid);
    var codon;
    for (var i = 0, len = codons.length; i < len; i++) {
      codon = codons[i];
      a[
        (nucleotideToBin(codon[2]) << 8) |
        (nucleotideToBin(codon[1]) << 4) |
        nucleotideToBin(codon[0])
      ] = __codonToAminoAcid[codon];
    }

    return a;

  })();

  function Seq(type) {

    if (type === undefined) {
      type = 'DNA';
    }

    if (!{'RNA': true, 'DNA': true}[type]) {
      throw new Error('Sequence type ' + type + ' not supported');
    }

    this.__type = type;
    this.__isRNA = (type === 'RNA');

    this.__endPadding = 0;

    this.__buffer = new ArrayBuffer(4);

    this.__complement = null;

    this.__content = null;
    this.__fractionalContent = null;
    this.__contentATGC = null;
    this.__fractionalContent = null;

  };

  Seq.prototype.read = function(strData) {

    var ntToByte = nucleotidesToByte;

    var nucleotideString = strData.toUpperCase()
      .replace(/\s/g, '')
      .replace('U', 'T')
      .replace(/[^ATGCBVHDMKRYSWN\-]/g, '-');

    var length = nucleotideString.length | 0;
    var max = length >>> 1;
    var odd = length & 1;

    var endPadding = (4 - (max + odd) % 4) % 4;
    this.__endPadding = endPadding;

    var buffer = new ArrayBuffer(max + odd + endPadding + 4);
    var dataArray = new Int8Array(buffer, 4);

    var n;
    var byte;
    var content;

    for (var i = 0; i < max; i++) {

      n = i << 1;
      dataArray[i] = ntToByte(nucleotideString[n] + nucleotideString[++n]);

    }

    if (odd) {

      dataArray[i] = __nucleotideTo4Bit[nucleotideString[i << 1]];

    }

    this.__buffer = buffer;
    this.__length = length;
    (new Uint32Array(buffer, 0, 1))[0] = length;

    this.__complement = null;

    this.__content = null;
    this.__fractionalContent = null;
    this.__contentATGC = null;
    this.__fractionalContent = null;

    return this;

  };

  Seq.prototype.readFASTA = function(strFASTA) {

    var data = strFASTA.split(/\n\r?/gi);

    while (data.length && data[0][0] === '>') {
      data.shift();
    }

    return this.read(data.join(''));

  };

  Seq.prototype.readBuffer = function(buffer) {

    this.__buffer = buffer;

    var length = (new Uint32Array(buffer, 0, 1))[0];

    var max = length >>> 1;
    var odd = length & 1;

    var endPadding = (4 - (max + odd) % 4) % 4;
    this.__endPadding = endPadding;

    this.__length = length;

    this.__complement = null;

    this.__content = null;
    this.__fractionalContent = null;
    this.__contentATGC = null;
    this.__fractionalContent = null;

    return this;

  };

  Seq.prototype.__byteComplement = function() {

    var bComp = __byteComplement;

    var fwdBuffer = this.__buffer;
    var len = fwdBuffer.byteLength;

    var n, i;

    var copyBuffer, fromArray, copyArray;

    var isOdd = this.__length & 1;

    if (isOdd) {

      copyBuffer = new ArrayBuffer(len);
      copyArray = new Uint32Array(copyBuffer, 4);
      fromArray = new Uint32Array(fwdBuffer, 4);

      n = (len - 4) >>> 2;
      while(n--) {
        copyArray[n] = (fromArray[n] << 4) | ((fromArray[n - 1]) >>> 28);
      }

    } else {

      copyBuffer = fwdBuffer;

    }

    var fwdArray = new Uint8Array(copyBuffer, 4);

    var buffer = new ArrayBuffer(len);
    var dataArray = new Uint8Array(buffer, 4);

    n = (len - 4) - this.__endPadding;
    i = 0;
    while(n--) {
      dataArray[i++] = bComp[fwdArray[n]];
    }

    (new Uint32Array(buffer, 0, 1))[0] = this.__length;

    return buffer;

  };

  Seq.prototype.size = function() {
    return this.__length;
  };

  Seq.prototype.sequence = function() {

    var byteToNt = __byteToNucleotides;
    var buffer = this.__buffer;

    if (buffer.byteLength < 4) {
      return '';
    }

    var dataArray = new Uint8Array(buffer, 4);
    var len = (buffer.byteLength - 4) - this.__endPadding;

    var nts = makeArray(len);

    for (var i = 0; i < len; i++) {
      nts[i] = byteToNt[dataArray[i]];
    }

    var returnString;

    i = nts.length - 1;
    if (this.__length & 1) {
      nts[i] = nts[i][0];
    }

    returnString = nts.join('');

    if (this.__isRNA) {
      returnString = returnString.replace(/T/gi, 'U');
    }

    return returnString;

  };

  Seq.prototype.complement = function() {

    if (!this.__complement) {
      this.__complement = this.__byteComplement();
    }

    var complement = new Seq(this.__type).readBuffer(this.__complement.slice(0));

    complement.__complement = this.__buffer.slice(0);

    return complement;

  };

  Seq.prototype.equivalent = function(seq) {

    if (!(seq instanceof Seq)) {
      throw new Error('Can only check for equivalence between sequences');
    }

    if (this.__type !== seq.__type) {
      return false;
    }

    var checkInts = new Uint32Array(this.__buffer);
    var compareInts = new Uint32Array(seq.__buffer);

    for (var i = 0, len = checkInts.length; i < len; i++) {
      if (checkInts[i] !== compareInts[i]) {
        return false;
      }
    }

    return true;

  };

  Seq.prototype.replicate = function(start, length) {

    start |= 0;

    if (start < 0) {
      start = this.__length + start;
    }

    if (length === undefined) {

      if (start === 0) {

        return this.__clone();

      }

      length = this.__length - start;

    } else {

      length |= 0;
      length = Math.min(length, this.__length - start);

    }

    length = Math.min(length, this.__length - start);

    if (length <= 0) {

      return this.__nullSeq();

    }

    return this.__slice(start, length);

  };

  Seq.prototype.polymerize = function(seq) {

    var seqLen = seq.__length;

    if (!(seq instanceof Seq)) {
      throw new Error('.polymerize requires valid sequence');
    }

    if (!this.__length) {
      return seq.__clone();
    }

    var offset = this.__length;
    var length = this.__length + seqLen;

    var max = length >>> 1;
    var odd = length & 1;

    var endPadding = (4 - (max + odd) % 4) % 4;
    var newBuffer = new ArrayBuffer(max + odd + endPadding + 4);
    var newArray = new Uint32Array(newBuffer, 4);

    var copyBuffer = this.__buffer;
    var copyArray = new Uint32Array(copyBuffer, 4);

    var seqBuffer = seq.__buffer;
    var seqArray = new Uint32Array(seqBuffer, 4);

    var copyPos = 0;
    var shift = (this.__length % 8) * 4;
    var shiftSeq = 32 - shift;

    for (var len = copyArray.length; copyPos < len; copyPos++) {
      newArray[copyPos] = copyArray[copyPos];
    }

    if (shift) {

      newArray[--copyPos] |= seqArray[0] << shift;

      for (var i = 0, len = seqArray.length; i < len; i++) {
        newArray[++copyPos] = (seqArray[i] >>> shiftSeq) | (seqArray[i + 1] << shift);
      }

    } else {

      for (var i = 0, len = seqArray.length; i < len; i++) {
        newArray[copyPos++] = seqArray[i];
      }

    }

    (new Uint32Array(newBuffer, 0, 1))[0] = length;

    return new Seq(this.__type).readBuffer(newBuffer);

  };

  Seq.prototype.insertion = function(seq, offset) {

    if (!(seq instanceof Seq)) {
      throw new Error('Insertion requires valid sequence');
    }

    offset |= 0;

    if (offset < 0) {
      offset = this.__length + offset;
    }

    offset = Math.min(offset, this.__length);

    return this.replicate(0, offset).polymerize(seq).polymerize(this.replicate(offset));

  };

  Seq.prototype.deletion = function(offset, count) {

    if (offset === undefined || count === undefined) {
      throw new Error('Must give valid offset and count for deletion');
    }

    offset |= 0;
    count |= 0;

    if (count === 0) {
      return this.__clone();
    }

    if (count < 0) {
      throw new Error('Invalid count for deletion');
    }

    if (offset < 0) {
      offset = this.__length + offset;
    }

    offset = Math.min(offset, this.__length);

    return this.replicate(0, offset).polymerize(this.replicate(offset + count));

  };

  Seq.prototype.repeat = function(count) {

    count |= 0;

    var copy = this.replicate();
    var base = new Seq(this.__type);

    if (count <= 0) {
      return base;
    }

    while(true) {
      if (count & 1) {
        base = base.polymerize(copy);
      }
      count >>>= 1;
      if (!count) {
        break;
      }
      copy = copy.polymerize(copy);
    }

    return base;

  };

  Seq.prototype.mask = function(seq) {

    if (!(seq instanceof Seq)) {
      throw new Error('Can only mask with valid sequence');
    }

    var newBuffer = this.__buffer.slice(0);
    var newArray = new Uint32Array(newBuffer, 4);
    var compareArray = new Uint32Array(seq.__buffer, 4);

    for (var i = 0, len = newArray.length; i < len; i++) {
      newArray[i] &= compareArray[i];
    }

    return new Seq(this.__type).readBuffer(newBuffer);

  };

  Seq.prototype.cover = function(seq) {

    if (!(seq instanceof Seq)) {
      throw new Error('Can only cover with valid sequence');
    }

    var newBuffer = this.__buffer.slice(0);
    var newArray = new Uint32Array(newBuffer, 4);
    var compareArray = new Uint32Array(seq.__buffer, 4);

    for (var i = 0, len = newArray.length; i < len; i++) {
      newArray[i] |= compareArray[i];
    }

    return new Seq(this.__type).readBuffer(newBuffer);

  };

  Seq.prototype.__nullSeq = function() {

    return new Seq(this.__type).readBuffer(new ArrayBuffer(4));

  };

  Seq.prototype.__clone = function() {

    return new Seq(this.__type).readBuffer(this.__buffer.slice(0));

  };

  Seq.prototype.__slice = function(start, length) {

    var max = length >>> 1;
    var odd = length & 1;

    var endPadding = (4 - (max + odd) % 4) % 4;
    var newBuffer = new ArrayBuffer(max + odd + endPadding + 4);
    var newArray = new Uint32Array(newBuffer, 4);

    var subBuffer = this.__buffer.slice(4 + (start >>> 1), 4 + (start >>> 1) + newBuffer.byteLength);
    var subInt32Length = subBuffer.byteLength >>> 2;
    var subArray = new Uint32Array(subBuffer, 0, subInt32Length);

    if (start & 1) {

      for (var i = 0, len = subArray.length; i < len; i++) {
        newArray[i] = (subArray[i] >>> 4) | (subArray[i + 1] << 28);
      }

      var remainder = subBuffer.byteLength - subArray.byteLength;
      if (remainder) {
        var remainderArray = new Uint8Array(newBuffer, 4 + (i << 2));
        var subRemainderArray = new Uint8Array(subBuffer, i << 2);
        if (newArray.length > 0) {
          newArray[i - 1] |= subRemainderArray[0] << 28;
        }
        for (var i = 0, len = subRemainderArray.length; i < len; i++) {
          remainderArray[i] = (subRemainderArray[i] >>> 4) | (subRemainderArray[i + 1] << 4);
        }
      }

    } else {

      for (var i = 0, len = subArray.length; i < len; i++) {
        newArray[i] = subArray[i];
      }

      var remainder = subArray.byteLength - subBuffer.byteLength;
      if (remainder) {
        var remainderArray = new Uint8Array(newBuffer, 4 + (i << 2));
        var subRemainderArray = new Uint8Array(subBuffer, i << 2);
        for (var i = 0, len = subRemainderArray.length; i < len; i++) {
          remainderArray[i] = subRemainderArray[i];
        }
      }

    }

    var clearShift = ((endPadding * 2) + odd) * 4;

    var clearOut = new Uint32Array(newBuffer, newBuffer.byteLength - 4);
    clearOut[0] = (clearOut[0] << clearShift) >>> clearShift;

    (new Uint32Array(newBuffer, 0, 1))[0] = length;

    return new Seq(this.__type).readBuffer(newBuffer);

  };

  Seq.prototype.content = function() {

    if (!this.__content) {

      var ntContentByte = makeArray(256);

      var buffer = this.__buffer;
      var dataArray = new Uint8Array(buffer);

      for(var i = 4; i < buffer.byteLength - this.__endPadding; i++) {
        ntContentByte[dataArray[i]]++;
      }

      var binToNt = binToNucleotide;
      var ntList = __nucleotideList;

      var ntContent = Object.create(null);
      for (var i = 0, len = ntList.length; i < len; i++) {
        ntContent[ntList[i]] = 0;
      }

      for (var i = 0, len = ntContentByte.length; i < len; i++) {
        if (ntContentByte[i]) {
          ntContent[binToNt(i & 0xF)] += ntContentByte[i];
          ntContent[binToNt(i >>> 4)] += ntContentByte[i];
        }
      }

      if (this.__length & 1) {
        ntContent['-']--;
      }

      if (this.__isRNA) {
        ntContent['U'] = ntContent['T'];
        delete ntContent['T'];
      }

      this.__content = ntContent;

    }

    var returnContent = Object.create(null);
    var keys = Object.keys(this.__content);
    for (var i = 0, len = keys.length; i < len; i++) {
      returnContent[keys[i]] = this.__content[keys[i]];
    }

    return returnContent;

  };

  Seq.prototype.fractionalContent = function() {

    if (!this.__fractionalContent) {

      var content = this.content();
      var nts = Object.keys(content);
      for (var i = 0, len = nts.length; i < len; i++) {
        content[nts[i]] = content[nts[i]] / this.__length;
      }

      this.__fractionalContent = content;

    }

    var returnContent = Object.create(null);
    var keys = Object.keys(this.__fractionalContent);
    for (var i = 0, len = keys.length; i < len; i++) {
      returnContent[keys[i]] = this.__fractionalContent[keys[i]];
    }

    return returnContent;

  };

  Seq.prototype.contentATGC = function() {

    if (!this.__contentATGC) {

      var ntToBin = nucleotideToBin;

      var content = this.content();
      var nts = Object.keys(content);
      var contentATGC = Object.create(null);
      contentATGC['A'] = 0;
      contentATGC['T'] = 0;
      contentATGC['G'] = 0;
      contentATGC['C'] = 0;

      var bits = 0;
      var nt;
      var ntBin;
      var n;
      var curContent;

      for (var i = 0, len = nts.length; i < len; i++) {
        nt = nts[i];
        n = ntToBin(nt);
        for (bits = 0; n; bits++) { n &= n - 1; }

        ntBin = ntToBin(nt);
        curContent = content[nts[i]] * (1 / bits);

        contentATGC['A'] += (((ntToBin('A') & ntBin) | 0) && curContent);
        contentATGC['T'] += (((ntToBin('T') & ntBin) | 0) && curContent);
        contentATGC['G'] += (((ntToBin('G') & ntBin) | 0) && curContent);
        contentATGC['C'] += (((ntToBin('C') & ntBin) | 0) && curContent);

      }

      if (this.__isRNA) {
        contentATGC['U'] = contentATGC['T'];
        delete contentATGC['T'];
      }

      this.__contentATGC = contentATGC;

    }

    var returnContent = Object.create(null);
    var keys = Object.keys(this.__contentATGC);
    for (var i = 0, len = keys.length; i < len; i++) {
      returnContent[keys[i]] = this.__contentATGC[keys[i]];
    }

    return returnContent;

  };

  Seq.prototype.fractionalContentATGC = function() {

    if (!this.__fractionalContentATGC) {

      var content = this.contentATGC();
      var nts = Object.keys(content);
      for (var i = 0, len = nts.length; i < len; i++) {
        content[nts[i]] = content[nts[i]] / this.__length;
      }

      this.__fractionalContentATGC = content;

    }

    var returnContent = Object.create(null);
    var keys = Object.keys(this.__fractionalContentATGC);
    for (var i = 0, len = keys.length; i < len; i++) {
      returnContent[keys[i]] = this.__fractionalContentATGC[keys[i]];
    }

    return returnContent;

  };

  Seq.prototype.translate = function(ntOffset, ntCount) {

    var binToAA = __12BitToAminoAcid;

    ntOffset |= 0;

    if (ntCount === undefined) {
      ntCount = this.__length - ntOffset;
    }

    ntCount |= 0;
    ntCount -= (ntCount % 3);

    var offset = (ntOffset >>> 1) + 4;
    var max = offset + (ntCount >>> 1) + (ntCount & 1);

    var dataArray = new Uint8Array(this.__buffer);

    var aminoAcids = makeArray(ntCount / 3);
    /**/
    var aa = 0;
    var lastByte, byte1, byte2, byte3;

    if ((ntOffset & 1) === 0) {

      for (var i = offset; i < max; i += 3) {

        var byte1 = dataArray[i];
        var byte2 = dataArray[i+1];
        var byte3 = dataArray[i+2];

        aminoAcids[aa++] = binToAA[byte1 | ((byte2 & 0xF) << 8)];
        aminoAcids[aa++] = binToAA[(byte3 << 4) | (byte2 >>> 4)];

      }

    } else {

      lastByte = dataArray[offset];

      for (var i = offset + 1; i < max; i += 3) {

        byte1 = dataArray[i];
        byte2 = dataArray[i+1];
        byte3 = dataArray[i+2];

        aminoAcids[aa++] = binToAA[(lastByte >> 4) | (byte1 << 4)];
        aminoAcids[aa++] = binToAA[byte2 | ((byte3 & 0xF) << 8)];

        lastByte = byte3;

      }

    }

    if (ntCount & 1) { aminoAcids.pop(); }

    return aminoAcids.join('');

  };

  Seq.prototype.translateFrame = function(frame, AAoffset, AAcount) {

    if (frame === undefined) {
      frame = 0;
    }

    if (frame !== 0 && frame !== 1 && frame !== 2) {
      throw new Error('Invalid translation frame, must be 0, 1 or 2.');
    }

    if (AAoffset === undefined) {
      return this.translate(frame);
    }

    if (AAcount === undefined) {
      return this.translate(frame + ((AAoffset | 0) * 3));
    }

    return this.translate(frame + ((AAoffset | 0) * 3), (AAcount * 3) | 0);

  };

  Seq.prototype.mapSequence = function(seq, offset) {

    if (!(seq instanceof Seq)) {
      throw new Error('.mapSequence requires valid Seq');
    }

    return new MatchMap(seq, this, offset);

  };

  /* MatchResult */

  function MatchResult(matchMap, pos, score) {

    Object.defineProperty(this, '__matchMap', {value: matchMap});

    this.position = pos;
    this.score = score;
    this.__align = null;

  };

  MatchResult.prototype.alignment = function() {

    if (!this.__align) {
      var map = this.__matchMap;
      if (this.position < 0) {
        this.__align = (new Nt.Seq())
          .read('-')
          .repeat(-this.position)
          .polymerize(map.__searchSpace.replicate(0, map.__query.__length + this.position));
      } else if (this.position + map.__query.__length > map.__searchSpace.__length) {
        this.__align = map.__searchSpace
          .replicate(this.position)
          .polymerize((new Nt.Seq()).read('-').repeat(map.__searchSpace.__length - this.position));
      } else {
        this.__align = map.__searchSpace.replicate(this.position, map.__query.__length);
      }
    }
    return this.__align;

  };

  MatchResult.prototype.alignmentMask = function() {

    return this.__matchMap.__query.mask(this.alignment());

  };

  MatchResult.prototype.alignmentCover = function() {

    return this.__matchMap.__query.cover(this.alignment());

  };

  /* MatchMap */

  function MatchMap(query, searchSpace, offset) {

    if (!(query instanceof Seq) || !(searchSpace instanceof Seq)) {
      throw new Error('MatchMap requires valid Seq');
    }

    this.__query = query.replicate();
    this.__searchSpace = searchSpace.replicate();
    this.__results = [];
    this.__orderedResults = [];
    this.__matchFrequencyData = null;
    this.__initialized = false;

    this.__offset = Math.max(0, offset | 0);
    this.__positionAdjustment = this.__offset - (this.__query.size() - 1);

    this.__debug = {
      searchTime: null,
      prepareTime: null,
      sortTime: null
    };

  };

  MatchMap.prototype.initialize = function(results) {

    this.__orderedResults = [];
    this.__matchFrequencyData = null;

    this.__initialized = true;

    var t = (new Date).valueOf();

    if (!results) {

      var dataArray = new Uint32Array(this.__execute(this.__query.__buffer, this.__searchSpace.__buffer));

      this.__debug.searchTime = (-t) + (t = (new Date).valueOf());

      var queryLen = this.__query.size();
      var searchLen = this.__searchSpace.size();
      var resultsLen = ((8 - (queryLen % 8)) % 8) + 1;
      var totalLen = searchLen + queryLen - 1;
      results = [].slice.call(dataArray, resultsLen, resultsLen + totalLen);

    }

    this.__results = results;
    this.__debug.prepareTime = (-t) + (t = (new Date).valueOf());

    return this;

  };

  MatchMap.prototype.sort = function() {

    if (!this.__initialized) {
      throw new Error('MatchMap must be initialized first.');
    }

    var t = new Date().valueOf();

    if (this.__debug.sortTime !== null) {
      return this;
    }

    var adjust = this.__positionAdjustment;
    this.__orderedResults = this.__results
      .map(function(v, i) { return {n: i + adjust, s: v}; })
      .sort(function(a, b) { return b.s - a.s; });
    this.__debug.sortTime = new Date().valueOf() - t;
    return this;

  };

  MatchMap.prototype.__calculate_p_match = function(query, searchSpace) {

    /*
      The approximate probability that two randomly chosen nucleotides
      from QUERY and SEARCH match each other
    */

    var queryContent = query.fractionalContentATGC();
    var searchSpaceContent = searchSpace.fractionalContentATGC();

    return (queryContent['A'] * searchSpaceContent['A']) +
      (queryContent['T'] * searchSpaceContent['T']) +
      (queryContent['G'] * searchSpaceContent['G']) +
      (queryContent['C'] * searchSpaceContent['C']);

  };

  MatchMap.prototype.results = function(offset, count) {

    if (!this.__initialized) {
      throw new Error('MatchMap must be initialized first.');
    }

    if (offset === undefined) {
      return this.__results.slice();
    }

    if (count === undefined) {
      return this.__results.slice(offset | 0);
    }

    return this.__results.slice(offset | 0, count | 0);

  };

  MatchMap.prototype.best = function() {

    if (!this.__initialized) {
      throw new Error('MatchMap must be initialized first.');
    }

    if (!this.__orderedResults.length) {
      throw new Error('MatchMap must be sorted first.');
    }

    var result = this.__orderedResults[0];
    return new MatchResult(this, result.n, result.s);

  };

  MatchMap.prototype.top = function(n) {

    if (!this.__initialized) {
      throw new Error('MatchMap must be initialized first.');
    }

    if (!this.__orderedResults.length) {
      throw new Error('MatchMap must be sorted first.');
    }

    var self = this;

    return this.__orderedResults.slice(0, n).map(function(result) {
      return new MatchResult(self, result.n, result.s);
    });

  };

  MatchMap.prototype.bottom = function(n) {

    if (!this.__initialized) {
      throw new Error('MatchMap must be initialized first.');
    }

    if (!this.__orderedResults.length) {
      throw new Error('MatchMap must be sorted first.');
    }

    var self = this;

    var adjust = this.__positionAdjustment;
    var len = this.__orderedResults.length;
    return this.__orderedResults.slice(this.__orderedResults.length - n, n).map(function(result) {
      return new MatchResult(self, result.n, result.s);
    });

  };

  /* Can be optimized with binary splitting */

  MatchMap.prototype.matchFrequencyData = function() {

    if (!this.__initialized) {
      throw new Error('MatchMap must be initialized first.');
    }

    if (!this.__orderedResults.length) {
      throw new Error('MatchMap must be sorted first.');
    }

    if (this.__matchFrequencyData) {
      return this.__matchFrequencyData;
    }

    var ordered = this.__orderedResults;
    var matchFrequencyData = makeArray(this.__query.size() + 1);

    var maxMatch = this.__query.size();
    var lastIndex = 0;
    var num;

    for (var i = 0, len = ordered.length; i < len; i++) {
      num = ordered[i].s;
      if (num < maxMatch) {
        matchFrequencyData[maxMatch] = i - lastIndex;
        lastIndex = i;
        maxMatch = num;
      }
      if (num === 0) {
        matchFrequencyData[0] = len - i;
        break;
      }
    }

    return (this.__matchFrequencyData = matchFrequencyData);

  };

  MatchMap.prototype.__countMatches = function(int, bitCount) {

    int |= int >>> 1;
    int |= int >>> 2;
    int &= 0x11111111;
    int |= int >>> 3;
    int |= int >>> 6;
    return bitCount[((int >>> 12) & 0xF0) | (int & 0xF)];

  };

  MatchMap.prototype.__execute = function(queryBuffer, searchSpaceBuffer) {

    var queryInts, spaceInts, queryIntsLength, spaceIntsLength,
      arrLen, mapBuffer, mapArray,
      A, B, A1, A2, T, cur, pos, move, i, k,
      adjustNeg, adjustPos,
      fnCountMatches, bitCount;

    queryInts = new Uint32Array(queryBuffer, 4);
    spaceInts = new Uint32Array(searchSpaceBuffer, 4);

    fnCountMatches = this.__countMatches;
    bitCount = __bitCount;

    queryIntsLength = queryInts.length|0;
    spaceIntsLength = spaceInts.length|0;

    arrLen = (queryIntsLength + spaceIntsLength) << 3;
    mapBuffer = new ArrayBuffer(4 * arrLen);
    mapArray = new Uint32Array(mapBuffer);

    for (k = 0|0; k < queryIntsLength; k++) {

      A = queryInts[k];
      cur = (queryIntsLength - k) << 3;

      for (i = 0|0; i < spaceIntsLength; i++) {
        (T = A & spaceInts[i]) && (mapArray[(i << 3) + cur] += fnCountMatches(T, bitCount));
      }

      A1 = A >>> 4;
      A2 = A << 4;

      adjustNeg = cur - 1;
      adjustPos = cur + 1;

      while(A1 || A2) {

        for (i = 0|0; i < spaceIntsLength; i++) {
          B = spaceInts[i];
          pos = (i << 3);

          (T = A1 & B) && (mapArray[pos + adjustNeg] += fnCountMatches(T, bitCount));
          (T = A2 & B) && (mapArray[pos + adjustPos] += fnCountMatches(T, bitCount));
        }

        A1 >>>= 4;
        A2 <<= 4;

        --adjustNeg;
        ++adjustPos;

      }

    }

    return mapBuffer;

  };

  return {
    Seq: Seq,
    MatchMap: MatchMap
  };

}();
```
