## Supplementary material for "Method for inferring migration routes from phylogenetic trees": Some route-trees: phylotree.js

```
!function(t){var e={};function n(r){if(e[r])return e[r].exports;var i=e[r]={i:r,l:!1,exports:{}};return t[r].call(i.exports,i,i.exports,n),i.l=!0,i.exports}n.m=t,n.c=e,n.d=function(t,e,r){n.o(t,e)||Object.defineProperty(t,e,{configurable:!1,enumerable:!0,get:r})},n.r=function(t){Object.defineProperty(t,"__esModule",{value:!0})},n.n=function(t){var e=t&&t.__esModule?function(){return t.default}:function(){return t};return n.d(e,"a",e),e},n.o=function(t,e){return Object.prototype.hasOwnProperty.call(t,e)},n.p="",n(n.s=60)}([function(t,e,n){(function(){var e,r,i,o,s,a,u,c,l,h,f,p,d={}.hasOwnProperty;p=n(1),f=p.isObject,h=p.isFunction,l=p.isEmpty,s=null,e=null,r=null,i=null,o=null,u=null,c=null,a=null,t.exports=function(){function t(t){this.parent=t,this.parent&&(this.options=this.parent.options,this.stringify=this.parent.stringify),this.children=[],s||(s=n(21),e=n(20),r=n(19),i=n(18),o=n(17),u=n(12),c=n(11),a=n(10))}return t.prototype.element=function(t,e,n){var r,i,o,s,a,u,c,p,y,g;if(u=null,null==e&&(e={}),e=e.valueOf(),f(e)||(n=(y=[e,n])[0],e=y[1]),null!=t&&(t=t.valueOf()),Array.isArray(t))for(o=0,c=t.length;o<c;o++)i=t[o],u=this.element(i);else if(h(t))u=this.element(t.apply());else if(f(t)){for(a in t)if(d.call(t,a))if(g=t[a],h(g)&&(g=g.apply()),f(g)&&l(g)&&(g=null),!this.options.ignoreDecorators&&this.stringify.convertAttKey&&0===a.indexOf(this.stringify.convertAttKey))u=this.attribute(a.substr(this.stringify.convertAttKey.length),g);else if(!this.options.separateArrayItems&&Array.isArray(g))for(s=0,p=g.length;s<p;s++)i=g[s],(r={})[a]=i,u=this.element(r);else f(g)?(u=this.element(a)).element(g):u=this.element(a,g)}else u=!this.options.ignoreDecorators&&this.stringify.convertTextKey&&0===t.indexOf(this.stringify.convertTextKey)?this.text(n):!this.options.ignoreDecorators&&this.stringify.convertCDataKey&&0===t.indexOf(this.stringify.convertCDataKey)?this.cdata(n):!this.options.ignoreDecorators&&this.stringify.convertCommentKey&&0===t.indexOf(this.stringify.convertCommentKey)?this.comment(n):!this.options.ignoreDecorators&&this.stringify.convertRawKey&&0===t.indexOf(this.stringify.convertRawKey)?this.raw(n):!this.options.ignoreDecorators&&this.stringify.convertPIKey&&0===t.indexOf(this.stringify.convertPIKey)?this.instruction(t.substr(this.stringify.convertPIKey.length),n):this.node(t,e,n);if(null==u)throw new Error("Could not create any elements with: "+t);return u},t.prototype.insertBefore=function(t,e,n){var r,i,o;if(this.isRoot)throw new Error("Cannot insert elements at root level");return i=this.parent.children.indexOf(this),o=this.parent.children.splice(i),r=this.parent.element(t,e,n),Array.prototype.push.apply(this.parent.children,o),r},t.prototype.insertAfter=function(t,e,n){var r,i,o;if(this.isRoot)throw new Error("Cannot insert elements at root level");return i=this.parent.children.indexOf(this),o=this.parent.children.splice(i+1),r=this.parent.element(t,e,n),Array.prototype.push.apply(this.parent.children,o),r},t.prototype.remove=function(){var t;if(this.isRoot)throw new Error("Cannot remove the root element");return t=this.parent.children.indexOf(this),[].splice.apply(this.parent.children,[t,t-t+1].concat([])),this.parent},t.prototype.node=function(t,e,n){var r,i;return null!=t&&(t=t.valueOf()),e||(e={}),e=e.valueOf(),f(e)||(n=(i=[e,n])[0],e=i[1]),r=new s(this,t,e),null!=n&&r.text(n),this.children.push(r),r},t.prototype.text=function(t){var e;return e=new c(this,t),this.children.push(e),this},t.prototype.cdata=function(t){var n;return n=new e(this,t),this.children.push(n),this},t.prototype.comment=function(t){var e;return e=new r(this,t),this.children.push(e),this},t.prototype.commentBefore=function(t){var e,n;return e=this.parent.children.indexOf(this),n=this.parent.children.splice(e),this.parent.comment(t),Array.prototype.push.apply(this.parent.children,n),this},t.prototype.commentAfter=function(t){var e,n;return e=this.parent.children.indexOf(this),n=this.parent.children.splice(e+1),this.parent.comment(t),Array.prototype.push.apply(this.parent.children,n),this},t.prototype.raw=function(t){var e;return e=new u(this,t),this.children.push(e),this},t.prototype.instruction=function(t,e){var n,r,i,o,s;if(null!=t&&(t=t.valueOf()),null!=e&&(e=e.valueOf()),Array.isArray(t))for(o=0,s=t.length;o<s;o++)n=t[o],this.instruction(n);else if(f(t))for(n in t)d.call(t,n)&&(r=t[n],this.instruction(n,r));else h(e)&&(e=e.apply()),i=new a(this,t,e),this.children.push(i);return this},t.prototype.instructionBefore=function(t,e){var n,r;return n=this.parent.children.indexOf(this),r=this.parent.children.splice(n),this.parent.instruction(t,e),Array.prototype.push.apply(this.parent.children,r),this},t.prototype.instructionAfter=function(t,e){var n,r;return n=this.parent.children.indexOf(this),r=this.parent.children.splice(n+1),this.parent.instruction(t,e),Array.prototype.push.apply(this.parent.children,r),this},t.prototype.declaration=function(t,e,n){var r,o;return r=this.document(),o=new i(r,t,e,n),r.children[0]instanceof i?r.children[0]=o:r.children.unshift(o),r.root()||r},t.prototype.doctype=function(t,e){var n,r,i,s,a,u,c,l,h;for(n=this.document(),r=new o(n,t,e),i=s=0,u=(l=n.children).length;s<u;i=++s)if(l[i]instanceof o)return n.children[i]=r,r;for(i=a=0,c=(h=n.children).length;a<c;i=++a)if(h[i].isRoot)return n.children.splice(i,0,r),r;return n.children.push(r),r},t.prototype.up=function(){if(this.isRoot)throw new Error("The root node has no parent. Use doc() if you need to get the document object.");return this.parent},t.prototype.root=function(){var t;for(t=this;t;){if(t.isDocument)return t.rootObject;if(t.isRoot)return t;t=t.parent}},t.prototype.document=function(){var t;for(t=this;t;){if(t.isDocument)return t;t=t.parent}},t.prototype.end=function(t){return this.document().end(t)},t.prototype.prev=function(){var t;if((t=this.parent.children.indexOf(this))<1)throw new Error("Already at the first node");return this.parent.children[t-1]},t.prototype.next=function(){var t;if(-1===(t=this.parent.children.indexOf(this))||t===this.parent.children.length-1)throw new Error("Already at the last node");return this.parent.children[t+1]},t.prototype.importDocument=function(t){var e;return(e=t.root().clone()).parent=this,e.isRoot=!1,this.children.push(e),this},t.prototype.ele=function(t,e,n){return this.element(t,e,n)},t.prototype.nod=function(t,e,n){return this.node(t,e,n)},t.prototype.txt=function(t){return this.text(t)},t.prototype.dat=function(t){return this.cdata(t)},t.prototype.com=function(t){return this.comment(t)},t.prototype.ins=function(t,e){return this.instruction(t,e)},t.prototype.doc=function(){return this.document()},t.prototype.dec=function(t,e,n){return this.declaration(t,e,n)},t.prototype.dtd=function(t,e){return this.doctype(t,e)},t.prototype.e=function(t,e,n){return this.element(t,e,n)},t.prototype.n=function(t,e,n){return this.node(t,e,n)},t.prototype.t=function(t){return this.text(t)},t.prototype.d=function(t){return this.cdata(t)},t.prototype.c=function(t){return this.comment(t)},t.prototype.r=function(t){return this.raw(t)},t.prototype.i=function(t,e){return this.instruction(t,e)},t.prototype.u=function(){return this.up()},t.prototype.importXMLBuilder=function(t){return this.importDocument(t)},t}()}).call(this)},function(t,e){(function(){var e,n,r,i,o,s,a=[].slice,u={}.hasOwnProperty;e=function(){var t,e,n,r,o,s;if(s=arguments[0],o=2<=arguments.length?a.call(arguments,1):[],i(Object.assign))Object.assign.apply(null,arguments);else for(t=0,n=o.length;t<n;t++)if(null!=(r=o[t]))for(e in r)u.call(r,e)&&(s[e]=r[e]);return s},i=function(t){return!!t&&"[object Function]"===Object.prototype.toString.call(t)},o=function(t){var e;return!!t&&("function"==(e=typeof t)||"object"===e)},n=function(t){return i(Array.isArray)?Array.isArray(t):"[object Array]"===Object.prototype.toString.call(t)},r=function(t){var e;if(n(t))return!t.length;for(e in t)if(u.call(t,e))return!1;return!0},s=function(t){var e,n;return o(t)&&(n=Object.getPrototypeOf(t))&&(e=n.constructor)&&"function"==typeof e&&e instanceof e&&Function.prototype.toString.call(e)===Function.prototype.toString.call(Object)},t.exports.assign=e,t.exports.isFunction=i,t.exports.isObject=o,t.exports.isArray=n,t.exports.isEmpty=r,t.exports.isPlainObject=s}).call(this)},function(t,e,n){"use strict";var r=n(7),i=Object.keys||function(t){var e=[];for(var n in t)e.push(n);return e};t.exports=h;var o=n(5);o.inherits=n(3);var s=n(33),a=n(23);o.inherits(h,s);for(var u=i(a.prototype),c=0;c<u.length;c++){var l=u[c];h.prototype[l]||(h.prototype[l]=a.prototype[l])}function h(t){if(!(this instanceof h))return new h(t);s.call(this,t),a.call(this,t),t&&!1===t.readable&&(this.readable=!1),t&&!1===t.writable&&(this.writable=!1),this.allowHalfOpen=!0,t&&!1===t.allowHalfOpen&&(this.allowHalfOpen=!1),this.once("end",f)}function f(){this.allowHalfOpen||this._writableState.ended||r.nextTick(p,this)}function p(t){t.end()}Object.defineProperty(h.prototype,"writableHighWaterMark",{enumerable:!1,get:function(){return this._writableState.highWaterMark}}),Object.defineProperty(h.prototype,"destroyed",{get:function(){return void 0!==this._readableState&&void 0!==this._writableState&&(this._readableState.destroyed&&this._writableState.destroyed)},set:function(t){void 0!==this._readableState&&void 0!==this._writableState&&(this._readableState.destroyed=t,this._writableState.destroyed=t)}}),h.prototype._destroy=function(t,e){this.push(null),this.end(),r.nextTick(e,t)}},function(t,e){"function"==typeof Object.create?t.exports=function(t,e){t.super_=e,t.prototype=Object.create(e.prototype,{constructor:{value:t,enumerable:!1,writable:!0,configurable:!0}})}:t.exports=function(t,e){t.super_=e;var n=function(){};n.prototype=e.prototype,t.prototype=new n,t.prototype.constructor=t}},function(t,e){var n;n=function(){return this}();try{n=n||Function("return this")()||(0,eval)("this")}catch(t){"object"==typeof window&&(n=window)}t.exports=n},function(t,e,n){(function(t){function n(t){return Object.prototype.toString.call(t)}e.isArray=function(t){return Array.isArray?Array.isArray(t):"[object Array]"===n(t)},e.isBoolean=function(t){return"boolean"==typeof t},e.isNull=function(t){return null===t},e.isNullOrUndefined=function(t){return null==t},e.isNumber=function(t){return"number"==typeof t},e.isString=function(t){return"string"==typeof t},e.isSymbol=function(t){return"symbol"==typeof t},e.isUndefined=function(t){return void 0===t},e.isRegExp=function(t){return"[object RegExp]"===n(t)},e.isObject=function(t){return"object"==typeof t&&null!==t},e.isDate=function(t){return"[object Date]"===n(t)},e.isError=function(t){return"[object Error]"===n(t)||t instanceof Error},e.isFunction=function(t){return"function"==typeof t},e.isPrimitive=function(t){return null===t||"boolean"==typeof t||"number"==typeof t||"string"==typeof t||"symbol"==typeof t||void 0===t},e.isBuffer=t.isBuffer}).call(this,n(25).Buffer)},function(t,e,n){var r=n(25),i=r.Buffer;function o(t,e){for(var n in t)e[n]=t[n]}function s(t,e,n){return i(t,e,n)}i.from&&i.alloc&&i.allocUnsafe&&i.allocUnsafeSlow?t.exports=r:(o(r,e),e.Buffer=s),o(i,s),s.from=function(t,e,n){if("number"==typeof t)throw new TypeError("Argument must not be a number");return i(t,e,n)},s.alloc=function(t,e,n){if("number"!=typeof t)throw new TypeError("Argument must be a number");var r=i(t);return void 0!==e?"string"==typeof n?r.fill(e,n):r.fill(e):r.fill(0),r},s.allocUnsafe=function(t){if("number"!=typeof t)throw new TypeError("Argument must be a number");return i(t)},s.allocUnsafeSlow=function(t){if("number"!=typeof t)throw new TypeError("Argument must be a number");return r.SlowBuffer(t)}},function(t,e,n){"use strict";(function(e){!e.version||0===e.version.indexOf("v0.")||0===e.version.indexOf("v1.")&&0!==e.version.indexOf("v1.8.")?t.exports={nextTick:function(t,n,r,i){if("function"!=typeof t)throw new TypeError('"callback" argument must be a function');var o,s,a=arguments.length;switch(a){case 0:case 1:return e.nextTick(t);case 2:return e.nextTick(function(){t.call(null,n)});case 3:return e.nextTick(function(){t.call(null,n,r)});case 4:return e.nextTick(function(){t.call(null,n,r,i)});default:for(o=new Array(a-1),s=0;s<o.length;)o[s++]=arguments[s];return e.nextTick(function(){t.apply(null,o)})}}}:t.exports=e}).call(this,n(8))},function(t,e){var n,r,i=t.exports={};function o(){throw new Error("setTimeout has not been defined")}function s(){throw new Error("clearTimeout has not been defined")}function a(t){if(n===setTimeout)return setTimeout(t,0);if((n===o||!n)&&setTimeout)return n=setTimeout,setTimeout(t,0);try{return n(t,0)}catch(e){try{return n.call(null,t,0)}catch(e){return n.call(this,t,0)}}}!function(){try{n="function"==typeof setTimeout?setTimeout:o}catch(t){n=o}try{r="function"==typeof clearTimeout?clearTimeout:s}catch(t){r=s}}();var u,c=[],l=!1,h=-1;function f(){l&&u&&(l=!1,u.length?c=u.concat(c):h=-1,c.length&&p())}function p(){if(!l){var t=a(f);l=!0;for(var e=c.length;e;){for(u=c,c=[];++h<e;)u&&u[h].run();h=-1,e=c.length}u=null,l=!1,function(t){if(r===clearTimeout)return clearTimeout(t);if((r===s||!r)&&clearTimeout)return r=clearTimeout,clearTimeout(t);try{r(t)}catch(e){try{return r.call(null,t)}catch(e){return r.call(this,t)}}}(t)}}function d(t,e){this.fun=t,this.array=e}function y(){}i.nextTick=function(t){var e=new Array(arguments.length-1);if(arguments.length>1)for(var n=1;n<arguments.length;n++)e[n-1]=arguments[n];c.push(new d(t,e)),1!==c.length||l||a(p)},d.prototype.run=function(){this.fun.apply(null,this.array)},i.title="browser",i.browser=!0,i.env={},i.argv=[],i.version="",i.versions={},i.on=y,i.addListener=y,i.once=y,i.off=y,i.removeListener=y,i.removeAllListeners=y,i.emit=y,i.prependListener=y,i.prependOnceListener=y,i.listeners=function(t){return[]},i.binding=function(t){throw new Error("process.binding is not supported")},i.cwd=function(){return"/"},i.chdir=function(t){throw new Error("process.chdir is not supported")},i.umask=function(){return 0}},function(t,e){function n(){this._events=this._events||{},this._maxListeners=this._maxListeners||void 0}function r(t){return"function"==typeof t}function i(t){return"object"==typeof t&&null!==t}function o(t){return void 0===t}t.exports=n,n.EventEmitter=n,n.prototype._events=void 0,n.prototype._maxListeners=void 0,n.defaultMaxListeners=10,n.prototype.setMaxListeners=function(t){if("number"!=typeof t||t<0||isNaN(t))throw TypeError("n must be a positive number");return this._maxListeners=t,this},n.prototype.emit=function(t){var e,n,s,a,u,c;if(this._events||(this._events={}),"error"===t&&(!this._events.error||i(this._events.error)&&!this._events.error.length)){if((e=arguments[1])instanceof Error)throw e;var l=new Error('Uncaught, unspecified "error" event. ('+e+")");throw l.context=e,l}if(o(n=this._events[t]))return!1;if(r(n))switch(arguments.length){case 1:n.call(this);break;case 2:n.call(this,arguments[1]);break;case 3:n.call(this,arguments[1],arguments[2]);break;default:a=Array.prototype.slice.call(arguments,1),n.apply(this,a)}else if(i(n))for(a=Array.prototype.slice.call(arguments,1),s=(c=n.slice()).length,u=0;u<s;u++)c[u].apply(this,a);return!0},n.prototype.addListener=function(t,e){var s;if(!r(e))throw TypeError("listener must be a function");return this._events||(this._events={}),this._events.newListener&&this.emit("newListener",t,r(e.listener)?e.listener:e),this._events[t]?i(this._events[t])?this._events[t].push(e):this._events[t]=[this._events[t],e]:this._events[t]=e,i(this._events[t])&&!this._events[t].warned&&(s=o(this._maxListeners)?n.defaultMaxListeners:this._maxListeners)&&s>0&&this._events[t].length>s&&(this._events[t].warned=!0,console.error("(node) warning: possible EventEmitter memory leak detected. %d listeners added. Use emitter.setMaxListeners() to increase limit.",this._events[t].length),"function"==typeof console.trace&&console.trace()),this},n.prototype.on=n.prototype.addListener,n.prototype.once=function(t,e){if(!r(e))throw TypeError("listener must be a function");var n=!1;function i(){this.removeListener(t,i),n||(n=!0,e.apply(this,arguments))}return i.listener=e,this.on(t,i),this},n.prototype.removeListener=function(t,e){var n,o,s,a;if(!r(e))throw TypeError("listener must be a function");if(!this._events||!this._events[t])return this;if(s=(n=this._events[t]).length,o=-1,n===e||r(n.listener)&&n.listener===e)delete this._events[t],this._events.removeListener&&this.emit("removeListener",t,e);else if(i(n)){for(a=s;a-- >0;)if(n[a]===e||n[a].listener&&n[a].listener===e){o=a;break}if(o<0)return this;1===n.length?(n.length=0,delete this._events[t]):n.splice(o,1),this._events.removeListener&&this.emit("removeListener",t,e)}return this},n.prototype.removeAllListeners=function(t){var e,n;if(!this._events)return this;if(!this._events.removeListener)return 0===arguments.length?this._events={}:this._events[t]&&delete this._events[t],this;if(0===arguments.length){for(e in this._events)"removeListener"!==e&&this.removeAllListeners(e);return this.removeAllListeners("removeListener"),this._events={},this}if(r(n=this._events[t]))this.removeListener(t,n);else if(n)for(;n.length;)this.removeListener(t,n[n.length-1]);return delete this._events[t],this},n.prototype.listeners=function(t){return this._events&&this._events[t]?r(this._events[t])?[this._events[t]]:this._events[t].slice():[]},n.prototype.listenerCount=function(t){if(this._events){var e=this._events[t];if(r(e))return 1;if(e)return e.length}return 0},n.listenerCount=function(t,e){return t.listenerCount(e)}},function(t,e,n){(function(){var e,r={}.hasOwnProperty;e=n(0),t.exports=function(t){function n(t,e,r){if(n.__super__.constructor.call(this,t),null==e)throw new Error("Missing instruction target");this.target=this.stringify.insTarget(e),r&&(this.value=this.stringify.insValue(r))}return function(t,e){for(var n in e)r.call(e,n)&&(t[n]=e[n]);function i(){this.constructor=t}i.prototype=e.prototype,t.prototype=new i,t.__super__=e.prototype}(n,e),n.prototype.clone=function(){return Object.create(this)},n.prototype.toString=function(t){return this.options.writer.set(t).processingInstruction(this)},n}()}).call(this)},function(t,e,n){(function(){var e,r={}.hasOwnProperty;e=n(0),t.exports=function(t){function n(t,e){if(n.__super__.constructor.call(this,t),null==e)throw new Error("Missing element text");this.value=this.stringify.eleText(e)}return function(t,e){for(var n in e)r.call(e,n)&&(t[n]=e[n]);function i(){this.constructor=t}i.prototype=e.prototype,t.prototype=new i,t.__super__=e.prototype}(n,e),n.prototype.clone=function(){return Object.create(this)},n.prototype.toString=function(t){return this.options.writer.set(t).text(this)},n}()}).call(this)},function(t,e,n){(function(){var e,r={}.hasOwnProperty;e=n(0),t.exports=function(t){function n(t,e){if(n.__super__.constructor.call(this,t),null==e)throw new Error("Missing raw text");this.value=this.stringify.raw(e)}return function(t,e){for(var n in e)r.call(e,n)&&(t[n]=e[n]);function i(){this.constructor=t}i.prototype=e.prototype,t.prototype=new i,t.__super__=e.prototype}(n,e),n.prototype.clone=function(){return Object.create(this)},n.prototype.toString=function(t){return this.options.writer.set(t).raw(this)},n}()}).call(this)},function(t,e,n){(function(){var e,r={}.hasOwnProperty;e=n(0),t.exports=function(t){function n(t,e,r){if(n.__super__.constructor.call(this,t),null==e)throw new Error("Missing notation name");if(!r.pubID&&!r.sysID)throw new Error("Public or system identifiers are required for an external entity");this.name=this.stringify.eleName(e),null!=r.pubID&&(this.pubID=this.stringify.dtdPubID(r.pubID)),null!=r.sysID&&(this.sysID=this.stringify.dtdSysID(r.sysID))}return function(t,e){for(var n in e)r.call(e,n)&&(t[n]=e[n]);function i(){this.constructor=t}i.prototype=e.prototype,t.prototype=new i,t.__super__=e.prototype}(n,e),n.prototype.toString=function(t){return this.options.writer.set(t).dtdNotation(this)},n}()}).call(this)},function(t,e,n){(function(){var e,r={}.hasOwnProperty;e=n(0),t.exports=function(t){function n(t,e,r){if(n.__super__.constructor.call(this,t),null==e)throw new Error("Missing DTD element name");r||(r="(#PCDATA)"),Array.isArray(r)&&(r="("+r.join(",")+")"),this.name=this.stringify.eleName(e),this.value=this.stringify.dtdElementValue(r)}return function(t,e){for(var n in e)r.call(e,n)&&(t[n]=e[n]);function i(){this.constructor=t}i.prototype=e.prototype,t.prototype=new i,t.__super__=e.prototype}(n,e),n.prototype.toString=function(t){return this.options.writer.set(t).dtdElement(this)},n}()}).call(this)},function(t,e,n){(function(){var e,r,i={}.hasOwnProperty;r=n(1).isObject,e=n(0),t.exports=function(t){function n(t,e,i,o){if(n.__super__.constructor.call(this,t),null==i)throw new Error("Missing entity name");if(null==o)throw new Error("Missing entity value");if(this.pe=!!e,this.name=this.stringify.eleName(i),r(o)){if(!o.pubID&&!o.sysID)throw new Error("Public and/or system identifiers are required for an external entity");if(o.pubID&&!o.sysID)throw new Error("System identifier is required for a public external entity");if(null!=o.pubID&&(this.pubID=this.stringify.dtdPubID(o.pubID)),null!=o.sysID&&(this.sysID=this.stringify.dtdSysID(o.sysID)),null!=o.nData&&(this.nData=this.stringify.dtdNData(o.nData)),this.pe&&this.nData)throw new Error("Notation declaration is not allowed in a parameter entity")}else this.value=this.stringify.dtdEntityValue(o)}return function(t,e){for(var n in e)i.call(e,n)&&(t[n]=e[n]);function r(){this.constructor=t}r.prototype=e.prototype,t.prototype=new r,t.__super__=e.prototype}(n,e),n.prototype.toString=function(t){return this.options.writer.set(t).dtdEntity(this)},n}()}).call(this)},function(t,e,n){(function(){var e,r={}.hasOwnProperty;e=n(0),t.exports=function(t){function n(t,e,r,i,o,s){if(n.__super__.constructor.call(this,t),null==e)throw new Error("Missing DTD element name");if(null==r)throw new Error("Missing DTD attribute name");if(!i)throw new Error("Missing DTD attribute type");if(!o)throw new Error("Missing DTD attribute default");if(0!==o.indexOf("#")&&(o="#"+o),!o.match(/^(#REQUIRED|#IMPLIED|#FIXED|#DEFAULT)$/))throw new Error("Invalid default value type; expected: #REQUIRED, #IMPLIED, #FIXED or #DEFAULT");if(s&&!o.match(/^(#FIXED|#DEFAULT)$/))throw new Error("Default value only applies to #FIXED or #DEFAULT");this.elementName=this.stringify.eleName(e),this.attributeName=this.stringify.attName(r),this.attributeType=this.stringify.dtdAttType(i),this.defaultValue=this.stringify.dtdAttDefault(s),this.defaultValueType=o}return function(t,e){for(var n in e)r.call(e,n)&&(t[n]=e[n]);function i(){this.constructor=t}i.prototype=e.prototype,t.prototype=new i,t.__super__=e.prototype}(n,e),n.prototype.toString=function(t){return this.options.writer.set(t).dtdAttList(this)},n}()}).call(this)},function(t,e,n){(function(){var e,r,i,o,s,a,u={}.hasOwnProperty;a=n(1).isObject,s=n(0),e=n(16),i=n(15),r=n(14),o=n(13),t.exports=function(t){function n(t,e,r){var i,o;n.__super__.constructor.call(this,t),this.documentObject=t,a(e)&&(e=(i=e).pubID,r=i.sysID),null==r&&(r=(o=[e,r])[0],e=o[1]),null!=e&&(this.pubID=this.stringify.dtdPubID(e)),null!=r&&(this.sysID=this.stringify.dtdSysID(r))}return function(t,e){for(var n in e)u.call(e,n)&&(t[n]=e[n]);function r(){this.constructor=t}r.prototype=e.prototype,t.prototype=new r,t.__super__=e.prototype}(n,s),n.prototype.element=function(t,e){var n;return n=new r(this,t,e),this.children.push(n),this},n.prototype.attList=function(t,n,r,i,o){var s;return s=new e(this,t,n,r,i,o),this.children.push(s),this},n.prototype.entity=function(t,e){var n;return n=new i(this,!1,t,e),this.children.push(n),this},n.prototype.pEntity=function(t,e){var n;return n=new i(this,!0,t,e),this.children.push(n),this},n.prototype.notation=function(t,e){var n;return n=new o(this,t,e),this.children.push(n),this},n.prototype.toString=function(t){return this.options.writer.set(t).docType(this)},n.prototype.ele=function(t,e){return this.element(t,e)},n.prototype.att=function(t,e,n,r,i){return this.attList(t,e,n,r,i)},n.prototype.ent=function(t,e){return this.entity(t,e)},n.prototype.pent=function(t,e){return this.pEntity(t,e)},n.prototype.not=function(t,e){return this.notation(t,e)},n.prototype.up=function(){return this.root()||this.documentObject},n}()}).call(this)},function(t,e,n){(function(){var e,r,i={}.hasOwnProperty;r=n(1).isObject,e=n(0),t.exports=function(t){function n(t,e,i,o){var s;n.__super__.constructor.call(this,t),r(e)&&(e=(s=e).version,i=s.encoding,o=s.standalone),e||(e="1.0"),this.version=this.stringify.xmlVersion(e),null!=i&&(this.encoding=this.stringify.xmlEncoding(i)),null!=o&&(this.standalone=this.stringify.xmlStandalone(o))}return function(t,e){for(var n in e)i.call(e,n)&&(t[n]=e[n]);function r(){this.constructor=t}r.prototype=e.prototype,t.prototype=new r,t.__super__=e.prototype}(n,e),n.prototype.toString=function(t){return this.options.writer.set(t).declaration(this)},n}()}).call(this)},function(t,e,n){(function(){var e,r={}.hasOwnProperty;e=n(0),t.exports=function(t){function n(t,e){if(n.__super__.constructor.call(this,t),null==e)throw new Error("Missing comment text");this.text=this.stringify.comment(e)}return function(t,e){for(var n in e)r.call(e,n)&&(t[n]=e[n]);function i(){this.constructor=t}i.prototype=e.prototype,t.prototype=new i,t.__super__=e.prototype}(n,e),n.prototype.clone=function(){return Object.create(this)},n.prototype.toString=function(t){return this.options.writer.set(t).comment(this)},n}()}).call(this)},function(t,e,n){(function(){var e,r={}.hasOwnProperty;e=n(0),t.exports=function(t){function n(t,e){if(n.__super__.constructor.call(this,t),null==e)throw new Error("Missing CDATA text");this.text=this.stringify.cdata(e)}return function(t,e){for(var n in e)r.call(e,n)&&(t[n]=e[n]);function i(){this.constructor=t}i.prototype=e.prototype,t.prototype=new i,t.__super__=e.prototype}(n,e),n.prototype.clone=function(){return Object.create(this)},n.prototype.toString=function(t){return this.options.writer.set(t).cdata(this)},n}()}).call(this)},function(t,e,n){(function(){var e,r,i,o,s,a={}.hasOwnProperty;s=n(1),o=s.isObject,i=s.isFunction,r=n(0),e=n(37),t.exports=function(t){function n(t,e,r){if(n.__super__.constructor.call(this,t),null==e)throw new Error("Missing element name");this.name=this.stringify.eleName(e),this.attributes={},null!=r&&this.attribute(r),t.isDocument&&(this.isRoot=!0,this.documentObject=t,t.rootObject=this)}return function(t,e){for(var n in e)a.call(e,n)&&(t[n]=e[n]);function r(){this.constructor=t}r.prototype=e.prototype,t.prototype=new r,t.__super__=e.prototype}(n,r),n.prototype.clone=function(){var t,e,n,r;for(e in(n=Object.create(this)).isRoot&&(n.documentObject=null),n.attributes={},r=this.attributes)a.call(r,e)&&(t=r[e],n.attributes[e]=t.clone());return n.children=[],this.children.forEach(function(t){var e;return(e=t.clone()).parent=n,n.children.push(e)}),n},n.prototype.attribute=function(t,n){var r,s;if(null!=t&&(t=t.valueOf()),o(t))for(r in t)a.call(t,r)&&(s=t[r],this.attribute(r,s));else i(n)&&(n=n.apply()),this.options.skipNullAttributes&&null==n||(this.attributes[t]=new e(this,t,n));return this},n.prototype.removeAttribute=function(t){var e,n,r;if(null==t)throw new Error("Missing attribute name");if(t=t.valueOf(),Array.isArray(t))for(n=0,r=t.length;n<r;n++)e=t[n],delete this.attributes[e];else delete this.attributes[t];return this},n.prototype.toString=function(t){return this.options.writer.set(t).element(this)},n.prototype.att=function(t,e){return this.attribute(t,e)},n.prototype.a=function(t,e){return this.attribute(t,e)},n}()}).call(this)},function(t,e,n){"use strict";var r=n(6).Buffer,i=r.isEncoding||function(t){switch((t=""+t)&&t.toLowerCase()){case"hex":case"utf8":case"utf-8":case"ascii":case"binary":case"base64":case"ucs2":case"ucs-2":case"utf16le":case"utf-16le":case"raw":return!0;default:return!1}};function o(t){var e;switch(this.encoding=function(t){var e=function(t){if(!t)return"utf8";for(var e;;)switch(t){case"utf8":case"utf-8":return"utf8";case"ucs2":case"ucs-2":case"utf16le":case"utf-16le":return"utf16le";case"latin1":case"binary":return"latin1";case"base64":case"ascii":case"hex":return t;default:if(e)return;t=(""+t).toLowerCase(),e=!0}}(t);if("string"!=typeof e&&(r.isEncoding===i||!i(t)))throw new Error("Unknown encoding: "+t);return e||t}(t),this.encoding){case"utf16le":this.text=u,this.end=c,e=4;break;case"utf8":this.fillLast=a,e=4;break;case"base64":this.text=l,this.end=h,e=3;break;default:return this.write=f,void(this.end=p)}this.lastNeed=0,this.lastTotal=0,this.lastChar=r.allocUnsafe(e)}function s(t){return t<=127?0:t>>5==6?2:t>>4==14?3:t>>3==30?4:t>>6==2?-1:-2}function a(t){var e=this.lastTotal-this.lastNeed,n=function(t,e,n){if(128!=(192&e[0]))return t.lastNeed=0,"�";if(t.lastNeed>1&&e.length>1){if(128!=(192&e[1]))return t.lastNeed=1,"�";if(t.lastNeed>2&&e.length>2&&128!=(192&e[2]))return t.lastNeed=2,"�"}}(this,t);return void 0!==n?n:this.lastNeed<=t.length?(t.copy(this.lastChar,e,0,this.lastNeed),this.lastChar.toString(this.encoding,0,this.lastTotal)):(t.copy(this.lastChar,e,0,t.length),void(this.lastNeed-=t.length))}function u(t,e){if((t.length-e)%2==0){var n=t.toString("utf16le",e);if(n){var r=n.charCodeAt(n.length-1);if(r>=55296&&r<=56319)return this.lastNeed=2,this.lastTotal=4,this.lastChar[0]=t[t.length-2],this.lastChar[1]=t[t.length-1],n.slice(0,-1)}return n}return this.lastNeed=1,this.lastTotal=2,this.lastChar[0]=t[t.length-1],t.toString("utf16le",e,t.length-1)}function c(t){var e=t&&t.length?this.write(t):"";if(this.lastNeed){var n=this.lastTotal-this.lastNeed;return e+this.lastChar.toString("utf16le",0,n)}return e}function l(t,e){var n=(t.length-e)%3;return 0===n?t.toString("base64",e):(this.lastNeed=3-n,this.lastTotal=3,1===n?this.lastChar[0]=t[t.length-1]:(this.lastChar[0]=t[t.length-2],this.lastChar[1]=t[t.length-1]),t.toString("base64",e,t.length-n))}function h(t){var e=t&&t.length?this.write(t):"";return this.lastNeed?e+this.lastChar.toString("base64",0,3-this.lastNeed):e}function f(t){return t.toString(this.encoding)}function p(t){return t&&t.length?this.write(t):""}e.StringDecoder=o,o.prototype.write=function(t){if(0===t.length)return"";var e,n;if(this.lastNeed){if(void 0===(e=this.fillLast(t)))return"";n=this.lastNeed,this.lastNeed=0}else n=0;return n<t.length?e?e+this.text(t,n):this.text(t,n):e||""},o.prototype.end=function(t){var e=t&&t.length?this.write(t):"";return this.lastNeed?e+"�":e},o.prototype.text=function(t,e){var n=function(t,e,n){var r=e.length-1;if(r<n)return 0;var i=s(e[r]);if(i>=0)return i>0&&(t.lastNeed=i-1),i;if(--r<n||-2===i)return 0;if((i=s(e[r]))>=0)return i>0&&(t.lastNeed=i-2),i;if(--r<n||-2===i)return 0;if((i=s(e[r]))>=0)return i>0&&(2===i?i=0:t.lastNeed=i-3),i;return 0}(this,t,e);if(!this.lastNeed)return t.toString("utf8",e);this.lastTotal=n;var r=t.length-(n-this.lastNeed);return t.copy(this.lastChar,0,r),t.toString("utf8",e,r)},o.prototype.fillLast=function(t){if(this.lastNeed<=t.length)return t.copy(this.lastChar,this.lastTotal-this.lastNeed,0,this.lastNeed),this.lastChar.toString(this.encoding,0,this.lastTotal);t.copy(this.lastChar,this.lastTotal-this.lastNeed,0,t.length),this.lastNeed-=t.length}},function(t,e,n){"use strict";(function(e,r,i){var o=n(7);function s(t){var e=this;this.next=null,this.entry=null,this.finish=function(){!function(t,e,n){var r=t.entry;t.entry=null;for(;r;){var i=r.callback;e.pendingcb--,i(n),r=r.next}e.corkedRequestsFree?e.corkedRequestsFree.next=t:e.corkedRequestsFree=t}(e,t)}}t.exports=_;var a,u=!e.browser&&["v0.10","v0.9."].indexOf(e.version.slice(0,5))>-1?r:o.nextTick;_.WritableState=m;var c=n(5);c.inherits=n(3);var l={deprecate:n(44)},h=n(32),f=n(6).Buffer,p=i.Uint8Array||function(){};var d,y=n(31);function g(){}function m(t,e){a=a||n(2),t=t||{};var r=e instanceof a;this.objectMode=!!t.objectMode,r&&(this.objectMode=this.objectMode||!!t.writableObjectMode);var i=t.highWaterMark,c=t.writableHighWaterMark,l=this.objectMode?16:16384;this.highWaterMark=i||0===i?i:r&&(c||0===c)?c:l,this.highWaterMark=Math.floor(this.highWaterMark),this.finalCalled=!1,this.needDrain=!1,this.ending=!1,this.ended=!1,this.finished=!1,this.destroyed=!1;var h=!1===t.decodeStrings;this.decodeStrings=!h,this.defaultEncoding=t.defaultEncoding||"utf8",this.length=0,this.writing=!1,this.corked=0,this.sync=!0,this.bufferProcessing=!1,this.onwrite=function(t){!function(t,e){var n=t._writableState,r=n.sync,i=n.writecb;if(function(t){t.writing=!1,t.writecb=null,t.length-=t.writelen,t.writelen=0}(n),e)!function(t,e,n,r,i){--e.pendingcb,n?(o.nextTick(i,r),o.nextTick(x,t,e),t._writableState.errorEmitted=!0,t.emit("error",r)):(i(r),t._writableState.errorEmitted=!0,t.emit("error",r),x(t,e))}(t,n,r,e,i);else{var s=E(n);s||n.corked||n.bufferProcessing||!n.bufferedRequest||v(t,n),r?u(w,t,n,s,i):w(t,n,s,i)}}(e,t)},this.writecb=null,this.writelen=0,this.bufferedRequest=null,this.lastBufferedRequest=null,this.pendingcb=0,this.prefinished=!1,this.errorEmitted=!1,this.bufferedRequestCount=0,this.corkedRequestsFree=new s(this)}function _(t){if(a=a||n(2),!(d.call(_,this)||this instanceof a))return new _(t);this._writableState=new m(t,this),this.writable=!0,t&&("function"==typeof t.write&&(this._write=t.write),"function"==typeof t.writev&&(this._writev=t.writev),"function"==typeof t.destroy&&(this._destroy=t.destroy),"function"==typeof t.final&&(this._final=t.final)),h.call(this)}function b(t,e,n,r,i,o,s){e.writelen=r,e.writecb=s,e.writing=!0,e.sync=!0,n?t._writev(i,e.onwrite):t._write(i,o,e.onwrite),e.sync=!1}function w(t,e,n,r){n||function(t,e){0===e.length&&e.needDrain&&(e.needDrain=!1,t.emit("drain"))}(t,e),e.pendingcb--,r(),x(t,e)}function v(t,e){e.bufferProcessing=!0;var n=e.bufferedRequest;if(t._writev&&n&&n.next){var r=e.bufferedRequestCount,i=new Array(r),o=e.corkedRequestsFree;o.entry=n;for(var a=0,u=!0;n;)i[a]=n,n.isBuf||(u=!1),n=n.next,a+=1;i.allBuffers=u,b(t,e,!0,e.length,i,"",o.finish),e.pendingcb++,e.lastBufferedRequest=null,o.next?(e.corkedRequestsFree=o.next,o.next=null):e.corkedRequestsFree=new s(e),e.bufferedRequestCount=0}else{for(;n;){var c=n.chunk,l=n.encoding,h=n.callback;if(b(t,e,!1,e.objectMode?1:c.length,c,l,h),n=n.next,e.bufferedRequestCount--,e.writing)break}null===n&&(e.lastBufferedRequest=null)}e.bufferedRequest=n,e.bufferProcessing=!1}function E(t){return t.ending&&0===t.length&&null===t.bufferedRequest&&!t.finished&&!t.writing}function T(t,e){t._final(function(n){e.pendingcb--,n&&t.emit("error",n),e.prefinished=!0,t.emit("prefinish"),x(t,e)})}function x(t,e){var n=E(e);return n&&(!function(t,e){e.prefinished||e.finalCalled||("function"==typeof t._final?(e.pendingcb++,e.finalCalled=!0,o.nextTick(T,t,e)):(e.prefinished=!0,t.emit("prefinish")))}(t,e),0===e.pendingcb&&(e.finished=!0,t.emit("finish"))),n}c.inherits(_,h),m.prototype.getBuffer=function(){for(var t=this.bufferedRequest,e=[];t;)e.push(t),t=t.next;return e},function(){try{Object.defineProperty(m.prototype,"buffer",{get:l.deprecate(function(){return this.getBuffer()},"_writableState.buffer is deprecated. Use _writableState.getBuffer instead.","DEP0003")})}catch(t){}}(),"function"==typeof Symbol&&Symbol.hasInstance&&"function"==typeof Function.prototype[Symbol.hasInstance]?(d=Function.prototype[Symbol.hasInstance],Object.defineProperty(_,Symbol.hasInstance,{value:function(t){return!!d.call(this,t)||this===_&&(t&&t._writableState instanceof m)}})):d=function(t){return t instanceof this},_.prototype.pipe=function(){this.emit("error",new Error("Cannot pipe, not readable"))},_.prototype.write=function(t,e,n){var r,i=this._writableState,s=!1,a=!i.objectMode&&(r=t,f.isBuffer(r)||r instanceof p);return a&&!f.isBuffer(t)&&(t=function(t){return f.from(t)}(t)),"function"==typeof e&&(n=e,e=null),a?e="buffer":e||(e=i.defaultEncoding),"function"!=typeof n&&(n=g),i.ended?function(t,e){var n=new Error("write after end");t.emit("error",n),o.nextTick(e,n)}(this,n):(a||function(t,e,n,r){var i=!0,s=!1;return null===n?s=new TypeError("May not write null values to stream"):"string"==typeof n||void 0===n||e.objectMode||(s=new TypeError("Invalid non-string/buffer chunk")),s&&(t.emit("error",s),o.nextTick(r,s),i=!1),i}(this,i,t,n))&&(i.pendingcb++,s=function(t,e,n,r,i,o){if(!n){var s=function(t,e,n){t.objectMode||!1===t.decodeStrings||"string"!=typeof e||(e=f.from(e,n));return e}(e,r,i);r!==s&&(n=!0,i="buffer",r=s)}var a=e.objectMode?1:r.length;e.length+=a;var u=e.length<e.highWaterMark;u||(e.needDrain=!0);if(e.writing||e.corked){var c=e.lastBufferedRequest;e.lastBufferedRequest={chunk:r,encoding:i,isBuf:n,callback:o,next:null},c?c.next=e.lastBufferedRequest:e.bufferedRequest=e.lastBufferedRequest,e.bufferedRequestCount+=1}else b(t,e,!1,a,r,i,o);return u}(this,i,a,t,e,n)),s},_.prototype.cork=function(){this._writableState.corked++},_.prototype.uncork=function(){var t=this._writableState;t.corked&&(t.corked--,t.writing||t.corked||t.finished||t.bufferProcessing||!t.bufferedRequest||v(this,t))},_.prototype.setDefaultEncoding=function(t){if("string"==typeof t&&(t=t.toLowerCase()),!(["hex","utf8","utf-8","ascii","binary","base64","ucs2","ucs-2","utf16le","utf-16le","raw"].indexOf((t+"").toLowerCase())>-1))throw new TypeError("Unknown encoding: "+t);return this._writableState.defaultEncoding=t,this},Object.defineProperty(_.prototype,"writableHighWaterMark",{enumerable:!1,get:function(){return this._writableState.highWaterMark}}),_.prototype._write=function(t,e,n){n(new Error("_write() is not implemented"))},_.prototype._writev=null,_.prototype.end=function(t,e,n){var r=this._writableState;"function"==typeof t?(n=t,t=null,e=null):"function"==typeof e&&(n=e,e=null),null!==t&&void 0!==t&&this.write(t,e),r.corked&&(r.corked=1,this.uncork()),r.ending||r.finished||function(t,e,n){e.ending=!0,x(t,e),n&&(e.finished?o.nextTick(n):t.once("finish",n));e.ended=!0,t.writable=!1}(this,r,n)},Object.defineProperty(_.prototype,"destroyed",{get:function(){return void 0!==this._writableState&&this._writableState.destroyed},set:function(t){this._writableState&&(this._writableState.destroyed=t)}}),_.prototype.destroy=y.destroy,_.prototype._undestroy=y.undestroy,_.prototype._destroy=function(t,e){this.end(),e(t)}}).call(this,n(8),n(30).setImmediate,n(4))},function(t,e,n){(e=t.exports=n(33)).Stream=e,e.Readable=e,e.Writable=n(23),e.Duplex=n(2),e.Transform=n(29),e.PassThrough=n(43)},function(t,e,n){"use strict";(function(t){
/*!
 * The buffer module from node.js, for the browser.
 *
 * @author   Feross Aboukhadijeh <> <http://feross.org>
 * @license  MIT
 */
var r=n(51),i=n(50),o=n(34);function s(){return u.TYPED_ARRAY_SUPPORT?2147483647:1073741823}function a(t,e){if(s()<e)throw new RangeError("Invalid typed array length");return u.TYPED_ARRAY_SUPPORT?(t=new Uint8Array(e)).__proto__=u.prototype:(null===t&&(t=new u(e)),t.length=e),t}function u(t,e,n){if(!(u.TYPED_ARRAY_SUPPORT||this instanceof u))return new u(t,e,n);if("number"==typeof t){if("string"==typeof e)throw new Error("If encoding is specified then the first argument must be a string");return h(this,t)}return c(this,t,e,n)}function c(t,e,n,r){if("number"==typeof e)throw new TypeError('"value" argument must not be a number');return"undefined"!=typeof ArrayBuffer&&e instanceof ArrayBuffer?function(t,e,n,r){if(e.byteLength,n<0||e.byteLength<n)throw new RangeError("'offset' is out of bounds");if(e.byteLength<n+(r||0))throw new RangeError("'length' is out of bounds");e=void 0===n&&void 0===r?new Uint8Array(e):void 0===r?new Uint8Array(e,n):new Uint8Array(e,n,r);u.TYPED_ARRAY_SUPPORT?(t=e).__proto__=u.prototype:t=f(t,e);return t}(t,e,n,r):"string"==typeof e?function(t,e,n){"string"==typeof n&&""!==n||(n="utf8");if(!u.isEncoding(n))throw new TypeError('"encoding" must be a valid string encoding');var r=0|d(e,n),i=(t=a(t,r)).write(e,n);i!==r&&(t=t.slice(0,i));return t}(t,e,n):function(t,e){if(u.isBuffer(e)){var n=0|p(e.length);return 0===(t=a(t,n)).length?t:(e.copy(t,0,0,n),t)}if(e){if("undefined"!=typeof ArrayBuffer&&e.buffer instanceof ArrayBuffer||"length"in e)return"number"!=typeof e.length||(r=e.length)!=r?a(t,0):f(t,e);if("Buffer"===e.type&&o(e.data))return f(t,e.data)}var r;throw new TypeError("First argument must be a string, Buffer, ArrayBuffer, Array, or array-like object.")}(t,e)}function l(t){if("number"!=typeof t)throw new TypeError('"size" argument must be a number');if(t<0)throw new RangeError('"size" argument must not be negative')}function h(t,e){if(l(e),t=a(t,e<0?0:0|p(e)),!u.TYPED_ARRAY_SUPPORT)for(var n=0;n<e;++n)t[n]=0;return t}function f(t,e){var n=e.length<0?0:0|p(e.length);t=a(t,n);for(var r=0;r<n;r+=1)t[r]=255&e[r];return t}function p(t){if(t>=s())throw new RangeError("Attempt to allocate Buffer larger than maximum size: 0x"+s().toString(16)+" bytes");return 0|t}function d(t,e){if(u.isBuffer(t))return t.length;if("undefined"!=typeof ArrayBuffer&&"function"==typeof ArrayBuffer.isView&&(ArrayBuffer.isView(t)||t instanceof ArrayBuffer))return t.byteLength;"string"!=typeof t&&(t=""+t);var n=t.length;if(0===n)return 0;for(var r=!1;;)switch(e){case"ascii":case"latin1":case"binary":return n;case"utf8":case"utf-8":case void 0:return j(t).length;case"ucs2":case"ucs-2":case"utf16le":case"utf-16le":return 2*n;case"hex":return n>>>1;case"base64":return Y(t).length;default:if(r)return j(t).length;e=(""+e).toLowerCase(),r=!0}}function y(t,e,n){var r=t[e];t[e]=t[n],t[n]=r}function g(t,e,n,r,i){if(0===t.length)return-1;if("string"==typeof n?(r=n,n=0):n>2147483647?n=2147483647:n<-2147483648&&(n=-2147483648),n=+n,isNaN(n)&&(n=i?0:t.length-1),n<0&&(n=t.length+n),n>=t.length){if(i)return-1;n=t.length-1}else if(n<0){if(!i)return-1;n=0}if("string"==typeof e&&(e=u.from(e,r)),u.isBuffer(e))return 0===e.length?-1:m(t,e,n,r,i);if("number"==typeof e)return e&=255,u.TYPED_ARRAY_SUPPORT&&"function"==typeof Uint8Array.prototype.indexOf?i?Uint8Array.prototype.indexOf.call(t,e,n):Uint8Array.prototype.lastIndexOf.call(t,e,n):m(t,[e],n,r,i);throw new TypeError("val must be string, number or Buffer")}function m(t,e,n,r,i){var o,s=1,a=t.length,u=e.length;if(void 0!==r&&("ucs2"===(r=String(r).toLowerCase())||"ucs-2"===r||"utf16le"===r||"utf-16le"===r)){if(t.length<2||e.length<2)return-1;s=2,a/=2,u/=2,n/=2}function c(t,e){return 1===s?t[e]:t.readUInt16BE(e*s)}if(i){var l=-1;for(o=n;o<a;o++)if(c(t,o)===c(e,-1===l?0:o-l)){if(-1===l&&(l=o),o-l+1===u)return l*s}else-1!==l&&(o-=o-l),l=-1}else for(n+u>a&&(n=a-u),o=n;o>=0;o--){for(var h=!0,f=0;f<u;f++)if(c(t,o+f)!==c(e,f)){h=!1;break}if(h)return o}return-1}function _(t,e,n,r){n=Number(n)||0;var i=t.length-n;r?(r=Number(r))>i&&(r=i):r=i;var o=e.length;if(o%2!=0)throw new TypeError("Invalid hex string");r>o/2&&(r=o/2);for(var s=0;s<r;++s){var a=parseInt(e.substr(2*s,2),16);if(isNaN(a))return s;t[n+s]=a}return s}function b(t,e,n,r){return z(j(e,t.length-n),t,n,r)}function w(t,e,n,r){return z(function(t){for(var e=[],n=0;n<t.length;++n)e.push(255&t.charCodeAt(n));return e}(e),t,n,r)}function v(t,e,n,r){return w(t,e,n,r)}function E(t,e,n,r){return z(Y(e),t,n,r)}function T(t,e,n,r){return z(function(t,e){for(var n,r,i,o=[],s=0;s<t.length&&!((e-=2)<0);++s)n=t.charCodeAt(s),r=n>>8,i=n%256,o.push(i),o.push(r);return o}(e,t.length-n),t,n,r)}function x(t,e,n){return 0===e&&n===t.length?r.fromByteArray(t):r.fromByteArray(t.slice(e,n))}function A(t,e,n){n=Math.min(t.length,n);for(var r=[],i=e;i<n;){var o,s,a,u,c=t[i],l=null,h=c>239?4:c>223?3:c>191?2:1;if(i+h<=n)switch(h){case 1:c<128&&(l=c);break;case 2:128==(192&(o=t[i+1]))&&(u=(31&c)<<6|63&o)>127&&(l=u);break;case 3:o=t[i+1],s=t[i+2],128==(192&o)&&128==(192&s)&&(u=(15&c)<<12|(63&o)<<6|63&s)>2047&&(u<55296||u>57343)&&(l=u);break;case 4:o=t[i+1],s=t[i+2],a=t[i+3],128==(192&o)&&128==(192&s)&&128==(192&a)&&(u=(15&c)<<18|(63&o)<<12|(63&s)<<6|63&a)>65535&&u<1114112&&(l=u)}null===l?(l=65533,h=1):l>65535&&(l-=65536,r.push(l>>>10&1023|55296),l=56320|1023&l),r.push(l),i+=h}return function(t){var e=t.length;if(e<=D)return String.fromCharCode.apply(String,t);var n="",r=0;for(;r<e;)n+=String.fromCharCode.apply(String,t.slice(r,r+=D));return n}(r)}e.Buffer=u,e.SlowBuffer=function(t){+t!=t&&(t=0);return u.alloc(+t)},e.INSPECT_MAX_BYTES=50,u.TYPED_ARRAY_SUPPORT=void 0!==t.TYPED_ARRAY_SUPPORT?t.TYPED_ARRAY_SUPPORT:function(){try{var t=new Uint8Array(1);return t.__proto__={__proto__:Uint8Array.prototype,foo:function(){return 42}},42===t.foo()&&"function"==typeof t.subarray&&0===t.subarray(1,1).byteLength}catch(t){return!1}}(),e.kMaxLength=s(),u.poolSize=8192,u._augment=function(t){return t.__proto__=u.prototype,t},u.from=function(t,e,n){return c(null,t,e,n)},u.TYPED_ARRAY_SUPPORT&&(u.prototype.__proto__=Uint8Array.prototype,u.__proto__=Uint8Array,"undefined"!=typeof Symbol&&Symbol.species&&u[Symbol.species]===u&&Object.defineProperty(u,Symbol.species,{value:null,configurable:!0})),u.alloc=function(t,e,n){return function(t,e,n,r){return l(e),e<=0?a(t,e):void 0!==n?"string"==typeof r?a(t,e).fill(n,r):a(t,e).fill(n):a(t,e)}(null,t,e,n)},u.allocUnsafe=function(t){return h(null,t)},u.allocUnsafeSlow=function(t){return h(null,t)},u.isBuffer=function(t){return!(null==t||!t._isBuffer)},u.compare=function(t,e){if(!u.isBuffer(t)||!u.isBuffer(e))throw new TypeError("Arguments must be Buffers");if(t===e)return 0;for(var n=t.length,r=e.length,i=0,o=Math.min(n,r);i<o;++i)if(t[i]!==e[i]){n=t[i],r=e[i];break}return n<r?-1:r<n?1:0},u.isEncoding=function(t){switch(String(t).toLowerCase()){case"hex":case"utf8":case"utf-8":case"ascii":case"latin1":case"binary":case"base64":case"ucs2":case"ucs-2":case"utf16le":case"utf-16le":return!0;default:return!1}},u.concat=function(t,e){if(!o(t))throw new TypeError('"list" argument must be an Array of Buffers');if(0===t.length)return u.alloc(0);var n;if(void 0===e)for(e=0,n=0;n<t.length;++n)e+=t[n].length;var r=u.allocUnsafe(e),i=0;for(n=0;n<t.length;++n){var s=t[n];if(!u.isBuffer(s))throw new TypeError('"list" argument must be an Array of Buffers');s.copy(r,i),i+=s.length}return r},u.byteLength=d,u.prototype._isBuffer=!0,u.prototype.swap16=function(){var t=this.length;if(t%2!=0)throw new RangeError("Buffer size must be a multiple of 16-bits");for(var e=0;e<t;e+=2)y(this,e,e+1);return this},u.prototype.swap32=function(){var t=this.length;if(t%4!=0)throw new RangeError("Buffer size must be a multiple of 32-bits");for(var e=0;e<t;e+=4)y(this,e,e+3),y(this,e+1,e+2);return this},u.prototype.swap64=function(){var t=this.length;if(t%8!=0)throw new RangeError("Buffer size must be a multiple of 64-bits");for(var e=0;e<t;e+=8)y(this,e,e+7),y(this,e+1,e+6),y(this,e+2,e+5),y(this,e+3,e+4);return this},u.prototype.toString=function(){var t=0|this.length;return 0===t?"":0===arguments.length?A(this,0,t):function(t,e,n){var r=!1;if((void 0===e||e<0)&&(e=0),e>this.length)return"";if((void 0===n||n>this.length)&&(n=this.length),n<=0)return"";if((n>>>=0)<=(e>>>=0))return"";for(t||(t="utf8");;)switch(t){case"hex":return S(this,e,n);case"utf8":case"utf-8":return A(this,e,n);case"ascii":return I(this,e,n);case"latin1":case"binary":return N(this,e,n);case"base64":return x(this,e,n);case"ucs2":case"ucs-2":case"utf16le":case"utf-16le":return O(this,e,n);default:if(r)throw new TypeError("Unknown encoding: "+t);t=(t+"").toLowerCase(),r=!0}}.apply(this,arguments)},u.prototype.equals=function(t){if(!u.isBuffer(t))throw new TypeError("Argument must be a Buffer");return this===t||0===u.compare(this,t)},u.prototype.inspect=function(){var t="",n=e.INSPECT_MAX_BYTES;return this.length>0&&(t=this.toString("hex",0,n).match(/.{2}/g).join(" "),this.length>n&&(t+=" ... ")),"<Buffer "+t+">"},u.prototype.compare=function(t,e,n,r,i){if(!u.isBuffer(t))throw new TypeError("Argument must be a Buffer");if(void 0===e&&(e=0),void 0===n&&(n=t?t.length:0),void 0===r&&(r=0),void 0===i&&(i=this.length),e<0||n>t.length||r<0||i>this.length)throw new RangeError("out of range index");if(r>=i&&e>=n)return 0;if(r>=i)return-1;if(e>=n)return 1;if(e>>>=0,n>>>=0,r>>>=0,i>>>=0,this===t)return 0;for(var o=i-r,s=n-e,a=Math.min(o,s),c=this.slice(r,i),l=t.slice(e,n),h=0;h<a;++h)if(c[h]!==l[h]){o=c[h],s=l[h];break}return o<s?-1:s<o?1:0},u.prototype.includes=function(t,e,n){return-1!==this.indexOf(t,e,n)},u.prototype.indexOf=function(t,e,n){return g(this,t,e,n,!0)},u.prototype.lastIndexOf=function(t,e,n){return g(this,t,e,n,!1)},u.prototype.write=function(t,e,n,r){if(void 0===e)r="utf8",n=this.length,e=0;else if(void 0===n&&"string"==typeof e)r=e,n=this.length,e=0;else{if(!isFinite(e))throw new Error("Buffer.write(string, encoding, offset[, length]) is no longer supported");e|=0,isFinite(n)?(n|=0,void 0===r&&(r="utf8")):(r=n,n=void 0)}var i=this.length-e;if((void 0===n||n>i)&&(n=i),t.length>0&&(n<0||e<0)||e>this.length)throw new RangeError("Attempt to write outside buffer bounds");r||(r="utf8");for(var o=!1;;)switch(r){case"hex":return _(this,t,e,n);case"utf8":case"utf-8":return b(this,t,e,n);case"ascii":return w(this,t,e,n);case"latin1":case"binary":return v(this,t,e,n);case"base64":return E(this,t,e,n);case"ucs2":case"ucs-2":case"utf16le":case"utf-16le":return T(this,t,e,n);default:if(o)throw new TypeError("Unknown encoding: "+r);r=(""+r).toLowerCase(),o=!0}},u.prototype.toJSON=function(){return{type:"Buffer",data:Array.prototype.slice.call(this._arr||this,0)}};var D=4096;function I(t,e,n){var r="";n=Math.min(t.length,n);for(var i=e;i<n;++i)r+=String.fromCharCode(127&t[i]);return r}function N(t,e,n){var r="";n=Math.min(t.length,n);for(var i=e;i<n;++i)r+=String.fromCharCode(t[i]);return r}function S(t,e,n){var r=t.length;(!e||e<0)&&(e=0),(!n||n<0||n>r)&&(n=r);for(var i="",o=e;o<n;++o)i+=F(t[o]);return i}function O(t,e,n){for(var r=t.slice(e,n),i="",o=0;o<r.length;o+=2)i+=String.fromCharCode(r[o]+256*r[o+1]);return i}function C(t,e,n){if(t%1!=0||t<0)throw new RangeError("offset is not uint");if(t+e>n)throw new RangeError("Trying to access beyond buffer length")}function P(t,e,n,r,i,o){if(!u.isBuffer(t))throw new TypeError('"buffer" argument must be a Buffer instance');if(e>i||e<o)throw new RangeError('"value" argument is out of bounds');if(n+r>t.length)throw new RangeError("Index out of range")}function k(t,e,n,r){e<0&&(e=65535+e+1);for(var i=0,o=Math.min(t.length-n,2);i<o;++i)t[n+i]=(e&255<<8*(r?i:1-i))>>>8*(r?i:1-i)}function L(t,e,n,r){e<0&&(e=4294967295+e+1);for(var i=0,o=Math.min(t.length-n,4);i<o;++i)t[n+i]=e>>>8*(r?i:3-i)&255}function R(t,e,n,r,i,o){if(n+r>t.length)throw new RangeError("Index out of range");if(n<0)throw new RangeError("Index out of range")}function M(t,e,n,r,o){return o||R(t,0,n,4),i.write(t,e,n,r,23,4),n+4}function B(t,e,n,r,o){return o||R(t,0,n,8),i.write(t,e,n,r,52,8),n+8}u.prototype.slice=function(t,e){var n,r=this.length;if(t=~~t,e=void 0===e?r:~~e,t<0?(t+=r)<0&&(t=0):t>r&&(t=r),e<0?(e+=r)<0&&(e=0):e>r&&(e=r),e<t&&(e=t),u.TYPED_ARRAY_SUPPORT)(n=this.subarray(t,e)).__proto__=u.prototype;else{var i=e-t;n=new u(i,void 0);for(var o=0;o<i;++o)n[o]=this[o+t]}return n},u.prototype.readUIntLE=function(t,e,n){t|=0,e|=0,n||C(t,e,this.length);for(var r=this[t],i=1,o=0;++o<e&&(i*=256);)r+=this[t+o]*i;return r},u.prototype.readUIntBE=function(t,e,n){t|=0,e|=0,n||C(t,e,this.length);for(var r=this[t+--e],i=1;e>0&&(i*=256);)r+=this[t+--e]*i;return r},u.prototype.readUInt8=function(t,e){return e||C(t,1,this.length),this[t]},u.prototype.readUInt16LE=function(t,e){return e||C(t,2,this.length),this[t]|this[t+1]<<8},u.prototype.readUInt16BE=function(t,e){return e||C(t,2,this.length),this[t]<<8|this[t+1]},u.prototype.readUInt32LE=function(t,e){return e||C(t,4,this.length),(this[t]|this[t+1]<<8|this[t+2]<<16)+16777216*this[t+3]},u.prototype.readUInt32BE=function(t,e){return e||C(t,4,this.length),16777216*this[t]+(this[t+1]<<16|this[t+2]<<8|this[t+3])},u.prototype.readIntLE=function(t,e,n){t|=0,e|=0,n||C(t,e,this.length);for(var r=this[t],i=1,o=0;++o<e&&(i*=256);)r+=this[t+o]*i;return r>=(i*=128)&&(r-=Math.pow(2,8*e)),r},u.prototype.readIntBE=function(t,e,n){t|=0,e|=0,n||C(t,e,this.length);for(var r=e,i=1,o=this[t+--r];r>0&&(i*=256);)o+=this[t+--r]*i;return o>=(i*=128)&&(o-=Math.pow(2,8*e)),o},u.prototype.readInt8=function(t,e){return e||C(t,1,this.length),128&this[t]?-1*(255-this[t]+1):this[t]},u.prototype.readInt16LE=function(t,e){e||C(t,2,this.length);var n=this[t]|this[t+1]<<8;return 32768&n?4294901760|n:n},u.prototype.readInt16BE=function(t,e){e||C(t,2,this.length);var n=this[t+1]|this[t]<<8;return 32768&n?4294901760|n:n},u.prototype.readInt32LE=function(t,e){return e||C(t,4,this.length),this[t]|this[t+1]<<8|this[t+2]<<16|this[t+3]<<24},u.prototype.readInt32BE=function(t,e){return e||C(t,4,this.length),this[t]<<24|this[t+1]<<16|this[t+2]<<8|this[t+3]},u.prototype.readFloatLE=function(t,e){return e||C(t,4,this.length),i.read(this,t,!0,23,4)},u.prototype.readFloatBE=function(t,e){return e||C(t,4,this.length),i.read(this,t,!1,23,4)},u.prototype.readDoubleLE=function(t,e){return e||C(t,8,this.length),i.read(this,t,!0,52,8)},u.prototype.readDoubleBE=function(t,e){return e||C(t,8,this.length),i.read(this,t,!1,52,8)},u.prototype.writeUIntLE=function(t,e,n,r){(t=+t,e|=0,n|=0,r)||P(this,t,e,n,Math.pow(2,8*n)-1,0);var i=1,o=0;for(this[e]=255&t;++o<n&&(i*=256);)this[e+o]=t/i&255;return e+n},u.prototype.writeUIntBE=function(t,e,n,r){(t=+t,e|=0,n|=0,r)||P(this,t,e,n,Math.pow(2,8*n)-1,0);var i=n-1,o=1;for(this[e+i]=255&t;--i>=0&&(o*=256);)this[e+i]=t/o&255;return e+n},u.prototype.writeUInt8=function(t,e,n){return t=+t,e|=0,n||P(this,t,e,1,255,0),u.TYPED_ARRAY_SUPPORT||(t=Math.floor(t)),this[e]=255&t,e+1},u.prototype.writeUInt16LE=function(t,e,n){return t=+t,e|=0,n||P(this,t,e,2,65535,0),u.TYPED_ARRAY_SUPPORT?(this[e]=255&t,this[e+1]=t>>>8):k(this,t,e,!0),e+2},u.prototype.writeUInt16BE=function(t,e,n){return t=+t,e|=0,n||P(this,t,e,2,65535,0),u.TYPED_ARRAY_SUPPORT?(this[e]=t>>>8,this[e+1]=255&t):k(this,t,e,!1),e+2},u.prototype.writeUInt32LE=function(t,e,n){return t=+t,e|=0,n||P(this,t,e,4,4294967295,0),u.TYPED_ARRAY_SUPPORT?(this[e+3]=t>>>24,this[e+2]=t>>>16,this[e+1]=t>>>8,this[e]=255&t):L(this,t,e,!0),e+4},u.prototype.writeUInt32BE=function(t,e,n){return t=+t,e|=0,n||P(this,t,e,4,4294967295,0),u.TYPED_ARRAY_SUPPORT?(this[e]=t>>>24,this[e+1]=t>>>16,this[e+2]=t>>>8,this[e+3]=255&t):L(this,t,e,!1),e+4},u.prototype.writeIntLE=function(t,e,n,r){if(t=+t,e|=0,!r){var i=Math.pow(2,8*n-1);P(this,t,e,n,i-1,-i)}var o=0,s=1,a=0;for(this[e]=255&t;++o<n&&(s*=256);)t<0&&0===a&&0!==this[e+o-1]&&(a=1),this[e+o]=(t/s>>0)-a&255;return e+n},u.prototype.writeIntBE=function(t,e,n,r){if(t=+t,e|=0,!r){var i=Math.pow(2,8*n-1);P(this,t,e,n,i-1,-i)}var o=n-1,s=1,a=0;for(this[e+o]=255&t;--o>=0&&(s*=256);)t<0&&0===a&&0!==this[e+o+1]&&(a=1),this[e+o]=(t/s>>0)-a&255;return e+n},u.prototype.writeInt8=function(t,e,n){return t=+t,e|=0,n||P(this,t,e,1,127,-128),u.TYPED_ARRAY_SUPPORT||(t=Math.floor(t)),t<0&&(t=255+t+1),this[e]=255&t,e+1},u.prototype.writeInt16LE=function(t,e,n){return t=+t,e|=0,n||P(this,t,e,2,32767,-32768),u.TYPED_ARRAY_SUPPORT?(this[e]=255&t,this[e+1]=t>>>8):k(this,t,e,!0),e+2},u.prototype.writeInt16BE=function(t,e,n){return t=+t,e|=0,n||P(this,t,e,2,32767,-32768),u.TYPED_ARRAY_SUPPORT?(this[e]=t>>>8,this[e+1]=255&t):k(this,t,e,!1),e+2},u.prototype.writeInt32LE=function(t,e,n){return t=+t,e|=0,n||P(this,t,e,4,2147483647,-2147483648),u.TYPED_ARRAY_SUPPORT?(this[e]=255&t,this[e+1]=t>>>8,this[e+2]=t>>>16,this[e+3]=t>>>24):L(this,t,e,!0),e+4},u.prototype.writeInt32BE=function(t,e,n){return t=+t,e|=0,n||P(this,t,e,4,2147483647,-2147483648),t<0&&(t=4294967295+t+1),u.TYPED_ARRAY_SUPPORT?(this[e]=t>>>24,this[e+1]=t>>>16,this[e+2]=t>>>8,this[e+3]=255&t):L(this,t,e,!1),e+4},u.prototype.writeFloatLE=function(t,e,n){return M(this,t,e,!0,n)},u.prototype.writeFloatBE=function(t,e,n){return M(this,t,e,!1,n)},u.prototype.writeDoubleLE=function(t,e,n){return B(this,t,e,!0,n)},u.prototype.writeDoubleBE=function(t,e,n){return B(this,t,e,!1,n)},u.prototype.copy=function(t,e,n,r){if(n||(n=0),r||0===r||(r=this.length),e>=t.length&&(e=t.length),e||(e=0),r>0&&r<n&&(r=n),r===n)return 0;if(0===t.length||0===this.length)return 0;if(e<0)throw new RangeError("targetStart out of bounds");if(n<0||n>=this.length)throw new RangeError("sourceStart out of bounds");if(r<0)throw new RangeError("sourceEnd out of bounds");r>this.length&&(r=this.length),t.length-e<r-n&&(r=t.length-e+n);var i,o=r-n;if(this===t&&n<e&&e<r)for(i=o-1;i>=0;--i)t[i+e]=this[i+n];else if(o<1e3||!u.TYPED_ARRAY_SUPPORT)for(i=0;i<o;++i)t[i+e]=this[i+n];else Uint8Array.prototype.set.call(t,this.subarray(n,n+o),e);return o},u.prototype.fill=function(t,e,n,r){if("string"==typeof t){if("string"==typeof e?(r=e,e=0,n=this.length):"string"==typeof n&&(r=n,n=this.length),1===t.length){var i=t.charCodeAt(0);i<256&&(t=i)}if(void 0!==r&&"string"!=typeof r)throw new TypeError("encoding must be a string");if("string"==typeof r&&!u.isEncoding(r))throw new TypeError("Unknown encoding: "+r)}else"number"==typeof t&&(t&=255);if(e<0||this.length<e||this.length<n)throw new RangeError("Out of range index");if(n<=e)return this;var o;if(e>>>=0,n=void 0===n?this.length:n>>>0,t||(t=0),"number"==typeof t)for(o=e;o<n;++o)this[o]=t;else{var s=u.isBuffer(t)?t:j(new u(t,r).toString()),a=s.length;for(o=0;o<n-e;++o)this[o+e]=s[o%a]}return this};var U=/[^+\/0-9A-Za-z-_]/g;function F(t){return t<16?"0"+t.toString(16):t.toString(16)}function j(t,e){var n;e=e||1/0;for(var r=t.length,i=null,o=[],s=0;s<r;++s){if((n=t.charCodeAt(s))>55295&&n<57344){if(!i){if(n>56319){(e-=3)>-1&&o.push(239,191,189);continue}if(s+1===r){(e-=3)>-1&&o.push(239,191,189);continue}i=n;continue}if(n<56320){(e-=3)>-1&&o.push(239,191,189),i=n;continue}n=65536+(i-55296<<10|n-56320)}else i&&(e-=3)>-1&&o.push(239,191,189);if(i=null,n<128){if((e-=1)<0)break;o.push(n)}else if(n<2048){if((e-=2)<0)break;o.push(n>>6|192,63&n|128)}else if(n<65536){if((e-=3)<0)break;o.push(n>>12|224,n>>6&63|128,63&n|128)}else{if(!(n<1114112))throw new Error("Invalid code point");if((e-=4)<0)break;o.push(n>>18|240,n>>12&63|128,n>>6&63|128,63&n|128)}}return o}function Y(t){return r.toByteArray(function(t){if((t=function(t){return t.trim?t.trim():t.replace(/^\s+|\s+$/g,"")}(t).replace(U,"")).length<2)return"";for(;t.length%4!=0;)t+="=";return t}(t))}function z(t,e,n,r){for(var i=0;i<r&&!(i+n>=e.length||i>=t.length);++i)e[i+n]=t[i];return i}}).call(this,n(4))},function(t,e,n){(function(){var e,r,i,o,s,a,u,c,l,h,f,p,d,y={}.hasOwnProperty;u=n(18),c=n(17),e=n(20),r=n(19),l=n(21),f=n(12),p=n(11),h=n(10),i=n(16),o=n(14),s=n(15),a=n(13),d=n(35),t.exports=function(t){function n(t){n.__super__.constructor.call(this,t)}return function(t,e){for(var n in e)y.call(e,n)&&(t[n]=e[n]);function r(){this.constructor=t}r.prototype=e.prototype,t.prototype=new r,t.__super__=e.prototype}(n,d),n.prototype.document=function(t){var e,n,i,o,s;for(this.textispresent=!1,o="",n=0,i=(s=t.children).length;n<i;n++)e=s[n],o+=function(){switch(!1){case!(e instanceof u):return this.declaration(e);case!(e instanceof c):return this.docType(e);case!(e instanceof r):return this.comment(e);case!(e instanceof h):return this.processingInstruction(e);default:return this.element(e,0)}}.call(this);return this.pretty&&o.slice(-this.newline.length)===this.newline&&(o=o.slice(0,-this.newline.length)),o},n.prototype.attribute=function(t){return" "+t.name+'="'+t.value+'"'},n.prototype.cdata=function(t,e){return this.space(e)+"<![CDATA["+t.text+"]]>"+this.newline},n.prototype.comment=function(t,e){return this.space(e)+"\x3c!-- "+t.text+" --\x3e"+this.newline},n.prototype.declaration=function(t,e){var n;return n=this.space(e),n+='<?xml version="'+t.version+'"',null!=t.encoding&&(n+=' encoding="'+t.encoding+'"'),null!=t.standalone&&(n+=' standalone="'+t.standalone+'"'),n+=this.spacebeforeslash+"?>",n+=this.newline},n.prototype.docType=function(t,n){var u,c,l,f,p;if(n||(n=0),f=this.space(n),f+="<!DOCTYPE "+t.root().name,t.pubID&&t.sysID?f+=' PUBLIC "'+t.pubID+'" "'+t.sysID+'"':t.sysID&&(f+=' SYSTEM "'+t.sysID+'"'),t.children.length>0){for(f+=" [",f+=this.newline,c=0,l=(p=t.children).length;c<l;c++)u=p[c],f+=function(){switch(!1){case!(u instanceof i):return this.dtdAttList(u,n+1);case!(u instanceof o):return this.dtdElement(u,n+1);case!(u instanceof s):return this.dtdEntity(u,n+1);case!(u instanceof a):return this.dtdNotation(u,n+1);case!(u instanceof e):return this.cdata(u,n+1);case!(u instanceof r):return this.comment(u,n+1);case!(u instanceof h):return this.processingInstruction(u,n+1);default:throw new Error("Unknown DTD node type: "+u.constructor.name)}}.call(this);f+="]"}return f+=this.spacebeforeslash+">",f+=this.newline},n.prototype.element=function(t,n){var i,o,s,a,u,c,d,g,m,_,b,w,v;for(d in n||(n=0),v=!1,this.textispresent?(this.newline="",this.pretty=!1):(this.newline=this.newlinedefault,this.pretty=this.prettydefault),g="",g+=(w=this.space(n))+"<"+t.name,m=t.attributes)y.call(m,d)&&(i=m[d],g+=this.attribute(i));if(0===t.children.length||t.children.every(function(t){return""===t.value}))this.allowEmpty?g+="></"+t.name+">"+this.newline:g+=this.spacebeforeslash+"/>"+this.newline;else if(this.pretty&&1===t.children.length&&null!=t.children[0].value)g+=">",g+=t.children[0].value,g+="</"+t.name+">"+this.newline;else{if(this.dontprettytextnodes)for(s=0,u=(_=t.children).length;s<u;s++)if(null!=(o=_[s]).value){this.textispresent++,v=!0;break}for(this.textispresent&&(this.newline="",this.pretty=!1,w=this.space(n)),g+=">"+this.newline,a=0,c=(b=t.children).length;a<c;a++)o=b[a],g+=function(){switch(!1){case!(o instanceof e):return this.cdata(o,n+1);case!(o instanceof r):return this.comment(o,n+1);case!(o instanceof l):return this.element(o,n+1);case!(o instanceof f):return this.raw(o,n+1);case!(o instanceof p):return this.text(o,n+1);case!(o instanceof h):return this.processingInstruction(o,n+1);default:throw new Error("Unknown XML node type: "+o.constructor.name)}}.call(this);v&&this.textispresent--,this.textispresent||(this.newline=this.newlinedefault,this.pretty=this.prettydefault),g+=w+"</"+t.name+">"+this.newline}return g},n.prototype.processingInstruction=function(t,e){var n;return n=this.space(e)+"<?"+t.target,t.value&&(n+=" "+t.value),n+=this.spacebeforeslash+"?>"+this.newline},n.prototype.raw=function(t,e){return this.space(e)+t.value+this.newline},n.prototype.text=function(t,e){return this.space(e)+t.value+this.newline},n.prototype.dtdAttList=function(t,e){var n;return n=this.space(e)+"<!ATTLIST "+t.elementName+" "+t.attributeName+" "+t.attributeType,"#DEFAULT"!==t.defaultValueType&&(n+=" "+t.defaultValueType),t.defaultValue&&(n+=' "'+t.defaultValue+'"'),n+=this.spacebeforeslash+">"+this.newline},n.prototype.dtdElement=function(t,e){return this.space(e)+"<!ELEMENT "+t.name+" "+t.value+this.spacebeforeslash+">"+this.newline},n.prototype.dtdEntity=function(t,e){var n;return n=this.space(e)+"<!ENTITY",t.pe&&(n+=" %"),n+=" "+t.name,t.value?n+=' "'+t.value+'"':(t.pubID&&t.sysID?n+=' PUBLIC "'+t.pubID+'" "'+t.sysID+'"':t.sysID&&(n+=' SYSTEM "'+t.sysID+'"'),t.nData&&(n+=" NDATA "+t.nData)),n+=this.spacebeforeslash+">"+this.newline},n.prototype.dtdNotation=function(t,e){var n;return n=this.space(e)+"<!NOTATION "+t.name,t.pubID&&t.sysID?n+=' PUBLIC "'+t.pubID+'" "'+t.sysID+'"':t.pubID?n+=' PUBLIC "'+t.pubID+'"':t.sysID&&(n+=' SYSTEM "'+t.sysID+'"'),n+=this.spacebeforeslash+">"+this.newline},n.prototype.openNode=function(t,e){var n,r,i,o;if(e||(e=0),t instanceof l){for(r in i=this.space(e)+"<"+t.name,o=t.attributes)y.call(o,r)&&(n=o[r],i+=this.attribute(n));return i+=(t.children?">":"/>")+this.newline}return i=this.space(e)+"<!DOCTYPE "+t.rootNodeName,t.pubID&&t.sysID?i+=' PUBLIC "'+t.pubID+'" "'+t.sysID+'"':t.sysID&&(i+=' SYSTEM "'+t.sysID+'"'),i+=(t.children?" [":">")+this.newline},n.prototype.closeNode=function(t,e){switch(e||(e=0),!1){case!(t instanceof l):return this.space(e)+"</"+t.name+">"+this.newline;case!(t instanceof c):return this.space(e)+"]>"+this.newline}},n}()}).call(this)},function(t,e){(function(){e.defaults={.1:{explicitCharkey:!1,trim:!0,normalize:!0,normalizeTags:!1,attrkey:"@",charkey:"#",explicitArray:!1,ignoreAttrs:!1,mergeAttrs:!1,explicitRoot:!1,validator:null,xmlns:!1,explicitChildren:!1,childkey:"@@",charsAsChildren:!1,includeWhiteChars:!1,async:!1,strict:!0,attrNameProcessors:null,attrValueProcessors:null,tagNameProcessors:null,valueProcessors:null,emptyTag:""},.2:{explicitCharkey:!1,trim:!1,normalize:!1,normalizeTags:!1,attrkey:"$",charkey:"_",explicitArray:!0,ignoreAttrs:!1,mergeAttrs:!1,explicitRoot:!0,validator:null,xmlns:!1,explicitChildren:!1,preserveChildrenOrder:!1,childkey:"$$",charsAsChildren:!1,includeWhiteChars:!1,async:!1,strict:!0,attrNameProcessors:null,attrValueProcessors:null,tagNameProcessors:null,valueProcessors:null,rootName:"root",xmldec:{version:"1.0",encoding:"UTF-8",standalone:!0},doctype:null,renderOpts:{pretty:!0,indent:"  ",newline:"\n"},headless:!1,chunkSize:1e4,emptyTag:"",cdata:!1}}}).call(this)},function(t,e){(function(){"use strict";var t;t=new RegExp(/(?!xmlns)^.*:/),e.normalize=function(t){return t.toLowerCase()},e.firstCharLowerCase=function(t){return t.charAt(0).toLowerCase()+t.slice(1)},e.stripPrefix=function(e){return e.replace(t,"")},e.parseNumbers=function(t){return isNaN(t)||(t=t%1==0?parseInt(t,10):parseFloat(t)),t},e.parseBooleans=function(t){return/^(?:true|false)$/i.test(t)&&(t="true"===t.toLowerCase()),t}}).call(this)},function(t,e,n){"use strict";t.exports=o;var r=n(2),i=n(5);function o(t){if(!(this instanceof o))return new o(t);r.call(this,t),this._transformState={afterTransform:function(t,e){var n=this._transformState;n.transforming=!1;var r=n.writecb;if(!r)return this.emit("error",new Error("write callback called multiple times"));n.writechunk=null,n.writecb=null,null!=e&&this.push(e),r(t);var i=this._readableState;i.reading=!1,(i.needReadable||i.length<i.highWaterMark)&&this._read(i.highWaterMark)}.bind(this),needTransform:!1,transforming:!1,writecb:null,writechunk:null,writeencoding:null},this._readableState.needReadable=!0,this._readableState.sync=!1,t&&("function"==typeof t.transform&&(this._transform=t.transform),"function"==typeof t.flush&&(this._flush=t.flush)),this.on("prefinish",s)}function s(){var t=this;"function"==typeof this._flush?this._flush(function(e,n){a(t,e,n)}):a(this,null,null)}function a(t,e,n){if(e)return t.emit("error",e);if(null!=n&&t.push(n),t._writableState.length)throw new Error("Calling transform done when ws.length != 0");if(t._transformState.transforming)throw new Error("Calling transform done when still transforming");return t.push(null)}i.inherits=n(3),i.inherits(o,r),o.prototype.push=function(t,e){return this._transformState.needTransform=!1,r.prototype.push.call(this,t,e)},o.prototype._transform=function(t,e,n){throw new Error("_transform() is not implemented")},o.prototype._write=function(t,e,n){var r=this._transformState;if(r.writecb=n,r.writechunk=t,r.writeencoding=e,!r.transforming){var i=this._readableState;(r.needTransform||i.needReadable||i.length<i.highWaterMark)&&this._read(i.highWaterMark)}},o.prototype._read=function(t){var e=this._transformState;null!==e.writechunk&&e.writecb&&!e.transforming?(e.transforming=!0,this._transform(e.writechunk,e.writeencoding,e.afterTransform)):e.needTransform=!0},o.prototype._destroy=function(t,e){var n=this;r.prototype._destroy.call(this,t,function(t){e(t),n.emit("close")})}},function(t,e,n){(function(t){var r=void 0!==t&&t||"undefined"!=typeof self&&self||window,i=Function.prototype.apply;function o(t,e){this._id=t,this._clearFn=e}e.setTimeout=function(){return new o(i.call(setTimeout,r,arguments),clearTimeout)},e.setInterval=function(){return new o(i.call(setInterval,r,arguments),clearInterval)},e.clearTimeout=e.clearInterval=function(t){t&&t.close()},o.prototype.unref=o.prototype.ref=function(){},o.prototype.close=function(){this._clearFn.call(r,this._id)},e.enroll=function(t,e){clearTimeout(t._idleTimeoutId),t._idleTimeout=e},e.unenroll=function(t){clearTimeout(t._idleTimeoutId),t._idleTimeout=-1},e._unrefActive=e.active=function(t){clearTimeout(t._idleTimeoutId);var e=t._idleTimeout;e>=0&&(t._idleTimeoutId=setTimeout(function(){t._onTimeout&&t._onTimeout()},e))},n(45),e.setImmediate="undefined"!=typeof self&&self.setImmediate||void 0!==t&&t.setImmediate||this&&this.setImmediate,e.clearImmediate="undefined"!=typeof self&&self.clearImmediate||void 0!==t&&t.clearImmediate||this&&this.clearImmediate}).call(this,n(4))},function(t,e,n){"use strict";var r=n(7);function i(t,e){t.emit("error",e)}t.exports={destroy:function(t,e){var n=this,o=this._readableState&&this._readableState.destroyed,s=this._writableState&&this._writableState.destroyed;return o||s?(e?e(t):!t||this._writableState&&this._writableState.errorEmitted||r.nextTick(i,this,t),this):(this._readableState&&(this._readableState.destroyed=!0),this._writableState&&(this._writableState.destroyed=!0),this._destroy(t||null,function(t){!e&&t?(r.nextTick(i,n,t),n._writableState&&(n._writableState.errorEmitted=!0)):e&&e(t)}),this)},undestroy:function(){this._readableState&&(this._readableState.destroyed=!1,this._readableState.reading=!1,this._readableState.ended=!1,this._readableState.endEmitted=!1),this._writableState&&(this._writableState.destroyed=!1,this._writableState.ended=!1,this._writableState.ending=!1,this._writableState.finished=!1,this._writableState.errorEmitted=!1)}}},function(t,e,n){t.exports=n(9).EventEmitter},function(t,e,n){"use strict";(function(e,r){var i=n(7);t.exports=b;var o,s=n(34);b.ReadableState=_;n(9).EventEmitter;var a=function(t,e){return t.listeners(e).length},u=n(32),c=n(6).Buffer,l=e.Uint8Array||function(){};var h=n(5);h.inherits=n(3);var f=n(48),p=void 0;p=f&&f.debuglog?f.debuglog("stream"):function(){};var d,y=n(47),g=n(31);h.inherits(b,u);var m=["error","close","destroy","pause","resume"];function _(t,e){o=o||n(2),t=t||{};var r=e instanceof o;this.objectMode=!!t.objectMode,r&&(this.objectMode=this.objectMode||!!t.readableObjectMode);var i=t.highWaterMark,s=t.readableHighWaterMark,a=this.objectMode?16:16384;this.highWaterMark=i||0===i?i:r&&(s||0===s)?s:a,this.highWaterMark=Math.floor(this.highWaterMark),this.buffer=new y,this.length=0,this.pipes=null,this.pipesCount=0,this.flowing=null,this.ended=!1,this.endEmitted=!1,this.reading=!1,this.sync=!0,this.needReadable=!1,this.emittedReadable=!1,this.readableListening=!1,this.resumeScheduled=!1,this.destroyed=!1,this.defaultEncoding=t.defaultEncoding||"utf8",this.awaitDrain=0,this.readingMore=!1,this.decoder=null,this.encoding=null,t.encoding&&(d||(d=n(22).StringDecoder),this.decoder=new d(t.encoding),this.encoding=t.encoding)}function b(t){if(o=o||n(2),!(this instanceof b))return new b(t);this._readableState=new _(t,this),this.readable=!0,t&&("function"==typeof t.read&&(this._read=t.read),"function"==typeof t.destroy&&(this._destroy=t.destroy)),u.call(this)}function w(t,e,n,r,i){var o,s=t._readableState;null===e?(s.reading=!1,function(t,e){if(e.ended)return;if(e.decoder){var n=e.decoder.end();n&&n.length&&(e.buffer.push(n),e.length+=e.objectMode?1:n.length)}e.ended=!0,x(t)}(t,s)):(i||(o=function(t,e){var n;r=e,c.isBuffer(r)||r instanceof l||"string"==typeof e||void 0===e||t.objectMode||(n=new TypeError("Invalid non-string/buffer chunk"));var r;return n}(s,e)),o?t.emit("error",o):s.objectMode||e&&e.length>0?("string"==typeof e||s.objectMode||Object.getPrototypeOf(e)===c.prototype||(e=function(t){return c.from(t)}(e)),r?s.endEmitted?t.emit("error",new Error("stream.unshift() after end event")):v(t,s,e,!0):s.ended?t.emit("error",new Error("stream.push() after EOF")):(s.reading=!1,s.decoder&&!n?(e=s.decoder.write(e),s.objectMode||0!==e.length?v(t,s,e,!1):D(t,s)):v(t,s,e,!1))):r||(s.reading=!1));return function(t){return!t.ended&&(t.needReadable||t.length<t.highWaterMark||0===t.length)}(s)}function v(t,e,n,r){e.flowing&&0===e.length&&!e.sync?(t.emit("data",n),t.read(0)):(e.length+=e.objectMode?1:n.length,r?e.buffer.unshift(n):e.buffer.push(n),e.needReadable&&x(t)),D(t,e)}Object.defineProperty(b.prototype,"destroyed",{get:function(){return void 0!==this._readableState&&this._readableState.destroyed},set:function(t){this._readableState&&(this._readableState.destroyed=t)}}),b.prototype.destroy=g.destroy,b.prototype._undestroy=g.undestroy,b.prototype._destroy=function(t,e){this.push(null),e(t)},b.prototype.push=function(t,e){var n,r=this._readableState;return r.objectMode?n=!0:"string"==typeof t&&((e=e||r.defaultEncoding)!==r.encoding&&(t=c.from(t,e),e=""),n=!0),w(this,t,e,!1,n)},b.prototype.unshift=function(t){return w(this,t,null,!0,!1)},b.prototype.isPaused=function(){return!1===this._readableState.flowing},b.prototype.setEncoding=function(t){return d||(d=n(22).StringDecoder),this._readableState.decoder=new d(t),this._readableState.encoding=t,this};var E=8388608;function T(t,e){return t<=0||0===e.length&&e.ended?0:e.objectMode?1:t!=t?e.flowing&&e.length?e.buffer.head.data.length:e.length:(t>e.highWaterMark&&(e.highWaterMark=function(t){return t>=E?t=E:(t--,t|=t>>>1,t|=t>>>2,t|=t>>>4,t|=t>>>8,t|=t>>>16,t++),t}(t)),t<=e.length?t:e.ended?e.length:(e.needReadable=!0,0))}function x(t){var e=t._readableState;e.needReadable=!1,e.emittedReadable||(p("emitReadable",e.flowing),e.emittedReadable=!0,e.sync?i.nextTick(A,t):A(t))}function A(t){p("emit readable"),t.emit("readable"),O(t)}function D(t,e){e.readingMore||(e.readingMore=!0,i.nextTick(I,t,e))}function I(t,e){for(var n=e.length;!e.reading&&!e.flowing&&!e.ended&&e.length<e.highWaterMark&&(p("maybeReadMore read 0"),t.read(0),n!==e.length);)n=e.length;e.readingMore=!1}function N(t){p("readable nexttick read 0"),t.read(0)}function S(t,e){e.reading||(p("resume read 0"),t.read(0)),e.resumeScheduled=!1,e.awaitDrain=0,t.emit("resume"),O(t),e.flowing&&!e.reading&&t.read(0)}function O(t){var e=t._readableState;for(p("flow",e.flowing);e.flowing&&null!==t.read(););}function C(t,e){return 0===e.length?null:(e.objectMode?n=e.buffer.shift():!t||t>=e.length?(n=e.decoder?e.buffer.join(""):1===e.buffer.length?e.buffer.head.data:e.buffer.concat(e.length),e.buffer.clear()):n=function(t,e,n){var r;t<e.head.data.length?(r=e.head.data.slice(0,t),e.head.data=e.head.data.slice(t)):r=t===e.head.data.length?e.shift():n?function(t,e){var n=e.head,r=1,i=n.data;t-=i.length;for(;n=n.next;){var o=n.data,s=t>o.length?o.length:t;if(s===o.length?i+=o:i+=o.slice(0,t),0===(t-=s)){s===o.length?(++r,n.next?e.head=n.next:e.head=e.tail=null):(e.head=n,n.data=o.slice(s));break}++r}return e.length-=r,i}(t,e):function(t,e){var n=c.allocUnsafe(t),r=e.head,i=1;r.data.copy(n),t-=r.data.length;for(;r=r.next;){var o=r.data,s=t>o.length?o.length:t;if(o.copy(n,n.length-t,0,s),0===(t-=s)){s===o.length?(++i,r.next?e.head=r.next:e.head=e.tail=null):(e.head=r,r.data=o.slice(s));break}++i}return e.length-=i,n}(t,e);return r}(t,e.buffer,e.decoder),n);var n}function P(t){var e=t._readableState;if(e.length>0)throw new Error('"endReadable()" called on non-empty stream');e.endEmitted||(e.ended=!0,i.nextTick(k,e,t))}function k(t,e){t.endEmitted||0!==t.length||(t.endEmitted=!0,e.readable=!1,e.emit("end"))}function L(t,e){for(var n=0,r=t.length;n<r;n++)if(t[n]===e)return n;return-1}b.prototype.read=function(t){p("read",t),t=parseInt(t,10);var e=this._readableState,n=t;if(0!==t&&(e.emittedReadable=!1),0===t&&e.needReadable&&(e.length>=e.highWaterMark||e.ended))return p("read: emitReadable",e.length,e.ended),0===e.length&&e.ended?P(this):x(this),null;if(0===(t=T(t,e))&&e.ended)return 0===e.length&&P(this),null;var r,i=e.needReadable;return p("need readable",i),(0===e.length||e.length-t<e.highWaterMark)&&p("length less than watermark",i=!0),e.ended||e.reading?p("reading or ended",i=!1):i&&(p("do read"),e.reading=!0,e.sync=!0,0===e.length&&(e.needReadable=!0),this._read(e.highWaterMark),e.sync=!1,e.reading||(t=T(n,e))),null===(r=t>0?C(t,e):null)?(e.needReadable=!0,t=0):e.length-=t,0===e.length&&(e.ended||(e.needReadable=!0),n!==t&&e.ended&&P(this)),null!==r&&this.emit("data",r),r},b.prototype._read=function(t){this.emit("error",new Error("_read() is not implemented"))},b.prototype.pipe=function(t,e){var n=this,o=this._readableState;switch(o.pipesCount){case 0:o.pipes=t;break;case 1:o.pipes=[o.pipes,t];break;default:o.pipes.push(t)}o.pipesCount+=1,p("pipe count=%d opts=%j",o.pipesCount,e);var u=(!e||!1!==e.end)&&t!==r.stdout&&t!==r.stderr?l:b;function c(e,r){p("onunpipe"),e===n&&r&&!1===r.hasUnpiped&&(r.hasUnpiped=!0,p("cleanup"),t.removeListener("close",m),t.removeListener("finish",_),t.removeListener("drain",h),t.removeListener("error",g),t.removeListener("unpipe",c),n.removeListener("end",l),n.removeListener("end",b),n.removeListener("data",y),f=!0,!o.awaitDrain||t._writableState&&!t._writableState.needDrain||h())}function l(){p("onend"),t.end()}o.endEmitted?i.nextTick(u):n.once("end",u),t.on("unpipe",c);var h=function(t){return function(){var e=t._readableState;p("pipeOnDrain",e.awaitDrain),e.awaitDrain&&e.awaitDrain--,0===e.awaitDrain&&a(t,"data")&&(e.flowing=!0,O(t))}}(n);t.on("drain",h);var f=!1;var d=!1;function y(e){p("ondata"),d=!1,!1!==t.write(e)||d||((1===o.pipesCount&&o.pipes===t||o.pipesCount>1&&-1!==L(o.pipes,t))&&!f&&(p("false write response, pause",n._readableState.awaitDrain),n._readableState.awaitDrain++,d=!0),n.pause())}function g(e){p("onerror",e),b(),t.removeListener("error",g),0===a(t,"error")&&t.emit("error",e)}function m(){t.removeListener("finish",_),b()}function _(){p("onfinish"),t.removeListener("close",m),b()}function b(){p("unpipe"),n.unpipe(t)}return n.on("data",y),function(t,e,n){if("function"==typeof t.prependListener)return t.prependListener(e,n);t._events&&t._events[e]?s(t._events[e])?t._events[e].unshift(n):t._events[e]=[n,t._events[e]]:t.on(e,n)}(t,"error",g),t.once("close",m),t.once("finish",_),t.emit("pipe",n),o.flowing||(p("pipe resume"),n.resume()),t},b.prototype.unpipe=function(t){var e=this._readableState,n={hasUnpiped:!1};if(0===e.pipesCount)return this;if(1===e.pipesCount)return t&&t!==e.pipes?this:(t||(t=e.pipes),e.pipes=null,e.pipesCount=0,e.flowing=!1,t&&t.emit("unpipe",this,n),this);if(!t){var r=e.pipes,i=e.pipesCount;e.pipes=null,e.pipesCount=0,e.flowing=!1;for(var o=0;o<i;o++)r[o].emit("unpipe",this,n);return this}var s=L(e.pipes,t);return-1===s?this:(e.pipes.splice(s,1),e.pipesCount-=1,1===e.pipesCount&&(e.pipes=e.pipes[0]),t.emit("unpipe",this,n),this)},b.prototype.on=function(t,e){var n=u.prototype.on.call(this,t,e);if("data"===t)!1!==this._readableState.flowing&&this.resume();else if("readable"===t){var r=this._readableState;r.endEmitted||r.readableListening||(r.readableListening=r.needReadable=!0,r.emittedReadable=!1,r.reading?r.length&&x(this):i.nextTick(N,this))}return n},b.prototype.addListener=b.prototype.on,b.prototype.resume=function(){var t=this._readableState;return t.flowing||(p("resume"),t.flowing=!0,function(t,e){e.resumeScheduled||(e.resumeScheduled=!0,i.nextTick(S,t,e))}(this,t)),this},b.prototype.pause=function(){return p("call pause flowing=%j",this._readableState.flowing),!1!==this._readableState.flowing&&(p("pause"),this._readableState.flowing=!1,this.emit("pause")),this},b.prototype.wrap=function(t){var e=this,n=this._readableState,r=!1;for(var i in t.on("end",function(){if(p("wrapped end"),n.decoder&&!n.ended){var t=n.decoder.end();t&&t.length&&e.push(t)}e.push(null)}),t.on("data",function(i){(p("wrapped data"),n.decoder&&(i=n.decoder.write(i)),!n.objectMode||null!==i&&void 0!==i)&&((n.objectMode||i&&i.length)&&(e.push(i)||(r=!0,t.pause())))}),t)void 0===this[i]&&"function"==typeof t[i]&&(this[i]=function(e){return function(){return t[e].apply(t,arguments)}}(i));for(var o=0;o<m.length;o++)t.on(m[o],this.emit.bind(this,m[o]));return this._read=function(e){p("wrapped _read",e),r&&(r=!1,t.resume())},this},Object.defineProperty(b.prototype,"readableHighWaterMark",{enumerable:!1,get:function(){return this._readableState.highWaterMark}}),b._fromList=C}).call(this,n(4),n(8))},function(t,e){var n={}.toString;t.exports=Array.isArray||function(t){return"[object Array]"==n.call(t)}},function(t,e){(function(){var e={}.hasOwnProperty;t.exports=function(){function t(t){var n,r,i,o,s,a,u,c,l;for(n in t||(t={}),this.pretty=t.pretty||!1,this.allowEmpty=null!=(r=t.allowEmpty)&&r,this.pretty?(this.indent=null!=(i=t.indent)?i:"  ",this.newline=null!=(o=t.newline)?o:"\n",this.offset=null!=(s=t.offset)?s:0,this.dontprettytextnodes=null!=(a=t.dontprettytextnodes)?a:0):(this.indent="",this.newline="",this.offset=0,this.dontprettytextnodes=0),this.spacebeforeslash=null!=(u=t.spacebeforeslash)?u:"",!0===this.spacebeforeslash&&(this.spacebeforeslash=" "),this.newlinedefault=this.newline,this.prettydefault=this.pretty,c=t.writer||{})e.call(c,n)&&(l=c[n],this[n]=l)}return t.prototype.set=function(t){var n,r,i;for(n in t||(t={}),"pretty"in t&&(this.pretty=t.pretty),"allowEmpty"in t&&(this.allowEmpty=t.allowEmpty),this.pretty?(this.indent="indent"in t?t.indent:"  ",this.newline="newline"in t?t.newline:"\n",this.offset="offset"in t?t.offset:0,this.dontprettytextnodes="dontprettytextnodes"in t?t.dontprettytextnodes:0):(this.indent="",this.newline="",this.offset=0,this.dontprettytextnodes=0),this.spacebeforeslash="spacebeforeslash"in t?t.spacebeforeslash:"",!0===this.spacebeforeslash&&(this.spacebeforeslash=" "),this.newlinedefault=this.newline,this.prettydefault=this.pretty,r=t.writer||{})e.call(r,n)&&(i=r[n],this[n]=i);return this},t.prototype.space=function(t){var e;return this.pretty&&(e=(t||0)+this.offset+1)>0?new Array(e).join(this.indent):""},t}()}).call(this)},function(t,e){(function(){var e=function(t,e){return function(){return t.apply(e,arguments)}},n={}.hasOwnProperty;t.exports=function(){function t(t){var r,i,o;for(r in this.assertLegalChar=e(this.assertLegalChar,this),t||(t={}),this.noDoubleEncoding=t.noDoubleEncoding,i=t.stringify||{})n.call(i,r)&&(o=i[r],this[r]=o)}return t.prototype.eleName=function(t){return t=""+t||"",this.assertLegalChar(t)},t.prototype.eleText=function(t){return t=""+t||"",this.assertLegalChar(this.elEscape(t))},t.prototype.cdata=function(t){return t=(t=""+t||"").replace("]]>","]]]]><![CDATA[>"),this.assertLegalChar(t)},t.prototype.comment=function(t){if((t=""+t||"").match(/--/))throw new Error("Comment text cannot contain double-hypen: "+t);return this.assertLegalChar(t)},t.prototype.raw=function(t){return""+t||""},t.prototype.attName=function(t){return""+t||""},t.prototype.attValue=function(t){return t=""+t||"",this.attEscape(t)},t.prototype.insTarget=function(t){return""+t||""},t.prototype.insValue=function(t){if((t=""+t||"").match(/\?>/))throw new Error("Invalid processing instruction value: "+t);return t},t.prototype.xmlVersion=function(t){if(!(t=""+t||"").match(/1\.[0-9]+/))throw new Error("Invalid version number: "+t);return t},t.prototype.xmlEncoding=function(t){if(!(t=""+t||"").match(/^[A-Za-z](?:[A-Za-z0-9._-])*$/))throw new Error("Invalid encoding: "+t);return t},t.prototype.xmlStandalone=function(t){return t?"yes":"no"},t.prototype.dtdPubID=function(t){return""+t||""},t.prototype.dtdSysID=function(t){return""+t||""},t.prototype.dtdElementValue=function(t){return""+t||""},t.prototype.dtdAttType=function(t){return""+t||""},t.prototype.dtdAttDefault=function(t){return null!=t?""+t||"":t},t.prototype.dtdEntityValue=function(t){return""+t||""},t.prototype.dtdNData=function(t){return""+t||""},t.prototype.convertAttKey="@",t.prototype.convertPIKey="?",t.prototype.convertTextKey="#text",t.prototype.convertCDataKey="#cdata",t.prototype.convertCommentKey="#comment",t.prototype.convertRawKey="#raw",t.prototype.assertLegalChar=function(t){var e;if(e=t.match(/[\0\uFFFE\uFFFF]|[\uD800-\uDBFF](?![\uDC00-\uDFFF])|(?:[^\uD800-\uDBFF]|^)[\uDC00-\uDFFF]/))throw new Error("Invalid character in string: "+t+" at index "+e.index);return t},t.prototype.elEscape=function(t){var e;return e=this.noDoubleEncoding?/(?!&\S+;)&/g:/&/g,t.replace(e,"&amp;").replace(/</g,"&lt;").replace(/>/g,"&gt;").replace(/\r/g,"&#xD;")},t.prototype.attEscape=function(t){var e;return e=this.noDoubleEncoding?/(?!&\S+;)&/g:/&/g,t.replace(e,"&amp;").replace(/</g,"&lt;").replace(/"/g,"&quot;").replace(/\t/g,"&#x9;").replace(/\n/g,"&#xA;").replace(/\r/g,"&#xD;")},t}()}).call(this)},function(t,e){(function(){t.exports=function(){function t(t,e,n){if(this.options=t.options,this.stringify=t.stringify,null==e)throw new Error("Missing attribute name of element "+t.name);if(null==n)throw new Error("Missing attribute value for attribute "+e+" of element "+t.name);this.name=this.stringify.attName(e),this.value=this.stringify.attValue(n)}return t.prototype.clone=function(){return Object.create(this)},t.prototype.toString=function(t){return this.options.writer.set(t).attribute(this)},t}()}).call(this)},function(t,e){(function(){"use strict";e.stripBOM=function(t){return"\ufeff"===t[0]?t.substring(1):t}}).call(this)},function(t,e,n){t.exports=n(24).PassThrough},function(t,e,n){t.exports=n(24).Transform},function(t,e,n){t.exports=n(2)},function(t,e,n){t.exports=n(23)},function(t,e,n){"use strict";t.exports=o;var r=n(29),i=n(5);function o(t){if(!(this instanceof o))return new o(t);r.call(this,t)}i.inherits=n(3),i.inherits(o,r),o.prototype._transform=function(t,e,n){n(null,t)}},function(t,e,n){(function(e){function n(t){try{if(!e.localStorage)return!1}catch(t){return!1}var n=e.localStorage[t];return null!=n&&"true"===String(n).toLowerCase()}t.exports=function(t,e){if(n("noDeprecation"))return t;var r=!1;return function(){if(!r){if(n("throwDeprecation"))throw new Error(e);n("traceDeprecation")?console.trace(e):console.warn(e),r=!0}return t.apply(this,arguments)}}}).call(this,n(4))},function(t,e,n){(function(t,e){!function(t,n){"use strict";if(!t.setImmediate){var r,i,o,s,a,u=1,c={},l=!1,h=t.document,f=Object.getPrototypeOf&&Object.getPrototypeOf(t);f=f&&f.setTimeout?f:t,"[object process]"==={}.toString.call(t.process)?r=function(t){e.nextTick(function(){d(t)})}:!function(){if(t.postMessage&&!t.importScripts){var e=!0,n=t.onmessage;return t.onmessage=function(){e=!1},t.postMessage("","*"),t.onmessage=n,e}}()?t.MessageChannel?((o=new MessageChannel).port1.onmessage=function(t){d(t.data)},r=function(t){o.port2.postMessage(t)}):h&&"onreadystatechange"in h.createElement("script")?(i=h.documentElement,r=function(t){var e=h.createElement("script");e.onreadystatechange=function(){d(t),e.onreadystatechange=null,i.removeChild(e),e=null},i.appendChild(e)}):r=function(t){setTimeout(d,0,t)}:(s="setImmediate$"+Math.random()+"$",a=function(e){e.source===t&&"string"==typeof e.data&&0===e.data.indexOf(s)&&d(+e.data.slice(s.length))},t.addEventListener?t.addEventListener("message",a,!1):t.attachEvent("onmessage",a),r=function(e){t.postMessage(s+e,"*")}),f.setImmediate=function(t){"function"!=typeof t&&(t=new Function(""+t));for(var e=new Array(arguments.length-1),n=0;n<e.length;n++)e[n]=arguments[n+1];var i={callback:t,args:e};return c[u]=i,r(u),u++},f.clearImmediate=p}function p(t){delete c[t]}function d(t){if(l)setTimeout(d,0,t);else{var e=c[t];if(e){l=!0;try{!function(t){var e=t.callback,r=t.args;switch(r.length){case 0:e();break;case 1:e(r[0]);break;case 2:e(r[0],r[1]);break;case 3:e(r[0],r[1],r[2]);break;default:e.apply(n,r)}}(e)}finally{p(t),l=!1}}}}}("undefined"==typeof self?void 0===t?this:t:self)}).call(this,n(4),n(8))},function(t,e){},function(t,e,n){"use strict";var r=n(6).Buffer,i=n(46);t.exports=function(){function t(){!function(t,e){if(!(t instanceof e))throw new TypeError("Cannot call a class as a function")}(this,t),this.head=null,this.tail=null,this.length=0}return t.prototype.push=function(t){var e={data:t,next:null};this.length>0?this.tail.next=e:this.head=e,this.tail=e,++this.length},t.prototype.unshift=function(t){var e={data:t,next:this.head};0===this.length&&(this.tail=e),this.head=e,++this.length},t.prototype.shift=function(){if(0!==this.length){var t=this.head.data;return 1===this.length?this.head=this.tail=null:this.head=this.head.next,--this.length,t}},t.prototype.clear=function(){this.head=this.tail=null,this.length=0},t.prototype.join=function(t){if(0===this.length)return"";for(var e=this.head,n=""+e.data;e=e.next;)n+=t+e.data;return n},t.prototype.concat=function(t){if(0===this.length)return r.alloc(0);if(1===this.length)return this.head.data;for(var e,n,i,o=r.allocUnsafe(t>>>0),s=this.head,a=0;s;)e=s.data,n=o,i=a,e.copy(n,i),a+=s.data.length,s=s.next;return o},t}(),i&&i.inspect&&i.inspect.custom&&(t.exports.prototype[i.inspect.custom]=function(){var t=i.inspect({length:this.length});return this.constructor.name+" "+t})},function(t,e){},function(t,e,n){t.exports=i;var r=n(9).EventEmitter;function i(){r.call(this)}n(3)(i,r),i.Readable=n(24),i.Writable=n(42),i.Duplex=n(41),i.Transform=n(40),i.PassThrough=n(39),i.Stream=i,i.prototype.pipe=function(t,e){var n=this;function i(e){t.writable&&!1===t.write(e)&&n.pause&&n.pause()}function o(){n.readable&&n.resume&&n.resume()}n.on("data",i),t.on("drain",o),t._isStdio||e&&!1===e.end||(n.on("end",a),n.on("close",u));var s=!1;function a(){s||(s=!0,t.end())}function u(){s||(s=!0,"function"==typeof t.destroy&&t.destroy())}function c(t){if(l(),0===r.listenerCount(this,"error"))throw t}function l(){n.removeListener("data",i),t.removeListener("drain",o),n.removeListener("end",a),n.removeListener("close",u),n.removeListener("error",c),t.removeListener("error",c),n.removeListener("end",l),n.removeListener("close",l),t.removeListener("close",l)}return n.on("error",c),t.on("error",c),n.on("end",l),n.on("close",l),t.on("close",l),t.emit("pipe",n),t}},function(t,e){e.read=function(t,e,n,r,i){var o,s,a=8*i-r-1,u=(1<<a)-1,c=u>>1,l=-7,h=n?i-1:0,f=n?-1:1,p=t[e+h];for(h+=f,o=p&(1<<-l)-1,p>>=-l,l+=a;l>0;o=256*o+t[e+h],h+=f,l-=8);for(s=o&(1<<-l)-1,o>>=-l,l+=r;l>0;s=256*s+t[e+h],h+=f,l-=8);if(0===o)o=1-c;else{if(o===u)return s?NaN:1/0*(p?-1:1);s+=Math.pow(2,r),o-=c}return(p?-1:1)*s*Math.pow(2,o-r)},e.write=function(t,e,n,r,i,o){var s,a,u,c=8*o-i-1,l=(1<<c)-1,h=l>>1,f=23===i?Math.pow(2,-24)-Math.pow(2,-77):0,p=r?0:o-1,d=r?1:-1,y=e<0||0===e&&1/e<0?1:0;for(e=Math.abs(e),isNaN(e)||e===1/0?(a=isNaN(e)?1:0,s=l):(s=Math.floor(Math.log(e)/Math.LN2),e*(u=Math.pow(2,-s))<1&&(s--,u*=2),(e+=s+h>=1?f/u:f*Math.pow(2,1-h))*u>=2&&(s++,u/=2),s+h>=l?(a=0,s=l):s+h>=1?(a=(e*u-1)*Math.pow(2,i),s+=h):(a=e*Math.pow(2,h-1)*Math.pow(2,i),s=0));i>=8;t[n+p]=255&a,p+=d,a/=256,i-=8);for(s=s<<i|a,c+=i;c>0;t[n+p]=255&s,p+=d,s/=256,c-=8);t[n+p-d]|=128*y}},function(t,e,n){"use strict";e.byteLength=function(t){var e=c(t),n=e[0],r=e[1];return 3*(n+r)/4-r},e.toByteArray=function(t){for(var e,n=c(t),r=n[0],s=n[1],a=new o(function(t,e,n){return 3*(e+n)/4-n}(0,r,s)),u=0,l=s>0?r-4:r,h=0;h<l;h+=4)e=i[t.charCodeAt(h)]<<18|i[t.charCodeAt(h+1)]<<12|i[t.charCodeAt(h+2)]<<6|i[t.charCodeAt(h+3)],a[u++]=e>>16&255,a[u++]=e>>8&255,a[u++]=255&e;2===s&&(e=i[t.charCodeAt(h)]<<2|i[t.charCodeAt(h+1)]>>4,a[u++]=255&e);1===s&&(e=i[t.charCodeAt(h)]<<10|i[t.charCodeAt(h+1)]<<4|i[t.charCodeAt(h+2)]>>2,a[u++]=e>>8&255,a[u++]=255&e);return a},e.fromByteArray=function(t){for(var e,n=t.length,i=n%3,o=[],s=0,a=n-i;s<a;s+=16383)o.push(l(t,s,s+16383>a?a:s+16383));1===i?(e=t[n-1],o.push(r[e>>2]+r[e<<4&63]+"==")):2===i&&(e=(t[n-2]<<8)+t[n-1],o.push(r[e>>10]+r[e>>4&63]+r[e<<2&63]+"="));return o.join("")};for(var r=[],i=[],o="undefined"!=typeof Uint8Array?Uint8Array:Array,s="ABCDEFGHIJKLMNOPQRSTUVWXYZabcdefghijklmnopqrstuvwxyz0123456789+/",a=0,u=s.length;a<u;++a)r[a]=s[a],i[s.charCodeAt(a)]=a;function c(t){var e=t.length;if(e%4>0)throw new Error("Invalid string. Length must be a multiple of 4");var n=t.indexOf("=");return-1===n&&(n=e),[n,n===e?0:4-n%4]}function l(t,e,n){for(var i,o,s=[],a=e;a<n;a+=3)i=(t[a]<<16&16711680)+(t[a+1]<<8&65280)+(255&t[a+2]),s.push(r[(o=i)>>18&63]+r[o>>12&63]+r[o>>6&63]+r[63&o]);return s.join("")}i["-".charCodeAt(0)]=62,i["_".charCodeAt(0)]=63},function(t,e,n){(function(t){!function(e){e.parser=function(t,e){return new o(t,e)},e.SAXParser=o,e.SAXStream=a,e.createStream=function(t,e){return new a(t,e)},e.MAX_BUFFER_LENGTH=65536;var r,i=["comment","sgmlDecl","textNode","tagName","doctype","procInstName","procInstBody","entity","attribName","attribValue","cdata","script"];function o(t,n){if(!(this instanceof o))return new o(t,n);!function(t){for(var e=0,n=i.length;e<n;e++)t[i[e]]=""}(this),this.q=this.c="",this.bufferCheckPosition=e.MAX_BUFFER_LENGTH,this.opt=n||{},this.opt.lowercase=this.opt.lowercase||this.opt.lowercasetags,this.looseCase=this.opt.lowercase?"toLowerCase":"toUpperCase",this.tags=[],this.closed=this.closedRoot=this.sawRoot=!1,this.tag=this.error=null,this.strict=!!t,this.noscript=!(!t&&!this.opt.noscript),this.state=A.BEGIN,this.strictEntities=this.opt.strictEntities,this.ENTITIES=this.strictEntities?Object.create(e.XML_ENTITIES):Object.create(e.ENTITIES),this.attribList=[],this.opt.xmlns&&(this.ns=Object.create(f)),this.trackPosition=!1!==this.opt.position,this.trackPosition&&(this.position=this.line=this.column=0),I(this,"onready")}e.EVENTS=["text","processinginstruction","sgmldeclaration","doctype","comment","opentagstart","attribute","opentag","closetag","opencdata","cdata","closecdata","error","end","ready","script","opennamespace","closenamespace"],Object.create||(Object.create=function(t){function e(){}return e.prototype=t,new e}),Object.keys||(Object.keys=function(t){var e=[];for(var n in t)t.hasOwnProperty(n)&&e.push(n);return e}),o.prototype={end:function(){P(this)},write:function(t){if(this.error)throw this.error;if(this.closed)return C(this,"Cannot write after close. Assign an onready handler.");if(null===t)return P(this);"object"==typeof t&&(t=t.toString());var n=0,r="";for(;r=Y(t,n++),this.c=r,r;)switch(this.trackPosition&&(this.position++,"\n"===r?(this.line++,this.column=0):this.column++),this.state){case A.BEGIN:if(this.state=A.BEGIN_WHITESPACE,"\ufeff"===r)continue;j(this,r);continue;case A.BEGIN_WHITESPACE:j(this,r);continue;case A.TEXT:if(this.sawRoot&&!this.closedRoot){for(var o=n-1;r&&"<"!==r&&"&"!==r;)(r=Y(t,n++))&&this.trackPosition&&(this.position++,"\n"===r?(this.line++,this.column=0):this.column++);this.textNode+=t.substring(o,n-1)}"<"!==r||this.sawRoot&&this.closedRoot&&!this.strict?(m(r)||this.sawRoot&&!this.closedRoot||k(this,"Text data outside of root node."),"&"===r?this.state=A.TEXT_ENTITY:this.textNode+=r):(this.state=A.OPEN_WAKA,this.startTagPosition=this.position);continue;case A.SCRIPT:"<"===r?this.state=A.SCRIPT_ENDING:this.script+=r;continue;case A.SCRIPT_ENDING:"/"===r?this.state=A.CLOSE_TAG:(this.script+="<"+r,this.state=A.SCRIPT);continue;case A.OPEN_WAKA:if("!"===r)this.state=A.SGML_DECL,this.sgmlDecl="";else if(m(r));else if(w(p,r))this.state=A.OPEN_TAG,this.tagName=r;else if("/"===r)this.state=A.CLOSE_TAG,this.tagName="";else if("?"===r)this.state=A.PROC_INST,this.procInstName=this.procInstBody="";else{if(k(this,"Unencoded <"),this.startTagPosition+1<this.position){var s=this.position-this.startTagPosition;r=new Array(s).join(" ")+r}this.textNode+="<"+r,this.state=A.TEXT}continue;case A.SGML_DECL:(this.sgmlDecl+r).toUpperCase()===u?(N(this,"onopencdata"),this.state=A.CDATA,this.sgmlDecl="",this.cdata=""):this.sgmlDecl+r==="--"?(this.state=A.COMMENT,this.comment="",this.sgmlDecl=""):(this.sgmlDecl+r).toUpperCase()===c?(this.state=A.DOCTYPE,(this.doctype||this.sawRoot)&&k(this,"Inappropriately located doctype declaration"),this.doctype="",this.sgmlDecl=""):">"===r?(N(this,"onsgmldeclaration",this.sgmlDecl),this.sgmlDecl="",this.state=A.TEXT):_(r)?(this.state=A.SGML_DECL_QUOTED,this.sgmlDecl+=r):this.sgmlDecl+=r;continue;case A.SGML_DECL_QUOTED:r===this.q&&(this.state=A.SGML_DECL,this.q=""),this.sgmlDecl+=r;continue;case A.DOCTYPE:">"===r?(this.state=A.TEXT,N(this,"ondoctype",this.doctype),this.doctype=!0):(this.doctype+=r,"["===r?this.state=A.DOCTYPE_DTD:_(r)&&(this.state=A.DOCTYPE_QUOTED,this.q=r));continue;case A.DOCTYPE_QUOTED:this.doctype+=r,r===this.q&&(this.q="",this.state=A.DOCTYPE);continue;case A.DOCTYPE_DTD:this.doctype+=r,"]"===r?this.state=A.DOCTYPE:_(r)&&(this.state=A.DOCTYPE_DTD_QUOTED,this.q=r);continue;case A.DOCTYPE_DTD_QUOTED:this.doctype+=r,r===this.q&&(this.state=A.DOCTYPE_DTD,this.q="");continue;case A.COMMENT:"-"===r?this.state=A.COMMENT_ENDING:this.comment+=r;continue;case A.COMMENT_ENDING:"-"===r?(this.state=A.COMMENT_ENDED,this.comment=O(this.opt,this.comment),this.comment&&N(this,"oncomment",this.comment),this.comment=""):(this.comment+="-"+r,this.state=A.COMMENT);continue;case A.COMMENT_ENDED:">"!==r?(k(this,"Malformed comment"),this.comment+="--"+r,this.state=A.COMMENT):this.state=A.TEXT;continue;case A.CDATA:"]"===r?this.state=A.CDATA_ENDING:this.cdata+=r;continue;case A.CDATA_ENDING:"]"===r?this.state=A.CDATA_ENDING_2:(this.cdata+="]"+r,this.state=A.CDATA);continue;case A.CDATA_ENDING_2:">"===r?(this.cdata&&N(this,"oncdata",this.cdata),N(this,"onclosecdata"),this.cdata="",this.state=A.TEXT):"]"===r?this.cdata+="]":(this.cdata+="]]"+r,this.state=A.CDATA);continue;case A.PROC_INST:"?"===r?this.state=A.PROC_INST_ENDING:m(r)?this.state=A.PROC_INST_BODY:this.procInstName+=r;continue;case A.PROC_INST_BODY:if(!this.procInstBody&&m(r))continue;"?"===r?this.state=A.PROC_INST_ENDING:this.procInstBody+=r;continue;case A.PROC_INST_ENDING:">"===r?(N(this,"onprocessinginstruction",{name:this.procInstName,body:this.procInstBody}),this.procInstName=this.procInstBody="",this.state=A.TEXT):(this.procInstBody+="?"+r,this.state=A.PROC_INST_BODY);continue;case A.OPEN_TAG:w(d,r)?this.tagName+=r:(L(this),">"===r?B(this):"/"===r?this.state=A.OPEN_TAG_SLASH:(m(r)||k(this,"Invalid character in tag name"),this.state=A.ATTRIB));continue;case A.OPEN_TAG_SLASH:">"===r?(B(this,!0),U(this)):(k(this,"Forward-slash in opening tag not followed by >"),this.state=A.ATTRIB);continue;case A.ATTRIB:if(m(r))continue;">"===r?B(this):"/"===r?this.state=A.OPEN_TAG_SLASH:w(p,r)?(this.attribName=r,this.attribValue="",this.state=A.ATTRIB_NAME):k(this,"Invalid attribute name");continue;case A.ATTRIB_NAME:"="===r?this.state=A.ATTRIB_VALUE:">"===r?(k(this,"Attribute without value"),this.attribValue=this.attribName,M(this),B(this)):m(r)?this.state=A.ATTRIB_NAME_SAW_WHITE:w(d,r)?this.attribName+=r:k(this,"Invalid attribute name");continue;case A.ATTRIB_NAME_SAW_WHITE:if("="===r)this.state=A.ATTRIB_VALUE;else{if(m(r))continue;k(this,"Attribute without value"),this.tag.attributes[this.attribName]="",this.attribValue="",N(this,"onattribute",{name:this.attribName,value:""}),this.attribName="",">"===r?B(this):w(p,r)?(this.attribName=r,this.state=A.ATTRIB_NAME):(k(this,"Invalid attribute name"),this.state=A.ATTRIB)}continue;case A.ATTRIB_VALUE:if(m(r))continue;_(r)?(this.q=r,this.state=A.ATTRIB_VALUE_QUOTED):(k(this,"Unquoted attribute value"),this.state=A.ATTRIB_VALUE_UNQUOTED,this.attribValue=r);continue;case A.ATTRIB_VALUE_QUOTED:if(r!==this.q){"&"===r?this.state=A.ATTRIB_VALUE_ENTITY_Q:this.attribValue+=r;continue}M(this),this.q="",this.state=A.ATTRIB_VALUE_CLOSED;continue;case A.ATTRIB_VALUE_CLOSED:m(r)?this.state=A.ATTRIB:">"===r?B(this):"/"===r?this.state=A.OPEN_TAG_SLASH:w(p,r)?(k(this,"No whitespace between attributes"),this.attribName=r,this.attribValue="",this.state=A.ATTRIB_NAME):k(this,"Invalid attribute name");continue;case A.ATTRIB_VALUE_UNQUOTED:if(!b(r)){"&"===r?this.state=A.ATTRIB_VALUE_ENTITY_U:this.attribValue+=r;continue}M(this),">"===r?B(this):this.state=A.ATTRIB;continue;case A.CLOSE_TAG:if(this.tagName)">"===r?U(this):w(d,r)?this.tagName+=r:this.script?(this.script+="</"+this.tagName,this.tagName="",this.state=A.SCRIPT):(m(r)||k(this,"Invalid tagname in closing tag"),this.state=A.CLOSE_TAG_SAW_WHITE);else{if(m(r))continue;v(p,r)?this.script?(this.script+="</"+r,this.state=A.SCRIPT):k(this,"Invalid tagname in closing tag."):this.tagName=r}continue;case A.CLOSE_TAG_SAW_WHITE:if(m(r))continue;">"===r?U(this):k(this,"Invalid characters in closing tag");continue;case A.TEXT_ENTITY:case A.ATTRIB_VALUE_ENTITY_Q:case A.ATTRIB_VALUE_ENTITY_U:var a,l;switch(this.state){case A.TEXT_ENTITY:a=A.TEXT,l="textNode";break;case A.ATTRIB_VALUE_ENTITY_Q:a=A.ATTRIB_VALUE_QUOTED,l="attribValue";break;case A.ATTRIB_VALUE_ENTITY_U:a=A.ATTRIB_VALUE_UNQUOTED,l="attribValue"}";"===r?(this[l]+=F(this),this.entity="",this.state=a):w(this.entity.length?g:y,r)?this.entity+=r:(k(this,"Invalid character in entity name"),this[l]+="&"+this.entity+r,this.entity="",this.state=a);continue;default:throw new Error(this,"Unknown state: "+this.state)}this.position>=this.bufferCheckPosition&&function(t){for(var n=Math.max(e.MAX_BUFFER_LENGTH,10),r=0,o=0,s=i.length;o<s;o++){var a=t[i[o]].length;if(a>n)switch(i[o]){case"textNode":S(t);break;case"cdata":N(t,"oncdata",t.cdata),t.cdata="";break;case"script":N(t,"onscript",t.script),t.script="";break;default:C(t,"Max buffer length exceeded: "+i[o])}r=Math.max(r,a)}var u=e.MAX_BUFFER_LENGTH-r;t.bufferCheckPosition=u+t.position}(this);return this}
/*! http://mths.be/fromcodepoint v0.1.0 by @mathias */,resume:function(){return this.error=null,this},close:function(){return this.write(null)},flush:function(){var t;S(t=this),""!==t.cdata&&(N(t,"oncdata",t.cdata),t.cdata=""),""!==t.script&&(N(t,"onscript",t.script),t.script="")}};try{r=n(49).Stream}catch(t){r=function(){}}var s=e.EVENTS.filter(function(t){return"error"!==t&&"end"!==t});function a(t,e){if(!(this instanceof a))return new a(t,e);r.apply(this),this._parser=new o(t,e),this.writable=!0,this.readable=!0;var n=this;this._parser.onend=function(){n.emit("end")},this._parser.onerror=function(t){n.emit("error",t),n._parser.error=null},this._decoder=null,s.forEach(function(t){Object.defineProperty(n,"on"+t,{get:function(){return n._parser["on"+t]},set:function(e){if(!e)return n.removeAllListeners(t),n._parser["on"+t]=e,e;n.on(t,e)},enumerable:!0,configurable:!1})})}a.prototype=Object.create(r.prototype,{constructor:{value:a}}),a.prototype.write=function(e){if("function"==typeof t&&"function"==typeof t.isBuffer&&t.isBuffer(e)){if(!this._decoder){var r=n(22).StringDecoder;this._decoder=new r("utf8")}e=this._decoder.write(e)}return this._parser.write(e.toString()),this.emit("data",e),!0},a.prototype.end=function(t){return t&&t.length&&this.write(t),this._parser.end(),!0},a.prototype.on=function(t,e){var n=this;return n._parser["on"+t]||-1===s.indexOf(t)||(n._parser["on"+t]=function(){var e=1===arguments.length?[arguments[0]]:Array.apply(null,arguments);e.splice(0,0,t),n.emit.apply(n,e)}),r.prototype.on.call(n,t,e)};var u="[CDATA[",c="DOCTYPE",l="http://www.w3.org/XML/1998/namespace",h="http://www.w3.org/2000/xmlns/",f={xml:l,xmlns:h},p=/[:_A-Za-z\u00C0-\u00D6\u00D8-\u00F6\u00F8-\u02FF\u0370-\u037D\u037F-\u1FFF\u200C-\u200D\u2070-\u218F\u2C00-\u2FEF\u3001-\uD7FF\uF900-\uFDCF\uFDF0-\uFFFD]/,d=/[:_A-Za-z\u00C0-\u00D6\u00D8-\u00F6\u00F8-\u02FF\u0370-\u037D\u037F-\u1FFF\u200C-\u200D\u2070-\u218F\u2C00-\u2FEF\u3001-\uD7FF\uF900-\uFDCF\uFDF0-\uFFFD\u00B7\u0300-\u036F\u203F-\u2040.\d-]/,y=/[#:_A-Za-z\u00C0-\u00D6\u00D8-\u00F6\u00F8-\u02FF\u0370-\u037D\u037F-\u1FFF\u200C-\u200D\u2070-\u218F\u2C00-\u2FEF\u3001-\uD7FF\uF900-\uFDCF\uFDF0-\uFFFD]/,g=/[#:_A-Za-z\u00C0-\u00D6\u00D8-\u00F6\u00F8-\u02FF\u0370-\u037D\u037F-\u1FFF\u200C-\u200D\u2070-\u218F\u2C00-\u2FEF\u3001-\uD7FF\uF900-\uFDCF\uFDF0-\uFFFD\u00B7\u0300-\u036F\u203F-\u2040.\d-]/;function m(t){return" "===t||"\n"===t||"\r"===t||"\t"===t}function _(t){return'"'===t||"'"===t}function b(t){return">"===t||m(t)}function w(t,e){return t.test(e)}function v(t,e){return!w(t,e)}var E,T,x,A=0;for(var D in e.STATE={BEGIN:A++,BEGIN_WHITESPACE:A++,TEXT:A++,TEXT_ENTITY:A++,OPEN_WAKA:A++,SGML_DECL:A++,SGML_DECL_QUOTED:A++,DOCTYPE:A++,DOCTYPE_QUOTED:A++,DOCTYPE_DTD:A++,DOCTYPE_DTD_QUOTED:A++,COMMENT_STARTING:A++,COMMENT:A++,COMMENT_ENDING:A++,COMMENT_ENDED:A++,CDATA:A++,CDATA_ENDING:A++,CDATA_ENDING_2:A++,PROC_INST:A++,PROC_INST_BODY:A++,PROC_INST_ENDING:A++,OPEN_TAG:A++,OPEN_TAG_SLASH:A++,ATTRIB:A++,ATTRIB_NAME:A++,ATTRIB_NAME_SAW_WHITE:A++,ATTRIB_VALUE:A++,ATTRIB_VALUE_QUOTED:A++,ATTRIB_VALUE_CLOSED:A++,ATTRIB_VALUE_UNQUOTED:A++,ATTRIB_VALUE_ENTITY_Q:A++,ATTRIB_VALUE_ENTITY_U:A++,CLOSE_TAG:A++,CLOSE_TAG_SAW_WHITE:A++,SCRIPT:A++,SCRIPT_ENDING:A++},e.XML_ENTITIES={amp:"&",gt:">",lt:"<",quot:'"',apos:"'"},e.ENTITIES={amp:"&",gt:">",lt:"<",quot:'"',apos:"'",AElig:198,Aacute:193,Acirc:194,Agrave:192,Aring:197,Atilde:195,Auml:196,Ccedil:199,ETH:208,Eacute:201,Ecirc:202,Egrave:200,Euml:203,Iacute:205,Icirc:206,Igrave:204,Iuml:207,Ntilde:209,Oacute:211,Ocirc:212,Ograve:210,Oslash:216,Otilde:213,Ouml:214,THORN:222,Uacute:218,Ucirc:219,Ugrave:217,Uuml:220,Yacute:221,aacute:225,acirc:226,aelig:230,agrave:224,aring:229,atilde:227,auml:228,ccedil:231,eacute:233,ecirc:234,egrave:232,eth:240,euml:235,iacute:237,icirc:238,igrave:236,iuml:239,ntilde:241,oacute:243,ocirc:244,ograve:242,oslash:248,otilde:245,ouml:246,szlig:223,thorn:254,uacute:250,ucirc:251,ugrave:249,uuml:252,yacute:253,yuml:255,copy:169,reg:174,nbsp:160,iexcl:161,cent:162,pound:163,curren:164,yen:165,brvbar:166,sect:167,uml:168,ordf:170,laquo:171,not:172,shy:173,macr:175,deg:176,plusmn:177,sup1:185,sup2:178,sup3:179,acute:180,micro:181,para:182,middot:183,cedil:184,ordm:186,raquo:187,frac14:188,frac12:189,frac34:190,iquest:191,times:215,divide:247,OElig:338,oelig:339,Scaron:352,scaron:353,Yuml:376,fnof:402,circ:710,tilde:732,Alpha:913,Beta:914,Gamma:915,Delta:916,Epsilon:917,Zeta:918,Eta:919,Theta:920,Iota:921,Kappa:922,Lambda:923,Mu:924,Nu:925,Xi:926,Omicron:927,Pi:928,Rho:929,Sigma:931,Tau:932,Upsilon:933,Phi:934,Chi:935,Psi:936,Omega:937,alpha:945,beta:946,gamma:947,delta:948,epsilon:949,zeta:950,eta:951,theta:952,iota:953,kappa:954,lambda:955,mu:956,nu:957,xi:958,omicron:959,pi:960,rho:961,sigmaf:962,sigma:963,tau:964,upsilon:965,phi:966,chi:967,psi:968,omega:969,thetasym:977,upsih:978,piv:982,ensp:8194,emsp:8195,thinsp:8201,zwnj:8204,zwj:8205,lrm:8206,rlm:8207,ndash:8211,mdash:8212,lsquo:8216,rsquo:8217,sbquo:8218,ldquo:8220,rdquo:8221,bdquo:8222,dagger:8224,Dagger:8225,bull:8226,hellip:8230,permil:8240,prime:8242,Prime:8243,lsaquo:8249,rsaquo:8250,oline:8254,frasl:8260,euro:8364,image:8465,weierp:8472,real:8476,trade:8482,alefsym:8501,larr:8592,uarr:8593,rarr:8594,darr:8595,harr:8596,crarr:8629,lArr:8656,uArr:8657,rArr:8658,dArr:8659,hArr:8660,forall:8704,part:8706,exist:8707,empty:8709,nabla:8711,isin:8712,notin:8713,ni:8715,prod:8719,sum:8721,minus:8722,lowast:8727,radic:8730,prop:8733,infin:8734,ang:8736,and:8743,or:8744,cap:8745,cup:8746,int:8747,there4:8756,sim:8764,cong:8773,asymp:8776,ne:8800,equiv:8801,le:8804,ge:8805,sub:8834,sup:8835,nsub:8836,sube:8838,supe:8839,oplus:8853,otimes:8855,perp:8869,sdot:8901,lceil:8968,rceil:8969,lfloor:8970,rfloor:8971,lang:9001,rang:9002,loz:9674,spades:9824,clubs:9827,hearts:9829,diams:9830},Object.keys(e.ENTITIES).forEach(function(t){var n=e.ENTITIES[t],r="number"==typeof n?String.fromCharCode(n):n;e.ENTITIES[t]=r}),e.STATE)e.STATE[e.STATE[D]]=D;function I(t,e,n){t[e]&&t[e](n)}function N(t,e,n){t.textNode&&S(t),I(t,e,n)}function S(t){t.textNode=O(t.opt,t.textNode),t.textNode&&I(t,"ontext",t.textNode),t.textNode=""}function O(t,e){return t.trim&&(e=e.trim()),t.normalize&&(e=e.replace(/\s+/g," ")),e}function C(t,e){return S(t),t.trackPosition&&(e+="\nLine: "+t.line+"\nColumn: "+t.column+"\nChar: "+t.c),e=new Error(e),t.error=e,I(t,"onerror",e),t}function P(t){return t.sawRoot&&!t.closedRoot&&k(t,"Unclosed root tag"),t.state!==A.BEGIN&&t.state!==A.BEGIN_WHITESPACE&&t.state!==A.TEXT&&C(t,"Unexpected end"),S(t),t.c="",t.closed=!0,I(t,"onend"),o.call(t,t.strict,t.opt),t}function k(t,e){if("object"!=typeof t||!(t instanceof o))throw new Error("bad call to strictFail");t.strict&&C(t,e)}function L(t){t.strict||(t.tagName=t.tagName[t.looseCase]());var e=t.tags[t.tags.length-1]||t,n=t.tag={name:t.tagName,attributes:{}};t.opt.xmlns&&(n.ns=e.ns),t.attribList.length=0,N(t,"onopentagstart",n)}function R(t,e){var n=t.indexOf(":")<0?["",t]:t.split(":"),r=n[0],i=n[1];return e&&"xmlns"===t&&(r="xmlns",i=""),{prefix:r,local:i}}function M(t){if(t.strict||(t.attribName=t.attribName[t.looseCase]()),-1!==t.attribList.indexOf(t.attribName)||t.tag.attributes.hasOwnProperty(t.attribName))t.attribName=t.attribValue="";else{if(t.opt.xmlns){var e=R(t.attribName,!0),n=e.prefix,r=e.local;if("xmlns"===n)if("xml"===r&&t.attribValue!==l)k(t,"xml: prefix must be bound to "+l+"\nActual: "+t.attribValue);else if("xmlns"===r&&t.attribValue!==h)k(t,"xmlns: prefix must be bound to "+h+"\nActual: "+t.attribValue);else{var i=t.tag,o=t.tags[t.tags.length-1]||t;i.ns===o.ns&&(i.ns=Object.create(o.ns)),i.ns[r]=t.attribValue}t.attribList.push([t.attribName,t.attribValue])}else t.tag.attributes[t.attribName]=t.attribValue,N(t,"onattribute",{name:t.attribName,value:t.attribValue});t.attribName=t.attribValue=""}}function B(t,e){if(t.opt.xmlns){var n=t.tag,r=R(t.tagName);n.prefix=r.prefix,n.local=r.local,n.uri=n.ns[r.prefix]||"",n.prefix&&!n.uri&&(k(t,"Unbound namespace prefix: "+JSON.stringify(t.tagName)),n.uri=r.prefix);var i=t.tags[t.tags.length-1]||t;n.ns&&i.ns!==n.ns&&Object.keys(n.ns).forEach(function(e){N(t,"onopennamespace",{prefix:e,uri:n.ns[e]})});for(var o=0,s=t.attribList.length;o<s;o++){var a=t.attribList[o],u=a[0],c=a[1],l=R(u,!0),h=l.prefix,f=l.local,p=""===h?"":n.ns[h]||"",d={name:u,value:c,prefix:h,local:f,uri:p};h&&"xmlns"!==h&&!p&&(k(t,"Unbound namespace prefix: "+JSON.stringify(h)),d.uri=h),t.tag.attributes[u]=d,N(t,"onattribute",d)}t.attribList.length=0}t.tag.isSelfClosing=!!e,t.sawRoot=!0,t.tags.push(t.tag),N(t,"onopentag",t.tag),e||(t.noscript||"script"!==t.tagName.toLowerCase()?t.state=A.TEXT:t.state=A.SCRIPT,t.tag=null,t.tagName=""),t.attribName=t.attribValue="",t.attribList.length=0}function U(t){if(!t.tagName)return k(t,"Weird empty close tag."),t.textNode+="</>",void(t.state=A.TEXT);if(t.script){if("script"!==t.tagName)return t.script+="</"+t.tagName+">",t.tagName="",void(t.state=A.SCRIPT);N(t,"onscript",t.script),t.script=""}var e=t.tags.length,n=t.tagName;t.strict||(n=n[t.looseCase]());for(var r=n;e--;){if(t.tags[e].name===r)break;k(t,"Unexpected close tag")}if(e<0)return k(t,"Unmatched closing tag: "+t.tagName),t.textNode+="</"+t.tagName+">",void(t.state=A.TEXT);t.tagName=n;for(var i=t.tags.length;i-- >e;){var o=t.tag=t.tags.pop();t.tagName=t.tag.name,N(t,"onclosetag",t.tagName);var s={};for(var a in o.ns)s[a]=o.ns[a];var u=t.tags[t.tags.length-1]||t;t.opt.xmlns&&o.ns!==u.ns&&Object.keys(o.ns).forEach(function(e){var n=o.ns[e];N(t,"onclosenamespace",{prefix:e,uri:n})})}0===e&&(t.closedRoot=!0),t.tagName=t.attribValue=t.attribName="",t.attribList.length=0,t.state=A.TEXT}function F(t){var e,n=t.entity,r=n.toLowerCase(),i="";return t.ENTITIES[n]?t.ENTITIES[n]:t.ENTITIES[r]?t.ENTITIES[r]:("#"===(n=r).charAt(0)&&("x"===n.charAt(1)?(n=n.slice(2),i=(e=parseInt(n,16)).toString(16)):(n=n.slice(1),i=(e=parseInt(n,10)).toString(10))),n=n.replace(/^0+/,""),isNaN(e)||i.toLowerCase()!==n?(k(t,"Invalid character entity"),"&"+t.entity+";"):String.fromCodePoint(e))}function j(t,e){"<"===e?(t.state=A.OPEN_WAKA,t.startTagPosition=t.position):m(e)||(k(t,"Non-whitespace before first tag."),t.textNode=e,t.state=A.TEXT)}function Y(t,e){var n="";return e<t.length&&(n=t.charAt(e)),n}A=e.STATE,String.fromCodePoint||(E=String.fromCharCode,T=Math.floor,x=function(){var t,e,n=[],r=-1,i=arguments.length;if(!i)return"";for(var o="";++r<i;){var s=Number(arguments[r]);if(!isFinite(s)||s<0||s>1114111||T(s)!==s)throw RangeError("Invalid code point: "+s);s<=65535?n.push(s):(t=55296+((s-=65536)>>10),e=s%1024+56320,n.push(t,e)),(r+1===i||n.length>16384)&&(o+=E.apply(null,n),n.length=0)}return o},Object.defineProperty?Object.defineProperty(String,"fromCodePoint",{value:x,configurable:!0,writable:!0}):String.fromCodePoint=x)}(e)}).call(this,n(25).Buffer)},function(t,e,n){(function(){"use strict";var t,r,i,o,s,a,u,c,l=function(t,e){return function(){return t.apply(e,arguments)}},h={}.hasOwnProperty;u=n(52),i=n(9),t=n(38),a=n(28),c=n(30).setImmediate,r=n(27).defaults,o=function(t){return"object"==typeof t&&null!=t&&0===Object.keys(t).length},s=function(t,e,n){var r,i;for(r=0,i=t.length;r<i;r++)e=(0,t[r])(e,n);return e},e.Parser=function(n){function f(t){var n,i,o;if(this.parseString=l(this.parseString,this),this.reset=l(this.reset,this),this.assignOrPush=l(this.assignOrPush,this),this.processAsync=l(this.processAsync,this),!(this instanceof e.Parser))return new e.Parser(t);for(n in this.options={},i=r[.2])h.call(i,n)&&(o=i[n],this.options[n]=o);for(n in t)h.call(t,n)&&(o=t[n],this.options[n]=o);this.options.xmlns&&(this.options.xmlnskey=this.options.attrkey+"ns"),this.options.normalizeTags&&(this.options.tagNameProcessors||(this.options.tagNameProcessors=[]),this.options.tagNameProcessors.unshift(a.normalize)),this.reset()}return function(t,e){for(var n in e)h.call(e,n)&&(t[n]=e[n]);function r(){this.constructor=t}r.prototype=e.prototype,t.prototype=new r,t.__super__=e.prototype}(f,i.EventEmitter),f.prototype.processAsync=function(){var t,e;try{return this.remaining.length<=this.options.chunkSize?(t=this.remaining,this.remaining="",this.saxParser=this.saxParser.write(t),this.saxParser.close()):(t=this.remaining.substr(0,this.options.chunkSize),this.remaining=this.remaining.substr(this.options.chunkSize,this.remaining.length),this.saxParser=this.saxParser.write(t),c(this.processAsync))}catch(t){if(e=t,!this.saxParser.errThrown)return this.saxParser.errThrown=!0,this.emit(e)}},f.prototype.assignOrPush=function(t,e,n){return e in t?(t[e]instanceof Array||(t[e]=[t[e]]),t[e].push(n)):this.options.explicitArray?t[e]=[n]:t[e]=n},f.prototype.reset=function(){var t,e,n,r,i;return this.removeAllListeners(),this.saxParser=u.parser(this.options.strict,{trim:!1,normalize:!1,xmlns:this.options.xmlns}),this.saxParser.errThrown=!1,this.saxParser.onerror=(i=this,function(t){if(i.saxParser.resume(),!i.saxParser.errThrown)return i.saxParser.errThrown=!0,i.emit("error",t)}),this.saxParser.onend=function(t){return function(){if(!t.saxParser.ended)return t.saxParser.ended=!0,t.emit("end",t.resultObject)}}(this),this.saxParser.ended=!1,this.EXPLICIT_CHARKEY=this.options.explicitCharkey,this.resultObject=null,r=[],t=this.options.attrkey,e=this.options.charkey,this.saxParser.onopentag=function(n){return function(i){var o,a,u,c,l;if((u={})[e]="",!n.options.ignoreAttrs)for(o in l=i.attributes)h.call(l,o)&&(t in u||n.options.mergeAttrs||(u[t]={}),a=n.options.attrValueProcessors?s(n.options.attrValueProcessors,i.attributes[o],o):i.attributes[o],c=n.options.attrNameProcessors?s(n.options.attrNameProcessors,o):o,n.options.mergeAttrs?n.assignOrPush(u,c,a):u[t][c]=a);return u["#name"]=n.options.tagNameProcessors?s(n.options.tagNameProcessors,i.name):i.name,n.options.xmlns&&(u[n.options.xmlnskey]={uri:i.uri,local:i.local}),r.push(u)}}(this),this.saxParser.onclosetag=function(t){return function(){var n,i,a,u,c,l,f,p,d,y;if(l=r.pop(),c=l["#name"],t.options.explicitChildren&&t.options.preserveChildrenOrder||delete l["#name"],!0===l.cdata&&(n=l.cdata,delete l.cdata),d=r[r.length-1],l[e].match(/^\s*$/)&&!n?(i=l[e],delete l[e]):(t.options.trim&&(l[e]=l[e].trim()),t.options.normalize&&(l[e]=l[e].replace(/\s{2,}/g," ").trim()),l[e]=t.options.valueProcessors?s(t.options.valueProcessors,l[e],c):l[e],1===Object.keys(l).length&&e in l&&!t.EXPLICIT_CHARKEY&&(l=l[e])),o(l)&&(l=""!==t.options.emptyTag?t.options.emptyTag:i),null!=t.options.validator&&(y="/"+function(){var t,e,n;for(n=[],t=0,e=r.length;t<e;t++)u=r[t],n.push(u["#name"]);return n}().concat(c).join("/"),function(){var e;try{l=t.options.validator(y,d&&d[c],l)}catch(n){return e=n,t.emit("error",e)}}()),t.options.explicitChildren&&!t.options.mergeAttrs&&"object"==typeof l)if(t.options.preserveChildrenOrder){if(d){for(a in d[t.options.childkey]=d[t.options.childkey]||[],f={},l)h.call(l,a)&&(f[a]=l[a]);d[t.options.childkey].push(f),delete l["#name"],1===Object.keys(l).length&&e in l&&!t.EXPLICIT_CHARKEY&&(l=l[e])}}else u={},t.options.attrkey in l&&(u[t.options.attrkey]=l[t.options.attrkey],delete l[t.options.attrkey]),!t.options.charsAsChildren&&t.options.charkey in l&&(u[t.options.charkey]=l[t.options.charkey],delete l[t.options.charkey]),Object.getOwnPropertyNames(l).length>0&&(u[t.options.childkey]=l),l=u;return r.length>0?t.assignOrPush(d,c,l):(t.options.explicitRoot&&(p=l,(l={})[c]=p),t.resultObject=l,t.saxParser.ended=!0,t.emit("end",t.resultObject))}}(this),n=function(t){return function(n){var i,o;if(o=r[r.length-1])return o[e]+=n,t.options.explicitChildren&&t.options.preserveChildrenOrder&&t.options.charsAsChildren&&(t.options.includeWhiteChars||""!==n.replace(/\\n/g,"").trim())&&(o[t.options.childkey]=o[t.options.childkey]||[],(i={"#name":"__text__"})[e]=n,t.options.normalize&&(i[e]=i[e].replace(/\s{2,}/g," ").trim()),o[t.options.childkey].push(i)),o}}(this),this.saxParser.ontext=n,this.saxParser.oncdata=function(t){var e;if(e=n(t))return e.cdata=!0}},f.prototype.parseString=function(e,n){var r;null!=n&&"function"==typeof n&&(this.on("end",function(t){return this.reset(),n(null,t)}),this.on("error",function(t){return this.reset(),n(t)}));try{return""===(e=e.toString()).trim()?(this.emit("end",null),!0):(e=t.stripBOM(e),this.options.async?(this.remaining=e,c(this.processAsync),this.saxParser):this.saxParser.write(e).close())}catch(t){if(r=t,!this.saxParser.errThrown&&!this.saxParser.ended)return this.emit("error",r),this.saxParser.errThrown=!0;if(this.saxParser.ended)throw r}},f}(),e.parseString=function(t,n,r){var i,o;return null!=r?("function"==typeof r&&(i=r),"object"==typeof n&&(o=n)):("function"==typeof n&&(i=n),o={}),new e.Parser(o).parseString(t,i)}}).call(this)},function(t,e,n){(function(){var e,r,i,o,s,a,u,c,l,h,f,p,d,y={}.hasOwnProperty;u=n(18),c=n(17),e=n(20),r=n(19),l=n(21),f=n(12),p=n(11),h=n(10),i=n(16),o=n(14),s=n(15),a=n(13),d=n(35),t.exports=function(t){function n(t,e){n.__super__.constructor.call(this,e),this.stream=t}return function(t,e){for(var n in e)y.call(e,n)&&(t[n]=e[n]);function r(){this.constructor=t}r.prototype=e.prototype,t.prototype=new r,t.__super__=e.prototype}(n,d),n.prototype.document=function(t){var e,n,i,o,s,a,l,f;for(n=0,o=(a=t.children).length;n<o;n++)(e=a[n]).isLastRootNode=!1;for(t.children[t.children.length-1].isLastRootNode=!0,f=[],i=0,s=(l=t.children).length;i<s;i++)switch(e=l[i],!1){case!(e instanceof u):f.push(this.declaration(e));break;case!(e instanceof c):f.push(this.docType(e));break;case!(e instanceof r):f.push(this.comment(e));break;case!(e instanceof h):f.push(this.processingInstruction(e));break;default:f.push(this.element(e))}return f},n.prototype.attribute=function(t){return this.stream.write(" "+t.name+'="'+t.value+'"')},n.prototype.cdata=function(t,e){return this.stream.write(this.space(e)+"<![CDATA["+t.text+"]]>"+this.endline(t))},n.prototype.comment=function(t,e){return this.stream.write(this.space(e)+"\x3c!-- "+t.text+" --\x3e"+this.endline(t))},n.prototype.declaration=function(t,e){return this.stream.write(this.space(e)),this.stream.write('<?xml version="'+t.version+'"'),null!=t.encoding&&this.stream.write(' encoding="'+t.encoding+'"'),null!=t.standalone&&this.stream.write(' standalone="'+t.standalone+'"'),this.stream.write(this.spacebeforeslash+"?>"),this.stream.write(this.endline(t))},n.prototype.docType=function(t,n){var u,c,l,f;if(n||(n=0),this.stream.write(this.space(n)),this.stream.write("<!DOCTYPE "+t.root().name),t.pubID&&t.sysID?this.stream.write(' PUBLIC "'+t.pubID+'" "'+t.sysID+'"'):t.sysID&&this.stream.write(' SYSTEM "'+t.sysID+'"'),t.children.length>0){for(this.stream.write(" ["),this.stream.write(this.endline(t)),c=0,l=(f=t.children).length;c<l;c++)switch(u=f[c],!1){case!(u instanceof i):this.dtdAttList(u,n+1);break;case!(u instanceof o):this.dtdElement(u,n+1);break;case!(u instanceof s):this.dtdEntity(u,n+1);break;case!(u instanceof a):this.dtdNotation(u,n+1);break;case!(u instanceof e):this.cdata(u,n+1);break;case!(u instanceof r):this.comment(u,n+1);break;case!(u instanceof h):this.processingInstruction(u,n+1);break;default:throw new Error("Unknown DTD node type: "+u.constructor.name)}this.stream.write("]")}return this.stream.write(this.spacebeforeslash+">"),this.stream.write(this.endline(t))},n.prototype.element=function(t,n){var i,o,s,a,u,c,d,g;for(u in n||(n=0),g=this.space(n),this.stream.write(g+"<"+t.name),c=t.attributes)y.call(c,u)&&(i=c[u],this.attribute(i));if(0===t.children.length||t.children.every(function(t){return""===t.value}))this.allowEmpty?this.stream.write("></"+t.name+">"):this.stream.write(this.spacebeforeslash+"/>");else if(this.pretty&&1===t.children.length&&null!=t.children[0].value)this.stream.write(">"),this.stream.write(t.children[0].value),this.stream.write("</"+t.name+">");else{for(this.stream.write(">"+this.newline),s=0,a=(d=t.children).length;s<a;s++)switch(o=d[s],!1){case!(o instanceof e):this.cdata(o,n+1);break;case!(o instanceof r):this.comment(o,n+1);break;case!(o instanceof l):this.element(o,n+1);break;case!(o instanceof f):this.raw(o,n+1);break;case!(o instanceof p):this.text(o,n+1);break;case!(o instanceof h):this.processingInstruction(o,n+1);break;default:throw new Error("Unknown XML node type: "+o.constructor.name)}this.stream.write(g+"</"+t.name+">")}return this.stream.write(this.endline(t))},n.prototype.processingInstruction=function(t,e){return this.stream.write(this.space(e)+"<?"+t.target),t.value&&this.stream.write(" "+t.value),this.stream.write(this.spacebeforeslash+"?>"+this.endline(t))},n.prototype.raw=function(t,e){return this.stream.write(this.space(e)+t.value+this.endline(t))},n.prototype.text=function(t,e){return this.stream.write(this.space(e)+t.value+this.endline(t))},n.prototype.dtdAttList=function(t,e){return this.stream.write(this.space(e)+"<!ATTLIST "+t.elementName+" "+t.attributeName+" "+t.attributeType),"#DEFAULT"!==t.defaultValueType&&this.stream.write(" "+t.defaultValueType),t.defaultValue&&this.stream.write(' "'+t.defaultValue+'"'),this.stream.write(this.spacebeforeslash+">"+this.endline(t))},n.prototype.dtdElement=function(t,e){return this.stream.write(this.space(e)+"<!ELEMENT "+t.name+" "+t.value),this.stream.write(this.spacebeforeslash+">"+this.endline(t))},n.prototype.dtdEntity=function(t,e){return this.stream.write(this.space(e)+"<!ENTITY"),t.pe&&this.stream.write(" %"),this.stream.write(" "+t.name),t.value?this.stream.write(' "'+t.value+'"'):(t.pubID&&t.sysID?this.stream.write(' PUBLIC "'+t.pubID+'" "'+t.sysID+'"'):t.sysID&&this.stream.write(' SYSTEM "'+t.sysID+'"'),t.nData&&this.stream.write(" NDATA "+t.nData)),this.stream.write(this.spacebeforeslash+">"+this.endline(t))},n.prototype.dtdNotation=function(t,e){return this.stream.write(this.space(e)+"<!NOTATION "+t.name),t.pubID&&t.sysID?this.stream.write(' PUBLIC "'+t.pubID+'" "'+t.sysID+'"'):t.pubID?this.stream.write(' PUBLIC "'+t.pubID+'"'):t.sysID&&this.stream.write(' SYSTEM "'+t.sysID+'"'),this.stream.write(this.spacebeforeslash+">"+this.endline(t))},n.prototype.endline=function(t){return t.isLastRootNode?"":this.newline},n}()}).call(this)},function(t,e,n){(function(){var e,r,i,o,s,a,u,c,l,h,f,p,d,y,g,m,_,b,w,v={}.hasOwnProperty;w=n(1),_=w.isObject,m=w.isFunction,b=w.isPlainObject,h=n(21),r=n(20),i=n(19),p=n(12),g=n(11),f=n(10),c=n(18),l=n(17),o=n(16),a=n(15),s=n(14),u=n(13),e=n(37),y=n(36),d=n(26),t.exports=function(){function t(t,e,n){var r;t||(t={}),t.writer?b(t.writer)&&(r=t.writer,t.writer=new d(r)):t.writer=new d(t),this.options=t,this.writer=t.writer,this.stringify=new y(t),this.onDataCallback=e||function(){},this.onEndCallback=n||function(){},this.currentNode=null,this.currentLevel=-1,this.openTags={},this.documentStarted=!1,this.documentCompleted=!1,this.root=null}return t.prototype.node=function(t,e,n){var r;if(null==t)throw new Error("Missing node name");if(this.root&&-1===this.currentLevel)throw new Error("Document can only have one root node");return this.openCurrent(),t=t.valueOf(),null==e&&(e={}),e=e.valueOf(),_(e)||(n=(r=[e,n])[0],e=r[1]),this.currentNode=new h(this,t,e),this.currentNode.children=!1,this.currentLevel++,this.openTags[this.currentLevel]=this.currentNode,null!=n&&this.text(n),this},t.prototype.element=function(t,e,n){return this.currentNode&&this.currentNode instanceof l?this.dtdElement.apply(this,arguments):this.node(t,e,n)},t.prototype.attribute=function(t,n){var r,i;if(!this.currentNode||this.currentNode.children)throw new Error("att() can only be used immediately after an ele() call in callback mode");if(null!=t&&(t=t.valueOf()),_(t))for(r in t)v.call(t,r)&&(i=t[r],this.attribute(r,i));else m(n)&&(n=n.apply()),this.options.skipNullAttributes&&null==n||(this.currentNode.attributes[t]=new e(this,t,n));return this},t.prototype.text=function(t){var e;return this.openCurrent(),e=new g(this,t),this.onData(this.writer.text(e,this.currentLevel+1)),this},t.prototype.cdata=function(t){var e;return this.openCurrent(),e=new r(this,t),this.onData(this.writer.cdata(e,this.currentLevel+1)),this},t.prototype.comment=function(t){var e;return this.openCurrent(),e=new i(this,t),this.onData(this.writer.comment(e,this.currentLevel+1)),this},t.prototype.raw=function(t){var e;return this.openCurrent(),e=new p(this,t),this.onData(this.writer.raw(e,this.currentLevel+1)),this},t.prototype.instruction=function(t,e){var n,r,i,o,s;if(this.openCurrent(),null!=t&&(t=t.valueOf()),null!=e&&(e=e.valueOf()),Array.isArray(t))for(n=0,o=t.length;n<o;n++)r=t[n],this.instruction(r);else if(_(t))for(r in t)v.call(t,r)&&(i=t[r],this.instruction(r,i));else m(e)&&(e=e.apply()),s=new f(this,t,e),this.onData(this.writer.processingInstruction(s,this.currentLevel+1));return this},t.prototype.declaration=function(t,e,n){var r;if(this.openCurrent(),this.documentStarted)throw new Error("declaration() must be the first node");return r=new c(this,t,e,n),this.onData(this.writer.declaration(r,this.currentLevel+1)),this},t.prototype.doctype=function(t,e,n){if(this.openCurrent(),null==t)throw new Error("Missing root node name");if(this.root)throw new Error("dtd() must come before the root node");return this.currentNode=new l(this,e,n),this.currentNode.rootNodeName=t,this.currentNode.children=!1,this.currentLevel++,this.openTags[this.currentLevel]=this.currentNode,this},t.prototype.dtdElement=function(t,e){var n;return this.openCurrent(),n=new s(this,t,e),this.onData(this.writer.dtdElement(n,this.currentLevel+1)),this},t.prototype.attList=function(t,e,n,r,i){var s;return this.openCurrent(),s=new o(this,t,e,n,r,i),this.onData(this.writer.dtdAttList(s,this.currentLevel+1)),this},t.prototype.entity=function(t,e){var n;return this.openCurrent(),n=new a(this,!1,t,e),this.onData(this.writer.dtdEntity(n,this.currentLevel+1)),this},t.prototype.pEntity=function(t,e){var n;return this.openCurrent(),n=new a(this,!0,t,e),this.onData(this.writer.dtdEntity(n,this.currentLevel+1)),this},t.prototype.notation=function(t,e){var n;return this.openCurrent(),n=new u(this,t,e),this.onData(this.writer.dtdNotation(n,this.currentLevel+1)),this},t.prototype.up=function(){if(this.currentLevel<0)throw new Error("The document node has no parent");return this.currentNode?(this.currentNode.children?this.closeNode(this.currentNode):this.openNode(this.currentNode),this.currentNode=null):this.closeNode(this.openTags[this.currentLevel]),delete this.openTags[this.currentLevel],this.currentLevel--,this},t.prototype.end=function(){for(;this.currentLevel>=0;)this.up();return this.onEnd()},t.prototype.openCurrent=function(){if(this.currentNode)return this.currentNode.children=!0,this.openNode(this.currentNode)},t.prototype.openNode=function(t){if(!t.isOpen)return!this.root&&0===this.currentLevel&&t instanceof h&&(this.root=t),this.onData(this.writer.openNode(t,this.currentLevel)),t.isOpen=!0},t.prototype.closeNode=function(t){if(!t.isClosed)return this.onData(this.writer.closeNode(t,this.currentLevel)),t.isClosed=!0},t.prototype.onData=function(t){return this.documentStarted=!0,this.onDataCallback(t)},t.prototype.onEnd=function(){return this.documentCompleted=!0,this.onEndCallback()},t.prototype.ele=function(){return this.element.apply(this,arguments)},t.prototype.nod=function(t,e,n){return this.node(t,e,n)},t.prototype.txt=function(t){return this.text(t)},t.prototype.dat=function(t){return this.cdata(t)},t.prototype.com=function(t){return this.comment(t)},t.prototype.ins=function(t,e){return this.instruction(t,e)},t.prototype.dec=function(t,e,n){return this.declaration(t,e,n)},t.prototype.dtd=function(t,e,n){return this.doctype(t,e,n)},t.prototype.e=function(t,e,n){return this.element(t,e,n)},t.prototype.n=function(t,e,n){return this.node(t,e,n)},t.prototype.t=function(t){return this.text(t)},t.prototype.d=function(t){return this.cdata(t)},t.prototype.c=function(t){return this.comment(t)},t.prototype.r=function(t){return this.raw(t)},t.prototype.i=function(t,e){return this.instruction(t,e)},t.prototype.att=function(){return this.currentNode&&this.currentNode instanceof l?this.attList.apply(this,arguments):this.attribute.apply(this,arguments)},t.prototype.a=function(){return this.currentNode&&this.currentNode instanceof l?this.attList.apply(this,arguments):this.attribute.apply(this,arguments)},t.prototype.ent=function(t,e){return this.entity(t,e)},t.prototype.pent=function(t,e){return this.pEntity(t,e)},t.prototype.not=function(t,e){return this.notation(t,e)},t}()}).call(this)},function(t,e,n){(function(){var e,r,i,o,s={}.hasOwnProperty;o=n(1).isPlainObject,e=n(0),i=n(36),r=n(26),t.exports=function(t){function n(t){n.__super__.constructor.call(this,null),t||(t={}),t.writer||(t.writer=new r),this.options=t,this.stringify=new i(t),this.isDocument=!0}return function(t,e){for(var n in e)s.call(e,n)&&(t[n]=e[n]);function r(){this.constructor=t}r.prototype=e.prototype,t.prototype=new r,t.__super__=e.prototype}(n,e),n.prototype.end=function(t){var e;return t?o(t)&&(e=t,t=this.options.writer.set(e)):t=this.options.writer,t.document(this)},n.prototype.toString=function(t){return this.options.writer.set(t).document(this)},n}()}).call(this)},function(t,e,n){(function(){var e,r,i,o,s,a,u;u=n(1),s=u.assign,a=u.isFunction,e=n(56),r=n(55),o=n(26),i=n(54),t.exports.create=function(t,n,r,i){var o,a;if(null==t)throw new Error("Root element needs a name");return i=s({},n,r,i),a=(o=new e(i)).element(t),i.headless||(o.declaration(i),null==i.pubID&&null==i.sysID||o.doctype(i)),a},t.exports.begin=function(t,n,i){var o;return a(t)&&(n=(o=[t,n])[0],i=o[1],t={}),n?new r(t,n,i):new e(t)},t.exports.stringWriter=function(t){return new o(t)},t.exports.streamWriter=function(t,e){return new i(t,e)}}).call(this)},function(t,e,n){(function(){"use strict";var t,r,i,o,s,a={}.hasOwnProperty;t=n(57),r=n(27).defaults,o=function(t){return"string"==typeof t&&(t.indexOf("&")>=0||t.indexOf(">")>=0||t.indexOf("<")>=0)},s=function(t){return"<![CDATA["+i(t)+"]]>"},i=function(t){return t.replace("]]>","]]]]><![CDATA[>")},e.Builder=function(){function e(t){var e,n,i;for(e in this.options={},n=r[.2])a.call(n,e)&&(i=n[e],this.options[e]=i);for(e in t)a.call(t,e)&&(i=t[e],this.options[e]=i)}return e.prototype.buildObject=function(e){var n,i,u,c,l,h;return n=this.options.attrkey,i=this.options.charkey,1===Object.keys(e).length&&this.options.rootName===r[.2].rootName?e=e[l=Object.keys(e)[0]]:l=this.options.rootName,h=this,u=function(t,e){var r,c,l,f,p,d;if("object"!=typeof e)h.options.cdata&&o(e)?t.raw(s(e)):t.txt(e);else if(Array.isArray(e)){for(f in e)if(a.call(e,f))for(p in c=e[f])l=c[p],t=u(t.ele(p),l).up()}else for(p in e)if(a.call(e,p))if(c=e[p],p===n){if("object"==typeof c)for(r in c)d=c[r],t=t.att(r,d)}else if(p===i)t=h.options.cdata&&o(c)?t.raw(s(c)):t.txt(c);else if(Array.isArray(c))for(f in c)a.call(c,f)&&(t="string"==typeof(l=c[f])?h.options.cdata&&o(l)?t.ele(p).raw(s(l)).up():t.ele(p,l).up():u(t.ele(p),l).up());else"object"==typeof c?t=u(t.ele(p),c).up():"string"==typeof c&&h.options.cdata&&o(c)?t=t.ele(p).raw(s(c)).up():(null==c&&(c=""),t=t.ele(p,c.toString()).up());return t},c=t.create(l,this.options.xmldec,this.options.doctype,{headless:this.options.headless,allowSurrogateChars:this.options.allowSurrogateChars}),u(c,e).end(this.options.renderOpts)},e}()}).call(this)},function(t,e,n){(function(){"use strict";var t,r,i,o,s={}.hasOwnProperty;r=n(27),t=n(58),i=n(53),o=n(28),e.defaults=r.defaults,e.processors=o,e.ValidationError=function(t){function e(t){this.message=t}return function(t,e){for(var n in e)s.call(e,n)&&(t[n]=e[n]);function r(){this.constructor=t}r.prototype=e.prototype,t.prototype=new r,t.__super__=e.prototype}(e,Error),e}(),e.Builder=t.Builder,e.Parser=i.Parser,e.parseString=i.parseString}).call(this)},function(t,e,n){const i=n(59).parseString;(function(){var t="d3.layout.phylotree.event";function e(t,e){return t[e]||!1}function n(t){return!(t.hidden||t.notshown)}function o(t){return t.notshown}function s(t){return!(t.target.hidden||t.target.notshown)}function u(t){return t.tag||!1}function c(t,e,n){var r=t.size();if(t.radial()){var i=t.pad_width(),o="fit-to-size"!=t.options()["top-bottom-spacing"]?t.pad_height():0;r=[r[1]+2*i,r[0]+2*i+o],e&&e.selectAll("."+t.css_classes()["tree-container"]).attr("transform","translate ("+i+","+(i+o)+")")}else r=[r[1]+("fit-to-size"!=t.options()["left-right-spacing"]?t.pad_width():0),r[0]+("fit-to-size"!=t.options()["top-bottom-spacing"]?t.pad_height():0)];return e&&(n&&(e=e.transition(100)),e.attr("height",r[1]).attr("width",r[0])),r}function l(t){return!(t.children&&t.children.length)}function h(t){return t.has_hidden_nodes||!1}function f(t){return t.collapsed||!1}function p(t){return[t.node,t["internal-node"],t["collapsed-node"],t["tagged-node"]].reduce(function(t,e,n,r){return t+"g."+e+(n<r.length-1?",":"")},"")}function d(t){return[t.branch,t["selected-branch"],t["tagged-branch"]].reduce(function(t,e,n,r){return t+"path."+e+(n<r.length-1?",":"")},"")}function y(t){return[t.clade].reduce(function(t,e,n,r){return t+"path."+e+(n<r.length-1?",":"")},"")}function g(t,e){var n=[];function r(){var t=n[n.length-1];"children"in t||(t.children=[]),n.push({name:null}),t.children.push(n[n.length-1]),n[n.length-1].original_child_order=t.children.length}function i(e){return{json:null,error:"Unexpected '"+t[e]+"' in '"+t.substring(e-20,e+1)+"[ERROR HERE]"+t.substring(e+1,e+20)+"'"}}var o=0,s="",a="",u="",c=null,l={"'":1,'"':1},h={name:"root"};n.push(h);for(var f,p=/\s/,d=0;d<t.length;d++)try{var y=t[d];switch(o){case 0:"("==y&&(r(),o=1);break;case 1:case 3:if(":"==y)o=3;else if(","==y||")"==y)try{f=void 0,f=n.pop(),e&&"children"in f?f.bootstrap_values=s:f.name=s,f.attribute=a,f.annotation=u,s="",a="",u="",o=1,","==y&&r()}catch(t){return i(d)}else if("("==y){if(s.length>0)return i(d);r()}else{if(y in l){if(1==o&&0===s.length&&0===a.length&&0===u.length){o=2,c=y;continue}return i(d)}if("["==y){if(u.length)return i(d);o=4}else if(3==o)a+=y;else{if(p.test(y))continue;if(";"==y){d=t.length;break}s+=y}}break;case 2:if(y==c){if(d<t.length-1&&t[d+1]==c){d++,s+=c;continue}c=0,o=1;continue}s+=y;break;case 4:if("]"==y)o=3;else{if("["==y)return i(d);u+=y}}}catch(t){return i(d)}return 1!=n.length?i(t.length-1):(s.length&&(h.name=s),{json:h,error:null})}function m(e){var n=new CustomEvent(t,{detail:["refresh",e]});document.dispatchEvent(n)}function b(e,n){var r=new CustomEvent(t,{detail:["count_update",n,e.count_handler()]});document.dispatchEvent(r)}function w(t){switch(t.detail[0]){case"refresh":t.detail[1].refresh();break;case"count_update":case"layout":t.detail[2](t.detail[1])}return!0}function v(){document.addEventListener(t,w,!1)}function E(t){return!t||null===t[0]&&null===t[1]?"":"translate ("+(null!==t[0]?t[0]:0)+","+(null!==t[1]?t[1]:0)+") "}function T(t){return null!==t?"rotate ("+t+") ":""}d3.layout.newick_parser=function(t,e){return g(t,e)},d3.layout.phylotree=function(w){var x=m,A={},D=d3.layout.hierarchy().sort(null).value(null),I=[1,1],N=[1,1],S=function(t,e){return 0},O=function(t){return 1},C=function(t){if("attribute"in t&&t.attribute&&t.attribute.length){var e=parseFloat(t.attribute);if(!isNaN(e))return Math.max(0,e)}},P=C,L=function(t){return(F["internal-names"]||l(t))&&t.name||""},R=L,M=!0,B=null,U=null,F={layout:"left-to-right",branches:"step",scaling:!0,bootstrap:!1,"color-fill":!0,"internal-names":!1,selectable:!0,"restricted-selectable":!1,collapsible:!0,"left-right-spacing":"fixed-step","top-bottom-spacing":"fixed-step","left-offset":0,"show-scale":"top","draw-size-bubbles":!1,"binary-selectable":!1,"is-radial":!1,"attribute-list":[],"max-radius":768,"annular-limit":.38196601125010515,compression:.2,"align-tips":!1,"maximum-per-node-spacing":100,"minimum-per-node-spacing":2,"maximum-per-level-spacing":100,"minimum-per-level-spacing":10,node_circle_size:d3.functor(3),transitions:null,brush:!0,reroot:!0,hide:!0,"label-nodes-with-name":!1,zoom:!1,"show-menu":!0,"include-bl-title":!0},j={"tree-container":"phylotree-container","tree-scale-bar":"tree-scale-bar",node:"node","internal-node":"internal-node","tagged-node":"node-tagged","selected-node":"node-selected","collapsed-node":"node-collapsed",branch:"branch","selected-branch":"branch-selected","tagged-branch":"branch-tagged","tree-selection-brush":"tree-selection-brush","branch-tracer":"branch-tracer",clade:"clade"},Y=[],z=[],V=[],q=function(t){return t.y},W=function(t){return t.x},G=[1,1],X=[15,20],H=12,K=12,Q=[0,H/2],Z=d3.svg.line().x(function(t){return q(t)}).y(function(t){return W(t)}).interpolate("step-before"),J=function(t){return"number"==typeof t?t+"px":t},tt=function(t){var e=radial_mapper(t[0].radius,t[0].angle),n=radial_mapper(t[0].radius,t[1].angle);return"M "+q(e)+","+W(e)+" A "+t[0].radius+","+t[0].radius+" 0,0, "+(t[1].angle>t[0].angle?1:0)+" "+q(n)+","+W(n)+" L "+q(t[1])+","+W(t[1])};function et(t,e){var n;if(v(),(n="root"==t.name?{json:t,error:null}:"string"!=typeof t?t:"<"==t[0]?function(t){function e(t,n){t.clade&&(t.clade.forEach(e),t.children=t.clade,delete t.clade),t.original_child_order=n+1,t.branch_length&&(t.attribute=t.branch_length[0]),t.taxonomy&&(t.name=t.taxonomy[0].scientific_name[0]),t.annotation=""}var n;return i(t,function(t,r){(n=r.phyloxml.phylogeny[0].clade[0]).name="root",e(n)}),{json:n,error:null}}(t):g(t,e)).json){t;var r={};(Y=D.call(this,n.json)).forEach(t=>{if(t.name){var e=t.name.indexOf("{");if(e>-1){var n=t.name.slice(e+1,t.name.length-1);t[n]=!0,r[n]=!0,t.name=t.name.slice(0,e)}}}),V=Object.keys(r)}else Y=[];return et.placenodes(),z=et.links(Y),et}return draw_branch=Z,draw_scale_bar=null,rescale_node_span=1,count_listener_handler=function(){},layout_listener_handler=function(){},node_styler=void 0,edge_styler=void 0,shown_font_size=H,selection_attribute_name="selected",popover_displayed=null,right_most_leaf=0,label_width=0,radial_center=0,radius=1,radius_pad_for_bubbles=0,radial_mapper=function(t,e){return{x:radial_center+t*Math.sin(e),y:radial_center+t*Math.cos(e)}},cartesian_mapper=function(t,e){return polar_to_cartesian(t-radial_center,e-radial_center)},cartesian_to_polar=function(t,e,n){t.radius=e*(t.radius+n),t.angle||(t.angle=2*Math.PI*t.x*G[0]/I[0]);var r=radial_mapper(t.radius,t.angle);return t.x=r.x,t.y=r.y,t},polar_to_cartesian=function(t,e){return r=Math.sqrt(Math.pow(t,2)+Math.pow(e,2)),a=Math.atan2(e,t),[r,a]},A.container=w||"body",et.placenodes=function(){var t=0,e=[[0,0],[0,0]],n=null,r=0,i=t,s=.5*r,a=F.scaling,u=!1,c=!1,h=1;function p(t){var e=0;if(t.x=t.children.map(d).reduce(function(t,n){return"number"==typeof n?t+n:(e+=1,t)},0),e==t.children.length)return t.notshown=!0,void(t.x=void 0);t.x/=t.children.length-e}function d(d){if(!o(d)){var y=l(d);if(d.text_angle=null,d.text_align=null,d.radius=null,d.angle=null,d.parent)if(a){if(u)return 0;if(d.y=P(d),void 0===d.y)return u=!0,0;d.y+=d.parent.y}else d.y=y?h:d.depth;else t=0,e=[[0,0],[0,0]],n=null,r=0,d.y=0;if(y){var g=O(d)/rescale_node_span;t=d.x=t+S(n,d)+.5*(r+g),e[1][1]=Math.max(e[1][1],d.y),e[1][0]=Math.min(e[1][0],d.y-.5*g),e[0][1]=c?Math.max(e[0][1],i+(d.x-i)*F.compression+s+(.5*g+S(n,d))*F.compression):Math.max(e[0][1],t+.5*g+S(n,d)),n=d,r=g}else{if(f(d)&&!c){if(i=t,s=.5*r,c=!0,p(d),c=!1,"number"==typeof d.x){d.x=i+(d.x-i)*F.compression+s,d.collapsed=[[d.x,d.y]];var m=function(e){e.hidden=!0,l(e)?(t=e.x=i+(e.x-i)*F.compression+s,d.collapsed.push([e.x,e.y])):e.children.map(m)};t=i,m(d),d.collapsed.splice(1,0,[i,d.y]),d.collapsed.push([t,d.y]),d.collapsed.push([d.x,d.y]),d.hidden=!1}}else p(d)}return d.x}}rescale_node_span=Y.map(function(t){return O(t)}).reduce(function(t,e){return Math.min(e,t||1e200)},null)||1,Y[0].x=d(Y[0]),h=d3.max(Y,function(t){return t.depth}),a&&u&&(a=!1,Y[0].x=d(Y[0]));var y=!1;function g(){if(et.radial()&&y&&(Q[1]=0),"fixed-step"==F["left-right-spacing"])I[1]=h*X[1],G[1]=(I[1]-Q[1]-F["left-offset"])/e[1][1],label_width=et._label_width(shown_font_size),et.radial();else{label_width=et._label_width(shown_font_size),y=!0;var t=I[1]-Q[1]-F["left-offset"];.5*t<label_width&&(shown_font_size*=.5*t/label_width,label_width=.5*t),G[1]=(I[1]-Q[1]-F["left-offset"]-label_width)/e[1][1]}}if(draw_scale_bar=F["show-scale"]&&a,"fixed-step"==F["top-bottom-spacing"]?(Q[1]=Math.max(H,-e[1][0]*X[0]),I[0]=e[0][1]*X[0],G[0]=X[0]):(G[0]=(I[0]-et.pad_height())/e[0][1],y=!0),shown_font_size=Math.min(H,G[0]),et.radial()){draw_branch=tt;var m=null,_=null,b=null,w=0,v=e[0][1]*G[0],E=0;Y.forEach(function(t){var e=t.x*G[0];t.angle=2*Math.PI*e/v,t.text_angle=t.angle-Math.PI/2,t.text_angle=t.text_angle>0&&t.text_angle<Math.PI,t.text_align=t.text_angle?"end":"start",t.text_angle=(t.text_angle?180:0)+180*t.angle/Math.PI}),g(),Y.forEach(function(t){t.radius=t.y*G[1]/I[1],E=Math.max(t.radius,E)});var T=0;et.align_tips()&&F["draw-size-bubbles"];Y.forEach(function(t){if(!t.children){var e=t.x*G[0];if(null!==m){var n=e-_,r=function(t,e,n,r,i){return i=i||0,Math.sqrt((e-t)*(e-t)+2*(t+i)*(e+i)*(1-Math.cos(n-r)))}(t.radius,b,t.angle,m,T),i=r>0?n/r:10*F["max-radius"];if(i>F["max-radius"]){var o=n/F["max-radius"],s=t.radius+b,a=t.radius*b-(o*o-(b-t.radius)*(b-t.radius))/2/(1-Math.cos(m-t.angle)),u=Math.sqrt(s*s-4*a);T=Math.min(F["annular-limit"]*E,(-s+u)/2),w=F["max-radius"]}else w=Math.max(w,i)}m=t.angle,_=e,b=t.radius}}),radius=Math.min(F["max-radius"],Math.max(v/2/Math.PI,w)),y&&(radius=Math.min(radius,.5*(Math.min(v,e[1][1]*G[1])-label_width)-radius*T)),radial_center=radius_pad_for_bubbles=radius;var x=1;T&&(x=E/(E+T),radius*=x),Y.forEach(function(t){if(cartesian_to_polar(t,radius,T),E=Math.max(E,t.radius),F["draw-size-bubbles"]?radius_pad_for_bubbles=Math.max(radius_pad_for_bubbles,t.radius+et.node_bubble_size(t)):radius_pad_for_bubbles=Math.max(radius_pad_for_bubbles,t.radius),t.collapsed){t.collapsed=t.collapsed.map(function(t){var e={};return e.x=t[0],e.y=t[1],[(e=cartesian_to_polar(e,radius,T)).x,e.y]});var e=t.collapsed[1];t.collapsed=t.collapsed.filter(function(n,r){return r<3||r>t.collapsed.length-4||Math.sqrt(Math.pow(n[0]-e[0],2)+Math.pow(n[1]-e[1],2))>3&&(e=n,!0)})}}),I[0]=radial_center+radius/x,I[1]=radial_center+radius/x}else g(),draw_branch=Z,right_most_leaf=0,Y.forEach(function(t){if(t.x*=G[0],t.y*=G[1],"right-to-left"==F.layout&&(t.y=e[1][1]*G[1]-t.y),l(t)&&(right_most_leaf=Math.max(right_most_leaf,t.y+et.node_bubble_size(t))),t.collapsed){t.collapsed.map(function(t){return[t[0]*=G[0],t[1]*=G[1]]});var n=t.collapsed[1][0];t.collapsed=t.collapsed.filter(function(e,r){return r<3||r>t.collapsed.length-4||e[0]-n>3&&(n=e[0],!0)})}});if(draw_scale_bar){var A,D;if(et.radial()){if(D=Math.min(radius/5,50),(D=(A=Math.pow(10,Math.ceil(Math.log(e[1][1]*D/radius)/Math.log(10))))*(radius/e[1][1]))<30){var N=Math.ceil(30/D);D*=N,A*=N}}else A=e[1][1],D=I[1]-Q[1]-F["left-offset"];var C=d3.scale.linear().domain([0,A]).range([shown_font_size,shown_font_size+D]),k=d3.format(".2g");if(draw_scale_bar=d3.svg.axis().scale(C).orient("top").tickFormat(function(t){return 0===t?"":k(t)}),et.radial())draw_scale_bar.tickValues([A]);else{var L=C.ticks();L=L.length>1?L[1]:L[0],draw_scale_bar.ticks(Math.min(10,d3.round(D/(shown_font_size*k(L).length*.8),0)))}}else draw_scale_bar=null;return et},et.size=function(t){return arguments.length&&(N=t),"fixed-step"!=F["top-bottom-spacing"]&&(I[0]=N[0]),"fixed-step"!=F["left-right-spacing"]&&(I[1]=N[1]),arguments.length?et:I},et.pad_height=function(){return draw_scale_bar?K+25:0},et.pad_width=function(){return Q[1]+F["left-offset"]+label_width},et.descendants=function(t){var e=[];return function t(n){l(n)?e.push(n):n.children.forEach(t)}(t),e},et.collapse_node=function(t){f(t)||(t.collapsed=!0)},et.separation=function(t){return arguments.length?(S=t,et):S},et.selection_label=function(t){return arguments.length?(selection_attribute_name=t,et.sync_edge_labels(),et):selection_attribute_name},et.handle_node_click=function(t){var e=d3.select(A.container).select("#d3_layout_phylotree_context_menu");if(e.empty()&&(e=d3.select(A.container).append("ul").attr("id","d3_layout_phylotree_context_menu").attr("class","dropdown-menu").attr("role","menu")),e.selectAll("li").remove(),t){if(!_.some([Boolean(t.menu_items),F.hide,F.selectable,F.collapsible])||!F["show-menu"])return;l(t)||(F.collapsible&&(e.append("li").append("a").attr("tabindex","-1").text(f(t)?"Expand Subtree":"Collapse Subtree").on("click",function(n){e.style("display","none"),et.toggle_collapse(t).update()}),F.selectable&&(e.append("li").attr("class","divider"),e.append("li").attr("class","dropdown-header").text("Toggle selection"))),F.selectable&&(e.append("li").append("a").attr("tabindex","-1").text("All descendant branches").on("click",function(n){e.style("display","none"),et.modify_selection(et.select_all_descendants(t,!0,!0))}),e.append("li").append("a").attr("tabindex","-1").text("All terminal branches").on("click",function(n){e.style("display","none"),et.modify_selection(et.select_all_descendants(t,!0,!1))}),e.append("li").append("a").attr("tabindex","-1").text("All internal branches").on("click",function(n){e.style("display","none"),et.modify_selection(et.select_all_descendants(t,!1,!0))}))),t.parent&&(F.selectable&&(e.append("li").append("a").attr("tabindex","-1").text("Incident branch").on("click",function(n){e.style("display","none"),et.modify_selection([t])}),e.append("li").append("a").attr("tabindex","-1").text("Path to root").on("click",function(n){e.style("display","none"),et.modify_selection(et.path_to_root(t))}),(F.reroot||F.hide)&&e.append("li").attr("class","divider")),F.reroot&&e.append("li").append("a").attr("tabindex","-1").text("Reroot on this node").on("click",function(n){e.style("display","none"),et.reroot(t).update()}),F.hide&&e.append("li").append("a").attr("tabindex","-1").text("Hide this "+(l(t)?"node":"subtree")).on("click",function(n){e.style("display","none"),et.modify_selection([t],"notshown",!0,!0).update_has_hidden_nodes().update()})),h(t)&&e.append("li").append("a").attr("tabindex","-1").text("Show all descendant nodes").on("click",function(n){e.style("display","none"),et.modify_selection(et.select_all_descendants(t,!0,!0),"notshown",!0,!0,"false").update_has_hidden_nodes().update()});var n=[];if("menu_items"in t&&"object"==typeof t.menu_items&&t.menu_items.forEach(function(e){3==e.length&&(e[2]&&!e[2](t)||n.push([e[0],e[1]]))}),n.length){const r=[F.hide,F.selectable,F.collapsible];_.some(r)&&e.append("li").attr("class","divider"),n.forEach(function(n){e.append("li").append("a").attr("tabindex","-1").text(n[0](t)).on("click",n[1])})}var r=$(A.container),i=d3.mouse(r[0]);e.style("position","absolute").style("left",i[0]+"px").style("top",i[1]+"px").style("display","block")}else e.style("display","none")},et.style_nodes=function(t){return arguments.length?(node_styler=t,et):node_styler},et.style_edges=function(t){return arguments.length?(edge_styler=t.bind(this),et):edge_styler},et.get_newick=function(t){t||(t=(t=>t.name));var e=[];return t=t||"",function n(r){var i;l(r)||(e.push("("),r.children.forEach(function(t,r){r&&e.push(","),n(t)}),e.push(")")),e.push((i=R(r),/[\s\[\]\,\)\(\:\'\"]/.test(i)?"'"+i.replace("'","''")+"'":i)),e.push(t(r));var o=P(r);void 0!==o&&e.push(":"+o)}(Y[0]),e.join("")},et.update_layout=function(e,n){var r,i;n&&(Y=D.call(this,e)).forEach(function(t){t.id=null}),et.placenodes(),z=et.links(Y),et.sync_edge_labels(),r=et,i=new CustomEvent(t,{detail:["layout",r,r.layout_handler()]}),document.dispatchEvent(i)},et.sync_edge_labels=function(){if(z.forEach(function(t){t[selection_attribute_name]=t.target[selection_attribute_name]||!1,t.tag=t.target.tag||!1}),m(et),et.count_handler()){var t={};t[selection_attribute_name]=z.reduce(function(t,e){return t+(e[selection_attribute_name]?1:0)},0),t.tagged=z.reduce(function(t,e){return t+(u(e)?1:0)},0),b(et,t,et.count_handler())}},et.predefined_selecters={all:t=>!0,none:t=>!1,"all-leaf-nodes":t=>l(t.target),"all-internal-nodes":t=>!l(t.target)},et.modify_selection=function(t,e,n,r,i){if(e=e||selection_attribute_name,i=i||"toggle",F["restricted-selectable"].length){if(!_.contains(_.keys(this.predefined_selecters),t))return;t=this.predefined_selecters[t]}if(!F["restricted-selectable"]&&!F.selectable||F["binary-selectable"]){if(F["binary-selectable"]&&("function"==typeof t?z.forEach(function(n){var r=t(n);n[e]=n[e]||!1,n[e]!=r&&(n[e]=r,o=!0,n.target[e]=r),F["attribute-list"].forEach(function(t){t!=e&&!0===n[e]&&(n[t]=!1,n.target[t]=!1)})}):(t.forEach(function(t){var n;n=!t[e],t[e]!=n&&(t[e]=n,o=!0)}),z.forEach(function(t){t[e]=t.target[e],F["attribute-list"].forEach(function(n){n!=e&&!0!==t[e]&&(t[n]=!1,t.target[n]=!1)})})),o)){if(r||m(et),et.count_handler())(s={})[e]=z.reduce(function(t,n){return t+(n[e]?1:0)},0),b(et,s,et.count_handler());n&&et.placenodes()}}else{var o=!1;if("function"==typeof t?z.forEach(function(n){var r=t(n);n[e]=n[e]||!1,n[e]!=r&&(n[e]=r,o=!0,n.target[e]=r)}):(t.forEach(function(t){var n;switch(i){case"true":n=!0;break;case"false":n=!1;break;default:n=!t[e]}t[e]!=n&&(t[e]=n,o=!0)}),z.forEach(function(t){t[e]=t.target[e]})),o){var s;if(r||m(et),et.count_handler())(s={})[e]=z.reduce(function(t,n){return t+(n[e]?1:0)},0),b(et,s,et.count_handler());n&&et.placenodes()}}return U&&"tag"!=e&&U(et.get_selection()),et},et.trigger_refresh=function(){x(et)},et.is_leafnode=l,et.radial=function(t){return arguments.length?(F["is-radial"]=t,et):F["is-radial"]},et.align_tips=function(t){return arguments.length?(F["align-tips"]=t,et):F["align-tips"]},et.node_bubble_size=function(t){return F["draw-size-bubbles"]?.5*(O(t)/rescale_node_span*G[0]):0},et.shift_tip=function(t){return F["is-radial"]?[("end"==t.text_align?-1:1)*(radius_pad_for_bubbles-t.radius),0]:(F["right-to-left"],[right_most_leaf-t.screen_x,0])},et.get_selection=function(){return Y.filter(function(t){return t[selection_attribute_name]})},et.count_handler=function(t){return arguments.length?(count_listener_handler=t,et):count_listener_handler},et.layout_handler=function(t){return arguments.length?(layout_listener_handler=t,et):layout_listener_handler},et.internal_label=function(t,n){et.clear_internal_nodes(n);for(var r=Y.length-1;r>=0;r--){var i=Y[r];l(i)||e(i,selection_attribute_name)||(i[selection_attribute_name]=t(i.children))}et.modify_selection(function(t,e){return l(t.target),t.target[selection_attribute_name]})},et.max_parsimony=function(t){et.clear_internal_nodes(t),function t(e){if(e.mp=[[0,0],[!1,!1]],l(e))e.mp[1][0]=e.mp[1][1]=e[selection_attribute_name]||!1,e.mp[0][0]=e.mp[1][0]?1:0,e.mp[0][1]=1-e.mp[0][0];else{e.children.forEach(t);var n=e.children.reduce(function(t,e){return e.mp[0][0]+t},0),r=e.children.reduce(function(t,e){return e.mp[0][1]+t},0);e[selection_attribute_name]?(e.mp[0][0]=r+1,e.mp[1][0]=!0,e.mp[0][1]=r,e.mp[1][1]=!0):(n<r+1?(e.mp[0][0]=n,e.mp[1][0]=!1):(e.mp[0][0]=r+1,e.mp[1][0]=!0),r<n+1?(e.mp[0][1]=r,e.mp[1][1]=!0):(e.mp[0][1]=n+1,e.mp[1][1]=!1))}}(Y[0]),Y.forEach(function(t){t.parent?t.mp=t.mp[1][t.parent.mp?1:0]:t.mp=t.mp[1][t.mp[0][0]<t.mp[0][1]?0:1]}),et.modify_selection(function(t,e){return l(t.target)?t.target[selection_attribute_name]:t.target.mp})},et.node_span=function(t){return arguments.length?(O="string"==typeof t&&"equal"==t?function(t){return 1}:t,et):O},et.resort_children=function(t){return function e(n){if(n.children){for(var r=0;r<n.children.length;r++)e(n.children[r]);n.children.sort(t)}}(Y[0]),et.update_layout(Y),et.update(),et},et.graft_a_node=function(t,e,n,r){if(t.parent&&Y.indexOf(t)>=0){var i=t.parent.children.indexOf(t),o={name:n,parent:t.parent,attribute:r?r[2]:null,original_child_order:t.original_child_order},s={name:e,parent:o,attribute:r?r[1]:null,original_child_order:2};o.children=[t,s],t.parent.children[i]=o,t.parent=o,t.attribute=r?r[0]:null,t.original_child_order=1,et.update_layout(Y[0],!0)}return et},et.delete_a_node=function(t){if("number"!=typeof t)return et.delete_a_node(Y.indexOf(t));if(t>0&&t<Y.length){var e=Y[t];if(e.parent){var n=e.parent.children.indexOf(e);n>=0&&(Y.splice(t,1),e.children&&e.children.forEach(function(t){t.original_child_order=e.parent.children.length,e.parent.children.push(t),t.parent=e.parent}),e.parent.children.length>2?e.parent.children.splice(n,1):e.parent.parent?(e.parent.parent.children[e.parent.parent.children.indexOf(e.parent)]=e.parent.children[1-n],e.parent.children[1-n].parent=e.parent.parent,Y.splice(Y.indexOf(e.parent),1)):(Y.splice(0,1),Y[0].parent=null,delete Y[0].attribute,delete Y[0].annotation,delete Y[0].original_child_order,Y[0].name="root"),et.update_layout(Y[0],!0))}}return et},et.traverse_and_compute=function(t,e){return"post-order"==(e=e||"post-order")&&(e=function e(n){if(n.children)for(var r=0;r<n.children.length;r++)e(n.children[r]);t(n)}),e(Y[0]),et},et.reroot=function(t){if(t.parent){new_json={name:"new_root",__mapped_bl:void 0,children:[t]},Y.forEach(function(t){t.__mapped_bl=P(t)}),et.branch_length(function(t){return t.__mapped_bl});var e=t,n=t.parent,r=(n.__mapped_bl,_.noop());if(n.parent){for(t.__mapped_bl=void 0===t.__mapped_bl?void 0:.5*t.__mapped_bl,r=n.__mapped_bl,n.__mapped_bl=t.__mapped_bl,new_json.children.push(n);n.parent;){var i=n.children.indexOf(e);n.parent.parent?n.children.splice(i,1,n.parent):n.children.splice(i,1);var o=n.parent.__mapped_bl;void 0!==o&&(n.parent.__mapped_bl=r,r=o),e=n,n=n.parent}i=n.children.indexOf(e);n.children.splice(i,1)}else{i=n.children.indexOf(e);n.children.splice(i,1),e=new_json}if(1==n.children.length)r&&(n.children[0].__mapped_bl+=r),e.children=e.children.concat(n.children);else{var s={name:"__reroot_top_clade"};s.__mapped_bl=r,s.children=n.children.map(function(t){return t}),e.children.push(s)}et.update_layout(new_json,!0)}return et},et.update_key_name=function(t,e){return Y.forEach(function(n){t in n&&(e&&(n[e]=n[t]),delete n[t])}),et.sync_edge_labels(),et},et.spacing_x=function(t,e){return arguments.length?(X[0]!=t&&t>=F["minimum-per-node-spacing"]&&t<=F["maximum-per-node-spacing"]&&(X[0]=t,e||et.placenodes()),et):X[0]},et.spacing_y=function(t,e){return arguments.length?(X[1]!=t&&t>=F["minimum-per-level-spacing"]&&t<=F["maximum-per-level-spacing"]&&(X[1]=t,e||et.placenodes()),et):X[1]},et.toggle_collapse=function(t){if(t.collapsed){t.collapsed=!1;var e=function(t){l(t)||t.collapsed||t.children.forEach(e),t.hidden=!1};e(t)}else t.collapsed=!0;return et.placenodes(),et},et.update_has_hidden_nodes=function(){for(k=Y.length-1;k>=0;k-=1)l(Y[k])?Y[k].has_hidden_nodes=Y[k].notshown:Y[k].has_hidden_nodes=Y[k].children.reduce(function(t,e){return e.notshown||t},!1);return et},et.branch_length=function(t){return arguments.length?(P=t||C,et):P},et.branch_name=function(t){return arguments.length?(R=t||L,et):R},et.length=function(t){return arguments.length?(default_length_attribute!=t&&(default_length_attribute=t,M=!0),et):default_length_attribute},et._label_width=function(t){t=t||shown_font_size;var e=0;return Y.filter(n).forEach(function(n){var r=R(n).length*t*.6;null!==n.angle&&(r*=Math.max(Math.abs(Math.cos(n.angle)),Math.abs(Math.sin(n.angle)))),e=Math.max(r,e)}),e},et.font_size=function(t){return arguments.length?(H=void 0===t?12:t,et):H},et.scale_bar_font_size=function(t){return arguments.length?(K=void 0===t?12:t,et):K},et.node_circle_size=function(t,e){return arguments.length?(F.node_circle_size=d3.functor(void 0===t?3:t),et):F.node_circle_size},et.needs_redraw=function(){return M},et.svg=function(t){return arguments.length?(B!==t&&(B=t,"phylotree-container"==j["tree-container"]&&(B.selectAll("*").remove(),svg_defs=B.append("defs")),d3.select(A.container).on("click",function(t){et.handle_node_click(null)},!0)),et):t},et.css=function(t){if(0===arguments.length)return j;if(arguments.length>2){var e={};return e[t[0]]=t[1],et.css(e)}for(var n in j)n in t&&t[n]!=j[n]&&(j[n]=t[n]);return et},et.options=function(t,e){if(!arguments.length)return F;var n=!1;for(var r in F)if(r in t&&t[r]!=F[r])switch(n=!0,F[r]=t[r],r){case"branches":switch(t[r]){case"straight":draw_branch.interpolate("linear");break;default:draw_branch.interpolate("step-before")}}return e&&n&&et.layout(),et},et.transitions=function(t){return void 0!==t?t:null!==F.transitions?F.transitions:Y.length<=300},et.update=function(t){if(!et.svg)return et;t=et.transitions(t);var e=0,r=B.selectAll("."+j["tree-container"]).data([0]);if(r.enter().append("g").attr("class",j["tree-container"]),r.attr("transform",function(t){return E([Q[1]+F["left-offset"],et.pad_height()])}),draw_scale_bar){var i=B.selectAll("."+j["tree-scale-bar"]).data([0]);i.enter().append("g"),i.attr("class",j["tree-scale-bar"]).style("font-size",J(K)).attr("transform",function(t){return E([Q[1]+F["left-offset"],et.pad_height()-10])}).call(draw_scale_bar),i.selectAll("text").style("text-anchor","end")}else B.selectAll("."+j["tree-scale-bar"]).remove();var o=r.selectAll(d(j)).data(z.filter(s),function(t){return t.target.id||(t.target.id=++e)});t?o.exit().transition().remove():o.exit().remove(),o.enter().insert("path",":first-child"),o.each(function(e){et.draw_edge(this,e,t)});var a=r.selectAll(y(j)).data(Y.filter(f),function(t){return t.id||(t.id=++e)}),u=function(){},h=_.noop();if(et.radial()){u=d3.svg.line().interpolate(function(t){t.pop();var e=t.shift(),n=t.join("L"),r=cartesian_mapper(e[0],e[1]),i=cartesian_mapper(t[0][0],t[0][1])[1],o=cartesian_mapper(t[t.length-1][0],t[t.length-1][1])[1];return n+"A "+r[0]+" "+r[0]+" "+(i>o?1:0)+" 0 0 "+t[0].join(",")}).y(function(t){return t[0]}).x(function(t){return t[1]}),h=function(t,e,n,r,i){return e?[n.screen_y+(t[0]-r)/50,n.screen_x+(t[1]-i)/50]:[n.screen_y,n.screen_x]}}else u=d3.svg.line().interpolate("basis").y(function(t){return t[0]}).x(function(t){return t[1]}),h=function(t,e,n,r,i){return e?[n.screen_y+(t[0]-r)/50,n.screen_x+(t[1]-i)/50]:[n.screen_y,n.screen_x]};a.exit().each(function(t){t.collapsed_clade=null}).remove();t?(a.enter().insert("path",":first-child"),a.attr("class",j.clade).attr("d",function(t){if(t.collapsed_clade)return t.collapsed_clade;var e=t.collapsed[0][0],n=t.collapsed[0][1];return u(t.collapsed.map(function(r,i){return h(r,i,t,e,n)}))}).transition().attr("d",function(t){return t.collapsed_clade=u(t.collapsed)})):(a.enter().insert("path",":first-child"),a.attr("class",j.clade).attr("d",function(t){return t.collapsed_clade=u(t.collapsed)}));var g=r.selectAll(p(j)).data(Y.filter(n),function(t){return t.id||(t.id=++e)});g.enter().append("g");t?(g.exit().transition().remove(),g=g.attr("transform",function(t){if(!_.isUndefined(t.screen_x)&&!_.isUndefined(t.screen_y))return"translate("+t.screen_x+","+t.screen_y+")"}).transition()):g.exit().remove(),g.attr("transform",function(t){const e="right-to-left"==F.layout&&l(t);return t.screen_x=q(t),t.screen_y=W(t),E([e?0:t.screen_x,t.screen_y])}).attr("class",et.reclass_node).each(function(e){et.draw_node(this,e,t)}),F["label-nodes-with-name"]&&g.attr("id",function(t){return"node-"+t.name});var m=c(et,B,t);if(F.brush){var b=r.selectAll("."+j["tree-selection-brush"]).data([0]);b.enter().insert("g",":first-child").attr("class",j["tree-selection-brush"]);var w=d3.svg.brush().x(d3.scale.identity().domain([0,m[0]-Q[1]-F["left-offset"]])).y(d3.scale.identity().domain([0,m[1]-et.pad_height()])).on("brush",function(){var t=d3.event.target.extent(),e=z.filter(s).filter(function(e,n){return e.source.screen_x>=t[0][0]&&e.source.screen_x<=t[1][0]&&e.source.screen_y>=t[0][1]&&e.source.screen_y<=t[1][1]&&e.target.screen_x>=t[0][0]&&e.target.screen_x<=t[1][0]&&e.target.screen_y>=t[0][1]&&e.target.screen_y<=t[1][1]}).map(function(t){return t.target});et.modify_selection(z.map(function(t){return t.target}),"tag",!1,e.length>0,"false"),et.modify_selection(e,"tag",!1,!1,"true")}).on("brushend",function(){b.call(d3.event.target.clear())});b.call(w)}if(et.sync_edge_labels(),F.zoom){var v=d3.behavior.zoom().scaleExtent([.1,10]).on("zoom",function(){var t=d3.event.translate;t[0]+=Q[1]+F["left-offset"],t[1]+=et.pad_height(),d3.select("."+j["tree-container"]).attr("transform","translate("+t+")scale("+d3.event.scale+")")});B.call(v)}return et},et.css_classes=function(t,e){if(!arguments.length)return j;var n=!1;for(var r in j)r in t&&t[r]!=j[r]&&(n=!0,j[r]=t[r]);return e&&n&&et.layout(),et},et.layout=function(t){return B?(B.selectAll("."+j["tree-container"]+",."+j["tree-scale-bar"]+",."+j["tree-selection-brush"]),et.update(t)):et},et.refresh=function(){var t=B.selectAll("."+j["tree-container"]),e=t.selectAll(d(j));e.attr("class",et.reclass_edge),edge_styler&&e.each(function(t){edge_styler(d3.select(this),t)});var n=t.selectAll(p(j));n.attr("class",et.reclass_node),node_styler&&n.each(function(t){node_styler(d3.select(this),t)})},et.reclass_edge=function(t){var n=j.branch;return u(t)&&(n+=" "+j["tagged-branch"]),e(t,selection_attribute_name)&&(n+=" "+j["selected-branch"]),n},et.reclass_node=function(t){var n=j[l(t)?"node":"internal-node"];return u(t)&&(n+=" "+j["tagged-node"]),e(t,selection_attribute_name)&&(n+=" "+j["selected-node"]),(f(t)||h(t))&&(n+=" "+j["collapsed-node"]),n},et.select_all_descendants=function(t,e,n){var r=[];return function i(o){l(o)?e&&o!=t&&r.push(o):(n&&o!=t&&r.push(o),o.children.forEach(i))}(t),r},et.path_to_root=function(t){for(var e=[];t;)e.push(t),t=t.parent;return e},et.draw_edge=function(t,e,n){(t=d3.select(t)).attr("class",et.reclass_edge).on("click",function(t){et.modify_selection([t.target],selection_attribute_name)});var r=draw_branch([e.source,e.target]);n?(t.datum().existing_path&&t.attr("d",function(t){return t.existing_path}),t.transition().attr("d",r)):t.attr("d",r),e.existing_path=r;var i=P(e.target);if(void 0!==i){if(F["include-bl-title"]){var o=t.selectAll("title");o.empty()&&(o=t.append("title")),o.text("Length = "+i)}}else t.selectAll("title").remove();return edge_styler&&edge_styler(t,e),et},et.clear_internal_nodes=function(t){t||Y.forEach(function(t){l(t)||(t[selection_attribute_name]=!1)})},et.draw_node=function(t,e,n){if(t=d3.select(t),l(e)){var r=t.selectAll("text").data([e]),i=t.selectAll("line");if(n?r.enter().append("text").style("opacity",0).transition().style("opacity",1):r.enter().append("text"),r.on("click",et.handle_node_click).attr("dy",function(t){return.33*shown_font_size}).text(function(t){return R(t)}).style("font-size",function(t){return J(shown_font_size)}),et.radial()?(n?r.transition():r).attr("transform",function(t){return T(t.text_angle)+E(et.align_tips()?et.shift_tip(t):null)}).attr("text-anchor",function(t){return t.text_align}):(n?r.transition():r).attr("text-anchor","start").attr("transform",function(t){return"right-to-left"==F.layout?E([-20,0]):E(et.align_tips()?et.shift_tip(t):null)}),et.align_tips()?(i=i.data([e]),n?(i.enter().append("line").style("opacity",0).transition().style("opacity",1),i.attr("x1",function(t){return("end"==t.text_align?-1:1)*et.node_bubble_size(e)}).attr("x2",0).attr("y1",0).attr("y2",0),i.transition().attr("x2",function(t){return"right-to-left"==F.layout?t.screen_x:et.shift_tip(t)[0]}).attr("transform",function(t){return T(t.text_angle)})):(i.enter().append("line"),i.attr("x1",function(t){return("end"==t.text_align?-1:1)*et.node_bubble_size(e)}).attr("y2",0).attr("y1",0).transition().attr("x2",function(t){return et.shift_tip(t)[0]}),i.attr("transform",function(t){return T(t.text_angle)})),i.classed(j["branch-tracer"],!0)):i.remove(),F["draw-size-bubbles"]){var o=et.node_bubble_size(e);(s=t.selectAll("circle").data([o])).enter().append("circle"),n&&(s=s.transition()),s.attr("r",function(t){return t}),shown_font_size>=5&&r.attr("dx",function(t){return("end"==t.text_align?-1:1)*((et.align_tips()?0:o)+.33*shown_font_size)})}else shown_font_size>=5&&r.attr("dx",function(t){return("end"==t.text_align?-1:1)*shown_font_size*.33})}else{var s=t.selectAll("circle").data([e]),a=et.node_circle_size()(e);a>0?(s.enter().append("circle"),s.attr("r",function(t){return Math.min(.75*shown_font_size,a)}).on("click",function(t){et.handle_node_click(t)})):s.remove()}return node_styler&&node_styler(t,e),e},et.get_nodes=function(){return Y},et.get_node_by_name=function(t){return _.findWhere(Y,{name:t})},et.assign_attributes=function(t){_.each(Y,function(e){_.indexOf(_.keys(t),e.name)>=0&&(e.annotations=t[e.name])})},et.set_partitions=function(t){this.partitions=t},et.get_partitions=function(t){return this.partitions},et.selection_callback=function(t){return t?(U=t,et):U},et.get_parsed_tags=function(){return V},d3.rebind(et,D,"sort","children","value"),et.nodes=et,et.links=d3.layout.cluster().links,et},d3.layout.phylotree.is_leafnode=l,d3.layout.phylotree.add_custom_menu=function(t,e,n,r){"menu_items"in t||(t.menu_items=[]),t.menu_items.some(function(t){return t[0]==e&&t[1]==n&&t[2]==r})||t.menu_items.push([e,n,r])},d3.layout.phylotree.trigger_refresh=m,d3.layout.phylotree.nexml_parser=function(t){var e;return i(t,function(t,n){e=n["nex:nexml"].trees[0].tree.map(function(t){var e=t.node.map(t=>t.$),n=e.reduce(function(t,e){return e.edges=[],e.name=e.id,t[e.id]=e,t},{}),r=e.filter(t=>t.root),i=r>0?r[0].id:e[0].id;return n[i].name="root",t.edge.map(t=>t.$).forEach(function(t){n[t.source].edges.push(t)}),function t(e,r){if(e.edges){var i=_.pluck(e.edges,"target");e.children=_.values(_.pick(n,i)),e.children.forEach(function(t,n){t.attribute=e.edges[n].length||""}),e.children.forEach(t),e.annotation=""}}(n[i]),n[i]})}),e}}).call(this)}]);
```
