## Supplementary material for "Method for inferring migration routes from phylogenetic trees": Some route-trees: underscore-min.js

```
//     Underscore.js 1.8.3
//     http://underscorejs.org
//     (c) 2009-2015 Jeremy Ashkenas, DocumentCloud and Investigative Reporters & Editors
//     Underscore may be freely distributed under the MIT license.
(function(){function n(n){function t(t,r,e,u,i,o){for(;i>=0&&o>i;i+=n){var a=u?u[i]:i;e=r(e,t[a],a,t)}return e}return function(r,e,u,i){e=b(e,i,4);var o=!k(r)&&m.keys(r),a=(o||r).length,c=n>0?0:a-1;return arguments.length<3&&(u=r[o?o[c]:c],c+=n),t(r,e,u,o,c,a)}}function t(n){return function(t,r,e){r=x(r,e);for(var u=O(t),i=n>0?0:u-1;i>=0&&u>i;i+=n)if(r(t[i],i,t))return i;return-1}}function r(n,t,r){return function(e,u,i){var o=0,a=O(e);if("number"==typeof i)n>0?o=i>=0?i:Math.max(i+a,o):a=i>=0?Math.min(i+1,a):i+a+1;else if(r&&i&&a)return i=r(e,u),e[i]===u?i:-1;if(u!==u)return i=t(l.call(e,o,a),m.isNaN),i>=0?i+o:-1;for(i=n>0?o:a-1;i>=0&&a>i;i+=n)if(e[i]===u)return i;return-1}}function e(n,t){var r=I.length,e=n.constructor,u=m.isFunction(e)&&e.prototype||a,i="constructor";for(m.has(n,i)&&!m.contains(t,i)&&t.push(i);r--;)i=I[r],i in n&&n[i]!==u[i]&&!m.contains(t,i)&&t.push(i)}var u=this,i=u._,o=Array.prototype,a=Object.prototype,c=Function.prototype,f=o.push,l=o.slice,s=a.toString,p=a.hasOwnProperty,h=Array.isArray,v=Object.keys,g=c.bind,y=Object.create,d=function(){},m=function(n){return n instanceof m?n:this instanceof m?void(this._wrapped=n):new m(n)};"undefined"!=typeof exports?("undefined"!=typeof module&&module.exports&&(exports=module.exports=m),exports._=m):u._=m,m.VERSION="1.8.3";var b=function(n,t,r){if(t===void 0)return n;switch(null==r?3:r){case 1:return function(r){return n.call(t,r)};case 2:return function(r,e){return n.call(t,r,e)};case 3:return function(r,e,u){return n.call(t,r,e,u)};case 4:return function(r,e,u,i){return n.call(t,r,e,u,i)}}return function(){return n.apply(t,arguments)}},x=function(n,t,r){return null==n?m.identity:m.isFunction(n)?b(n,t,r):m.isObject(n)?m.matcher(n):m.property(n)};m.iteratee=function(n,t){return x(n,t,1/0)};var _=function(n,t){return function(r){var e=arguments.length;if(2>e||null==r)return r;for(var u=1;e>u;u++)for(var i=arguments[u],o=n(i),a=o.length,c=0;a>c;c++){var f=o[c];t&&r[f]!==void 0||(r[f]=i[f])}return r}},j=function(n){if(!m.isObject(n))return{};if(y)return y(n);d.prototype=n;var t=new d;return d.prototype=null,t},w=function(n){return function(t){return null==t?void 0:t[n]}},A=Math.pow(2,53)-1,O=w("length"),k=function(n){var t=O(n);return"number"==typeof t&&t>=0&&A>=t};m.each=m.forEach=function(n,t,r){t=b(t,r);var e,u;if(k(n))for(e=0,u=n.length;u>e;e++)t(n[e],e,n);else{var i=m.keys(n);for(e=0,u=i.length;u>e;e++)t(n[i[e]],i[e],n)}return n},m.map=m.collect=function(n,t,r){t=x(t,r);for(var e=!k(n)&&m.keys(n),u=(e||n).length,i=Array(u),o=0;u>o;o++){var a=e?e[o]:o;i[o]=t(n[a],a,n)}return i},m.reduce=m.foldl=m.inject=n(1),m.reduceRight=m.foldr=n(-1),m.find=m.detect=function(n,t,r){var e;return e=k(n)?m.findIndex(n,t,r):m.findKey(n,t,r),e!==void 0&&e!==-1?n[e]:void 0},m.filter=m.select=function(n,t,r){var e=[];return t=x(t,r),m.each(n,function(n,r,u){t(n,r,u)&&e.push(n)}),e},m.reject=function(n,t,r){return m.filter(n,m.negate(x(t)),r)},m.every=m.all=function(n,t,r){t=x(t,r);for(var e=!k(n)&&m.keys(n),u=(e||n).length,i=0;u>i;i++){var o=e?e[i]:i;if(!t(n[o],o,n))return!1}return!0},m.some=m.any=function(n,t,r){t=x(t,r);for(var e=!k(n)&&m.keys(n),u=(e||n).length,i=0;u>i;i++){var o=e?e[i]:i;if(t(n[o],o,n))return!0}return!1},m.contains=m.includes=m.include=function(n,t,r,e){return k(n)||(n=m.values(n)),("number"!=typeof r||e)&&(r=0),m.indexOf(n,t,r)>=0},m.invoke=function(n,t){var r=l.call(arguments,2),e=m.isFunction(t);return m.map(n,function(n){var u=e?t:n[t];return null==u?u:u.apply(n,r)})},m.pluck=function(n,t){return m.map(n,m.property(t))},m.where=function(n,t){return m.filter(n,m.matcher(t))},m.findWhere=function(n,t){return m.find(n,m.matcher(t))},m.max=function(n,t,r){var e,u,i=-1/0,o=-1/0;if(null==t&&null!=n){n=k(n)?n:m.values(n);for(var a=0,c=n.length;c>a;a++)e=n[a],e>i&&(i=e)}else t=x(t,r),m.each(n,function(n,r,e){u=t(n,r,e),(u>o||u===-1/0&&i===-1/0)&&(i=n,o=u)});return i},m.min=function(n,t,r){var e,u,i=1/0,o=1/0;if(null==t&&null!=n){n=k(n)?n:m.values(n);for(var a=0,c=n.length;c>a;a++)e=n[a],i>e&&(i=e)}else t=x(t,r),m.each(n,function(n,r,e){u=t(n,r,e),(o>u||1/0===u&&1/0===i)&&(i=n,o=u)});return i},m.shuffle=function(n){for(var t,r=k(n)?n:m.values(n),e=r.length,u=Array(e),i=0;e>i;i++)t=m.random(0,i),t!==i&&(u[i]=u[t]),u[t]=r[i];return u},m.sample=function(n,t,r){return null==t||r?(k(n)||(n=m.values(n)),n[m.random(n.length-1)]):m.shuffle(n).slice(0,Math.max(0,t))},m.sortBy=function(n,t,r){return t=x(t,r),m.pluck(m.map(n,function(n,r,e){return{value:n,index:r,criteria:t(n,r,e)}}).sort(function(n,t){var r=n.criteria,e=t.criteria;if(r!==e){if(r>e||r===void 0)return 1;if(e>r||e===void 0)return-1}return n.index-t.index}),"value")};var F=function(n){return function(t,r,e){var u={};return r=x(r,e),m.each(t,function(e,i){var o=r(e,i,t);n(u,e,o)}),u}};m.groupBy=F(function(n,t,r){m.has(n,r)?n[r].push(t):n[r]=[t]}),m.indexBy=F(function(n,t,r){n[r]=t}),m.countBy=F(function(n,t,r){m.has(n,r)?n[r]++:n[r]=1}),m.toArray=function(n){return n?m.isArray(n)?l.call(n):k(n)?m.map(n,m.identity):m.values(n):[]},m.size=function(n){return null==n?0:k(n)?n.length:m.keys(n).length},m.partition=function(n,t,r){t=x(t,r);var e=[],u=[];return m.each(n,function(n,r,i){(t(n,r,i)?e:u).push(n)}),[e,u]},m.first=m.head=m.take=function(n,t,r){return null==n?void 0:null==t||r?n[0]:m.initial(n,n.length-t)},m.initial=function(n,t,r){return l.call(n,0,Math.max(0,n.length-(null==t||r?1:t)))},m.last=function(n,t,r){return null==n?void 0:null==t||r?n[n.length-1]:m.rest(n,Math.max(0,n.length-t))},m.rest=m.tail=m.drop=function(n,t,r){return l.call(n,null==t||r?1:t)},m.compact=function(n){return m.filter(n,m.identity)};var S=function(n,t,r,e){for(var u=[],i=0,o=e||0,a=O(n);a>o;o++){var c=n[o];if(k(c)&&(m.isArray(c)||m.isArguments(c))){t||(c=S(c,t,r));var f=0,l=c.length;for(u.length+=l;l>f;)u[i++]=c[f++]}else r||(u[i++]=c)}return u};m.flatten=function(n,t){return S(n,t,!1)},m.without=function(n){return m.difference(n,l.call(arguments,1))},m.uniq=m.unique=function(n,t,r,e){m.isBoolean(t)||(e=r,r=t,t=!1),null!=r&&(r=x(r,e));for(var u=[],i=[],o=0,a=O(n);a>o;o++){var c=n[o],f=r?r(c,o,n):c;t?(o&&i===f||u.push(c),i=f):r?m.contains(i,f)||(i.push(f),u.push(c)):m.contains(u,c)||u.push(c)}return u},m.union=function(){return m.uniq(S(arguments,!0,!0))},m.intersection=function(n){for(var t=[],r=arguments.length,e=0,u=O(n);u>e;e++){var i=n[e];if(!m.contains(t,i)){for(var o=1;r>o&&m.contains(arguments[o],i);o++);o===r&&t.push(i)}}return t},m.difference=function(n){var t=S(arguments,!0,!0,1);return m.filter(n,function(n){return!m.contains(t,n)})},m.zip=function(){return m.unzip(arguments)},m.unzip=function(n){for(var t=n&&m.max(n,O).length||0,r=Array(t),e=0;t>e;e++)r[e]=m.pluck(n,e);return r},m.object=function(n,t){for(var r={},e=0,u=O(n);u>e;e++)t?r[n[e]]=t[e]:r[n[e][0]]=n[e][1];return r},m.findIndex=t(1),m.findLastIndex=t(-1),m.sortedIndex=function(n,t,r,e){r=x(r,e,1);for(var u=r(t),i=0,o=O(n);o>i;){var a=Math.floor((i+o)/2);r(n[a])<u?i=a+1:o=a}return i},m.indexOf=r(1,m.findIndex,m.sortedIndex),m.lastIndexOf=r(-1,m.findLastIndex),m.range=function(n,t,r){null==t&&(t=n||0,n=0),r=r||1;for(var e=Math.max(Math.ceil((t-n)/r),0),u=Array(e),i=0;e>i;i++,n+=r)u[i]=n;return u};var E=function(n,t,r,e,u){if(!(e instanceof t))return n.apply(r,u);var i=j(n.prototype),o=n.apply(i,u);return m.isObject(o)?o:i};m.bind=function(n,t){if(g&&n.bind===g)return g.apply(n,l.call(arguments,1));if(!m.isFunction(n))throw new TypeError("Bind must be called on a function");var r=l.call(arguments,2),e=function(){return E(n,e,t,this,r.concat(l.call(arguments)))};return e},m.partial=function(n){var t=l.call(arguments,1),r=function(){for(var e=0,u=t.length,i=Array(u),o=0;u>o;o++)i[o]=t[o]===m?arguments[e++]:t[o];for(;e<arguments.length;)i.push(arguments[e++]);return E(n,r,this,this,i)};return r},m.bindAll=function(n){var t,r,e=arguments.length;if(1>=e)throw new Error("bindAll must be passed function names");for(t=1;e>t;t++)r=arguments[t],n[r]=m.bind(n[r],n);return n},m.memoize=function(n,t){var r=function(e){var u=r.cache,i=""+(t?t.apply(this,arguments):e);return m.has(u,i)||(u[i]=n.apply(this,arguments)),u[i]};return r.cache={},r},m.delay=function(n,t){var r=l.call(arguments,2);return setTimeout(function(){return n.apply(null,r)},t)},m.defer=m.partial(m.delay,m,1),m.throttle=function(n,t,r){var e,u,i,o=null,a=0;r||(r={});var c=function(){a=r.leading===!1?0:m.now(),o=null,i=n.apply(e,u),o||(e=u=null)};return function(){var f=m.now();a||r.leading!==!1||(a=f);var l=t-(f-a);return e=this,u=arguments,0>=l||l>t?(o&&(clearTimeout(o),o=null),a=f,i=n.apply(e,u),o||(e=u=null)):o||r.trailing===!1||(o=setTimeout(c,l)),i}},m.debounce=function(n,t,r){var e,u,i,o,a,c=function(){var f=m.now()-o;t>f&&f>=0?e=setTimeout(c,t-f):(e=null,r||(a=n.apply(i,u),e||(i=u=null)))};return function(){i=this,u=arguments,o=m.now();var f=r&&!e;return e||(e=setTimeout(c,t)),f&&(a=n.apply(i,u),i=u=null),a}},m.wrap=function(n,t){return m.partial(t,n)},m.negate=function(n){return function(){return!n.apply(this,arguments)}},m.compose=function(){var n=arguments,t=n.length-1;return function(){for(var r=t,e=n[t].apply(this,arguments);r--;)e=n[r].call(this,e);return e}},m.after=function(n,t){return function(){return--n<1?t.apply(this,arguments):void 0}},m.before=function(n,t){var r;return function(){return--n>0&&(r=t.apply(this,arguments)),1>=n&&(t=null),r}},m.once=m.partial(m.before,2);var M=!{toString:null}.propertyIsEnumerable("toString"),I=["valueOf","isPrototypeOf","toString","propertyIsEnumerable","hasOwnProperty","toLocaleString"];m.keys=function(n){if(!m.isObject(n))return[];if(v)return v(n);var t=[];for(var r in n)m.has(n,r)&&t.push(r);return M&&e(n,t),t},m.allKeys=function(n){if(!m.isObject(n))return[];var t=[];for(var r in n)t.push(r);return M&&e(n,t),t},m.values=function(n){for(var t=m.keys(n),r=t.length,e=Array(r),u=0;r>u;u++)e[u]=n[t[u]];return e},m.mapObject=function(n,t,r){t=x(t,r);for(var e,u=m.keys(n),i=u.length,o={},a=0;i>a;a++)e=u[a],o[e]=t(n[e],e,n);return o},m.pairs=function(n){for(var t=m.keys(n),r=t.length,e=Array(r),u=0;r>u;u++)e[u]=[t[u],n[t[u]]];return e},m.invert=function(n){for(var t={},r=m.keys(n),e=0,u=r.length;u>e;e++)t[n[r[e]]]=r[e];return t},m.functions=m.methods=function(n){var t=[];for(var r in n)m.isFunction(n[r])&&t.push(r);return t.sort()},m.extend=_(m.allKeys),m.extendOwn=m.assign=_(m.keys),m.findKey=function(n,t,r){t=x(t,r);for(var e,u=m.keys(n),i=0,o=u.length;o>i;i++)if(e=u[i],t(n[e],e,n))return e},m.pick=function(n,t,r){var e,u,i={},o=n;if(null==o)return i;m.isFunction(t)?(u=m.allKeys(o),e=b(t,r)):(u=S(arguments,!1,!1,1),e=function(n,t,r){return t in r},o=Object(o));for(var a=0,c=u.length;c>a;a++){var f=u[a],l=o[f];e(l,f,o)&&(i[f]=l)}return i},m.omit=function(n,t,r){if(m.isFunction(t))t=m.negate(t);else{var e=m.map(S(arguments,!1,!1,1),String);t=function(n,t){return!m.contains(e,t)}}return m.pick(n,t,r)},m.defaults=_(m.allKeys,!0),m.create=function(n,t){var r=j(n);return t&&m.extendOwn(r,t),r},m.clone=function(n){return m.isObject(n)?m.isArray(n)?n.slice():m.extend({},n):n},m.tap=function(n,t){return t(n),n},m.isMatch=function(n,t){var r=m.keys(t),e=r.length;if(null==n)return!e;for(var u=Object(n),i=0;e>i;i++){var o=r[i];if(t[o]!==u[o]||!(o in u))return!1}return!0};var N=function(n,t,r,e){if(n===t)return 0!==n||1/n===1/t;if(null==n||null==t)return n===t;n instanceof m&&(n=n._wrapped),t instanceof m&&(t=t._wrapped);var u=s.call(n);if(u!==s.call(t))return!1;switch(u){case"[object RegExp]":case"[object String]":return""+n==""+t;case"[object Number]":return+n!==+n?+t!==+t:0===+n?1/+n===1/t:+n===+t;case"[object Date]":case"[object Boolean]":return+n===+t}var i="[object Array]"===u;if(!i){if("object"!=typeof n||"object"!=typeof t)return!1;var o=n.constructor,a=t.constructor;if(o!==a&&!(m.isFunction(o)&&o instanceof o&&m.isFunction(a)&&a instanceof a)&&"constructor"in n&&"constructor"in t)return!1}r=r||[],e=e||[];for(var c=r.length;c--;)if(r[c]===n)return e[c]===t;if(r.push(n),e.push(t),i){if(c=n.length,c!==t.length)return!1;for(;c--;)if(!N(n[c],t[c],r,e))return!1}else{var f,l=m.keys(n);if(c=l.length,m.keys(t).length!==c)return!1;for(;c--;)if(f=l[c],!m.has(t,f)||!N(n[f],t[f],r,e))return!1}return r.pop(),e.pop(),!0};m.isEqual=function(n,t){return N(n,t)},m.isEmpty=function(n){return null==n?!0:k(n)&&(m.isArray(n)||m.isString(n)||m.isArguments(n))?0===n.length:0===m.keys(n).length},m.isElement=function(n){return!(!n||1!==n.nodeType)},m.isArray=h||function(n){return"[object Array]"===s.call(n)},m.isObject=function(n){var t=typeof n;return"function"===t||"object"===t&&!!n},m.each(["Arguments","Function","String","Number","Date","RegExp","Error"],function(n){m["is"+n]=function(t){return s.call(t)==="[object "+n+"]"}}),m.isArguments(arguments)||(m.isArguments=function(n){return m.has(n,"callee")}),"function"!=typeof/./&&"object"!=typeof Int8Array&&(m.isFunction=function(n){return"function"==typeof n||!1}),m.isFinite=function(n){return isFinite(n)&&!isNaN(parseFloat(n))},m.isNaN=function(n){return m.isNumber(n)&&n!==+n},m.isBoolean=function(n){return n===!0||n===!1||"[object Boolean]"===s.call(n)},m.isNull=function(n){return null===n},m.isUndefined=function(n){return n===void 0},m.has=function(n,t){return null!=n&&p.call(n,t)},m.noConflict=function(){return u._=i,this},m.identity=function(n){return n},m.constant=function(n){return function(){return n}},m.noop=function(){},m.property=w,m.propertyOf=function(n){return null==n?function(){}:function(t){return n[t]}},m.matcher=m.matches=function(n){return n=m.extendOwn({},n),function(t){return m.isMatch(t,n)}},m.times=function(n,t,r){var e=Array(Math.max(0,n));t=b(t,r,1);for(var u=0;n>u;u++)e[u]=t(u);return e},m.random=function(n,t){return null==t&&(t=n,n=0),n+Math.floor(Math.random()*(t-n+1))},m.now=Date.now||function(){return(new Date).getTime()};var B={"&":"&amp;","<":"&lt;",">":"&gt;",'"':"&quot;","'":"&#x27;","`":"&#x60;"},T=m.invert(B),R=function(n){var t=function(t){return n[t]},r="(?:"+m.keys(n).join("|")+")",e=RegExp(r),u=RegExp(r,"g");return function(n){return n=null==n?"":""+n,e.test(n)?n.replace(u,t):n}};m.escape=R(B),m.unescape=R(T),m.result=function(n,t,r){var e=null==n?void 0:n[t];return e===void 0&&(e=r),m.isFunction(e)?e.call(n):e};var q=0;m.uniqueId=function(n){var t=++q+"";return n?n+t:t},m.templateSettings={evaluate:/<%([\s\S]+?)%>/g,interpolate:/<%=([\s\S]+?)%>/g,escape:/<%-([\s\S]+?)%>/g};var K=/(.)^/,z={"'":"'","\\":"\\","\r":"r","\n":"n","\u2028":"u2028","\u2029":"u2029"},D=/\\|'|\r|\n|\u2028|\u2029/g,L=function(n){return"\\"+z[n]};m.template=function(n,t,r){!t&&r&&(t=r),t=m.defaults({},t,m.templateSettings);var e=RegExp([(t.escape||K).source,(t.interpolate||K).source,(t.evaluate||K).source].join("|")+"|$","g"),u=0,i="__p+='";n.replace(e,function(t,r,e,o,a){return i+=n.slice(u,a).replace(D,L),u=a+t.length,r?i+="'+\n((__t=("+r+"))==null?'':_.escape(__t))+\n'":e?i+="'+\n((__t=("+e+"))==null?'':__t)+\n'":o&&(i+="';\n"+o+"\n__p+='"),t}),i+="';\n",t.variable||(i="with(obj||{}){\n"+i+"}\n"),i="var __t,__p='',__j=Array.prototype.join,"+"print=function(){__p+=__j.call(arguments,'');};\n"+i+"return __p;\n";try{var o=new Function(t.variable||"obj","_",i)}catch(a){throw a.source=i,a}var c=function(n){return o.call(this,n,m)},f=t.variable||"obj";return c.source="function("+f+"){\n"+i+"}",c},m.chain=function(n){var t=m(n);return t._chain=!0,t};var P=function(n,t){return n._chain?m(t).chain():t};m.mixin=function(n){m.each(m.functions(n),function(t){var r=m[t]=n[t];m.prototype[t]=function(){var n=[this._wrapped];return f.apply(n,arguments),P(this,r.apply(m,n))}})},m.mixin(m),m.each(["pop","push","reverse","shift","sort","splice","unshift"],function(n){var t=o[n];m.prototype[n]=function(){var r=this._wrapped;return t.apply(r,arguments),"shift"!==n&&"splice"!==n||0!==r.length||delete r[0],P(this,r)}}),m.each(["concat","join","slice"],function(n){var t=o[n];m.prototype[n]=function(){return P(this,t.apply(this._wrapped,arguments))}}),m.prototype.value=function(){return this._wrapped},m.prototype.valueOf=m.prototype.toJSON=m.prototype.value,m.prototype.toString=function(){return""+this._wrapped},"function"==typeof define&&define.amd&&define("underscore",[],function(){return m})}).call(this);
//# sourceMappingURL=underscore-min.map
```
